# Evolutionary analysis of genome-specific duplications in flatworm genomes

**DOI:** 10.1101/2024.02.05.578899

**Authors:** Mauricio Langleib, Javier Calvelo, Alicia Costábile, Estela Castillo, José F. Tort, Federico G. Hoffmann, Anna V. Protasio, Uriel Koziol, Andrés Iriarte

## Abstract

Platyhelminthes, also known as flatworms, is a phylum of bilaterian invertebrates infamous for their parasitic representatives. The classes Cestoda, Monogenea, and Trematoda comprise parasitic helminths inhabiting multiple hosts, including fishes, humans, and livestock, and are responsible for considerable economic damage and burden on human health. As in other animals, the genomes of flatworms have a wide variety of paralogs, genes related via duplication, whose origins could be mapped throughout the evolution of the phylum. Through *in-silico* analysis, we studied inparalogs, *i.e*., genome-specific duplications, focusing on their biological functions, expression changes, and evolutionary rate. These genes are assumed to be key players in the adaptation process of species to each particular niche. Our results showed that genes associated with specific functions, such as response to stress, ion binding, oxidoreductase activity, and peptidase, are overrepresented among inparalogs. This trend is conserved among species from different classes, including free-living species. Available expression data from *Schistosoma mansoni*, a parasite from the trematode class, demonstrated high conservation of the expression patterns between inparalogs, but with notable exceptions, which also display evidence of rapid evolution. We discuss how natural selection may operate to maintain these genes and the particular duplication models that fit better to the observations. Our results support the critical role of gene duplication in the evolution of flatworms.

## 1. Introduction

The phylum Platyhelminthes comprises invertebrate bilateral metazoans commonly known as flatworms (Hickman et al., 2008). Phylogenetic studies show that this phylum belongs to the Lophotrochozoa clade (e.g. Aguinaldo et al., 1997; Egger et al., 2015), which also includes other invertebrates such as Annelids (segmented worms) and Mollusks (Dunn et al., 2014).

Platyhelminthes were classically divided into four classes: Turbellaria (composed of a paraphyletic group of largely free-living species), Cestoda, Monogenea, and Trematoda. The latter three form a monophyletic group of parasitic species named Neodermata, which share the presence of a tegumental syncytium covering the outermost layer of these organisms as a defining synapomorphy (Egger et al., 2015; Hickman et al., 2008). The tegument protects these parasites from the immune response of their hosts and constitutes a functional interphase for the acquisition of nutrients and the excretion of products (Dalton et al., 2004). Indeed, recent works on excreted/secreted products from flatworms have drawn attention to their role in the crosstalk between the parasite and their hosts. These excretory/secretory products are rich in highly immunogenic proteins and proteases involved in parasite invasion and survival. Typical examples are EgAgB and Eg95, secreted by the cestode *Echinococcus granulosus* (Sánchez et al., 1993; Silva-álvarez et al., 2015); SmVal, secreted by the trematode *Schistosoma mansoni* (Curwen et al., 2006) and cathepsins L and B, secreted by *F. hepatica* (see for example Cancela et al., (2008). Interestingly, most of these proteins are members of gene families that have experienced lineage-specific expansions (Coghlan et al., 2019).

It is generally accepted that Neodermata represents an extreme case of adaptive radiation in parasites(Littlewood, 2006). While monogeneans have relatively simple life cycles being most of them ectoparasites of fishes, trematodes and cestodes have complex life cycles involving different hosts (including vertebrates and invertebrates), and undergo extensive anatomical and physiological changes among developmental stages (Hickman et al., 2008). Many parasitic flatworms are infective to humans and domestic animals, resulting in a considerable burden on human health and livestock productivity, especially in developing countries (Feasey et al., 2010).

This work analyzed inparalogs in Platyhelminthes, that is, paralogs that arose after the species split (Remm et al., 2001; Sonnhammer and Koonin, 2002). These genome-specific duplicated genes, which most probably appeared exclusively in each recently diverged lineage analyzed, could have played a vital role in the evolution of traits in each flatworm species. In other words, among all paralogs, we expect that inparalogs may best reflect the most recent biases in duplicability and selective constraints that operate idiosyncratically on each genome, reflecting the particular process that differentiates each species from the other. We focused on the evolutionary rate and the enriched functional categories of inparalogs in parasitic and free-living flatworms. In addition, the relative role of positive natural selection in recent duplications, sequence divergence, and gene expression level is discussed under family evolution models consistent with the observed results.

## 2. Materials and Methods

### 2.1. Sequences and Organisms

Genome sequences, coding sequences, protein sequences, and annotation data of flatworms were downloaded from WormBase ParaSite v11.0 (Howe et al., 2017) available at ftp.ebi.ac.uk/pub/databases/wormbase/parasite/releases/WBPS11/. For coding and protein sequences, when multiple isoforms were annotated, only the longest one was considered for further analysis. The final dataset of Platyhelminthes comprises 30 genomes: 14 tapeworms, 12 flukes, two monogeneans, and two free-living species. Analyses were also performed on genomes belonging to seven Mollusca species and one Annelida species (Supplementary Table 1). Sequences from these organisms were obtained from the NCBI RefSeq database.

### 2.2. Identification of homologous gene families

Clusters of homologous sequences were inferred by means of the get_homologues software v18042017 (Contreras-Moreira and Vinuesa, 2013). A minimum amino acid identity of 40% and coverage of 75% were used as thresholds for BLAST searches. Homology relationships were solved using the OrthoMCL algorithm (Li et al., 2003). The general strategy followed in this manuscript is shown as a workflow in Supplementary Figure 1.

### 2.3. Gene tree inferences

Clusters of homologous protein sequences were aligned with MUSCLE v3.8.31 (Edgar, 2004). Model selection and phylogenetic inferences were made using IQ-Tree v1.5.3 (Nguyen et al., 2015), evaluating support for nodes with the ultrafast bootstrap routine (Hoang et al., 2018). This software integrates model testing, maximum likelihood inference, and computation of branch support into a single run.

### 2.4. Identification of inparalogs in each genome

Inparalogs are a subtype of paralogous genes duplicated after a given speciation event (Remm et al., 2001; Sonnhammer and Koonin, 2002). From a practical point of view, two or more genes in the same genome encoding for proteins displaying higher identity to each other than to any other sequence from other species are considered inparalogs (Koonin, 2005). Following this idea, we employed a phylogenetic approach to identifying inparalogs (Nehrt et al., 2011). Thus, inparalogs were defined as those sequences that cluster into a species-specific monophyletic group within each homologous cluster. A homemade script developed in Python was used to parse phylograms to identify inparalogs clusters, making use of the ete3 v3.1.2 (Huerta-Cepas et al., 2016a) and networkx v2.8.6 libraries (Hagberg et al., 2008).

### 2.5. Prediction of secreted and transmembrane proteins

Secreted proteins (from henceforth the “secretome”) were predicted following (Garg and Ranganathan (2011). Briefly, protein sequences secreted by the canonical pathway were predicted employing SignalP v4.1 (Petersen et al., 2011), setting a D-score cut-off of 0.34 to ensure high specificity. Proteins secreted by the alternative pathways were predicted by SecretomeP v1.0 (Bendtsen et al., 2004). As per published methods in Cuesta-Astroz et al. (2017), a minimum NN-Score of 0.9 was set as a threshold for significance. Transmembrane proteins were identified using TMHMM 2.0c (Krogh et al., 2001)with default parameters and excluded from the secretome. Finally, proteins predicted to reside in the mitochondria by TargetP v1.1 (Emanuelsson et al., 2007) were also excluded from the secretome, employing an NN-score cut-off of 0.95 (Garg and Ranganathan, 2011).

### 2.6. Functional annotation and enrichment analysis

Gene Ontology (GO) term annotation for each protein was done using the eggNOG-mapper tool v0.12.7 (Huerta-Cepas et al., 2016b) with default parameters. The DIAMOND algorithm (Buchfink et al., 2014) was used to infer homology between queries and sequences in the eggNOG database (Huerta-Cepas et al., 2016b). Terms were derived from the hit in the database with the lower e-value (*i.e.,* ‘one2one’ orthology inference modality). Fisher test with a multiple-testing FDR correction was applied for enrichment analysis as implemented in the topGO R library v1.0 (Alexa and Rahnenfuhrer, 2019). Enriched ontologies with an adjusted p-value (FDR) of 0.05 or lower were considered significant. A broader view of molecular function GO enrichment was obtained by mapping enriched GO terms to GO Slims. The map_to_slim.py script from the GOATOOLS package (Klopfenstein et al., 2018) was employed for this purpose, mapping each term to the generic GO Slim set (available at www.geneontology.org).

In order to test if secreted or transmembrane protein-coding genes are overrepresented among inparalog clusters, we applied a Fisher test for every organism with an FDR multiple test correction in R. Briefly, all homologous clusters with at least two sequences were inspected. Then, a 2×2 contingency table was filled classifying homologous genes in four categories: i) inparalog, secreted/transmembrane protein-coding genes, ii) non-inparalog, secreted/transmembrane protein-coding genes, iii) inparalog, non-secreted/non-transmembrane proteins coding genes and iv) non-inparalog, non-secreted/non-transmembrane protein-coding genes. The cumulative result of this procedure (that is, classifying every homolog into these categories for all phylogenetic trees) for each species was used to perform the enrichment test.

Data wrangling was performed using R’s tidyverse package (Wickham et al., 2019), and the results were visualized with the ggplot2 library of this package (Wickham, 2016).

### 2.7. Test for positive selection

Protein sequences for each cluster of inparalogs were aligned with MUSCLE v3.8.31(Edgar, 2004), and low-quality zones were removed with GBlocks v0.91b (Castresana, 2000). The resulting protein alignment was transformed into codon-aligned multiple sequences using PAL2NAL v14 (Suyama et al., 2006). The ratio of nonsynonymous substitutions per non-synonymous sites to the synonymous substitutions per synonymous sites (dN/dS rate or ω) was estimated with the CodeML program from the PAML package v4.9g (Yang, 2007) employing the ete-evol tool from the ETE3 toolkit(Huerta-Cepas et al., 2016a). Site-specific models M1a (nearly neutral model: two ω categories, ω < 1 and ω = 1) and M2a (positive selection allowed: three ω categories, ω < 1, ω = 1, and ω > 1) (Yang et al., 2005) were then applied to each cluster of inparalogs. The likelihood ratio test (LRT), with p < 0.05, was used to identify the more probable model in each cluster (Yang and Bielawski, 2000). Finally, the Bayes Empirical Bayes (BEB) test was used to identify sites evolving under positive selection (ω >1), using a posterior probability >= 0.95 (Yang et al., 2005).

### 2.8. Gene expression analyses of S. mansoni

Raw expression data available at the Sequence Read Archive (Leinonen et al., 2011) was downloaded and formatted using fastq-dump v2.9.1 from the SRA Toolkit (available at ncbi.github.io/sra-tools). RNA sequencing data from the Wellcome Sanger Institute, Protasio et al. (2012), and Wangwiwatsin et al. (2020), are available at Wormbase ParaSite (Supplementary Table 1). Expression divergence among inparalogs was assessed using Pearson’s and Spearman’s correlation coefficients. Trimming FastQ files and sequencing adaptor removal was performed with Trimmomatic v0.38 (Bolger et al., 2014) (parameters: LEADING:3 TRAILING:3 MINLEN:36). Quality control was performed with FastQC v0.11.5 available at www.bioinformatics.babraham.ac.uk, and quality reports were summarized with MultiQC v1.9 (Ewels et al., 2016). Read mapping against the S. mansoni genome was performed with STAR v2.5.2b (Dobin et al., 2013).

Multi-mapped reads were discarded (i.e., option –outFilterMultimapNmax 1) to ignore ambiguously mapped reads to inparalogs. Feature count was performed with htseq-count v2.0.1 (Putri et al., 2022) with “-s no” option, and normalized pseudocounts were calculated with edgeR (Robinson et al., 2009) using the Trimmed Mean of M-values (TMM) method (Robinson and Oshlack, 2010). Then, the median values were calculated among replicates. Correlation coefficients were calculated employing R’s library corrr (Kuhn et al., 2020). These correlation coefficients were plotted against the total sum of cophenetic dS (separately) for every pair of genes constituting inparalogous clusters in order to evaluate expression divergence in relation to sequence divergence. Sequence divergence measures were calculated by processing CodeML calculations and inferred gene trees with the R libraries treeio (Wang et al., 2020) and ape (Paradis and Schliep, 2019). R ggplot2 library (Wickham, 2016) was used to visualize expression data.

### 2.9. Data availability

Sequence alignments, phylogenetic trees, dN/dS results, transmembrane and secreted proteins predictions, RNA-seq mapping statistics, and functional annotation (GO terms) are available at Zenodo.org (doi.org/10.5281/zenodo.10257625).

## 3. Results and Discussion

### 3.1. Almost twenty-six thousand inparalogous groups were identified across flatworm genomes

In the analysis of 30 flatworm genomes we identified 263,858 protein-coding genes as singletons, single sequences belonging to a unique genome (Figure 1A). When the identity and coverage thresholds are relaxed, this number remains around 250,000, suggesting that this result is relatively independent of the specific search parameters (data not shown). In addition, 67,269 clusters of homologous genes of two or more sequences were assembled. These clusters comprised 304,784 genes, representing 53.5% of the analyzed gene set (Figure 1B).

**Figure 1.**
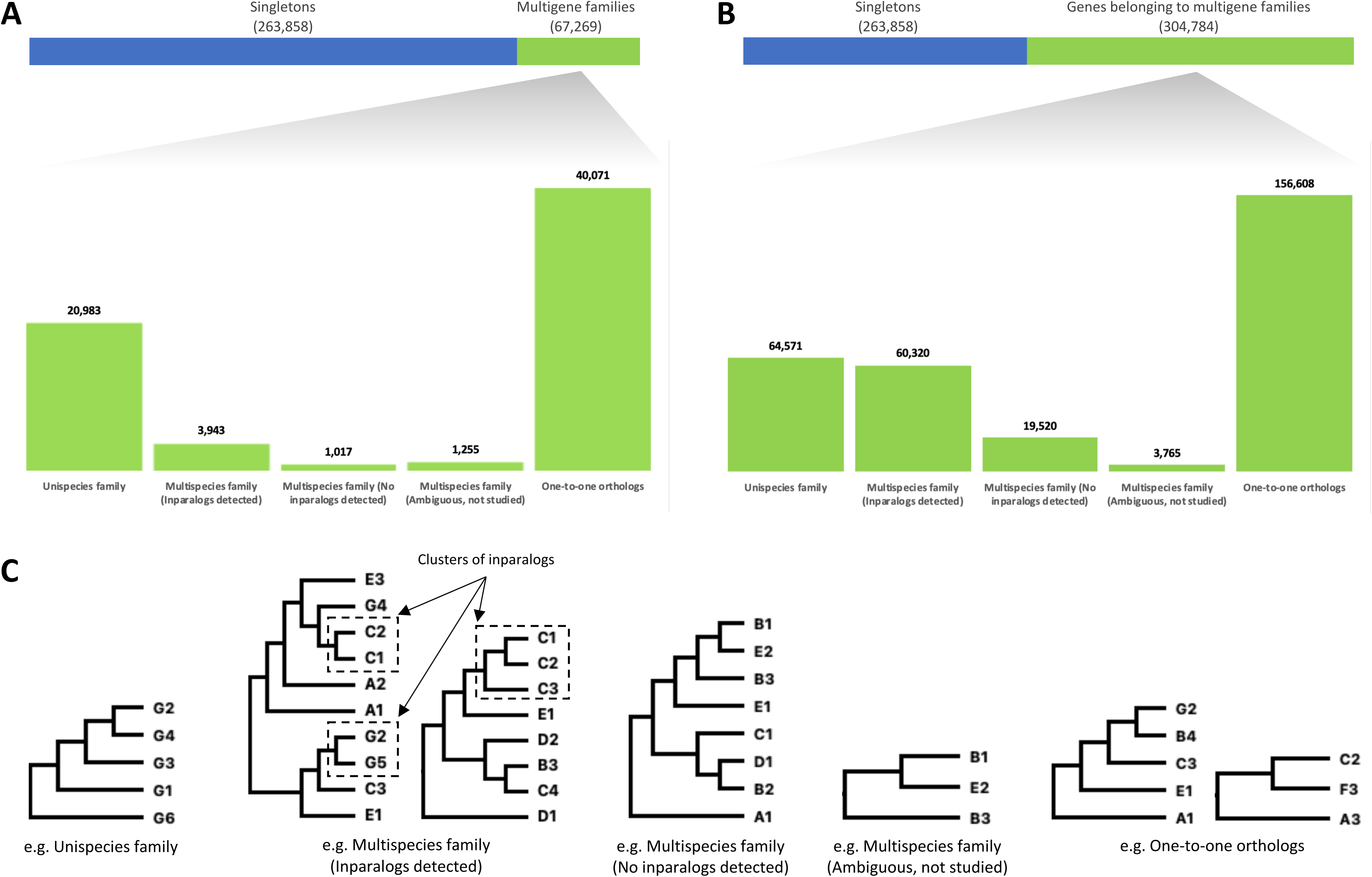
Distribution of homologous clusters (A) and genes (B) according to categories defined by species distribution within each cluster. “Singletons”: clusters comprising only one gene. “Unispecies families”: homologous clusters with more than one gene, all belonging to a unique species. Multispecies family with inparalogs: clusters comprising homologous genes from more than one species with inparalogs detected. Multispecies families without inparalogs: homologous clusters comprising genes from more than one species without inparalogs detected. Ambiguous multispecies family: clusters of three homologous genes with three sequences belonging to two different species (no inparalogs could be defined in the unrooted tree). “One-to-one orthologs”: clusters with orthologous genes. C) Examples of phylogenetic trees of different categories of multigene families. Letters indicate species, and numbers indicate gene copies within each species.

Genes belonging to clusters composed exclusively by sequences coded in a single genome were considered inparalogs without being subject to any phylogenetic analysis, as they appear to have diverged significantly from other putative homologs coded in other genomes (N of clusters=20,983, N of genes=64,571; named as “Unispecies family” in Figure 1). The remaining clusters of homologous sequences were subjected to gene tree inference and topological analysis (see Materials and Methods).

The pipeline defined 25,995 groups of inparalogs, distributed in 24,926 clusters of homologs. The number of inparalogs in each species ranged from 163 to 40,944 in E. granulosus and M. lignano, respectively (Table 1). This considerable variation is expected given that our strategy for identifying inparalogs mainly depends on the evolutionary distance among available genomes. In agreement with the recent genome duplication event proposed in M. lignano (Zadesenets et al., 2016), we find a very high, yet not surprising, number of inparalogs in this free-living species. Furthermore, a second factor to consider is the quality of genome assembly and annotation in these non-model species, which clearly affects in some cases, the inparalogs and singletons counts (e.g. S. erinaceieuropaei) (Supplementary Table 2).

**Table 1.** The number of inparalogs and singletons identified in each species.

| Class | Species | Total annotated protein-coding genes | Homologs* | Singletons | Inparalogs | Inparalogous clusters |
| --- | --- | --- | --- | --- | --- | --- |
| Cestoda | *Hymenolepis diminuta* | 11,271 | 7,884 | 3,387 | 239 | 84 |
|  | *Hymenolepis microstoma* | 12,373 | 8,704 | 3,669 | 1,436 | 570 |
|  | *Hymenolepis nana* | 13,777 | 8,592 | 5,185 | 768 | 257 |
|  | *Mesocostoides corti* | 10,614 | 6,651 | 3,963 | 396 | 139 |
|  | *Echinococcus granulosus* | 10,273 | 9,316 | 957 | 163 | 80 |
|  | *Echinococcus multilocularis* | 10,669 | 9,558 | 1,111 | 603 | 225 |
|  | *Echinococcus canadensis* | 11,432 | 7,559 | 3,873 | 632 | 174 |
|  | *Taenia solium* | 12,481 | 8,435 | 4,046 | 189 | 86 |
|  | *Taenia asiatica* | 13,322 | 11,222 | 2,100 | 1,352 | 563 |
|  | *Taenia saginata* | 13,161 | 10,873 | 2,288 | 987 | 421 |
|  | *Hydatigera taeniformis* | 11,649 | 7,313 | 4,336 | 177 | 82 |
|  | *Diphyllbothrium latum* | 19,966 | 6,191 | 13,775 | 639 | 273 |
|  | *Schistocephalus solidus* | 20,228 | 8,608 | 11,620 | 2,776 | 551 |
|  | *Spirometra erinaceieuropaei* | 39,557 | 14,467 | 25,090 | 8,974 | 3,950 |
| Trematoda | *Clonorchis sinensis* | 13,634 | 6,494 | 7,140 | 544 | 142 |
|  | *Opisthorchis viverrini* | 16,356 | 6,397 | 9,959 | 394 | 151 |
|  | *Echinostoma caproni* | 18,607 | 6,596 | 12,011 | 1,396 | 350 |
|  | *Fasciola hepatica* | 9,708 | 5,329 | 4,379 | 218 | 96 |
|  | *Trichobilharzia regenti* | 22,185 | 7,730 | 14,455 | 595 | 192 |
|  | *Schistosoma curassoni* | 23,546 | 11,495 | 12,051 | 316 | 130 |
|  | *Schistosoma haematobium* | 11,140 | 8,532 | 2,608 | 375 | 104 |
|  | *Schistosoma japonicum* | 12,738 | 7,375 | 5,363 | 450 | 136 |
|  | *Schistosoma mansoni* | 14,519 | 12,996 | 1,523 | 728 | 289 |
|  | *Schistosoma margrebowiei* | 26,189 | 13,047 | 13,142 | 2,419 | 627 |
|  | *Schistosoma mattheei* | 22,997 | 11,572 | 11,425 | 180 | 81 |
|  | *Schistosoma rodhaini* | 24,089 | 11,109 | 12,980 | 1,228 | 272 |
| Monogenea | *Gyrodactylus salaris* | 15,436 | 2,812 | 12,624 | 334 | 139 |
|  | *Protopolystoma xenopodis* | 37,906 | 4,362 | 33,544 | 1,470 | 378 |
| Turbellaria | *Macrostomum lignano* | 49,013 | 41,410 | 7,603 | 40,944 | 12,944 |
|  | *Schmidtea mediterranea* | 29,850 | 12,199 | 17,651 | 9,533 | 2,509 |

\* Number of genes that present homologs in other species or within the same species.

### 3.2. Functionally related groups of genes are systematically enriched among inparalogs in different species

Since inparalogs are genome-specific and, therefore, the product of duplications that occur independently and recently in the lineages, we first hypothesized that these genes were involved in different biological processes or functions in each parasite species. On average, 41% of the homologs detected were devoid of any annotation, so no inferences could be made about their function. The analysis of the remaining annotated genes by evaluating their GO and Interpro assignments showed that, in general, the same higher-rank gene ontology terms were systematically enriched among inparalogs in different species (Supplementary Table 3, Supplementary Figure 2). Many of these enriched functional terms have been previously identified as relevant factors in the interaction with the host and parasite survival, dealing with processes such as infection, immunomodulation, and energy interchange (see Coghlan et al., 2019; Cuesta-Astroz et al., 2017; Silva-álvarez et al., 2015; among many others). Almost all lineages analyzed show inparalog enrichment in functional terms such as “Peptidase activity” (GO:0008233) or “Lipid binding” (GO:0008289), which have been experimentally linked to the parasite infection process (Franchini et al., 2015; McKerrow et al., 2006). Other high-order GO terms also enriched among members of the four classes of the phylum are “Oxidoreductase activity” (GO:0016491), “Response to stress” (GO:0006950), “Ion binding” (GO:0043167), “Transmembrane transporter activity” (GO:0022857), “Transferase activity, transferring glycosyl groups” (GO:0016757) and “Kinase activity” (GO:0016301). Coherently, many of these families went through complex expansion processes throughout diverse parasitic helminth lineages, as reported recently by Coghlan et al. (2019) and others. Thus, we believe that a more detailed analysis within these terms and functional categories, for instance, at the level of each homologous group, could show differences among free-living and parasitic species.

The term “Peptidase activity” (GO:0008233) is linked to many identified inparalogs across flatworms. Diverse proteases such as cathepsins, leucine aminopeptidases, legumains, elastases, and metallopeptidases and their inhibitors have been identified as relevant in the parasite-host interphase in diverse lineages. To analyze the possible differences among flatworm classes, we classified 2748 peptidases out of 3434 candidates coded by inparalogs using the MEROPS database (Rawlings et al., 2018) (best blastp hit against the pepunit database, ID > 40%, e-value < 10^-5^). Some characteristic trends among some families were observed, with ubiquitin peptidases (C19 family) or Zinc Aminopeptidases (M01 family), showing higher inparalog numbers in cestodes, while the well-known cathepsins of the papain-like proteases (family C01) are enriched in inparalogs within Trematodes. Interestingly, some of these protease families were also amplified in free-living species (Supplementary Figure 3).

Inparalogs coding for proteins involved in the biosynthesis of glycan moieties were also found and can be linked to the enrichment term “Transferase activity, transferring glycosyl groups” (GO:0016757) or to more general terms related to carbohydrate biosynthesis. These include glycosyl transferases of the families GT-10 and GT-23 (fucosyltransferases), GT-14 (I-branching enzyme and core-2 branching enzyme), and beta-1,4-galactosyltransferases, as well as glycosyl hydrolases of the lactate dehydrogenase/glycoside hydrolase family 4 and GT-13 (O-glycosyl hydrolases). Species-specific duplication of many copies of glycosyltransferases has been reported in E. granulosus and E. multilocularis (del Puerto et al., 2016; Tsai et al., 2013). In particular, del Puerto et al. (2016) proposed that observed differences in the biosynthesis of mucin O-glycans between both species can be partially explained by expression differences in a glycosyltransferase gene supporting a relevant role of this gene in the differences among species. In the trematode S. mansoni, changes in glycan composition between the different life stages have been well described, a phenomenon that has been postulated as a mechanism for the evasion of the immune system of different hosts (Hokke et al., 2007).

When the InterPro functional classification of inparalogs was analyzed, we found highly frequent superfamilies overlapped with the previously mentioned GO terms and that were also mentioned by Coghlan et al. (2019) (Supplementary Table 4). That is, G protein-coupled receptor, rhodopsin-like family (IPR000276), Protein kinase-like domain (IPR011009), Ion transport domain (e.g., IPR005821, IPR006201), Galactose-binding-like domain (IPR008979), Glycosyltransferase families (e.g., family 31 (IPR002659), 10 (IPR001503), 14 (IPR003406), 23 (IPR027350)), Glycoside hydrolase family (IPR017853), Ankyrin repeat-containing domain (IPR020683), Alpha-2-macroglobulin family (IPR001599) and transporters, mainly form the Major facilitator superfamily domain (IPR020846) and the ABC transporters superfamily (IPR003439), which were repeatedly found across species.

Interestingly, among highly frequent groups, we identified CAP family members (IPR035940), Kunitz (IPR002223), EF-hand pair (IPR011992), and the Fibronectin type III (IPR036116) domains. CAP superfamily (an acronym for CRISP, Ag5, and PR-1) comprises a functionally diverse group of proteins involved in various biological processes. In flatworm parasites, CAPs are expressed throughout the life cycle and are associated with invasion, establishment, remodeling, among other functions (Chalmers et al., 2008; Rofatto et al., 2012; Yoshino et al., 2014). The Kunitz-type family of peptidase inhibitors, has also been proposed to have roles in the interaction between flatworm parasites and their host (see Fló et al., 2017; Mambelli et al., 2021; Smith et al., 2020; among others). Many lineage-specific expansions have been described in trematodes and cestodes for CAP and Kunitz-type families (e.g. Costábile et al., 2018; González et al., 2009). EF-hand and Fibronectin type III domains are present in well-known parasitic antigens (see, for instance, Chow et al., 2001; Emmanoch et al., 2018; Fitzsimmons et al., 2012; Haag et al., 2009; Zheng et al., 2018), making them highly interesting for more comprehensive future studies. It is important to remark that these analyses are highly dependent on functional annotation and prediction, which as previously stated is relatively poor in flatworms compared to model organisms (Supplementary Table 2).

High duplicability, an increased likelihood of duplication (mutation), retention, and fixation during evolution (He and Zhang, 2005), has been described for the same functional categories in different eukaryotes (Kirkness et al., 2010; Waterhouse et al., 2010). Here we focused exclusively on genome-specific duplications and found common trends between the functional categories enriched among these recent gene duplications and those of gene families expanded throughout the phylogenetic tree of flatworms (Coghlan et al., 2019). This result suggests that the adaptive strategies of different flatworm species (ancestral and extant species) may depend on a similar set of molecular functions or biological processes. A significant but minor effect of phylogenetic inertia on duplication trends of particular gene families throughout the evolution of Platyhelminthes cannot be discarded. Further analysis within each functional category could help identify species-specific expansions driven by positive selection.

### 3.3. Secreted proteins are overrepresented among genome-specific duplications

Several of the genes involved in the interaction with the host that have been repeatedly reported as expanded gene families in helminths are part of the excretory/secretory proteome (Chiumiento and Bruschi, 2009; Wang et al., 2015). In this work, we identified a total of 63,050 secreted proteins in flatworms’ genomes (Supplementary Table 2). The fraction of secreted proteins in each genome ranges from 5.05% to 16.16% and agrees with previously published studies (Cuesta-Astroz et al., 2017). No substantial differences in the amount of secreted inparalogs were found when comparing among classes, between parasitic and free-living species, nor between flatworms and the closely related phyla Annelida and Mollusca (Supplementary Table 2). In parasitic species, a large amount of experimental evidence highlights the roles of multigenic families of secreted and surface molecules in host-parasite and environment-parasite interaction. These involve very diverse functions including invasion, immune evasion, signaling, detoxification, and energy intake (Hewitson et al., 2009; Lightowlers and Rickard, 1988).

Our analyses show that secreted-protein coding genes are significantly overrepresented among inparalogs in 21 of the 30 flatworm species analyzed, including both parasitic and free-living species (Figure 2). Only in Echinococcus canadensis, and the poorly annotated species Schistosoma curassoni, Schistosoma margrebowiei, and Schistosoma matthei, the secreted proteins are less frequent among inparalogs than in non-inparalogs. This suggests that this group of genes has higher duplicability than the average gene set, at least with regard to more recent duplication events. New copies of secreted protein-coding genes may be at least partially associated with the niche preferences or particularities of each species.

**Figure 2.**
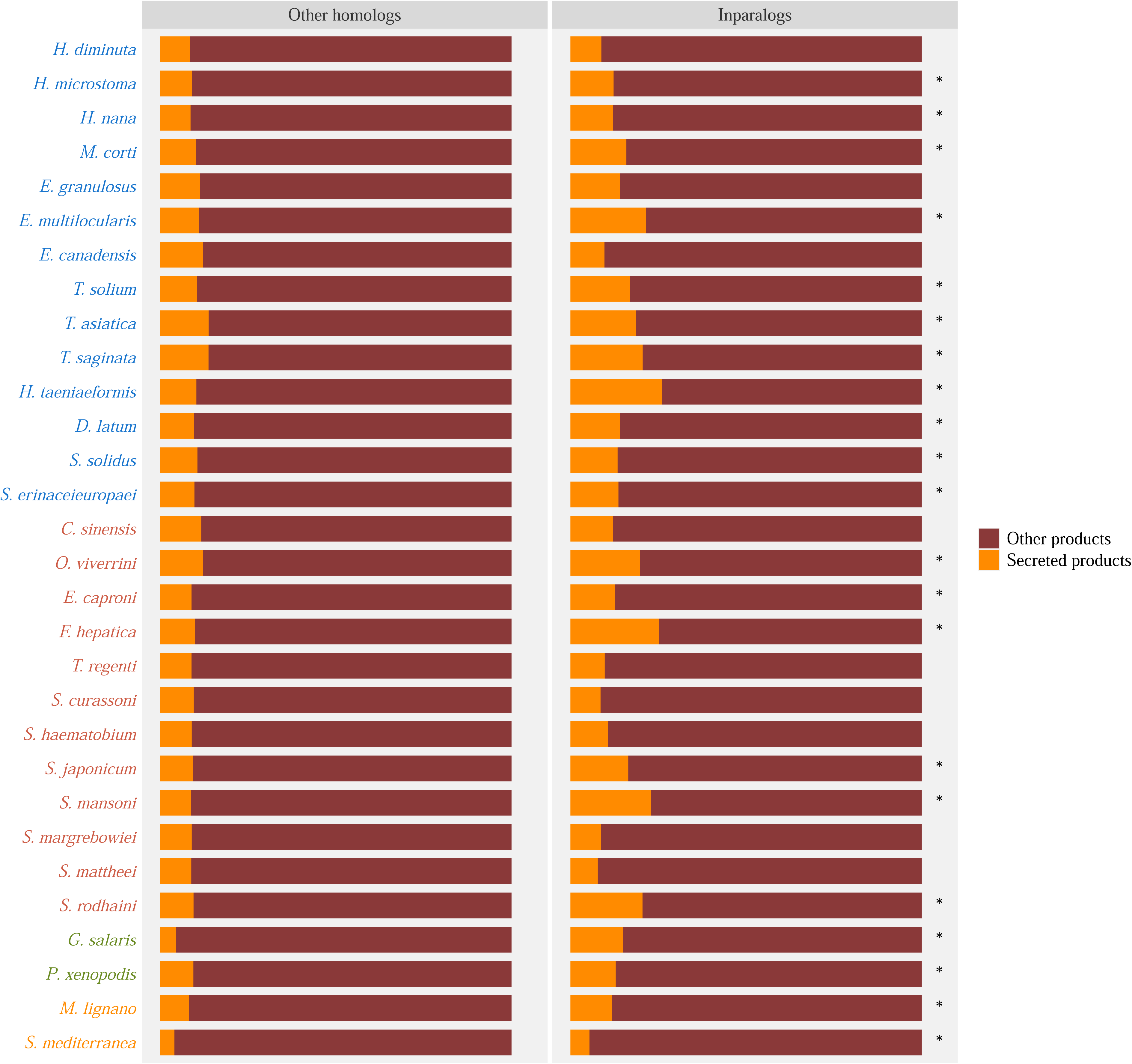
Distribution of secreted protein-coding genes within inparalogs and other homologs in each flatworm genome. Significantly overrepresentation of this “functional category” among inparalogs is indicated with “*” (Fisher exact test, FDR < 1%). Note that homologous genes (all but singletons) are the background for enrichment tests.

High duplicability is believed to be a ubiquitous characteristic of secreted proteins protein-coding genes in all organisms (e.g., Kondrashov et al., 2002; Waterhouse et al., 2010). Despite the inherent heterogeneity within this group, various features may account for this observation in platyhelminths and other eukaryotic phyla. First, an increased copy number of copies could lead to a high dose of the protein products that play vital roles in the interaction with the host or environment, a process known as gene dosage effect (Kondrashov et al., 2002). Second, secreted proteins, which are folded in the endoplasmic reticulum and are restricted to the extracellular space, have a reduced fitness cost of misfolding and miss interactions (Feyertag et al., 2016). Finally, the effect of negative auto-regulation may also be reduced compared to non-secreted proteins.

Protein families involved in nutrient uptake, peptidase activity, and lipid binding are good examples of selective-driven processes in which the increased dosage benefits the organism. Thus, in a fixation phase, new secreted protein-coding genes can be maintained in the population by selection because of increased gene dosage. On the other hand, in many cases, the function of secreted proteins depends on the interaction with specific targets, nutrients, or substrates. Thus, feasibly while keeping the functional core of the protein, these new duplicated genes can acquire slightly different functions by a few amino acid changes. Indeed, due to relaxed purifying selection, duplicated genes would quickly accumulate new mutations eventually leading to a new function, which may then be fixed and maintained by selection (Kondrashov et al., 2002).

An archetypical example in flatworms is the liver fluke cathepsins, where distinct changes in active site residues result in diverse enzyme specificity (Corvo et al., 2013, 2009). Another example can also be found in Glutathione S-transferases and other proteins involved in detoxification. A few specific amino acid changes in the substrate-binding site can increase the repertoire of possible second substrates increasing detoxification capacity (Hayes et al., 2005). Moreover, in the case of parasite species, the duplication of specific secreted protein-coding genes followed by divergence may be favored as it increases genetic variability and thus increases the immune evasion capacity.

The high frequency of inparalogs coding for secreted proteins is conserved among most flatworm parasites, including ecto- and endo-parasitic species, in free-living species (Figure 2), and also in Annelida and Mollusca species (Supplementary Figure 4A). This supports the idea that high duplicability in secreted proteins is a general rule for Platytrochozoa as found in other main eukaryotic lineages (Kondrashov et al., 2002).

We observed that transmembrane-proteins coding genes do not show the same trend as secreted-protein coding genes (Supplementary Figures 4B and C). Thus, further analyses are needed to assess whether these groups possess different evolutionary dynamics or regulatory mechanisms that may underlie the observed differences.

### 3.4. Most inparalogs show a conserved expression pattern among developmental stages in **S. mansoni**

Life cycles of parasitic flatworms are complex, and regulatory differences in gene expression across different developmental stages are evident (Huang et al., 2016; Olson et al., 2018; Protasio et al., 2012; Tsai et al., 2013). To evaluate the differences in the expression of inparalogs we selected the model trematode S. mansoni, the more thoroughly studied species within parasitic flatworms. Because of the denser sampling within the genus Schistosoma, the analyzed species have diverged more recently than in other cross-genera comparisons. Therefore, the inparalogs identified in these species are more recently duplicated genes or, in other words, species-specific duplications.

RNA-seq data of S. mansoni was taken from 46 different sequencing runs from 3 studies (Supplementary Table 5, Supplementary Figures 5, 6, and 7). Gene expression levels for 728 inparalogs from 289 clusters in different life cycle stages were estimated using unique mapped reads. Among these 76% display median Trimmed Mean of M-values (TMM) values above 5 in at least one of the different developmental stages analyzed indicating that these consist of functional expressed genes. Since S. mansoni transcriptomic data is relatively comprehensive of the diverse life stages of the cycle, it is reasonable to propose that at least part of the 84 (11.5%) inparalogous genes with marginal or null expression value (median TMM < 1) in all the time points analyzed might be putative pseudogenes or miss-annotated genes (Supplementary Table 5). However, some genes might have expression restricted to particular tissues or moments during the cycle, not tested or obscured in whole organism transcriptomic approaches like the ones here analyzed. Thus, this number could be considered an estimated upper value for the actual number of pseudogenes among inparalogs in S. mansoni.

We estimated the Pearson and Spearman correlations of expression values of inparalogs among time points. Genes with a median TMM value lower than 1 in all stages were removed, leaving 740 pairwise comparisons (Supplementary Table 6). Results showed that the correlation values within groups of inparalogs tend to be high or very high for the majority of the comparisons (Figure 3), suggesting that in most cases, new gene copies tend to maintain the regulatory constraints of their ancestral gene. This is consistent with the idea that inparalogs analyzed correspond to recent duplication events in S. mansoni. Many published studies analyzed the expression pattern of duplicated genes and agreed on a positive correlation among inparalogs that is at the same time stronger than outparalogs (paralogs originated in a duplication that occurred before the speciation event) and possibly than orthologs. This latter observation had been extensively discussed in the context of the ortholog conjecture (Chen and Zhang, 2012; Gabaldón and Koonin, 2013; Nehrt et al., 2011; Stamboulian et al., 2020) as it has implications at the level of homology-based functional annotation.

**Figure 3.**
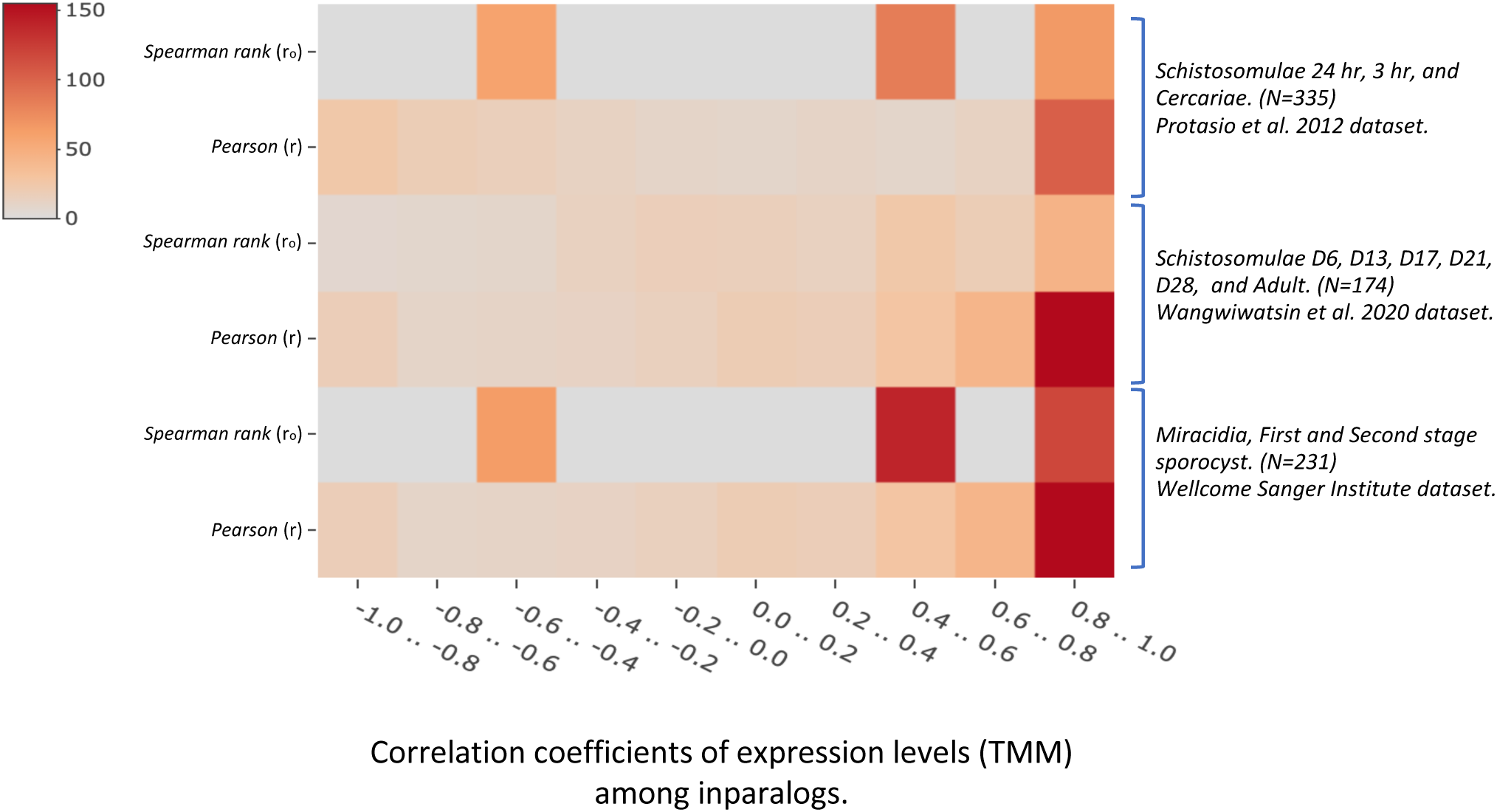
Heatmap showing the distribution of Pearson (r) and Spearman (r_o_) values of correlation of expression levels (i.e., TMM) among inparalogs identified in S. mansoni. Twelve developmental stages from three different studies were included. Pairwise correlations were estimated within each study. Genes with TMM values in all-time points below 1 were not included in the analysis. The total number of pairwise correlations estimated within each study is indicated in brackets.

Gene expression divergence is often used as a proxy indicator of the divergence of gene function (Wang et al., 2012), hence we could conclude that in S. mansoni, most putative functional inparalogs maintain gene function after duplication, as predicted by the “Positive dosage” duplication model (Kondrashov et al., 2002). Under this model, duplicated copies are maintained by selection for increased gene dosage of an original function that increases organismal fitness. This is consistent with the observation that functions that depend on the dosage, like stress and environmental response, and functions related to secretion (required in large doses) are overrepresented among inparalogs in most analyzed flatworm species (see above).

Despite this, subtle changes in the duplicated genes might be relevant to produce diversity within proteins exposed to the interaction with the host, providing adaptative advantage at vital processes in parasitic species such as immune evasion, invasion and detoxification. Examples of these are cathepsins and GSTs as already discussed (Hayes et al., 2005).

Qian et al. (2011) proposed a subfunctionalization-like model that may also explain the observed highly correlated expression patterns of recently duplicated genes. According to this model, following duplication there is a reduction of expression in the joint expression level of both copies to reach the pre-duplication optimal level. This model implies a deleterious effect of protein production excess due to waste of energy and anabolites, the toxicity of misfolded proteins, mis-interaction, or a broken stoichiometry among molecules (Drummond and Wilke, 2009; Pa and Hurst, 2003; Qian and Zhang, 2008; Wagner, 2005; Zhang and Yang, 2015). In this study, the expression values of ancestral genes were not estimated, so a reduction in the joint expression level of duplicated genes could not be discarded. However, as previously mentioned, for many overrepresented functions among inparalogs, e.g., functions typically associated with secreted proteins, the predicted deleterious effects may not apply. New expression data of Platyhelminthes species may be helpful to assess which of the proposed models better fits the observations.

Many genes with highly correlated expression values showed stage-specific expression. For example, Smp_303390 and Smp_303420, a pair of genes coding for HSP70, showed expression mainly in the Schistosomulae 3 hrs stage of S. mansoni (Figure 4A), while other inparalogs of this group were not expressed. Other genes showed similar expression trends among many studied stages; see, for instance, the case of Smp_058690 and Smp_058700, two inparalogs coding for glutathione peroxidases that tend to increase the expression towards the adult stage (D35) (Figure 4B). Highly correlated expression can be seen in more than two inparalogs, as shown for Saposin B type domain proteins (Figure 5C).

**Figure 4.**
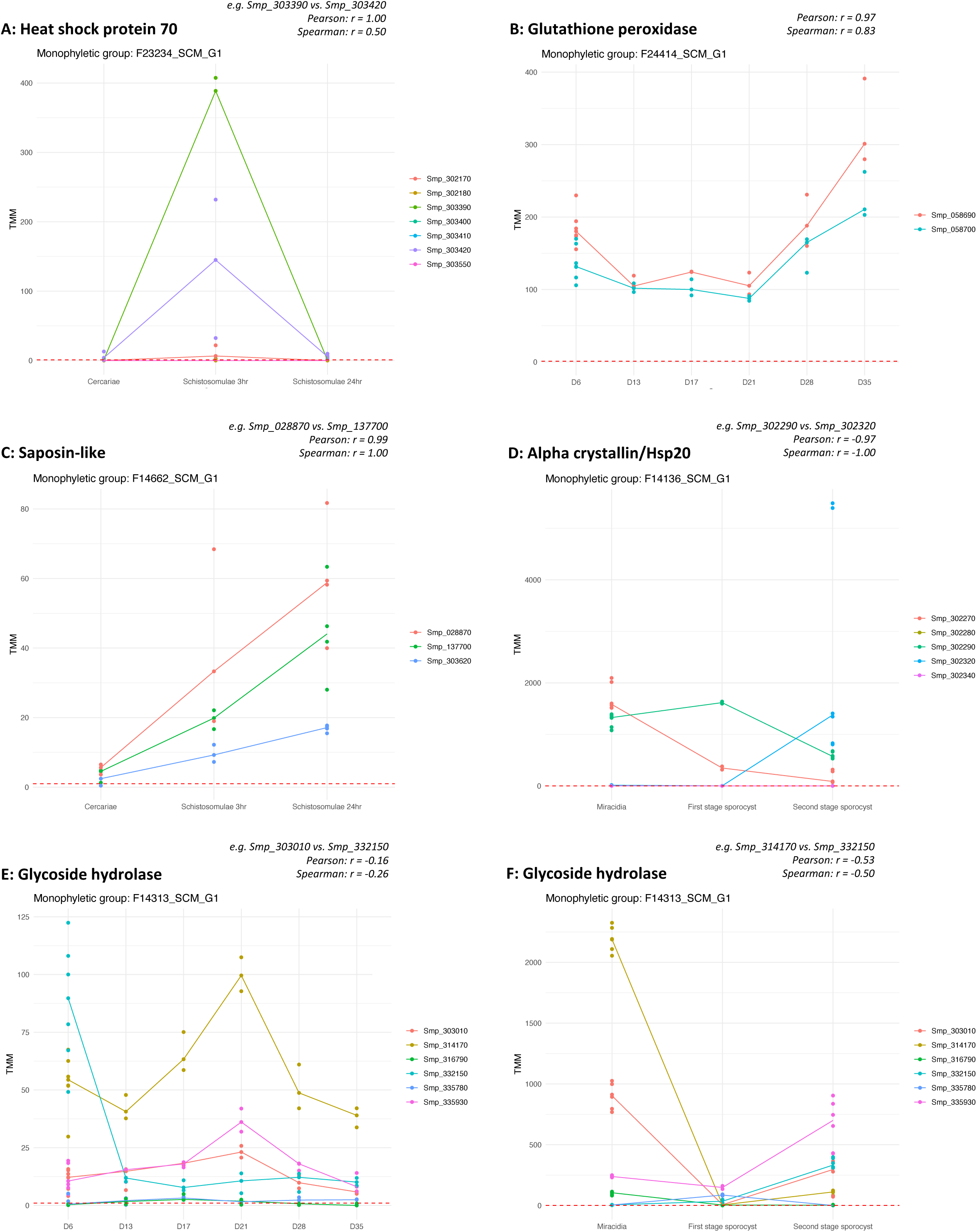
Expression patterns were observed in six selected examples of inparalogous clusters. The expression of each inparalog in each replicate analyzed is indicated with a dot, while the median value of expression among replicates for each inparalog is plotted. A, B, and C are examples of positive correlations among inparalogs. A is an example of expression restricted to a specific stage or time point. B and C are examples of similar expression trends throughout different conditions. E and F are examples of negative correlations. The correlation coefficients Pearson and Spearman, between selected pairs of genes, are indicated for each selected pair of inparalogs.

We studied the association between the estimated correlation values of expression and the physical distance between inparalogs. Based on gene location, all pairs of inparalogs were classified into three categories: “close”, less than 10 kbp, “distant”, more than 10 kbp, and “different chromosomes”. Expectedly, close inparalogs tend to show a higher expression correlation than distant inparalogs and inparalogs in different chromosomes, although the difference is not significant between close and distant inparalogs (U-test, p < 0.05, Supplementary Figure 8). This result suggests that regulatory constraints linked to genome location may contribute to the observed pattern. To analyze if expression correlation declined with increasing time since duplication (Casneuf et al., 2006; Rogozin, 2014), we studied the association between the estimated correlation values and the synonymous cophenetic distance among inparalogs in S. mansoni. As a result, we could not find conclusive evidence to support this association (data not shown).

Although to a minor extent, we identified inparalogs that considerably change expression patterns among analyzed stages. This pattern can be evidenced in the negative correlation values shown in Supplementary Table 6 or the plots of TMM values shown in Supplementary Figures 5, 6, and 7. We believe that these inparalogs are good candidates for further studies since they probably represent S. mansoni species-specific duplications that functionally and idiosyncratically diverged. See, for instance, the expression pattern of inparalogs in S. mansoni that belongs to the family of the putative major egg antigen (IPR002068: Alpha crystallin/Hsp20 domain) (Figure 4D), or the Glycoside hydrolase superfamily (Figure 4E, F). Different expression patterns throughout the development and functional divergence in duplicated genes have been reported previously in cestodes and flukes (Choi et al., 2020; Costábile et al., 2018; Cwiklinski et al., 2015; Protasio et al., 2012).

Transcriptional regulation of new gene copies may have diverged quickly to cope with specific functions at different time points during the life cycle of this species. Thus, these new copies most likely perform slightly different functions, depending on the conserved functional core that characterizes each gene family. According to the “Modified duplication” model, these rapid regulatory changes, which in some cases may have originated at the time of duplication itself, may be favored by natural selection (Lynch and Katju, 2004). However, other selectionist models that predict functional (and genetic) variability at a population level at a pre-duplication phase cannot be completely discarded (e.g., the Adaptive radiation model proposed by Francino (2005) or the IAC model proposed by Andersson et al. (2015).

Taken together, our results support the idea that the evolutionary trajectories of functional inparalogs may be explained by the beneficial effects of an increased gene dosage (an immediate benefit of duplication per se) and functional divergence. Which model is more suitable in each case depends on the specific gene family analyzed.

### 3.5. Evidence of positive selection is limited to a small proportion of codon positions in the inparalogs

To assess the strength and mode of natural selection in the evolution of coding sequences within each inparalog group, we estimated the dN/dS ratio (ω) (Supplementary Table 4). Two site models, M1a (nearly neutral evolution model) and M2a (positive selection model), were used, and a global ω among all inparalogs was estimated for each group using the model with the highest likelihood. Since the dN/dS ratio can be a misleading selection estimator when divergence time from the common ancestral sequence is too short, we only considered those inparalogs groups with a global dS > 0.1 for further analysis.

A significant fraction of inparalogs groups studied have a dS value below 0.1 (Average 54.6%, SD +/- 26.2%), indicating that the sequence of these new gene copies remained highly similar after duplication (Supplementary Figure 9A, Supplementary Table 4). Of course, this divergence is also dependent on the phylogenetic sampling of the study, so the observed differences among species are not surprising. On average, only 11.2% (SD +/- 6.8%) of all groups of inparalogs are evolving with one or more codon sites with ω > 1 (Supplementary Figure 9A), many of which are linked to host interaction. Interestingly, the global ω estimation on inparalogous groups showed that only a few analyzed cases are under strong purifying selection (ω < 0.25) (Supplementary Figure 9B).

Duplicated genes evolve faster than orthologs with the same level of synonymous sequence divergence. The acceleration of evolution is an inherent feature of duplicated genes, which could be explained in terms of positive selection or relaxation of negative selection (Kondrashov et al., 2002). We observed that in inparalogs where a model that allows sites to evolve under positive selection is significantly more probable than the nearly neutral model only a few sites evolve with a ω > 1 (Supplementary Figure 9C). However, this pattern is not an indicator of a minor effect of positive selection since, in most cases, positively selected amino acid substitutions occur only in a small region of the coding sequences (Wagner, 2002).

We further analyzed the expression patterns of the 16 inparalogs groups of S. mansoni for which evidence of positive selection was found (highlighted in yellow in Supplementary Table 4). Most inparalogs in these 16 groups (46 out of 56) have no sign of pseudogenization (Median TMM among replicates in at le analyzed experiment >= 1). Only 8 of these groups have InterPro annotations, with 3 of them corresponding to Peptidase M13 groups (neprilysin family), and the remaining to Glycoside hydrolase family 5; P-type ATPase; Short-chain dehydrogenase/reductase SDR; and Bax inhibitor 1-related, Cadherin-like. Many of these families and functions have been associated with the infection process and survival of Schistosoma and other parasite species (see above) and all are secreted or surface-associated proteins.

Expression divergence among stages can be observed in the inparalogs groups F14490_SCM_G1 (Peptidase M13), F14313_SCM_G1 (Glycoside hydrolase, family 5), F14720_SCM_G1 (Peptidase M13), and F14724_SCM_G1 (Supplementary Table 6, Supplementary Figures 4, 5, and 6). In particular, we analyzed the expression pattern of the inparalogs from the F14313_SCM_G1 group (Glycoside hydrolase, family 5) in the SchistoCyte Atlas single-cell expression database (Wendt et al., 2020). This function, including these and other gene families, is critical to glycosylation homeostasis in Schistosoma and other eukaryotes (Hulme et al., 2022). We found that most inparalogs of this group are co-expressed in the same clusters of cells, which correspond to the early neoblast (stem cell) progeny and later precursors of the tegumental lineage. Thus, they may be expressed in adults mainly when neoblast cells exit the cycle and begin differentiation towards a tegumental fate. It is possible that the protein products may also be found in the differentiated tegument itself, as has been described for Tetraspanin-2 and Sm13, although their mRNAs are expressed almost exclusively in tegumental progenitors (Wendt et al., 2018). Note, however, that our results suggest that some inparalogs from this group, e.g., Smp_314170, are also highly expressed in other developmental stages (Figure 4F). This observation is not surprising given that the production of specific glycans is tightly regulated throughout the cycle of Schistosoma (for a review, see Hokke et al., 2007)

In summary, these 16 groups identified in S. mansoni, as well as others identified in the present study in which positive selection was detected, are good candidates to be involved in the adaptation process of each species and deserve further investigation.

## 4. Conclusions

Through in-silico analyses, we studied the evolution of species-specific duplicated genes within flatworms. Our results showed a conserved set of functional terms enriched among inparalogs in different species from all classes. As previously published studies showed, genes coding for these functions were systematically linked to duplications in flatworms. Thus, we proposed that these functions represent a repertoire of tools that different species of flatworms, current and ancestral, have been using recurrently throughout the evolution of the phylum, implying that selection contributes to the conservation of new duplicated copies of genes from these functional categories. Many observed overrepresented functions among inparalogs are linked to secretion, suggesting that both results are not independent. The trends observed here have been reported in many eukaryotic lineages and are widely distributed in flatworms. Therefore, this observation may not be unique to a particular niche or lifestyle and supports the idea that inherent characteristics of these functional categories may make them more prone to high duplicability. Nevertheless, this does not rule out the possibility that a selective-driven process can direct the evolution of recently duplicated genes in the specific evolutionary path of each parasitic species. A more in-depth sequence and phylogenetic analysis of each gene family within each functional category are needed to identify this kind of process.

Our analysis of transcriptomic data of S. mansoni showed that most inparalogs tend to keep the ancestral regulation constraints and probably the ancestral function of the gene, at least in short timescales. A positive dosage model, which predicts a selectively driven maintenance of duplicated copies to increase protein product, may fit better to these observations considering the specific functions overrepresented among inparalogs. However, we also identified some inparalogs that display considerable changes in the expression pattern. This observation may be explained in terms of positive or relaxed negative selection, allowing quick regulatory changes. Furthermore, applying site molecular evolution models, we could identify protein positions evolving with a dN/dS rate above one. We think this sequence divergence, probably driven by positive selection, occurs in these specific protein positions when “exploring” the functional space around the original function. Therefore, these genes should be studied functionally since they represent good candidates to be involved in the particular adaptation process of each species.

## Supporting information

Supplementary Figure 1. Workflow diagram of the bioinformatic analyses done in the present study.

Supplementary Figure 2. Heatmap of functional GO terms enrichment among inparalogous genes in each species.

Supplementary Figure 3. Distribution of peptidase families coded by inparalogs.

Distribution of secreted and transmembrane protein-coding genes within inparalogs and other homologs in studied phyla.

Supplementary Figure 5. Plots of the expression level (measured as TMM, Trimmed Means of M-values) of inparalogous groups identified in S. mansoni.

Supplementary Figure 6. Plots of the expression level (measured as TMM, Trimmed Means of M-values) of inparalogous groups identified in S. mansoni.

Supplementary Figure 7. Plots of the expression level (measured as TMM, Trimmed Means of M-values) of inparalogous groups identified in S. mansoni.

Supplementary Figure 8. Box-plots showing the distribution of Pearson correlation coefficients (r) of expression level of inparalogs by distance.

Supplementary Figure 9. Summary of results of the molecular evolution tests.

Supplementary Table 1. Accession codes of genomics and transcriptomic data used in the presented study.

Supplementary Table 2. Summary of results for secreted and transmembrane protein-coding genes in each species analyzed.

Supplementary Table 3. GO terms enrichment analysis results.

Supplementary Table 4. InterPro annotation and results for the site test of positive selection within inparalogs clusters.

Supplementary Table 5. Estimated expression level (TMM, Trimmed Means of M-values) of inparalogs identified in S. mansoni.

Supplementary Table 6. Comparative expression patterns among inparalogs in S. mansoni.

## Credit authorship contribution statement

Mauricio Langleib: Conceptualization, Validation, Software, Formal Analysis, Investigation, Data curation, Visualization, Writing - original draft. Javier Calvelo: Validation, Software, Formal Analysis, Data curation, Visualization. Alicia Costábile: Investigation, Writing - review & editing. Estela Castillo: Conceptualization, Writing - review & editing. José F. Tort: Conceptualization, Writing - review & editing. Federico G. Hoffmann: Conceptualization, Writing - review & editing. Anna V. Protasio: Conceptualization, Writing - review & editing. Uriel Koziol: Conceptualization, Investigation, Writing - review & editing. Andrés Iriarte: Conceptualization, Methodology, Investigation, Resources Writing - original draft, Writing - review & editing, Supervision, Project administration, Funding acquisition.

## Acknowledgments

This work was supported by Agencia Nacional de Investigación e Innovación (ANII), Uruguay [FCE_3_2016_1_125297] and Programa de Desarrollo de las Ciencias Básicas (PEDECIBA), Uruguay. A.C., E.C., J.F.T., U.K. and A.I. are members of the Sistema Nacional de Investigadores, Uruguay.

---

**Supplementary Table 1.** Accession codes of genomics and transcriptomic data used in the presented study. Genomic data, including genome sequence, annotation, and predicted coding sequences, were downloaded from WormBase ParaSite (parasite.wormbase.org) and the NCBI RefSeq database (www.ncbi.nlm.nih.gov/refseq). Raw reads from transcriptomic experiments were downloaded from the NCBI SRA database (www.ncbi.nlm.nih.gov/sra).

**Supplementary Table 2.** Summary of results for secreted and transmembrane protein-coding genes in each species analyzed. Mollusca and Annelida phyla were included.

**Supplementary Table 3.** GO terms enrichment analysis results. Significant enriched terms are shown (Fisher test, adjusted p-value (FDR) =< 0.05). A broader view of GO enrichment was obtained by mapping enriched GO terms to GO Slims. Ontology, Biological process (BP), Cellular component (CC), or Molecular Function (MF), is indicated for each significantly enriched functional term.

**Supplementary Table 4.** InterPro annotation and results for the site test of positive selection within inparalogs clusters identified in the analyzed Flatworm genomes. The site models M1a (neutral) vs. M2a (selection) were implemented and compared by means of the Likelihood ratio test (LRT) (Yang et al., 2005). The monophyletic section of the phylogenetic tree of the homologous gene family was used as input. The Bayes empirical Bayes (BEB) method was used for calculating the posterior probabilities (pp) for site classes. Thus, the number of identified positive-selected sites is indicated for pp > 95% and 99%. LRT higher than six is considered significant and indicated with “***”.

**Supplementary Table 5.** Estimated expression level (TMM, Trimmed Means of M-values) of inparalogs identified in S. mansoni. Expression levels were estimated from three datasets: 1) developmental stages Cercariae, Schistosomulae at 3 hr., and Schistosomulae at 24 hr. (Protasio et al., 2012); 2) Schistosomulae at the days (D) 6, 13, 17, 21 and 28, and Adults (Wangwiwatsin et al., 2020); and 3) developmental stages Miracidia, first stage Sporocyst, and second stage Sporocyst (direct submissions from the Wellcome Sanger Institute) (Supplementary Table 1).

**Supplementary Table 6.** Comparative expression patterns among inparalogs in S. mansoni. The pairwise correlation coefficients (Pearson r and Spearman rank (r_o_)) of the median TMM were estimated between inparalogs within each inparalogous cluster. Expression data of datasets were independently analyzed. The cophenetic dS estimated between inparalogs, the distance in the genome, the distance-based classification, and the InterPro annotation for each inparalog were included. A pair of inparalogs with TMM values below one in one or more of the analyzed time points within each experiment were not considered in the analysis (see column "Genes above threshold (TMM >= 1)")

**Supplementary Figure 1.** Workflow diagram of the bioinformatic analyses done in the present study.

**Supplementary Figure 2.** Heatmap of functional GO terms enrichment among inparalogous genes in each species. The categories, biological process (BP), molecular function (MF), and cellular component (CC), are indicated. Functional GO terms are ordered according to semantic similarity. Black arrows indicate terms mentioned in the main text.

**Supplementary Figure 3.** Distribution of peptidase families coded by inparalogs. Peptidases were classified based on sequence similarity to members of the MEROPS database, available at ebi.ac.uk/merops. The number of peptidases was normalized within each species and shown as a heatmap. Thus, the peptidase families with the higher frequency have a value of 1.

**Supplementary Figure 4.** Distribution of secreted and transmembrane protein-coding genes within inparalogs and other homologs in Mollusca and Annelida (A and C) and transmembrane protein-coding genes in Platyhelminthes (B). Significantly over- or under-representation of these functional categories among inparalogs is indicated with “*” (Fisher exact test, FDR < 1%). Only homologous genes (no singletons) were considered as background for this enrichment test.

**Supplementary Figure 5.** Plots of the expression level (measured as TMM, Trimmed Means of M-values) of inparalogous groups identified in S. mansoni throughout the developmental stages studied in Protasio et al. (2012). Each dot corresponds to a replicate of each stage of each inparalog. The median TMM among all replicates is plotted for each inparalog. Expression data was taken from the NCBI SRA database as indicated in Supplementary Table 1.

**Supplementary Figure 6.** Plots of the expression level (measured as TMM, Trimmed Means of M-values) of inparalogous groups identified in S. mansoni throughout the developmental stages studied in Wangwiwatsin et al. (2020). Each dot corresponds to a replicate of each stage of each inparalog. The median TMM among all replicates is plotted for each inparalog. Expression data was taken from the NCBI SRA database as indicated in Supplementary Table 1.

**Supplementary Figure 7.** Plots of the expression level (measured as TMM, Trimmed Means of M-values) of inparalogous groups identified in S. mansoni throughout the developmental stages Miracidia, First, and Second stage sporocyst. Data was directly submitted to NCBI by Wellcome Sanger Institute. Each dot corresponds to a replicate of each stage of each inparalog. The median TMM among all replicates is plotted for each inparalog. Expression data was taken from the NCBI SRA database as indicated in Supplementary Table 1.

**Supplementary Figure 8.** Box-plots showing the distribution of Pearson correlation coefficients (r) of expression level (TMM) of inparalogs classified according to physical distance in the genome. All pairs of inparalogs were classified in “Close”, “Distant”, and “Diff. chromosome” using a threshold of 10 kbp. Only genes showing a median TMM above one in one or more developmental stages, within each study, were considered in the analysis. The number of pairwise comparisons included is shown between brackets. Statistical significance was assessed by Mann-Whitney U-test and indicated by asterisks: p⍰≤⍰0.1 (*), p⍰≤⍰0.05 (**), and p⍰≤⍰0.01 (***).

**Supplementary Figure 9.** Summary of results of the molecular evolution tests. A) Distribution of inparalogous groups according to the dN/dS test results. All the inparalogous groups with a total dS estimation below 0.1 were not considered for the dN/dS test and are indicated as a green bar. The neutral model M1a, with no sites evolving under positive selection, was compared with the positive selection model M2a using the Likelihood Ratio Test in each inparalogous group. The sky blue bar indicates those groups in which the neutral model fits significantly better than the positive selection model. The magenta bar indicates those groups in which the positive selection model fits significantly better than the neutral model. Missing data, groups in which no test could be applied, were indicated in pink. B) Histogram showing the dN/dS rate distribution estimated in inparalogous groups in each species. Note that the dN/dS was estimated in each group according to the site model that fits better and that groups with dS < 0.1 were not analyzed. C) Box plot of the number of sites evolving under positive selection in each group is shown. The number of sites with dN/dS > 1 in a cluster was only considered if the positive selection model was significantly more probable than the neutral model.

## References

Aguinaldo, A.M.A., Turbeville, J.M., Linford, L.S., Rivera, M.C., Garey, J.R., Raff, R.A., Lake, J.A., 1997. Evidence for a clade of nematodes, arthropods and other moulting animals. Nature 387, 489–493. 10.1038/387489a0

Alexa, A., Rahnenfuhrer, J., 2019. topGO: Enrichment Analysis for Gene Ontology. R package version 2.36.0.

Andersson, D.I., Jerlström-Hultqvist, J., Näsvall, J., 2015. Evolution of new functions de novo and from preexisting genes. Perspectives in Biology 7, a017996. 10.1101/cshperspect.a017996

Bendtsen, J.D., Jensen, L.J., Blom, N., Von Heijne, G., Brunak, S., 2004. Feature-based prediction of non-classical and leaderless protein secretion. Protein Engineering, Design and Selection 17, 349–356. 10.1093/protein/gzh037

Bolger, A.M., Lohse, M., Usadel, B., Planck, M., Plant, M., Mühlenberg, A., 2014. Trimmomatic: A flexible trimmer for Illumina Sequence Data. Bioinformatics. 30, 2114–2120. doi: 10.1093/bioinformatics/btu170

Buchfink, B., Xie, C., Huson, D.H., 2014. Fast and sensitive protein alignment using DIAMOND. Nat Methods 12, 59–60. 10.1038/nmeth.3176

Cancela, M., Acosta, D., Rinaldi, G., Silva, E., Durán, R., Roche, L., Zaha, A., Carmona, C., Tort, J.F., 2008. A distinctive repertoire of cathepsins is expressed by juvenile invasive Fasciola hepatica. Biochimie 90, 1461–1475. 10.1016/j.biochi.2008.04.020

Casneuf, T., De Bodt, S., Raes, J., Maere, S., Van de Peer, Y., 2006. Nonrandom divergence of gene expression following gene and genome duplications in the flowering plant Arabidopsis thaliana. Genome Biol 7, R13. 10.1186/gb-2006-7-2-r13

Castresana, J., 2000. Selection of conserved blocks from multiple alignments for their use in phylogenetic analysis. Mol Biol Evol 17, 540–552. 10.1093/oxfordjournals.molbev.a026334

Chalmers, I.W., Mcardle, A.J., Coulson, R., Wagner, M.A., Schmid, R., Hirai, H., Hoffmann, K.F., 2008. Developmentally regulated expression, alternative splicing and distinct sub-groupings in members of the Schistosoma mansoni venom allergen-like (SmVAL) gene family. BMC Genomics 20, 89. 10.1186/1471-2164-9-89

Chen, X., Zhang, J., 2012. The Ortholog Conjecture Is Untestable by the Current Gene Ontology but Is Supported by RNA Sequencing Data. PLoS Comput Biol 8, e1002784. 10.1371/journal.pcbi.1002784

Chiumiento, L., Bruschi, F., 2009. Enzymatic antioxidant systems in helminth parasites. Parasitol Res 105, 593–603. 10.1007/s00436-009-1483-0

Choi, Y.J., Fontenla, S., Fischer, P.U., Le, T.H., Costábile, A., Blair, D., Brindley, P.J., Tort, J.F., Cabada, M.M., Mitreva, M., 2020. Adaptive Radiation of the Flukes of the Family Fasciolidae Inferred from Genome-Wide Comparisons of Key Species. Mol Biol Evol 37, 84–99. 10.1093/molbev/msz204

Chow, C., Gauci, C.G., Cowman, A.F., Lightowlers, M.W., 2001. A gene family expressing a host-protective antigen of Echinococcus granulosus. Mol Biochem Parasitol 118, 83–88. 10.1016/S0166-6851(01)00373-5

Coghlan, A., Tyagi, R., Cotton, J.A., Holroyd, N., Rosa, B.A., Tsai, I.J., Laetsch, D.R., Beech, R.N., Day, T.A., Hallsworth-Pepin, K., Ke, H.M., Kuo, T.H., Lee, T.J., Martin, J., Maizels, R.M., Mutowo, P., Ozersky, P., Parkinson, J., Reid, A.J., Rawlings, N.D., Ribeiro, D.M., Swapna, L.S., Stanley, E., Taylor, D.W., Wheeler, N.J., Zamanian, M., Zhang, X., Allan, F., Allen, J.E., Asano, K., Babayan, S.A., Bah, G., Beasley, H., Bennett, H.M., Bisset, S.A., Castillo, E., Cook, J., Cooper, P.J., Cruz-Bustos, T., Cuéllar, C., Devaney, E., Doyle, S.R., Eberhard, M.L., Emery, A., Eom, K.S., Gilleard, J.S., Gordon, D., Harcus, Y., Harsha, B., Hawdon, J.M., Hill, D.E., Hodgkinson, J., Horák, P., Howe, K.L., Huckvale, T., Kalbe, M., Kaur, G., Kikuchi, T., Koutsovoulos, G., Kumar, S., Leach, A.R., Lomax, J., Makepeace, B., Matthews, J.B., Muro, A., O’Boyle, N.M., Olson, P.D., Osuna, A., Partono, F., Pfarr, K., Rinaldi, G., Foronda, P., Rollinson, D., Samblas, M.G., Sato, H., Schnyder, M., Scholz, T., Shafie, M., Tanya, V.N., Toledo, R., Tracey, A., Urban, J.F., Wang, L.C., Zarlenga, D., Blaxter, M.L., Mitreva, M., Berriman, M., 2019. Comparative genomics of the major parasitic worms. Nat Genet 51, 163–174. 10.1038/s41588-018-0262-1

Contreras-Moreira, B., Vinuesa, P., 2013. GET_HOMOLOGUES, a versatile software package for scalable and robust microbial pangenome analysis. Appl Environ Microbiol 79, 7696–7701. 10.1128/AEM.02411-13

Corvo, I., Cancela, M., Cappetta, M., Pi-Denis, N., Tort, J.F., Roche, L., 2009. The major cathepsin L secreted by the invasive juvenile Fasciola hepatica prefers proline in the S2 subsite and can cleave collagen. Mol Biochem Parasitol 167, 41–47. 10.1016/j.molbiopara.2009.04.005

Corvo, I., O’Donoghue, A.J., Pastro, L., Pi-Denis, N., Eroy-Reveles, A., Roche, L., McKerrow, J.H., Dalton, J.P., Craik, C.S., Caffrey, C.R., Tort, J.F., 2013. Dissecting the Active Site of the Collagenolytic Cathepsin L3 Protease of the Invasive Stage of Fasciola hepatica. PLoS Negl Trop Dis 7, e2269. 10.1371/journal.pntd.0002269

Costábile, A., Koziol, U., Tort, J.F., Iriarte, A., Castillo, E., 2018. Expansion of cap superfamily proteins in the genome of Mesocestoides corti: An extreme case of a general bilaterian trend. Gene Rep 11, 110–120. 10.1016/j.genrep.2018.03.010

Cuesta-Astroz, Y., de Oliveira, F.S., Nahum, L.A., Oliveira, G., 2017. Helminth secretomes reflect different lifestyles and parasitized hosts. Int J Parasitol 47, 529–544. 10.1016/j.ijpara.2017.01.007

Curwen, R.S., Ashton, P.D., Sundaralingam, S., Wilson, R.A., 2006. Identification of novel proteases and immunomodulators in the secretions of schistosome cercariae that facilitate host entry. Molecular & Cellular Proteomics 5, 835–844. 10.1074/mcp.M500313-MCP200

Cwiklinski, K., Dalton, J.P., Dufresne, P.J., La Course, J., Williams, D.J.L., Hodgkinson, J., Paterson, S., 2015. The Fasciola hepatica genome: Gene duplication and polymorphism reveals adaptation to the host environment and the capacity for rapid evolution. Genome Biol 16, 71. 10.1186/s13059-015-0632-2

Dalton, J.P., Skelly, P., Halton, D.W., 2004. Role of the tegument and gut in nutrient uptake by parasitic platyhelminths. Can J Zool 82, 211–232. 10.1139/z03-213

del Puerto, L., Rovetta, R., Navatta, M., Fontana, C., Lin, G., Moyna, G., Dematteis, S., Brehm, K., Koziol, U., Ferreira, F., Díaz, A., 2016. Negligible elongation of mucin glycans with Gal β1-3 units distinguishes the laminated layer of Echinococcus multilocularis from that of Echinococcus granulosus. Int J Parasitol 46, 311–321. 10.1016/j.ijpara.2015.12.009

Dobin, A., Davis, C.A., Schlesinger, F., Drenkow, J., Zaleski, C., Jha, S., Batut, P., Chaisson, M., Gingeras, T.R., 2013. STAR: Ultrafast universal RNA-seq aligner. Bioinformatics 29, 15–21. 10.1093/bioinformatics/bts635

Drummond, D.A., Wilke, C.O., 2009. Mistranslation-induced protein misfolding as a dominant constraint on coding-sequence evolution. Cell 134, 341–352. 10.1016/j.cell.2008.05.042.Mistranslation-induced

Dunn, C.W., Giribet, G., Edgecombe, G.D., Hejnol, A., 2014. Animal Phylogeny and Its Evolutionary Implications. Annu Rev Ecol Evol Syst 45, 371–395. 10.1146/annurev-ecolsys-120213-091627

Edgar, R.C., 2004. MUSCLE: Multiple sequence alignment with high accuracy and high throughput. Nucleic Acids Res 32, 1792–1797. 10.1093/nar/gkh340

Egger, B., Lapraz, F., Tomiczek, B., Müller, S., Dessimoz, C., Girstmair, J., Škunca, N., Rawlinson, K.A., Cameron, C.B., Beli, E., Todaro, M.A., Gammoudi, M., Noreña, C., Telford, M.J., 2015. A transcriptomic-phylogenomic analysis of the evolutionary relationships of flatworms. Current Biology 25, 1347–1353. 10.1016/j.cub.2015.03.034

Emanuelsson, O., Brunak, S., von Heijne, G., Nielsen, H., 2007. Locating proteins in the cell using TargetP, SignalP and related tools. Nat Protoc 2, 953–971. 10.1038/nprot.2007.131

Emmanoch, P., Kosa, N., Vichasri-Grams, S., Tesana, S., Grams, R., Geadkaew-Krenc, A., 2018. Comparative characterization of four calcium-binding EF hand proteins from opisthorchis viverrini. Korean Journal of Parasitology 56, 81–86. 10.3347/kjp.2018.56.1.81

Ewels, P., Magnusson, M., Lundin, S., Käller, M., 2016. MultiQC: Summarize analysis results for multiple tools and samples in a single report. Bioinformatics 32, 3047–3048. 10.1093/bioinformatics/btw354

Feasey, N., Wansbrough-Jones, M., Mabey, D.C.W., Solomon, A.W., 2010. Neglected tropical diseases. Br Med Bull 93, 179–200. 10.1093/bmb/ldp046

Feyertag, F., Berninsone, P.M., Alvarez-ponce, D., 2016. Secreted Proteins Defy the Expression Level – Evolutionary Rate Anticorrelation. Mol Biol Evol 34, 692–706. 10.1093/molbev/msw268

Fitzsimmons, C.M., Jones, F.M., Stearn, A., Chalmers, I.W., Hoffmann, K.F., Wawrzyniak, J., Wilson, S., Kabatereine, N.B., Dunne, D.W., 2012. The Schistosoma mansoni Tegumental-Allergen-Like (TAL) Protein Family: Influence of Developmental Expression on Human IgE Responses. PLoS Negl Trop Dis 6, e1593. 10.1371/journal.pntd.0001593

Fló, M., Margenat, M., Pellizza, L., Graña, M., Durán, R., Báez, A., Salceda, E., Soto, E., Alvarez, B., Fernández, C., 2017. Functional diversity of secreted cestode Kunitz proteins: Inhibition of serine peptidases and blockade of cation channels. PLoS Pathog 13, e1006169. 10.1371/journal.ppat.1006169

Franchini, G.R., Pórfido, J.L., Ibáñez Shimabukuro, M., Rey Burusco, M.F., Bélgamo, J.A., Smith, B.O., Kennedy, M.W., Córsico, B., 2015. The unusual lipid binding proteins of parasitic helminths and their potential roles in parasitism and as therapeutic targets. Prostaglandins Leukot Essent Fatty Acids 93, 31–36. 10.1016/j.plefa.2014.08.003

Francino, M.P., 2005. An adaptive radiation model for the origin of new gene functions. Nat Genet 37, 537–537. 10.1038/ng1579

Gabaldón, T., Koonin, E. V, 2013. Functional and evolutionary implications of gene orthology. Nat Rev Genet 14, 360–366. 10.1038/nrg3456

Garg, G., Ranganathan, S., 2011. In silico secretome analysis approach for next generation sequencing transcriptomic data. BMC Genomics 12, S14. 10.1186/1471-2164-12-S3-S14

González, S., Fló, M., Margenat, M., Durán, R., González-Sapienza, G., Graña, M., Parkinson, J., Maizels, R.M., Salinas, G., Alvarez, B., Fernández, C., 2009. A family of diverse Kunitz inhibitors from Echinococcus granulosus potentially involved in host-parasite cross-talk. PLoS One 4, e7009. 10.1371/journal.pone.0007009

Haag, K.L., Gottstein, B., Ayala, F.J., 2009. The EG95 antigen of Echinococcus spp. contains positively selected amino acids, which may influence host specificity and vaccine efficacy. PLoS One 4, e5362. 10.1371/journal.pone.0005362

Hagberg, A.A., Schult, D.A., Swart, P.J., 2008. Exploring network structure, dynamics, and function using NetworkX, in: Varoquaux, G., Vaught, T., Millman, J. (Eds.), Proceedings of the 7th Python in Science Conference (SciPy2008). Pasadena, CA USA, pp. 11–15.

Hayes, J.D., Flanagan, J.U., Jowsey, I.R., 2005. Glutathione transferases. Annu Rev Pharmacol Toxicol 45, 51–88. 10.1146/annurev.pharmtox.45.120403.095857

He, X., Zhang, J., 2005. Rapid subfunctionalization accompanied by prolonged and substantial neofunctionalization in duplicate gene evolution. Genetics 169, 1157–1164. 10.1534/genetics.104.037051

Hewitson, J.P., Grainger, J.R., Maizels, R.M., 2009. Helminth immunoregulation: The role of parasite secreted proteins in modulating host immunity. Mol Biochem Parasitol 167, 1–11. 10.1016/j.molbiopara.2009.04.008

Hickman, C., Roberts, L., Keen, S., Larson, A., Anson, H., Eisenhour, D., 2008. Integrated Principles of Zoology, 14th ed. McGraw-Hill Higher Education, Boston, Massachusetts.

Hoang, D.T., Chernomor, O., von Haeseler, A., Minh, B.Q., Vinh, L.S., 2018. UFBoot2: Improving the Ultrafast Bootstrap Approximation. Mol Biol Evol 35, 518–522. 10.5281/zenodo.854445

Hokke, C.H., Fitzpatrick, J.M., Hoffmann, K.F., 2007. Integrating transcriptome, proteome and glycome analyses of Schistosoma biology. Trends Parasitol 23, 165–174. 10.1016/j.pt.2007.02.007

Howe, K.L., Bolt, B.J., Shafie, M., Kersey, P., Berriman, M., 2017. WormBase ParaSite − a comprehensive resource for helminth genomics. Mol Biochem Parasitol 215, 2–10. 10.1016/j.molbiopara.2016.11.005

Huang, F., Dang, Z., Suzuki, Y., Horiuchi, T., Yagi, K., 2016. Analysis on Gene Expression Profile in Oncospheres and Early Stage Metacestodes from Echinococcus multilocularis. PLoS Negl Trop Dis 10, e0004634. 10.1371/journal.pntd.0004634

Huerta-Cepas, J., Serra, F., Bork, P., 2016a. ETE 3: Reconstruction, Analysis, and Visualization of Phylogenomic Data. Mol Biol Evol 33, 1635–1638. 10.1093/molbev/msw046

Huerta-Cepas, J., Szklarczyk, D., Forslund, K., Cook, H., Heller, D., Walter, M.C., Rattei, T., Mende, D.R., Sunagawa, S., Kuhn, M., Jensen, L.J., Von Mering, C., Bork, P., 2016b. EGGNOG 4.5: A hierarchical orthology framework with improved functional annotations for eukaryotic, prokaryotic and viral sequences. Nucleic Acids Res 44, D286–D293. 10.1093/nar/gkv1248

Hulme, B.J., Geyer, K.K., Forde-Thomas, J.E., Padalino, G., Phillips, D.W., Ittiprasert, W., Karinshak, S.E., Mann, V.H., Chalmers, I.W., Brindley, P.J., Hokke, C.H., Hoffmann, K.F., 2022. Schistosoma mansoni α-N-acetylgalactosaminidase (SmNAGAL) regulates coordinated parasite movement and egg production. PLoS Pathog 18, e1009828. 10.1371/journal.ppat.1009828

Kirkness, E.F., Haas, B.J., Sun, W., Braig, H.R., Perotti, M.A., Clark, J.M., Lee, S.H., Robertson, H.M., Kennedy, R.C., Elhaik, E., Gerlach, D., Kriventseva, E. V, Elsik, C.G., Graur, D., Hill, C.A., Veenstra, J.A., Walenz, B., Tubío, J.M.C., Ribeiro, J.M.C., Rozas, J., Johnston, J.S., Reese, J.T., Popadic, A., Tojo, M., Raoult, D., Reed, D.L., Tomoyasu, Y., Krause, E., Mittapalli, O., Margam, V.M., Li, H.M., Meyer, J.M., Johnson, R.M., Romero-Severson, J., VanZee, J.P., Alvarez-Ponce, D., Vieira, F.G., Aguadé, M., Guirao-Rico, S., Anzola, J.M., Yoon, K.S., Strycharz, J.P., Unger, M.F., Christley, S., Lobo, N.F., Seufferheld, M.J., Wang, N.K., Dasch, G.A., Struchiner, C.J., Madey, G., Hannick, L.I., Bidwell, S., Joardar, V., Caler, E., Shao, R., Barker, S.C., Cameron, S., Bruggner, R. V, Regier, A., Johnson, J., Viswanathan, L., Utterback, T.R., Sutton, G.G., Lawson, D., Waterhouse, R.M., Venter, J.C., Strausberg, R.L., Berenbaum, M.R., Collins, F.H., Zdobnov, E.M., Pittendrigh, B.R., 2010. Genome sequences of the human body louse and its primary endosymbiont provide insights into the permanent parasitic lifestyle. Proc Natl Acad Sci U S A 107, 12168–12173. 10.1073/pnas.1003379107

Klopfenstein, D. V., Zhang, L., Pedersen, B.S., Ramírez, F., Vesztrocy, A.W., Naldi, A., Mungall, C.J., Yunes, J.M., Botvinnik, O., Weigel, M., Dampier, W., Dessimoz, C., Flick, P., Tang, H., 2018. GOATOOLS: A Python library for Gene Ontology analyses. Sci Rep 8, 10872. 10.1038/s41598-018-28948-z

Kondrashov, F.A., Rogozin, I.B., Wolf, Y.I., Koonin, E. V, 2002. Selection in the evolution of gene duplications. Genome Biol 3, RESEARCH0008. 10.1186/gb-2002-3-2-research0008

Koonin, E. V, 2005. Orthologs, Paralogs, and Evolutionary Genomics. Annu Rev Genet 39, 309–338. 10.1146/annurev.genet.39.073003.114725

Krogh, A., Larsson, B., von Heijne, G., Sonnhammer, E.L., 2001. Predicting transmembrane protein topology with a hidden Markov model: application to complete genomes. J Mol Biol 305, 567–580. 10.1006/jmbi.2000.4315

Kuhn, M., Jackson, S., Cimentad, J., 2020. corrr: Correlations in R. R package version 0.4.3.

Leinonen, R., Sugawara, H., Shumway, M., 2011. The sequence read archive. Nucleic Acids Res 39, 2010–2012. 10.1093/nar/gkq1019

Li, L., Stoeckert, C.J.J., Roos, D.S., 2003. OrthoMCL: Identification of Ortholog Groups for Eukaryotic Genomes. Genome Res 13, 2178–2189. 10.1101/gr.1224503

Lightowlers, M.W., Rickard, M.D., 1988. Excretory-secretory products of helminth parasites: Effects on host immune responses. Parasitology 96, S123–S166. 10.1017/S0031182000086017

Littlewood, D.T.J., 2006. Parasitic flatworms: molecular biology, biochemistry, immunology and physiology. CABI, Wallingford. 10.1079/9780851990279.0001

Lynch, M., Katju, V., 2004. The altered evolutionary trajectories of gene duplicates. Trends in Genetics 20, 544–549. 10.1016/j.tig.2004.09.001

Mambelli, F., Santos, B.P.O., Morais, S.B., Gimenez, E.G.T., Astoni, D.C. dos S., Braga, A.D., Ferreira, R.S., Amaral, F.A., Magalhães, M.T.Q. de, Oliveira, S.C., 2021. S. mansoni Sm KI-1 Kunitz-domain: Leucine point mutation at P1 site generates enhanced neutrophil elastase inhibitory activity. PLoS Negl Trop Dis 15, e0009007. 10.1371/journal.pntd.0009007

McKerrow, J.H., Caffrey, C., Kelly, B., Loke, P., Sajid, M., 2006. Proteases in parasitic diseases. Annual Review of Pathology: Mechanisms of Disease 1, 497–536. 10.1146/annurev.pathol.1.110304.100151

Nehrt, N.L., Clark, W.T., Radivojac, P., Hahn, M.W., 2011. Testing the ortholog conjecture with comparative functional genomic data from mammals. PLoS Comput Biol 7, e1002073. 10.1371/journal.pcbi.1002073

Nguyen, L.T., Schmidt, H.A., Von Haeseler, A., Minh, B.Q., 2015. IQ-TREE: A fast and effective stochastic algorithm for estimating maximum-likelihood phylogenies. Mol Biol Evol 32, 268–274. 10.1093/molbev/msu300

Olson, P.D., Zarowiecki, M., James, K., Baillie, A., Bartl, G., Burchell, P., Chellappoo, A., Jarero, F., Tan, L.Y., Holroyd, N., Berriman, M., 2018. Genome-wide transcriptome profiling and spatial expression analyses identify signals and switches of development in tapeworms. Evodevo 9, 1–29. 10.1186/s13227-018-0110-5

Pa, C., Hurst, L.D., 2003. Dosage sensitivity and the evolution of gene families in yeast. Nature 424, 194–197. 10.1038/nature01713.1.

Paradis, E., Schliep, K., 2019. Ape 5.0: An environment for modern phylogenetics and evolutionary analyses in R. Bioinformatics 35, 526–528. 10.1093/bioinformatics/bty633

Petersen, T.N., Brunak, S., Von Heijne, G., Nielsen, H., 2011. SignalP 4.0: Discriminating signal peptides from transmembrane regions. Nat Methods 8, 785–786. 10.1038/nmeth.1701

Protasio, A. V., Tsai, I.J., Babbage, A., Nichol, S., Hunt, M., Aslett, M.A., de Silva, N., Velarde, G.S., Anderson, T.J.C., Clark, R.C., Davidson, C., Dillon, G.P., Holroyd, N.E., LoVerde, P.T., Lloyd, C., McQuillan, J., Oliveira, G., Otto, T.D., Parker-Manuel, S.J., Quail, M.A., Wilson, R.A., Zerlotini, A., Dunne, D.W., Berriman, M., 2012. A systematically improved high quality genome and transcriptome of the human blood fluke Schistosoma mansoni. PLoS Negl Trop Dis 6, e1455. 10.1371/journal.pntd.0001455

Putri, G.H., Anders, S., Pyl, P.T., Pimanda, J.E., Zanini, F., 2022. Analysing high-throughput sequencing data in Python with HTSeq 2.0. Bioinformatics 38, 2943–2945. 10.1093/bioinformatics/btac166

Qian, W., Liao, B.-Y., Chang, A.Y.-F., Zhang, J., 2011. Maintenance of duplicate genes and their functional redundancy by reduced expression. Trends in Genetics 26, 425–430. 10.1016/j.tig.2010.07.002.Maintenance

Qian, W., Zhang, J., 2008. Gene Dosage and Gene Duplicability Wenfeng. Genetics 179, 2319–2324. 10.1534/genetics.108.090936

Rawlings, N.D., Barrett, A.J., Thomas, P.D., Huang, X., Bateman, A., Finn, R.D., 2018. The MEROPS database of proteolytic enzymes, their substrates and inhibitors in 2017 and a comparison with peptidases in the PANTHER database. Nucleic Acids Res 46, D624– D632. 10.1093/nar/gkx1134

Remm, M., Storm, C.E. V, Sonnhammer, E.L.L., 2001. Automatic clustering of orthologs and in-paralogs from pairwise species comparisons. J Mol Biol 314, 1041–1052. 10.1006/jmbi.2000.5197

Robinson, M.D., McCarthy, D.J., Smyth, G.K., 2009. edgeR: A Bioconductor package for differential expression analysis of digital gene expression data. Bioinformatics 26, 139–140. 10.1093/bioinformatics/btp616

Robinson, M.D., Oshlack, A., 2010. A scaling normalization method for differential expression analysis of RNA-seq data. Genome Biol 11, R25. 10.1186/gb-2010-11-3-r25

Rofatto, H.K., Parker-manuel, S.J., Barbosa, T.C., Tararam, C.A., Wilson, R.A., Leite, L.C.C., Farias, L.P., 2012. Tissue expression patterns of Schistosoma mansoni Venom Allergen-Like proteins 6 and 7. Int J Parasitol 42, 613–620. 10.1016/j.ijpara.2012.04.008

Rogozin, I.B., 2014. Complexity of gene expression evolution after duplication: Protein dosage rebalancing. Genet Res Int 2014, 516508. 10.1155/2014/516508

Sánchez, F., Garcia, J., March, F., Cardeñosa, N., Coll, P., Muñoz, C., Auladell, C., Prats, G., 1993. Ultrastructural localization of major hydatid fluid antigens in brood capsules and protoscoleces of Echinococcus granulosus of human origin. Parasite Immunol 15, 441–447. 10.1111/j.1365-3024.1993.tb00629.x

Silva-álvarez, V., Maite, A., Lía, A., Zamarreño, F., Costabel, D., García-zepeda, E., Salinas, G., Córsico, B., María, A., 2015. Echinococcus granulosus antigen B: A Hydrophobic Ligand Binding Protein at the host – parasite interface. Prostaglandins Leukot Essent Fatty Acids 93, 17–23. 10.1016/j.plefa.2014.09.008

Smith, D., Cwiklinski, K., Jewhurst, H., Tikhonova, I.G., Dalton, J.P., 2020. An atypical and functionally diverse family of Kunitz - type cysteine/serine proteinase inhibitors secreted by the helminth parasite Fasciola hepatica. Scientific Reportseports 10, 20657. 10.1038/s41598-020-77687-7

Sonnhammer, E.L.L., Koonin, E. V, 2002. Orthology, paralogy and proposed classification for paralog subtypes. Trends in G 18, 619–620. 10.1016/S0168-9525(02)02793-2

Stamboulian, M., Guerrero, R.F., Hahn, M.W., Radivojac, P., 2020. The ortholog conjecture revisited: the value of orthologs and paralogs in function prediction. Bioinformatics 36, i219–i226. 10.1093/bioinformatics/btaa468

Suyama, M., Torrents, D., Bork, P., 2006. PAL2NAL: Robust conversion of protein sequence alignments into the corresponding codon alignments. Nucleic Acids Res 34, W609– W612. 10.1093/nar/gkl315

Tsai, I.J., Zarowiecki, M., Holroyd, N., Garciarrubio, A., Sanchez-Flores, A., Brooks, K.L., Tracey, A., Bobes, R.J., Fragoso, G., Sciutto, E., Aslett, M., Beasley, H., Bennett, H.M., Cai, J., Camicia, F., Clark, R., Cucher, M., De Silva, N., Day, T.A., Deplazes, P., Estrada, K., Fernández, C., Holland, P.W.H., Hou, J., Hu, S., Huckvale, T., Hung, S.S., Kamenetzky, L., Keane, J.A., Kiss, F., Koziol, U., Lambert, O., Liu, K., Luo, X., Luo, Y., MacChiaroli, N., Nichol, S., Paps, J., Parkinson, J., Pouchkina-Stantcheva, N., Riddiford, N., Rosenzvit, M., Salinas, G., Wasmuth, J.D., Zamanian, M., Zheng, Y., Cai, X., Soberon, X., Olson, P.D., Laclette, J.P., Brehm, K., Berriman, M., Morett, E., Portillo, T., Jose, M. V., Carrero, J.C., Larralde, C., Morales-Montor, J., Limon-Lason, J., Cevallos, M.A., Gonzalez, V., Ochoa-Leyva, A., Landa, A., Jimenez, L., Valdes, V., 2013. The genomes of four tapeworm species reveal adaptations to parasitism. Nature 496, 57–63. 10.1038/nature12031

Wagner, A., 2005. Energy constraints on the evolution of gene expression. Mol Biol Evol 22, 1365–1374. 10.1093/molbev/msi126

Wagner, A., 2002. Selection and gene duplication: A view from the genome. Genome Biol 3, reviews1012.1–reviews1012.3. 10.1186/gb-2002-3-5-reviews1012

Wang, L.G., Lam, T.T.Y., Xu, S., Dai, Z., Zhou, L., Feng, T., Guo, P., Dunn, C.W., Jones, B.R., Bradley, T., Zhu, H., Guan, Y., Jiang, Y., Yu, G., 2020. Treeio: An R Package for Phylogenetic Tree Input and Output with Richly Annotated and Associated Data. Mol Biol Evol 37, 599–603. 10.1093/molbev/msz240

Wang, Y., Wang, X., Paterson, A.H., 2012. Genome and gene duplications and gene expression divergence: A view from plants. Ann N Y Acad Sci 1256, 1–14. 10.1111/j.1749-6632.2011.06384.x

Wang, Y., Xiao, D., Shen, Y., Han, X., Zhao, F., Li, X., Wu, W., Zhou, H., Zhang, J., Cao, J., 2015. Proteomic analysis of the excretory/secretory products and antigenic proteins of Echinococcus granulosus adult worms from infected dogs. BMC Vet Res 11, 119. 10.1186/s12917-015-0423-8

Wangwiwatsin, A., Protasio, A. V., Wilson, S., Owusu, C., Holroyd, N.E., Sanders, M.J., Keane, J., Doenhoff, M.J., Rinaldi, G., Berriman, M., 2020. Transcriptome of the parasitic flatworm Schistosoma mansoni during intra-mammalian development. PLoS Negl Trop Dis 14, e0007743. 10.1371/journal.pntd.0007743

Waterhouse, R.M., Zdobnov, E.M., Kriventseva, E. V, 2010. Correlating Traits of Gene Retention, Sequence. Genome Biol Evol 2, 75–86. 10.1093/gbe/evq083

Wendt, G., Lu Zhao, Chen, R., Liu, C., O’Donoghue, A.J., Caffrey, C.R., Reese, M.L., Collins III, J.J., 2020. A single-cell RNAseq atlas of Schistosoma mansoni identifies a key regulator of blood feeding. Physiol Behav 369, 1644–1649. doi:10.1126/science.abb7709

Wendt, G.R., Collins, J.N., Pei, J., Pearson, M.S., Bennett, H.M., Loukas, A., Berriman, M., Grishin, N. V, Collins, J.J., 2018. Flatworm-specific transcriptional regulators promote the specification of tegumental progenitors in Schistosoma mansoni. Elife 7, e33221. 10.7554/eLife.33221

Wickham, H., 2016. ggplot2: Elegant Graphics for Data Analysis, 2nd ed, Journal of the Royal Statistical Society: Series A (Statistics in Society). Springer International Publishing, New York. 10.1007/978-0-387-98141-3

Wickham, H., Averick, M., Bryan, J., Chang, W., McGowan, L., François, R., Grolemund, G., Hayes, A., Henry, L., Hester, J., Kuhn, M., Pedersen, T., Miller, E., Bache, S., Müller, K., Ooms, J., Robinson, D., Seidel, D., Spinu, V., Takahashi, K., Vaughan, D., Wilke, C., Woo, K., Yutani, H., 2019. Welcome to the Tidyverse. J Open Source Softw 4, 1686. 10.21105/joss.01686

Yang, Z., 2007. PAML 4: Phylogenetic analysis by maximum likelihood. Mol Biol Evol 24, 1586–1591. 10.1093/molbev/msm088

Yang, Z., Bielawski, J.P., 2000. Statistical methods for detecting molecular adaptation. Trends Ecol Evol 15, 496–503. 10.1016/s0169-5347(00)01994-7

Yang, Z., Wong, W.S.W., Nielsen, R., 2005. Bayes empirical Bayes inference of amino acid sites under positive selection. Mol Biol Evol 22, 1107–1118. 10.1093/molbev/msi097

Yoshino, T.P., Brown, M., Wu, X., Jackson, C.J., Ocadiz-ruiz, R., Chalmers, I.W., Kolb, M., Hokke, C.H., Hoffmann, K.F., 2014. Excreted/secreted Schistosoma mansoni venom allergen-like 9 (SmVAL9) modulates host extracellular matrix remodelling gene expression. Int J Parasitol 9, 18–29. 10.1016/j.ijpara.2014.04.002

Zadesenets, K.S., Vizoso, D.B., Schlatter, A., Konopatskaia, I.D., Berezikov, E., Schärer, L., Rubtsov, N.B., 2016. Evidence for karyotype polymorphism in the free-living flatworm, macrostomum lignano, a model organism for evolutionary and developmental biology. PLoS One 11, e0164915. 10.1371/journal.pone.0164915

Zhang, J., Yang, J.-R., 2015. Determinants of the rate of protein sequence evolution. Physiol Behav 16, 409–420. 10.1038/nrg3950

Zheng, A.Y., Guo, X., Su, M., Jin, X., Ding, J., Wang, Z., Ayaz, M., Kutyrev, I., Jia, W., 2018. Identification of emu-TegP11, an EF-hand domain-containing tegumental protein of Echinococcus multilocularis. Vet Parasitol 255, 107–113. 10.1016/j.vetpar.2018.04.006

