## Supplementary Figure 1. Workflow diagram of the bioinformatic analyses done in the present study. for "Evolutionary analysis of genome-specific duplications in flatworm genomes"

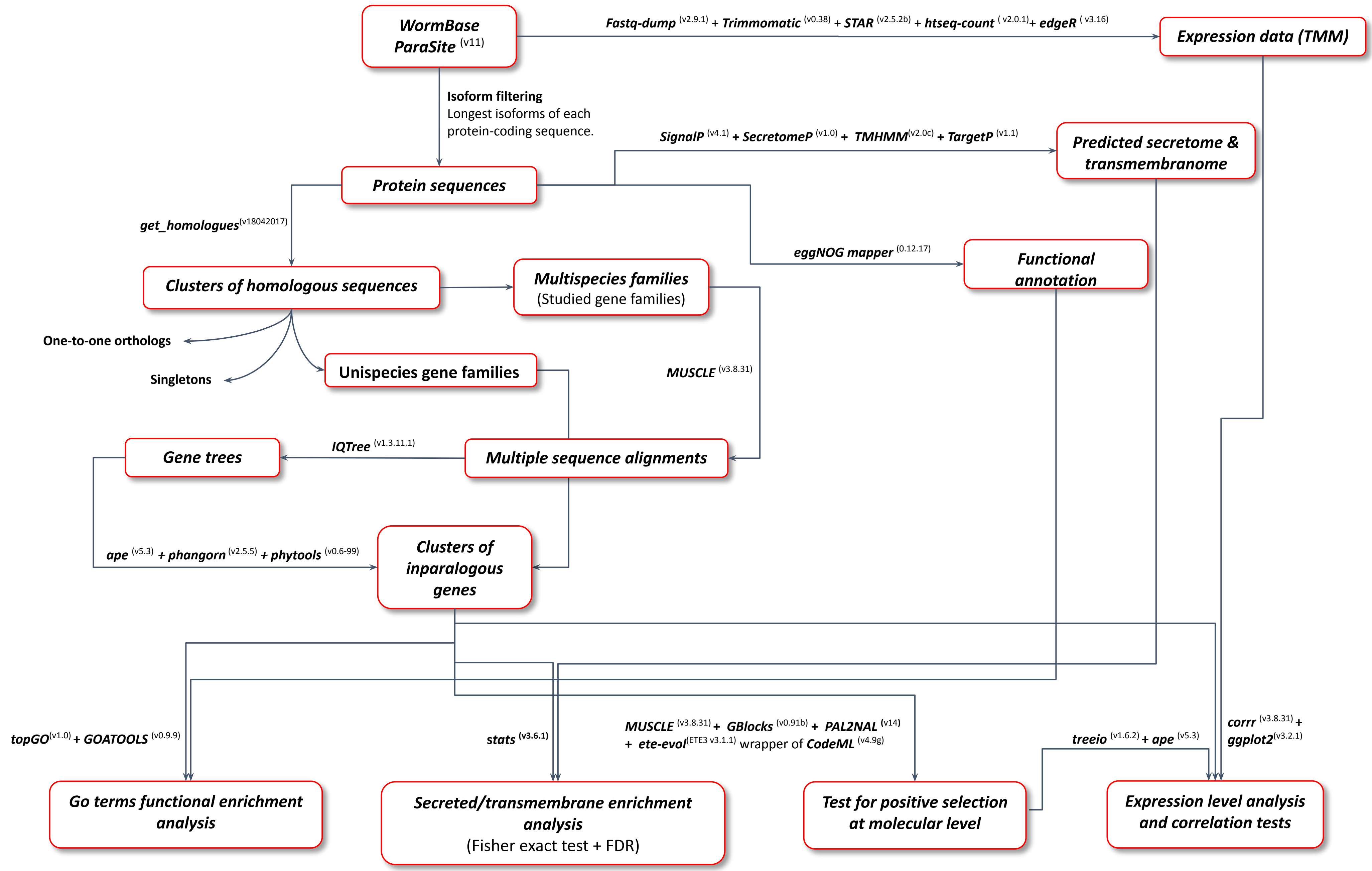


Supplementary Figure 1. Workflow diagram of the bioinformatic analyses done in the present study.
