## Supplementary Figure 2. Heatmap of functional GO terms enrichment among inparalogous genes in each species. for "Evolutionary analysis of genome-specific duplications in flatworm genomes"

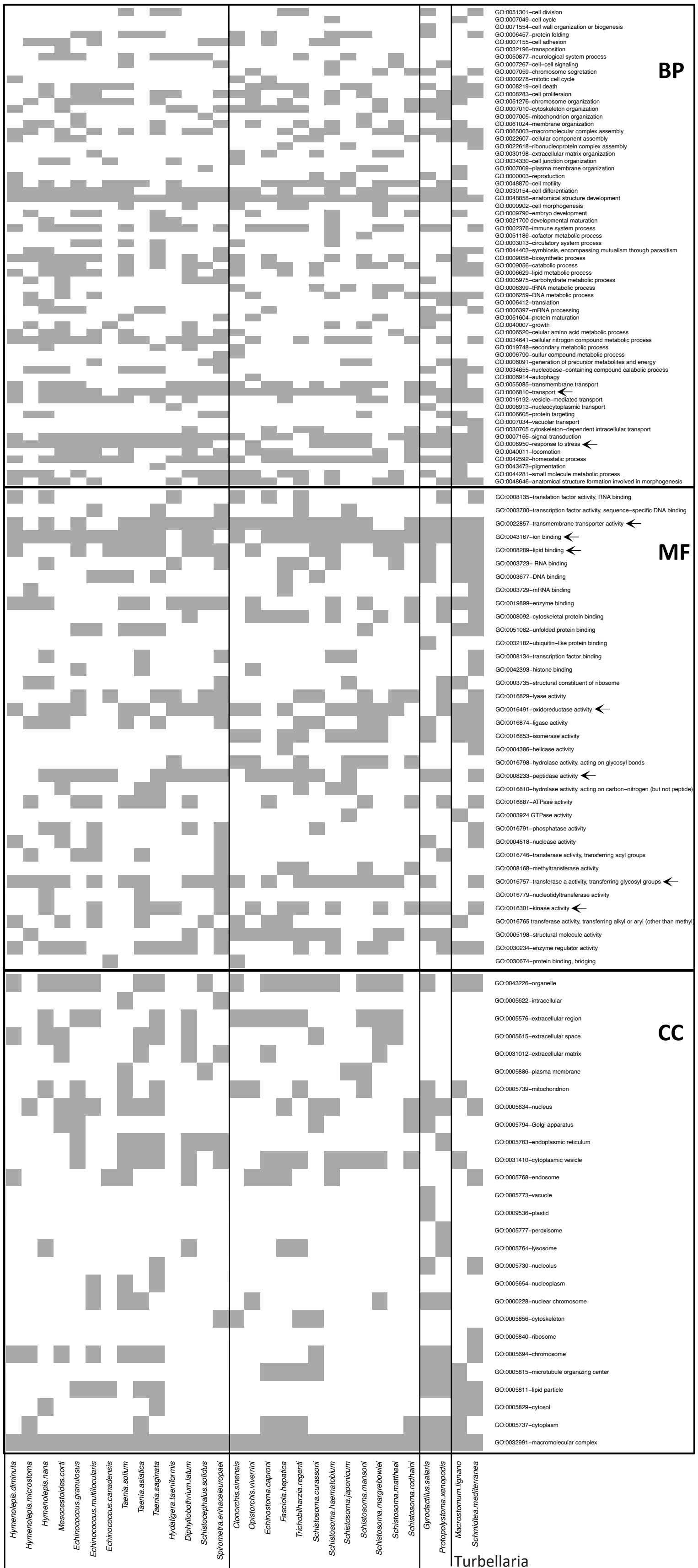


Supplementary Figure 2. Heatmap of functional GO terms enrichment among inparalogous genes in each species. The categories, biological process (BP), molecular function (MF), and cellular component (CC), are indicated. Functional GO terms are ordered according to semantic similarity. Black arrows indicate terms mentioned in the main text.
