## Supplementary Figure 3. Distribution of peptidase families coded by inparalogs. for "Evolutionary analysis of genome-specific duplications in flatworm genomes"

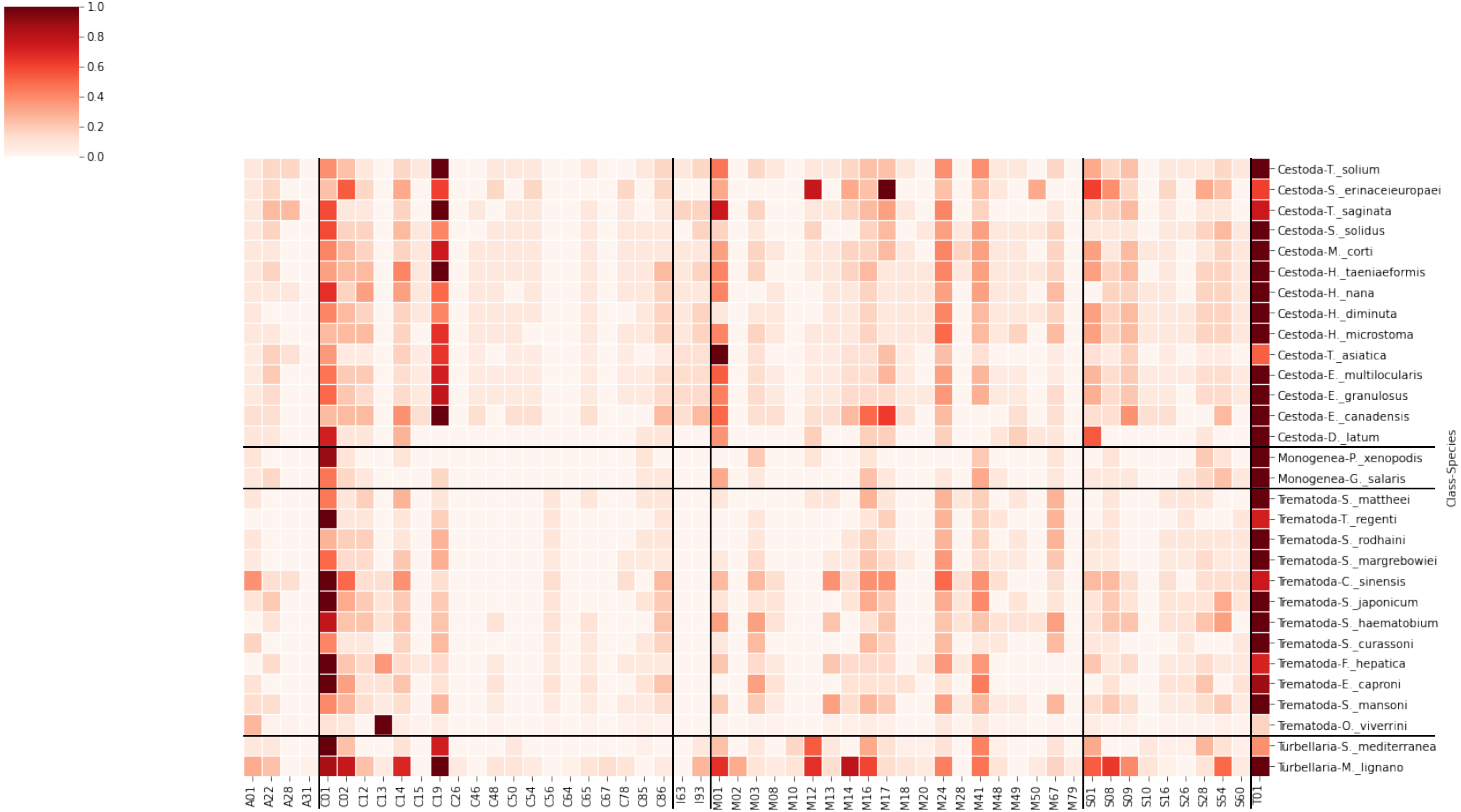


Supplementary Figure 3. Distribution of peptidase families coded by inparalogs. Peptidases were classified based on sequence similarity to members of the MEROPS database, available at ebi.ac.uk/merops. The number of peptidases was normalized within each species and shown as a heatmap. Thus, the peptidase families with the higher frequency have a value of 1.
