## Supplementary material for "Evolutionary analysis of genome-specific duplications in flatworm genomes": Distribution of secreted and transmembrane protein-coding genes within inparalogs and other homologs in studied phyla.

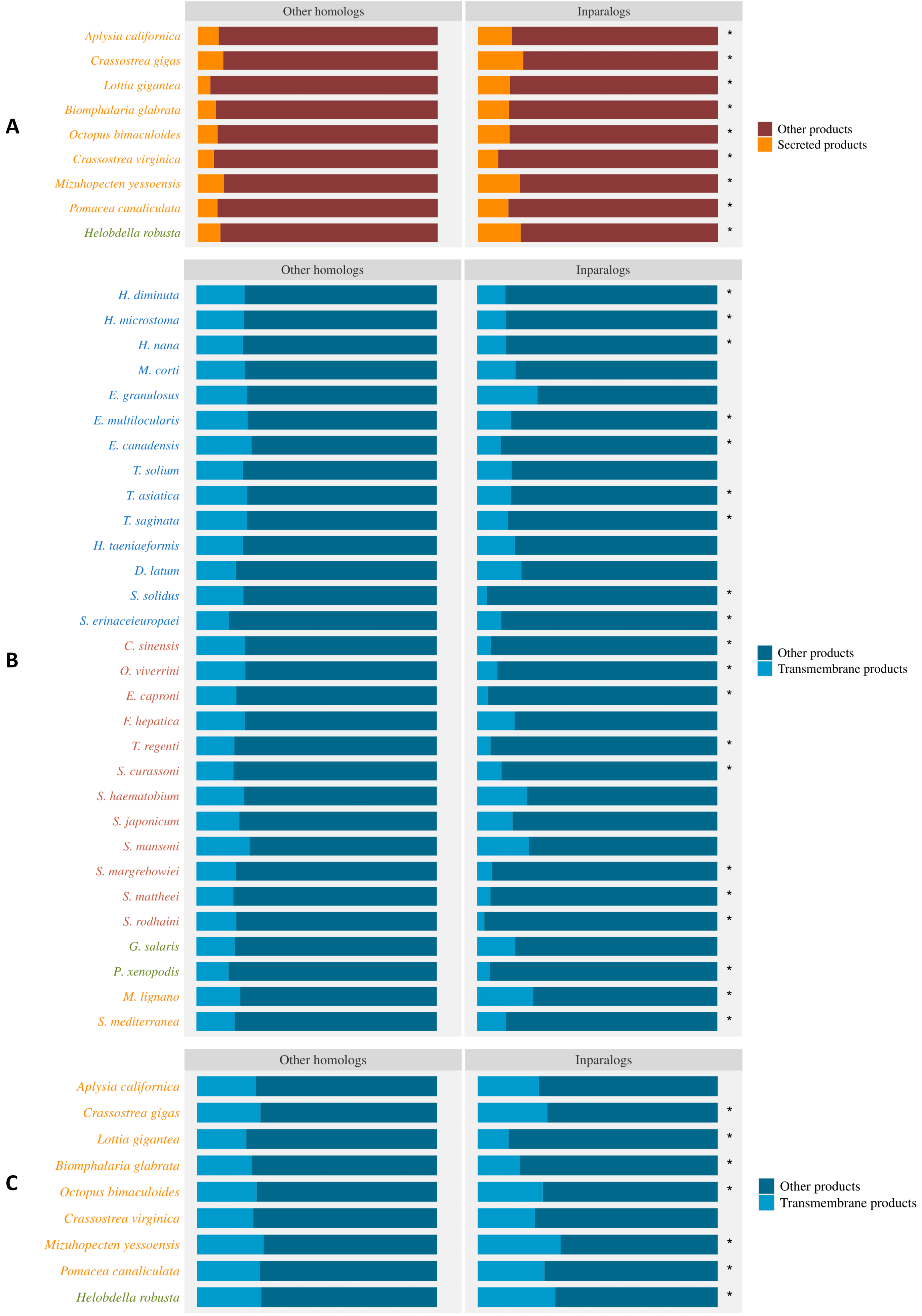


Supplementary Figure 4. Distribution of secreted and transmembrane protein-coding genes within inparalogs and other homologs in Mollusca and Annelida (A and C) and transmembrane protein-coding genes in Platyhelminthes (B). Significantly over- or under-representation of these functional categories among inparalogs is indicated with "*" (Fisher exact test, FDR < 1%). Only homologous genes (no singletons) were considered as background for this enrichment test.
