## Supplementary Figure 5. Plots of the expression level (measured as TMM, Trimmed Means of M-values) of inparalogous groups identified in S. mansoni. for "Evolutionary analysis of genome-specific duplications in flatworm genomes"

**Supplementary Figure 5**. Plots of the expression level (measured as TMM, Trimmed Means of M-values) of inparalogous groups identified in *S. mansoni* throughout the developmental stages studied in Protasio et al. 2012. Each dot corresponds to a replicate of each stage of each inparalog. The median TMM among all replicates is plotted for each inparalog. Expression data was taken from the NCBI SRA database as indicated in Supplementary Table 1.

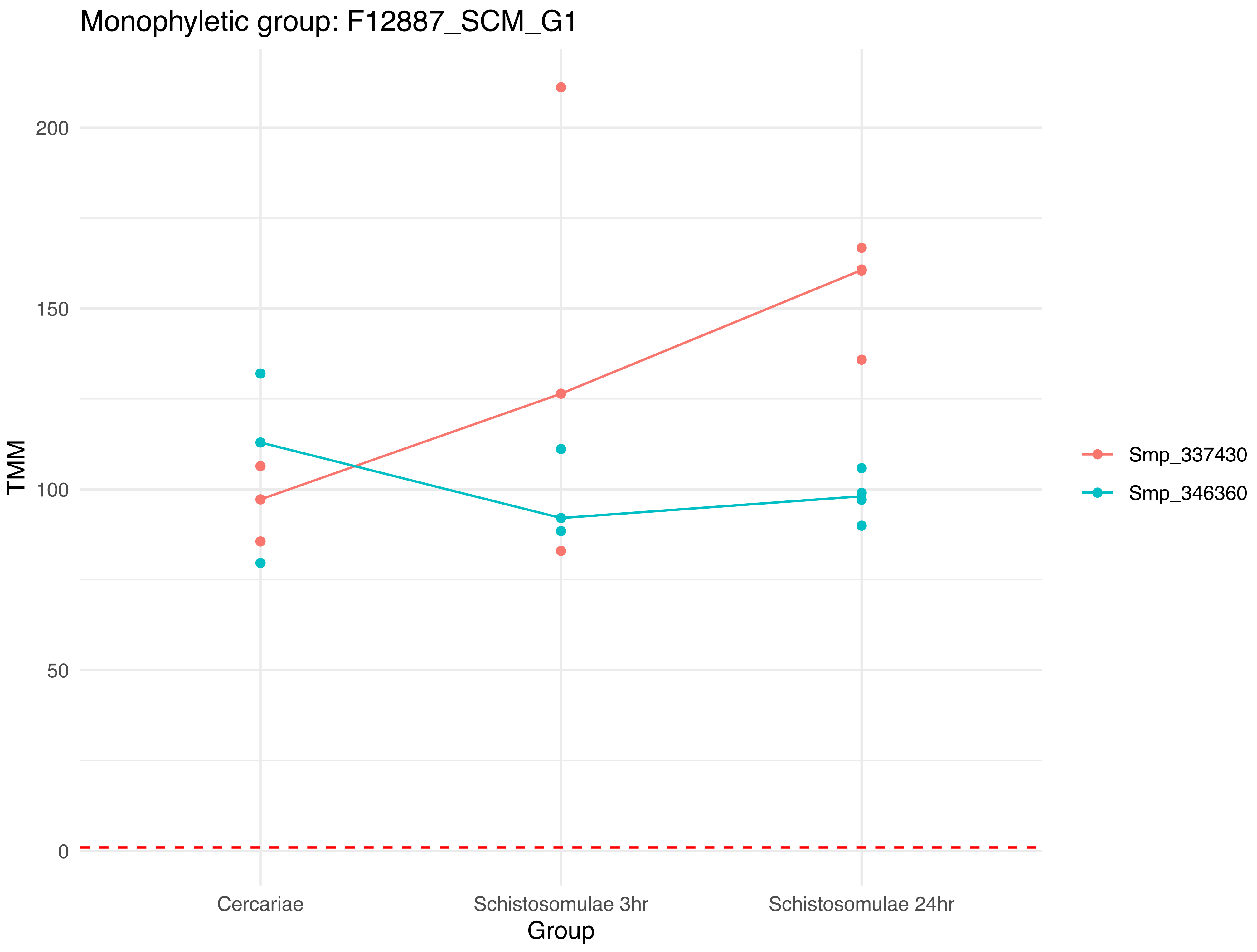

Monophyletic group: F12887_SCM_G1
TMM
200
150
100
50
0
Cercariae
Schistosomulae 3hr
Schistosomulae 24hr
Group
Smp_337430
Smp_346360

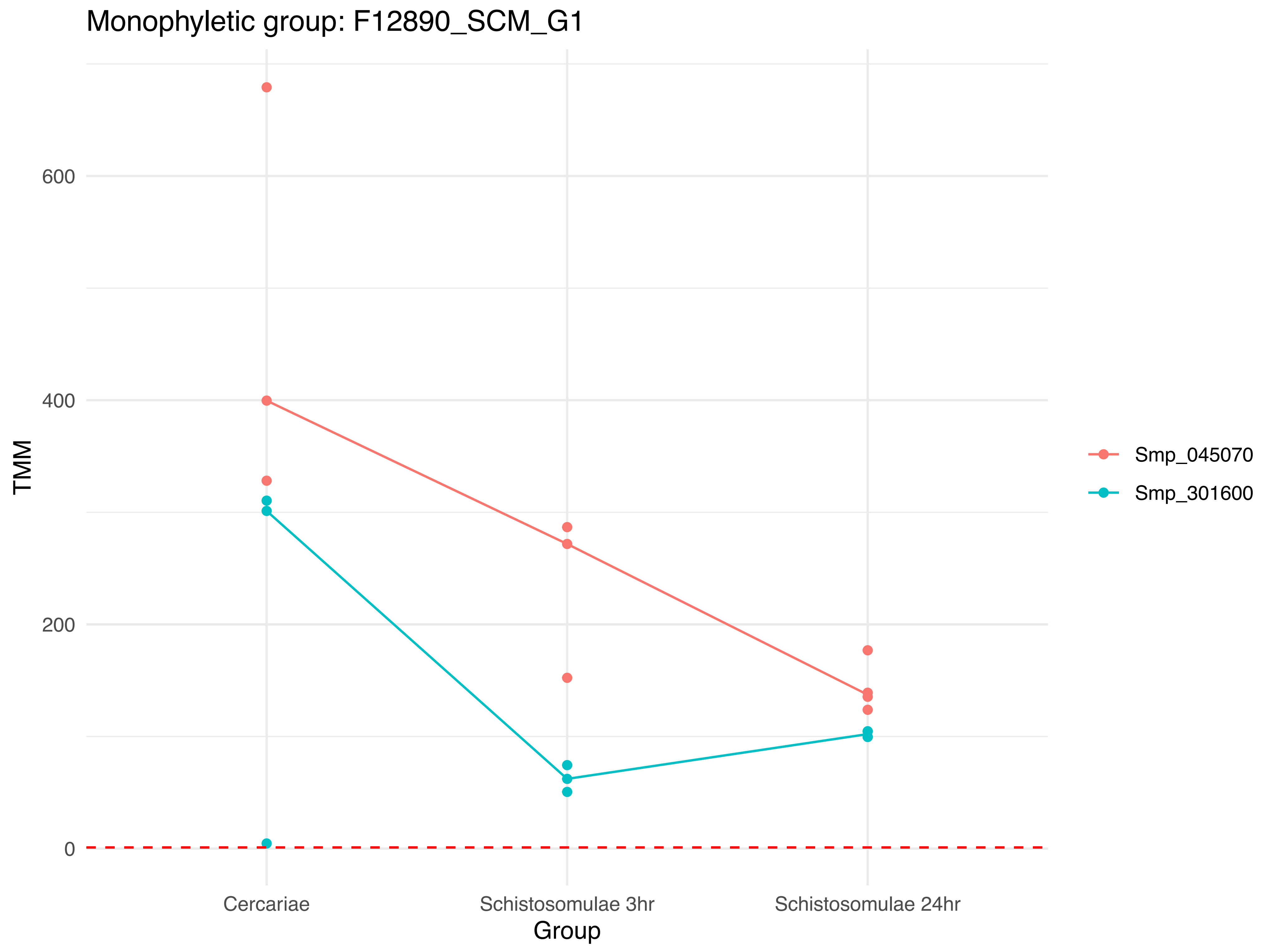

Monophyletic group: F12890_SCM_G1
600
400
200
0
TMM
Cercariae
Schistosomulae 3hr
Schistosomulae 24hr
Group
Smp_045070
Smp_301600

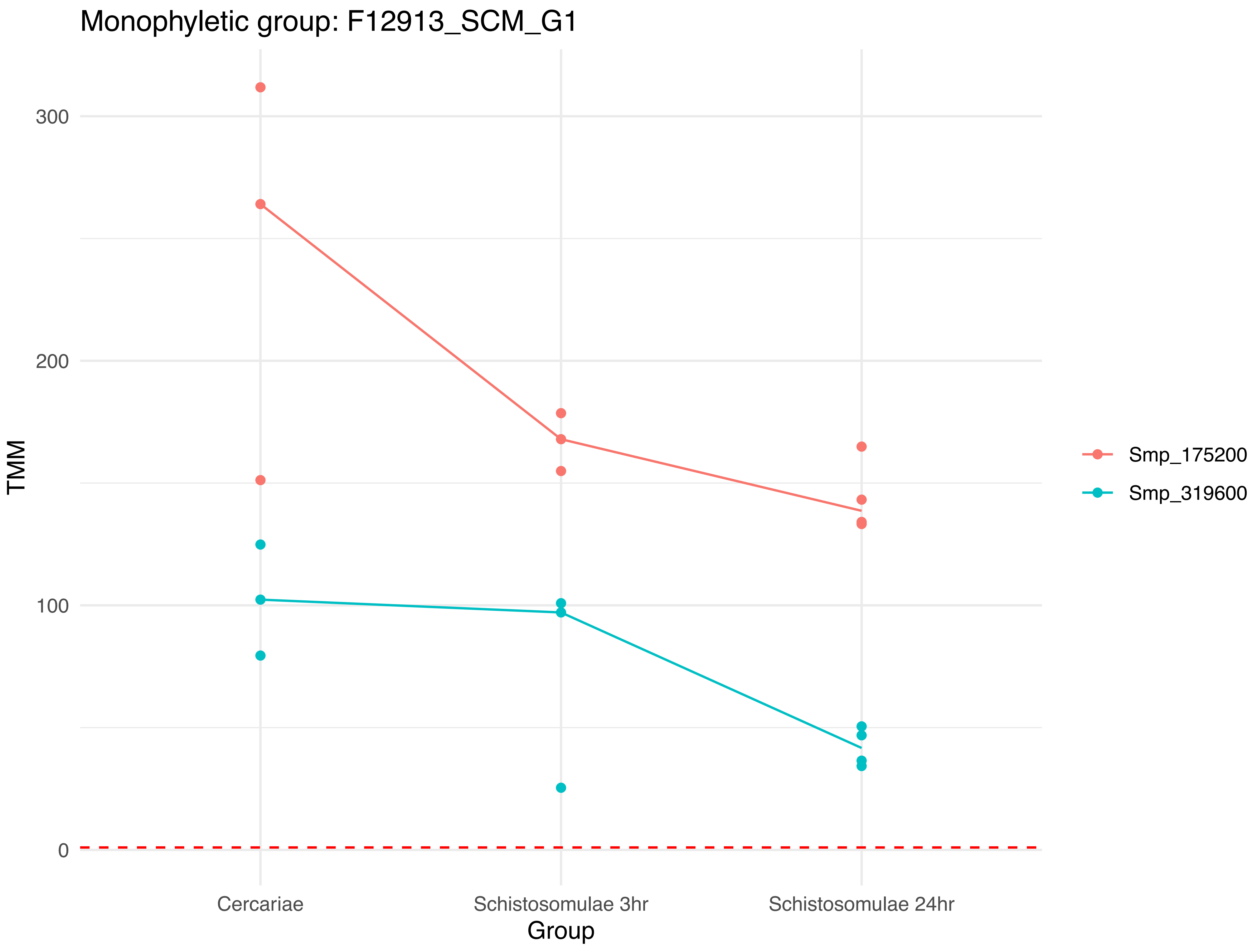
Monophyletic group: F12913_SCM_G1
300
200
100
0
TMM
Cercariae
Schistosomulae 3hr
Schistosomulae 24hr
Group
Smp_175200
Smp_319600

Monophyletic group: F13161_SCM_G1

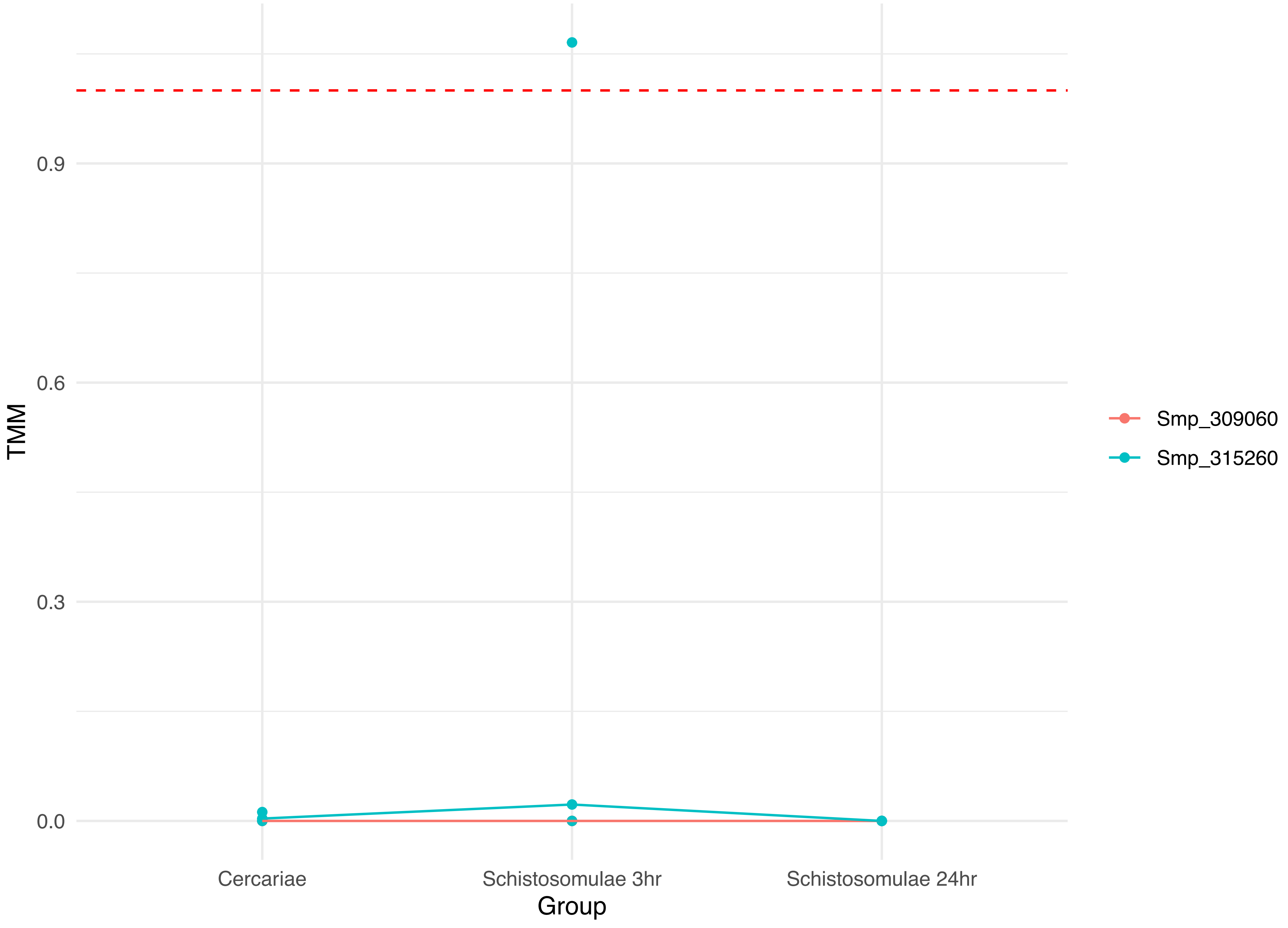

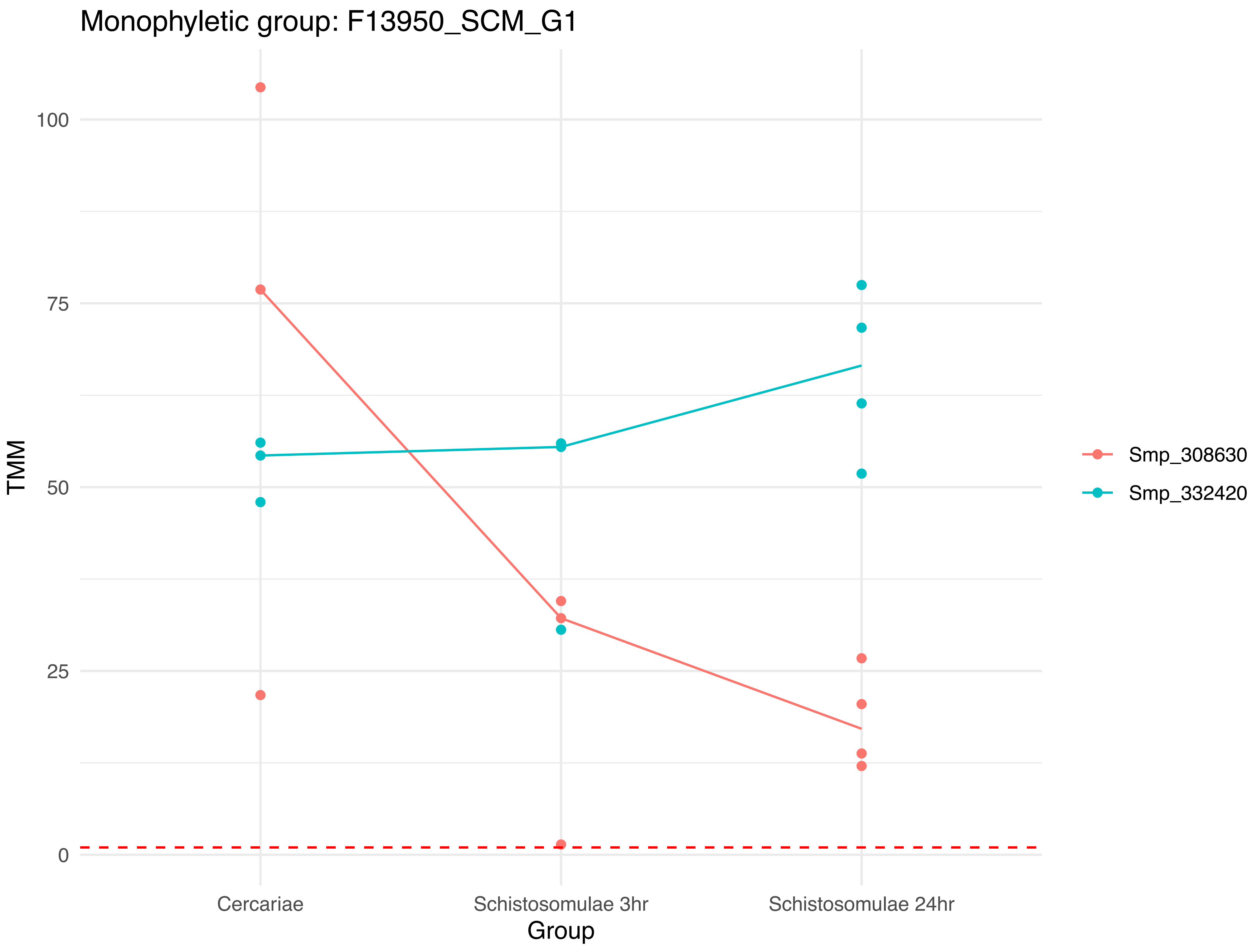
Monophyletic group: F13950_SCM_G1
TMM
100
75
50
25
0
Cercariae
Schistosomulae 3hr
Schistosomulae 24hr
Group
Smp_308630
Smp_332420

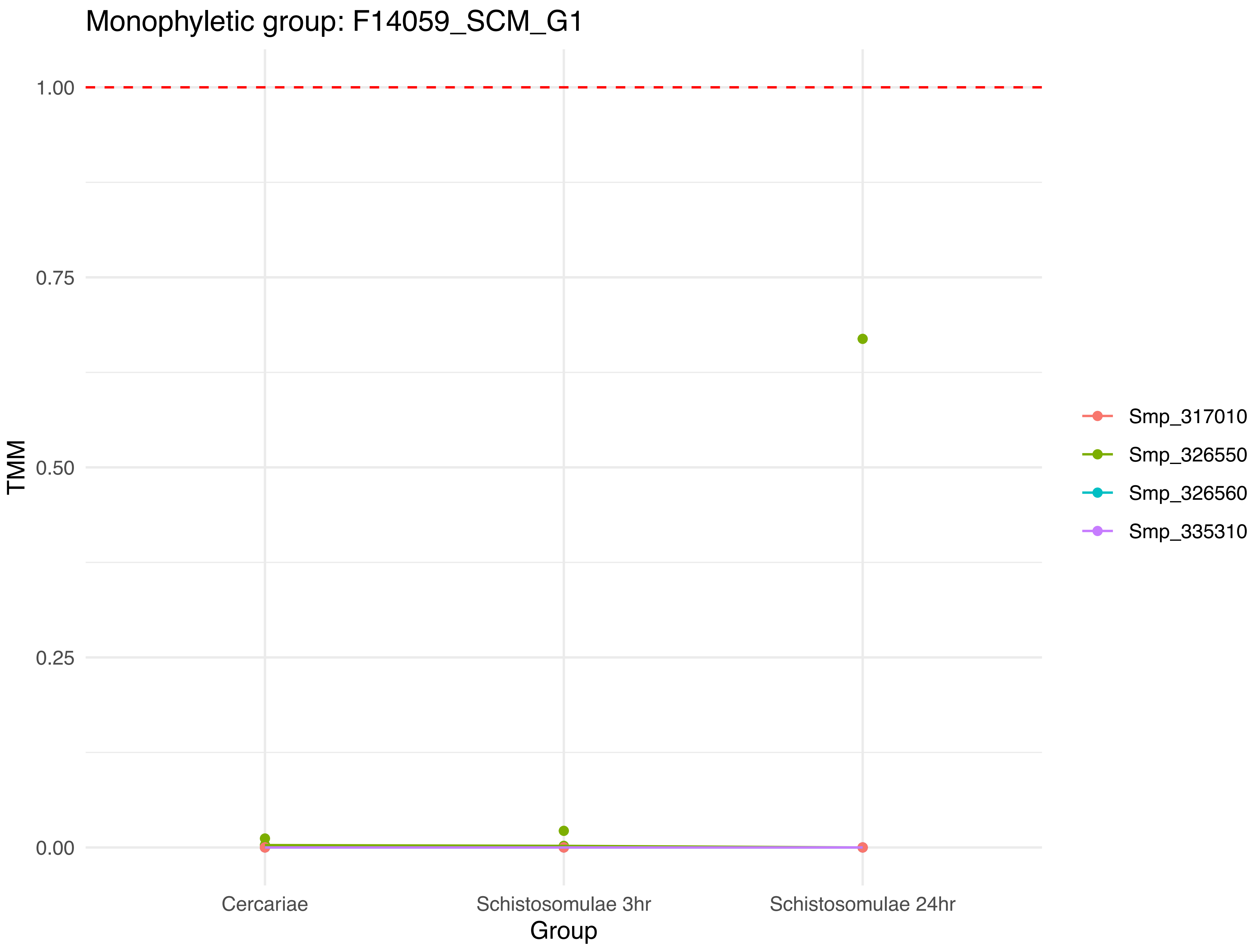
Monophyletic group: F14059_SCM_G1
1.00
0.75
0.50
0.25
0.00
TMM
Cercariae
Schistosomulae 3hr
Schistosomulae 24hr
Group
Smp_317010
Smp_326550
Smp_326560
Smp_335310

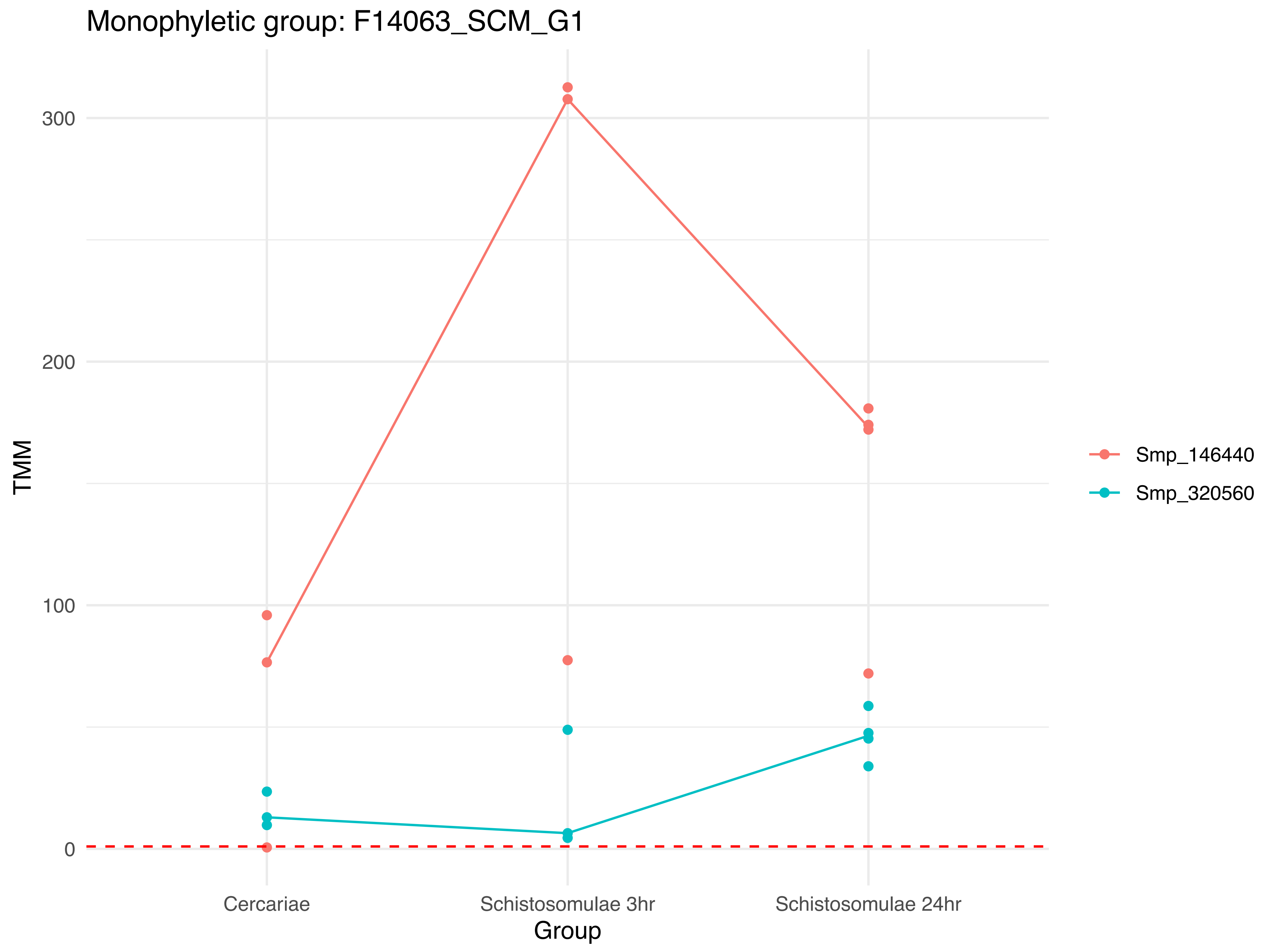

Monophyletic group: F14063_SCM_G1
300
200
100
0
TMM
Cercariae
Schistosomulae 3hr
Schistosomulae 24hr
Group
Smp_146440
Smp_320560

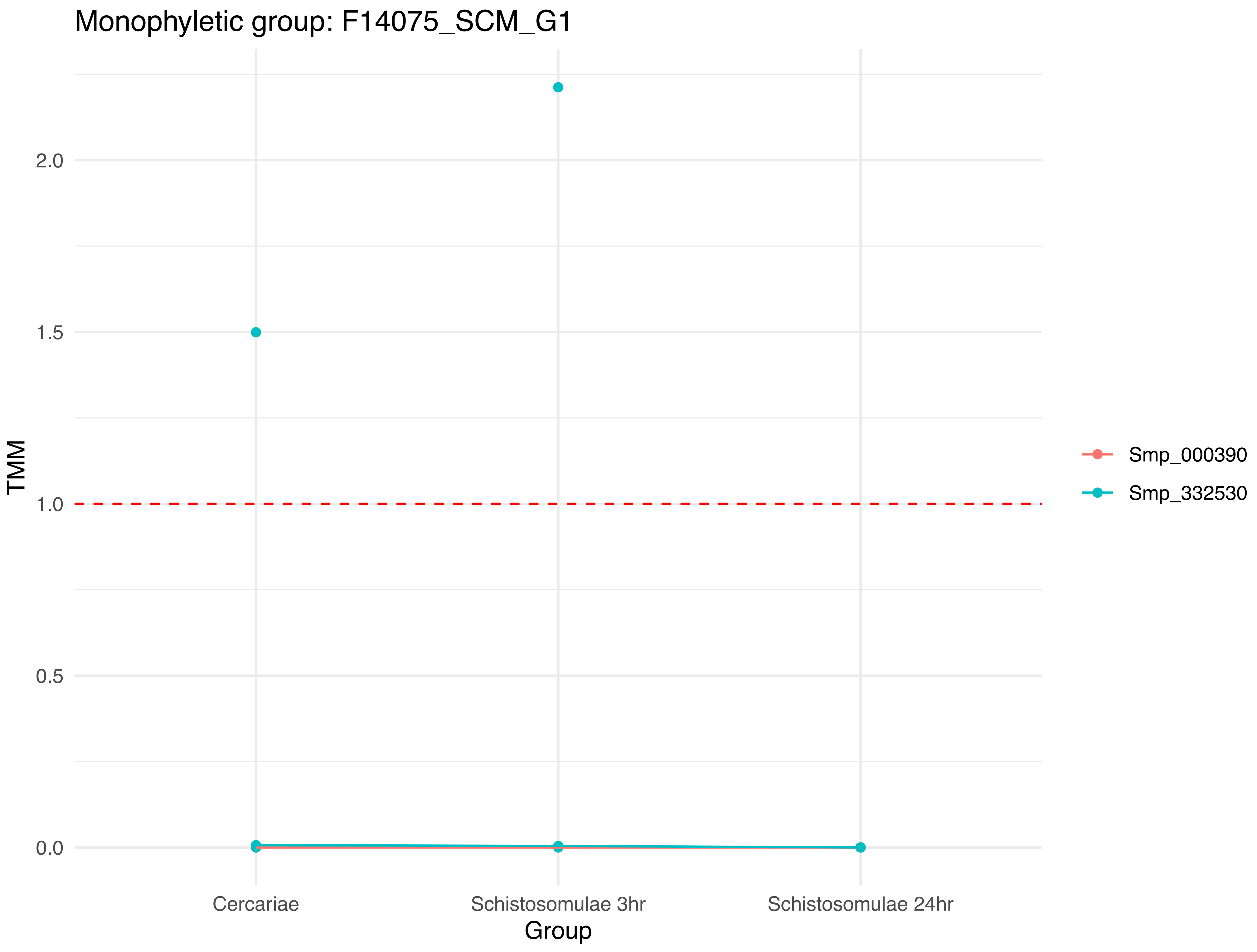
Monophyletic group: F14075_SCM_G1
TMM
2.0
1.5
1.0
0.5
0.0
Cercariae
Schistosomulae 3hr
Schistosomulae 24hr
Group
Smp_000390
Smp_332530

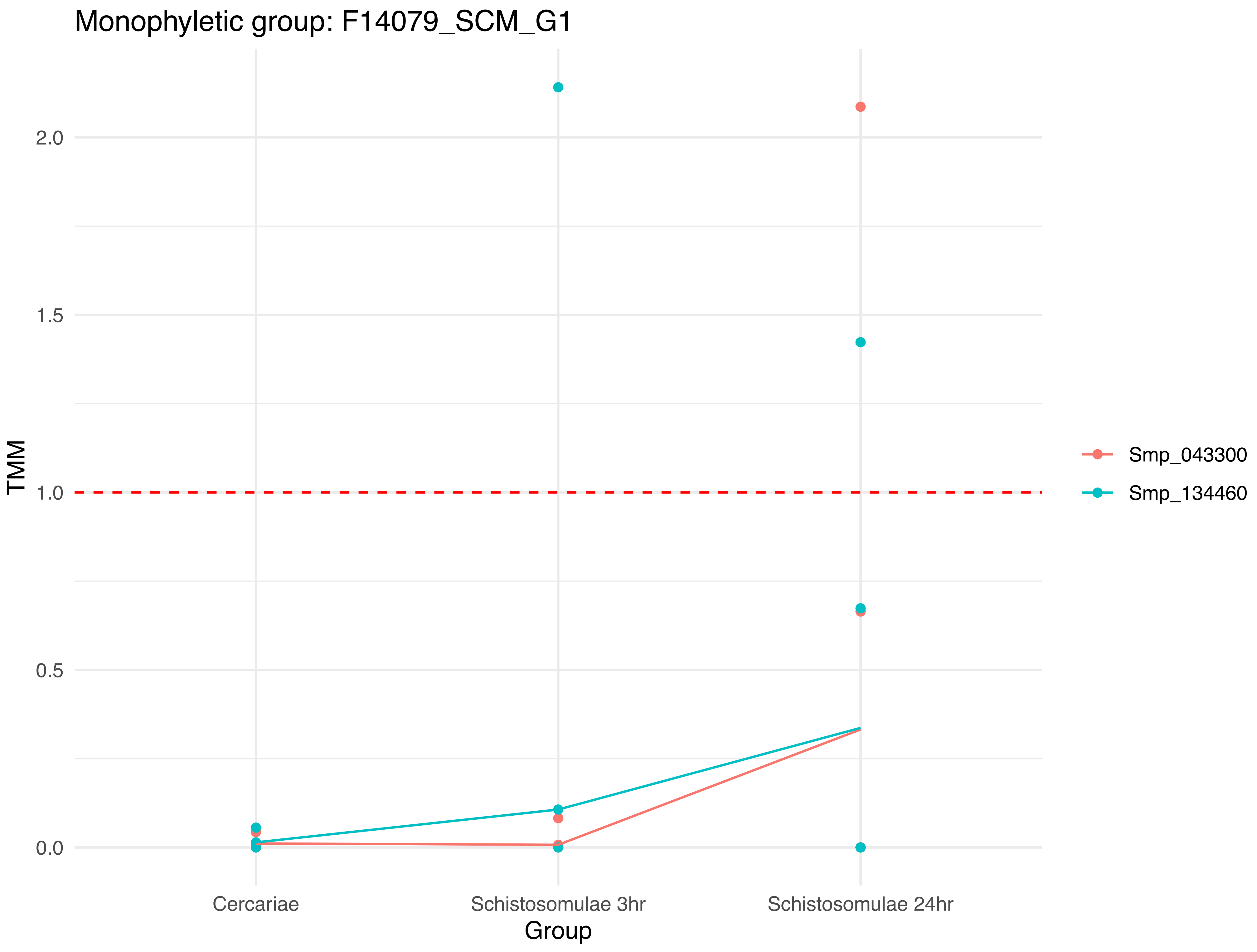
Monophyletic group: F14079_SCM_G1
TMM
2.0
1.5
1.0
0.5
0.0
Cercariae
Schistosomulae 3hr
Schistosomulae 24hr
Group
Smp_043300
Smp_134460

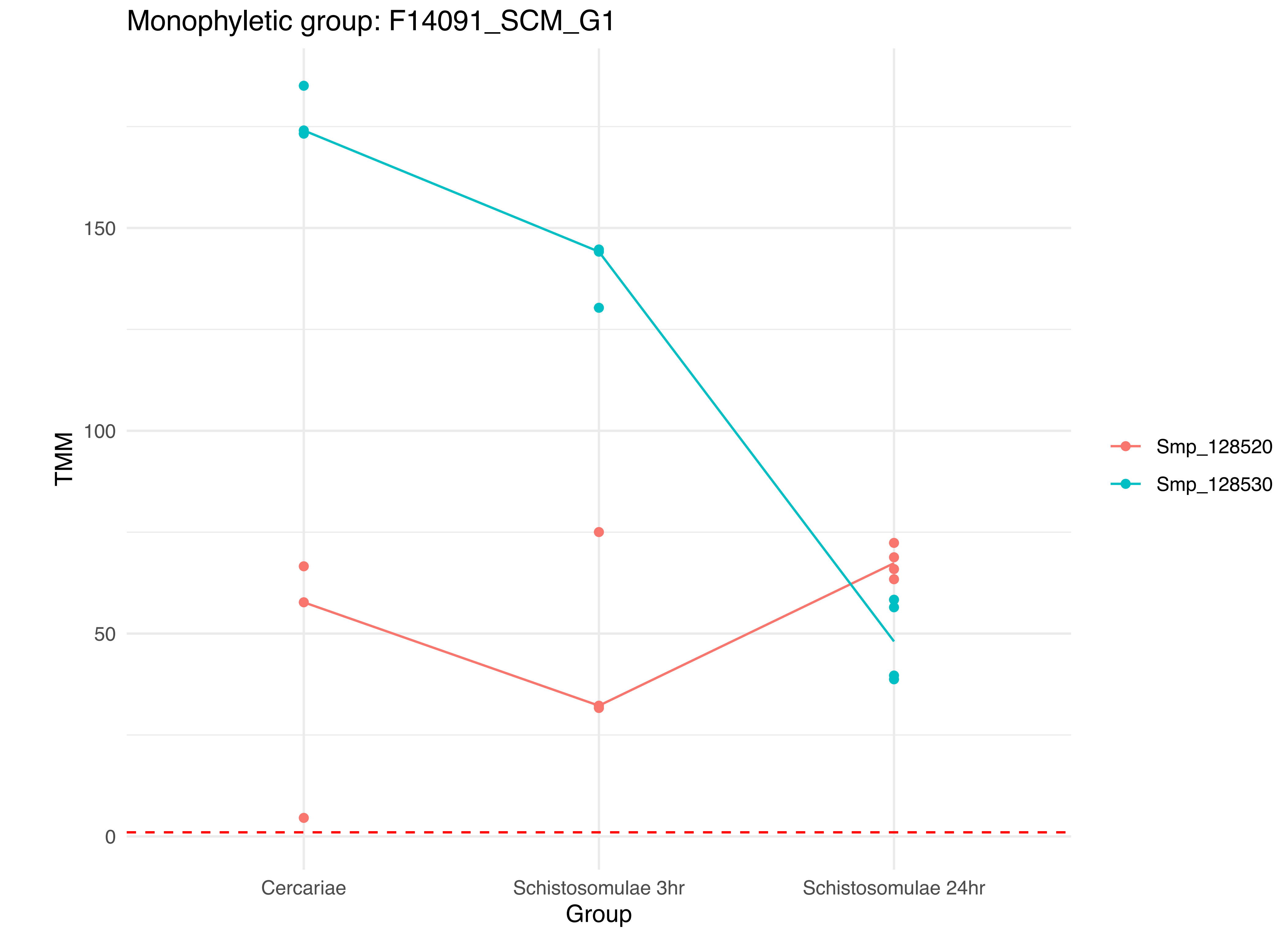

Monophyletic group: F14091_SCM_G1
TMM
150
100
50
0
Cercariae
Schistosomulae 3hr
Schistosomulae 24hr
Group
Smp_128520
Smp_128530

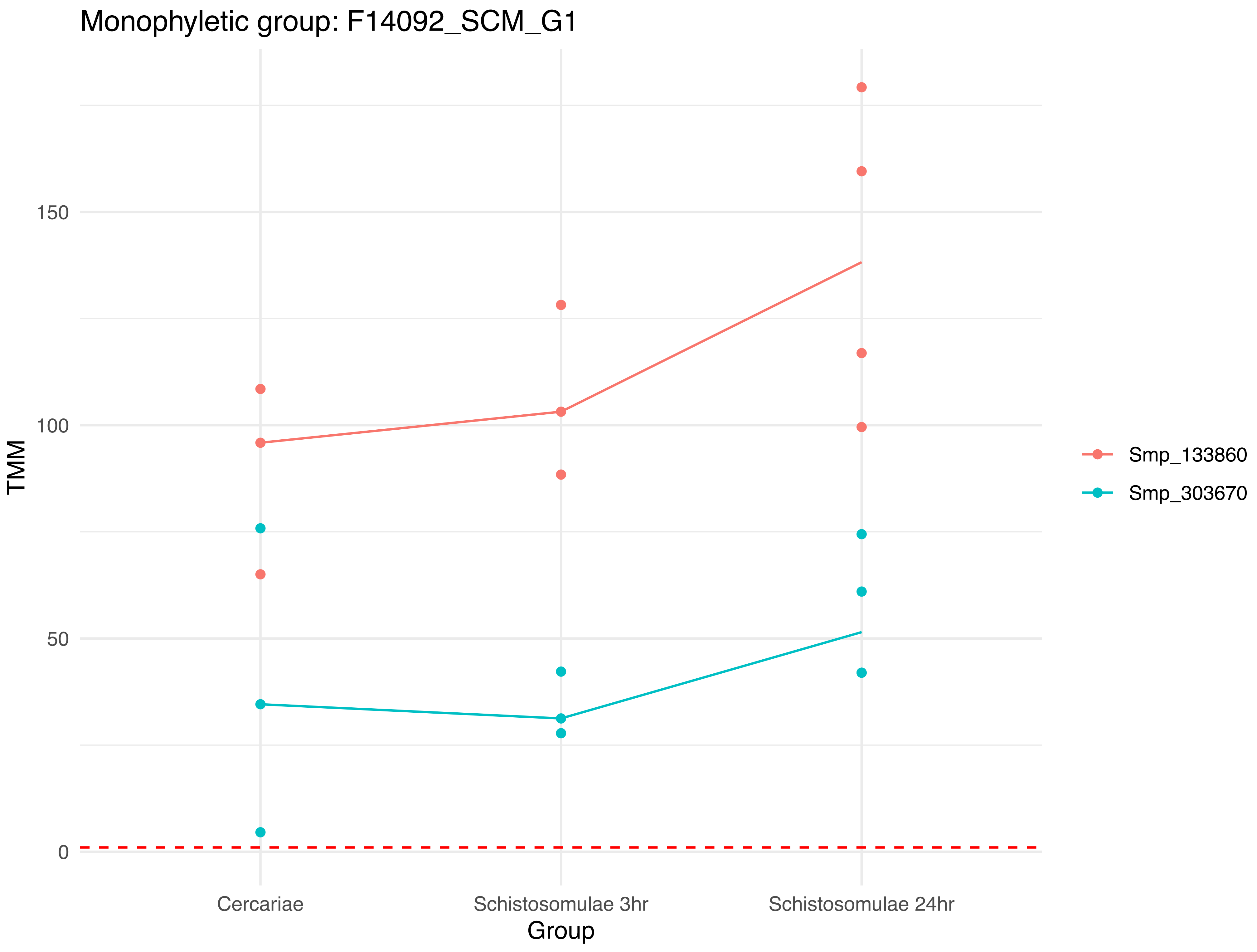

Monophyletic group: F14092_SCM_G1
150
100
50
0
TMM
Cercariae
Schistosomulae 3hr
Schistosomulae 24hr
Group
Smp_133860
Smp_303670

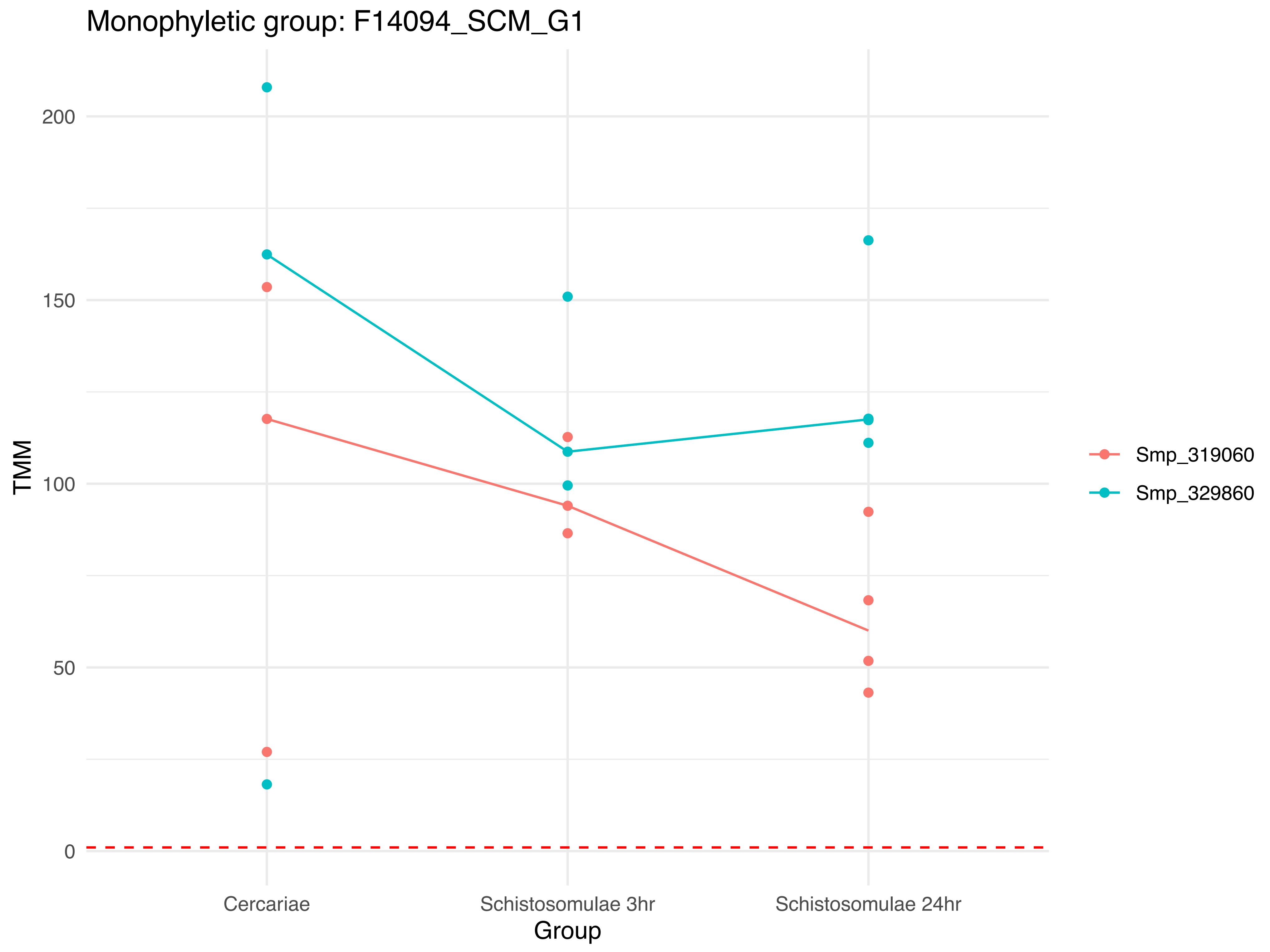

Monophyletic group: F14094_SCM_G1
TMM
200
150
100
50
0
Cercariae
Schistosomulae 3hr
Schistosomulae 24hr
Group
Smp_319060
Smp_329860

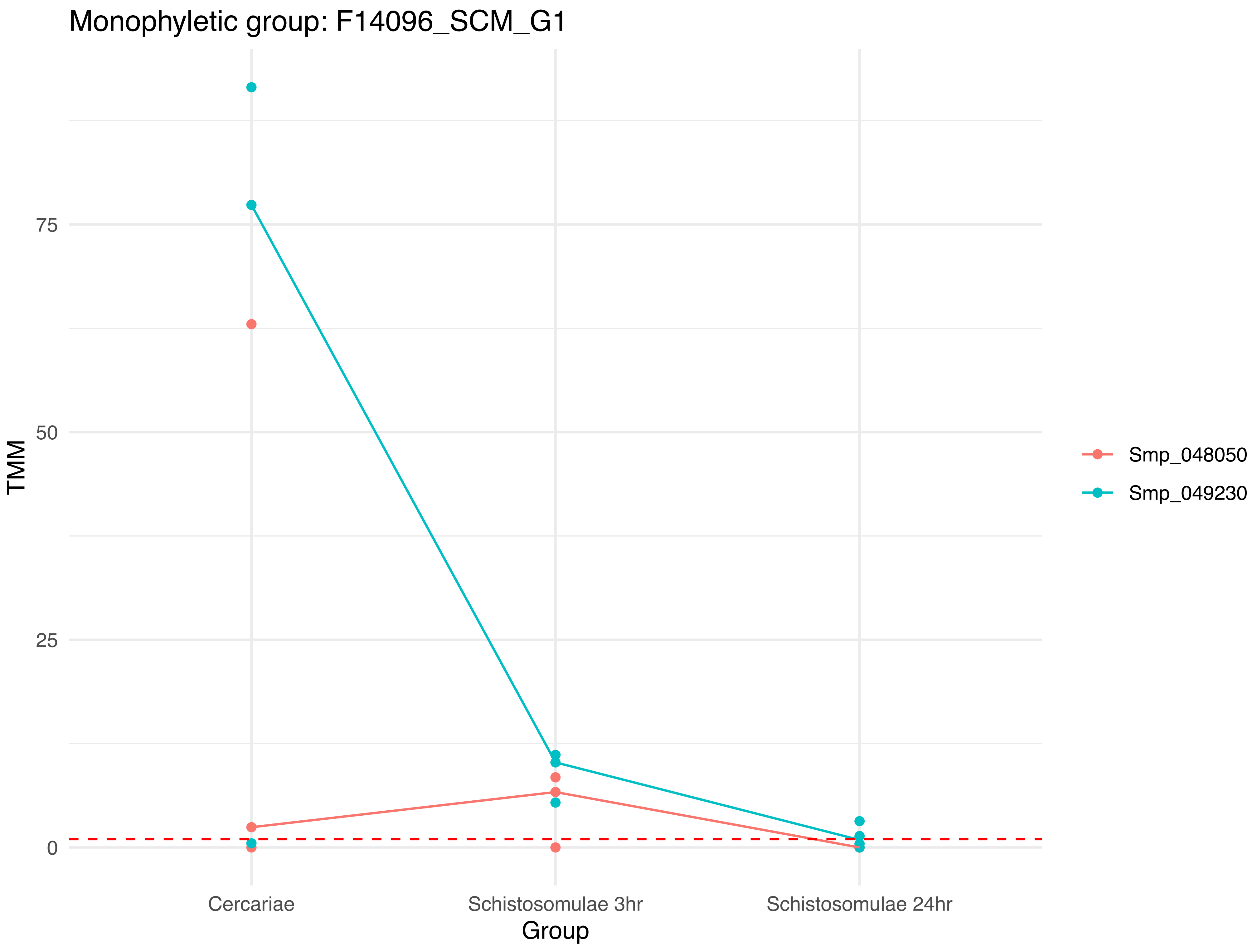
Monophyletic group: F14096_SCM_G1
TMM
75
50
25
0
Cercariae
Schistosomulae 3hr
Schistosomulae 24hr
Group
Smp_048050
Smp_049230

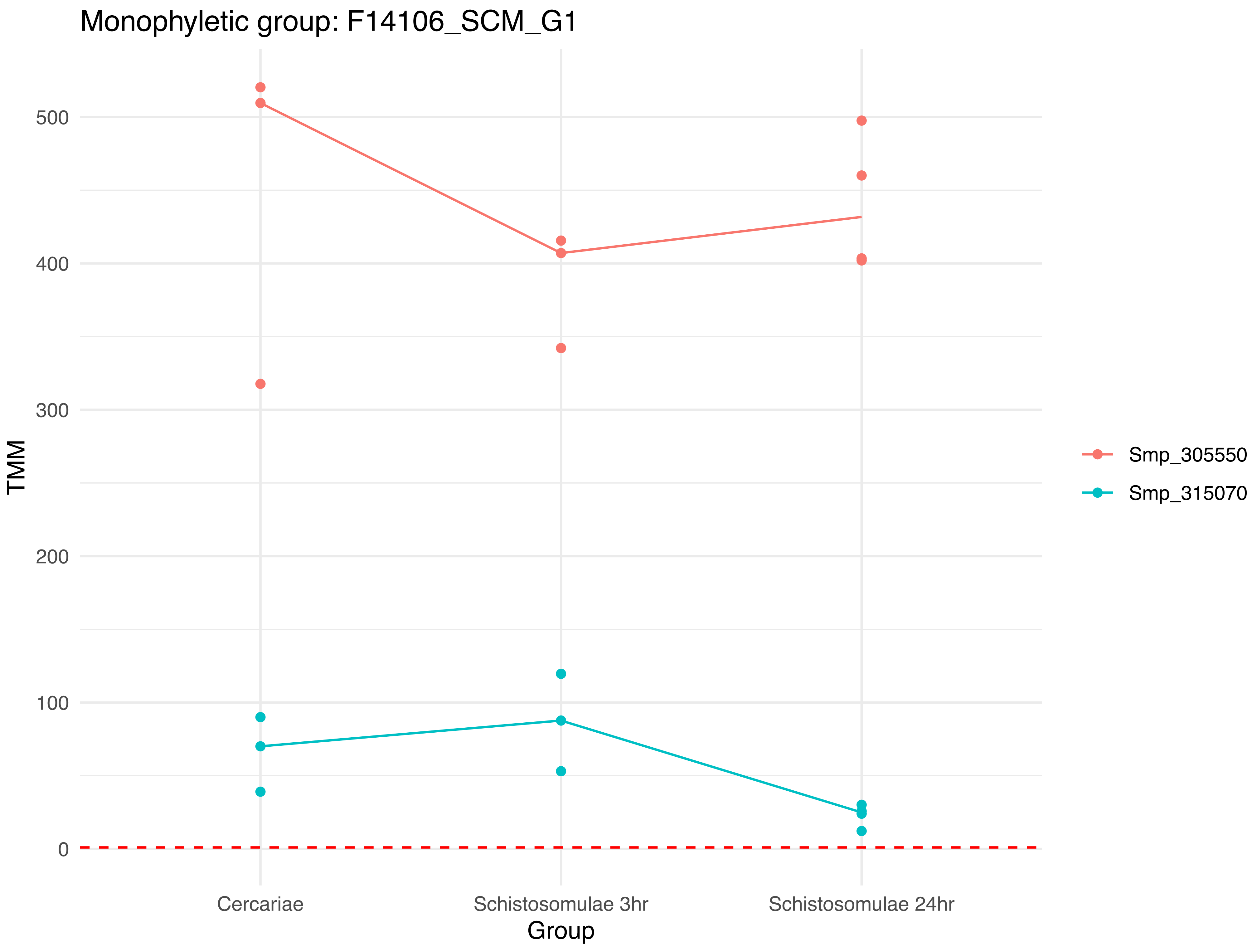
Monophyletic group: F14106_SCM_G1
TMM
500
400
300
200
100
0
Cercariae
Schistosomulae 3hr
Schistosomulae 24hr
Group
Smp_305550
Smp_315070

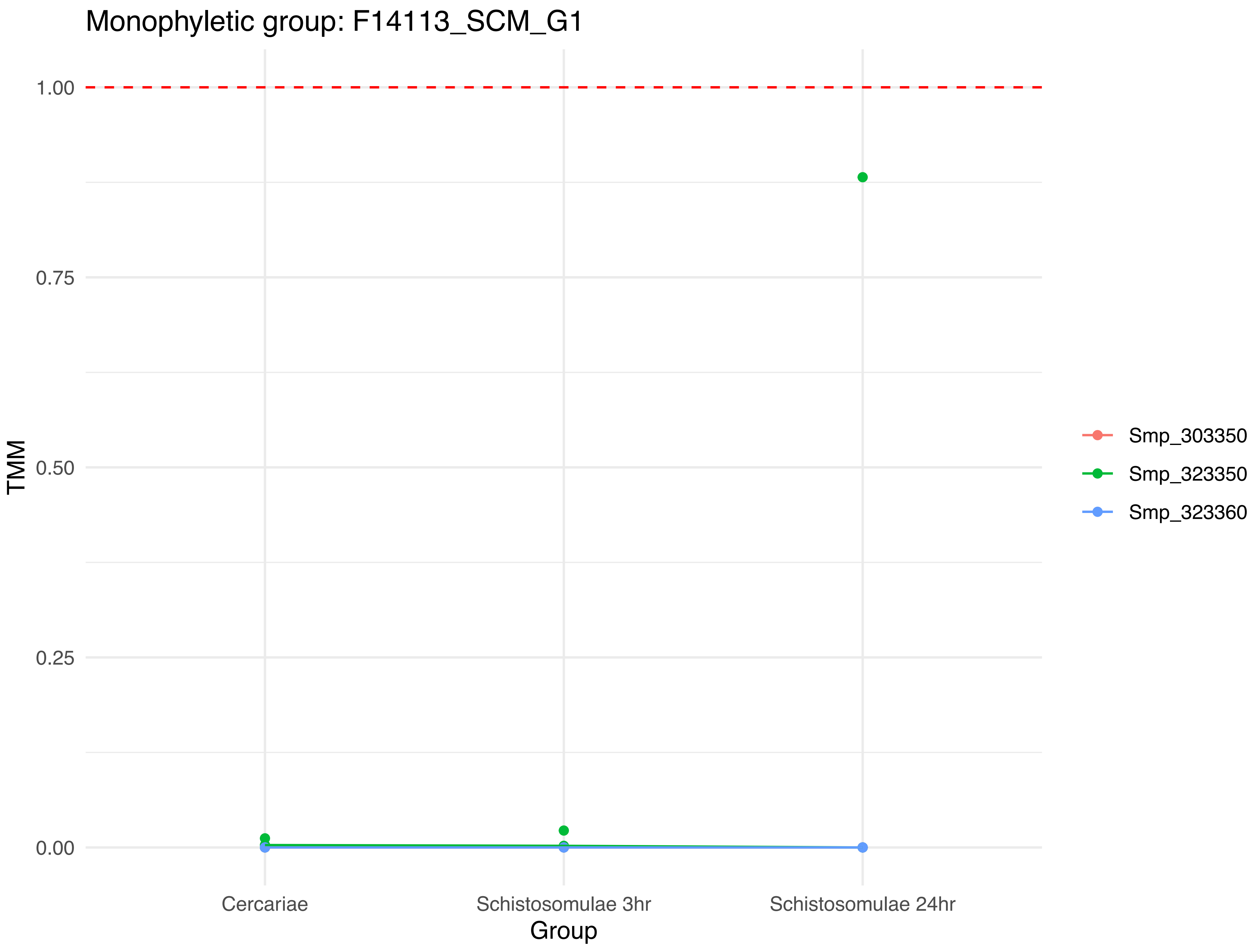
Monophyletic group: F14113_SCM_G1
1.00
0.75
0.50
0.25
0.00
TMM
Cercariae
Schistosomulae 3hr
Schistosomulae 24hr
Group
Smp_303350
Smp_323350
Smp_323360

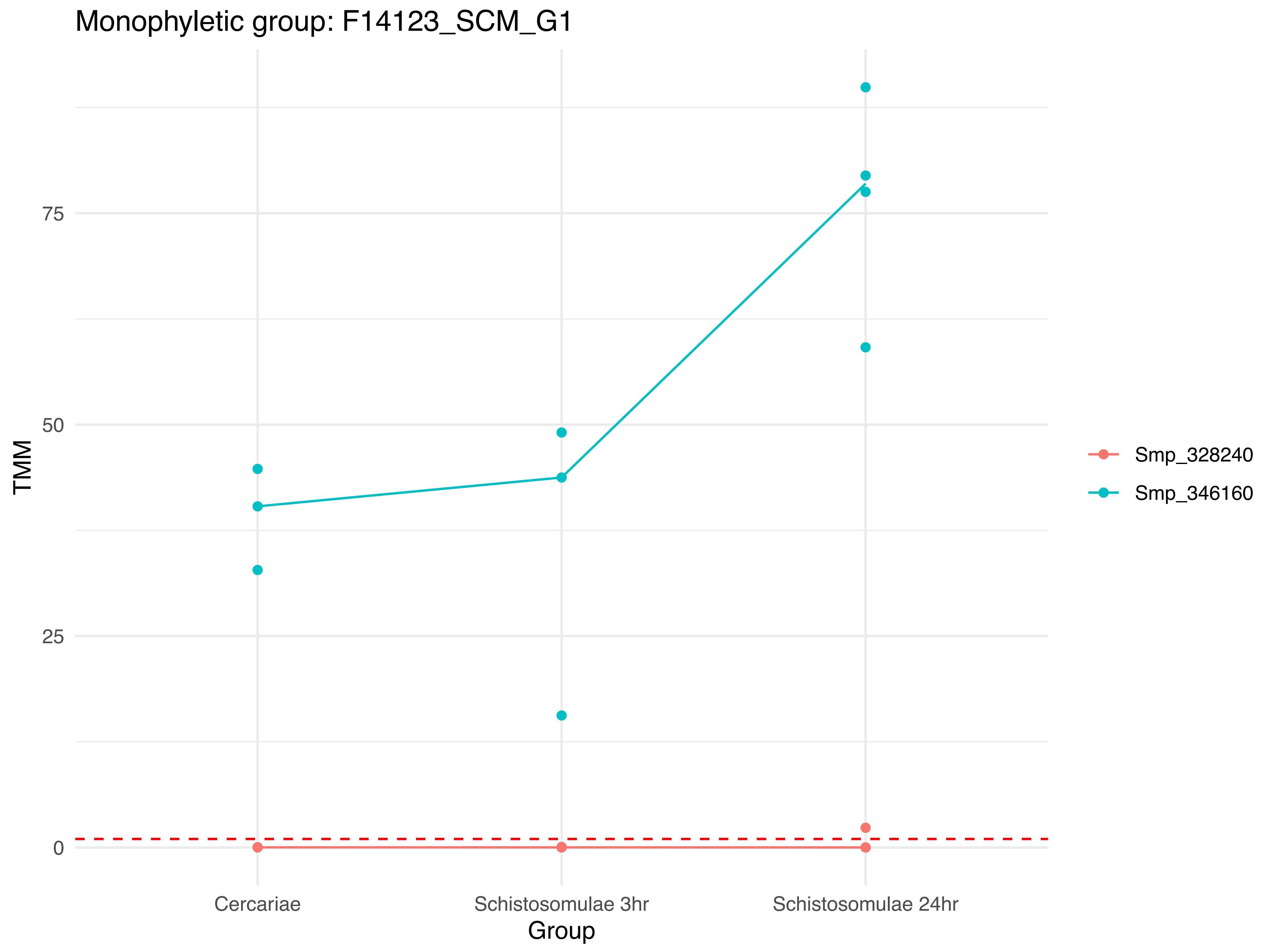

Monophyletic group: F14123_SCM_G1
TMM
75
50
25
0
Cercariae
Schistosomulae 3hr
Schistosomulae 24hr
Group
Smp_328240
Smp_346160

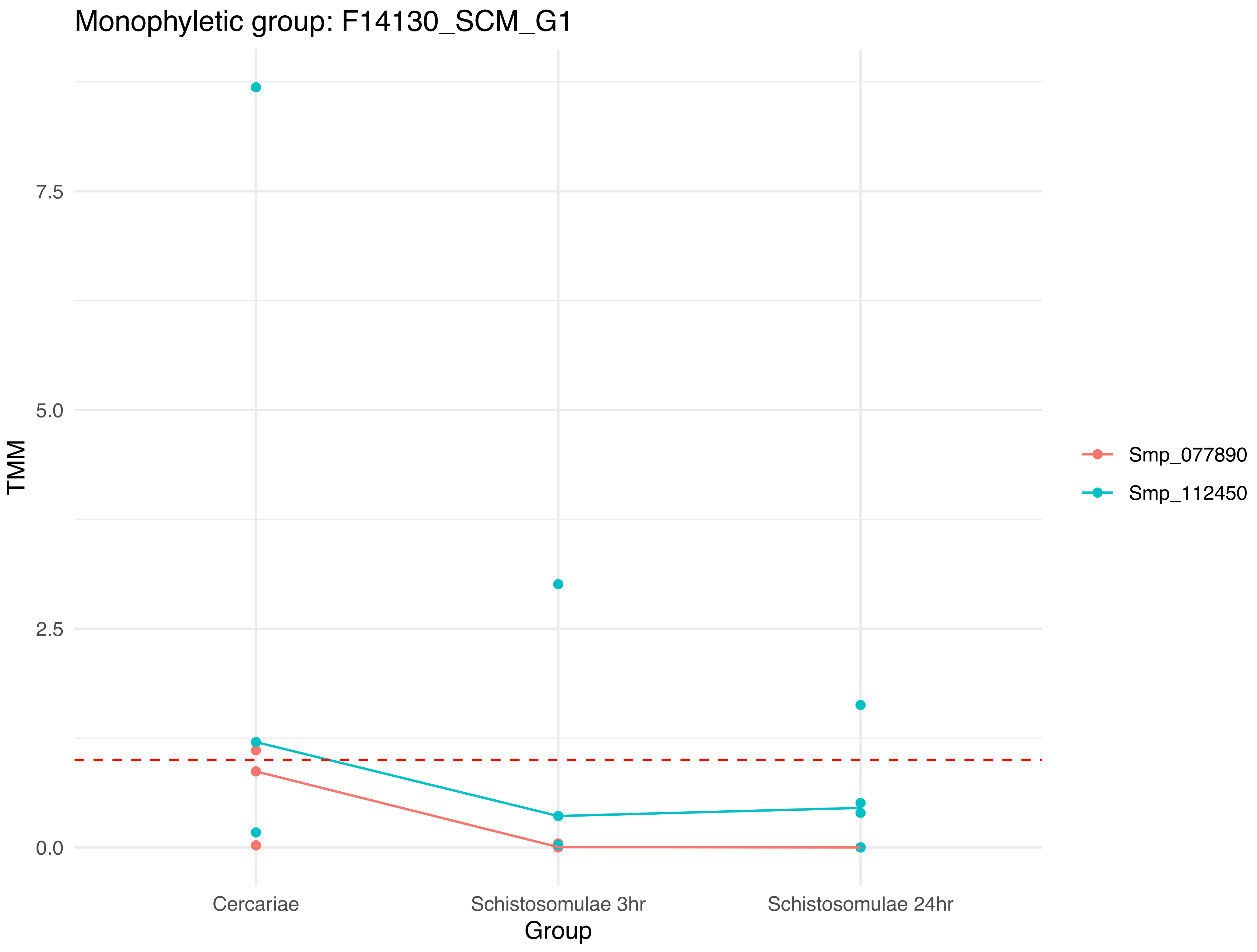
Monophyletic group: F14130_SCM_G1
TMM
7.5
5.0
2.5
0.0
Cercariae
Schistosomulae 3hr
Schistosomulae 24hr
Group
Smp_077890
Smp_112450

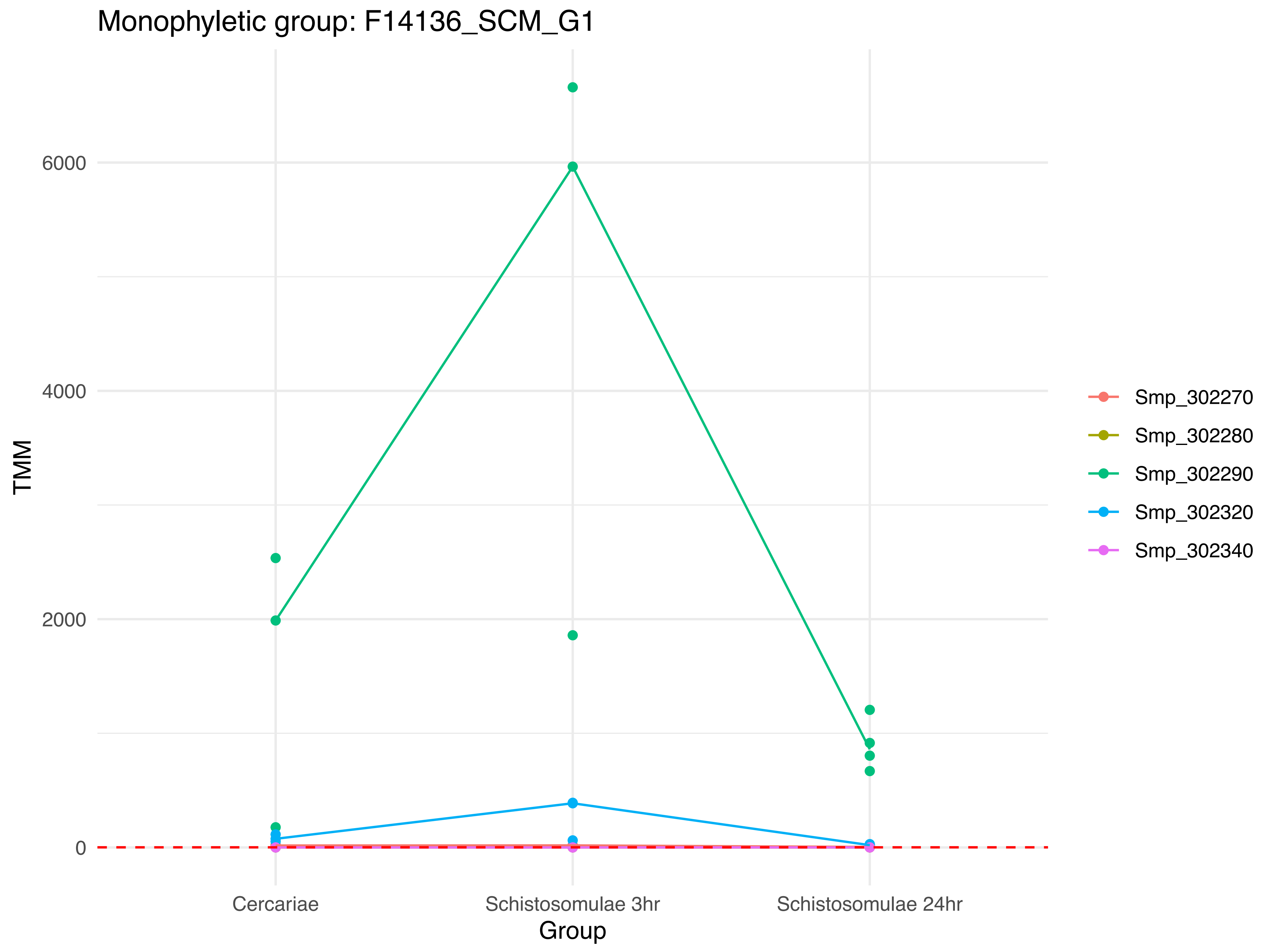
Monophyletic group: F14136_SCM_G1
TMM
6000
4000
2000
0
Cercariae
Schistosomulae 3hr
Schistosomulae 24hr
Group
Smp_302270
Smp_302280
Smp_302290
Smp_302320
Smp_302340

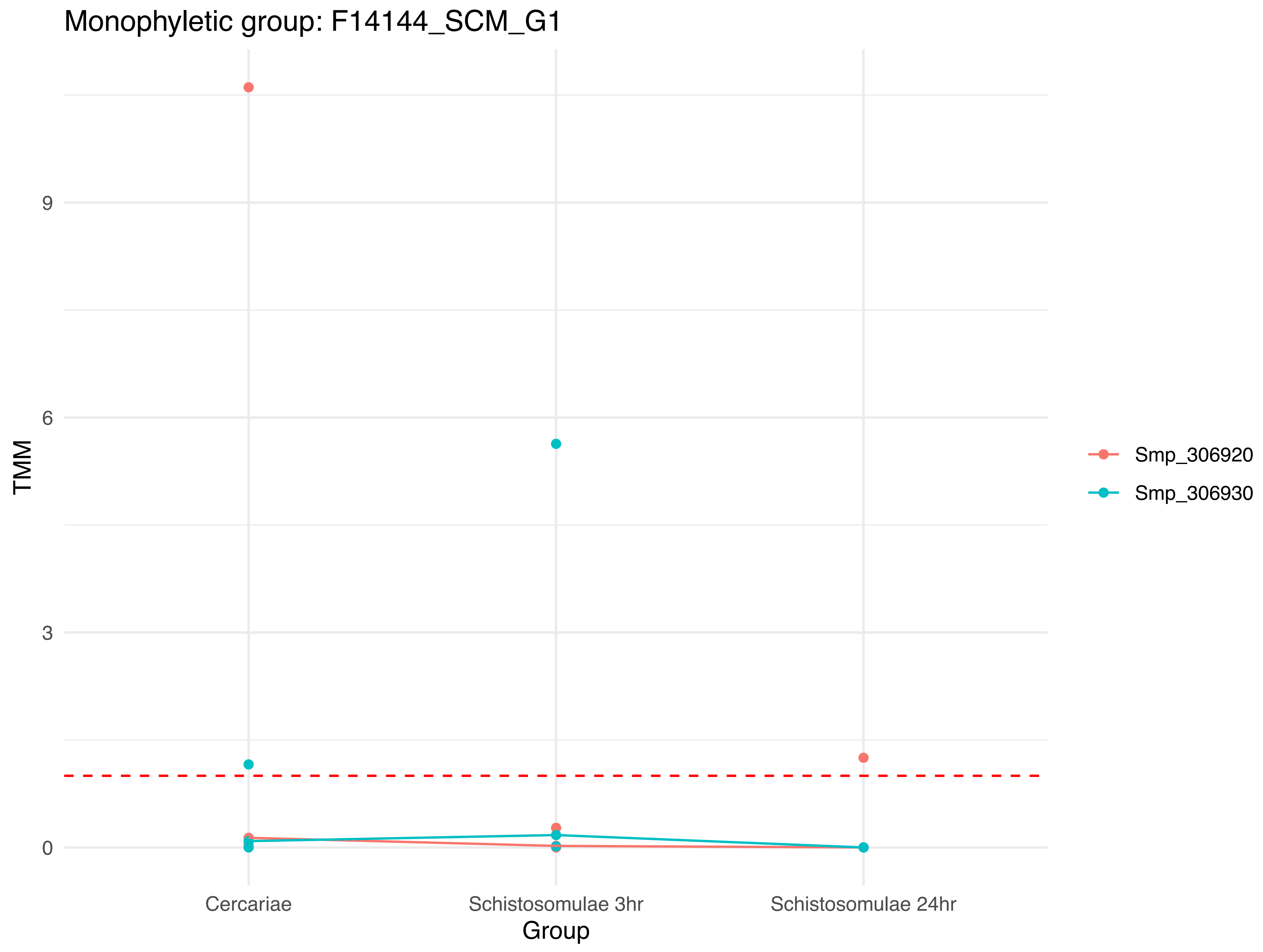

Monophyletic group: F14144_SCM_G1
TMM
9
6
3
0
Cercariae
Schistosomulae 3hr
Schistosomulae 24hr
Group
Smp_306920
Smp_306930

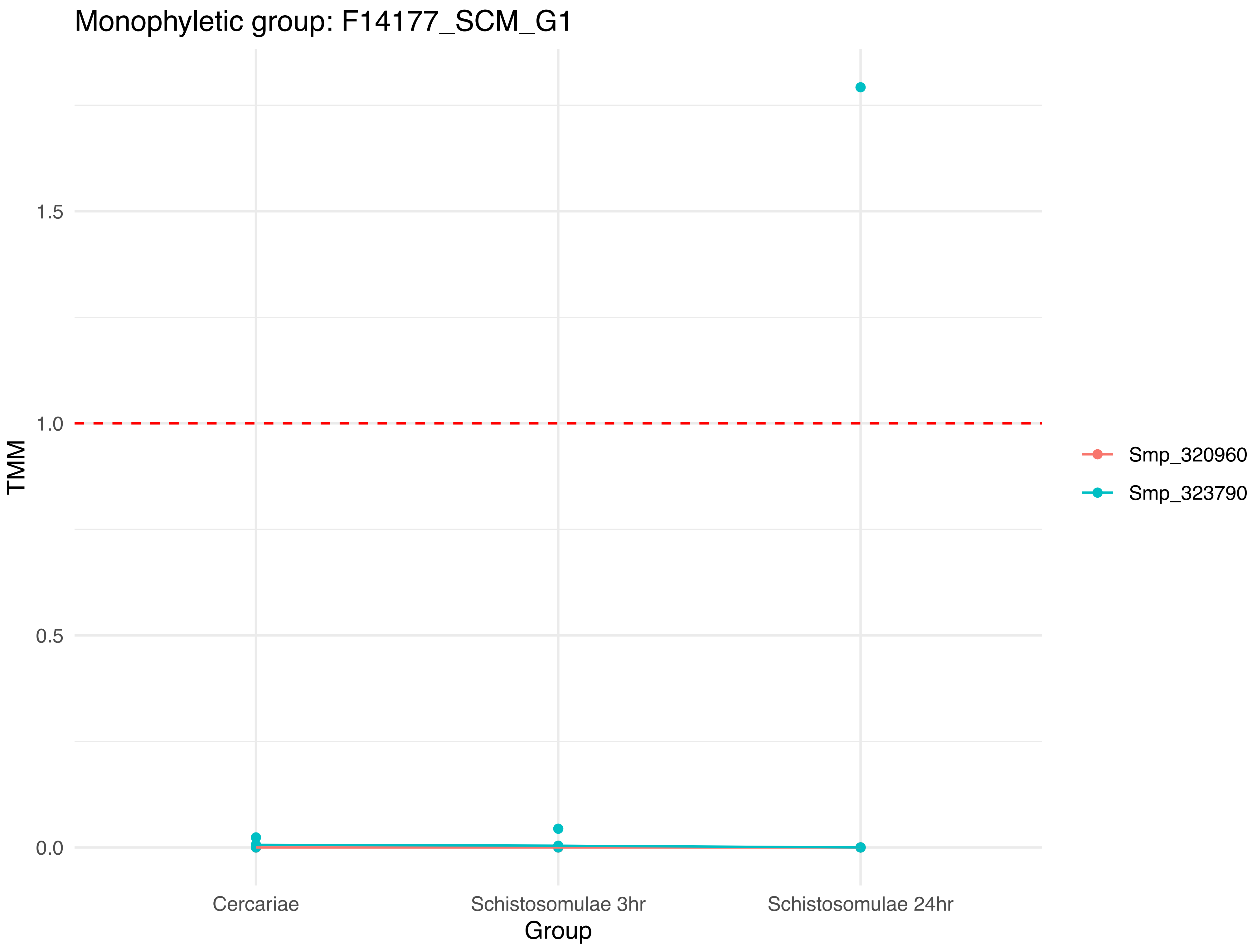

Monophyletic group: F14177_SCM_G1
TMM
1.5
1.0
0.5
0.0
Cercariae
Schistosomulae 3hr
Schistosomulae 24hr
Group
Smp_320960
Smp_323790

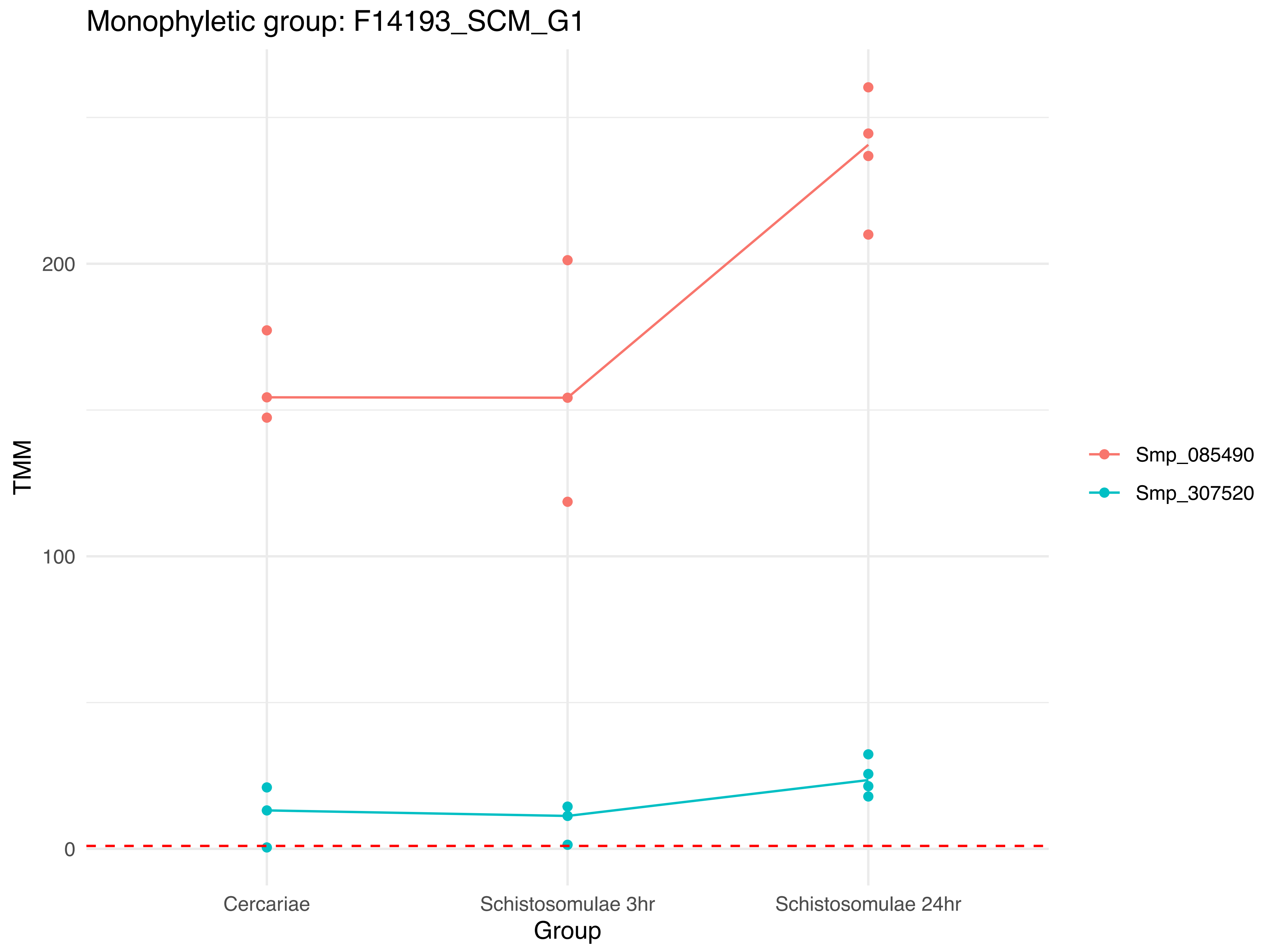
Monophyletic group: F14193_SCM_G1
TMM
200
100
0
Cercariae
Schistosomulae 3hr
Schistosomulae 24hr
Group
Smp_085490
Smp_307520

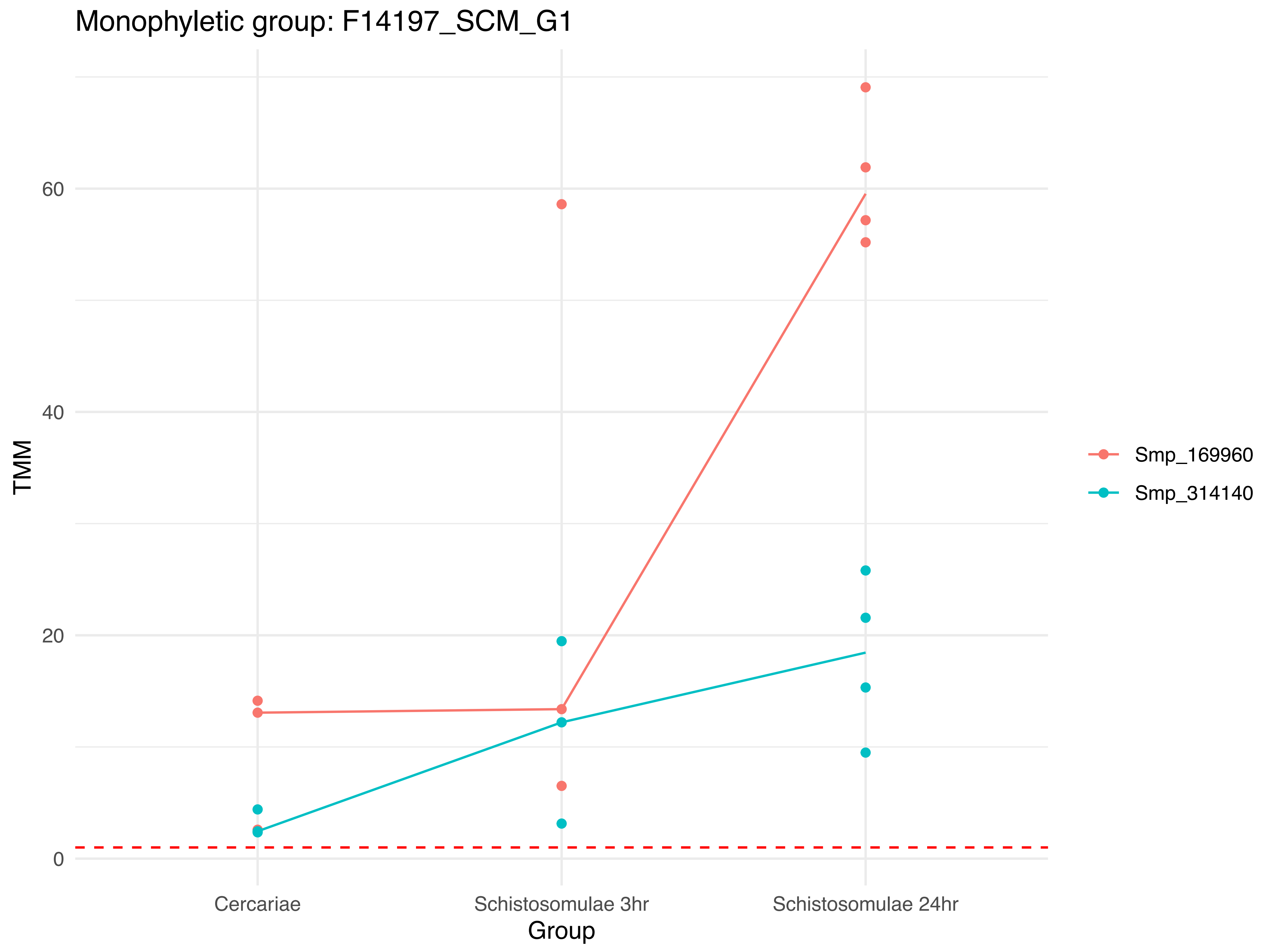
Monophyletic group: F14197_SCM_G1
TMM
60
40
20
0
Cercariae
Schistosomulae 3hr
Schistosomulae 24hr
Group
Smp_169960
Smp_314140

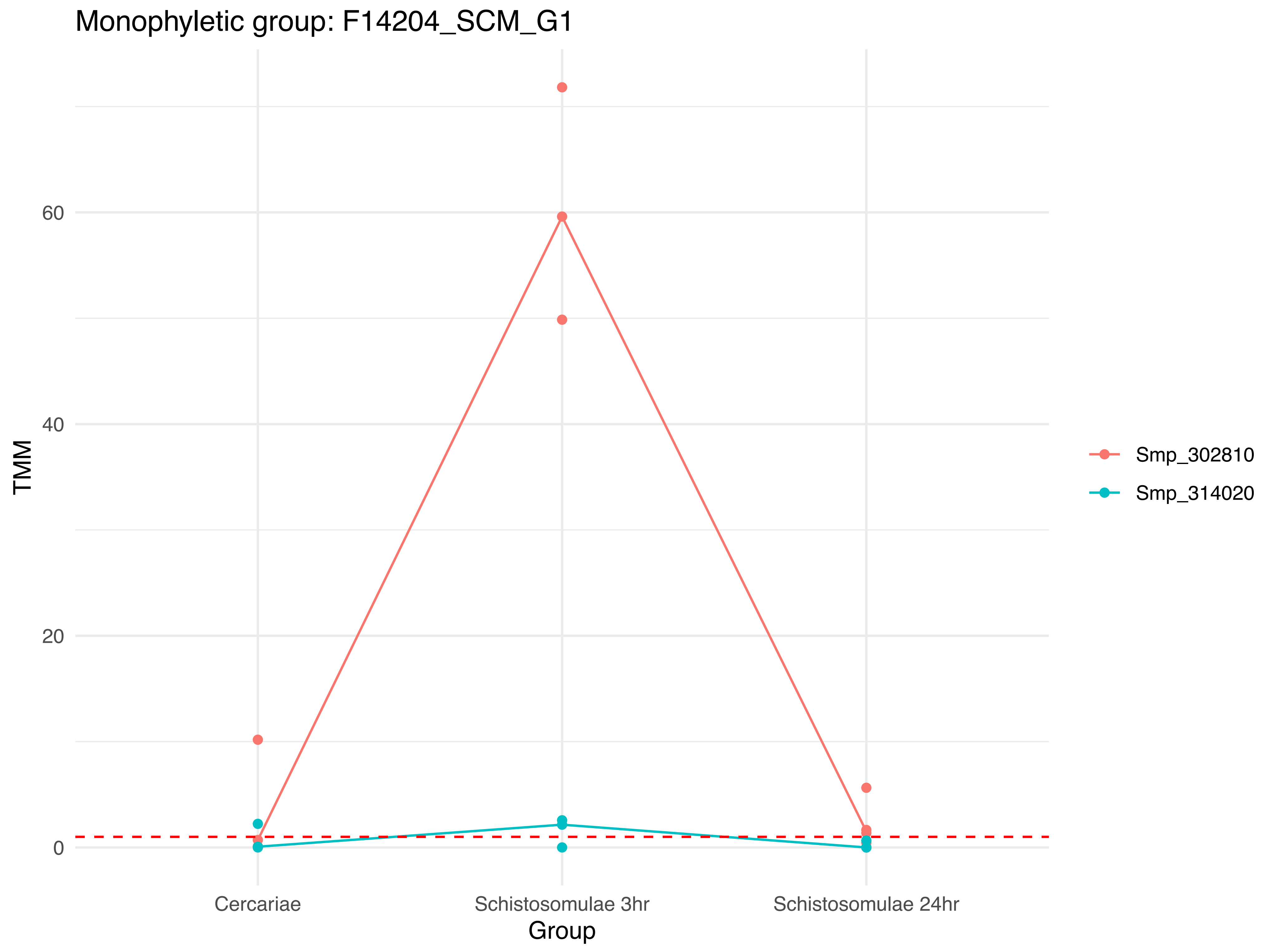

Monophyletic group: F14204_SCM_G1
TMM
60
40
20
0
Cercariae
Schistosomulae 3hr
Schistosomulae 24hr
Group
Smp_302810
Smp_314020

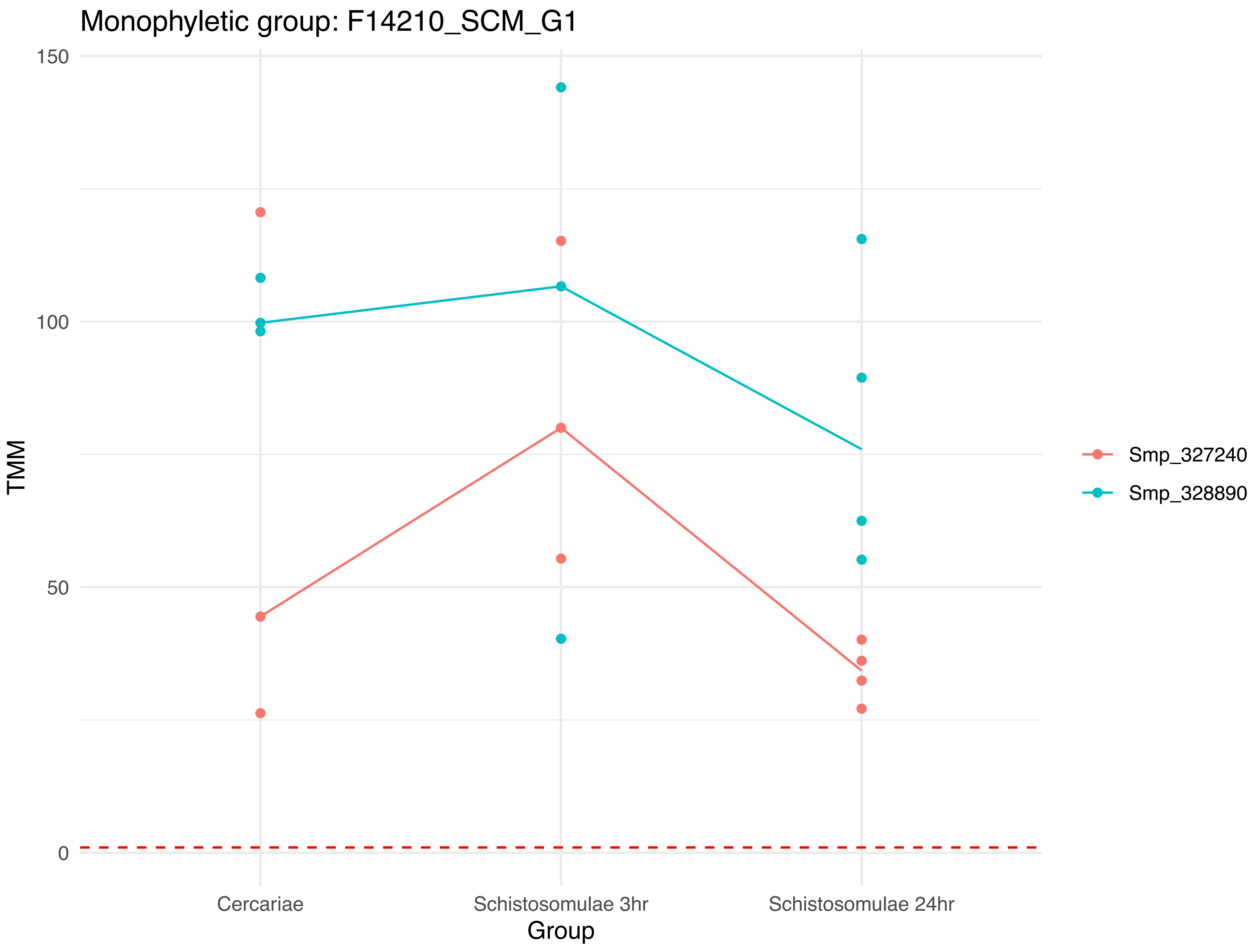

Monophyletic group: F14210_SCM_G1
150
100
50
0
TMM
Cercariae
Schistosomulae 3hr
Schistosomulae 24hr
Group
Smp_327240
Smp_328890

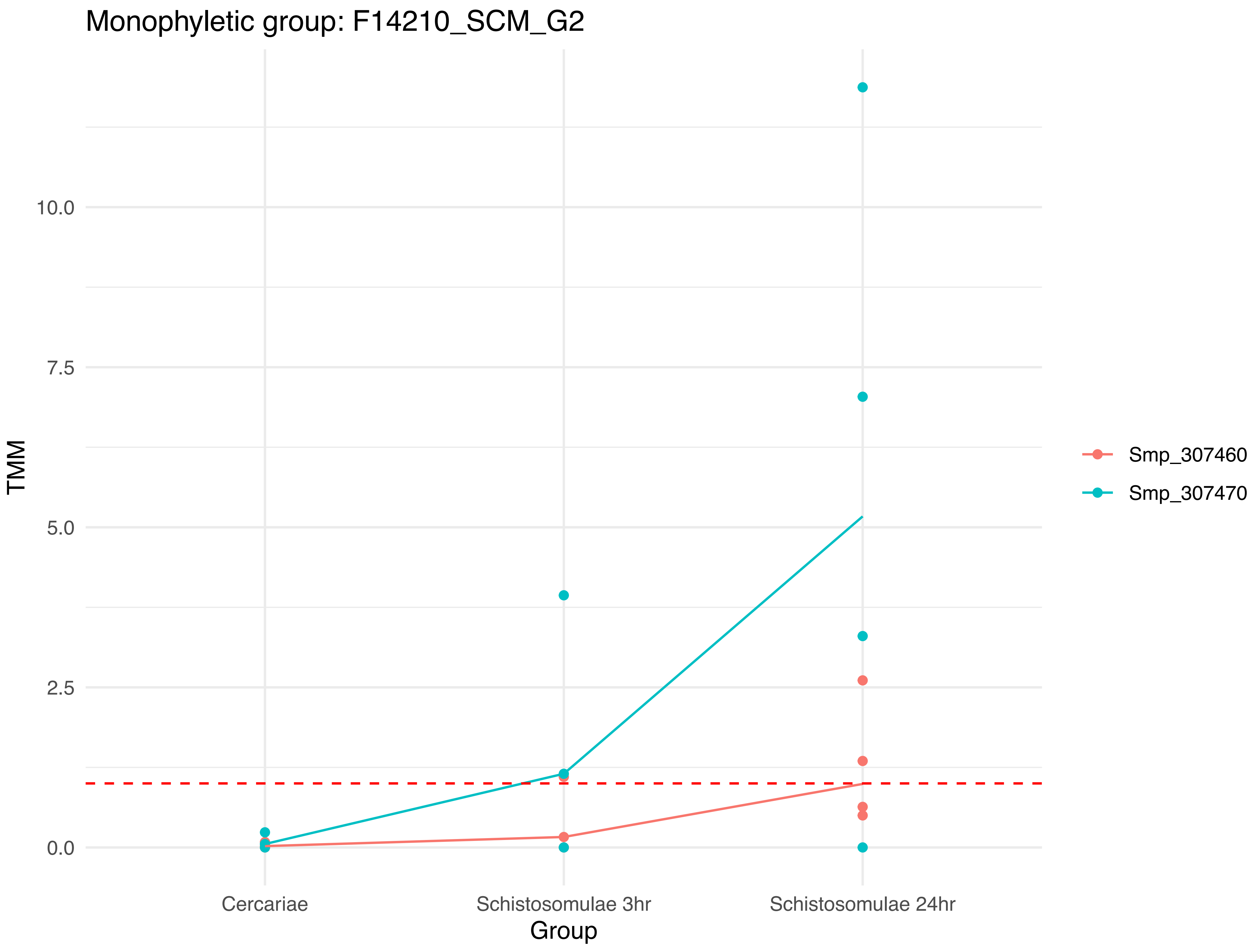
Monophyletic group: F14210_SCM_G2
TMM
10.0
7.5
5.0
2.5
0.0
Cercariae
Schistosomulae 3hr
Schistosomulae 24hr
Group
Smp_307460
Smp_307470

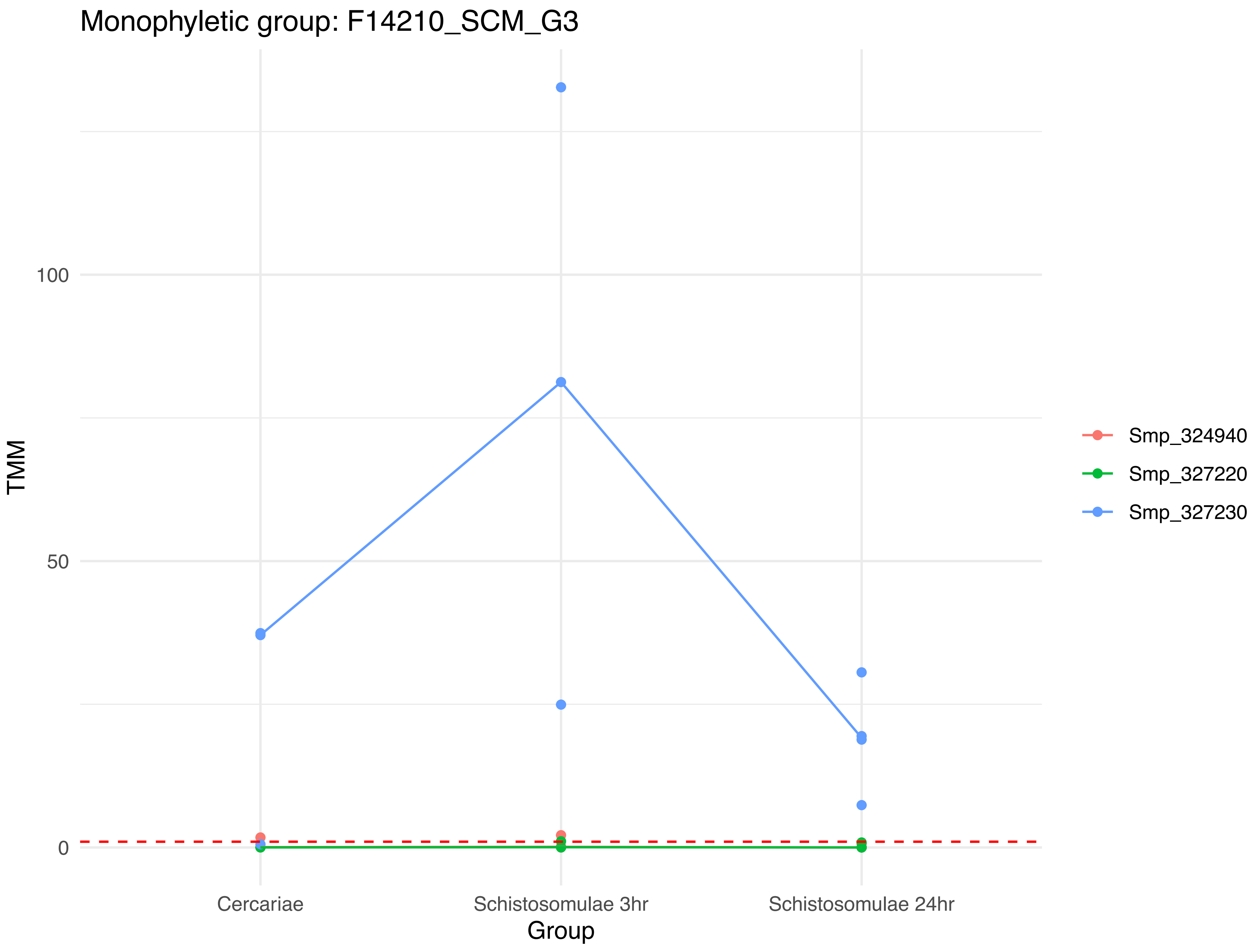
Monophyletic group: F14210_SCM_G3
100
50
0
TMM
Cercariae
Schistosomulae 3hr
Schistosomulae 24hr
Group
Smp_324940
Smp_327220
Smp_327230

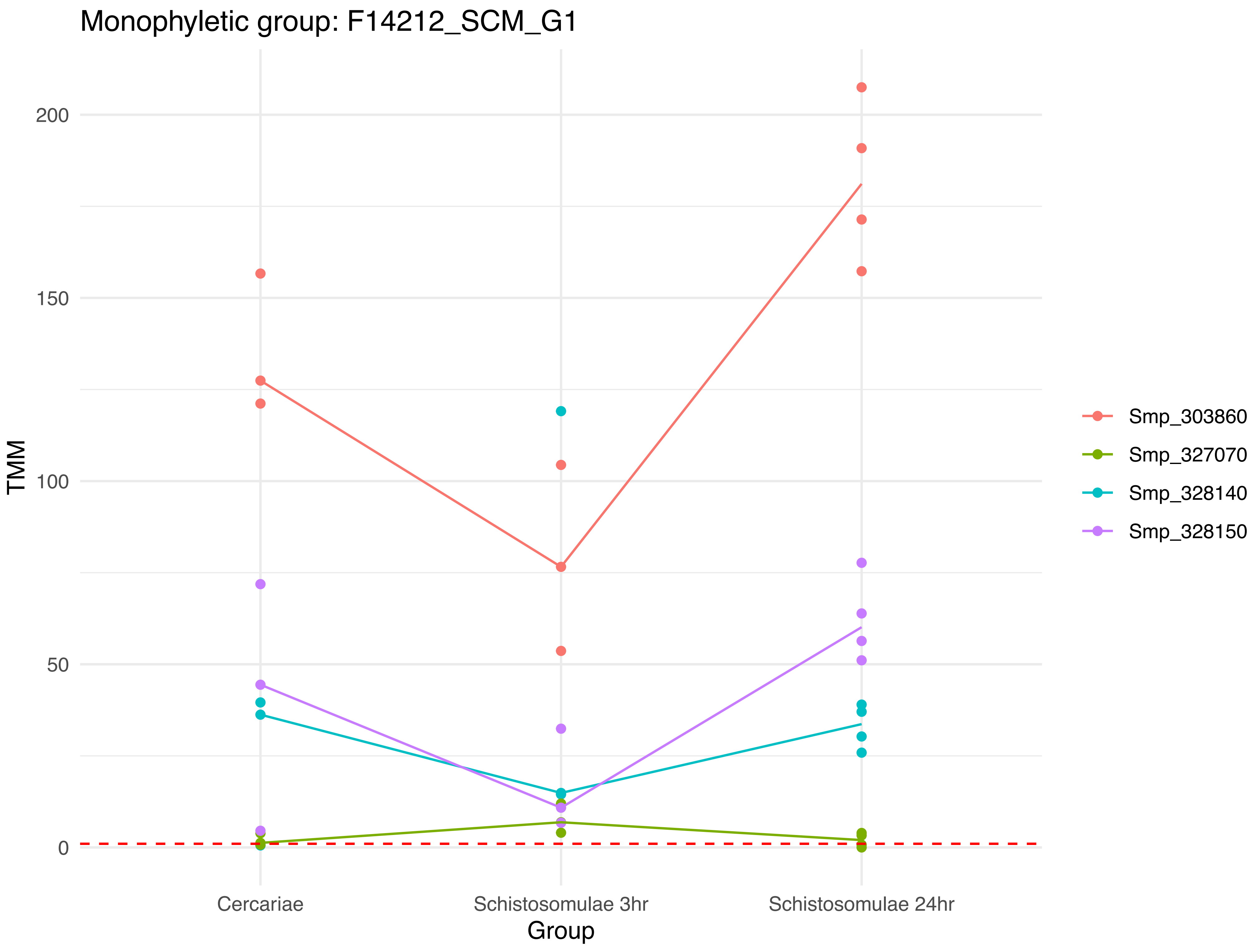

Monophyletic group: F14212_SCM_G1
TMM
200
150
100
50
0
Cercariae
Schistosomulae 3hr
Schistosomulae 24hr
Group
Smp_303860
Smp_327070
Smp_328140
Smp_328150

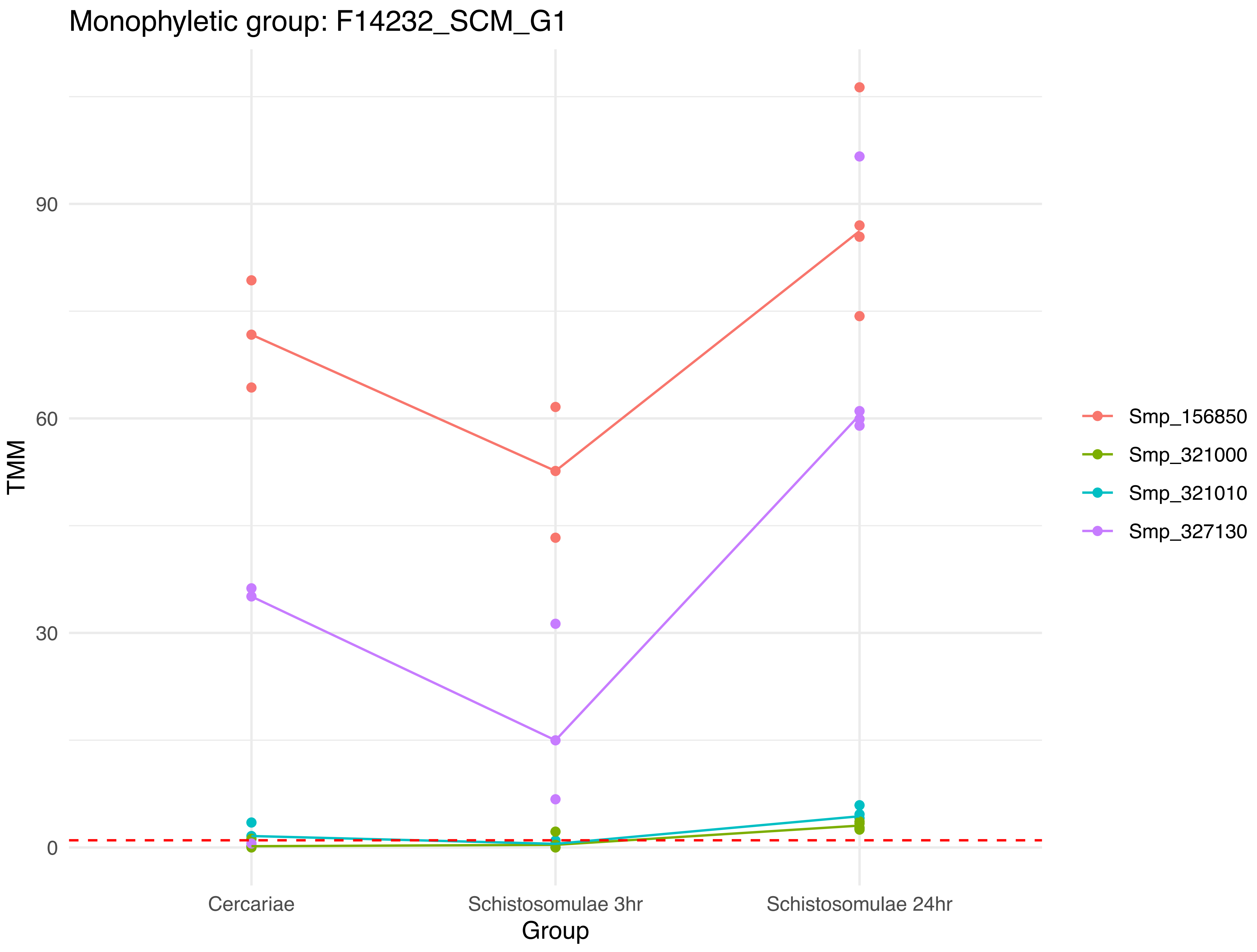

Monophyletic group: F14232_SCM_G1
TMM
90
60
30
0
Cercariae
Schistosomulae 3hr
Schistosomulae 24hr
Group
Smp_156850
Smp_321000
Smp_321010
Smp_327130

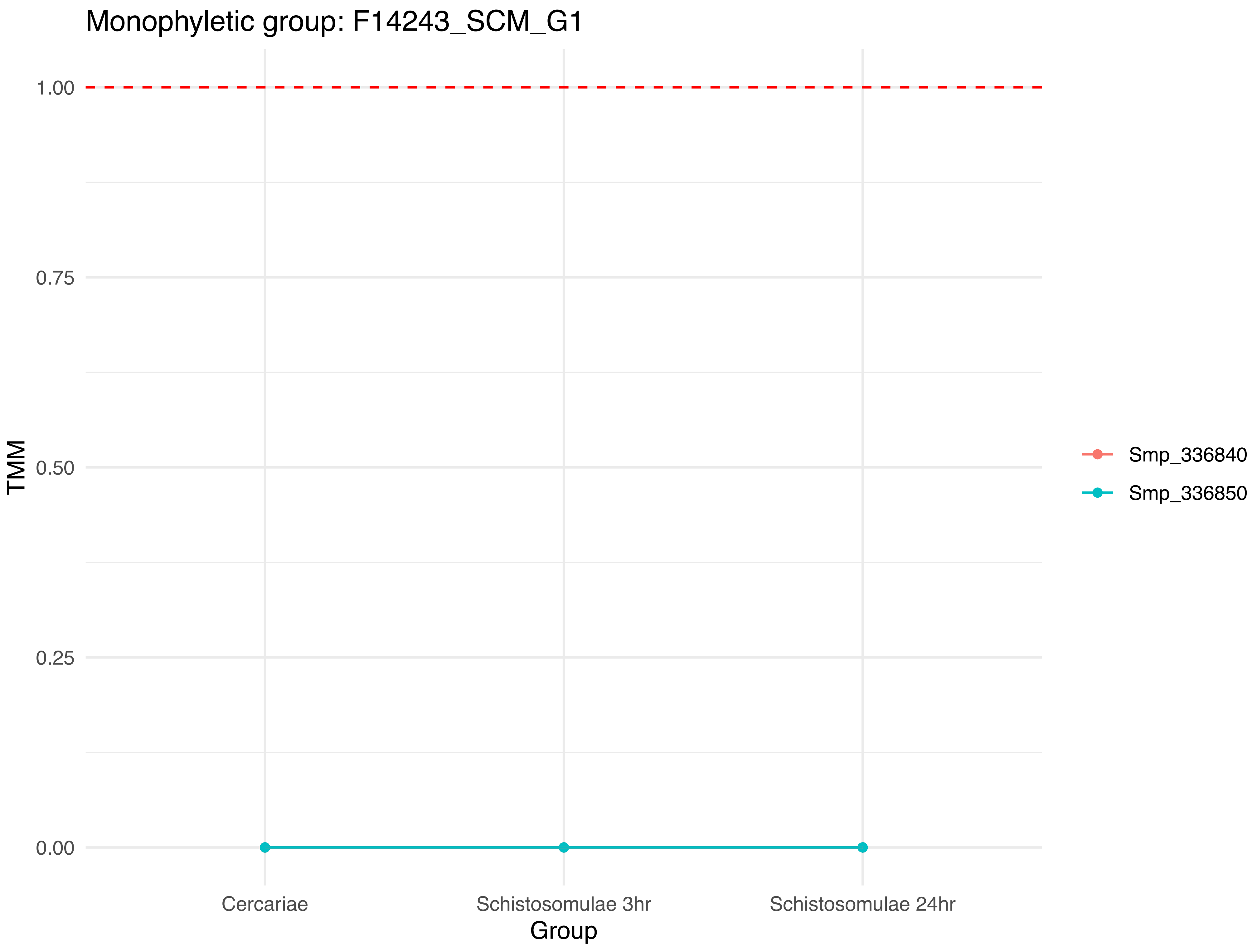

Monophyletic group: F14243_SCM_G1
TMM
1.00
0.75
0.50
0.25
0.00
Cercariae
Schistosomulae 3hr
Schistosomulae 24hr
Group
Smp_336840
Smp_336850

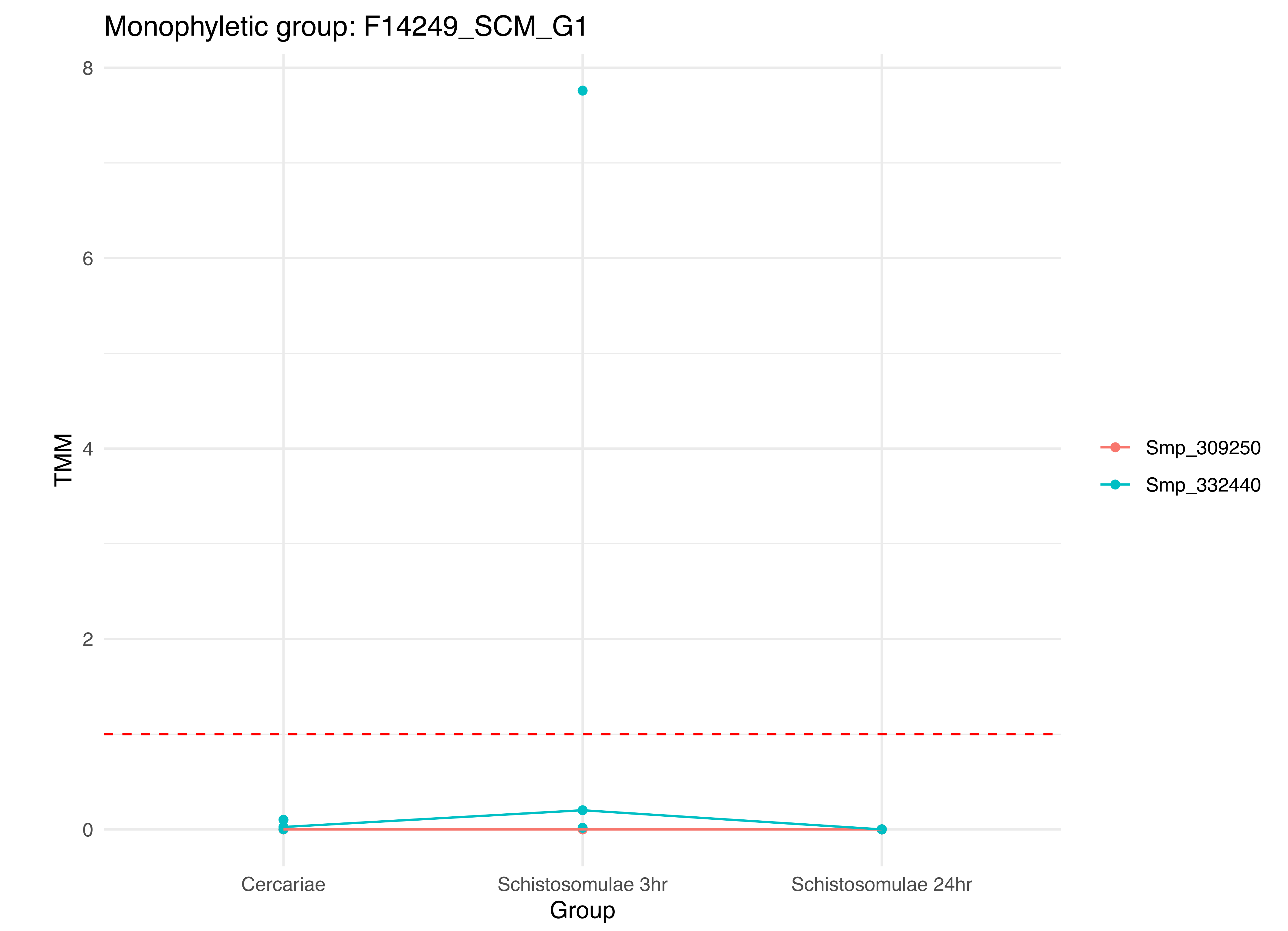
Monophyletic group: F14249_SCM_G1
TMM
0
2
4
6
8
Cercariae
Schistosomulae 3hr
Schistosomulae 24hr
Group
Smp_309250
Smp_332440

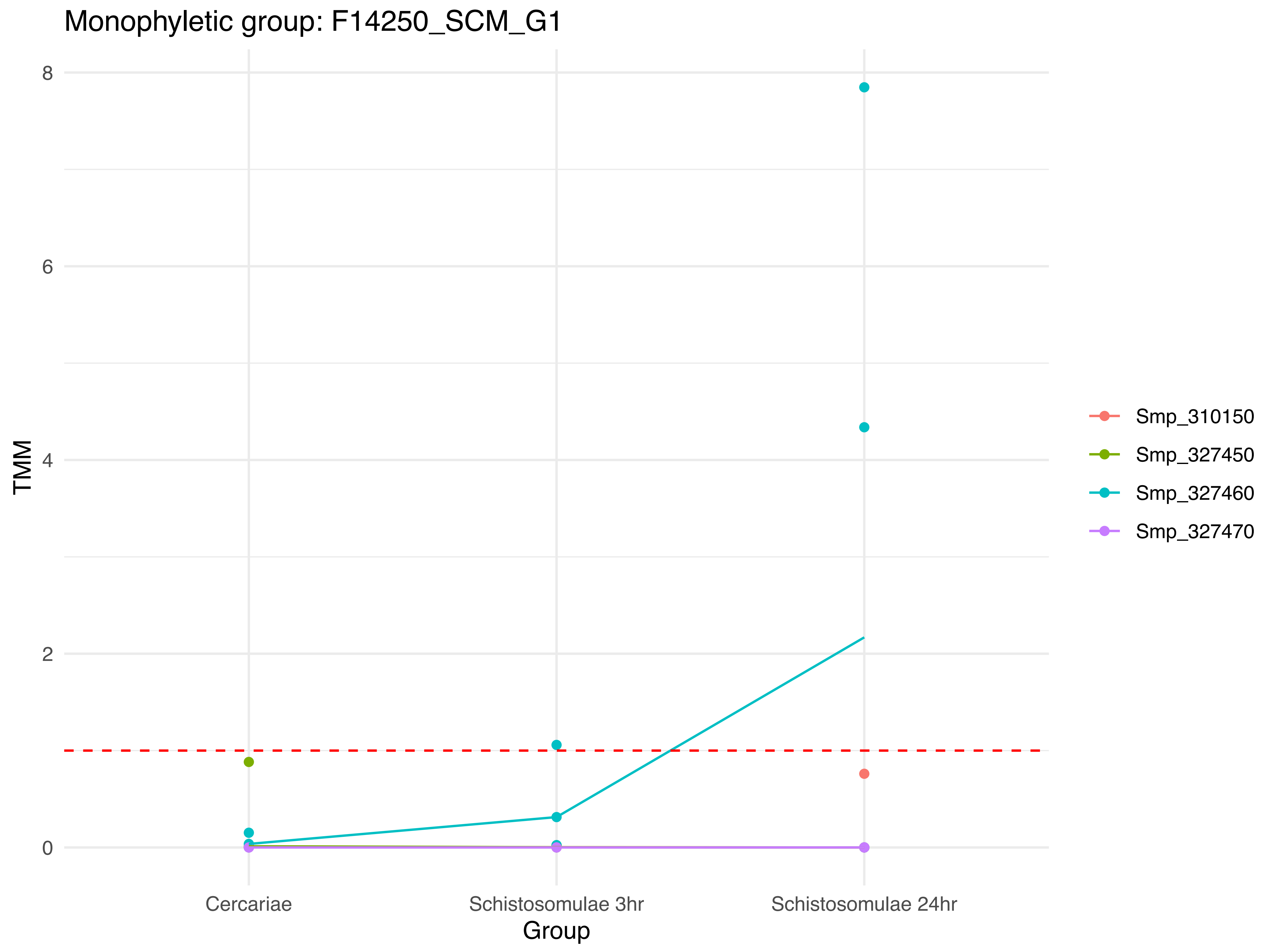

Monophyletic group: F14250_SCM_G1
TMM
8
6
4
2
0
Cercariae
Schistosomulae 3hr
Schistosomulae 24hr
Group
Smp_310150
Smp_327450
Smp_327460
Smp_327470

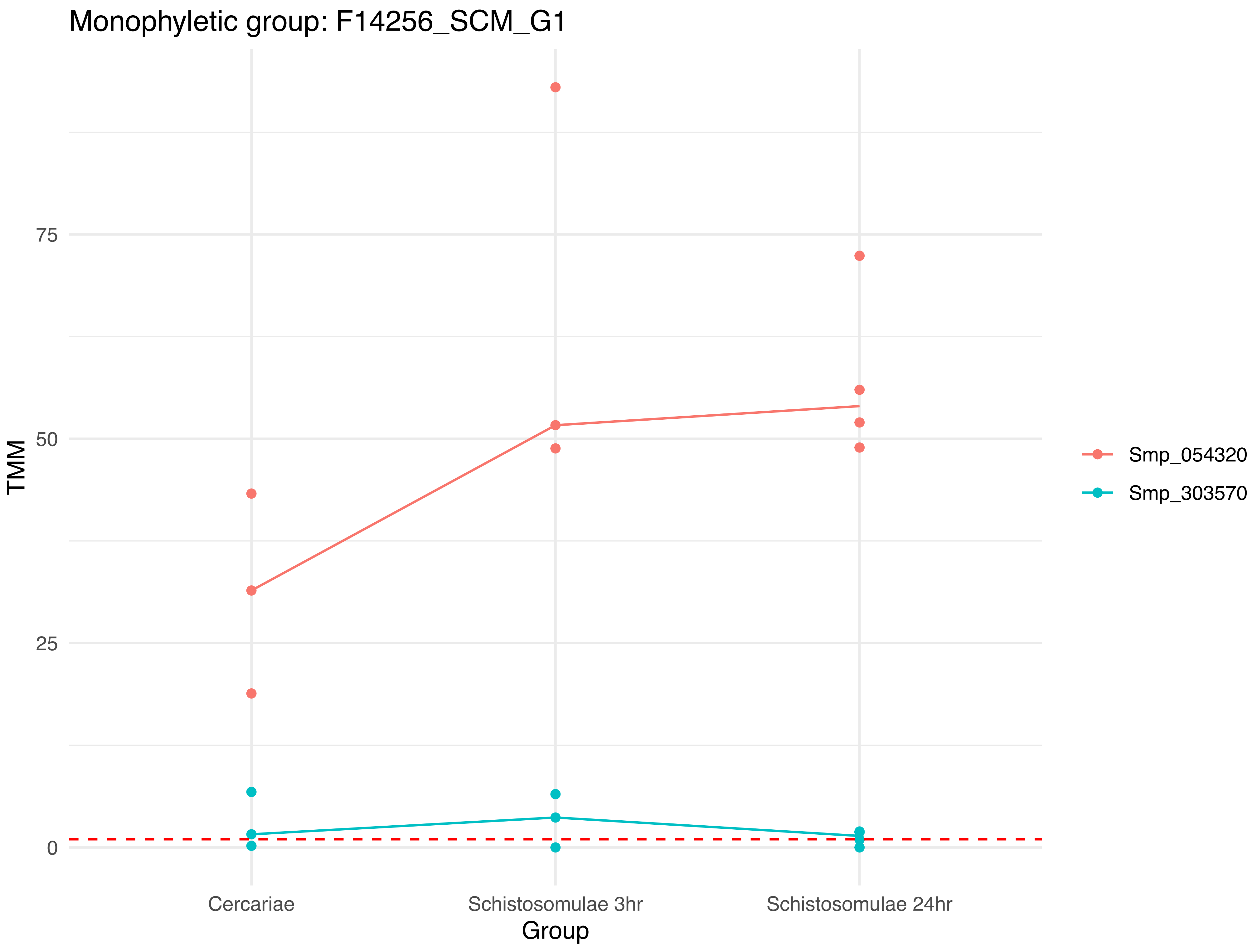

Monophyletic group: F14256_SCM_G1
TMM
75
50
25
0
Cercariae
Schistosomulae 3hr
Schistosomulae 24hr
Group
Smp_054320
Smp_303570

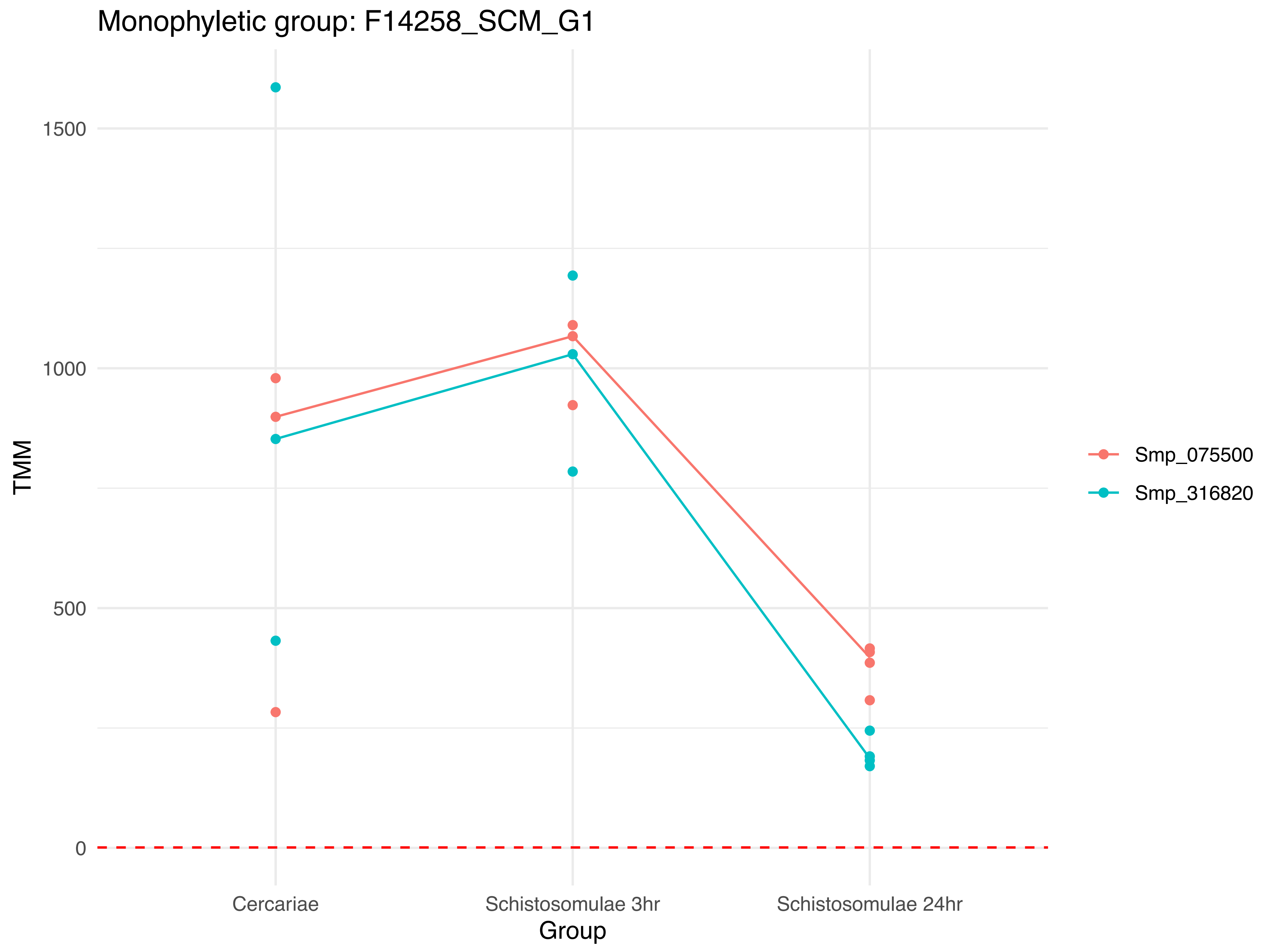

Monophyletic group: F14258_SCM_G1
TMM
1500
1000
500
0
Cercariae
Schistosomulae 3hr
Schistosomulae 24hr
Group
Smp_075500
Smp_316820

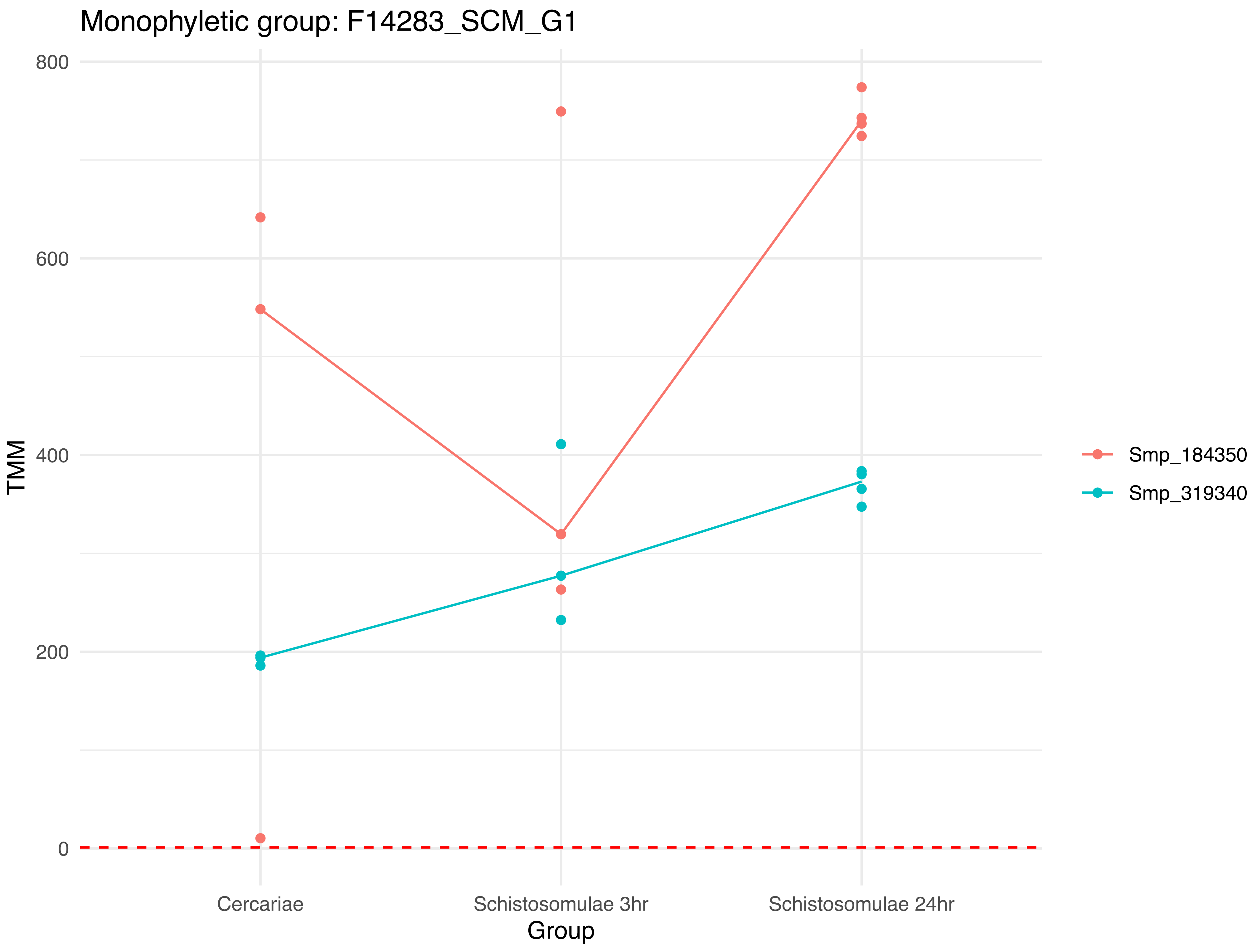
Monophyletic group: F14283_SCM_G1
TMM
800
600
400
200
0
Cercariae
Schistosomulae 3hr
Schistosomulae 24hr
Group
Smp_184350
Smp_319340

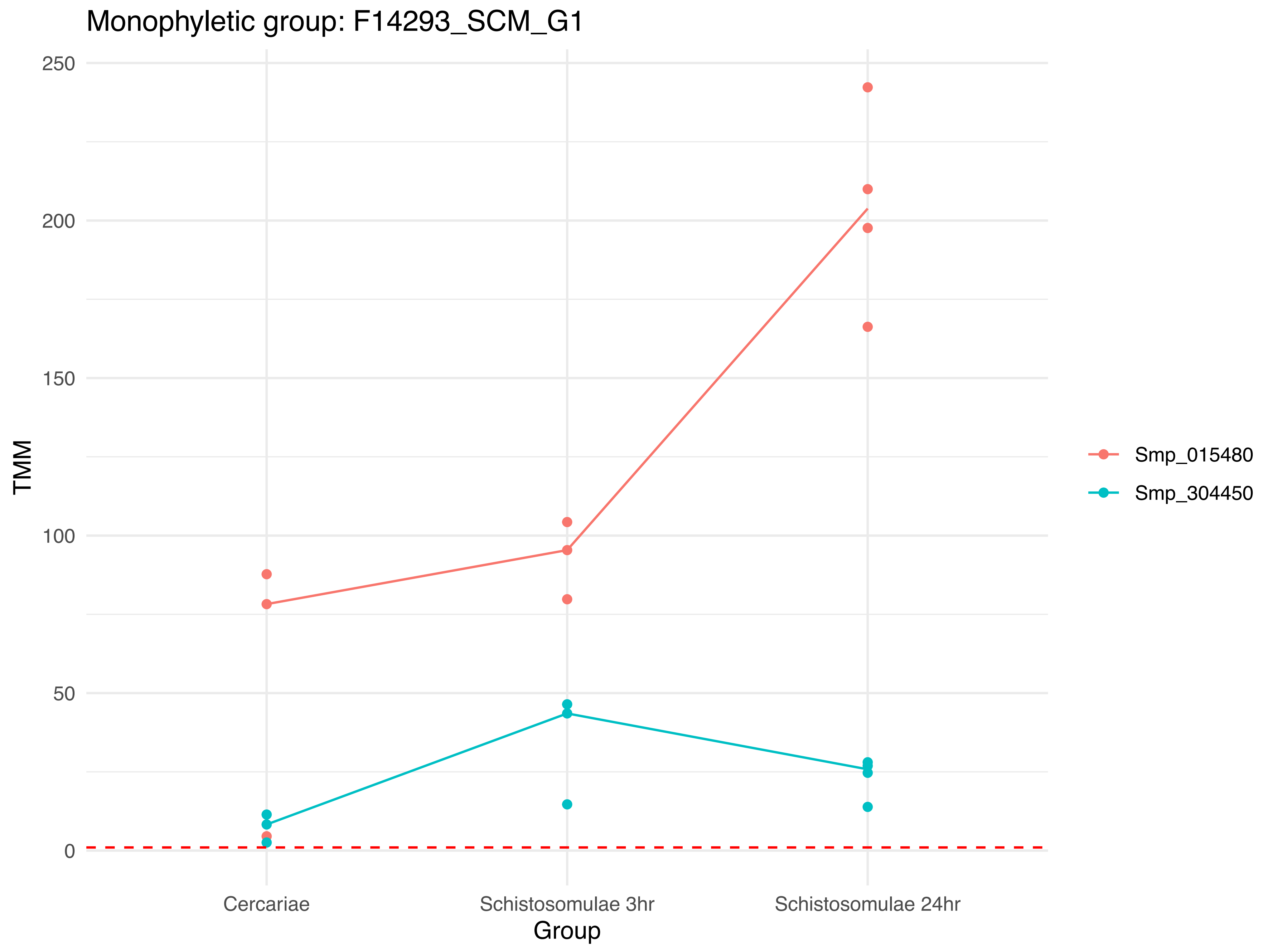
Monophyletic group: F14293_SCM_G1
250
200
150
100
50
0
TMM
Cercariae
Schistosomulae 3hr
Schistosomulae 24hr
Group
Smp_015480
Smp_304450

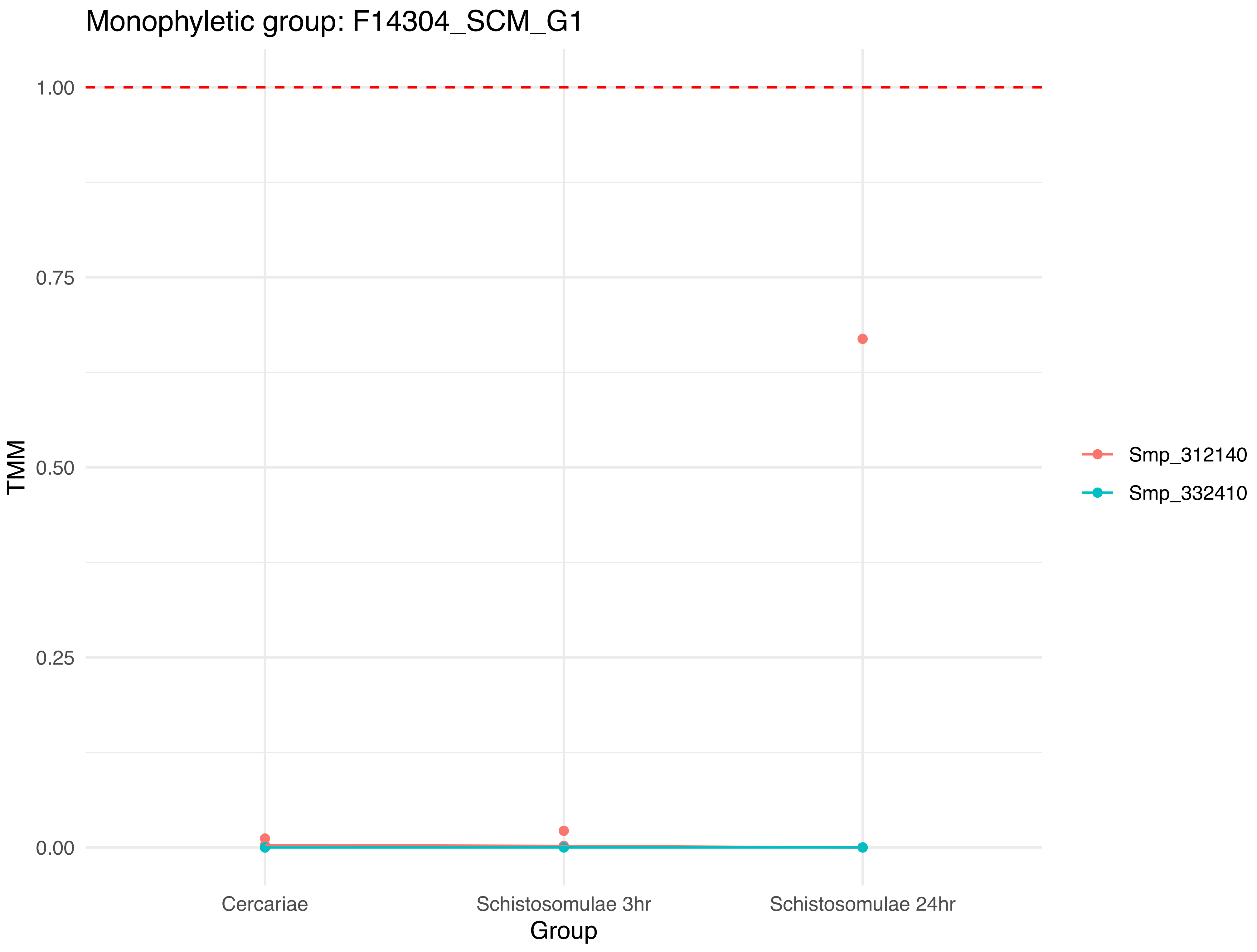

Monophyletic group: F14304_SCM_G1
1.00
0.75
0.50
0.25
0.00
TMM
Cercariae
Schistosomulae 3hr
Schistosomulae 24hr
Group
Smp_312140
Smp_332410

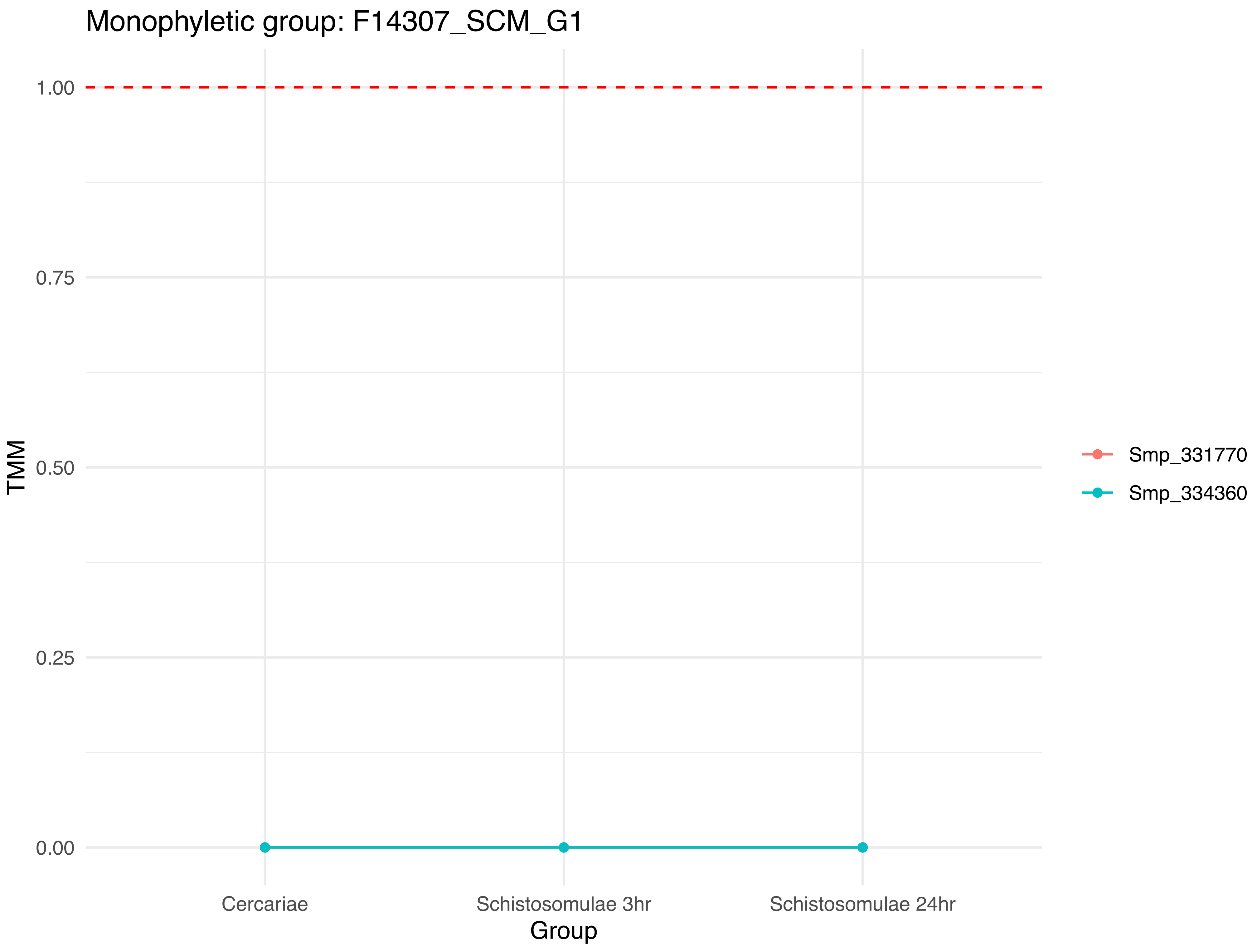
Monophyletic group: F14307_SCM_G1
1.00
0.75
0.50
0.25
0.00
TMM
Cercariae
Schistosomulae 3hr
Schistosomulae 24hr
Group
Smp_331770
Smp_334360

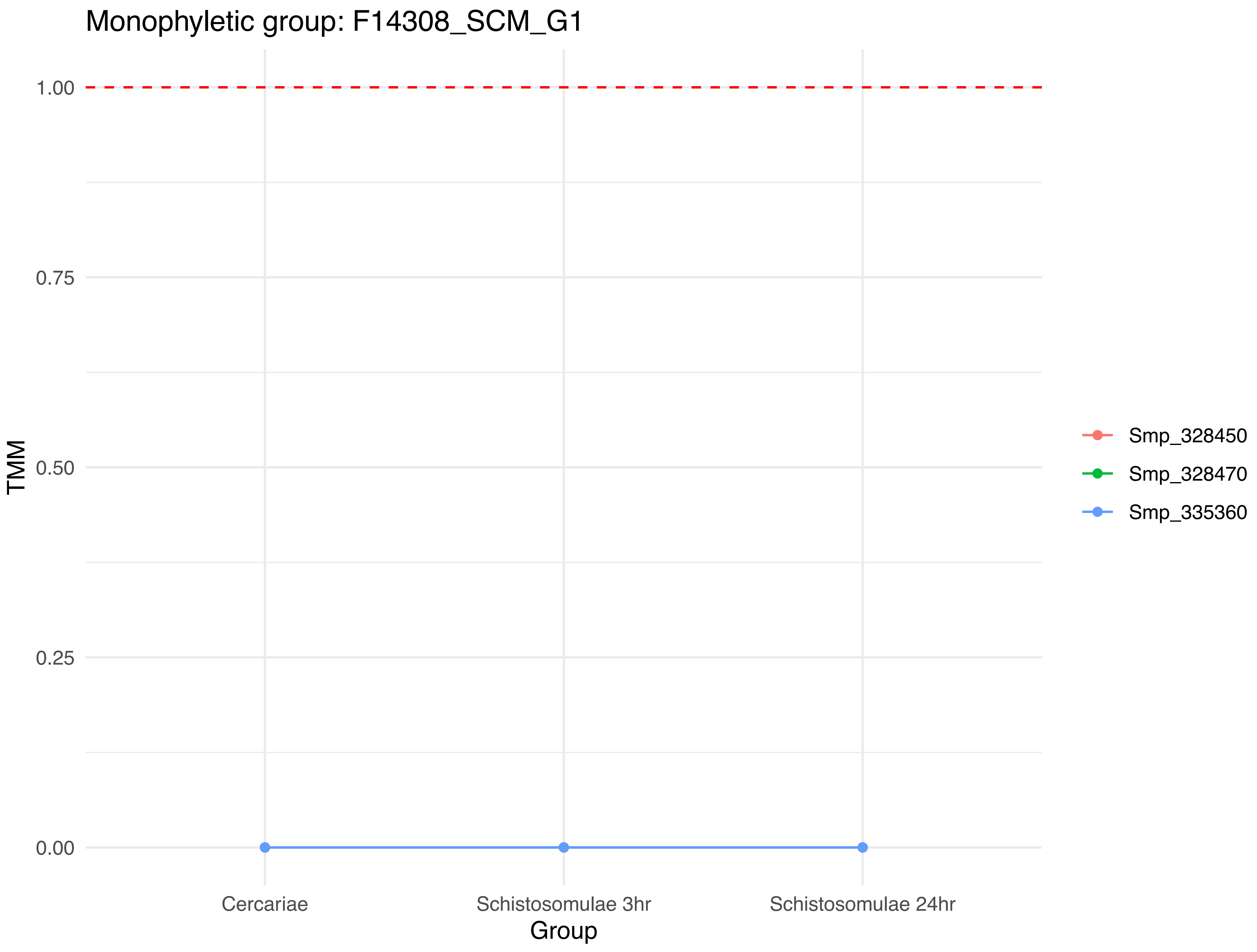

Monophyletic group: F14308_SCM_G1
TMM
1.00
0.75
0.50
0.25
0.00
Cercariae
Schistosomulae 3hr
Schistosomulae 24hr
Group
Smp_328450
Smp_328470
Smp_335360

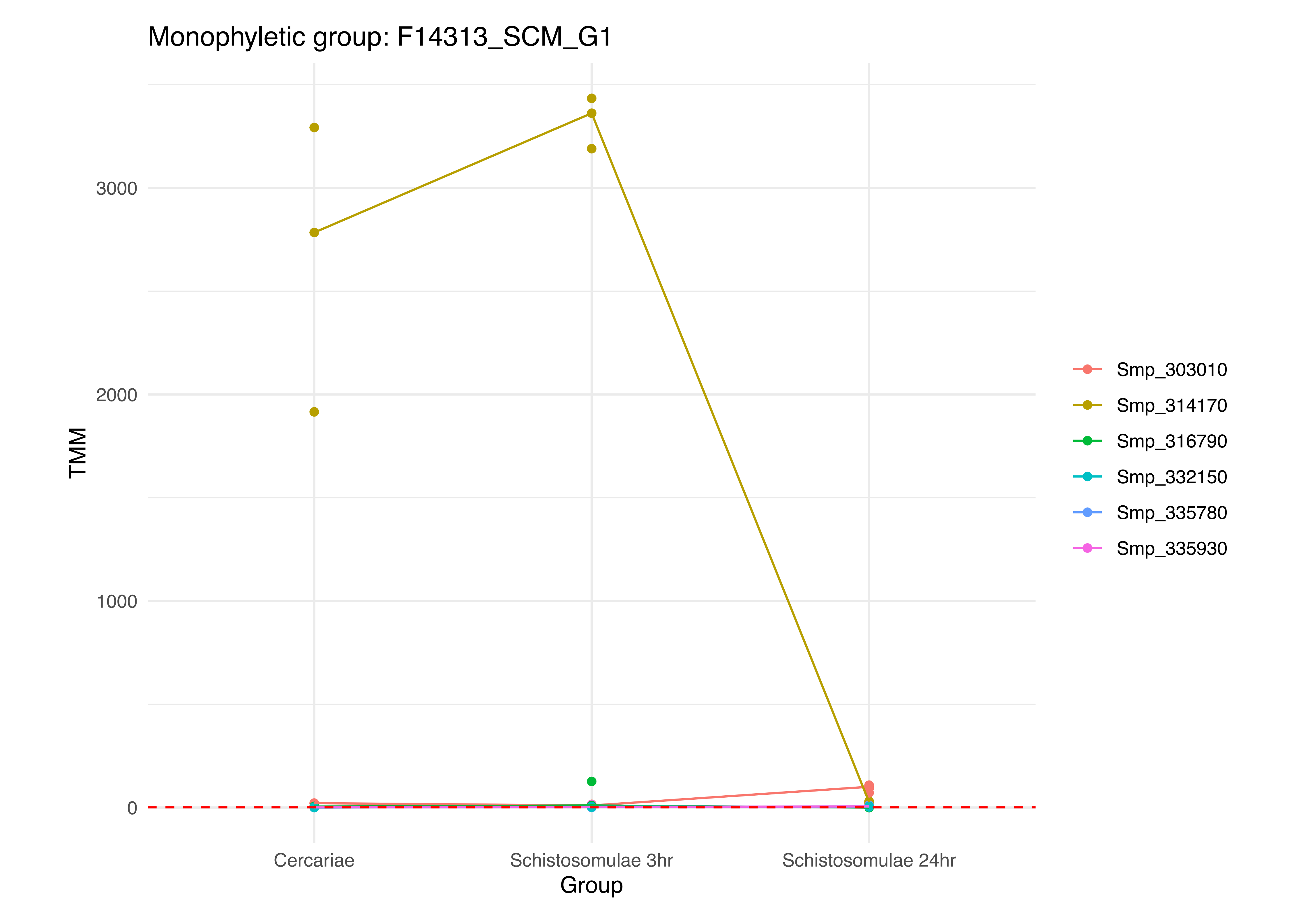

Monophyletic group: F14313_SCM_G1
TMM
3000
2000
1000
0
Cercariae
Schistosomulae 3hr
Schistosomulae 24hr
Group
Smp_303010
Smp_314170
Smp_316790
Smp_332150
Smp_335780
Smp_335930

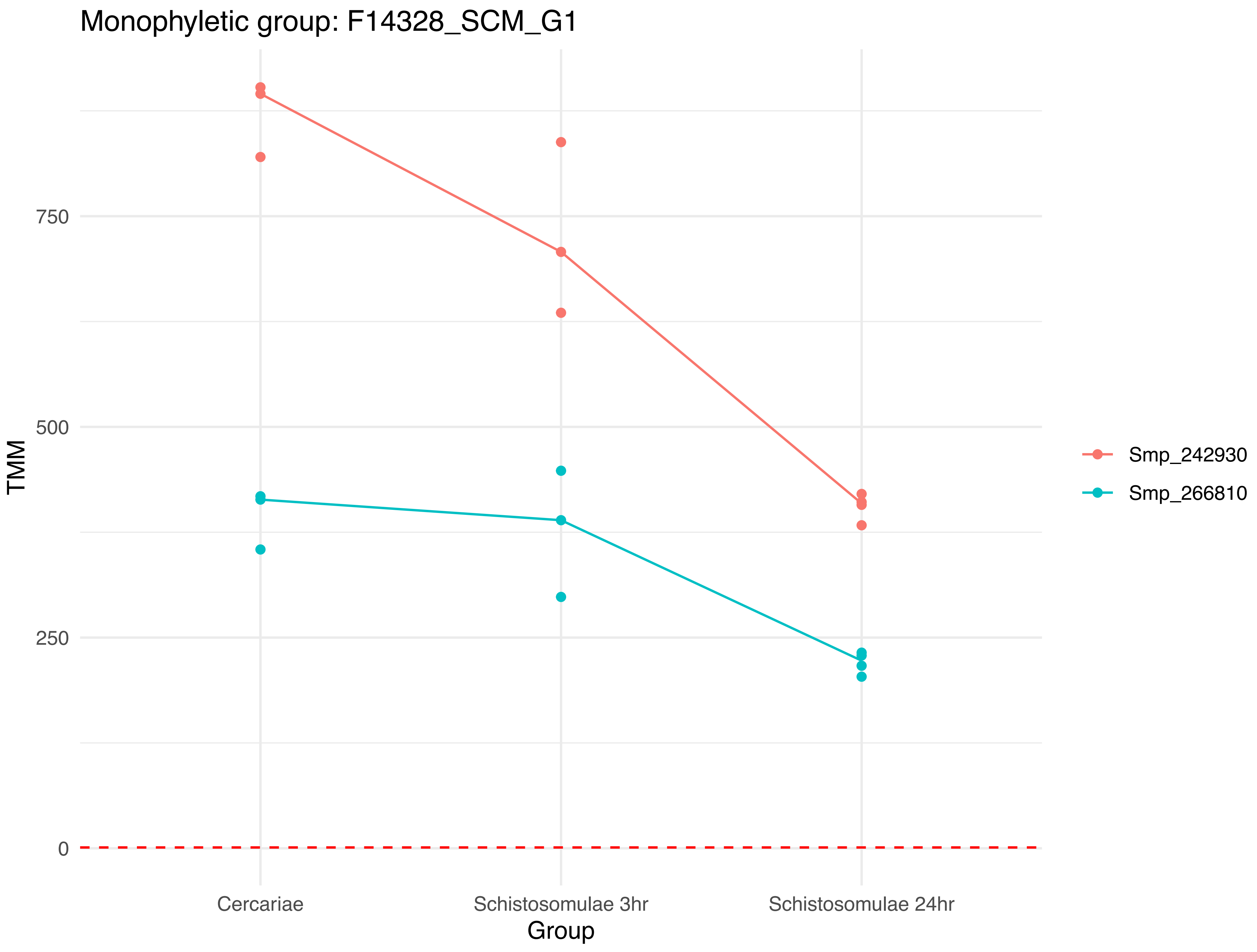

Monophyletic group: F14328_SCM_G1
TMM
750
500
250
0
Cercariae
Schistosomulae 3hr
Schistosomulae 24hr
Group
Smp_242930
Smp_266810

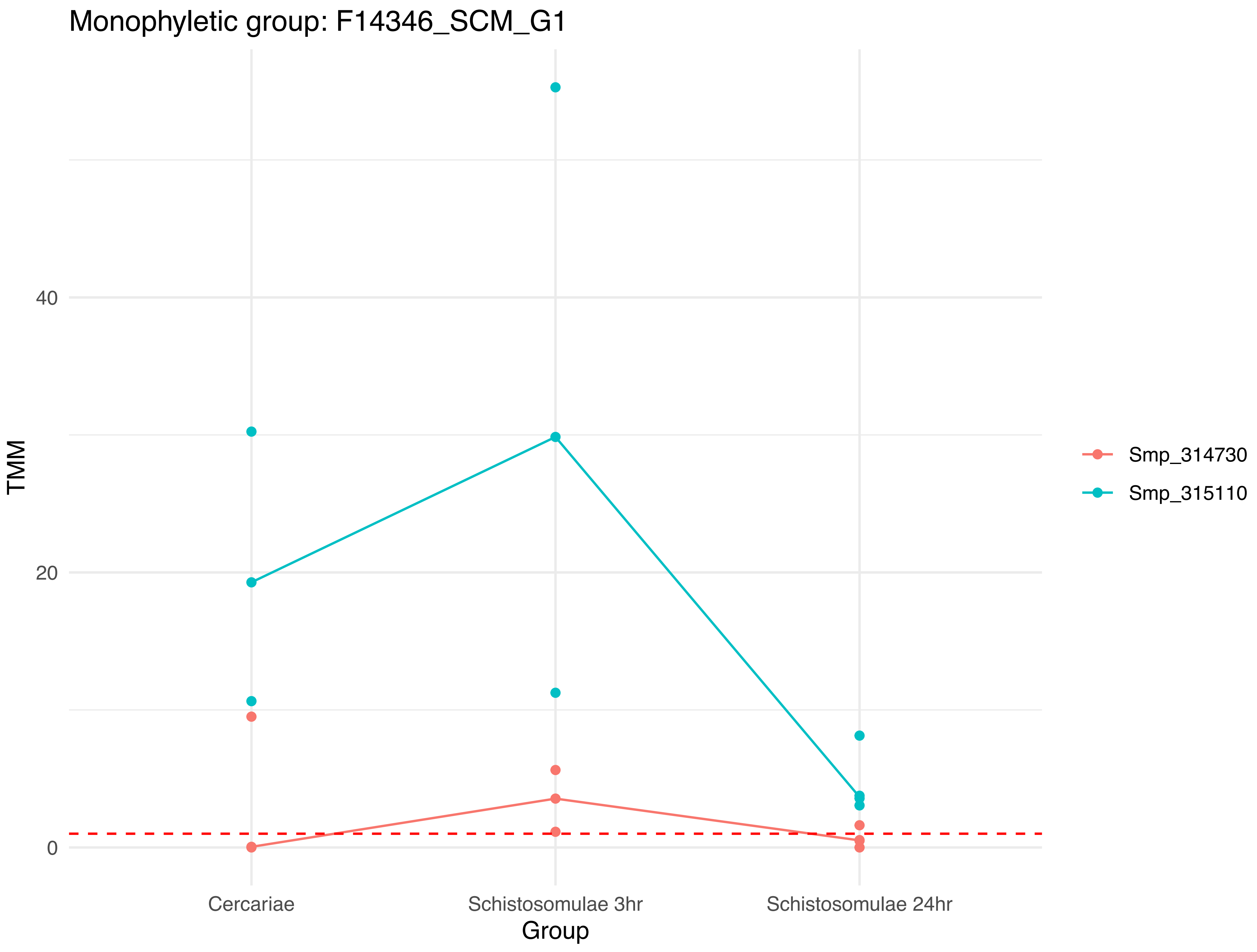

Monophyletic group: F14346_SCM_G1
TMM
0
20
40
Cercariae
Schistosomulae 3hr
Schistosomulae 24hr
Group
Smp_314730
Smp_315110

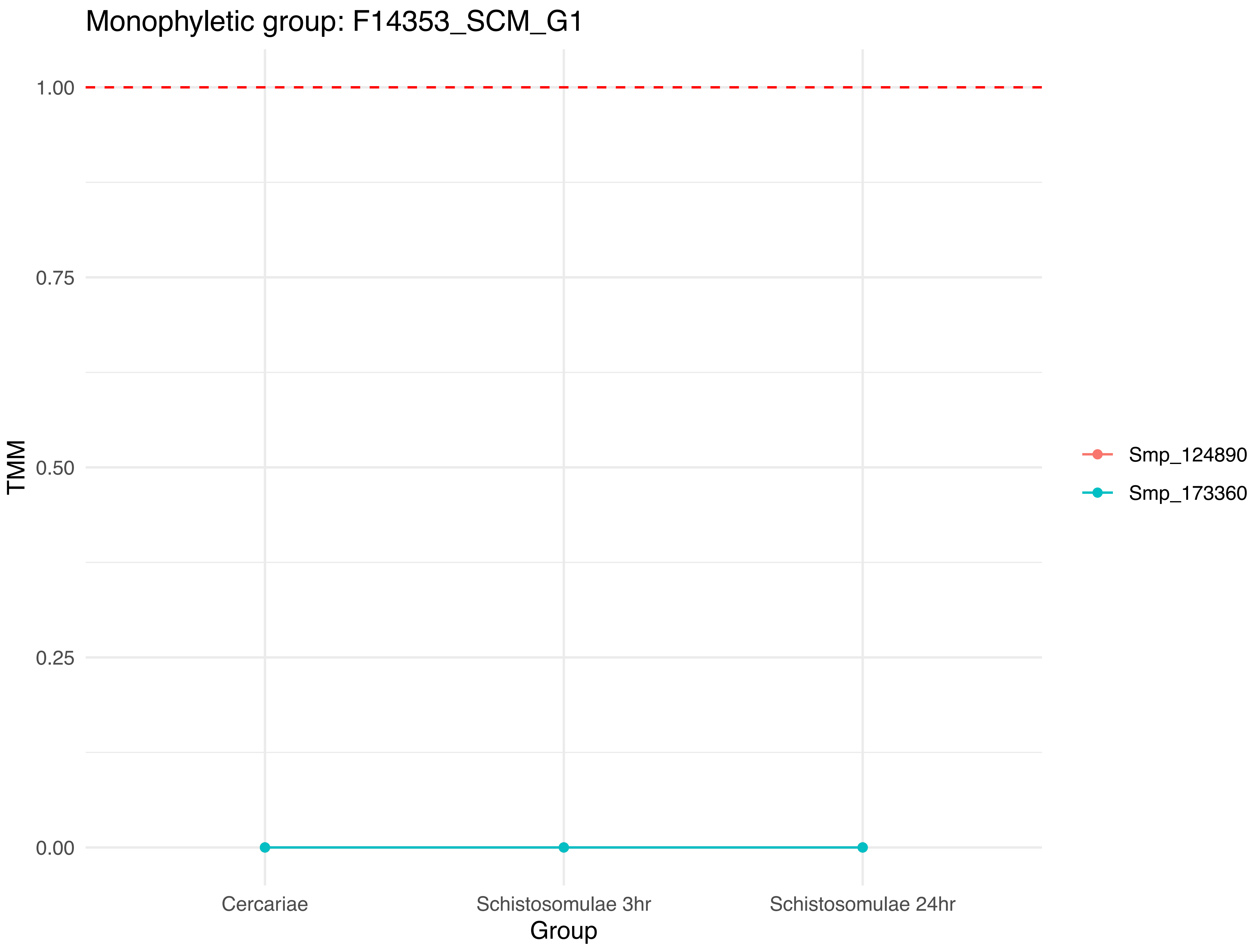

Monophyletic group: F14353_SCM_G1
TMM
1.00
0.75
0.50
0.25
0.00
Cercariae
Schistosomulae 3hr
Schistosomulae 24hr
Group
Smp_124890
Smp_173360

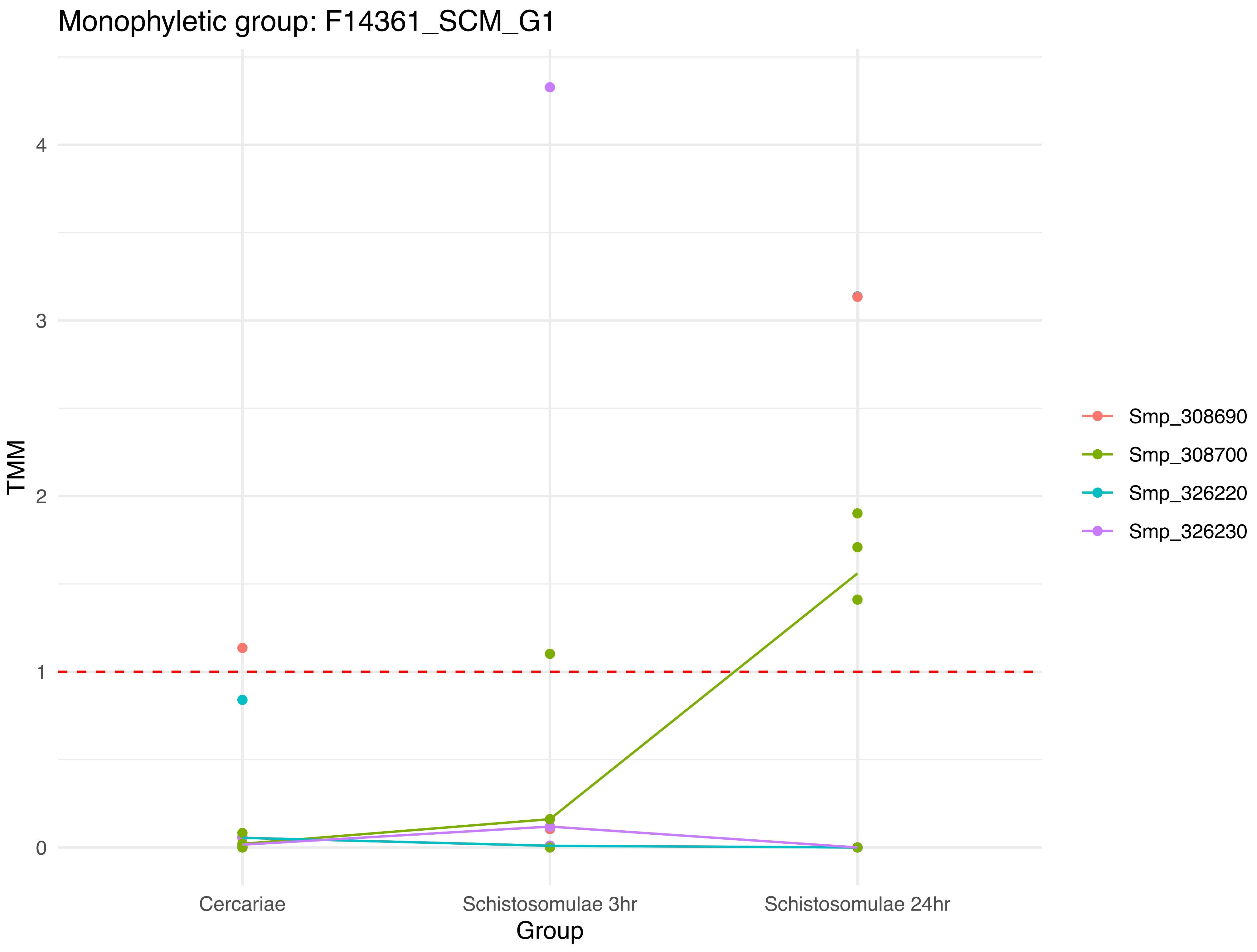

Monophyletic group: F14361_SCM_G1
TMM
4
3
2
1
0
Cercariae
Schistosomulae 3hr
Schistosomulae 24hr
Group
Smp_308690
Smp_308700
Smp_326220
Smp_326230

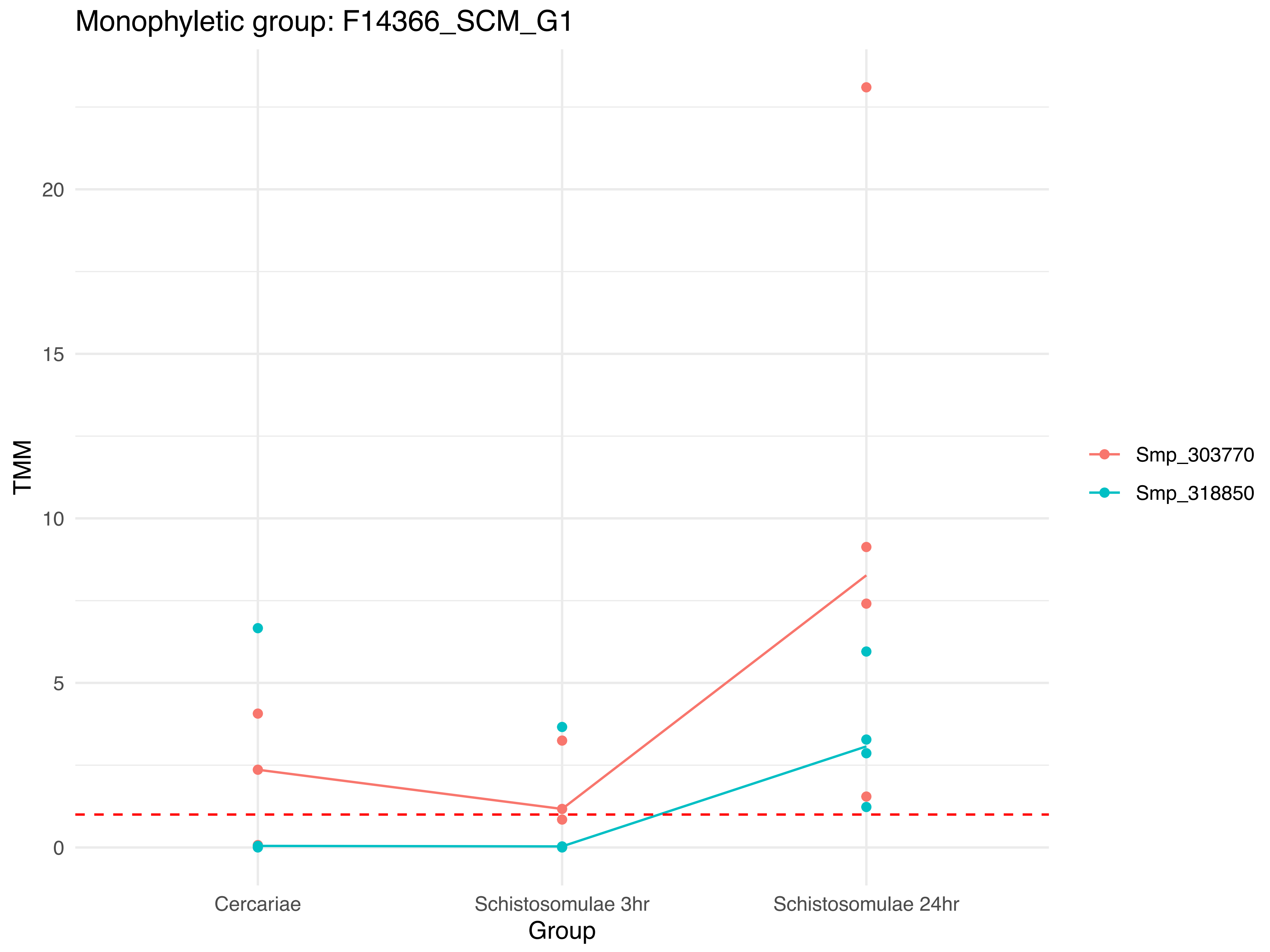
Monophyletic group: F14366_SCM_G1
TMM
0
5
10
15
20
Cercariae
Schistosomulae 3hr
Schistosomulae 24hr
Group
Smp_303770
Smp_318850

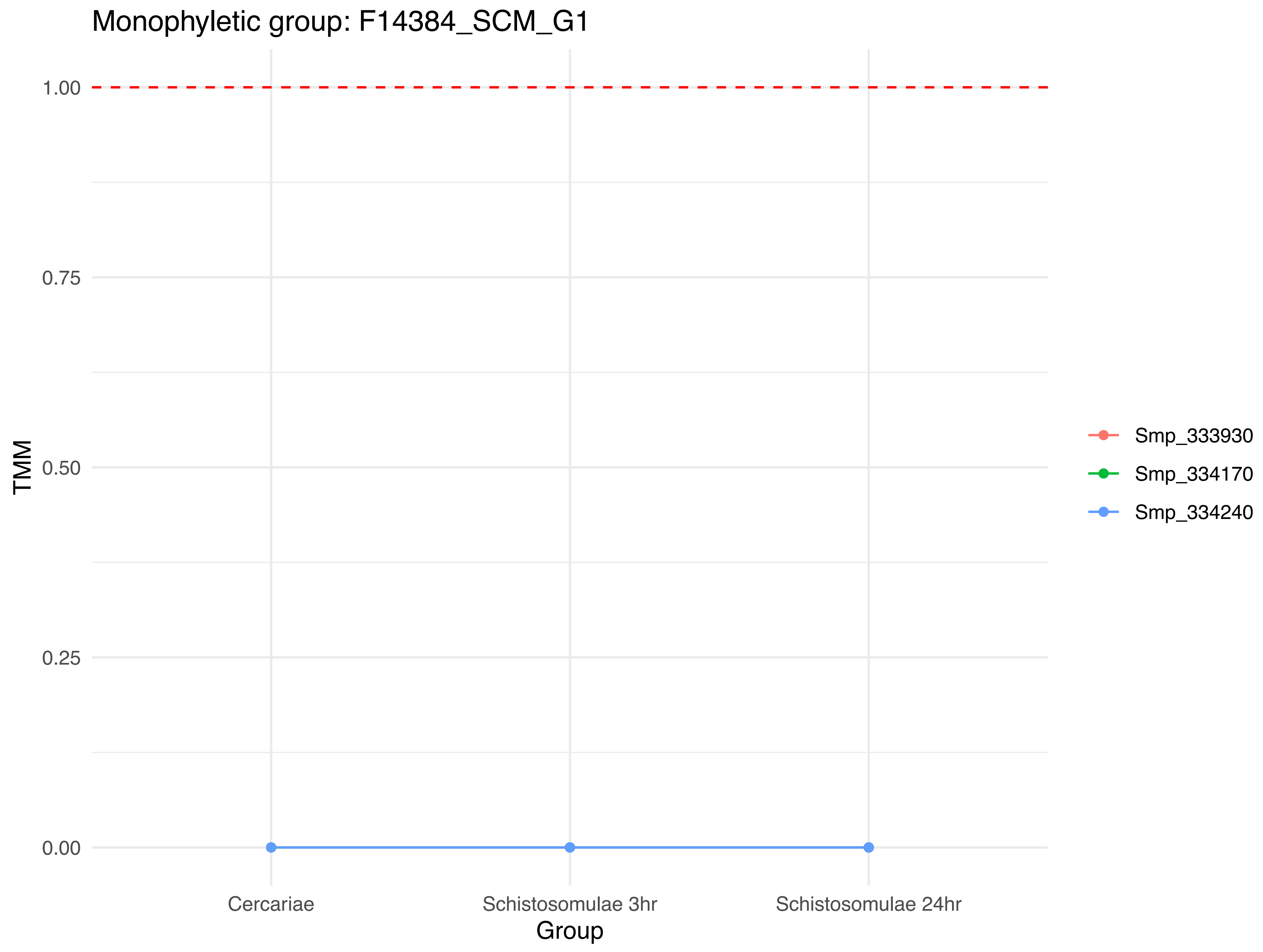

Monophyletic group: F14384_SCM_G1
TMM
1.00
0.75
0.50
0.25
0.00
Cercariae
Schistosomulae 3hr
Schistosomulae 24hr
Group
Smp_333930
Smp_334170
Smp_334240

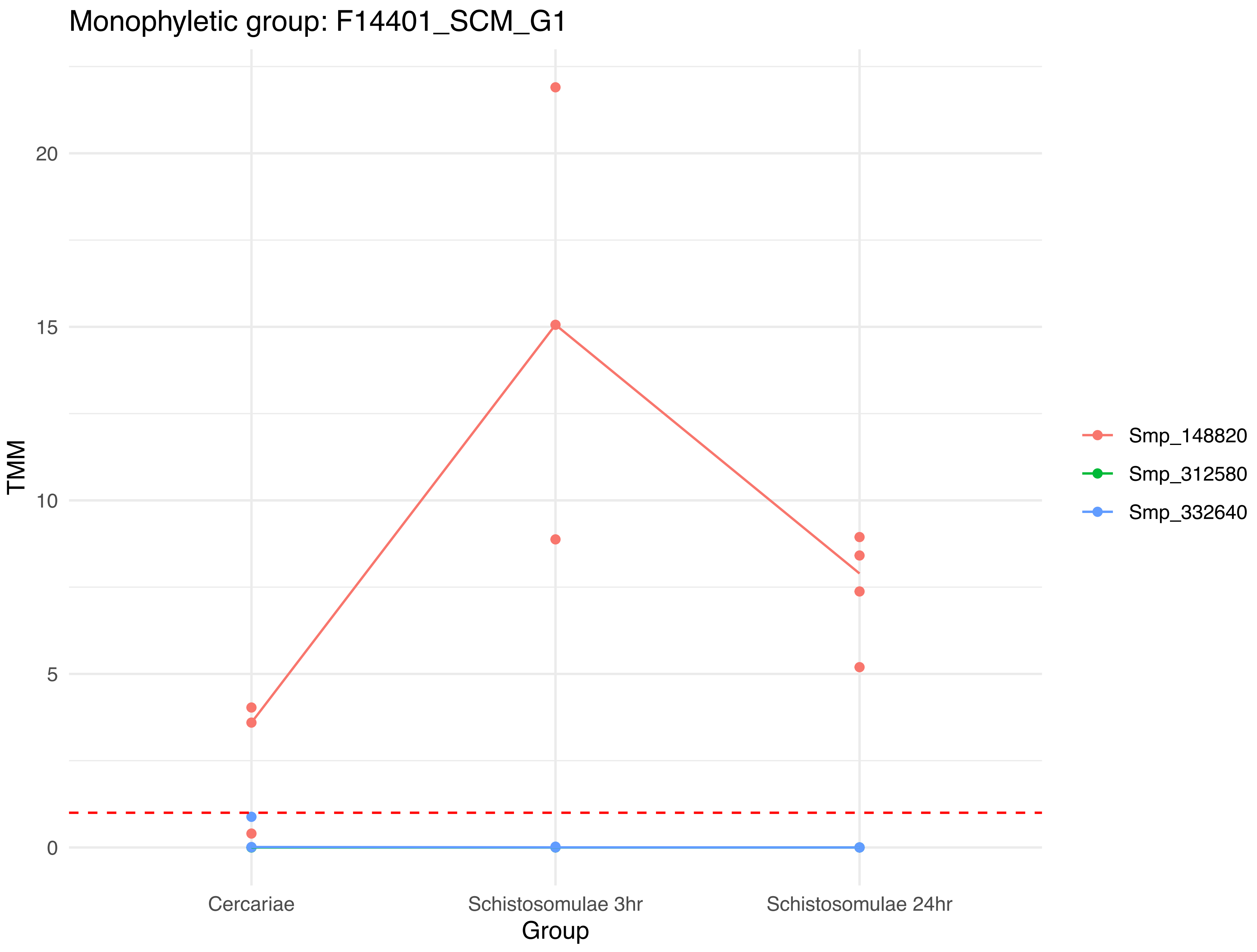

Monophyletic group: F14401_SCM_G1
TMM
20
15
10
5
0
Cercariae
Schistosomulae 3hr
Schistosomulae 24hr
Group
Smp_148820
Smp_312580
Smp_332640

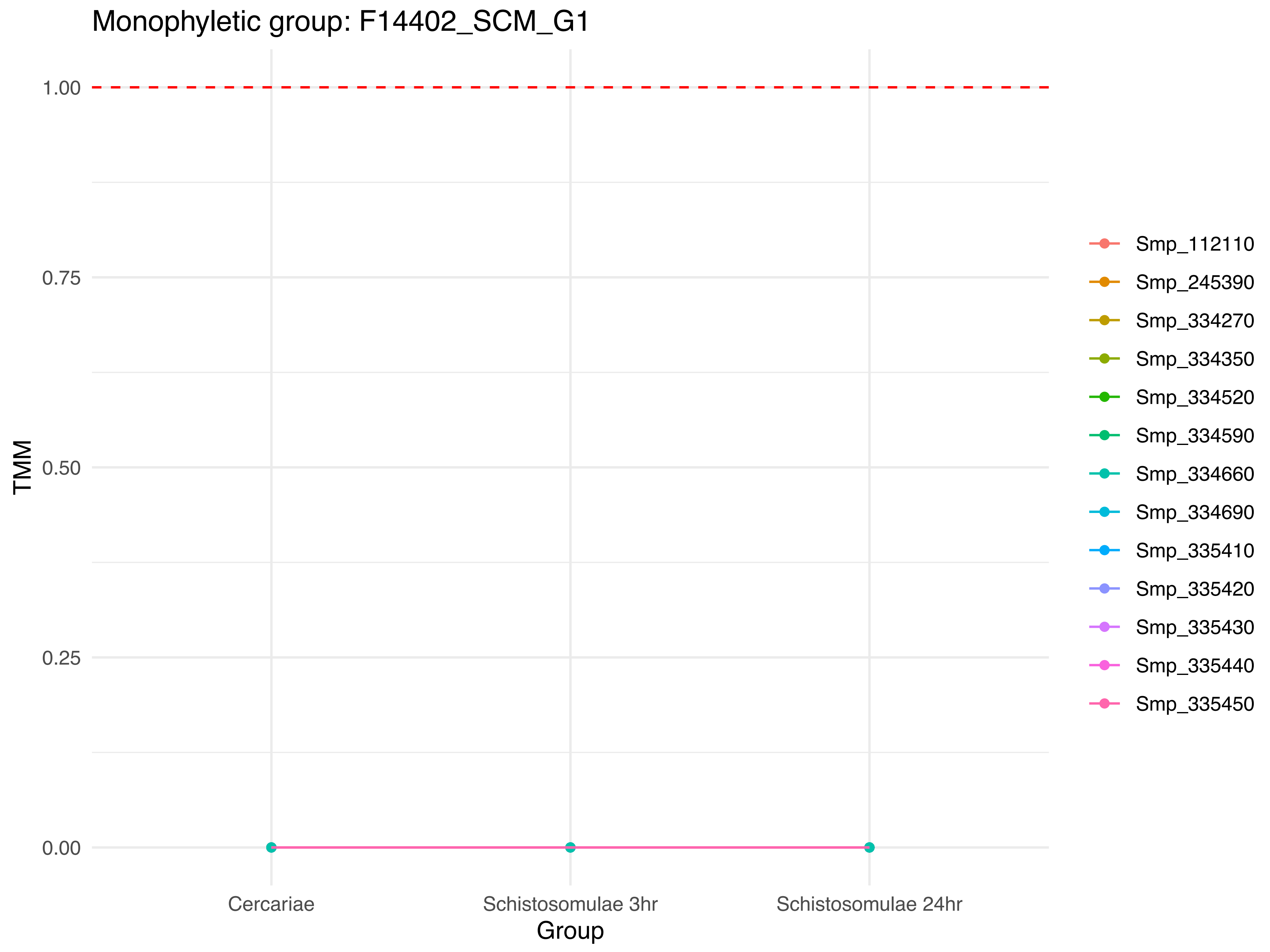
Monophyletic group: F14402_SCM_G1
TMM
1.00
0.75
0.50
0.25
0.00
Cercariae
Schistosomulae 3hr
Schistosomulae 24hr
Group
Smp_112110
Smp_245390
Smp_334270
Smp_334350
Smp_334520
Smp_334590
Smp_334660
Smp_334690
Smp_335410
Smp_335420
Smp_335430
Smp_335440
Smp_335450

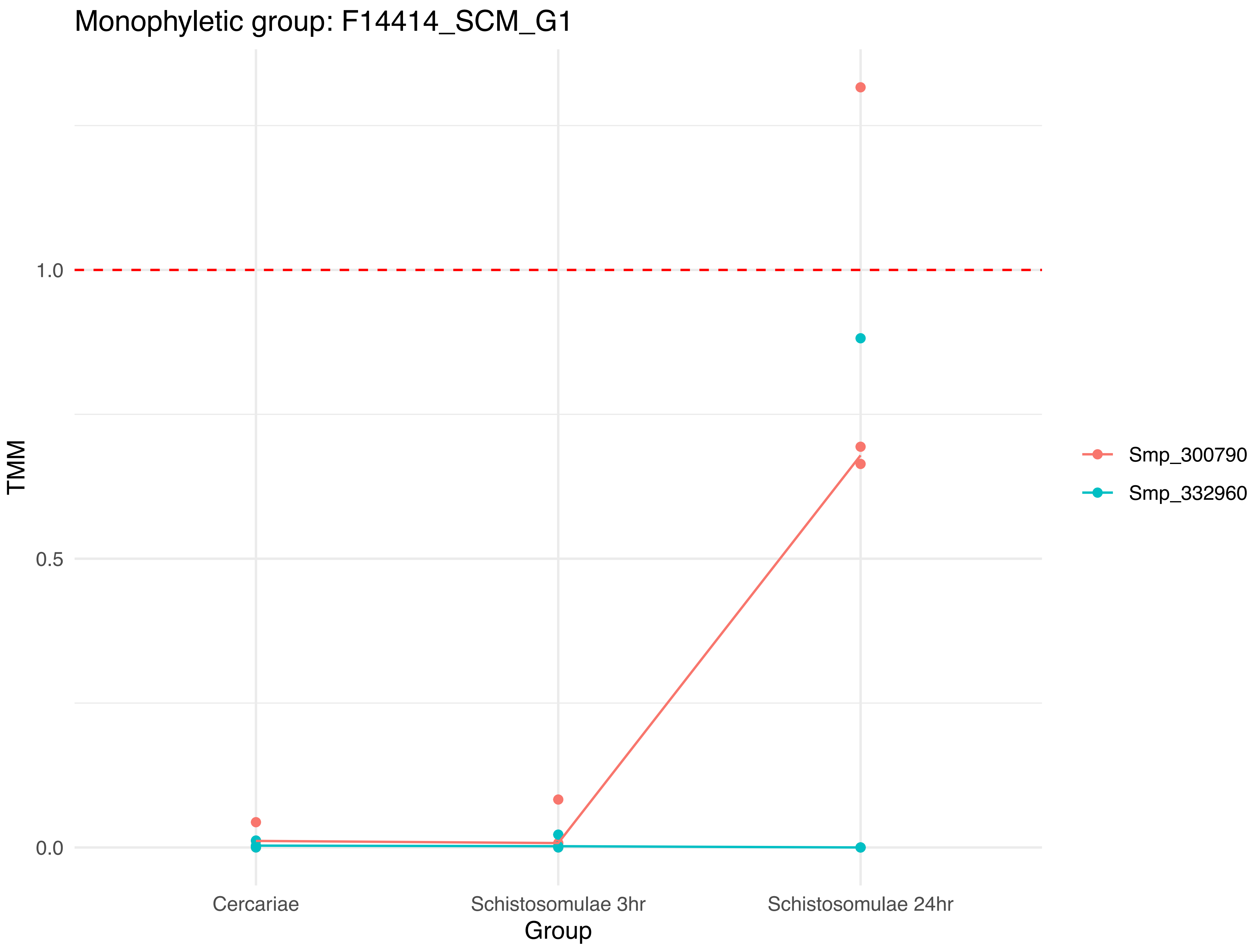

Monophyletic group: F14414_SCM_G1
1.0
0.5
0.0
TMM
Cercariae
Schistosomulae 3hr
Schistosomulae 24hr
Group
Smp_300790
Smp_332960

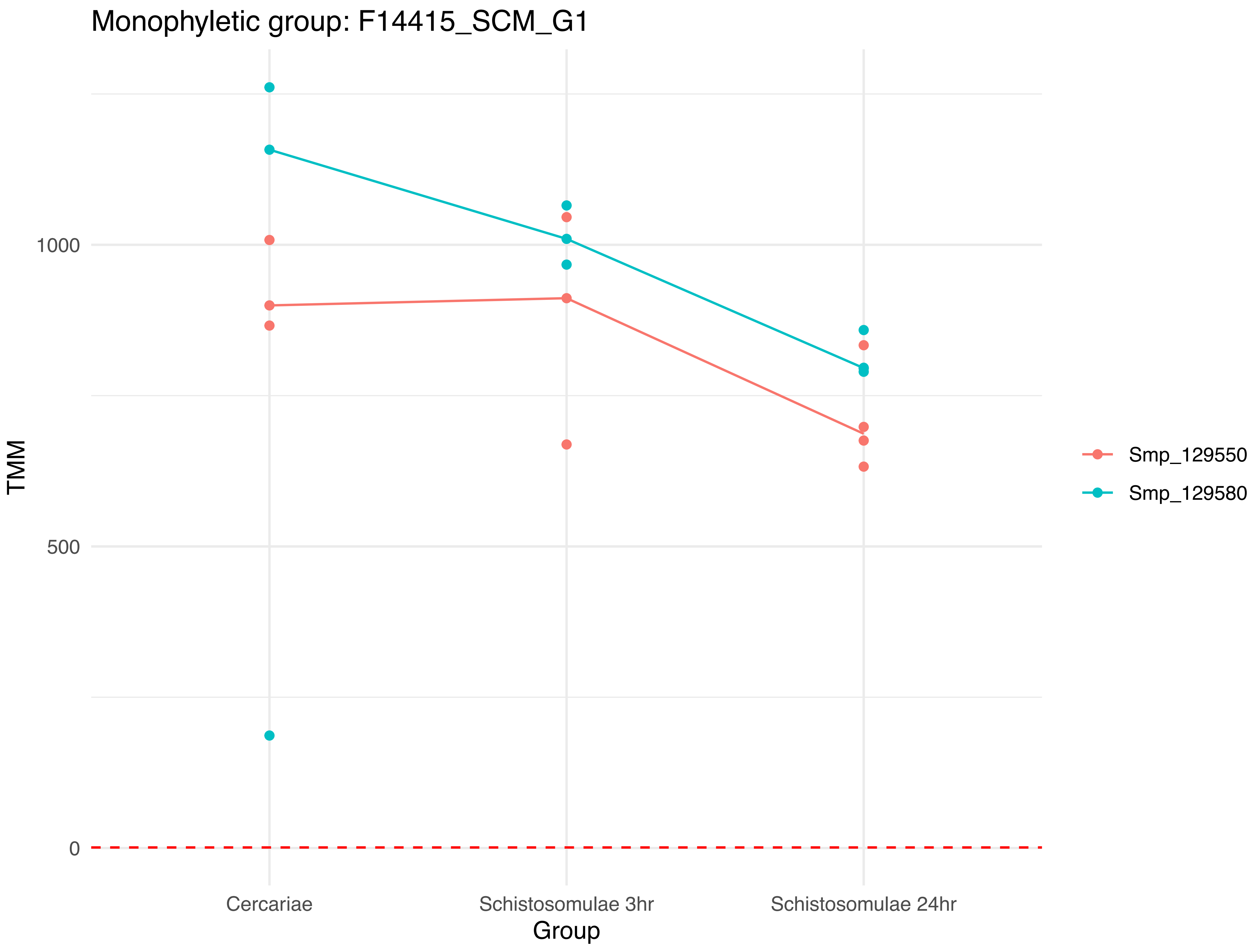

Monophyletic group: F14415_SCM_G1
TMM
1000
500
0
Cercariae
Schistosomulae 3hr
Schistosomulae 24hr
Group
Smp_129550
Smp_129580

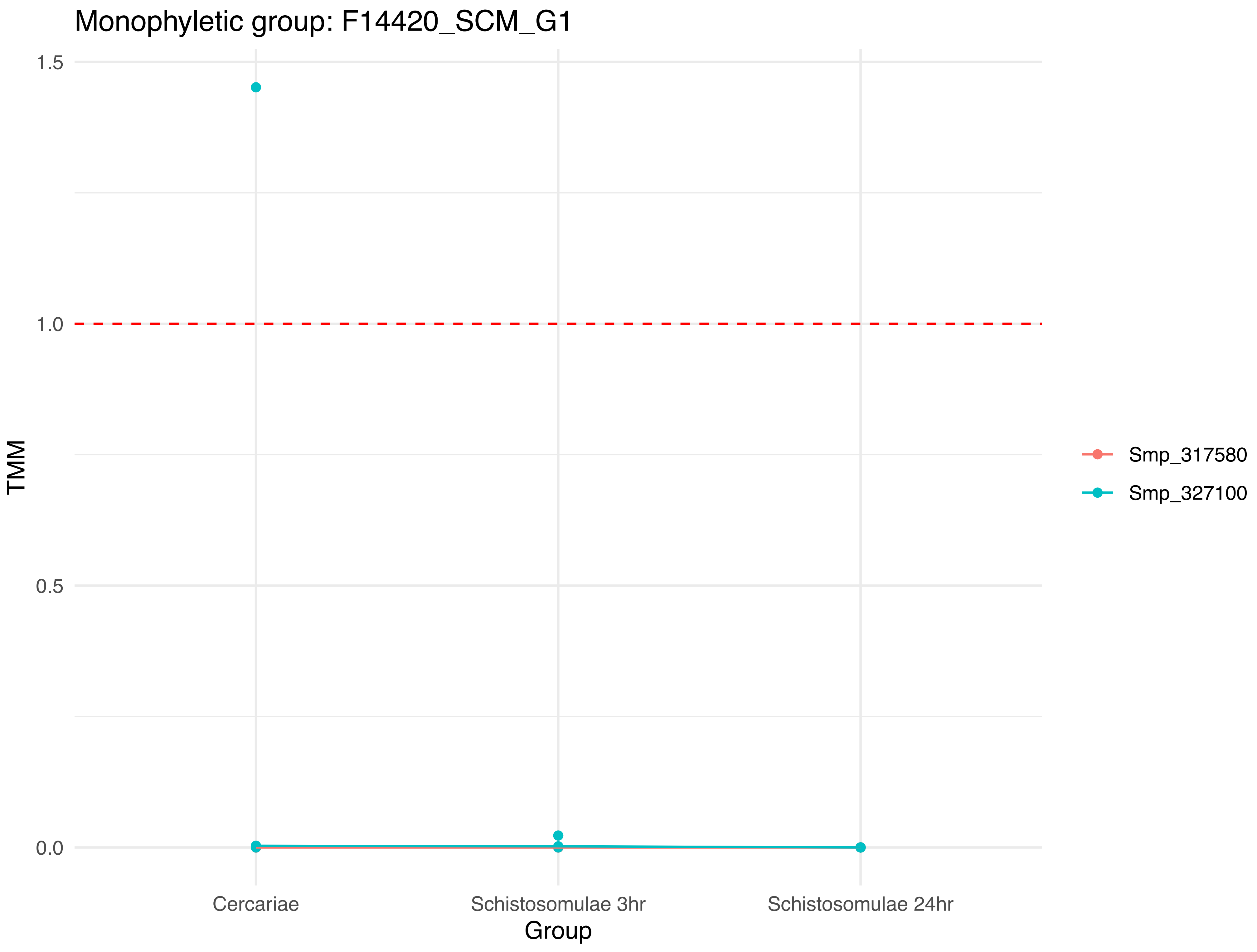
Monophyletic group: F14420_SCM_G1
1.5
1.0
0.5
0.0
TMM
Cercariae
Schistosomulae 3hr
Schistosomulae 24hr
Group
Smp_317580
Smp_327100

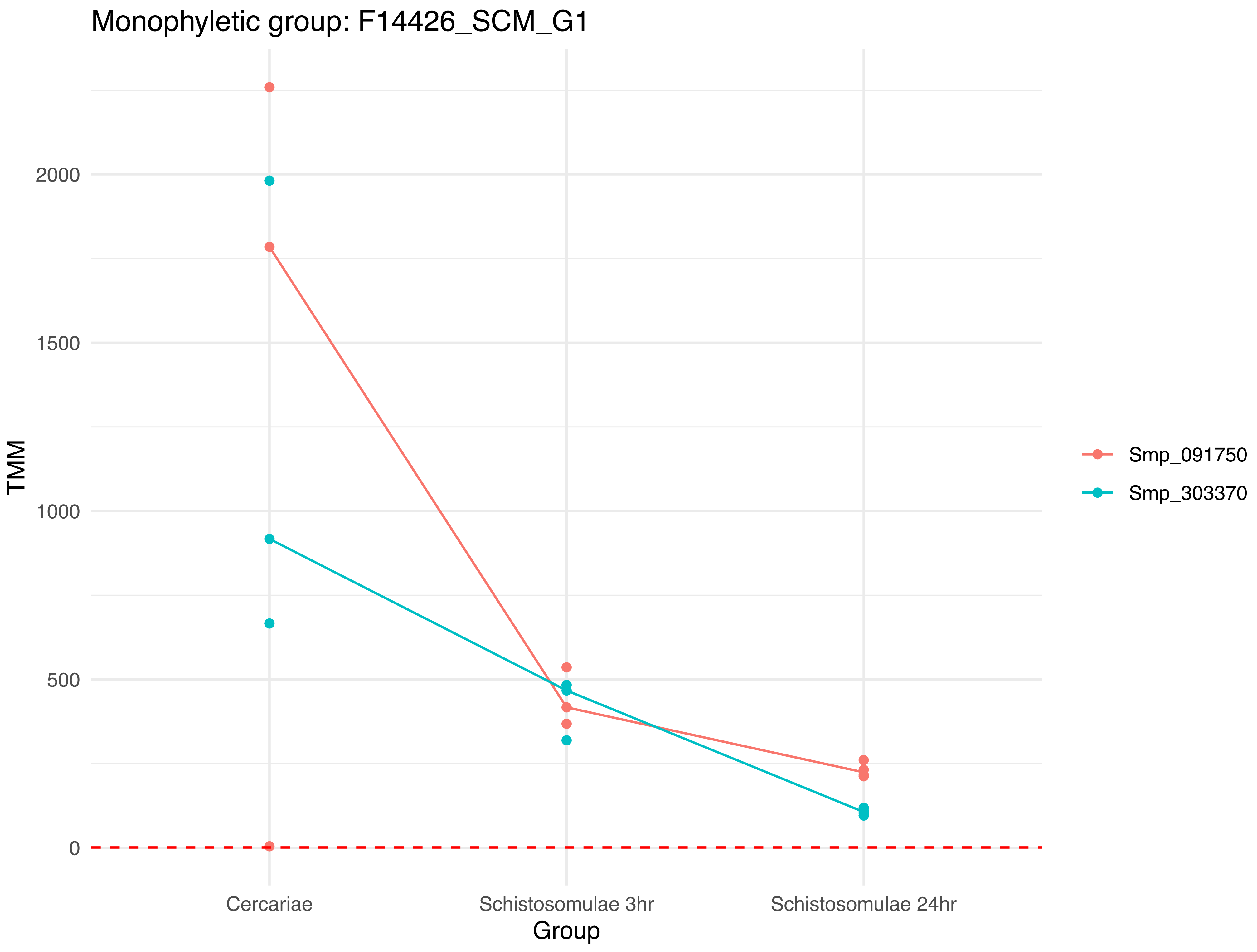

Monophyletic group: F14426_SCM_G1
2000
1500
1000
500
0
TMM
Cercariae
Schistosomulae 3hr
Schistosomulae 24hr
Group
Smp_091750
Smp_303370

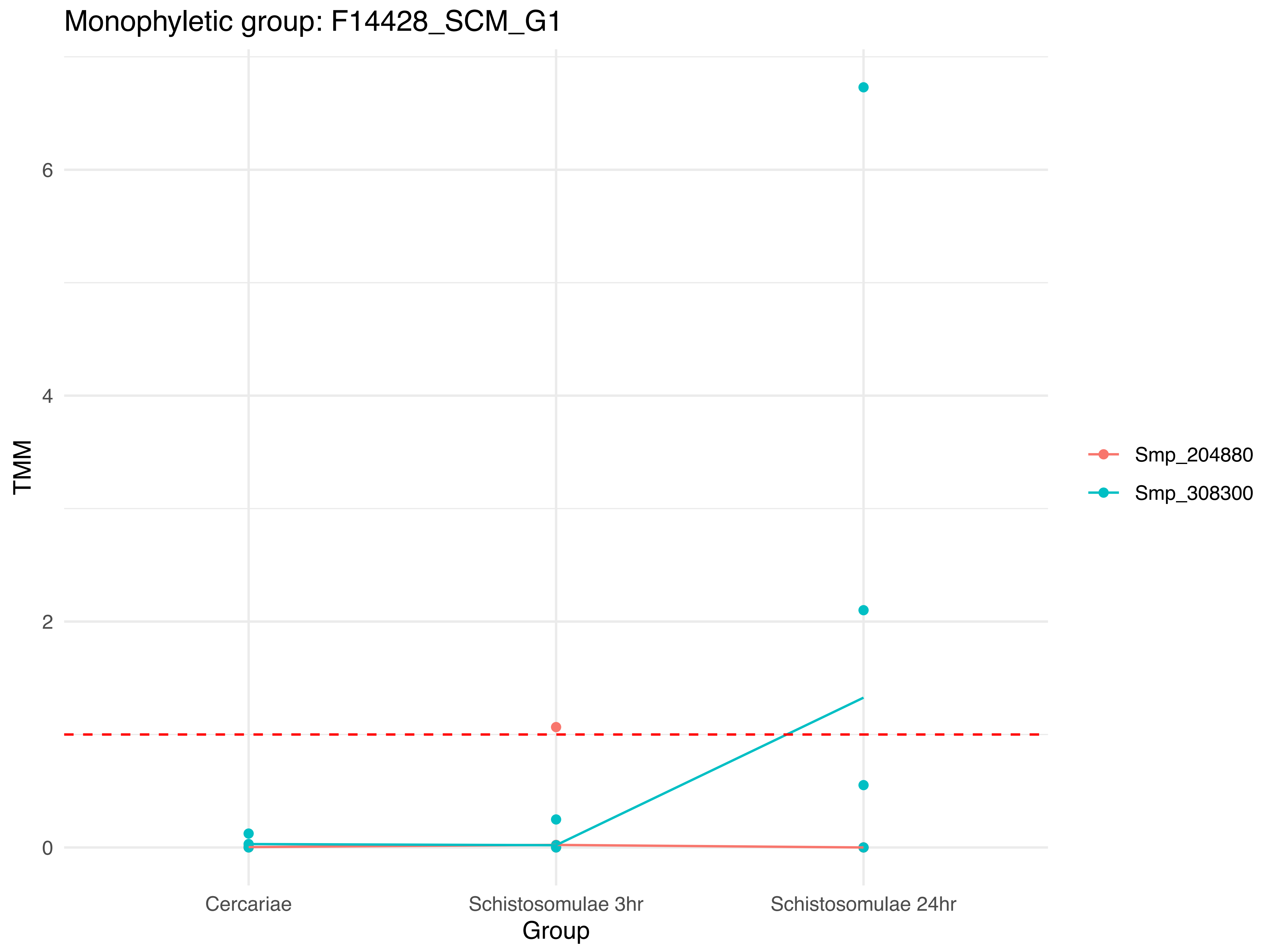

Monophyletic group: F14428_SCM_G1
TMM
6
4
2
0
Cercariae
Schistosomulae 3hr
Schistosomulae 24hr
Group
Smp_204880
Smp_308300

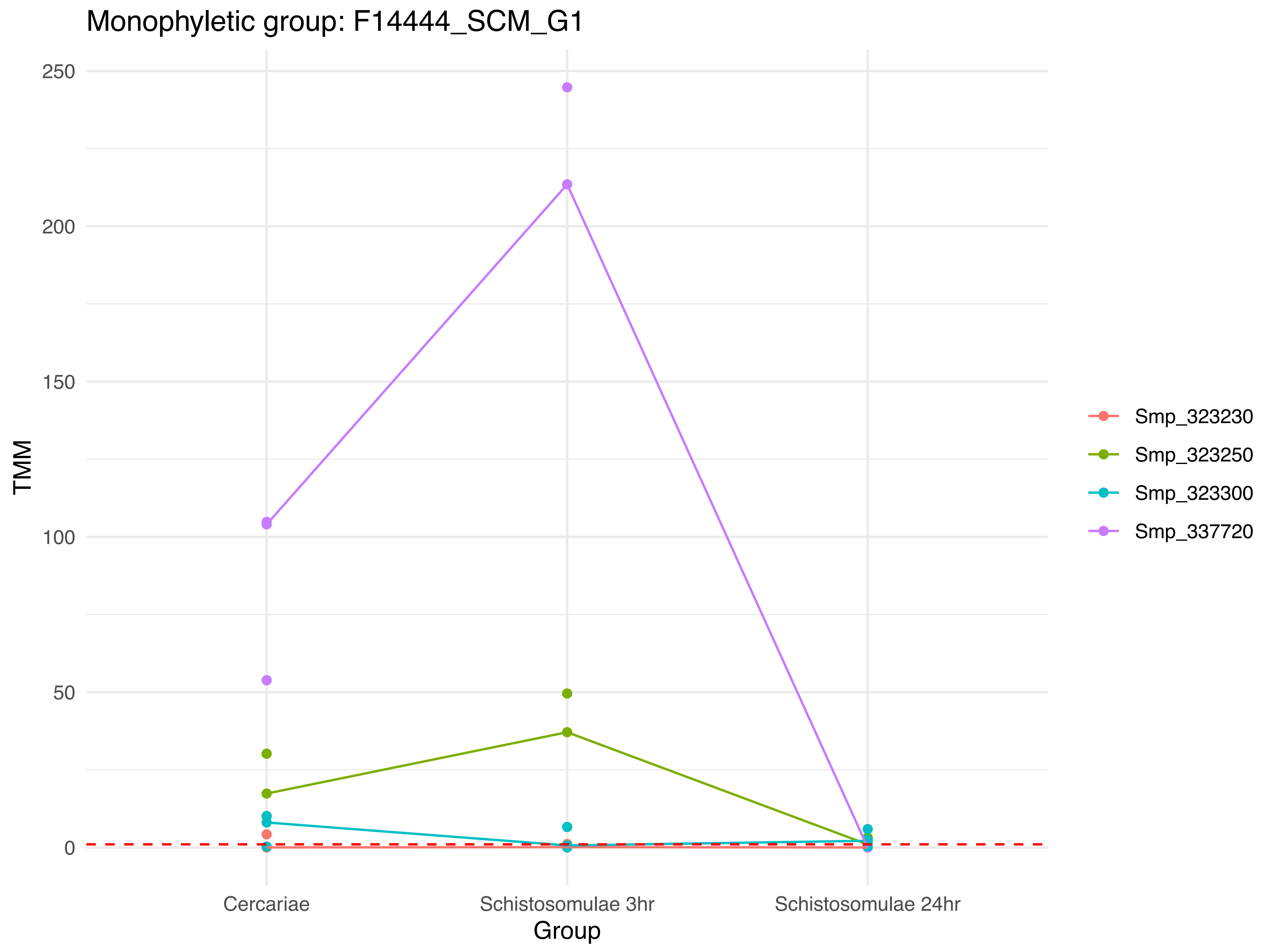

Monophyletic group: F14444_SCM_G1
250
200
150
100
50
0
TMM
Cercariae
Schistosomulae 3hr
Schistosomulae 24hr
Group
Smp_323230
Smp_323250
Smp_323300
Smp_337720

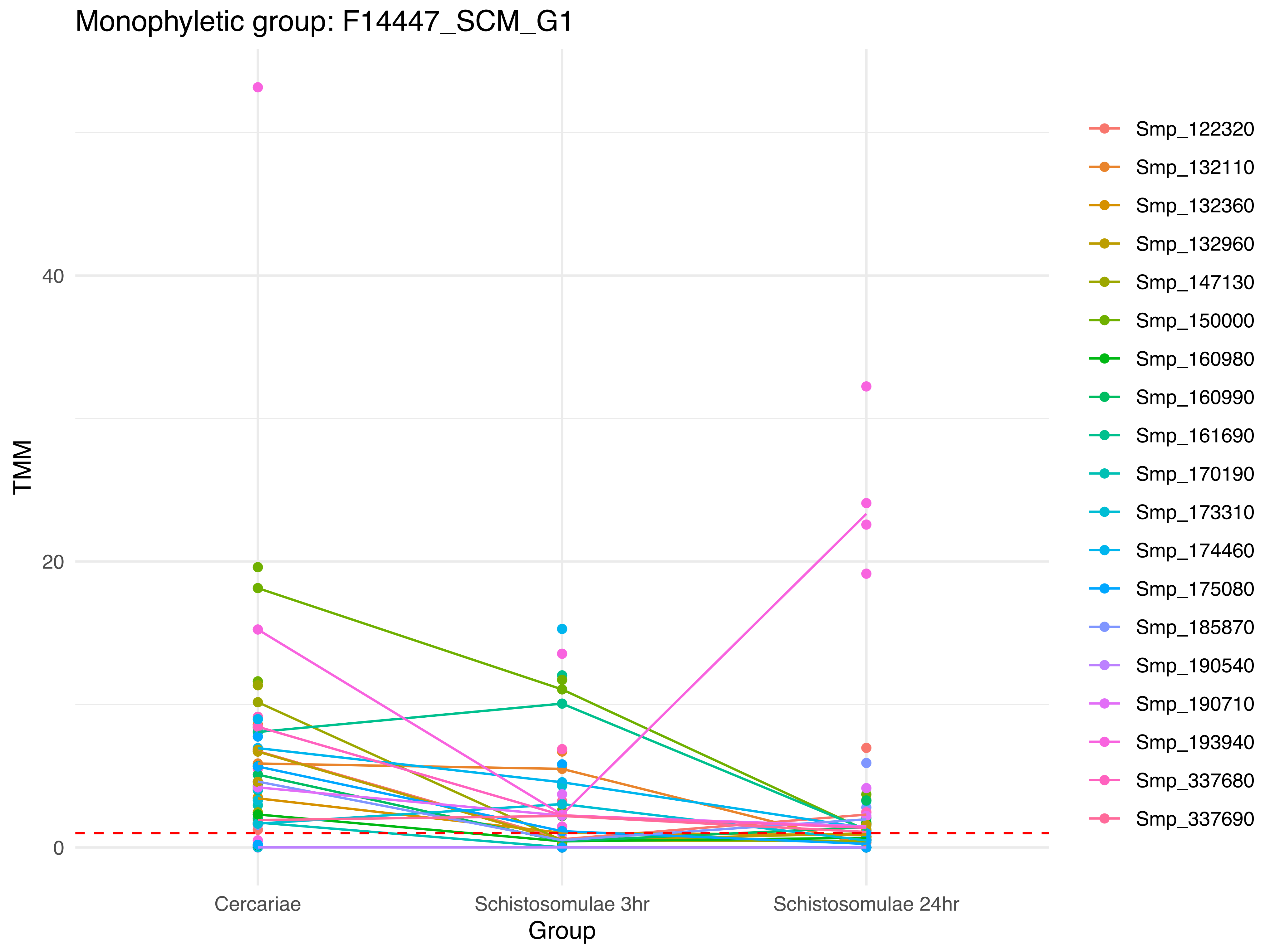

Monophyletic group: F14447_SCM_G1
TMM
0
20
40
Group
Cercariae
Schistosomulae 3hr
Schistosomulae 24hr
Smp_122320
Smp_132110
Smp_132360
Smp_132960
Smp_147130
Smp_150000
Smp_160980
Smp_160990
Smp_161690
Smp_170190
Smp_173310
Smp_174460
Smp_175080
Smp_185870
Smp_190540
Smp_190710
Smp_193940
Smp_337680
Smp_337690

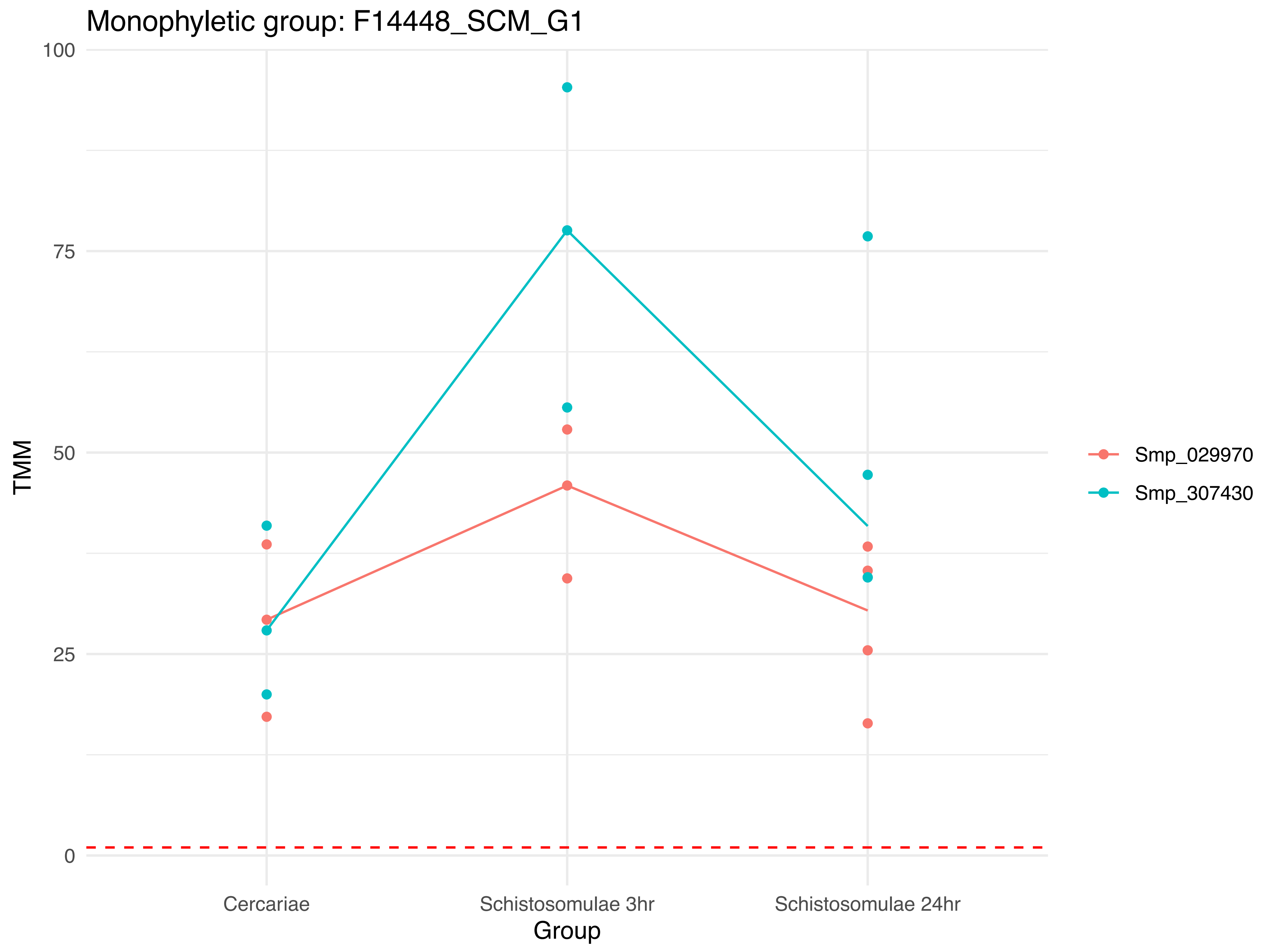

Monophyletic group: F14448_SCM_G1
100
75
50
25
0
TMM
Cercariae
Schistosomulae 3hr
Schistosomulae 24hr
Group
Smp_029970
Smp_307430

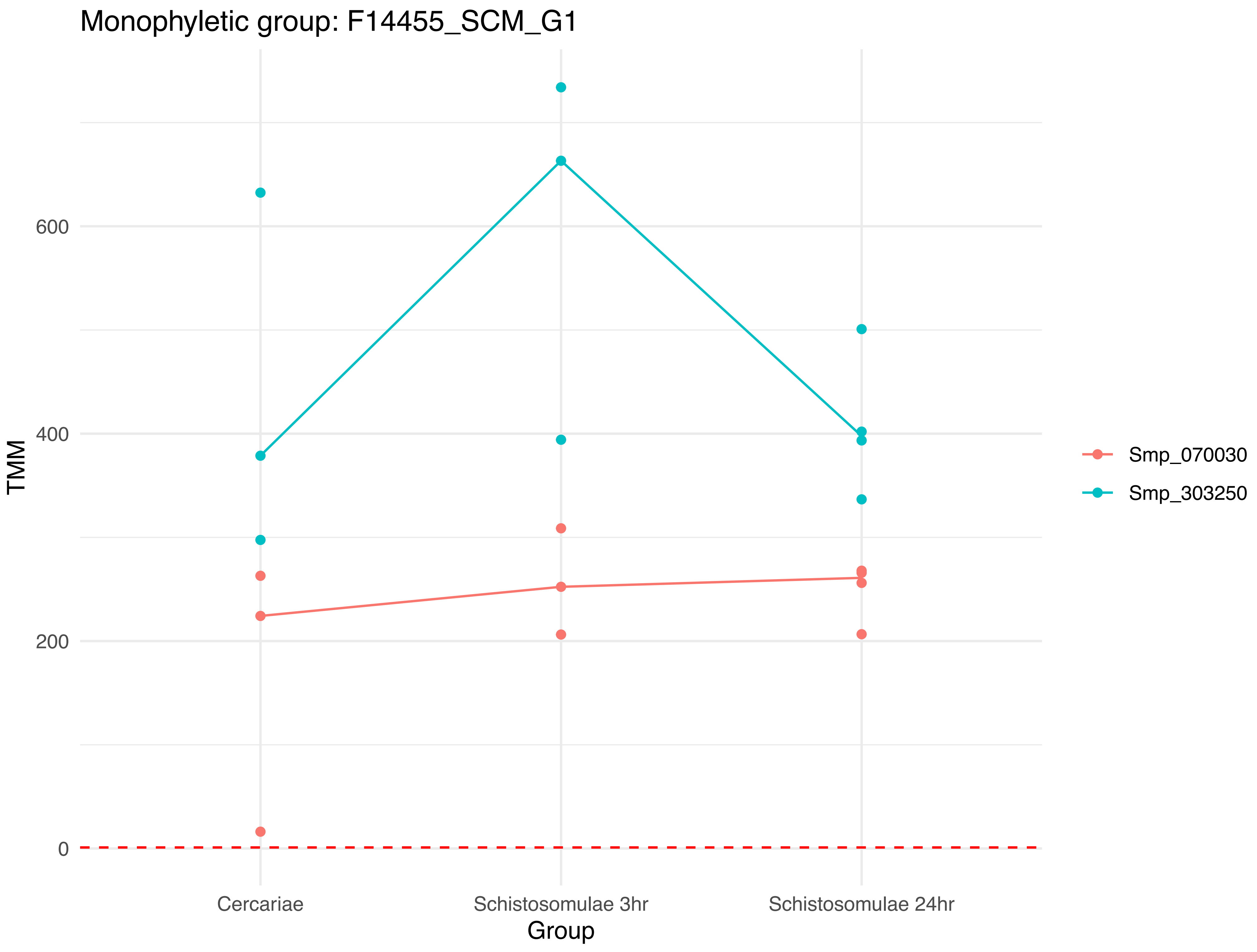

Monophyletic group: F14455_SCM_G1
TMM
600
400
200
0
Cercariae
Schistosomulae 3hr
Schistosomulae 24hr
Group
Smp_070030
Smp_303250

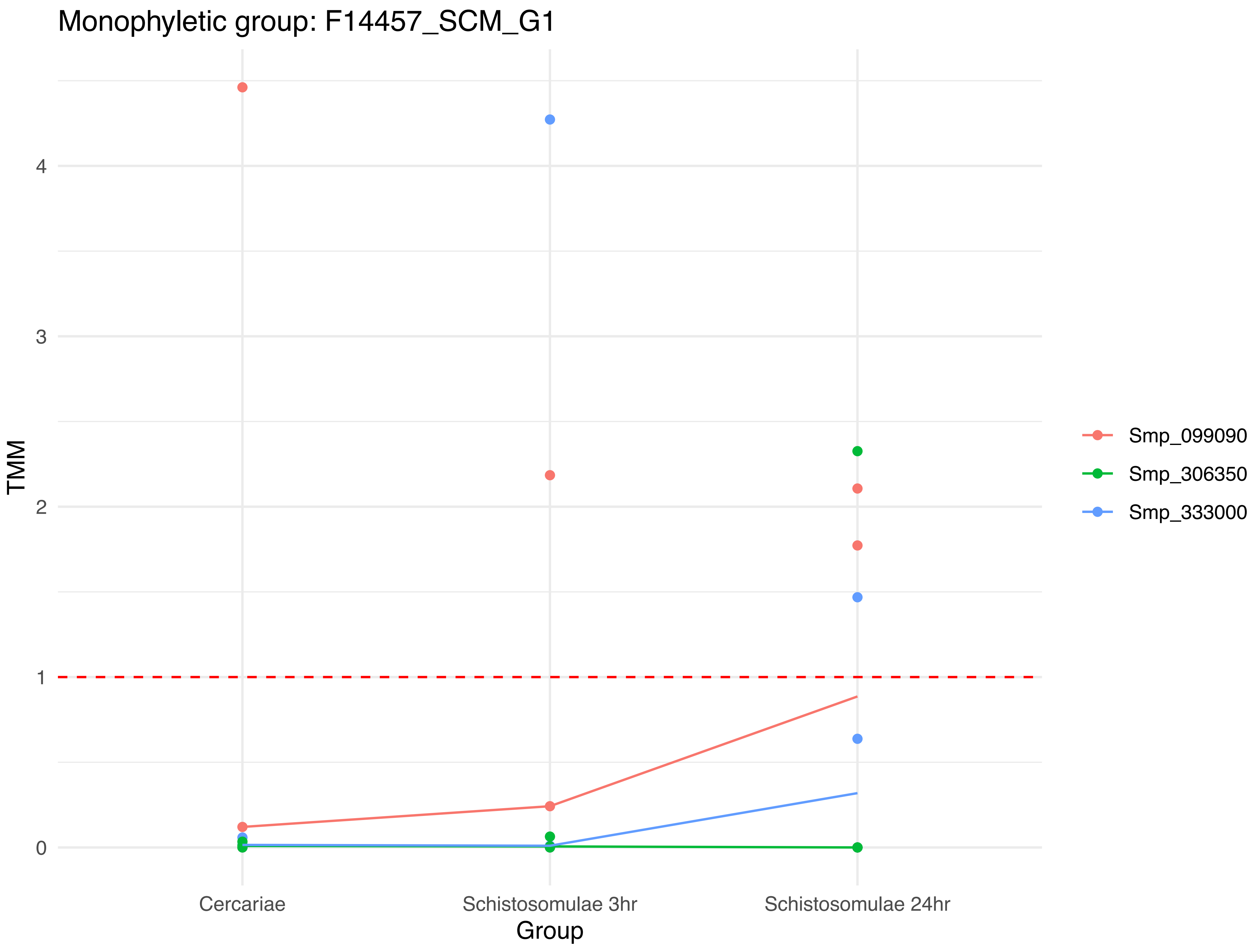
Monophyletic group: F14457_SCM_G1
TMM
0
1
2
3
4
Cercariae
Schistosomulae 3hr
Schistosomulae 24hr
Group
Smp_099090
Smp_306350
Smp_333000

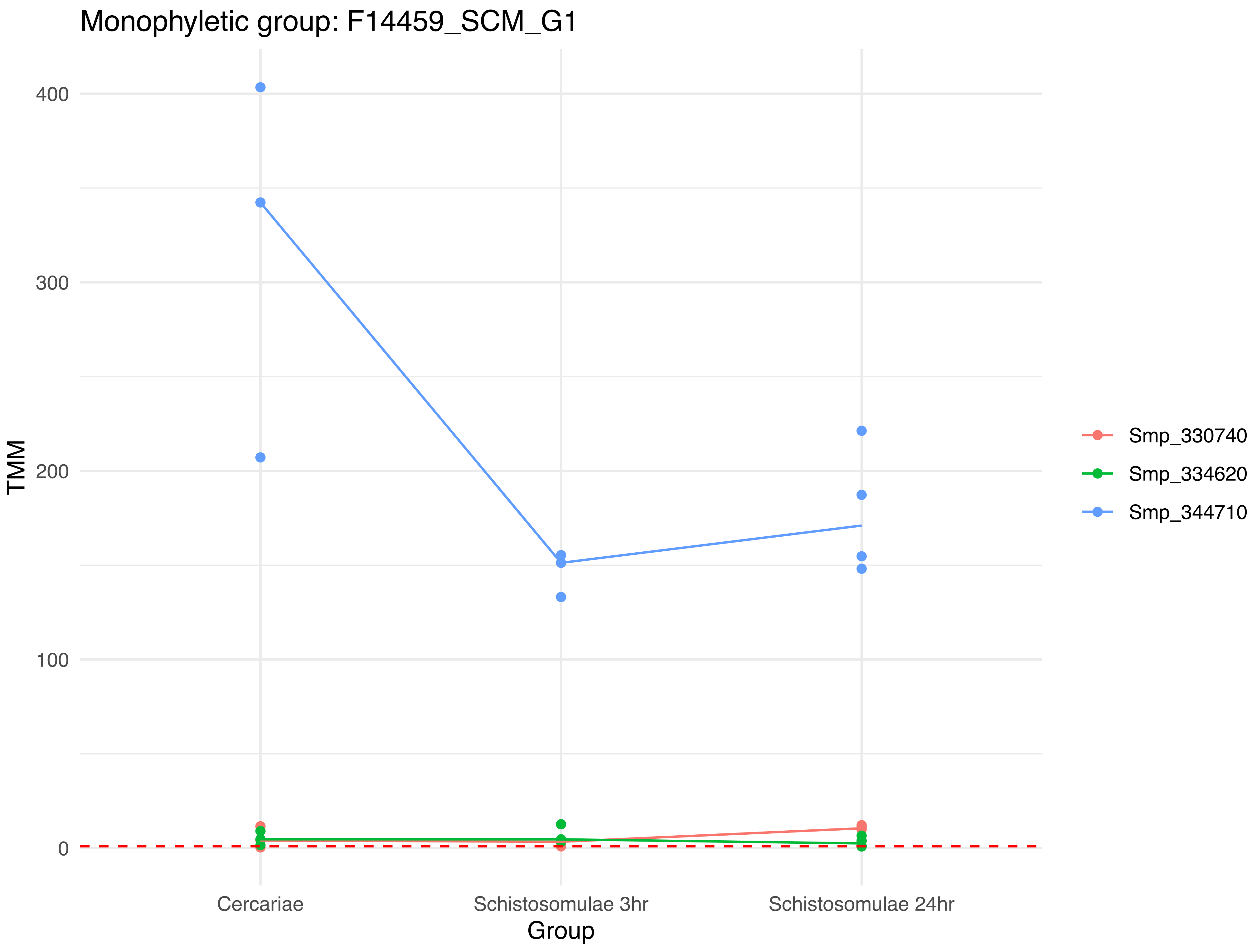

Monophyletic group: F14459_SCM_G1
TMM
400
300
200
100
0
Cercariae
Schistosomulae 3hr
Schistosomulae 24hr
Group
Smp_330740
Smp_334620
Smp_344710

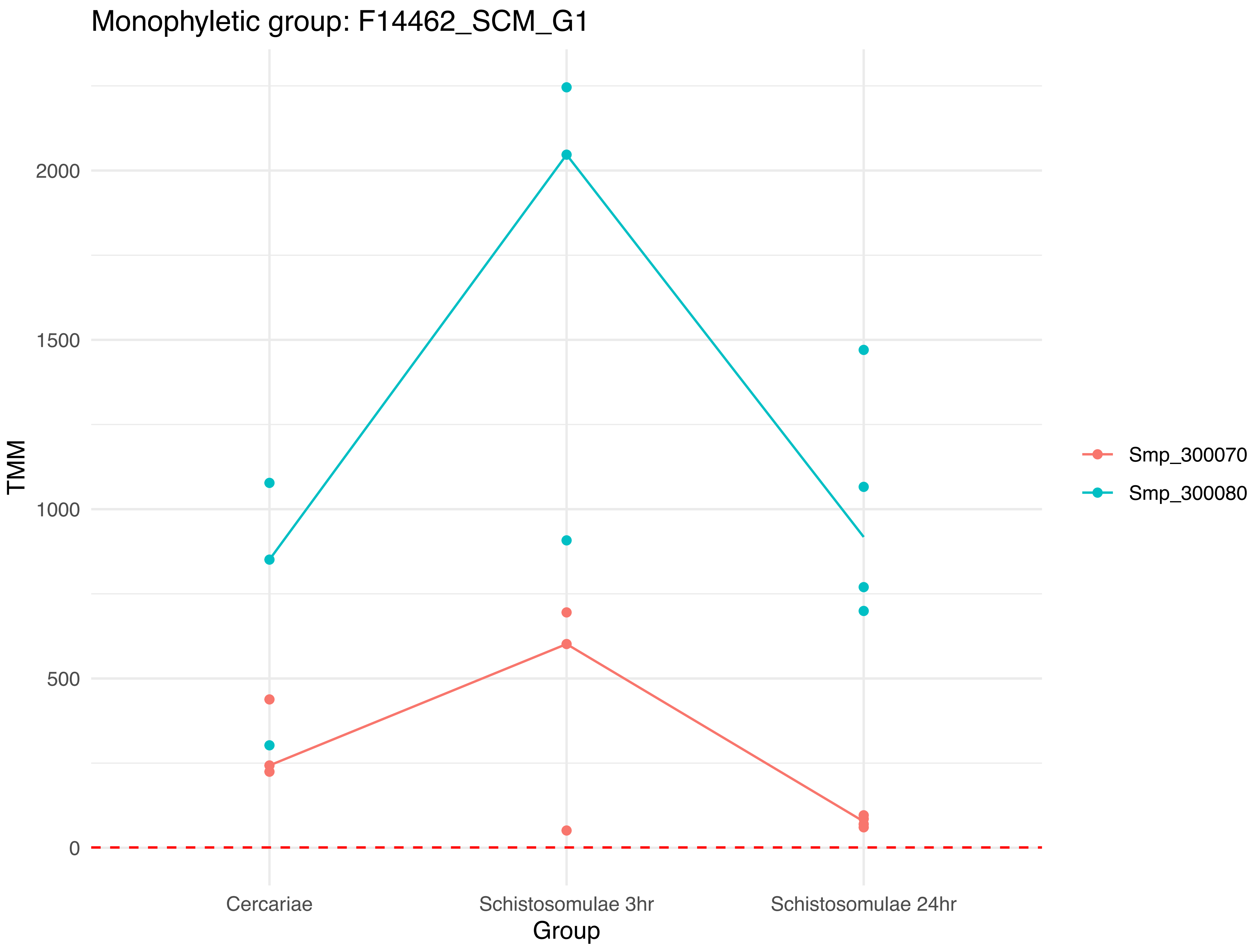
Monophyletic group: F14462_SCM_G1
TMM
0
500
1000
1500
2000
Cercariae
Schistosomulae 3hr
Schistosomulae 24hr
Group
Smp_300070
Smp_300080

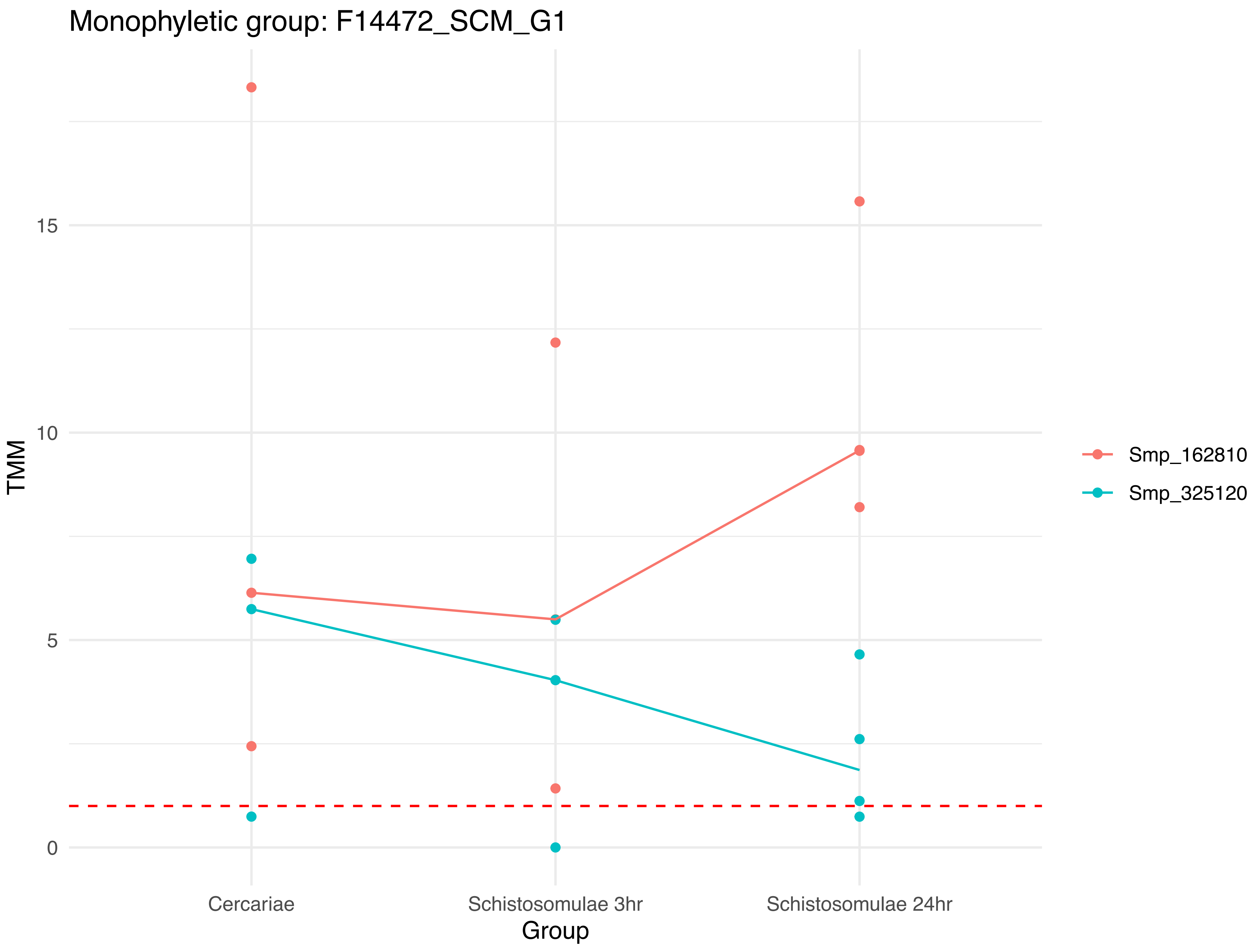
Monophyletic group: F14472_SCM_G1
TMM
15
10
5
0
Cercariae
Schistosomulae 3hr
Schistosomulae 24hr
Group
Smp_162810
Smp_325120

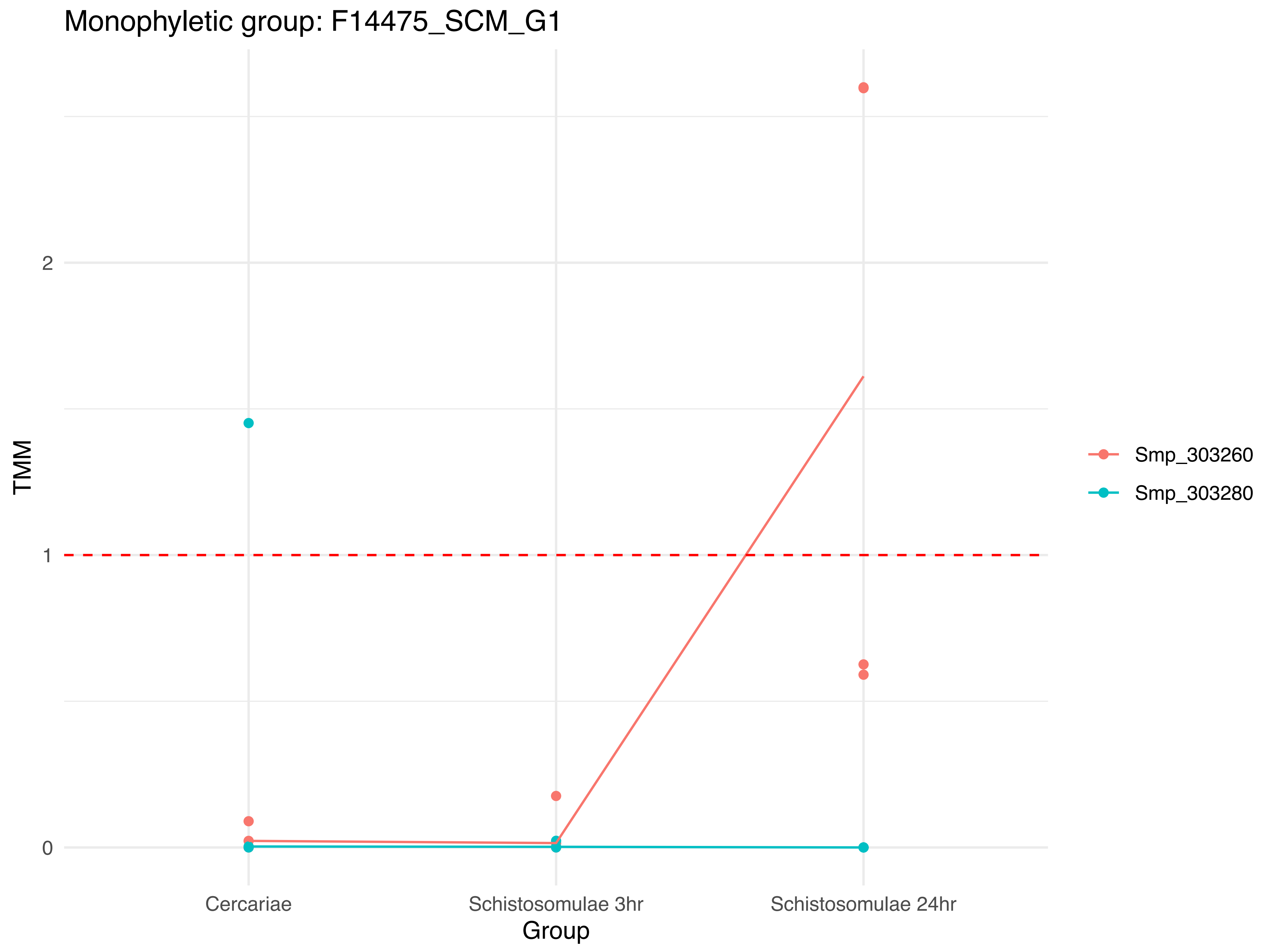
Monophyletic group: F14475_SCM_G1
TMM
2
1
0
Cercariae
Schistosomulae 3hr
Schistosomulae 24hr
Group
Smp_303260
Smp_303280

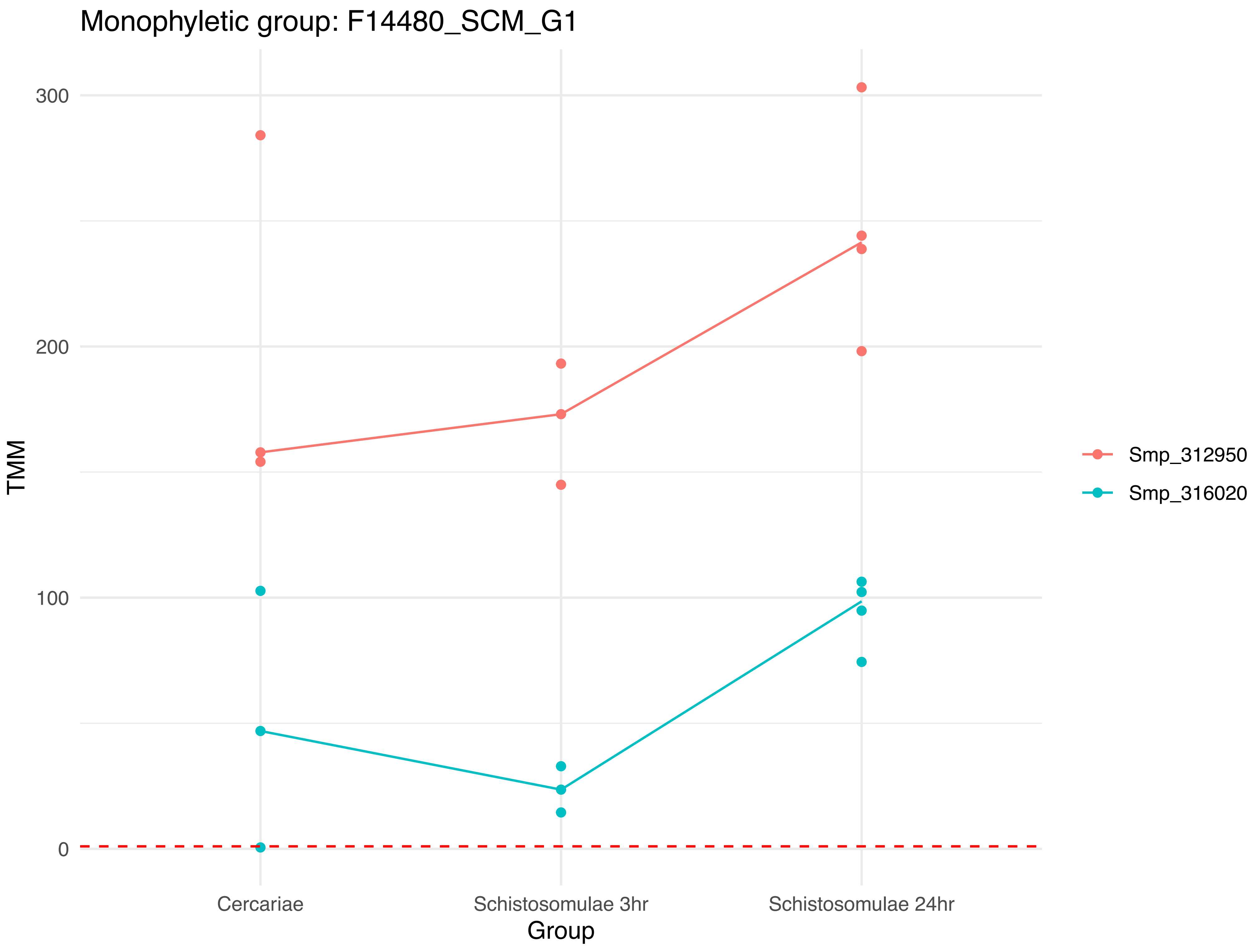

Monophyletic group: F14480_SCM_G1
300
200
100
0
TMM
Cercariae
Schistosomulae 3hr
Schistosomulae 24hr
Group
Smp_312950
Smp_316020

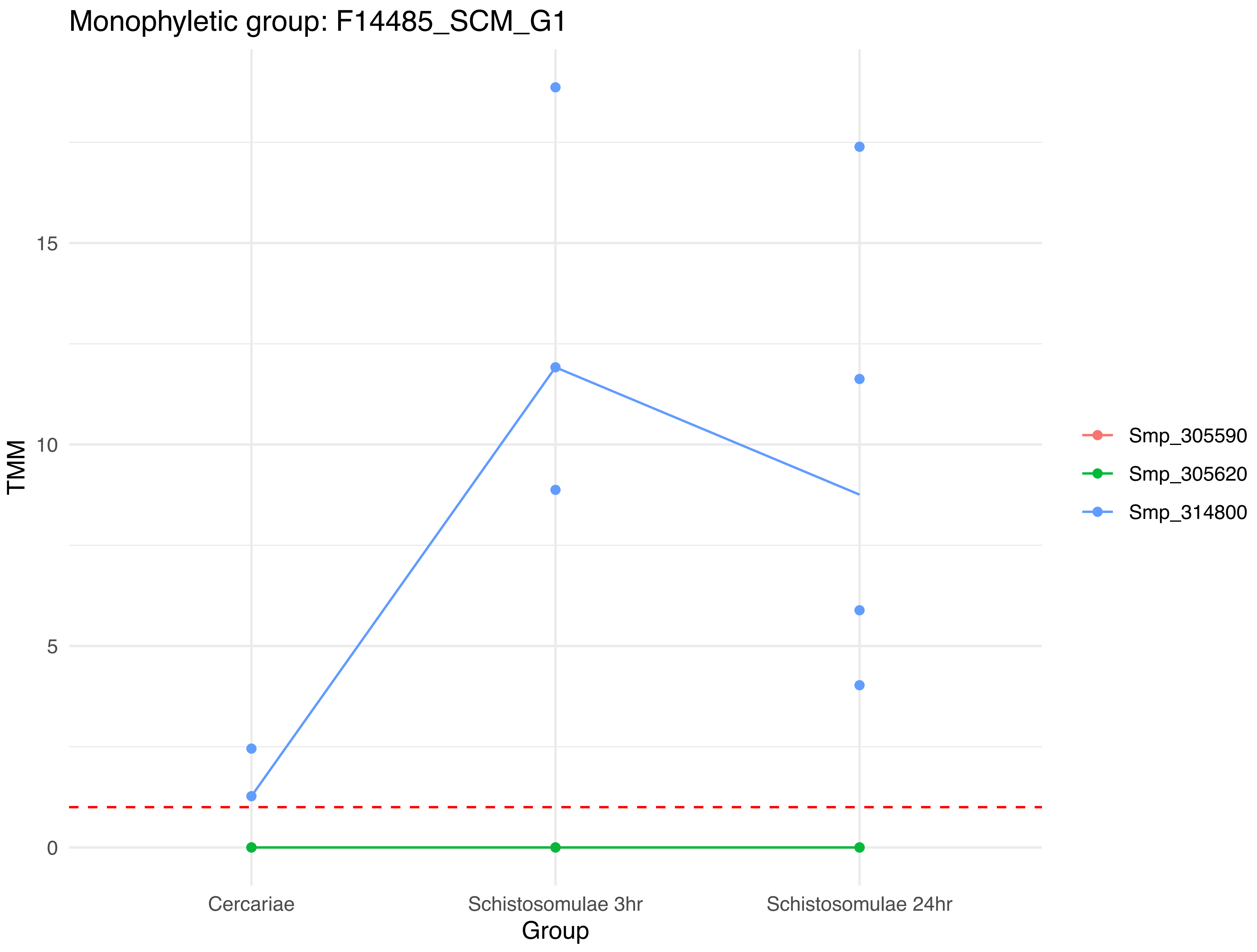
Monophyletic group: F14485_SCM_G1
TMM
15
10
5
0
Cercariae
Schistosomulae 3hr
Schistosomulae 24hr
Group
Smp_305590
Smp_305620
Smp_314800

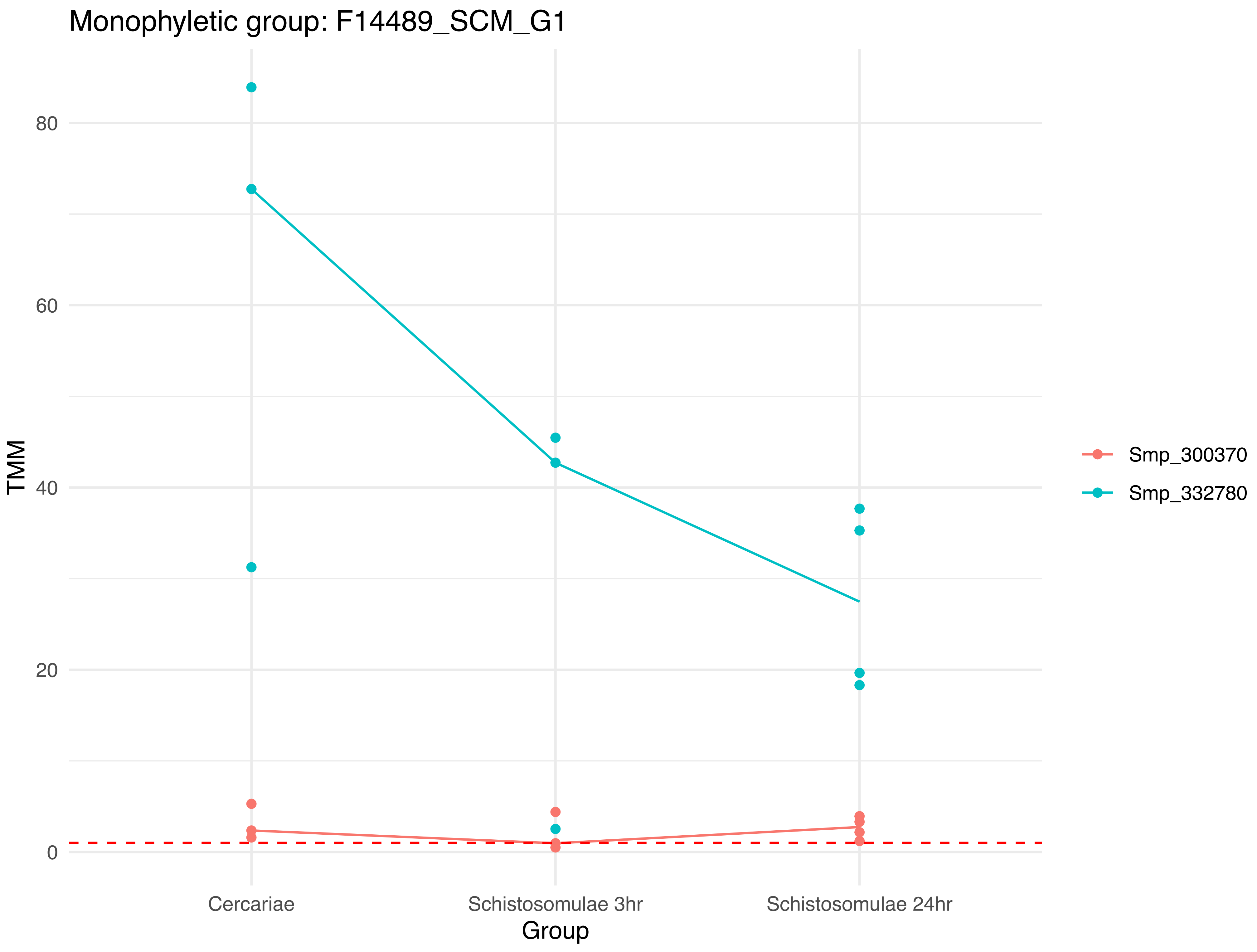

Monophyletic group: F14489_SCM_G1
80
60
40
20
0
TMM
Cercariae
Schistosomulae 3hr
Schistosomulae 24hr
Group
Smp_300370
Smp_332780

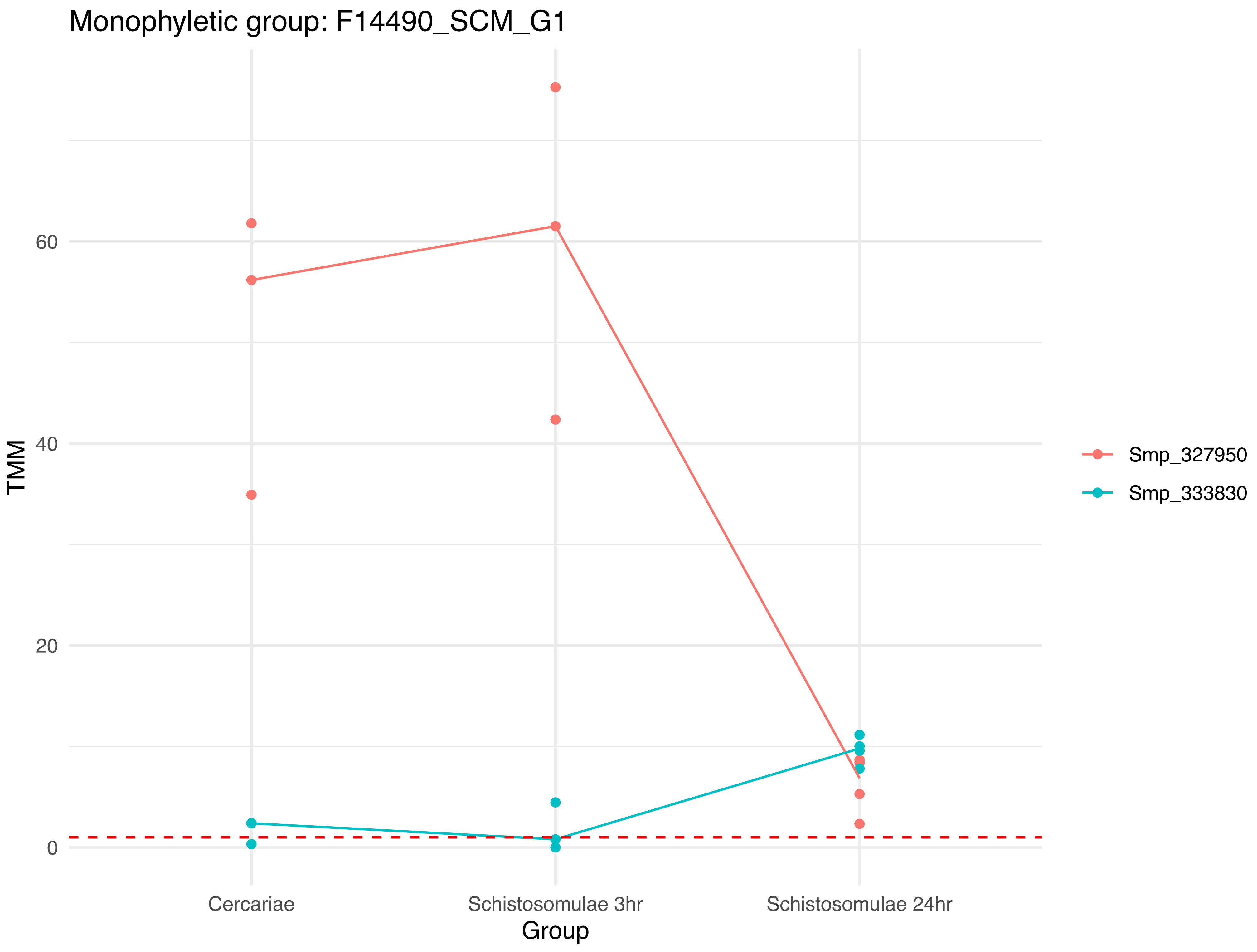
Monophyletic group: F14490_SCM_G1
TMM
60
40
20
0
Cercariae
Schistosomulae 3hr
Schistosomulae 24hr
Group
Smp_327950
Smp_333830

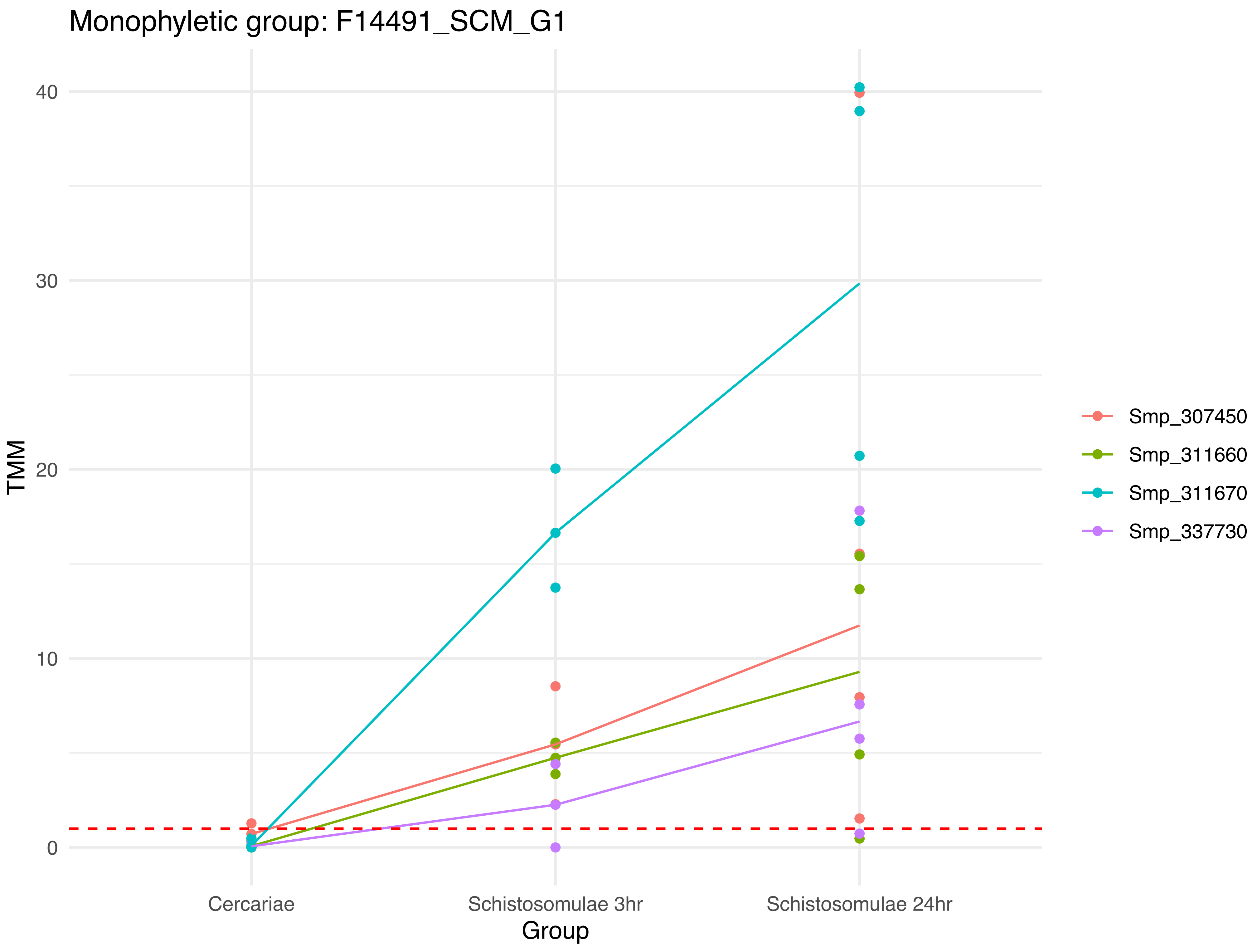

Monophyletic group: F14491_SCM_G1
TMM
40
30
20
10
0
Cercariae
Schistosomulae 3hr
Schistosomulae 24hr
Group
Smp_307450
Smp_311660
Smp_311670
Smp_337730

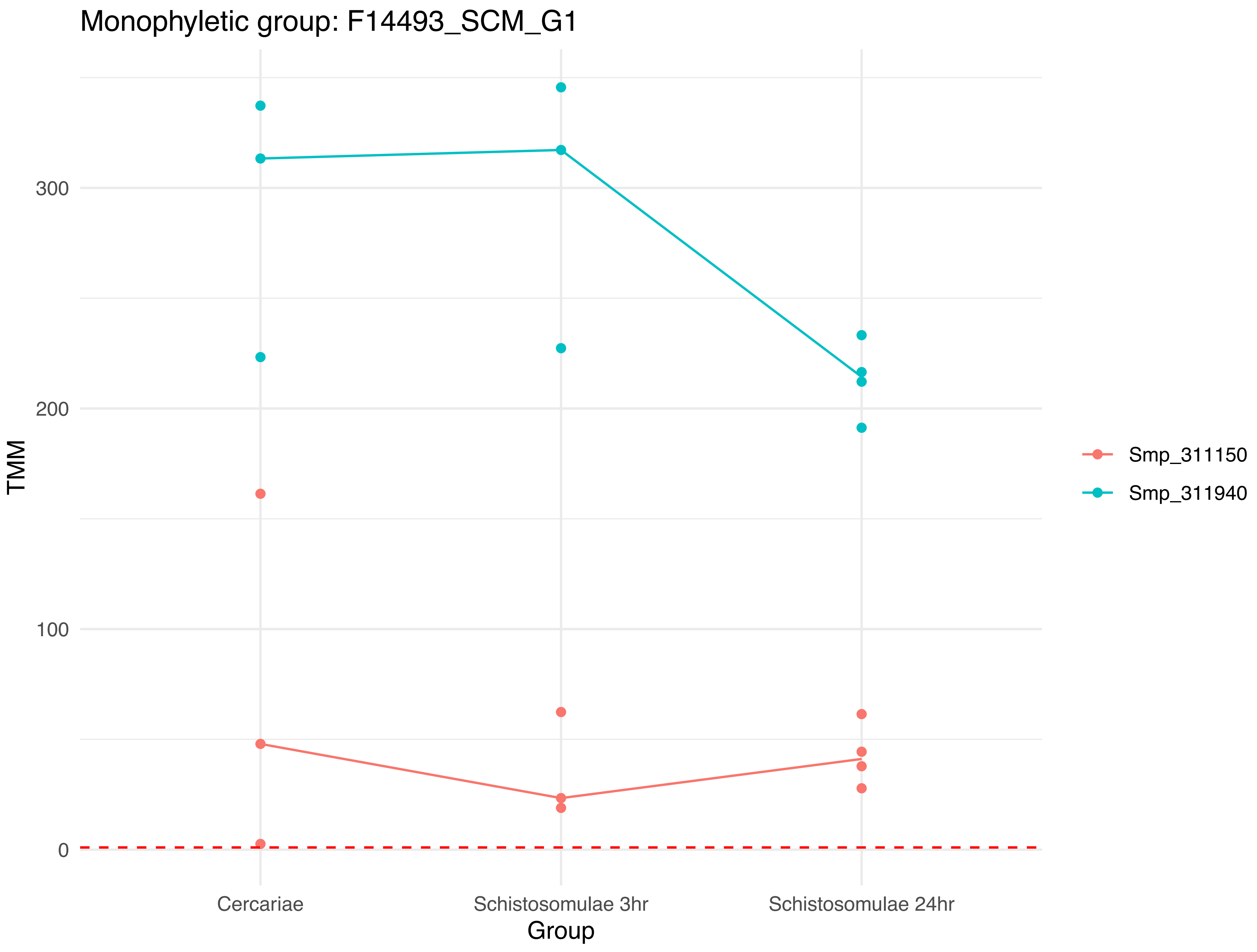

Monophyletic group: F14493_SCM_G1
TMM
300
200
100
0
Cercariae
Schistosomulae 3hr
Schistosomulae 24hr
Group
Smp_311150
Smp_311940

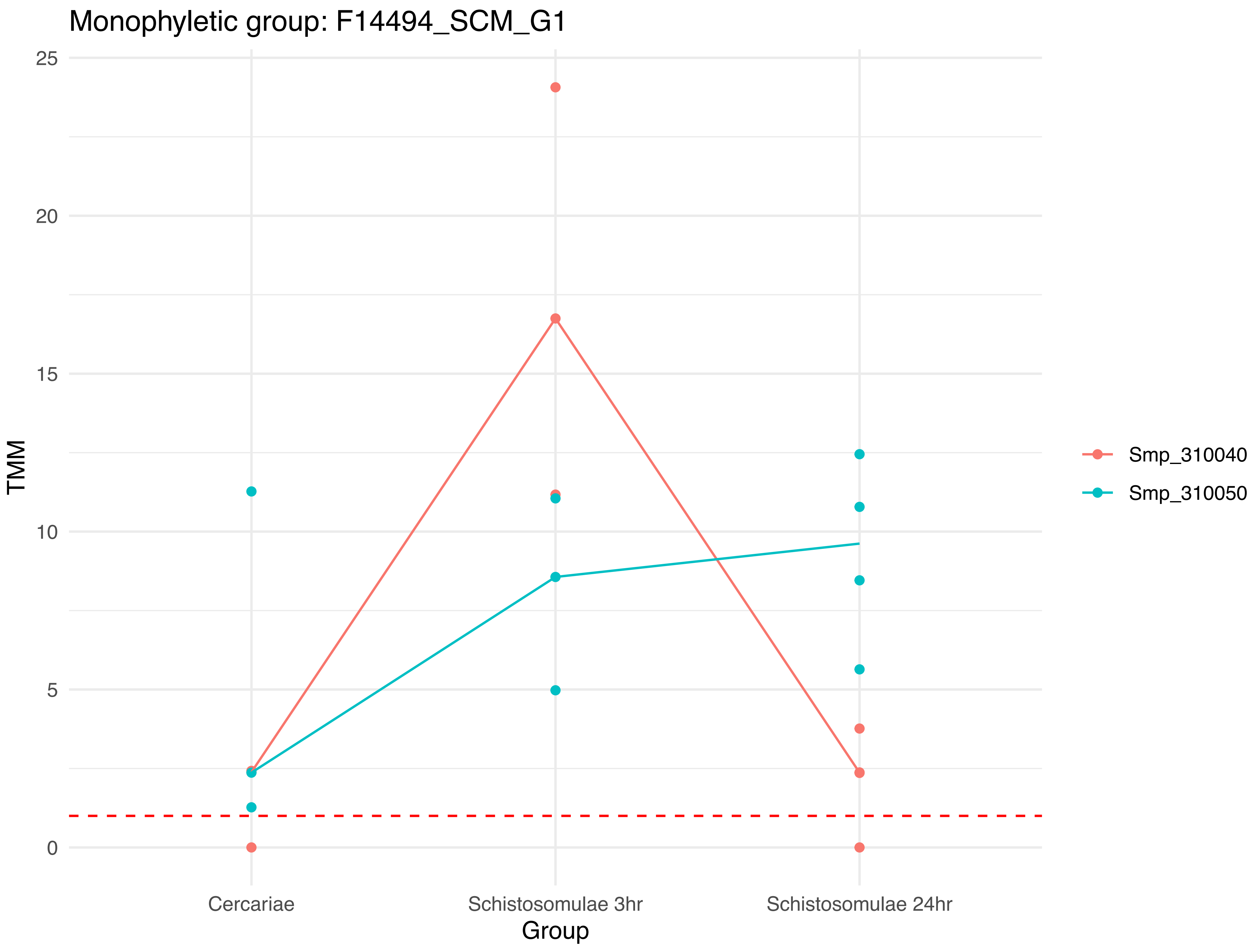
Monophyletic group: F14494_SCM_G1
25
20
15
10
5
0
TMM
Cercariae
Schistosomulae 3hr
Schistosomulae 24hr
Group
Smp_310040
Smp_310050

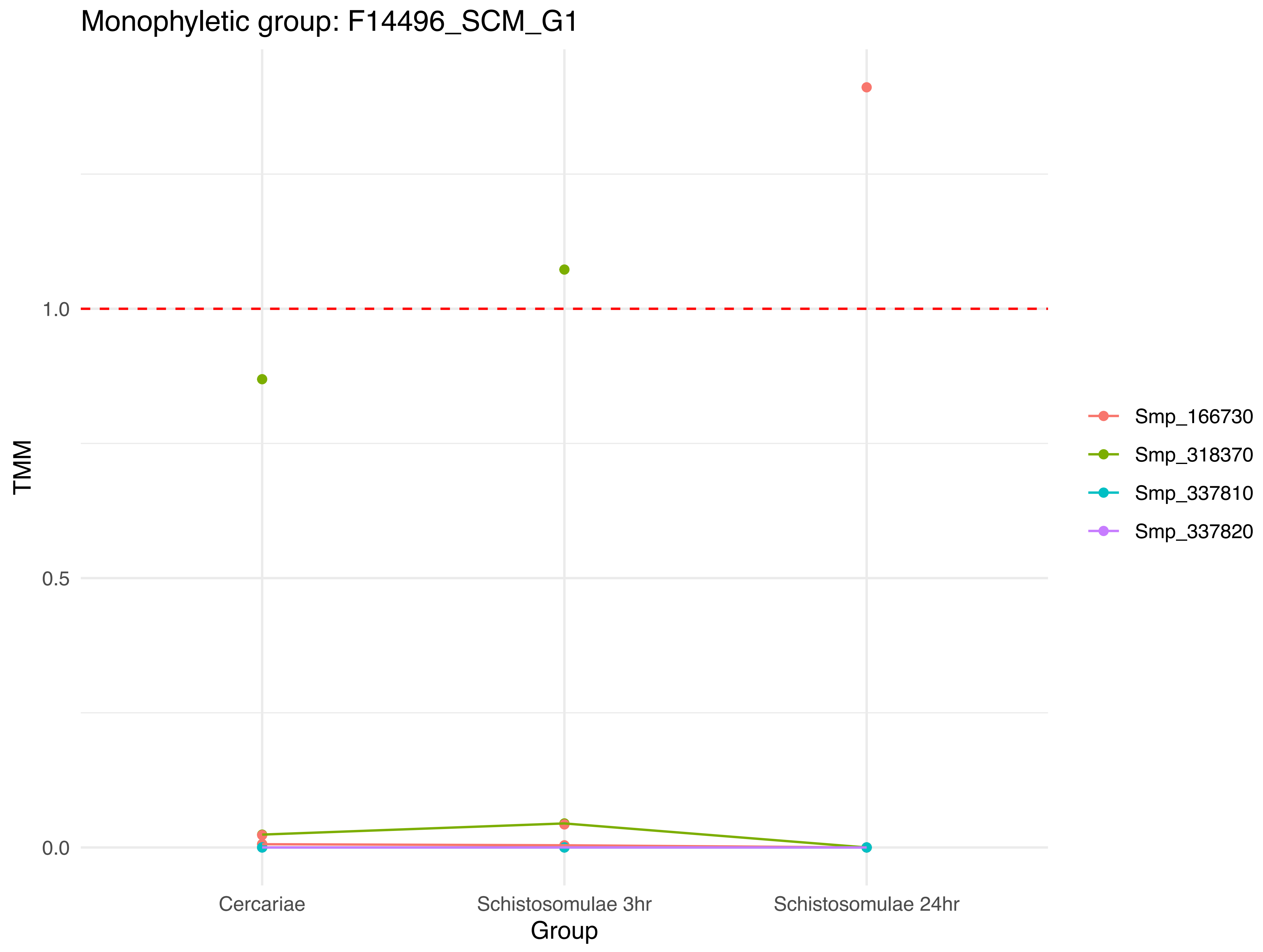
Monophyletic group: F14496_SCM_G1
TMM
1.0
0.5
0.0
Cercariae
Schistosomulae 3hr
Schistosomulae 24hr
Group
Smp_166730
Smp_318370
Smp_337810
Smp_337820

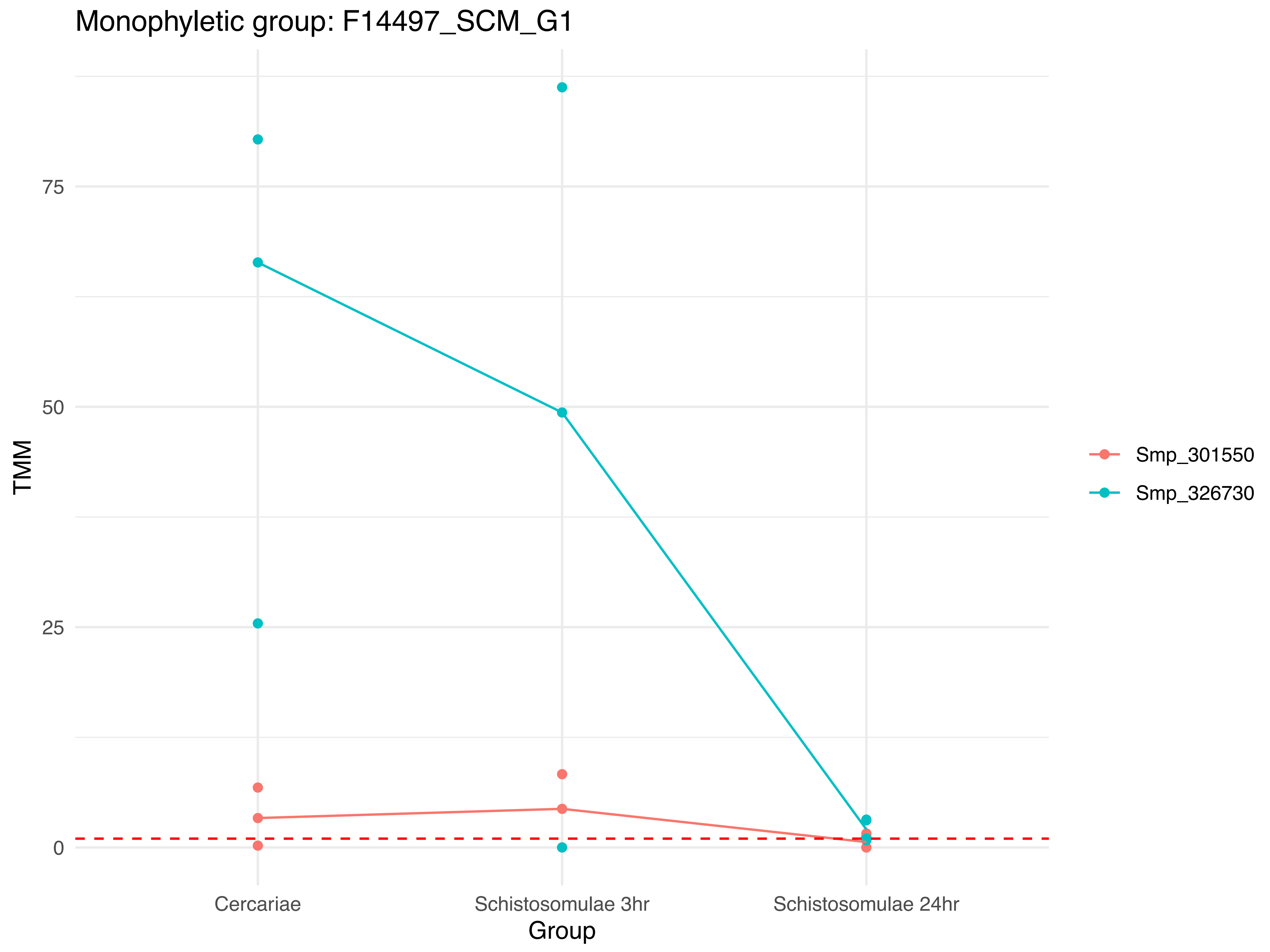

Monophyletic group: F14497_SCM_G1
75
50
25
0
TMM
Cercariae
Schistosomulae 3hr
Schistosomulae 24hr
Group
Smp_301550
Smp_326730

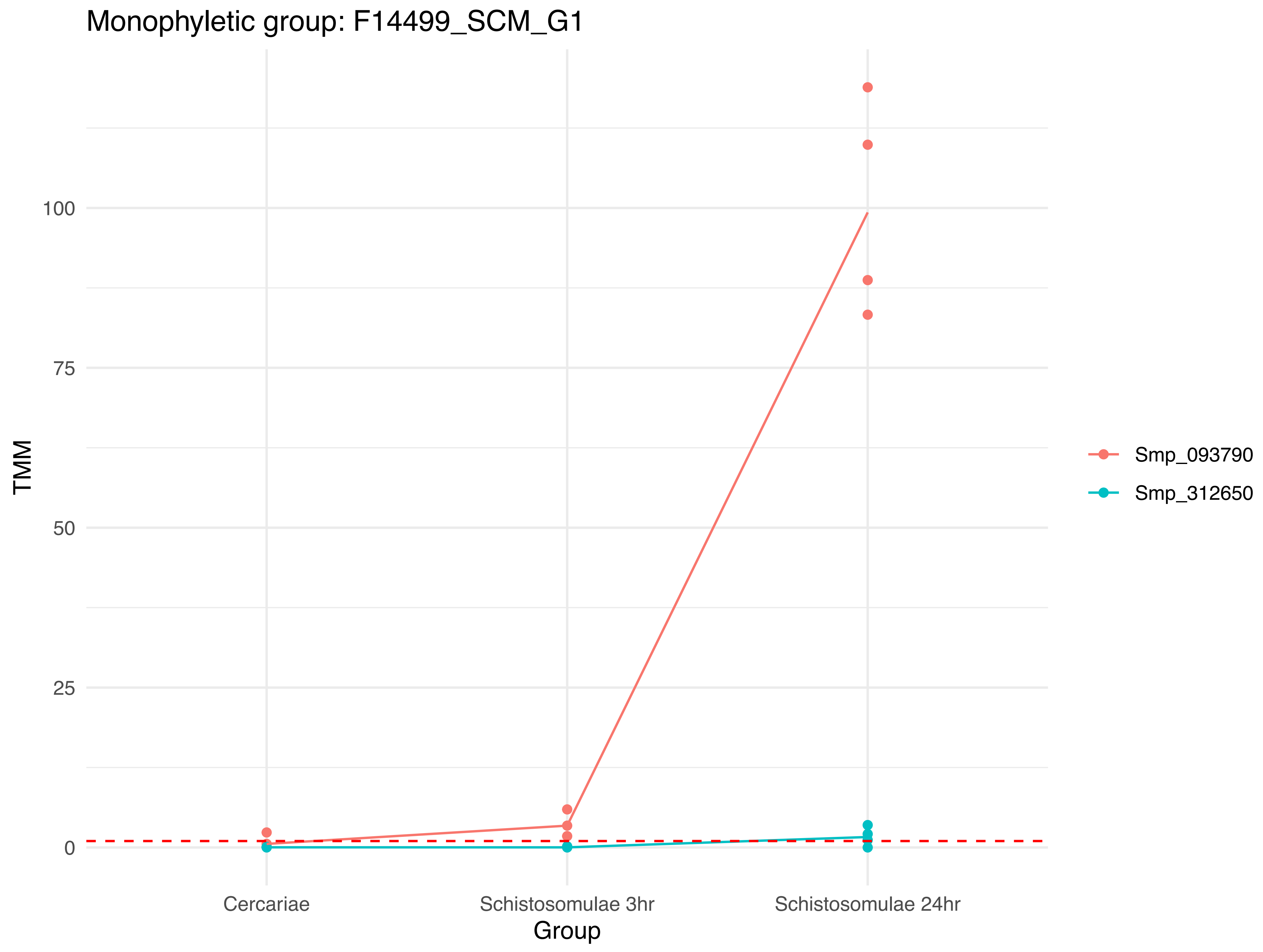

Monophyletic group: F14499_SCM_G1
TMM
100
75
50
25
0
Cercariae
Schistosomulae 3hr
Schistosomulae 24hr
Group
Smp_093790
Smp_312650

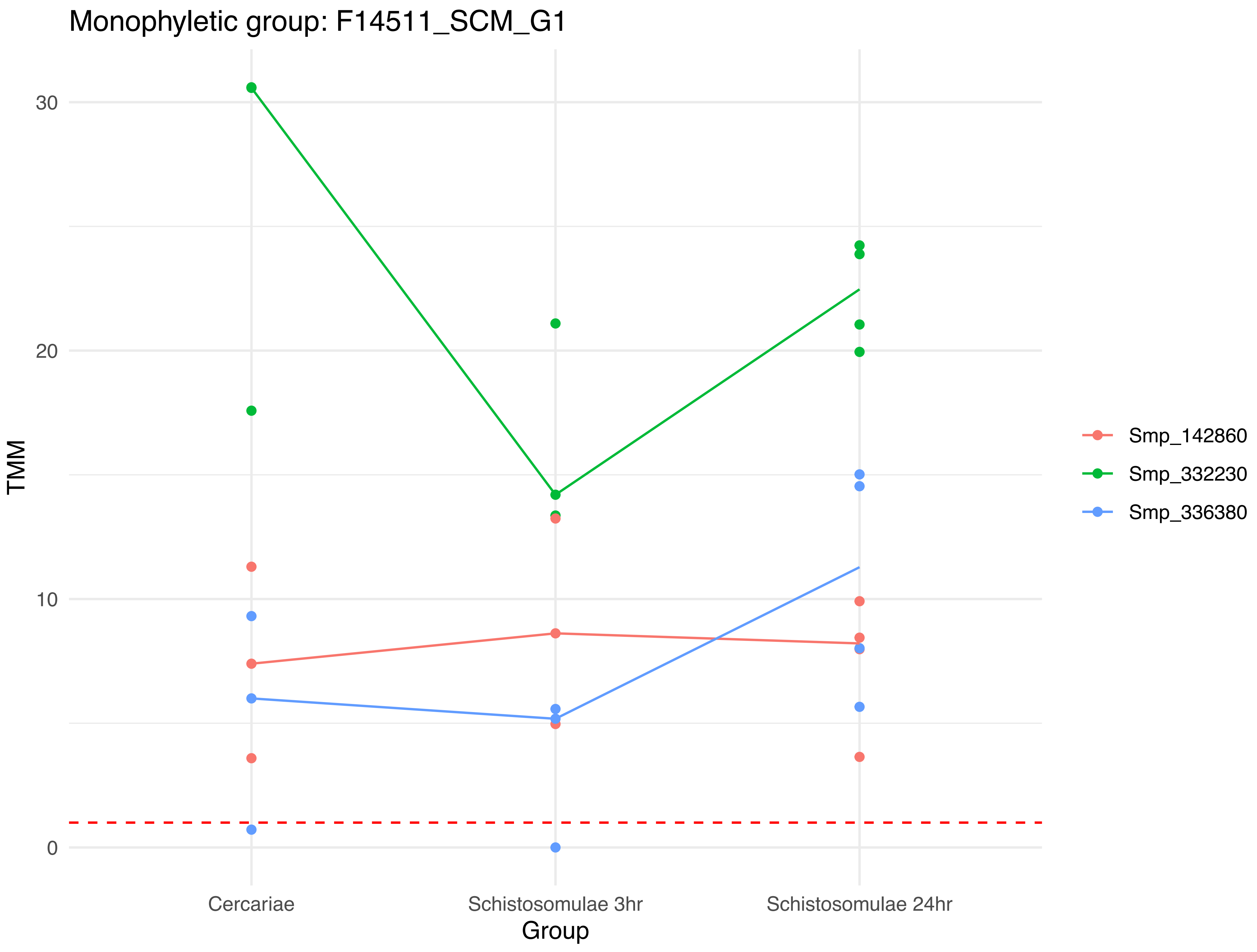

Monophyletic group: F14511_SCM_G1
TMM
30
20
10
0
Cercariae
Schistosomulae 3hr
Schistosomulae 24hr
Group
Smp_142860
Smp_332230
Smp_336380

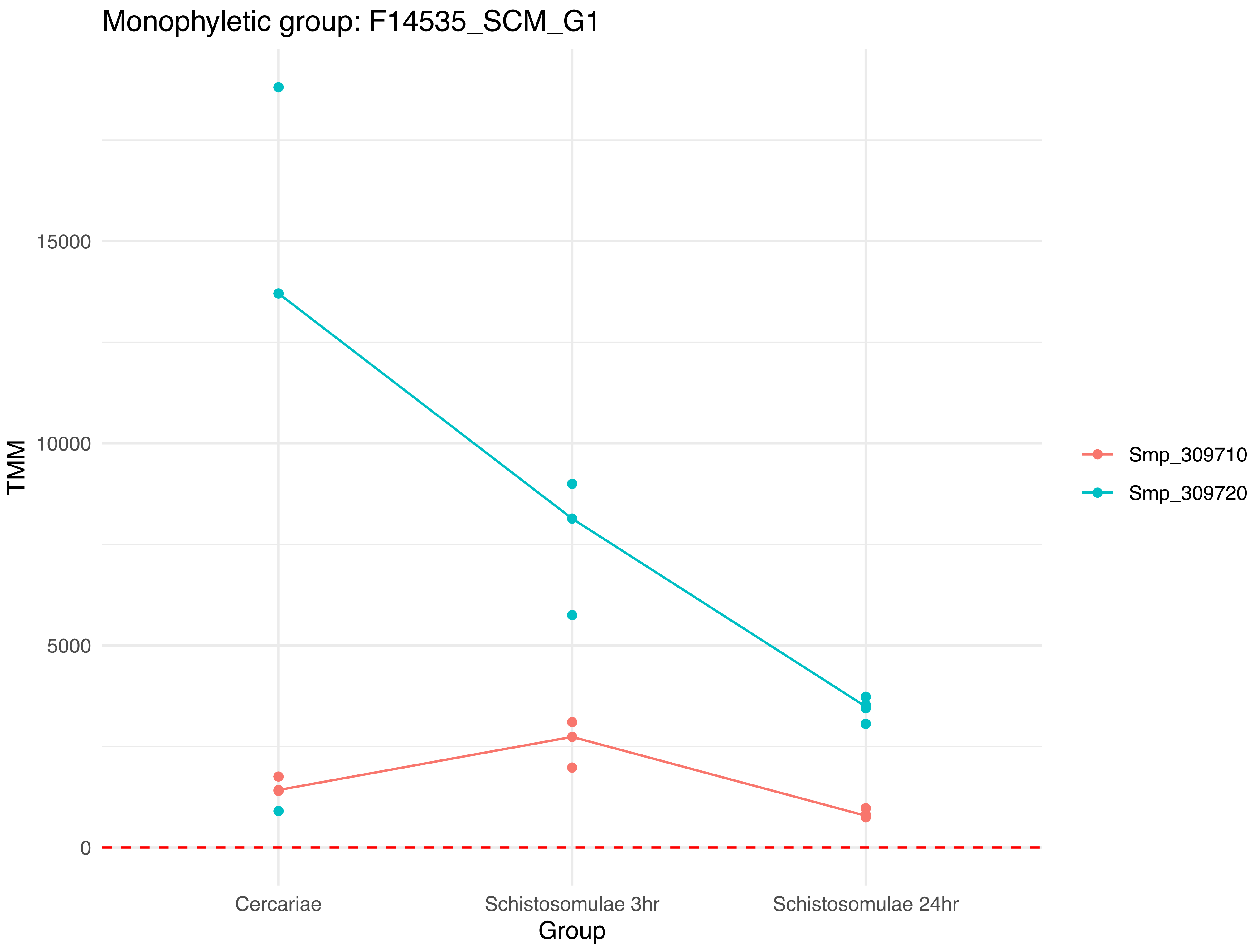
Monophyletic group: F14535_SCM_G1
TMM
15000
10000
5000
0
Cercariae
Schistosomulae 3hr
Schistosomulae 24hr
Group
Smp_309710
Smp_309720

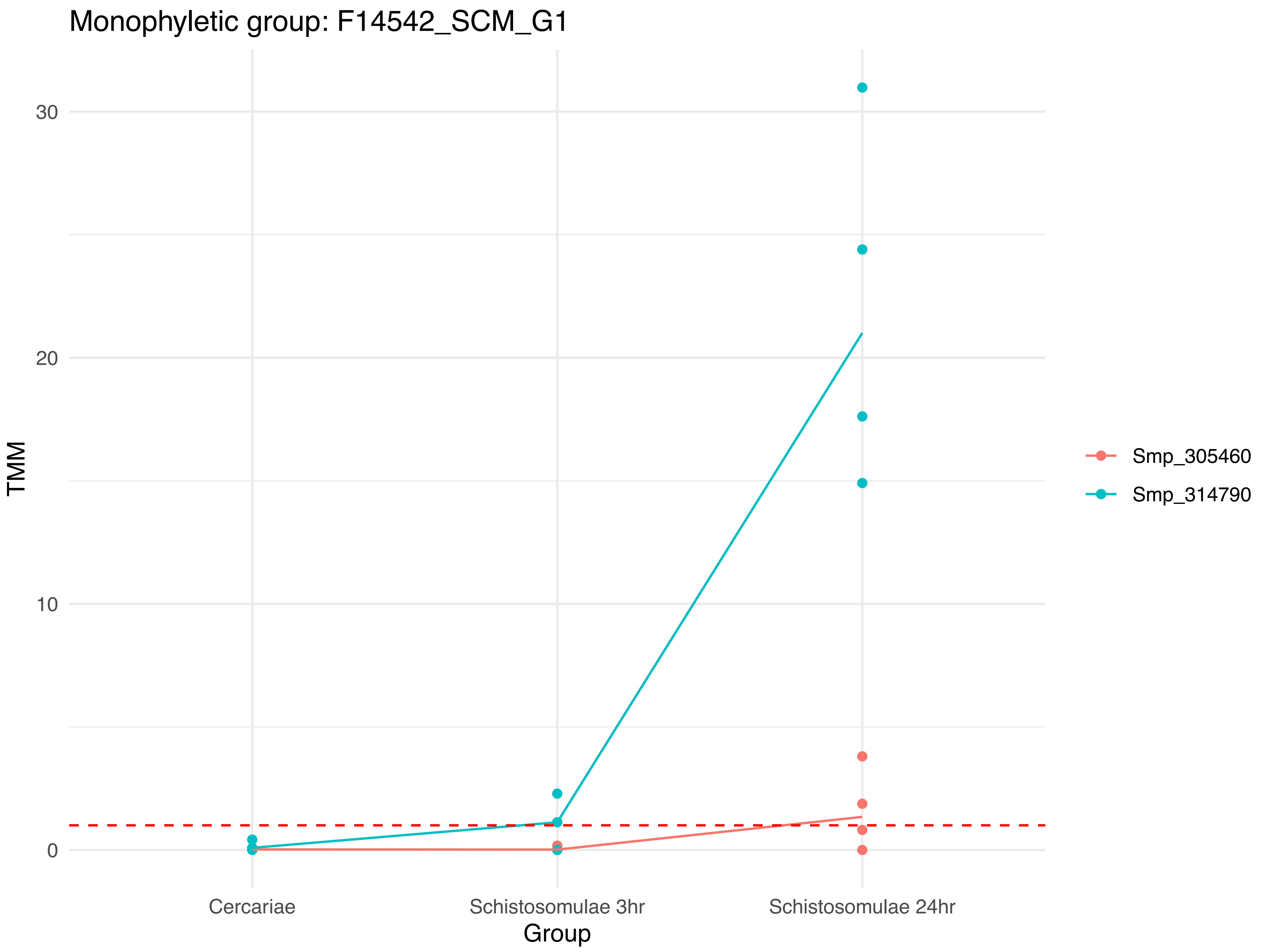

Monophyletic group: F14542_SCM_G1
30
20
10
0
TMM
Cercariae
Schistosomulae 3hr
Schistosomulae 24hr
Group
Smp_305460
Smp_314790

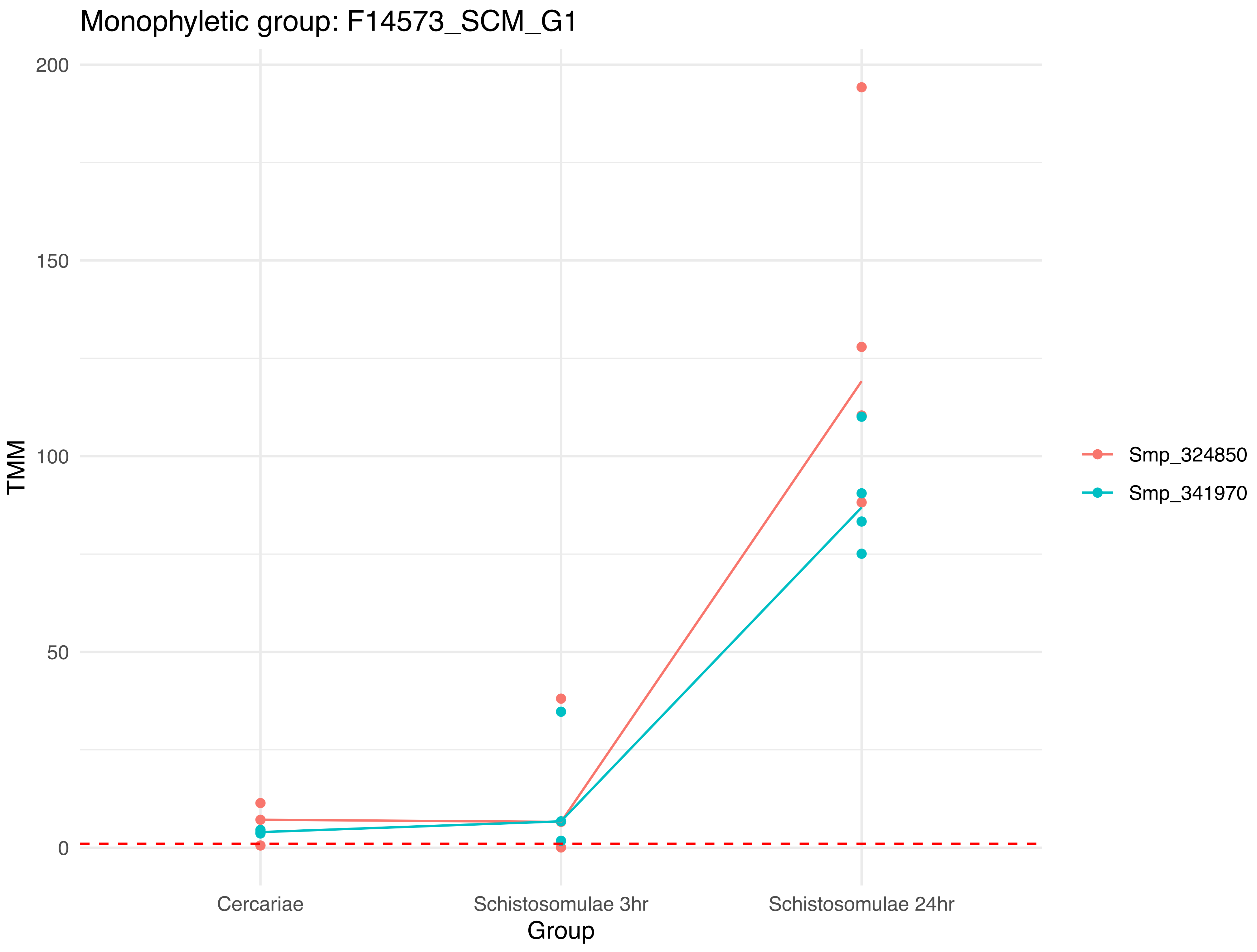
Monophyletic group: F14573_SCM_G1
TMM
200
150
100
50
0
Cercariae
Schistosomulae 3hr
Schistosomulae 24hr
Group
Smp_324850
Smp_341970

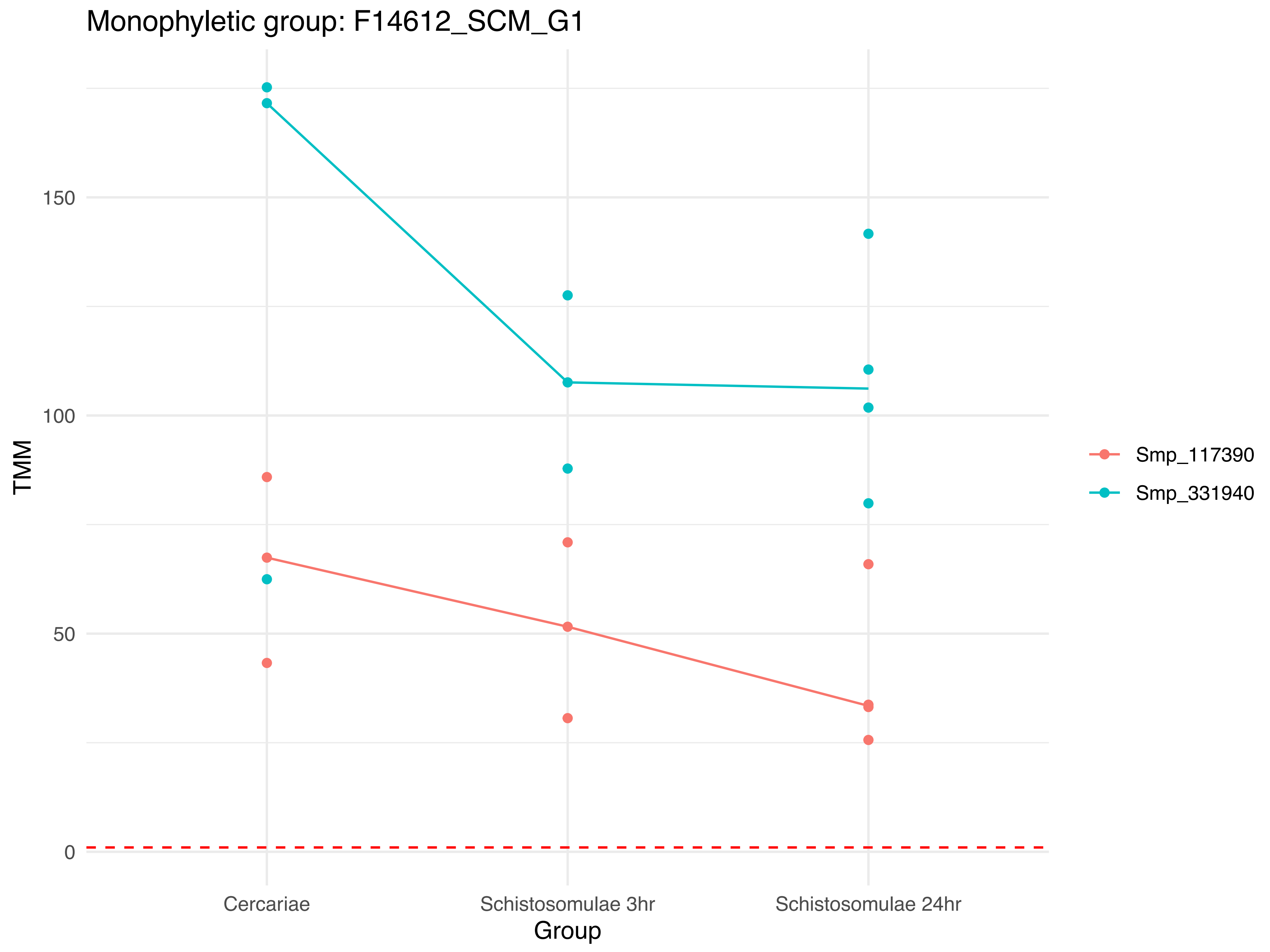

Monophyletic group: F14612_SCM_G1
150
100
50
0
TMM
Cercariae
Schistosomulae 3hr
Schistosomulae 24hr
Group
Smp_117390
Smp_331940

### Monophyletic group: F14613_SCM_G1

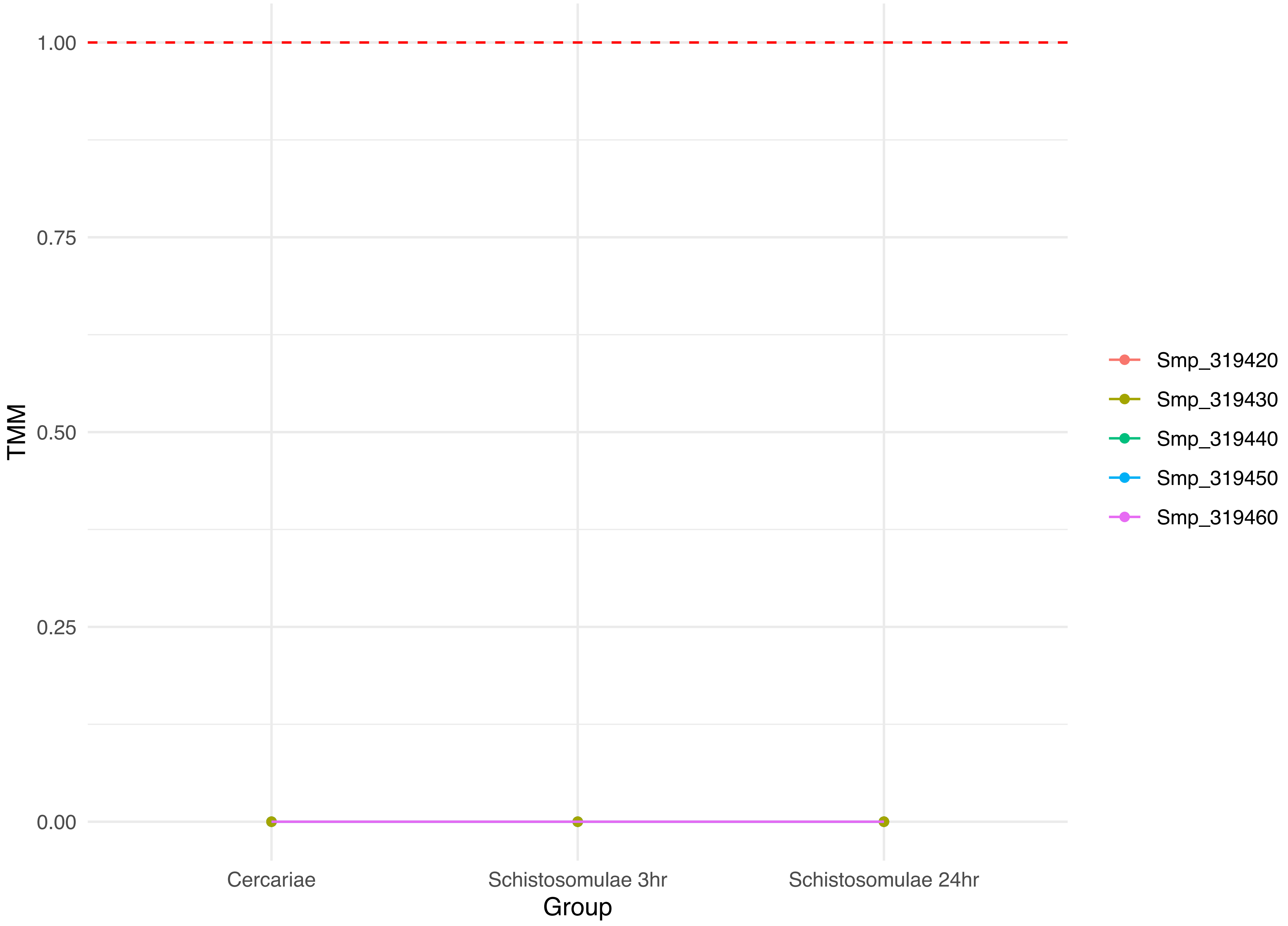

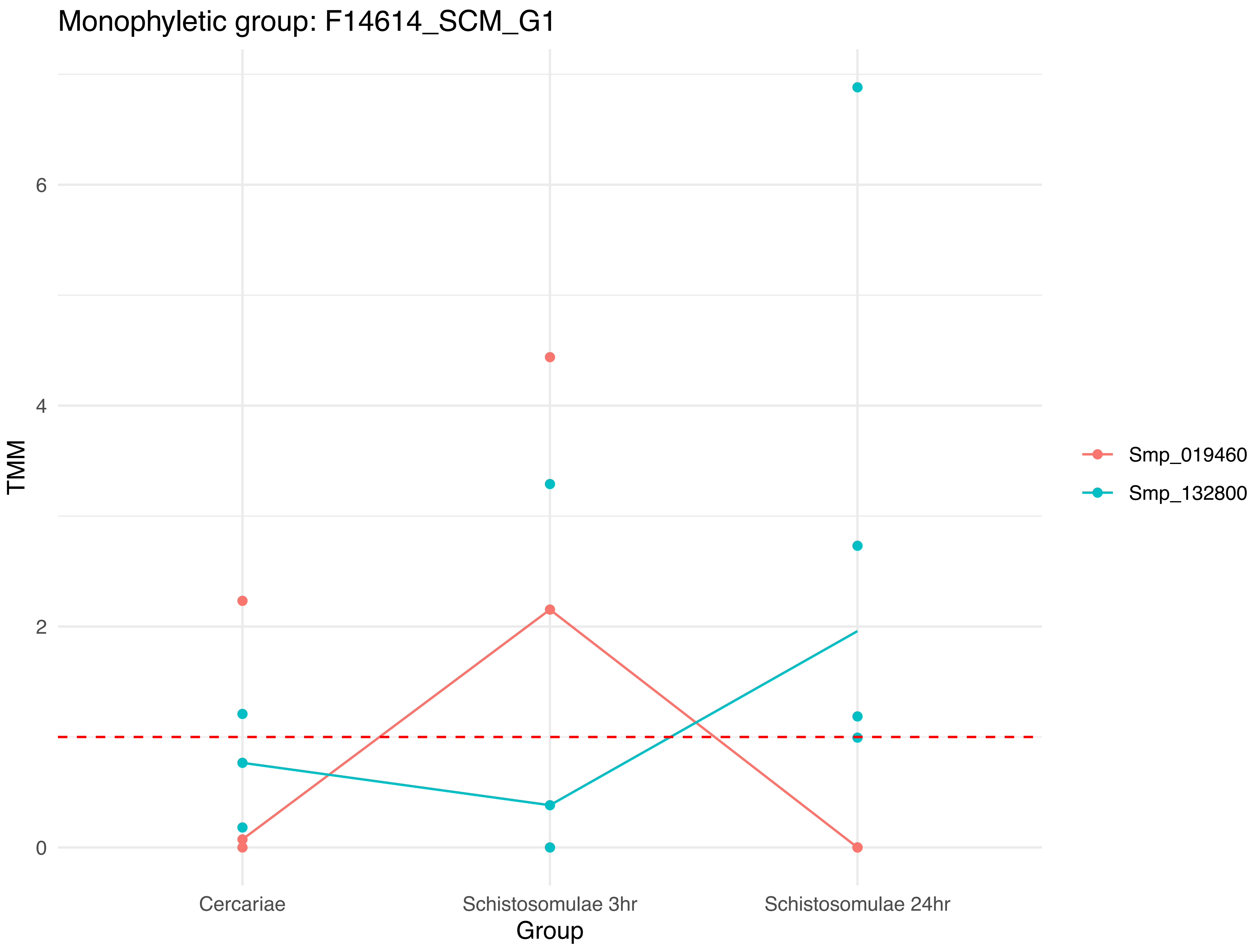

Monophyletic group: F14614_SCM_G1
TMM
6
4
2
0
Cercariae
Schistosomulae 3hr
Schistosomulae 24hr
Group
Smp_019460
Smp_132800

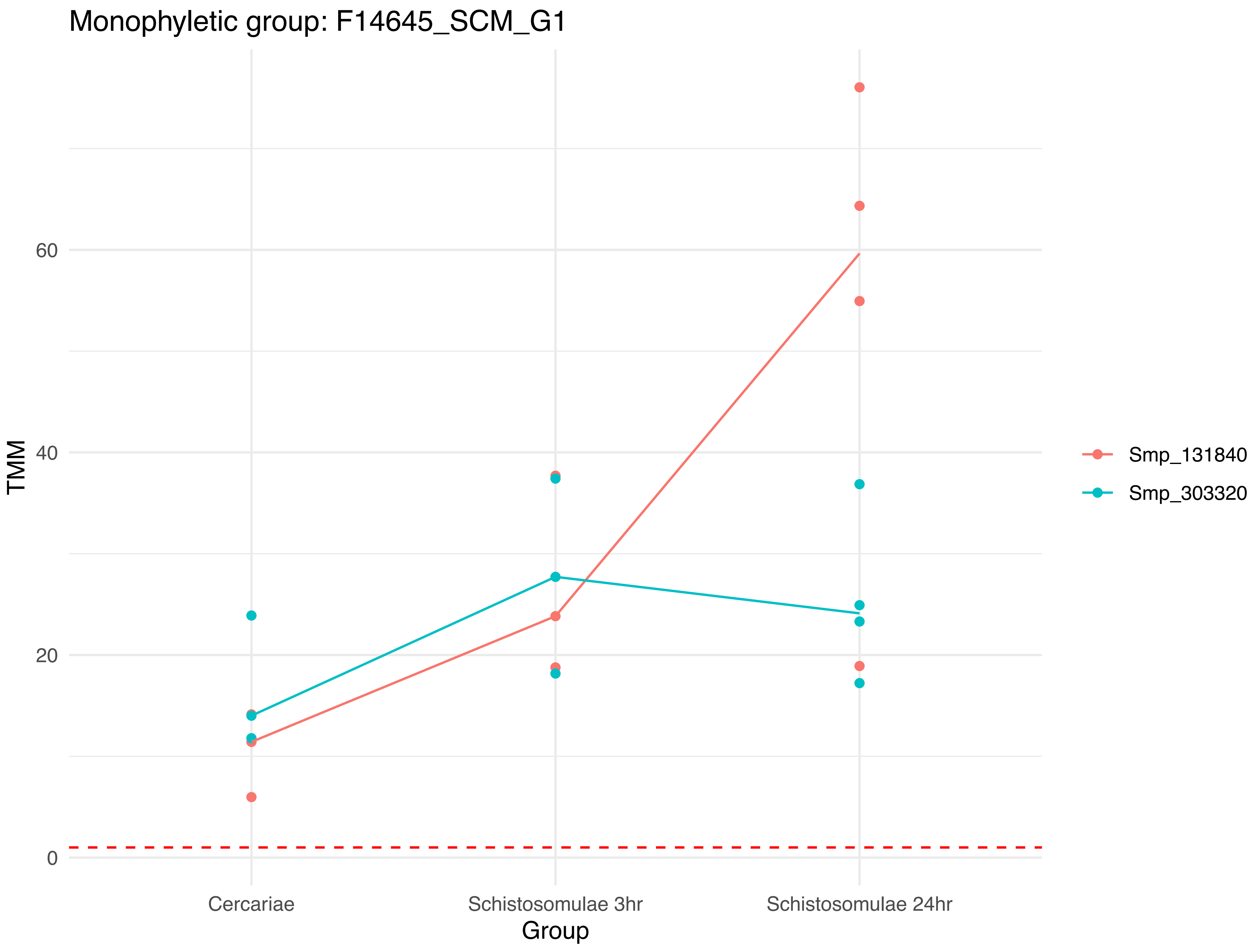
Monophyletic group: F14645_SCM_G1
TMM
60
40
20
0
Cercariae
Schistosomulae 3hr
Schistosomulae 24hr
Group
Smp_131840
Smp_303320

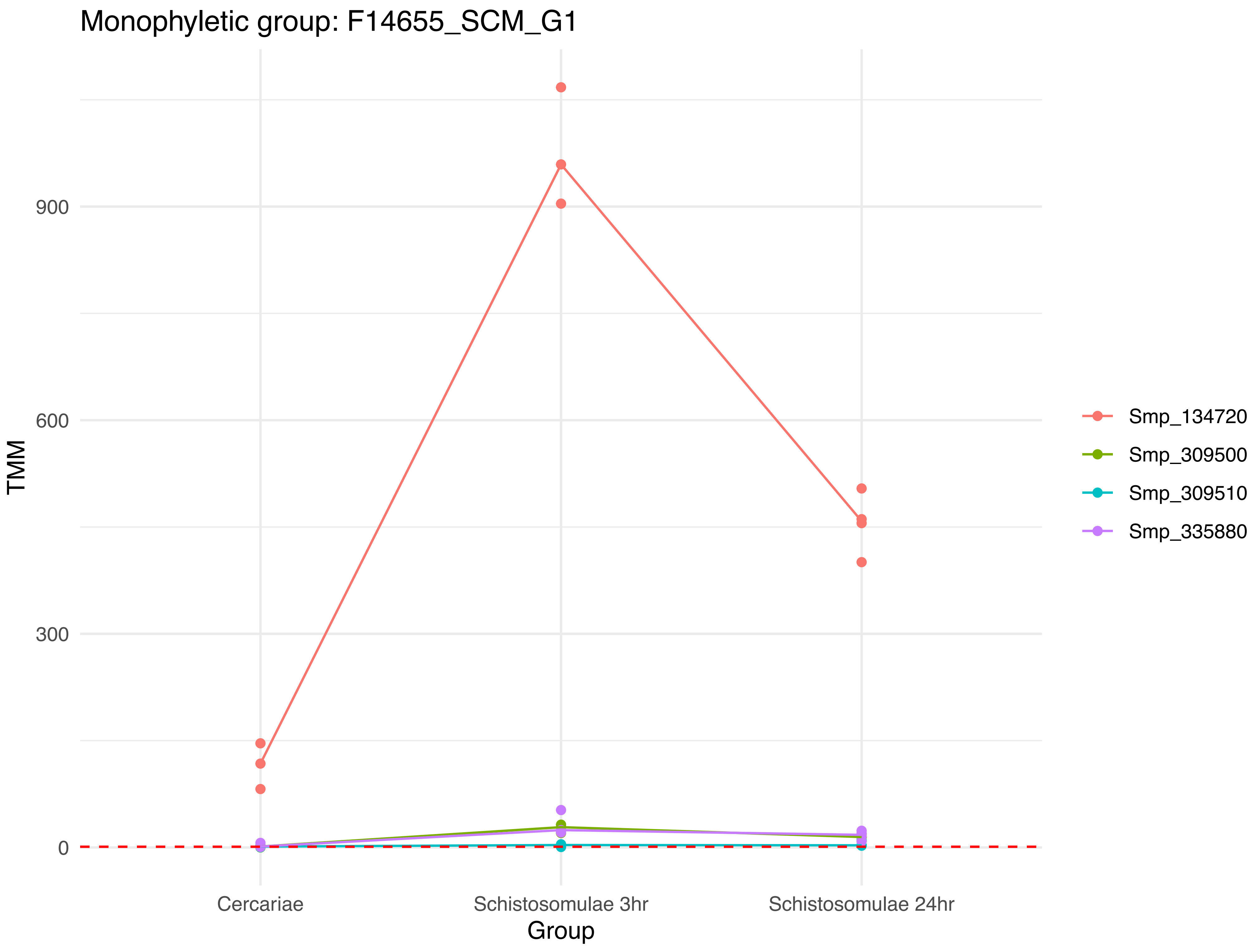

Monophyletic group: F14655_SCM_G1
TMM
900
600
300
0
Cercariae
Schistosomulae 3hr
Schistosomulae 24hr
Group
Smp_134720
Smp_309500
Smp_309510
Smp_335880

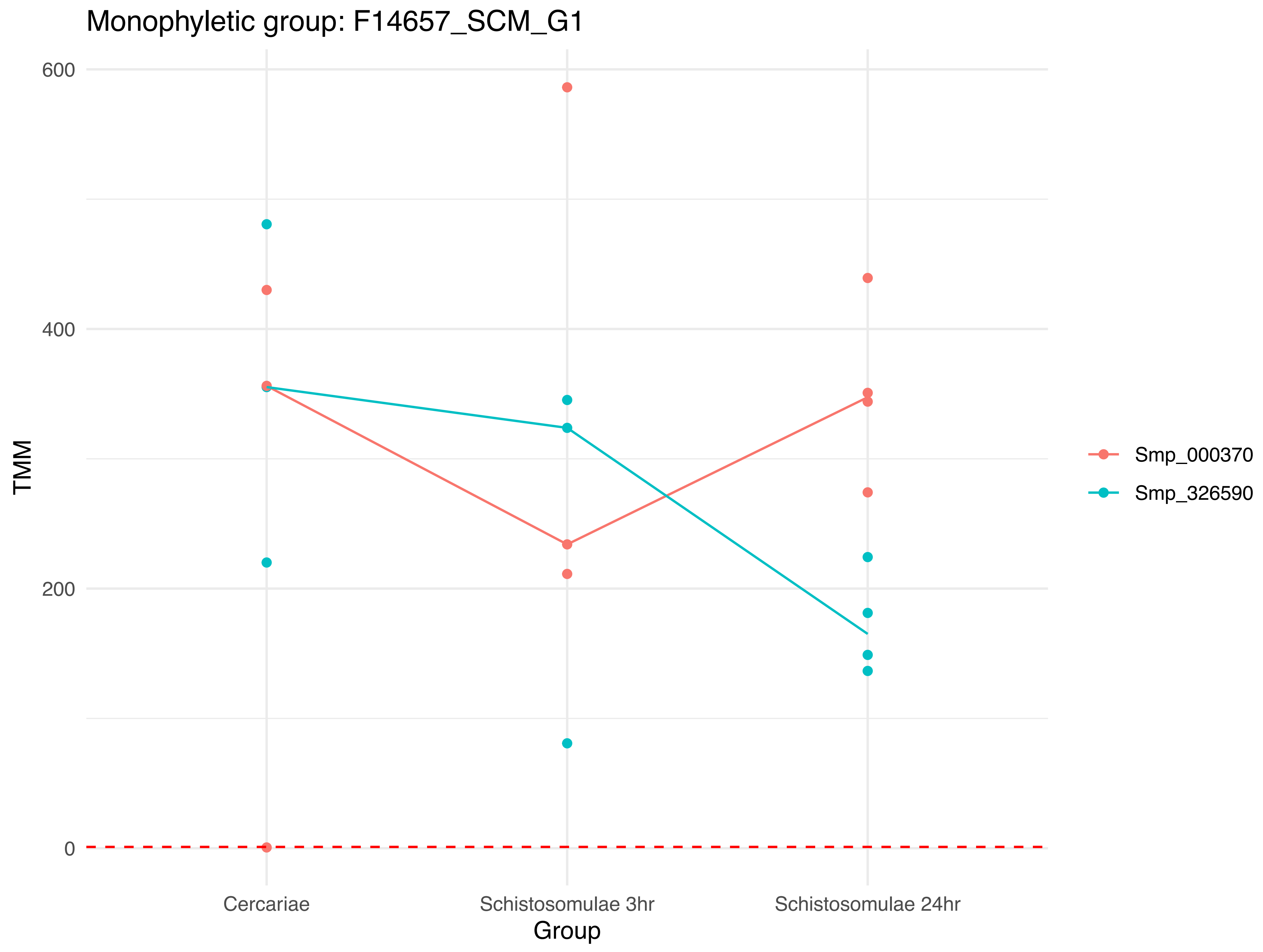

Monophyletic group: F14657_SCM_G1
600
400
200
0
TMM
Cercariae
Schistosomulae 3hr
Schistosomulae 24hr
Group
Smp_000370
Smp_326590

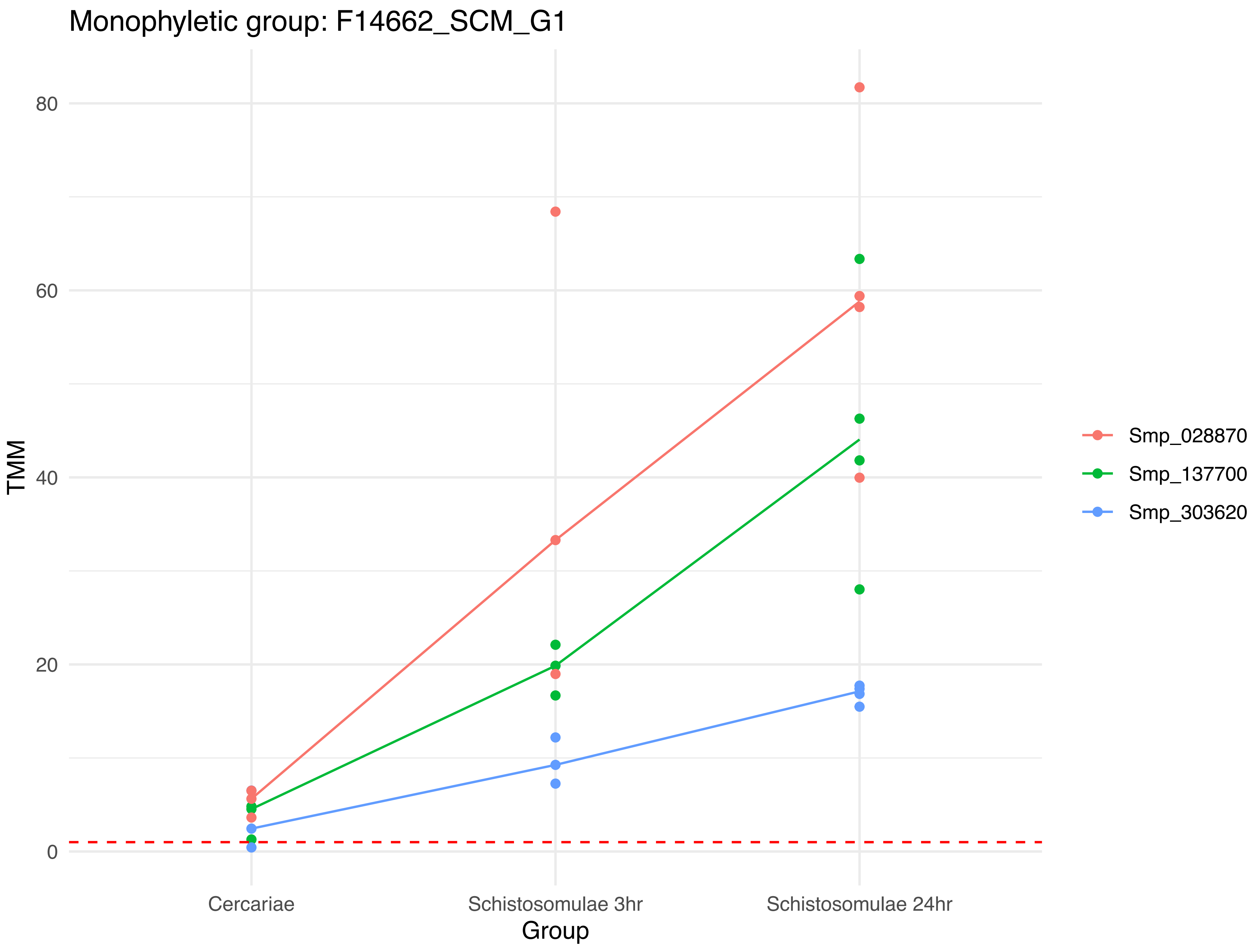
Monophyletic group: F14662_SCM_G1
TMM
80
60
40
20
0
Cercariae
Schistosomulae 3hr
Schistosomulae 24hr
Group
Smp_028870
Smp_137700
Smp_303620

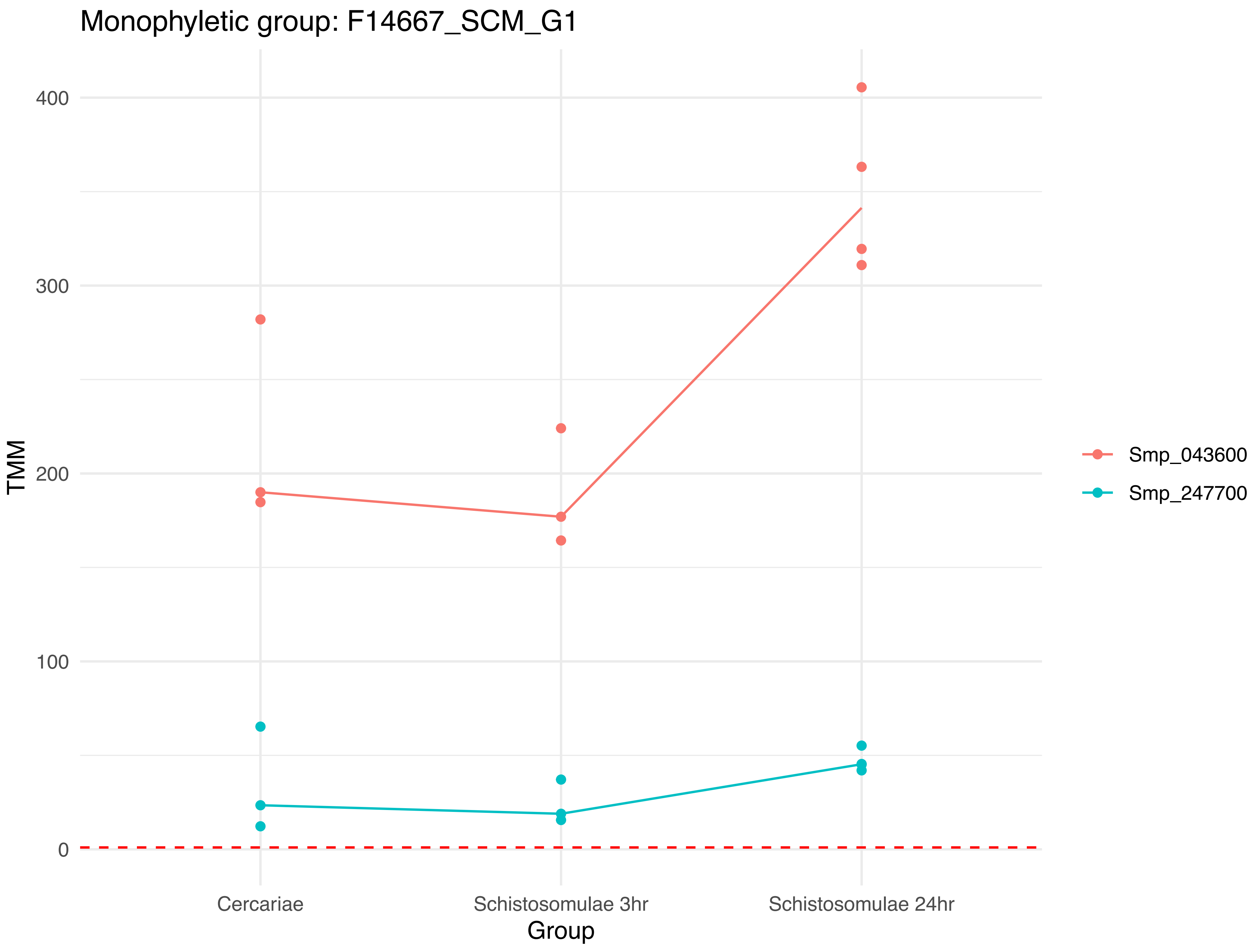
Monophyletic group: F14667_SCM_G1
400
300
200
100
0
TMM
Cercariae
Schistosomulae 3hr
Schistosomulae 24hr
Group
Smp_043600
Smp_247700

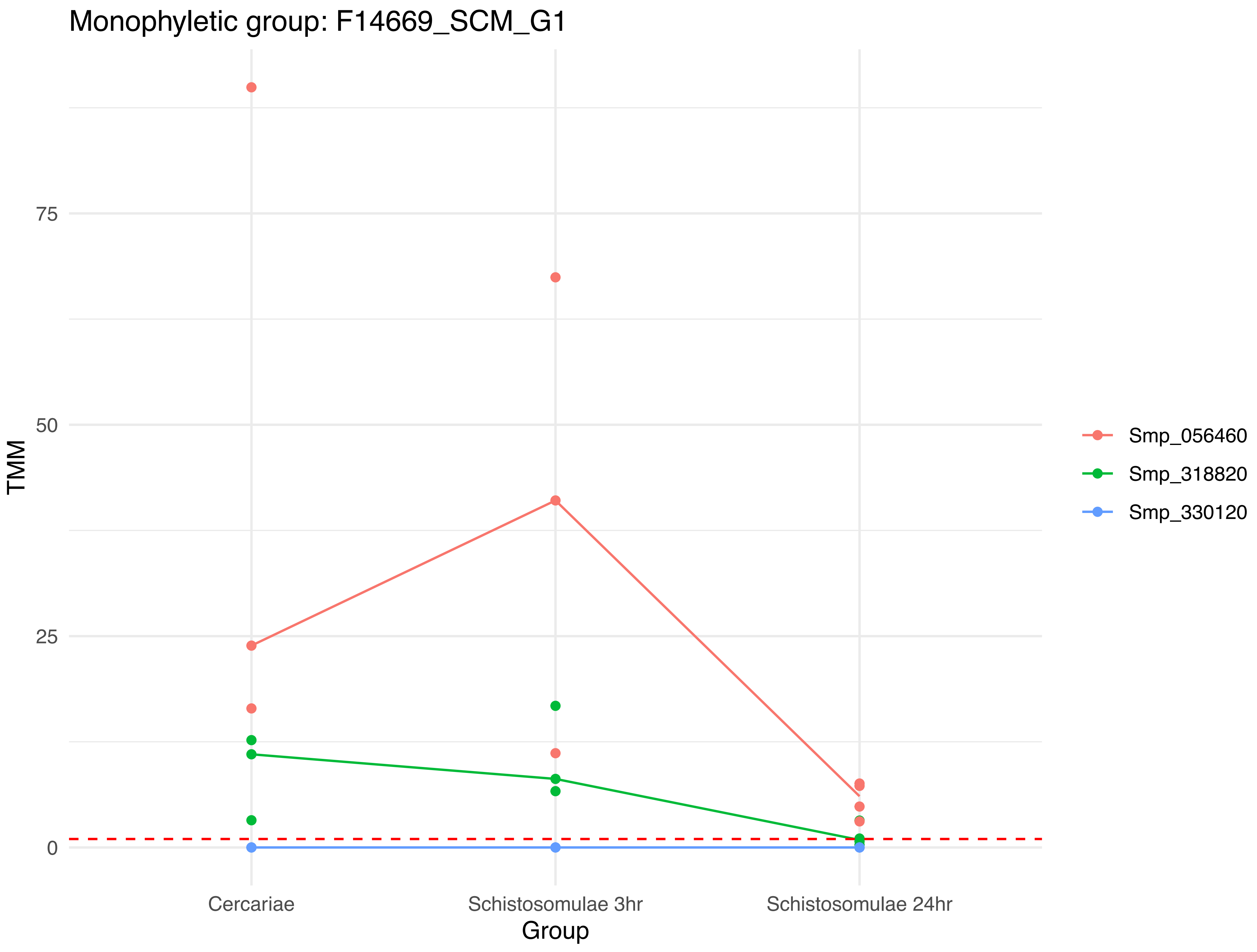
Monophyletic group: F14669_SCM_G1
TMM
75
50
25
0
Cercariae
Schistosomulae 3hr
Schistosomulae 24hr
Group
Smp_056460
Smp_318820
Smp_330120

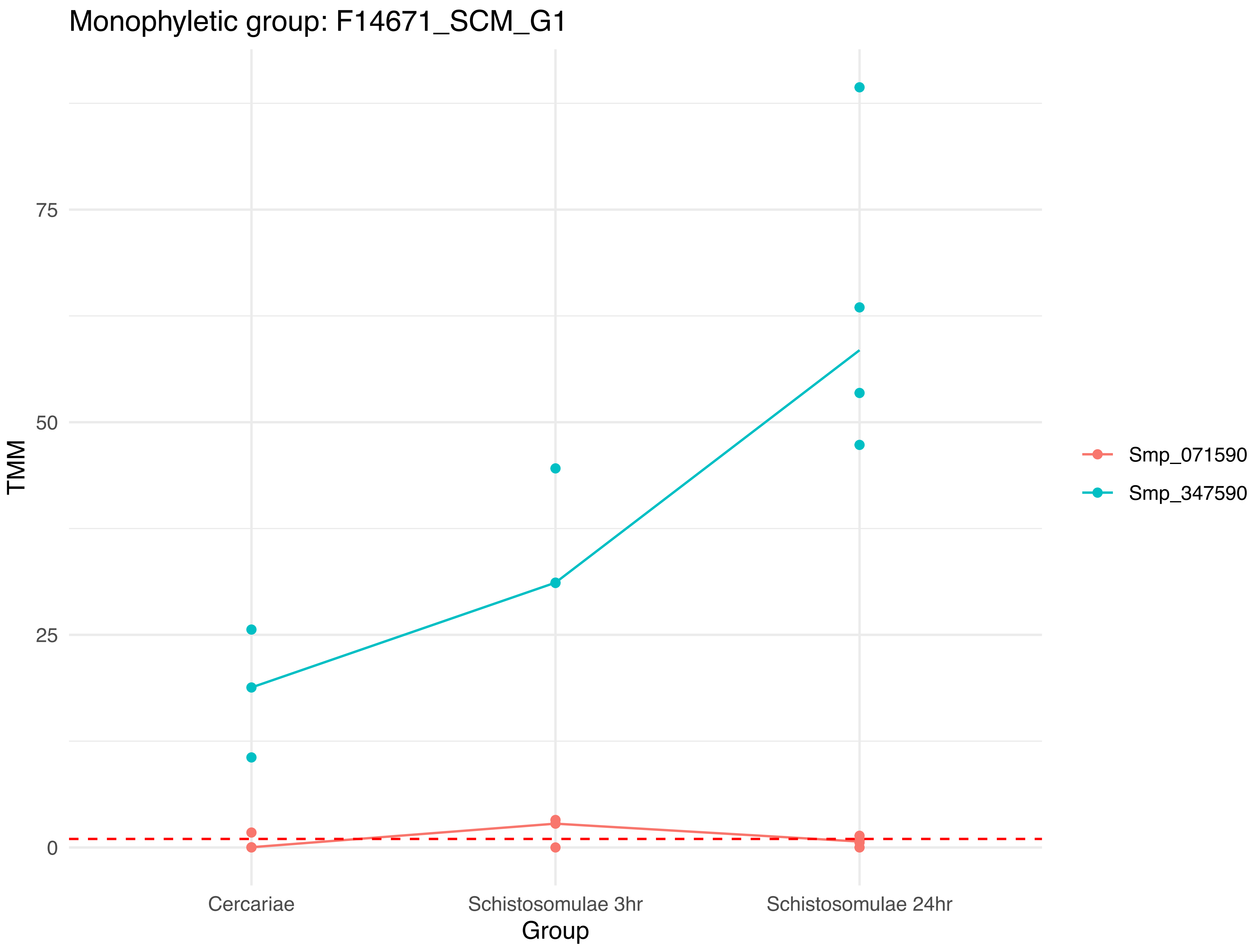

Monophyletic group: F14671_SCM_G1
TMM
0
25
50
75
Cercariae
Schistosomulae 3hr
Schistosomulae 24hr
Group
Smp_071590
Smp_347590

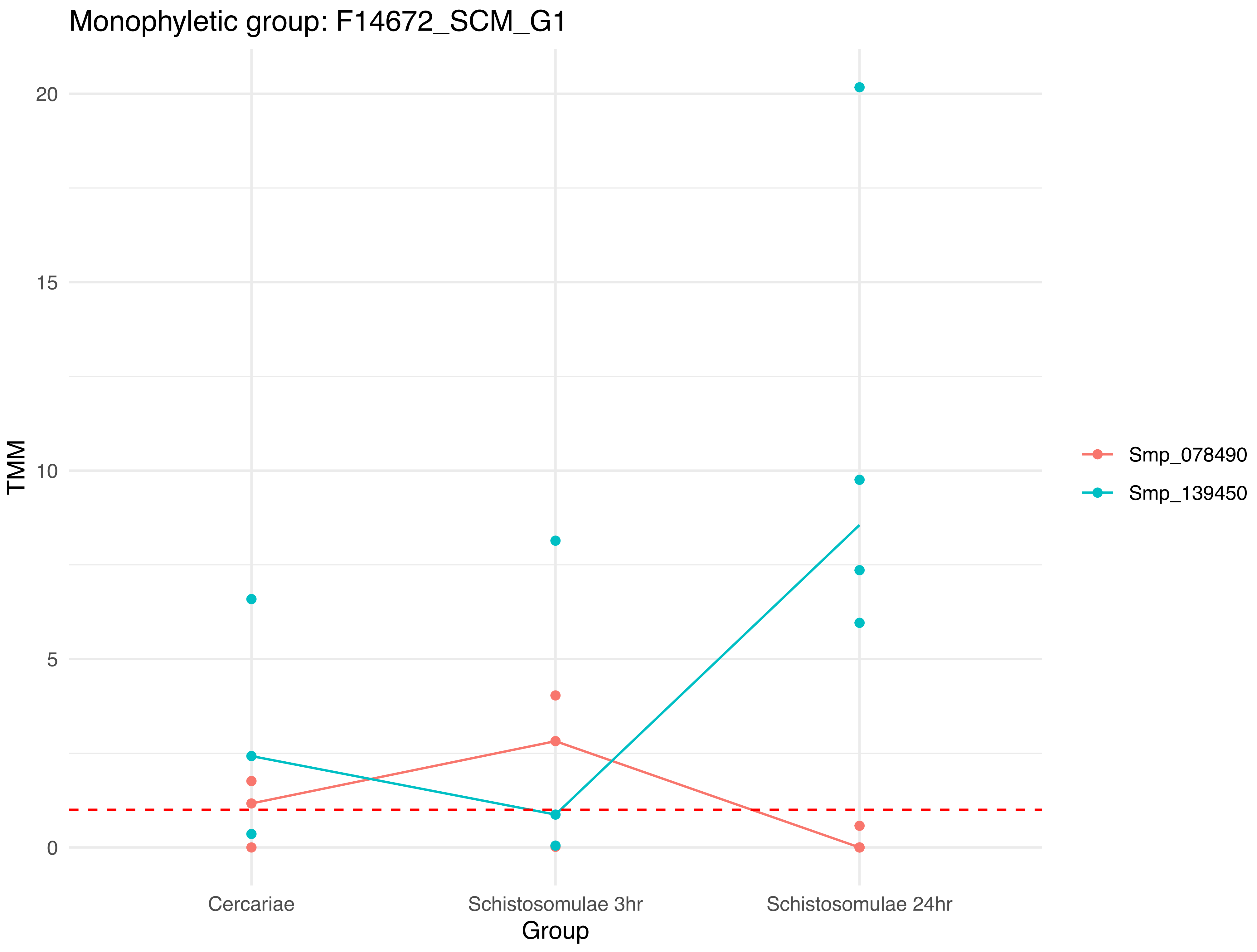
Monophyletic group: F14672_SCM_G1
20
15
10
5
0
TMM
Cercariae
Schistosomulae 3hr
Schistosomulae 24hr
Group
Smp_078490
Smp_139450

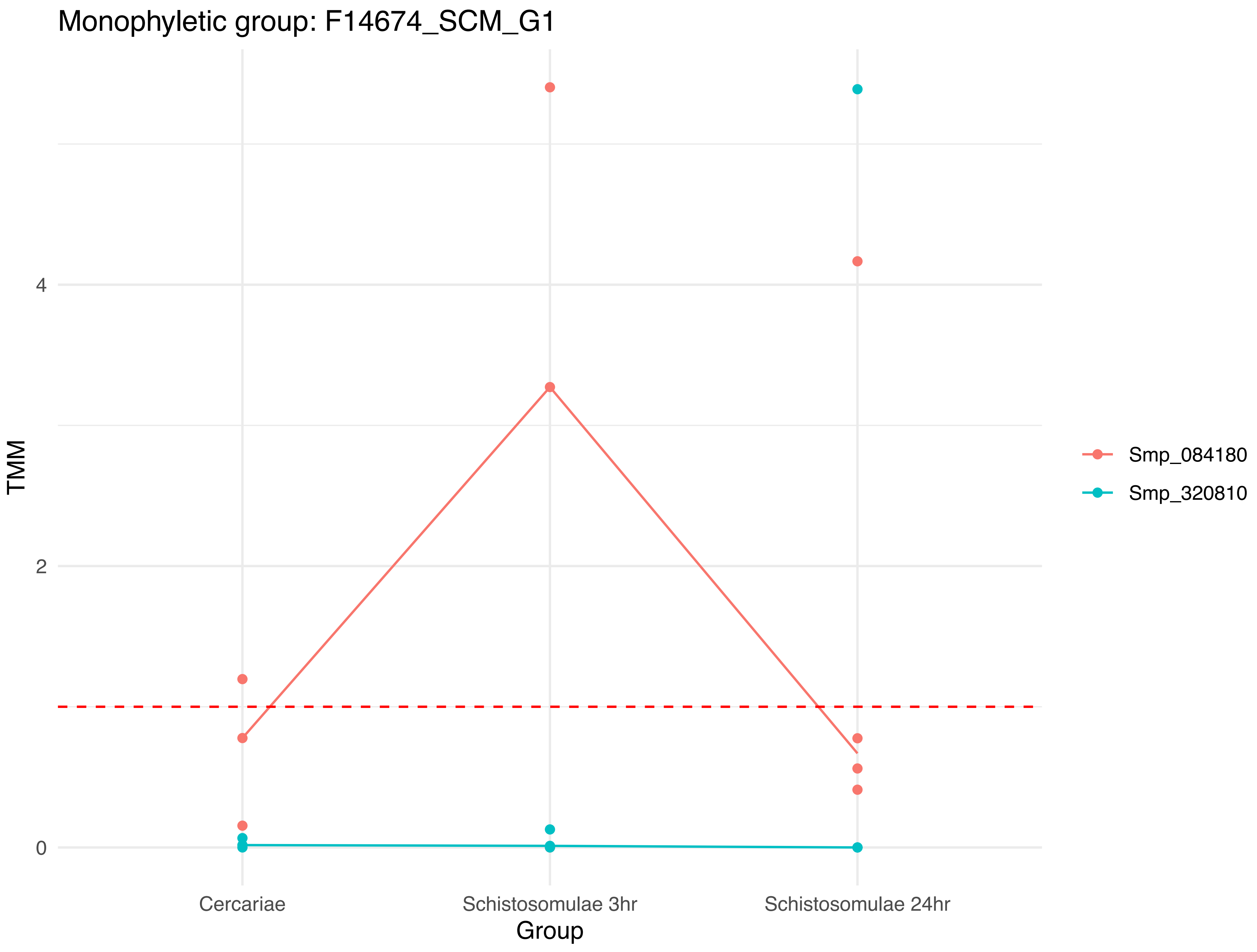

Monophyletic group: F14674_SCM_G1
TMM
0
2
4
Cercariae
Schistosomulae 3hr
Schistosomulae 24hr
Group
Smp_084180
Smp_320810

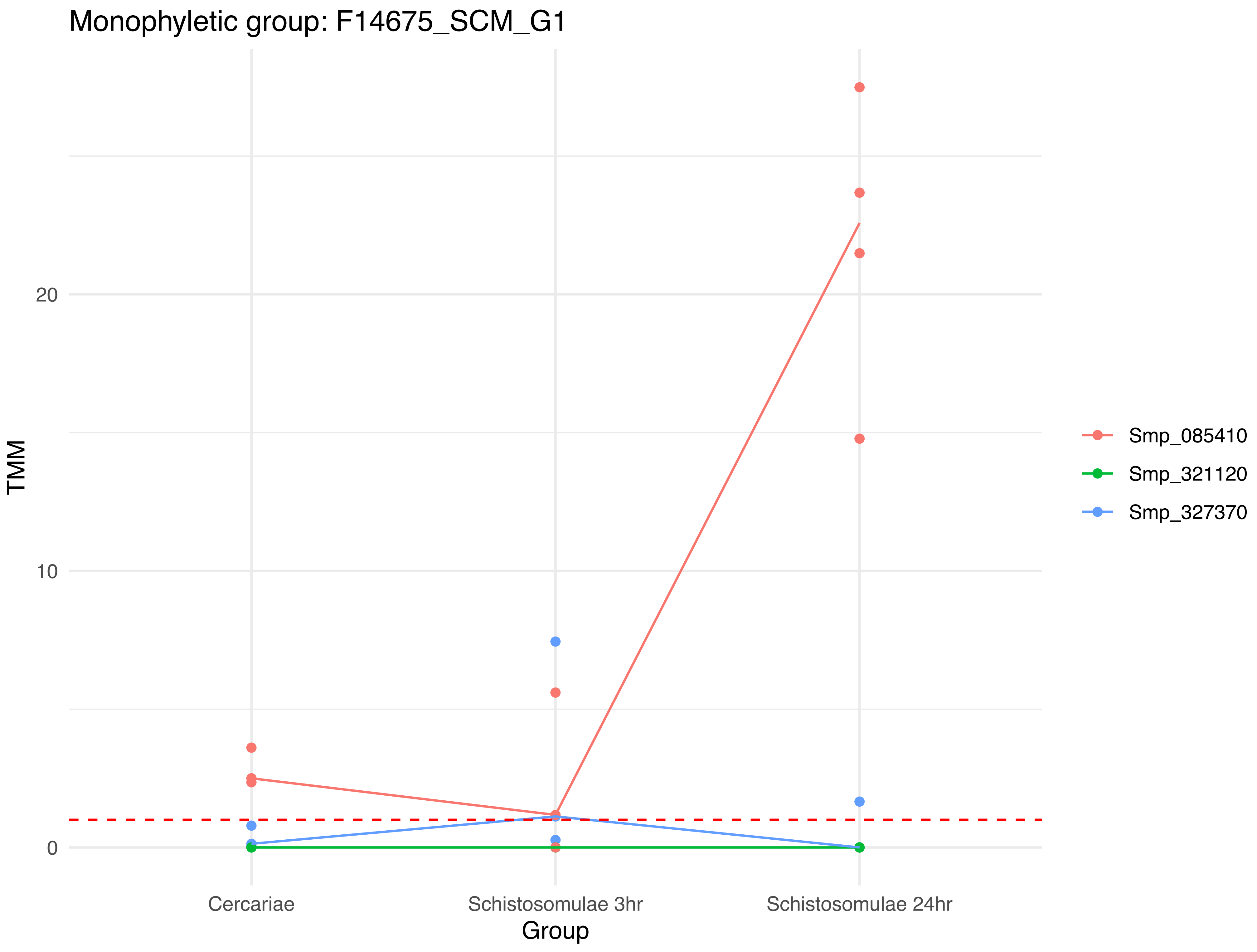

Monophyletic group: F14675_SCM_G1
TMM
20
10
0
Cercariae
Schistosomulae 3hr
Schistosomulae 24hr
Group
Smp_085410
Smp_321120
Smp_327370

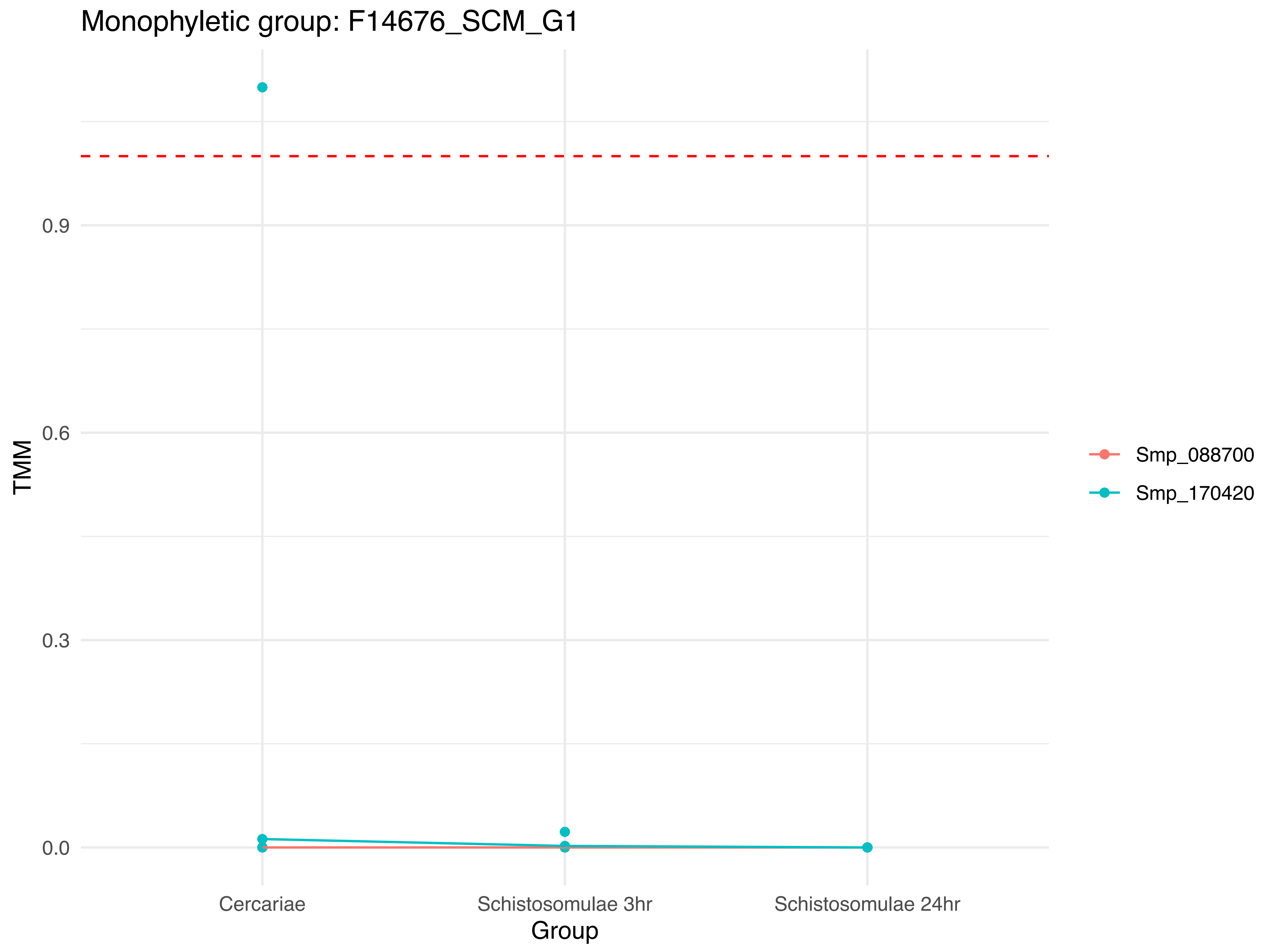

Monophyletic group: F14676_SCM_G1
TMM
0.9
0.6
0.3
0.0
Cercariae
Schistosomulae 3hr
Schistosomulae 24hr
Group
Smp_088700
Smp_170420

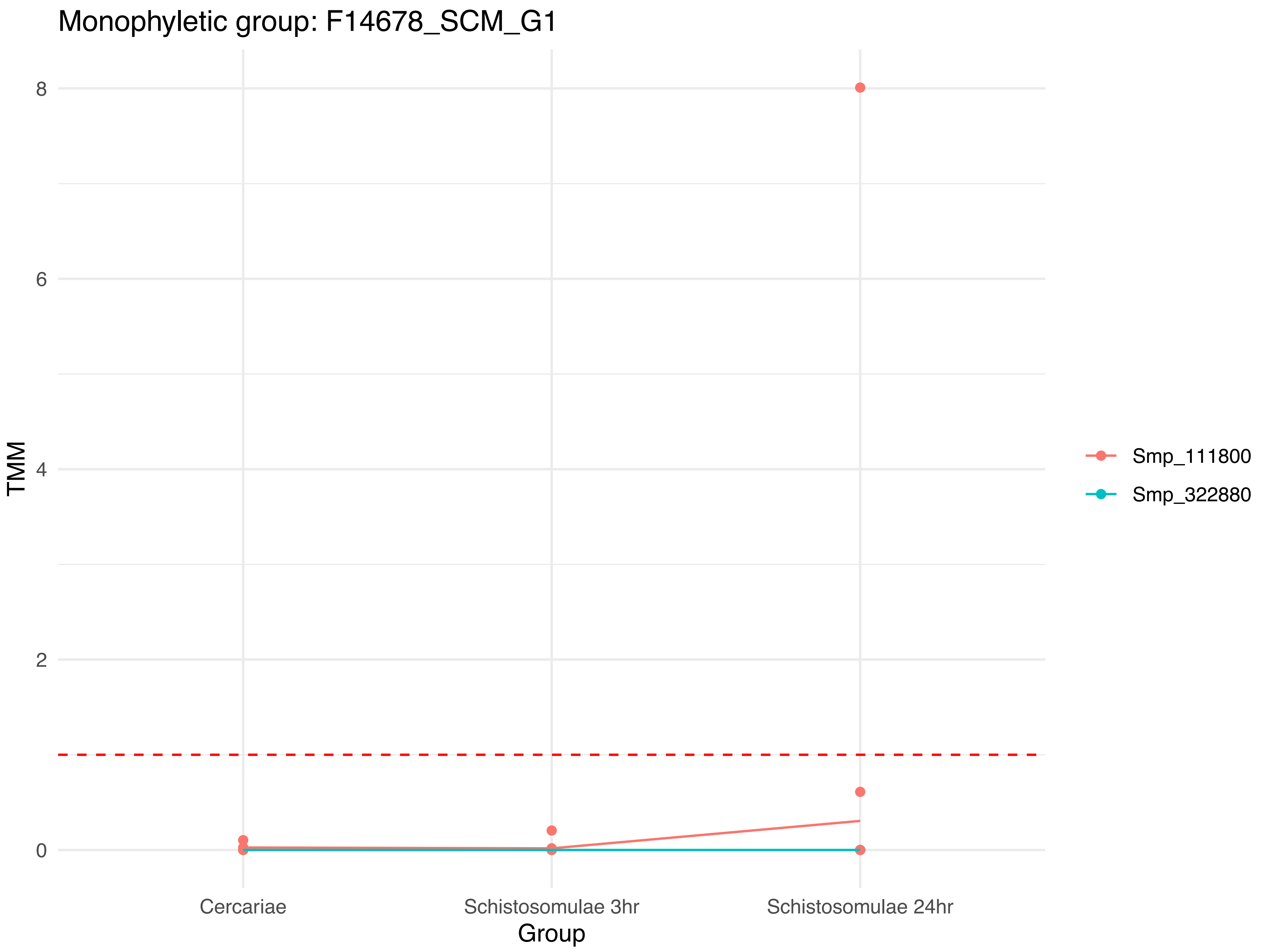
Monophyletic group: F14678_SCM_G1
8
6
4
2
0
TMM
Cercariae
Schistosomulae 3hr
Schistosomulae 24hr
Group
Smp_111800
Smp_322880

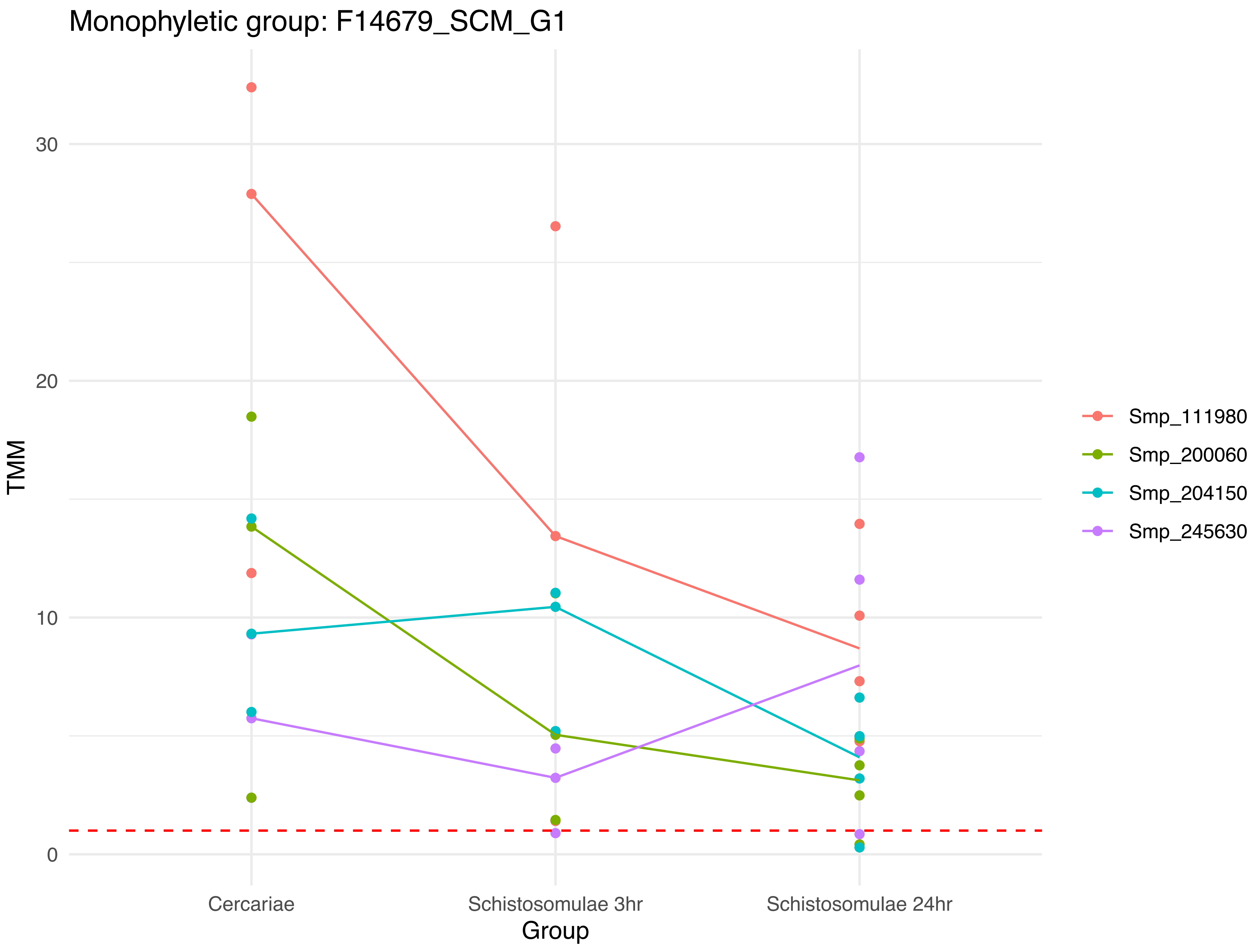

Monophyletic group: F14679_SCM_G1
TMM
30
20
10
0
Cercariae
Schistosomulae 3hr
Schistosomulae 24hr
Group
Smp_111980
Smp_200060
Smp_204150
Smp_245630

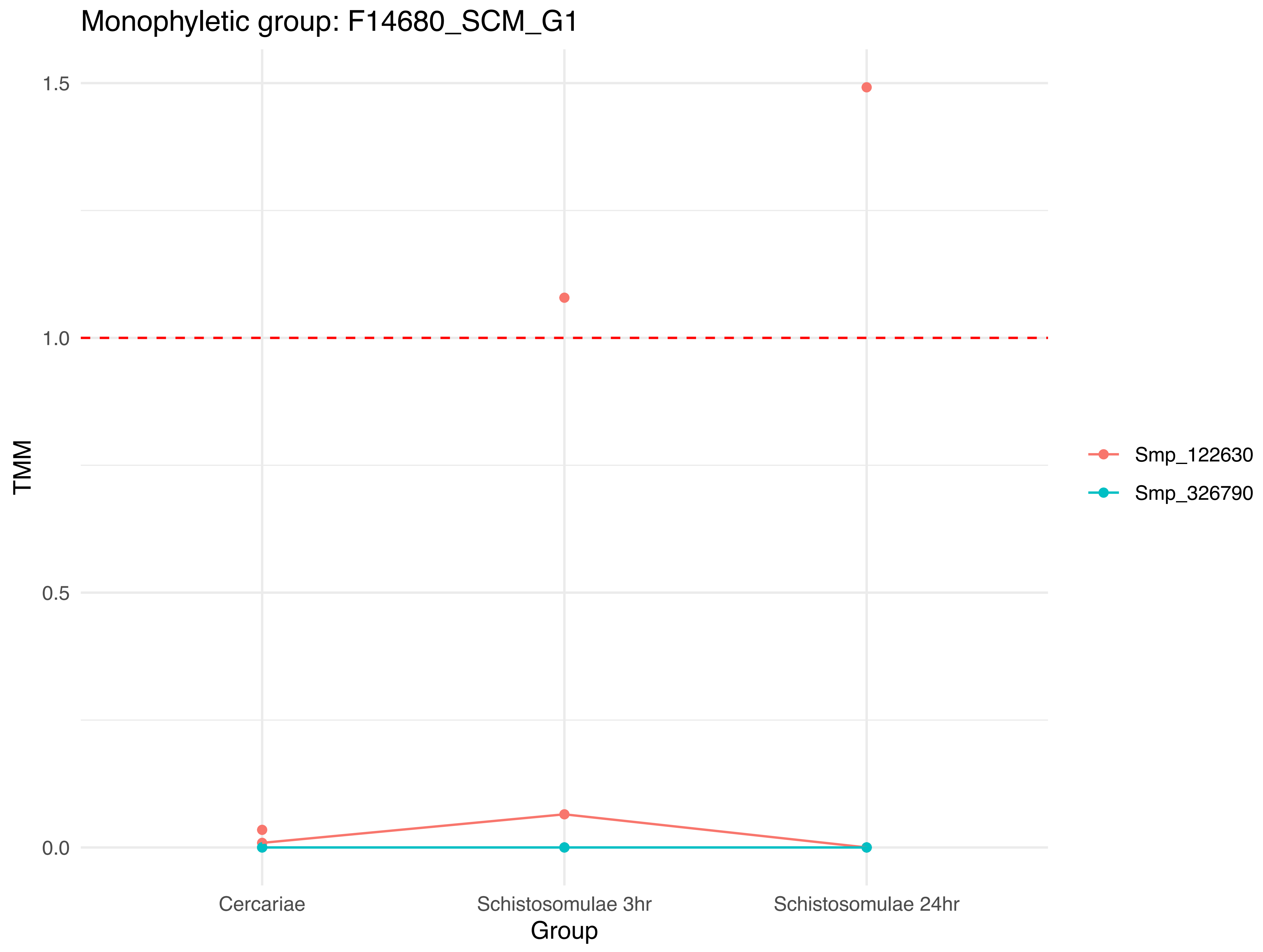

Monophyletic group: F14680_SCM_G1
TMM
1.5
1.0
0.5
0.0
Cercariae
Schistosomulae 3hr
Schistosomulae 24hr
Group
Smp_122630
Smp_326790

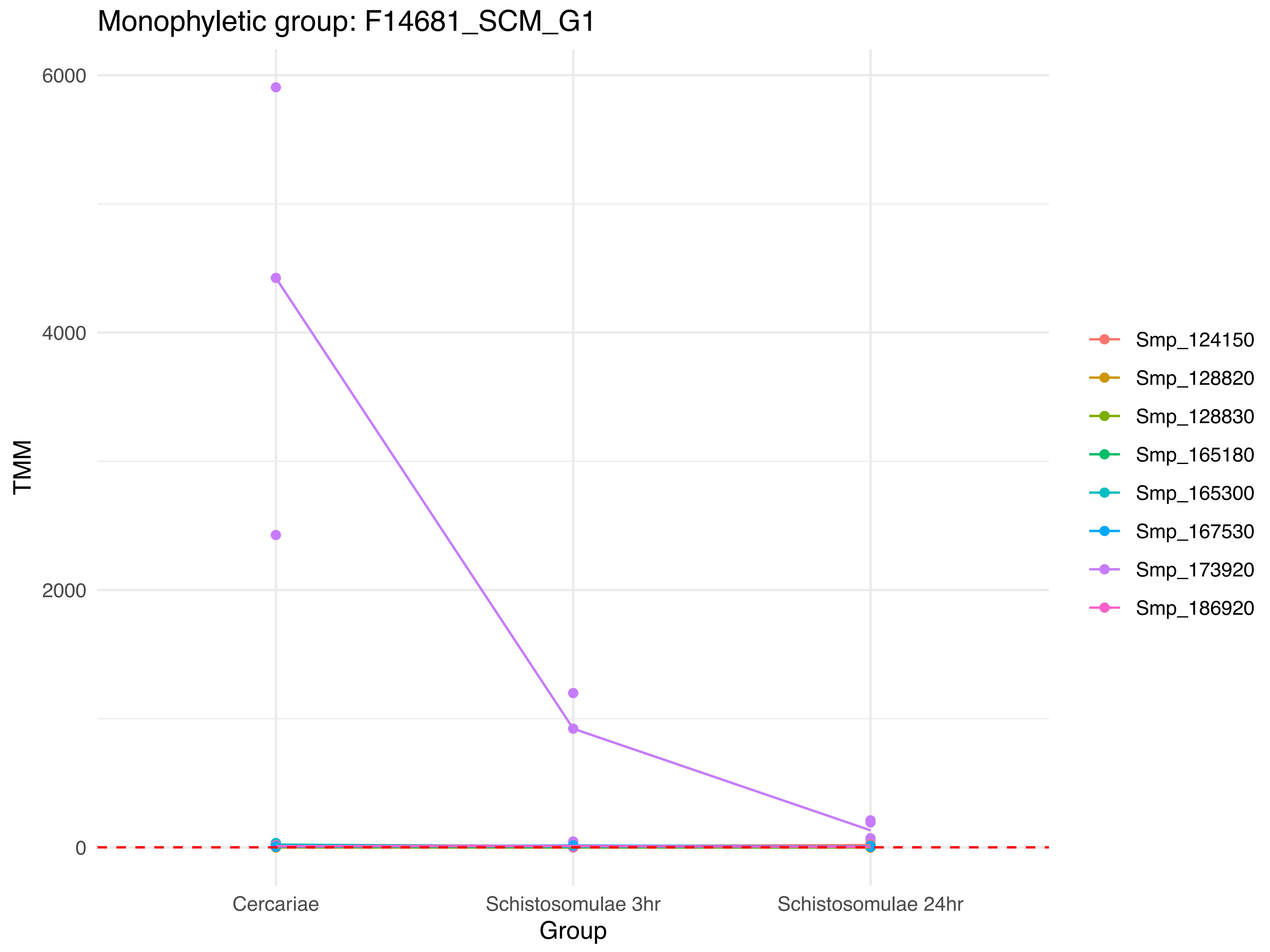

Monophyletic group: F14681_SCM_G1
6000
4000
2000
0
TMM
Cercariae
Schistosomulae 3hr
Schistosomulae 24hr
Group
Smp_124150
Smp_128820
Smp_128830
Smp_165180
Smp_165300
Smp_167530
Smp_173920
Smp_186920

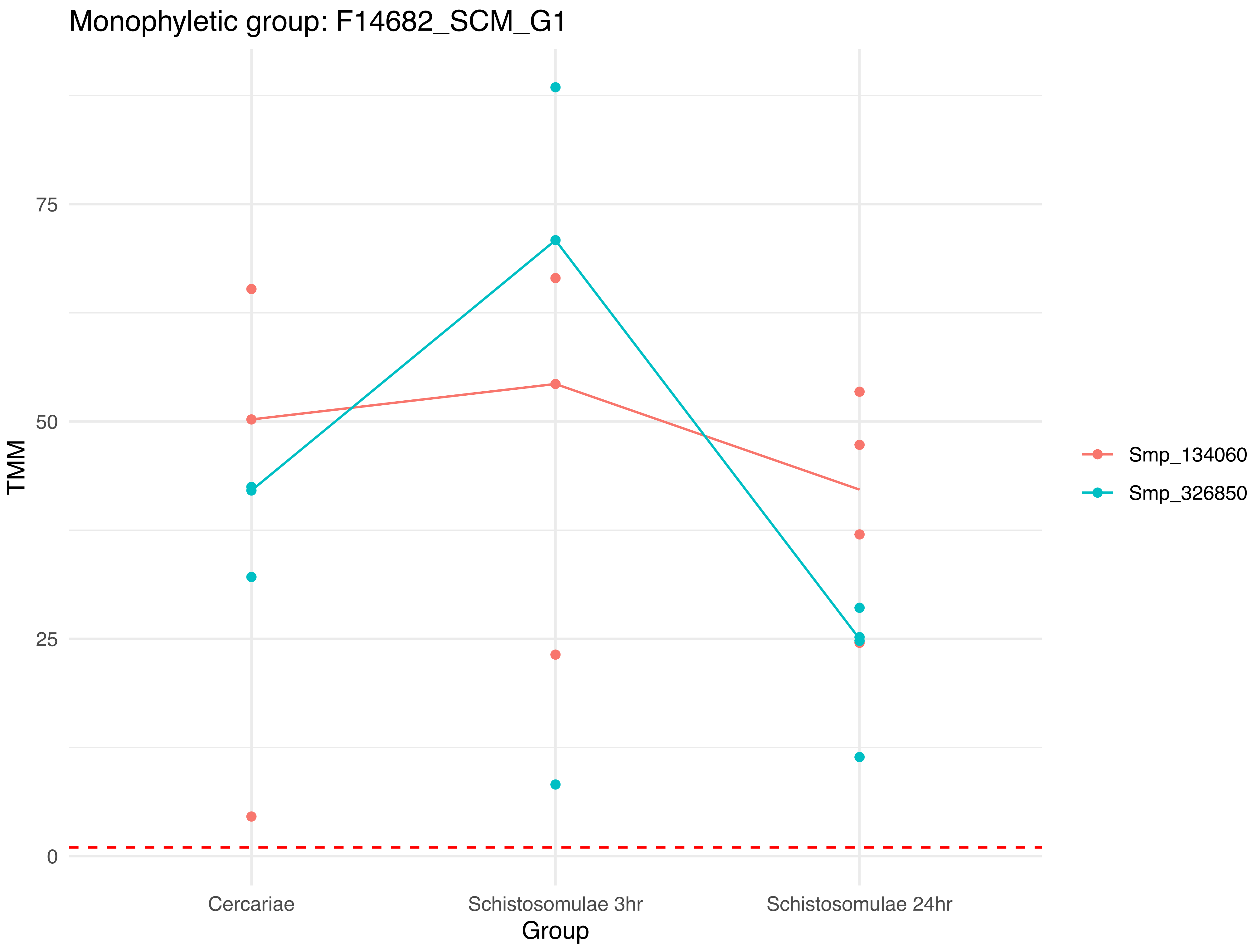

Monophyletic group: F14682_SCM_G1
75
50
25
0
TMM
Cercariae
Schistosomulae 3hr
Schistosomulae 24hr
Group
Smp_134060
Smp_326850

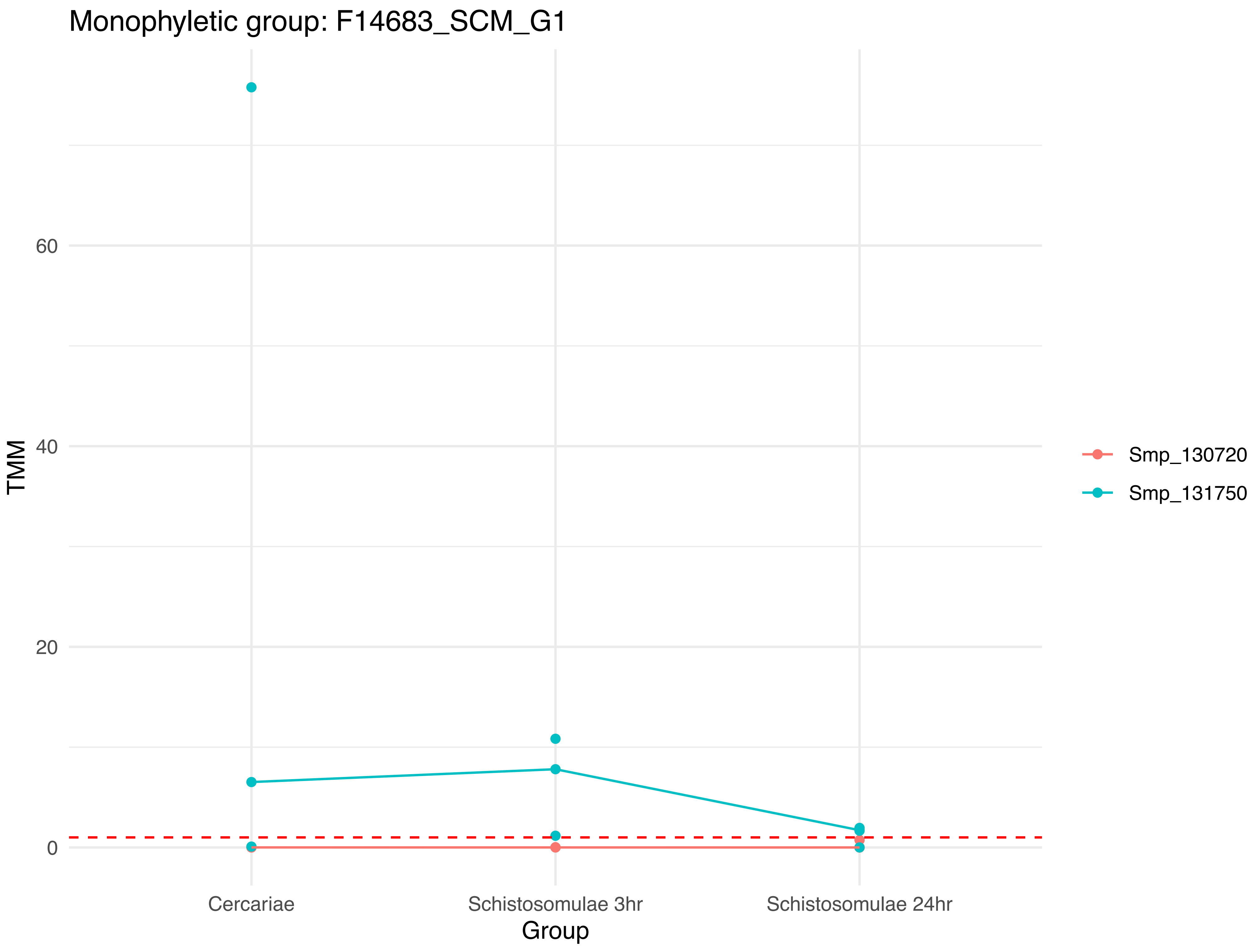

Monophyletic group: F14683_SCM_G1
TMM
60
40
20
0
Cercariae
Schistosomulae 3hr
Schistosomulae 24hr
Group
Smp_130720
Smp_131750

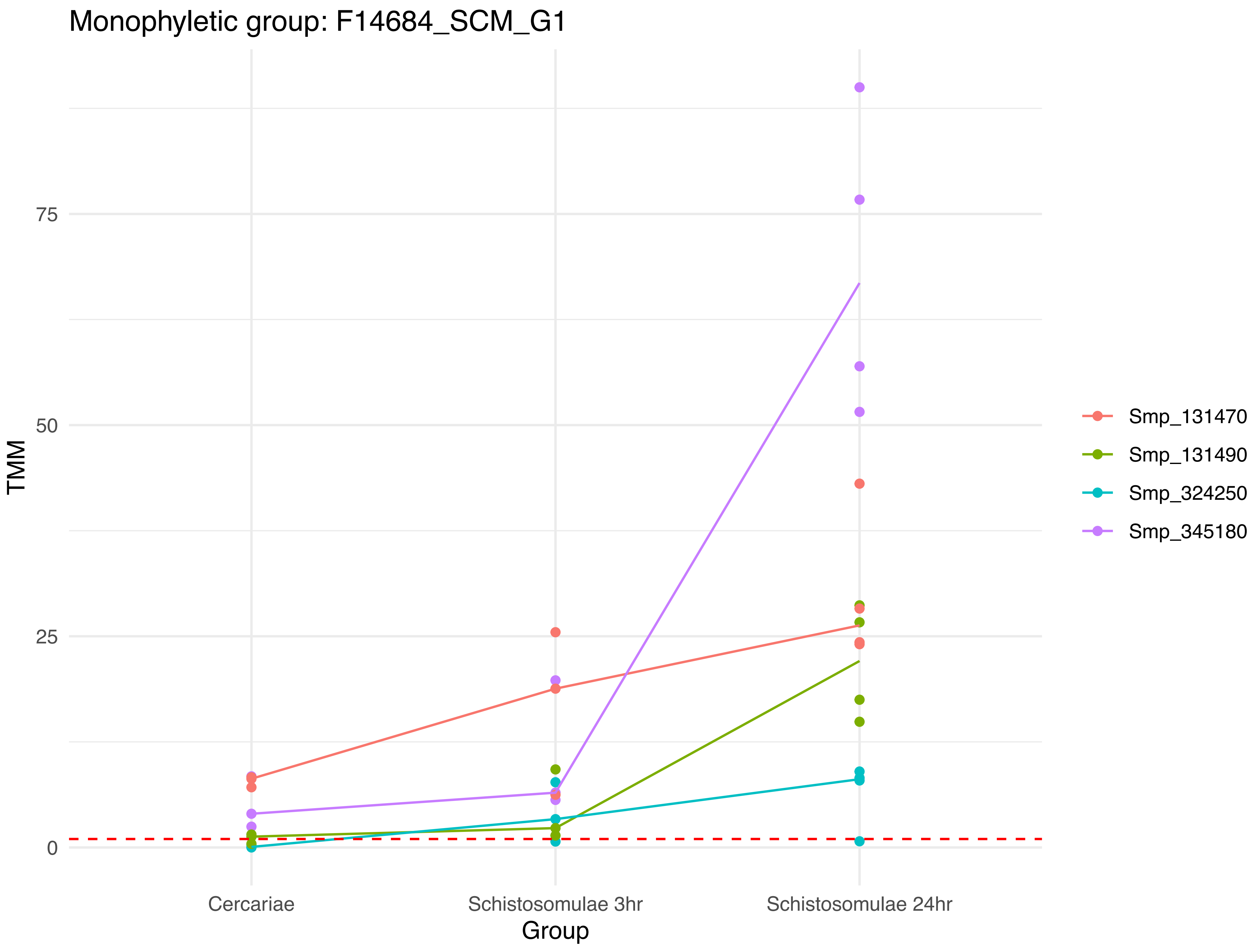

Monophyletic group: F14684_SCM_G1
TMM
75
50
25
0
Cercariae
Schistosomulae 3hr
Schistosomulae 24hr
Group
Smp_131470
Smp_131490
Smp_324250
Smp_345180

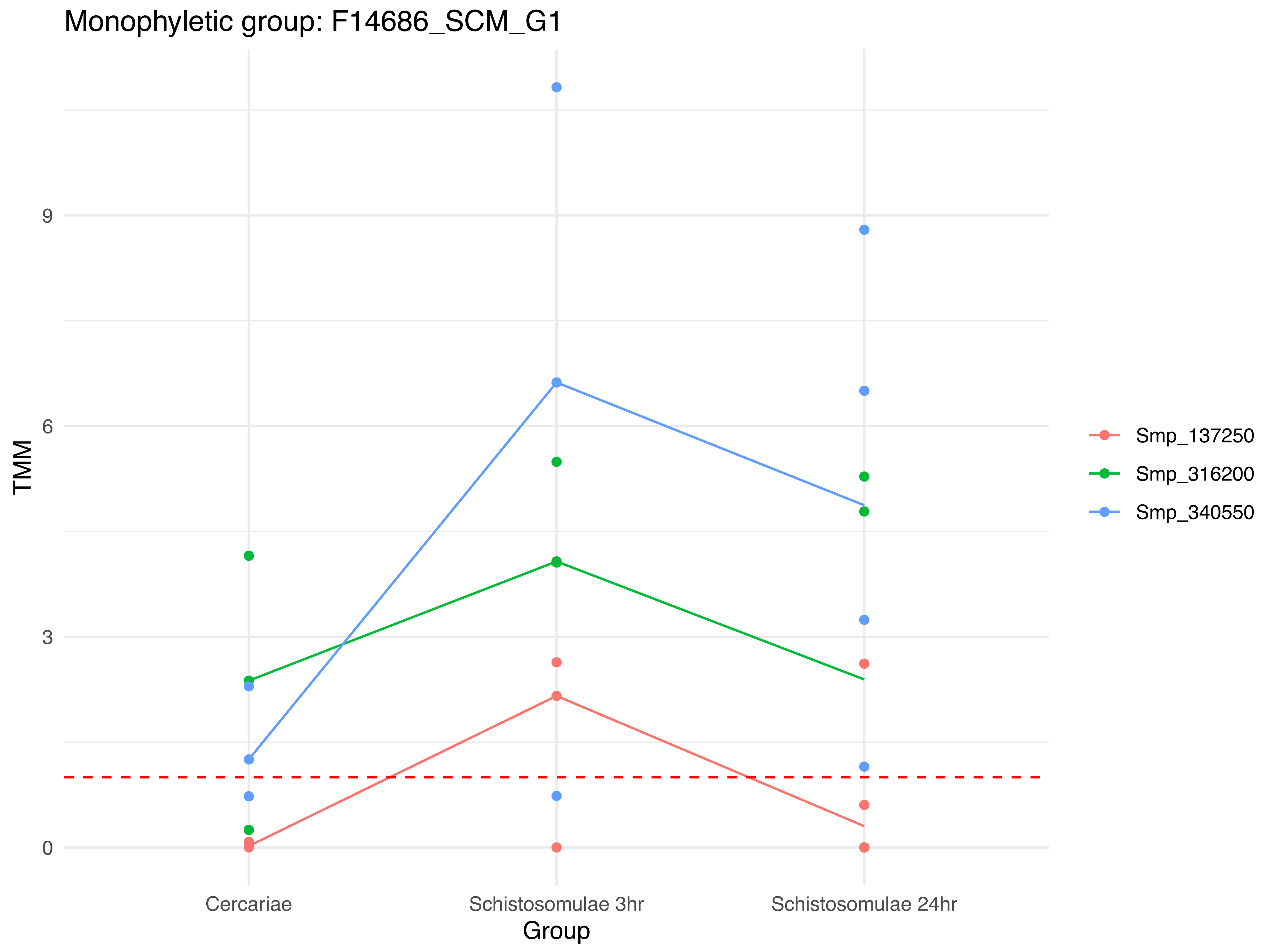

Monophyletic group: F14686_SCM_G1
TMM
9
6
3
0
Cercariae
Schistosomulae 3hr
Schistosomulae 24hr
Group
Smp_137250
Smp_316200
Smp_340550

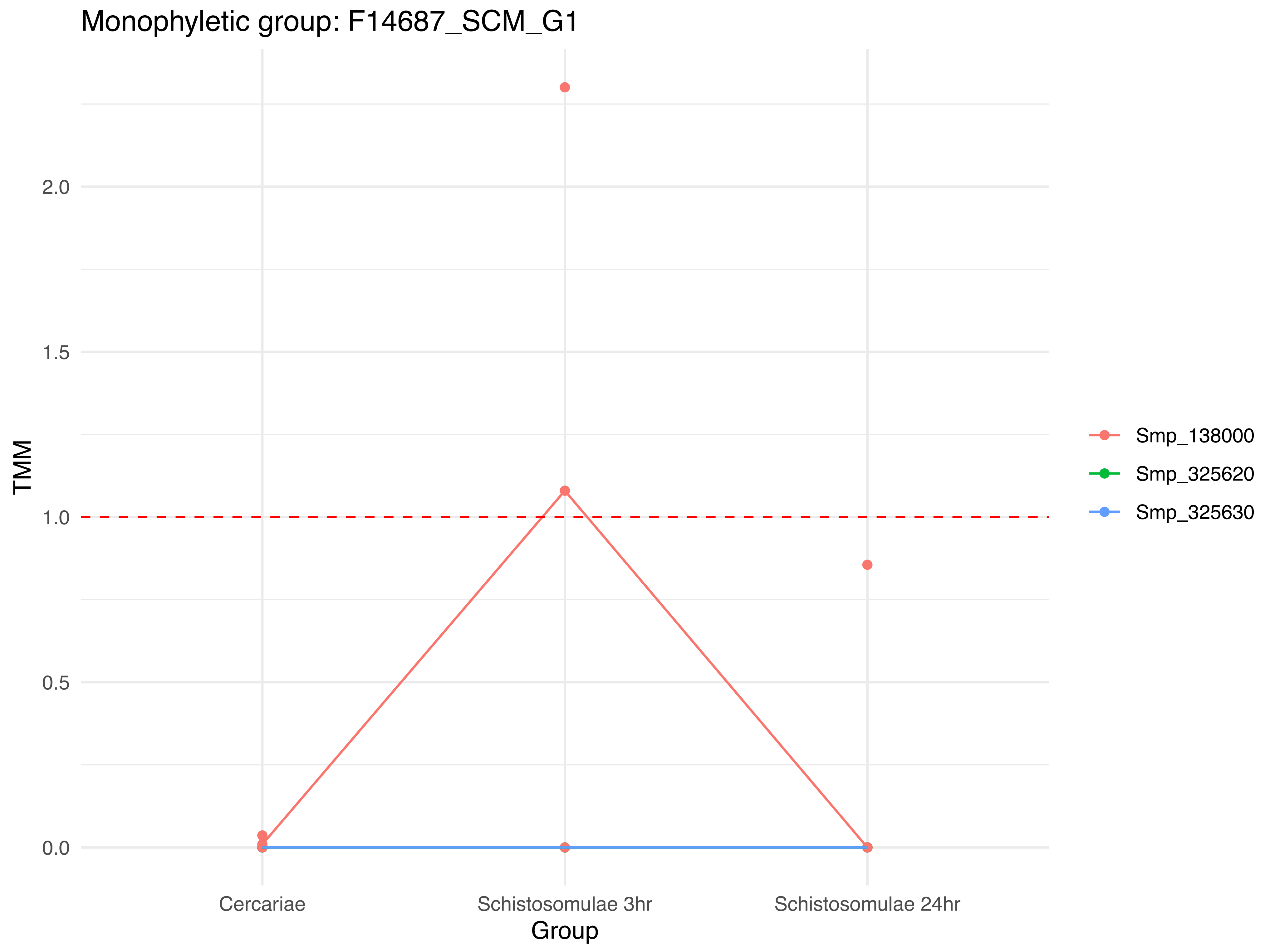

Monophyletic group: F14687_SCM_G1
TMM
2.0
1.5
1.0
0.5
0.0
Cercariae
Schistosomulae 3hr
Schistosomulae 24hr
Group
Smp_138000
Smp_325620
Smp_325630

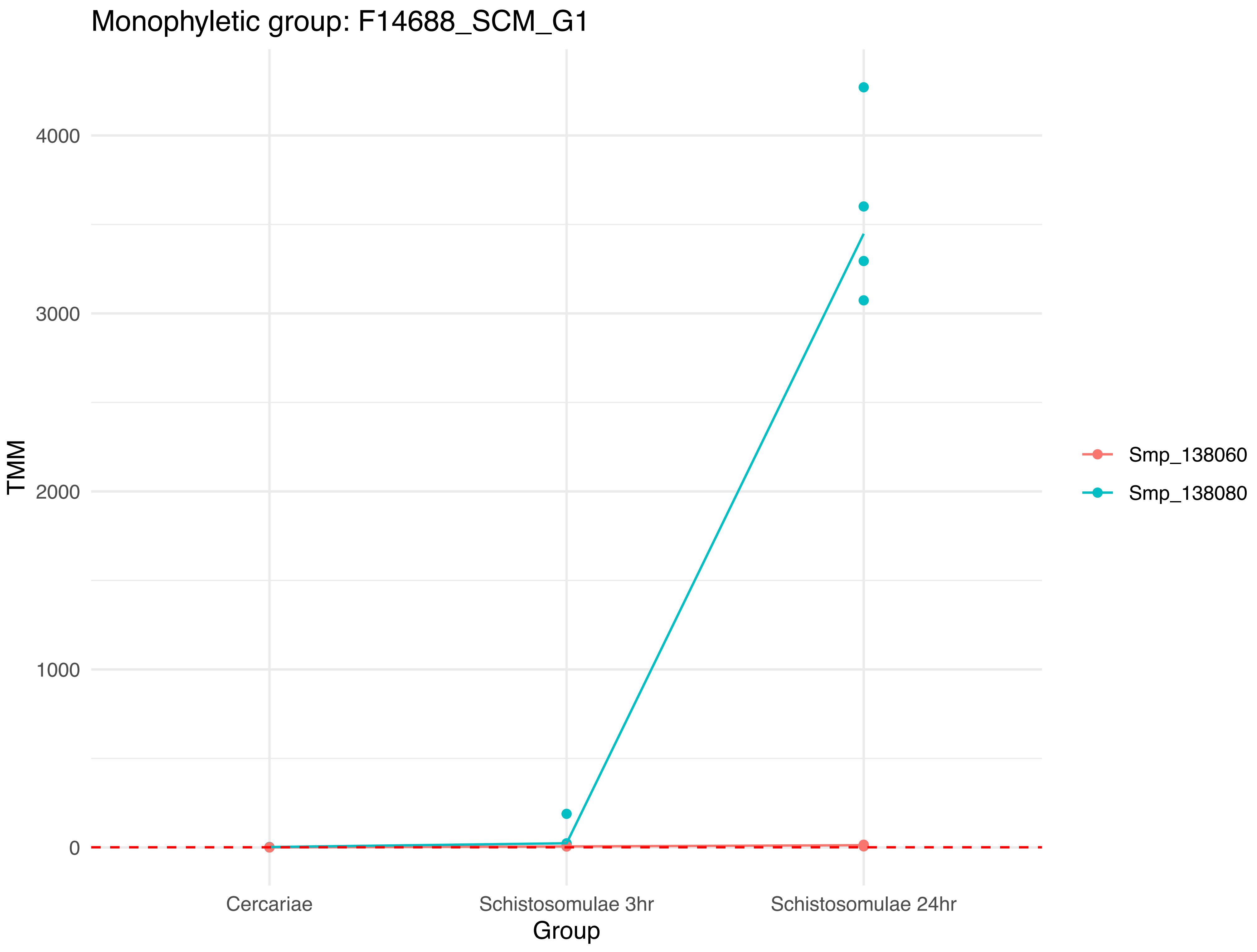

Monophyletic group: F14688_SCM_G1
4000
3000
2000
1000
0
TMM
Cercariae
Schistosomulae 3hr
Schistosomulae 24hr
Group
Smp_138060
Smp_138080

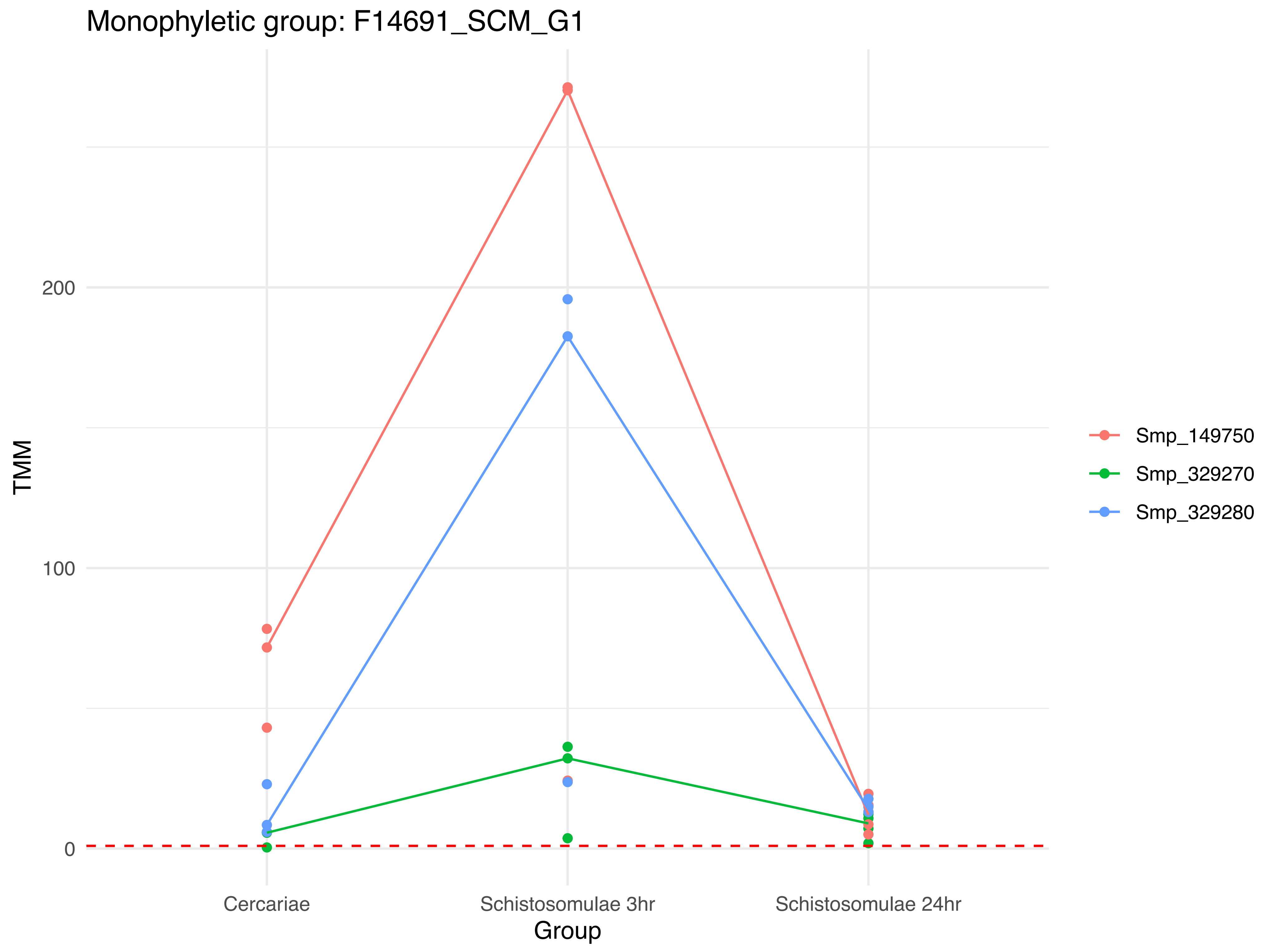

Monophyletic group: F14691_SCM_G1
TMM
200
100
0
Cercariae
Schistosomulae 3hr
Schistosomulae 24hr
Group
Smp_149750
Smp_329270
Smp_329280

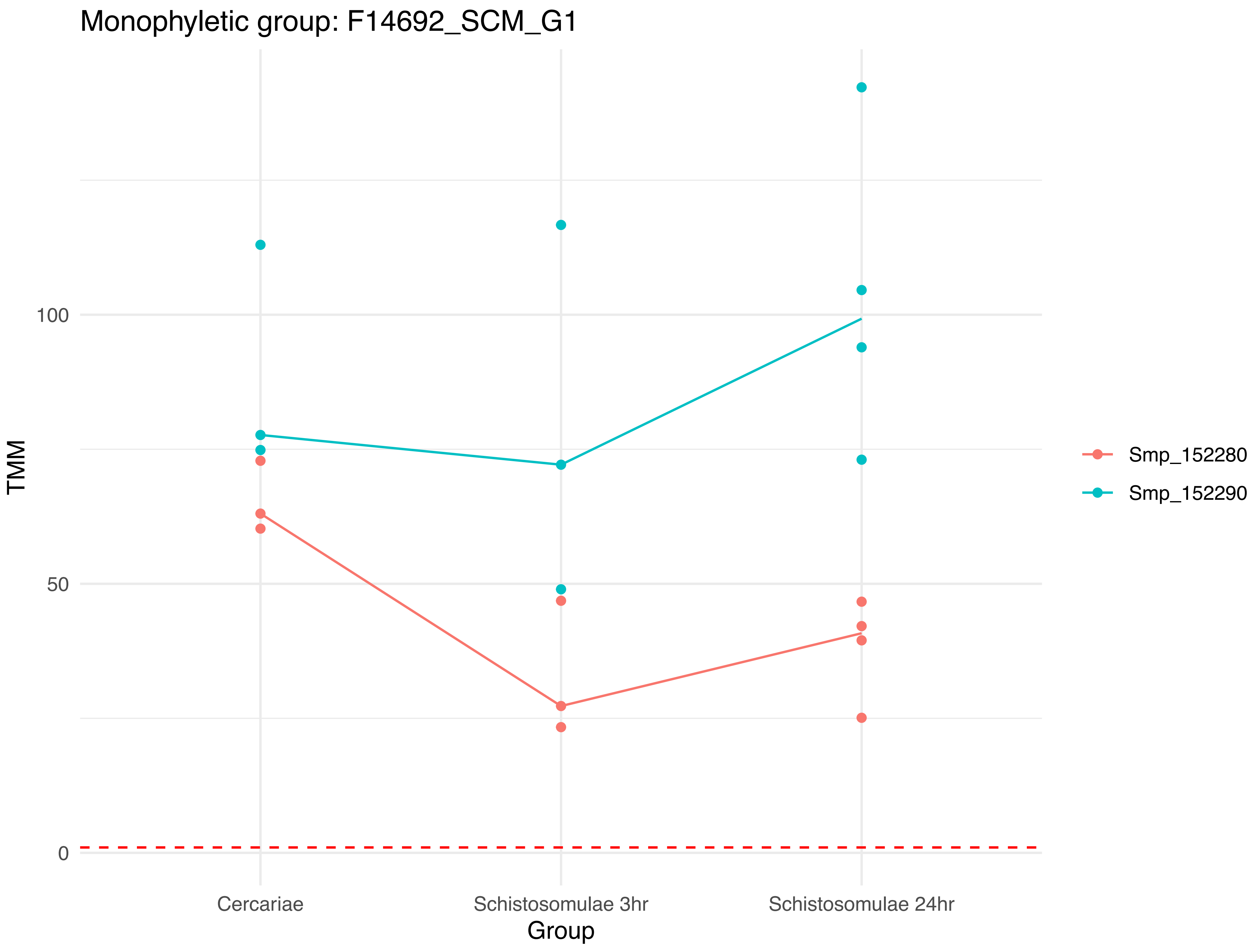
Monophyletic group: F14692_SCM_G1
TMM
100
50
0
Cercariae
Schistosomulae 3hr
Schistosomulae 24hr
Group
Smp_152280
Smp_152290

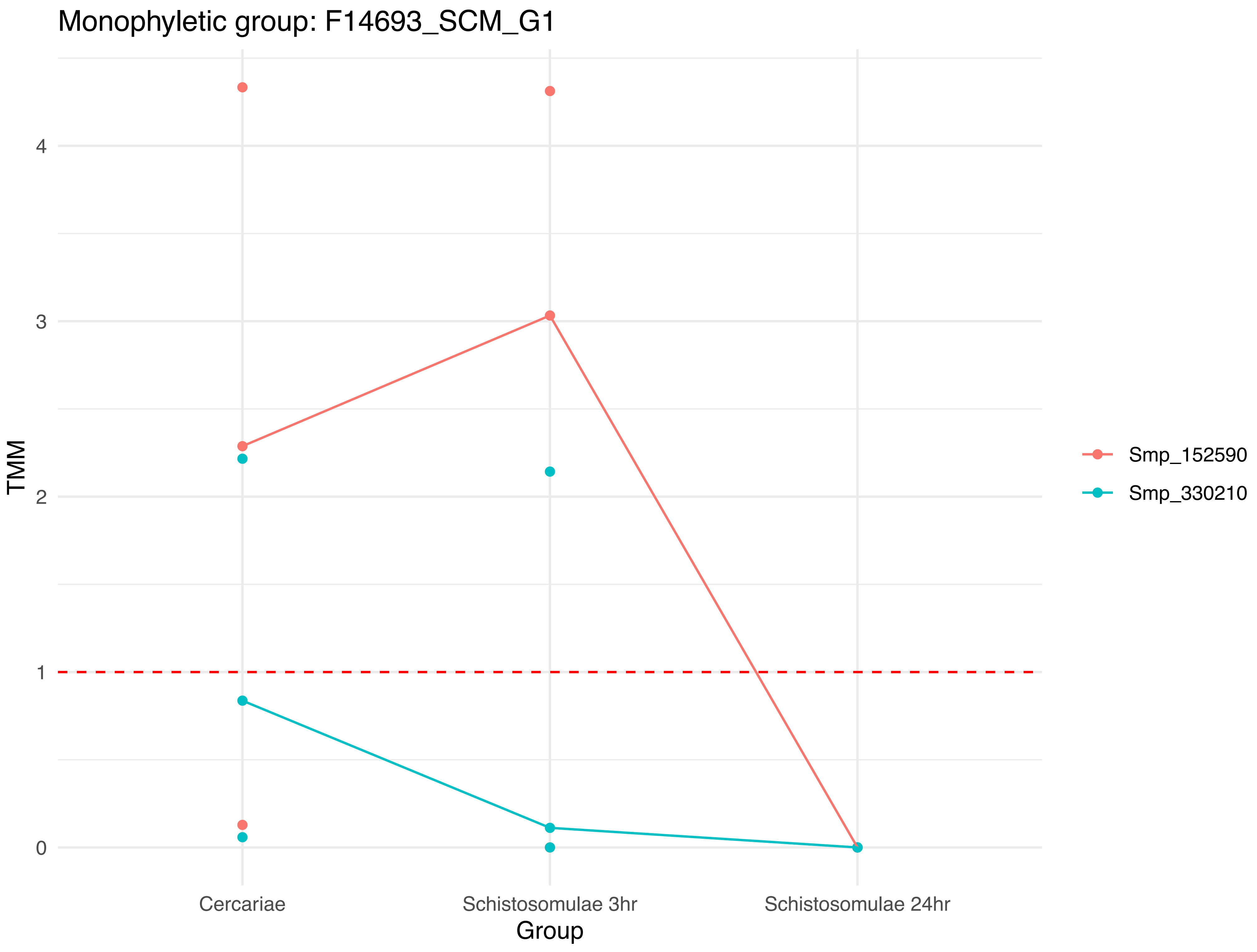

Monophyletic group: F14693_SCM_G1
TMM
4
3
2
1
0
Cercariae
Schistosomulae 3hr
Schistosomulae 24hr
Group
Smp_152590
Smp_330210

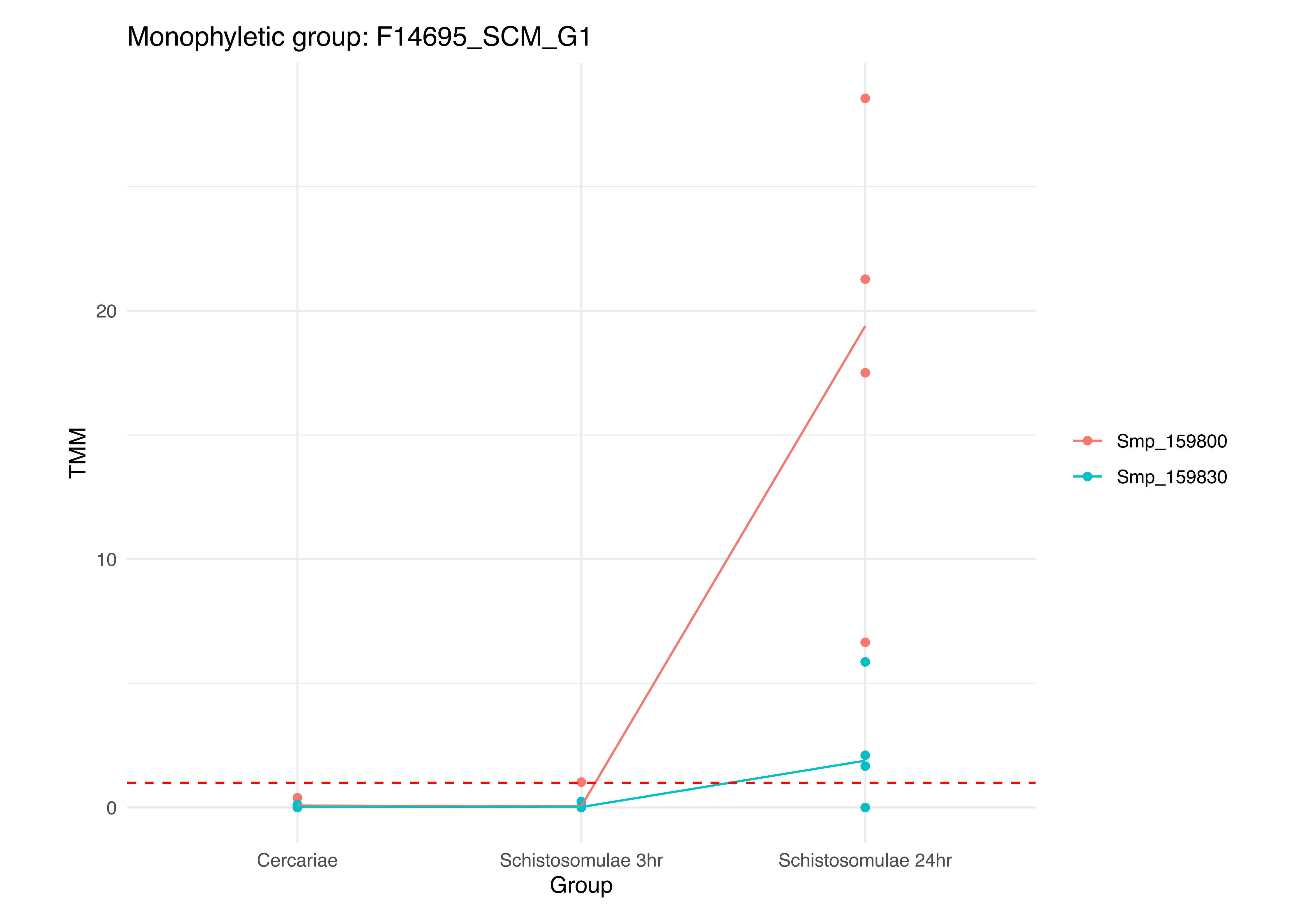

Monophyletic group: F14695_SCM_G1
TMM
20
10
0
Cercariae
Schistosomulae 3hr
Schistosomulae 24hr
Group
Smp_159800
Smp_159830

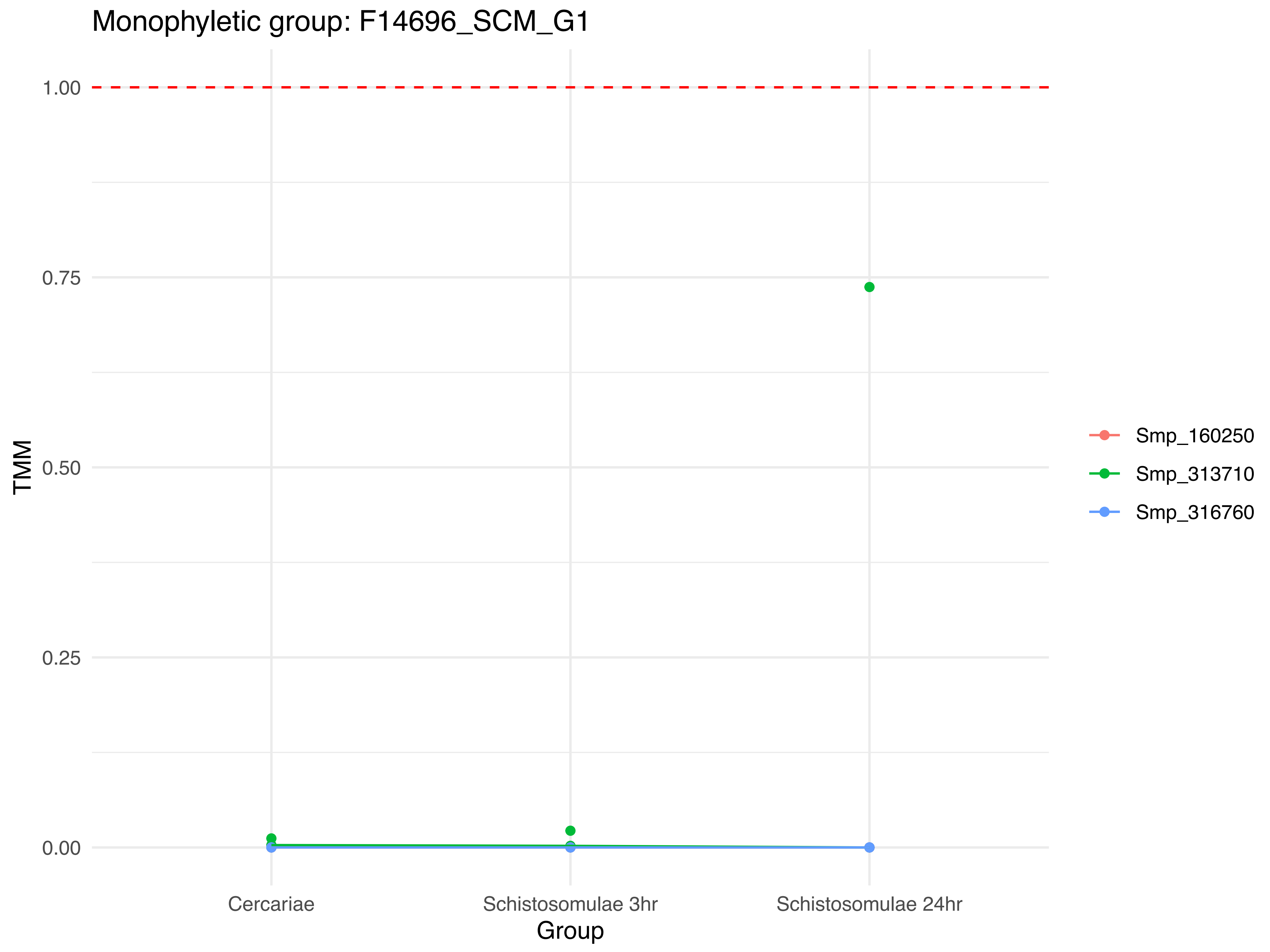

Monophyletic group: F14696_SCM_G1
1.00
0.75
0.50
0.25
0.00
TMM
Cercariae
Schistosomulae 3hr
Schistosomulae 24hr
Group
Smp_160250
Smp_313710
Smp_316760

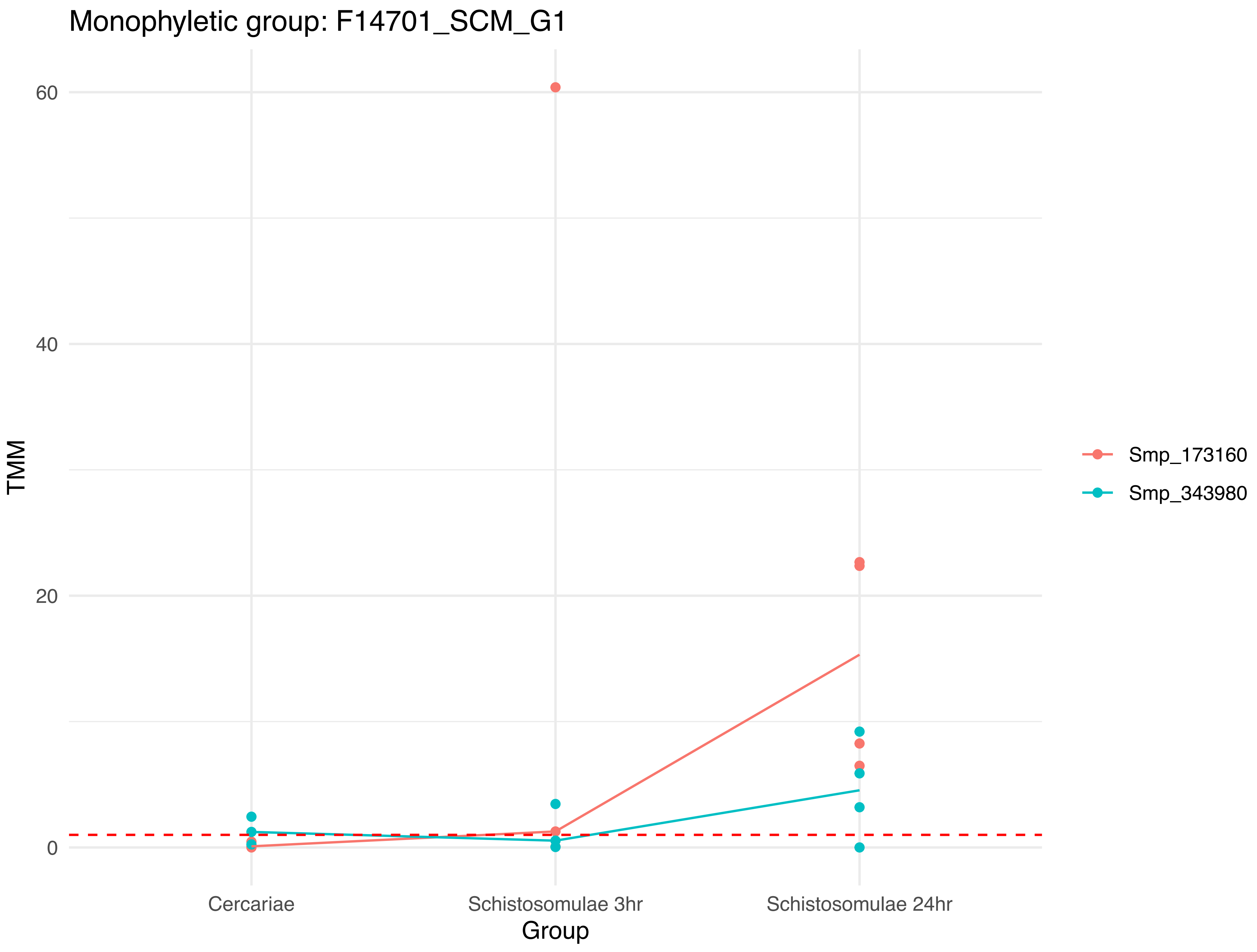

Monophyletic group: F14701_SCM_G1
TMM
60
40
20
0
Cercariae
Schistosomulae 3hr
Schistosomulae 24hr
Group
Smp_173160
Smp_343980

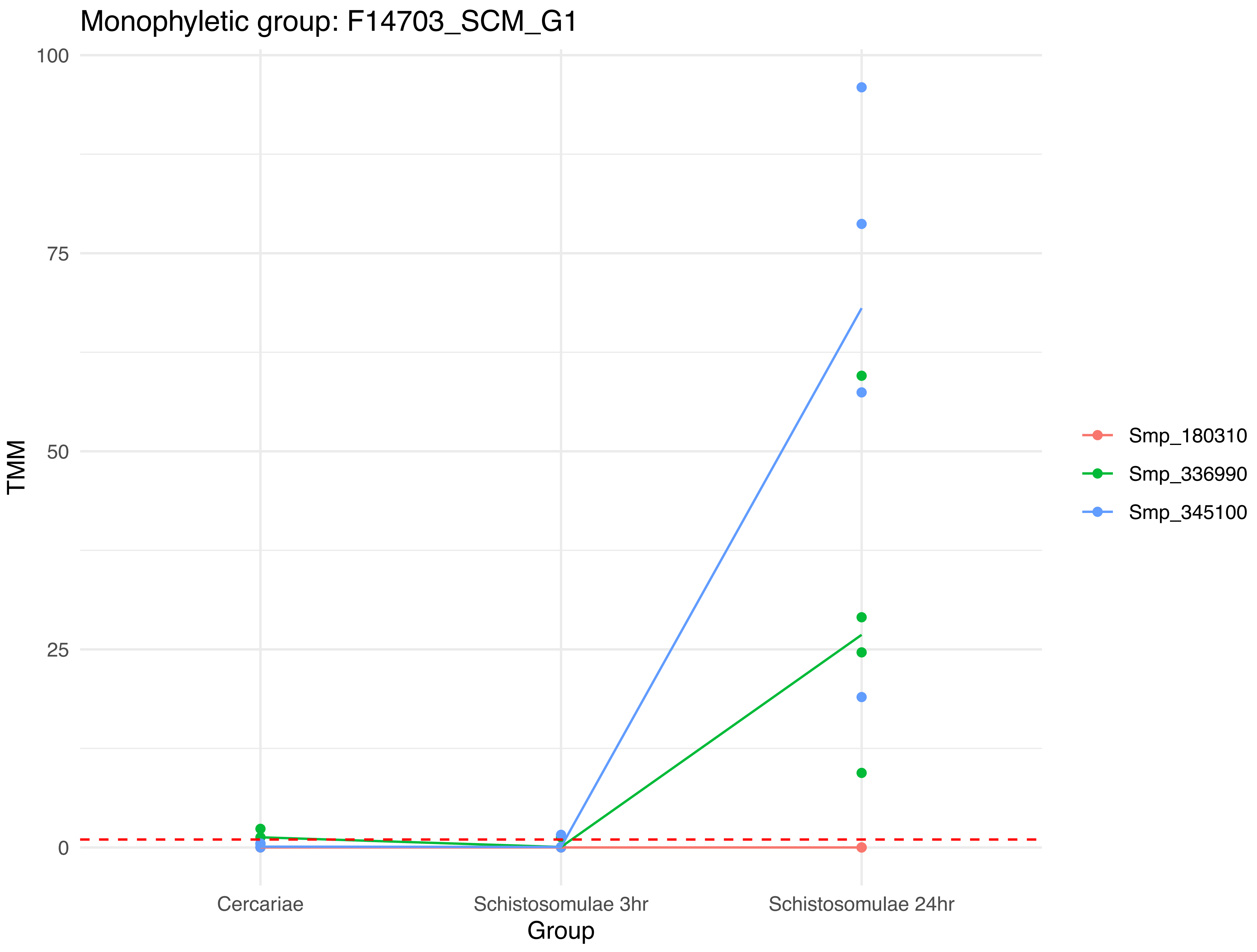
Monophyletic group: F14703_SCM_G1
100
75
50
25
0
TMM
Cercariae
Schistosomulae 3hr
Schistosomulae 24hr
Group
Smp_180310
Smp_336990
Smp_345100

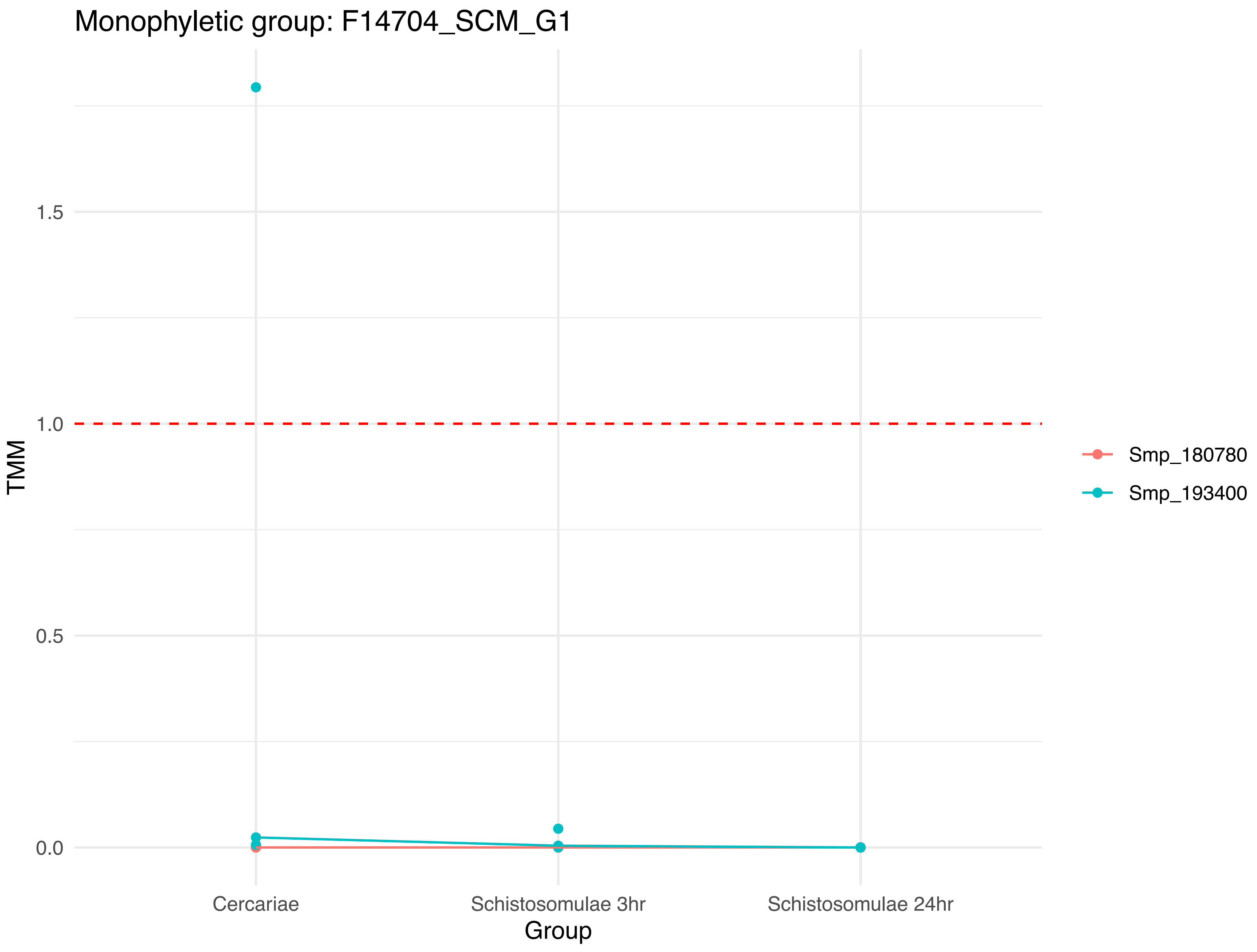
Monophyletic group: F14704_SCM_G1
TMM
1.5
1.0
0.5
0.0
Cercariae
Schistosomulae 3hr
Schistosomulae 24hr
Group
Smp_180780
Smp_193400

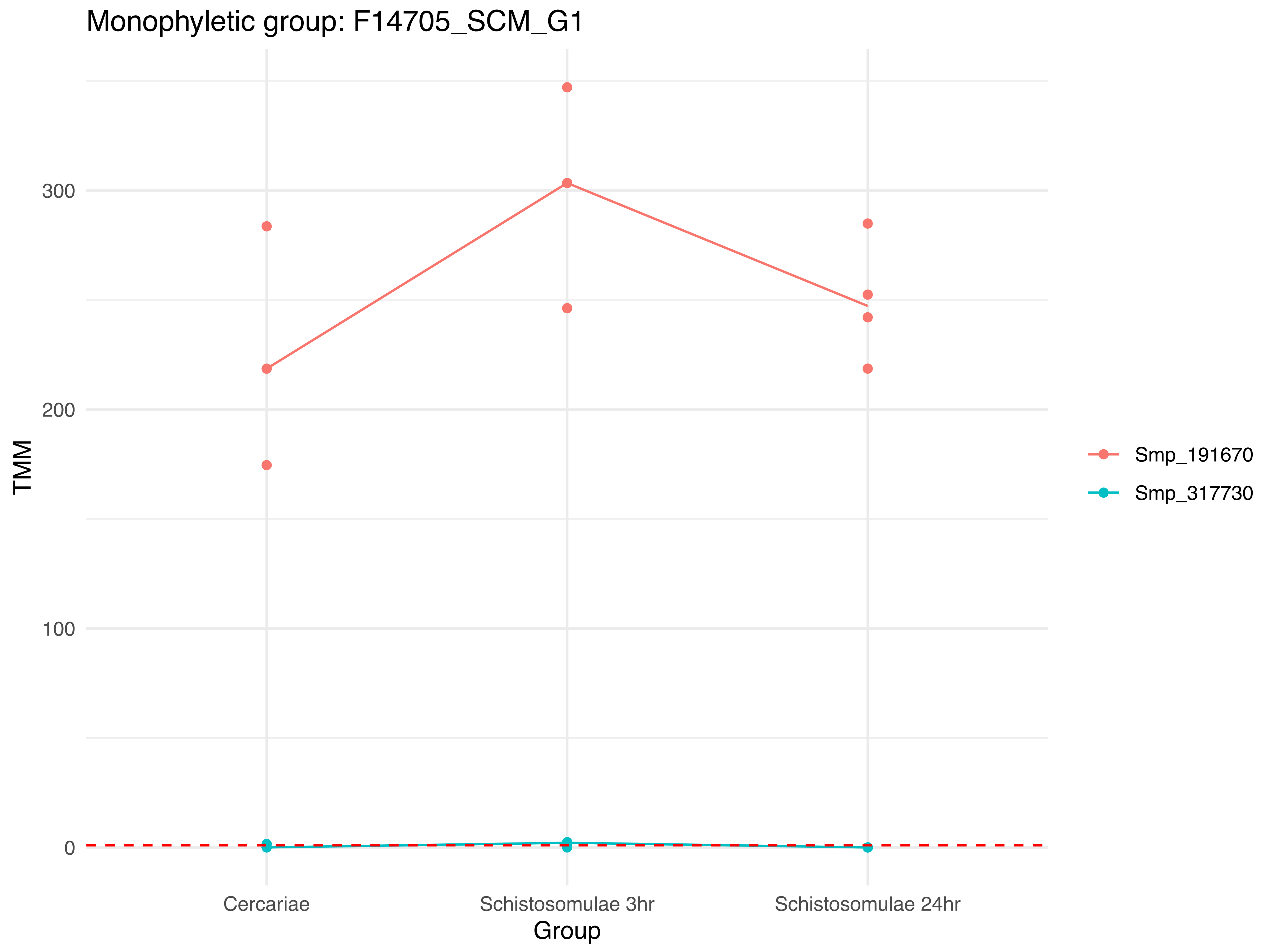
Monophyletic group: F14705_SCM_G1
TMM
300
200
100
0
Cercariae
Schistosomulae 3hr
Schistosomulae 24hr
Group
Smp_191670
Smp_317730

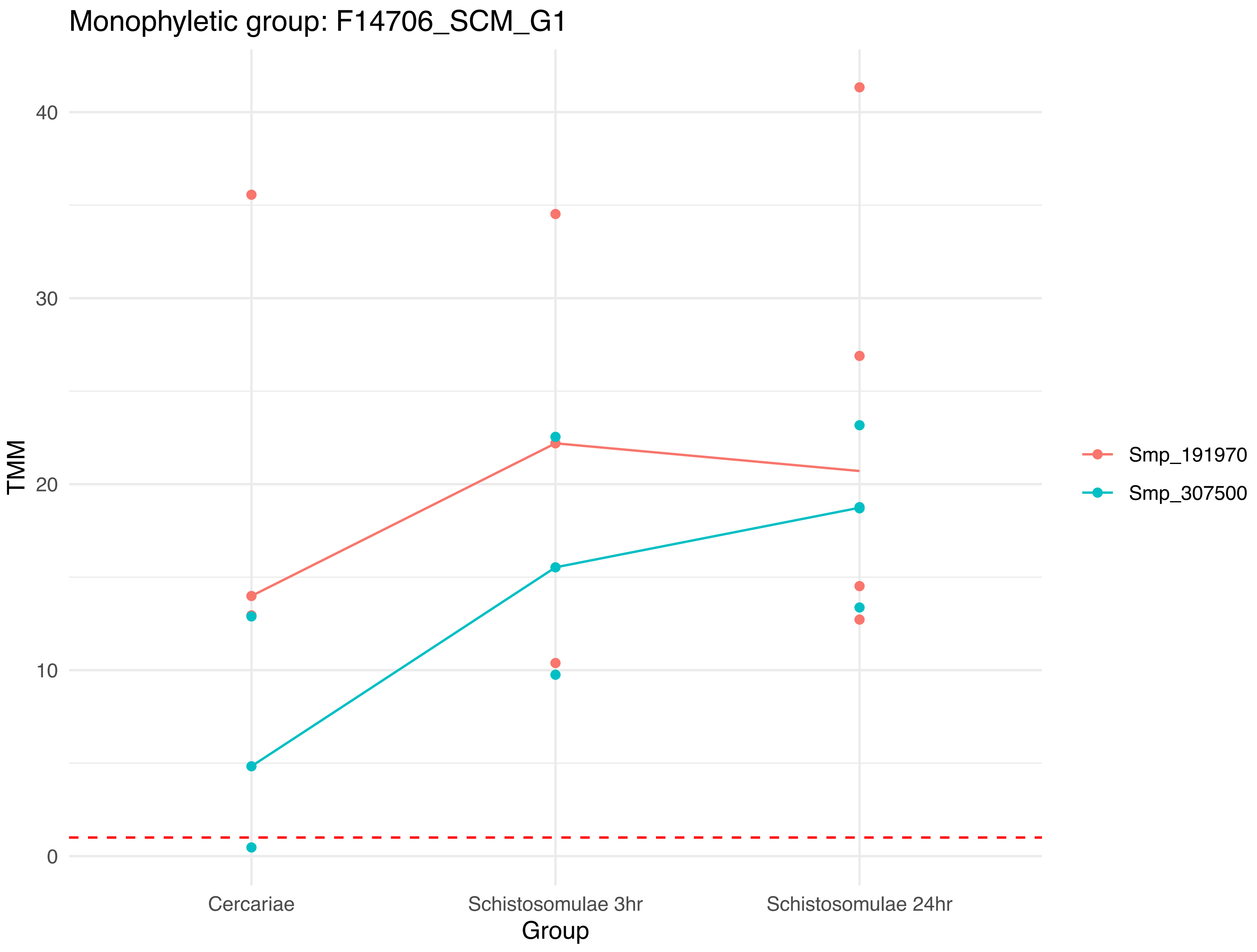
Monophyletic group: F14706_SCM_G1
TMM
40
30
20
10
0
Cercariae
Schistosomulae 3hr
Schistosomulae 24hr
Group
Smp_191970
Smp_307500

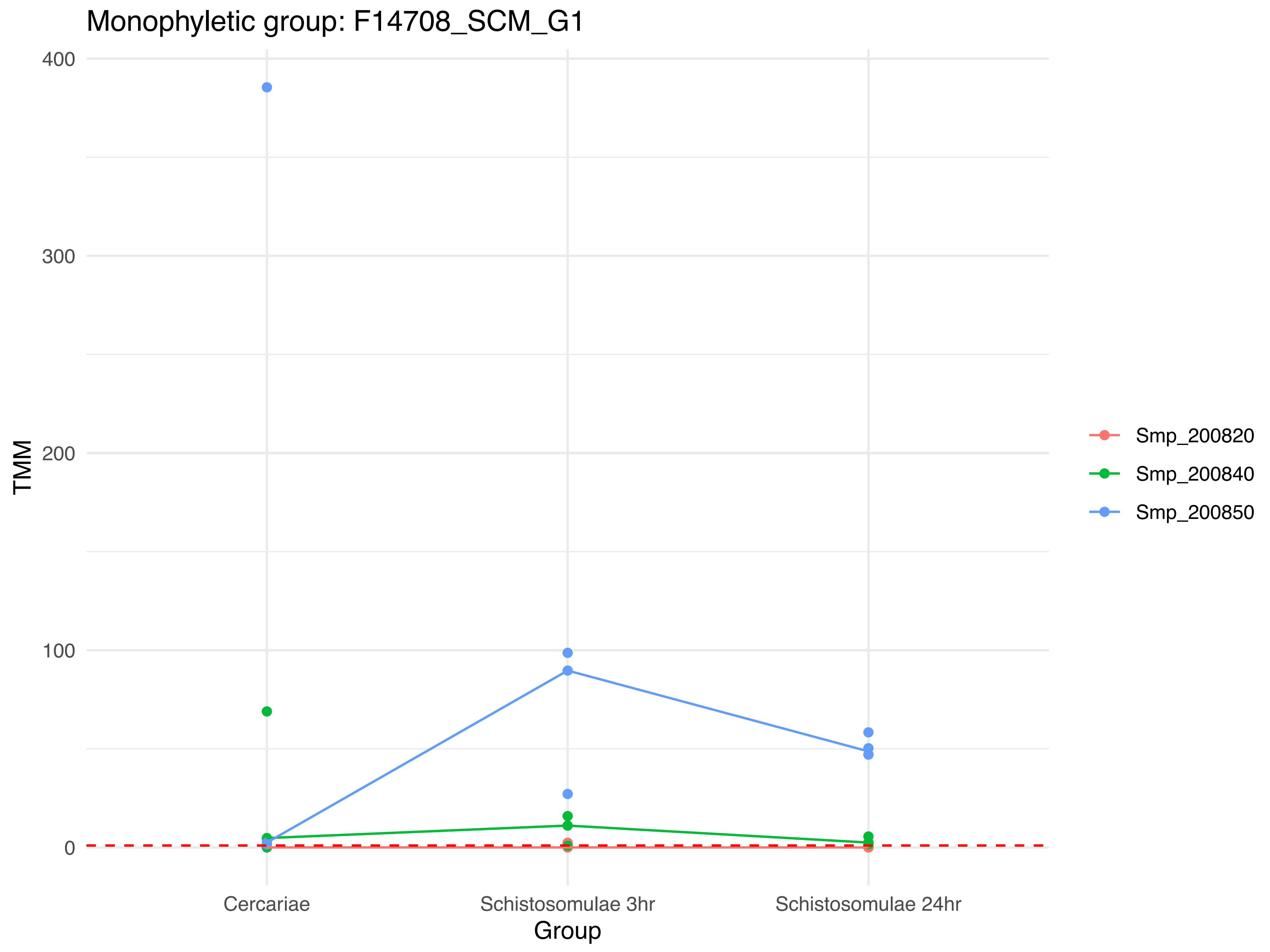

Monophyletic group: F14708_SCM_G1
400
300
200
100
0
TMM
Cercariae
Schistosomulae 3hr
Schistosomulae 24hr
Group
Smp_200820
Smp_200840
Smp_200850

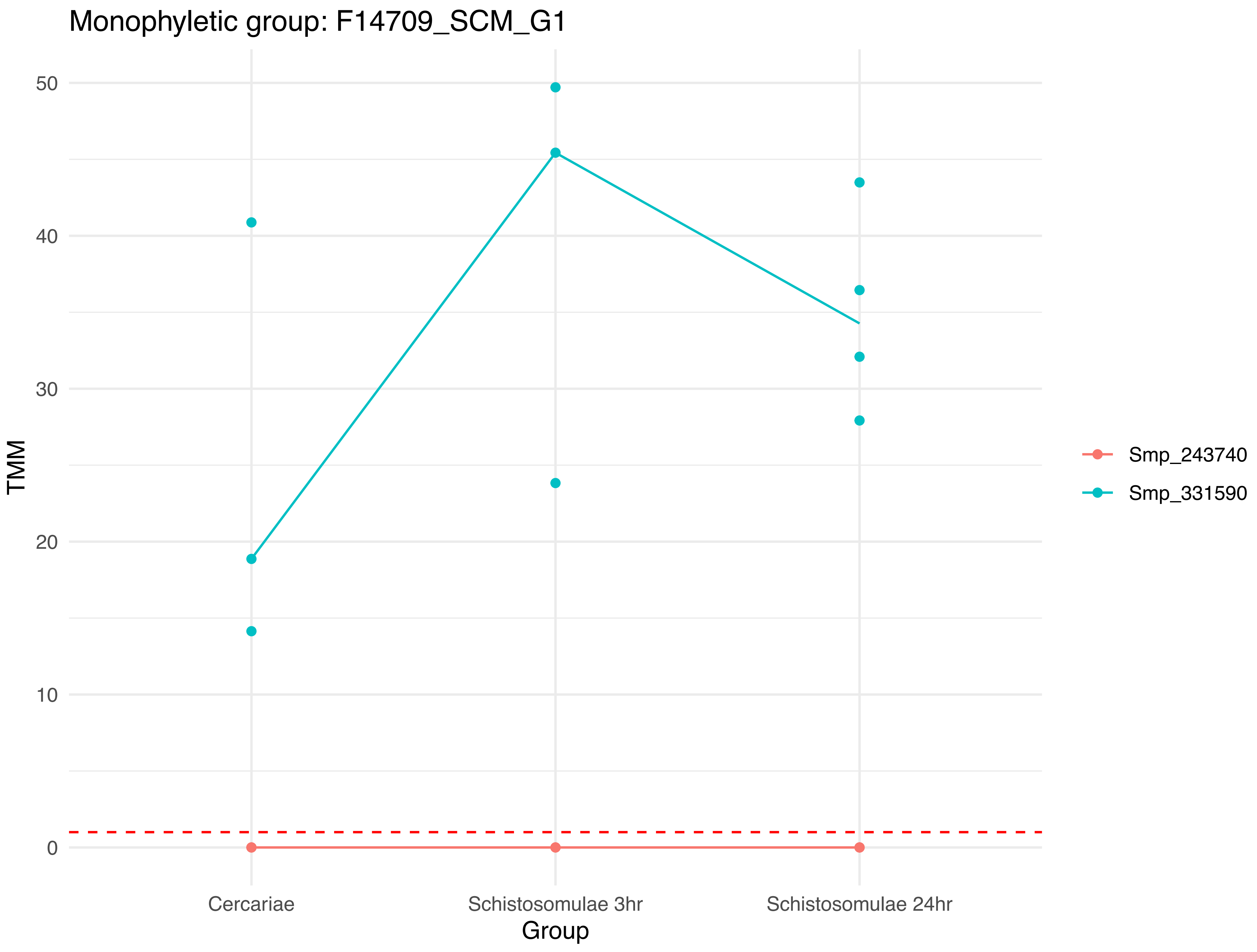

Monophyletic group: F14709_SCM_G1
TMM
50
40
30
20
10
0
Cercariae
Schistosomulae 3hr
Schistosomulae 24hr
Group
Smp_243740
Smp_331590

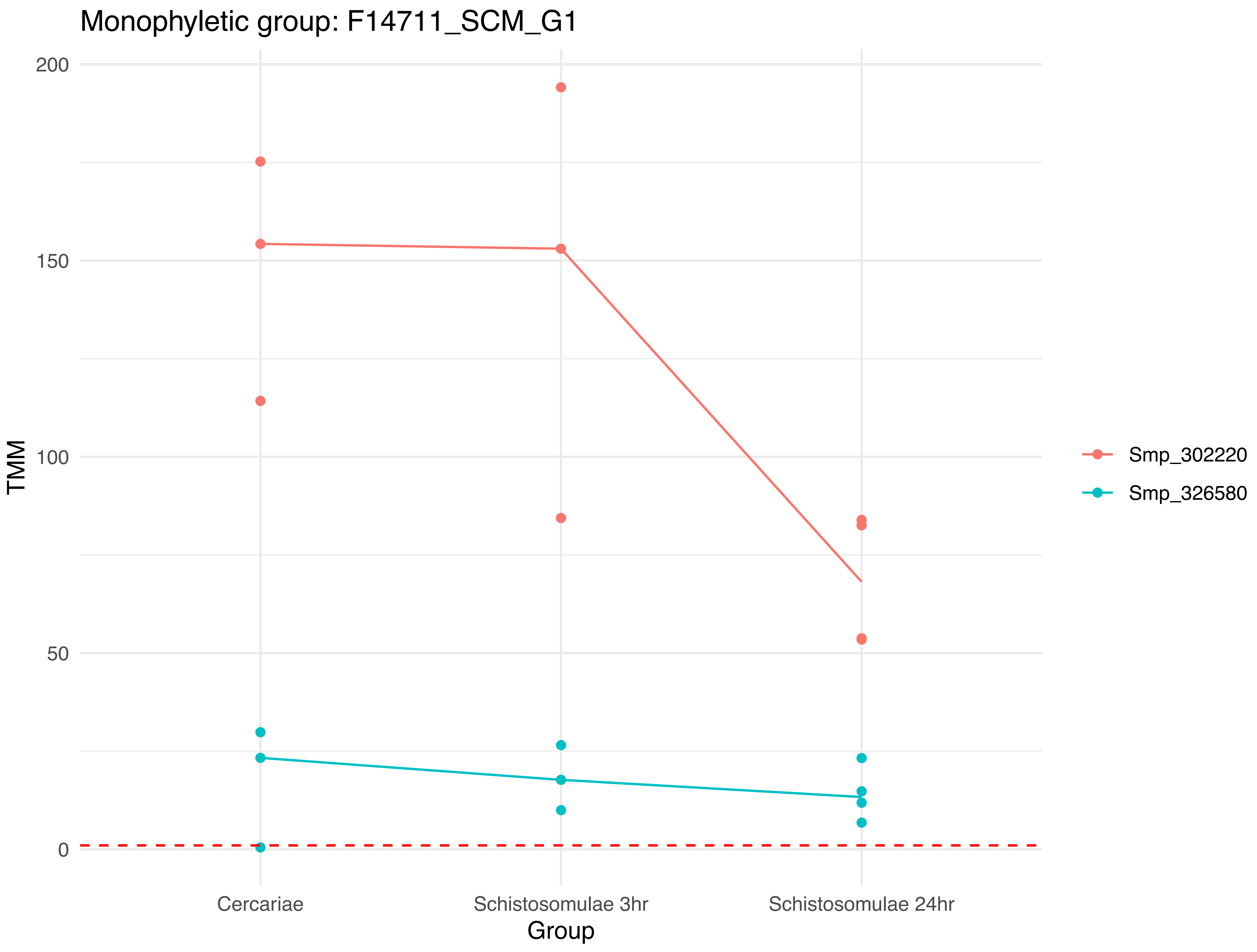
Monophyletic group: F14711_SCM_G1
TMM
200
150
100
50
0
Cercariae
Schistosomulae 3hr
Schistosomulae 24hr
Group
Smp_302220
Smp_326580

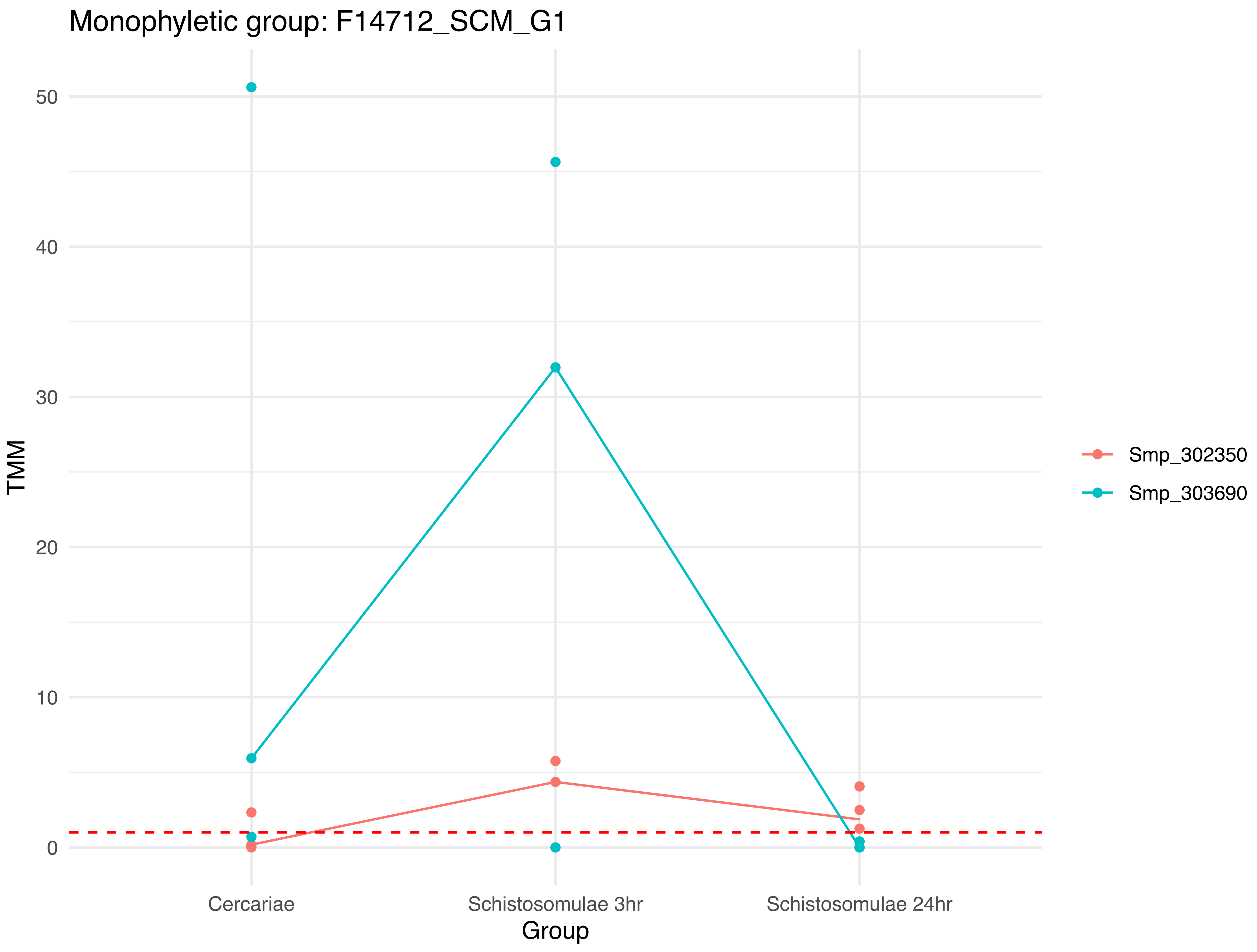
Monophyletic group: F14712_SCM_G1
TMM
50
40
30
20
10
0
Cercariae
Schistosomulae 3hr
Schistosomulae 24hr
Group
Smp_302350
Smp_303690

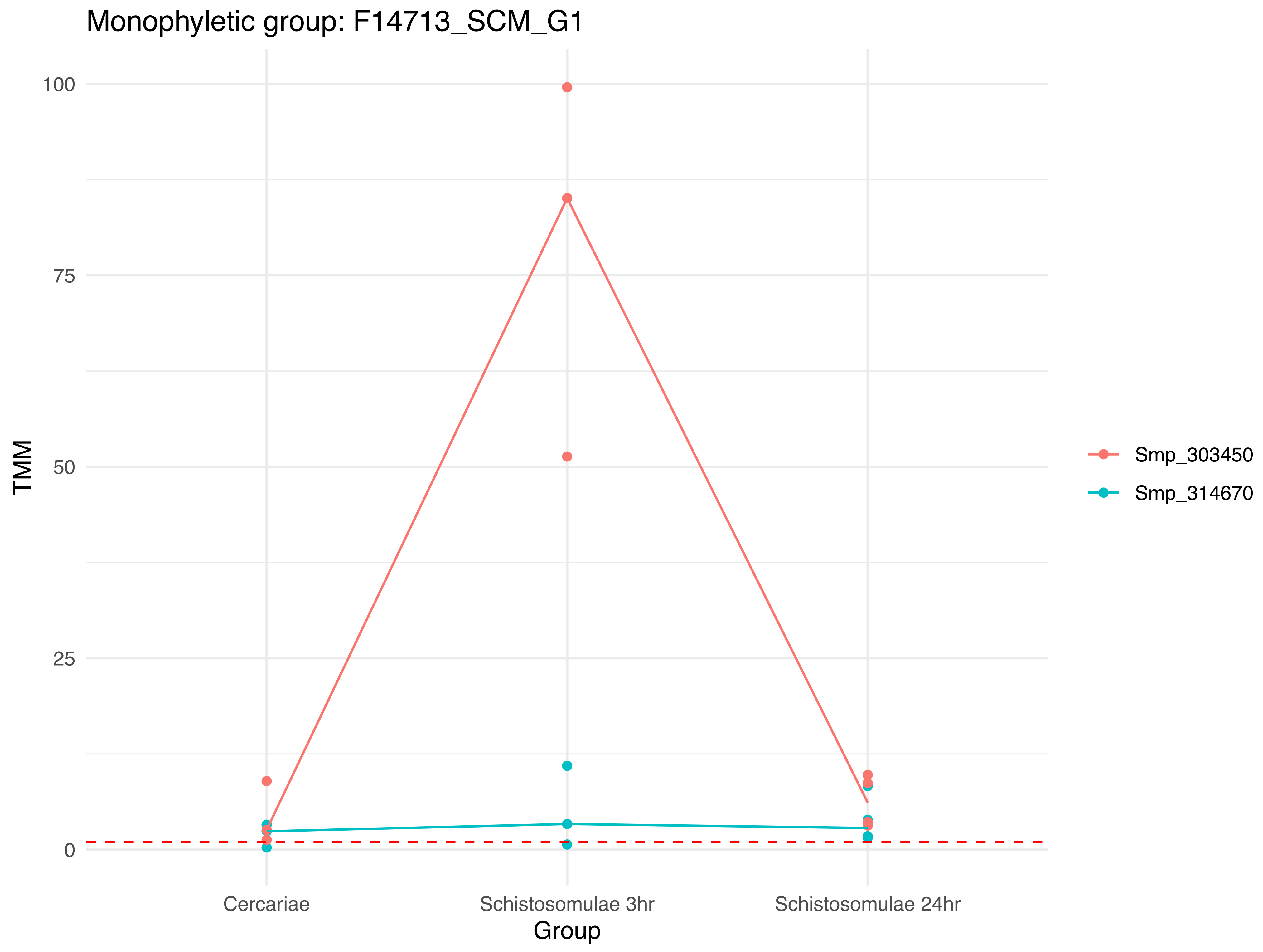
Monophyletic group: F14713_SCM_G1
TMM
100
75
50
25
0
Cercariae
Schistosomulae 3hr
Schistosomulae 24hr
Group
Smp_303450
Smp_314670

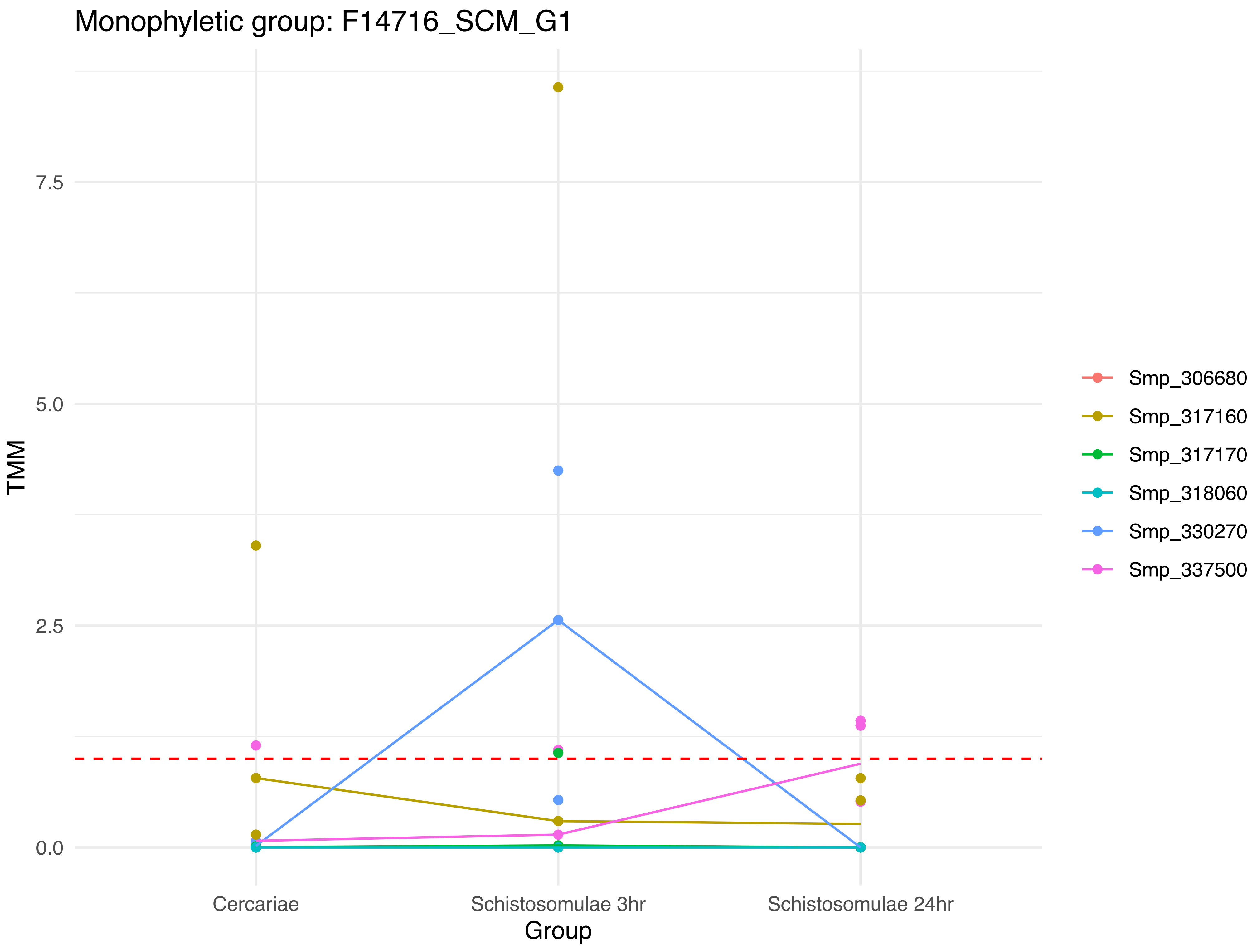

Monophyletic group: F14716_SCM_G1
TMM
0.0
2.5
5.0
7.5
Cercariae
Schistosomulae 3hr
Schistosomulae 24hr
Group
Smp_306680
Smp_317160
Smp_317170
Smp_318060
Smp_330270
Smp_337500

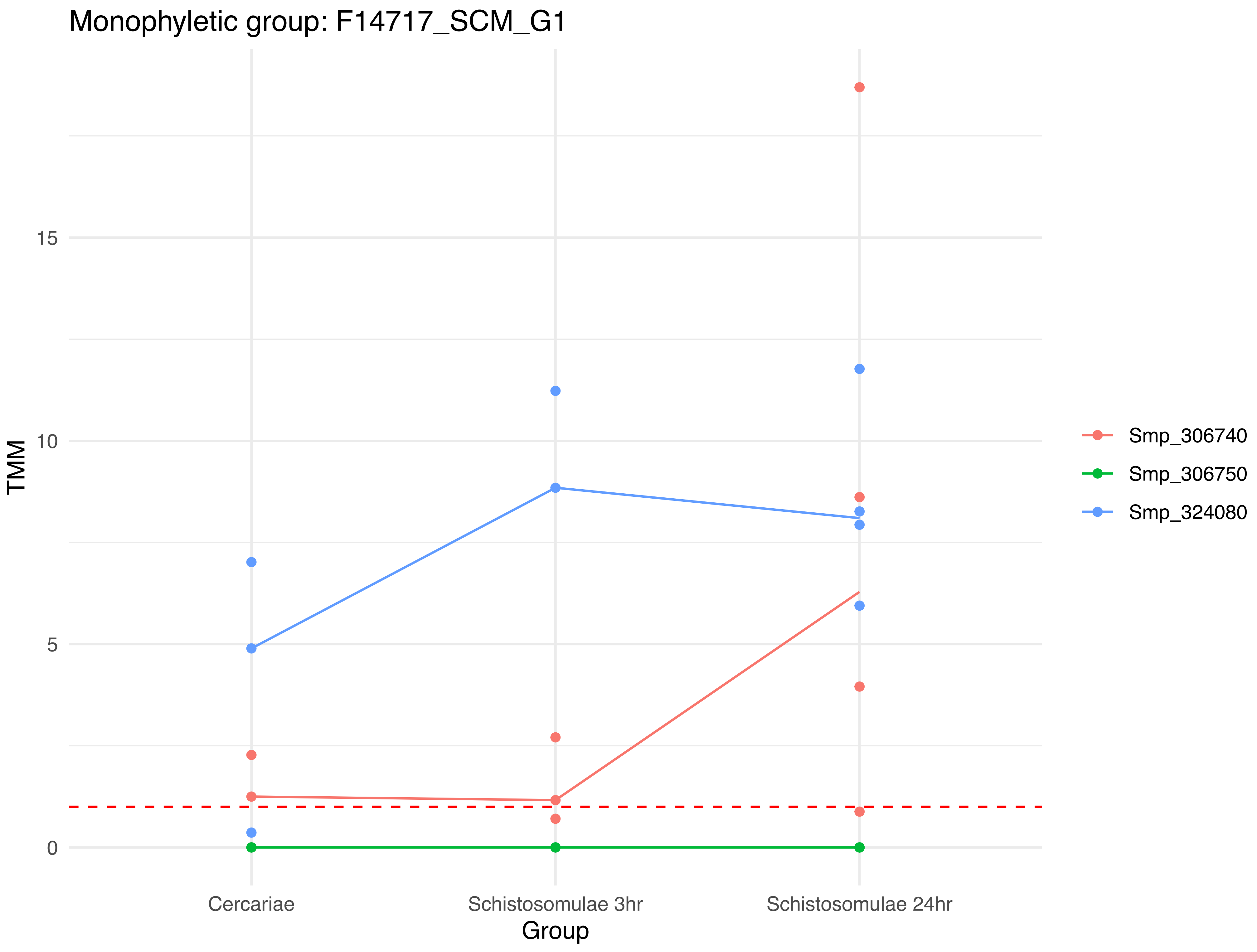

Monophyletic group: F14717_SCM_G1
15
10
5
0
TMM
Cercariae
Schistosomulae 3hr
Schistosomulae 24hr
Group
Smp_306740
Smp_306750
Smp_324080

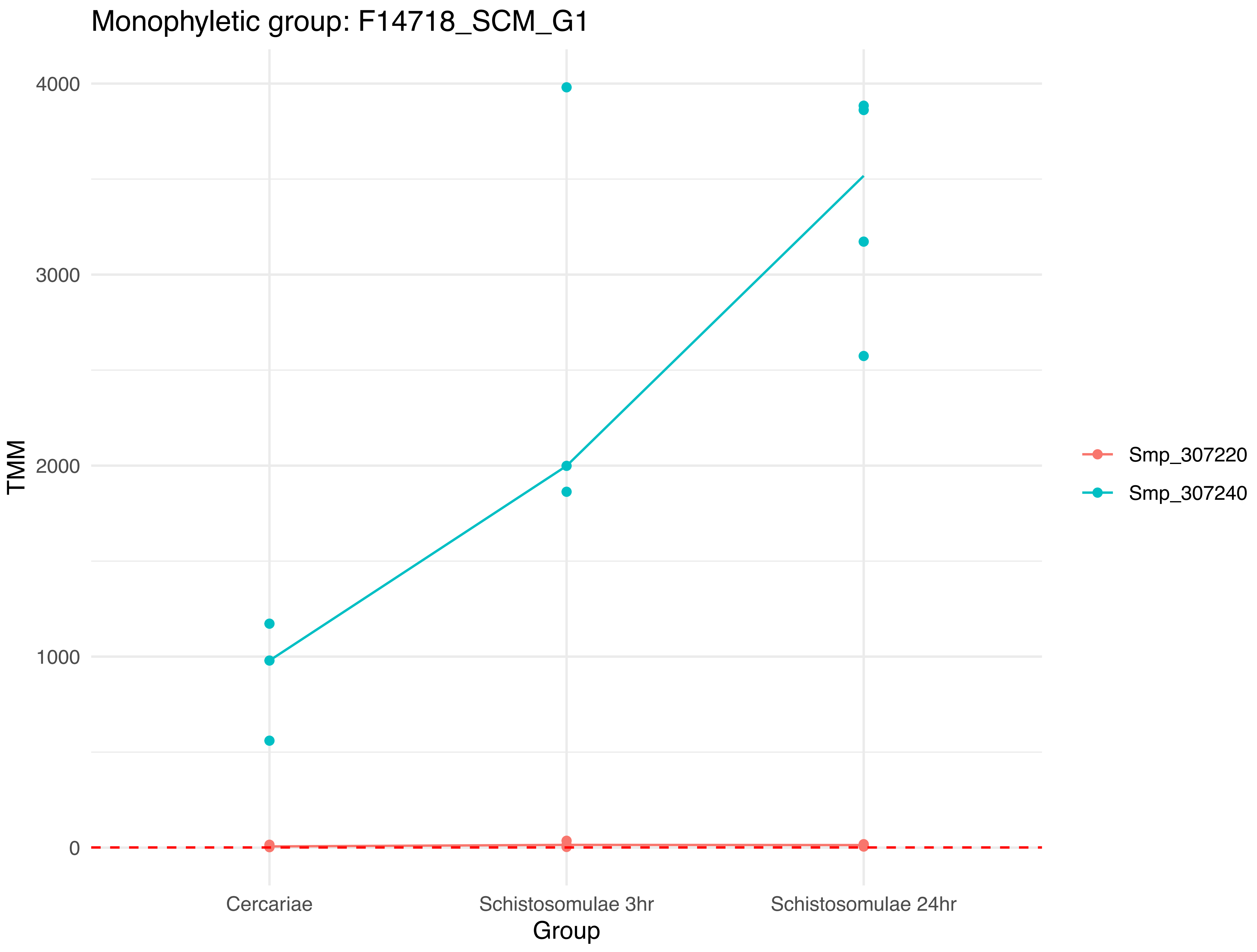

Monophyletic group: F14718_SCM_G1
4000
3000
2000
1000
0
TMM
Cercariae
Schistosomulae 3hr
Schistosomulae 24hr
Group
Smp_307220
Smp_307240

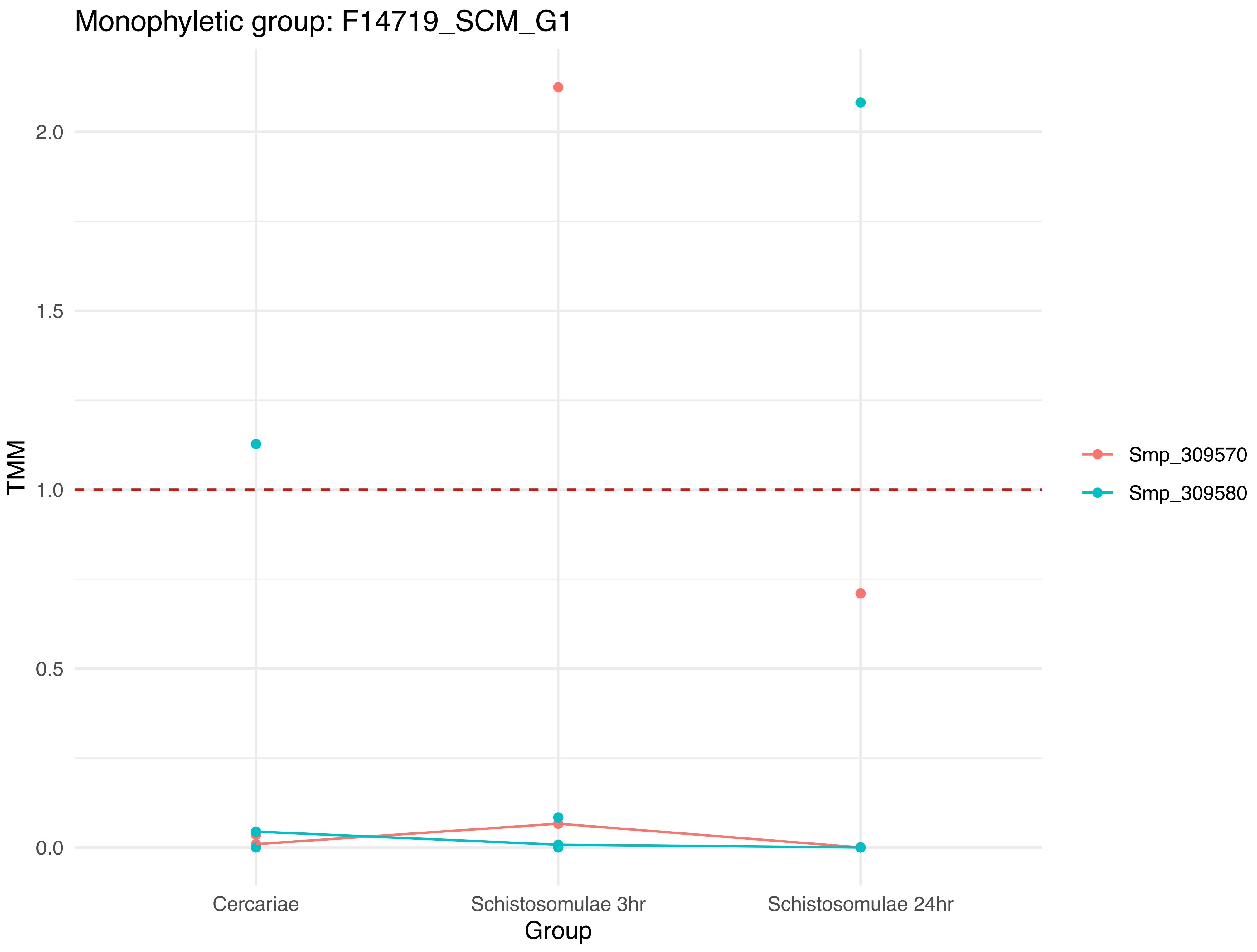
Monophyletic group: F14719_SCM_G1
TMM
2.0
1.5
1.0
0.5
0.0
Cercariae
Schistosomulae 3hr
Schistosomulae 24hr
Group
Smp_309570
Smp_309580

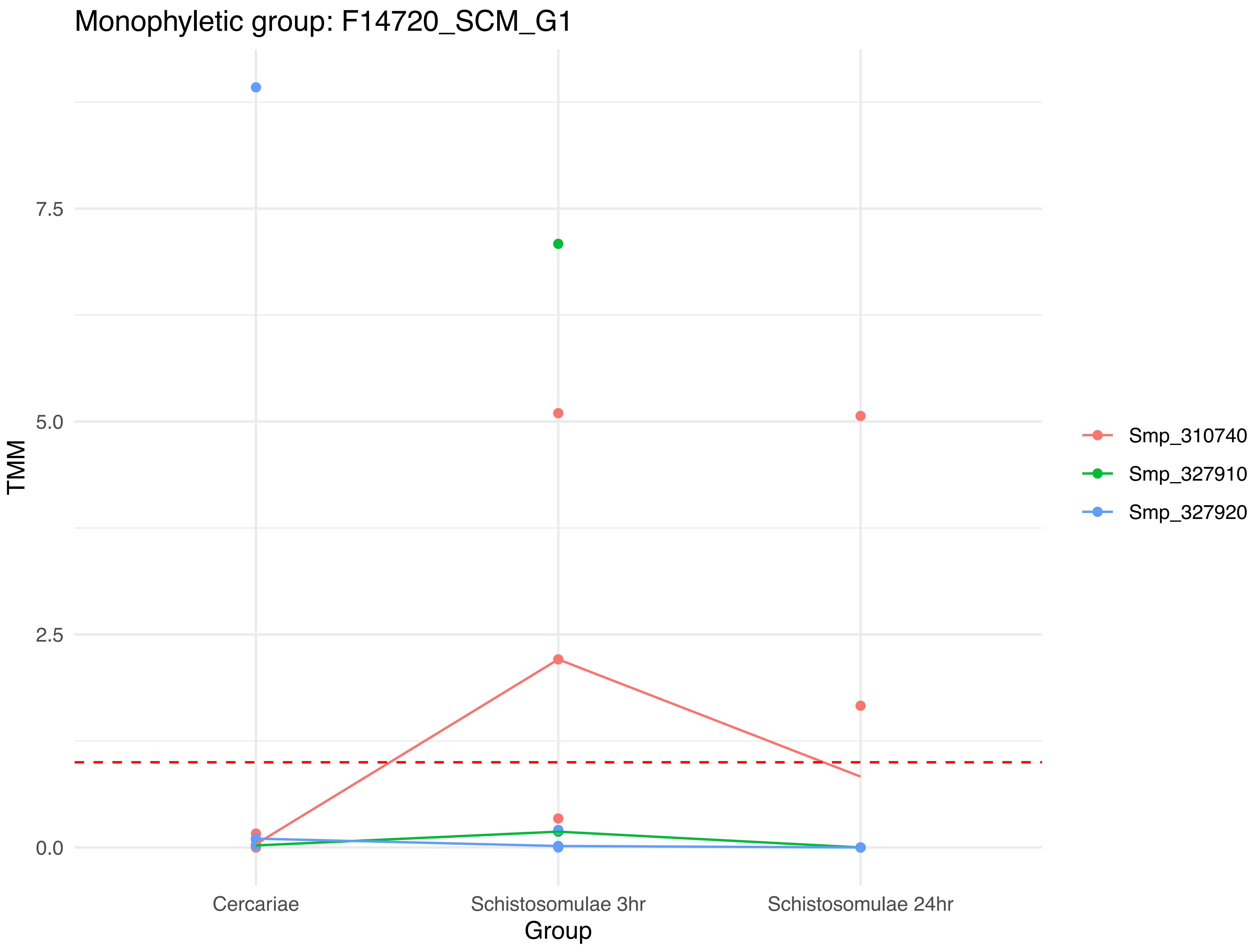

Monophyletic group: F14720_SCM_G1
TMM
0.0
2.5
5.0
7.5
Cercariae
Schistosomulae 3hr
Schistosomulae 24hr
Group
Smp_310740
Smp_327910
Smp_327920

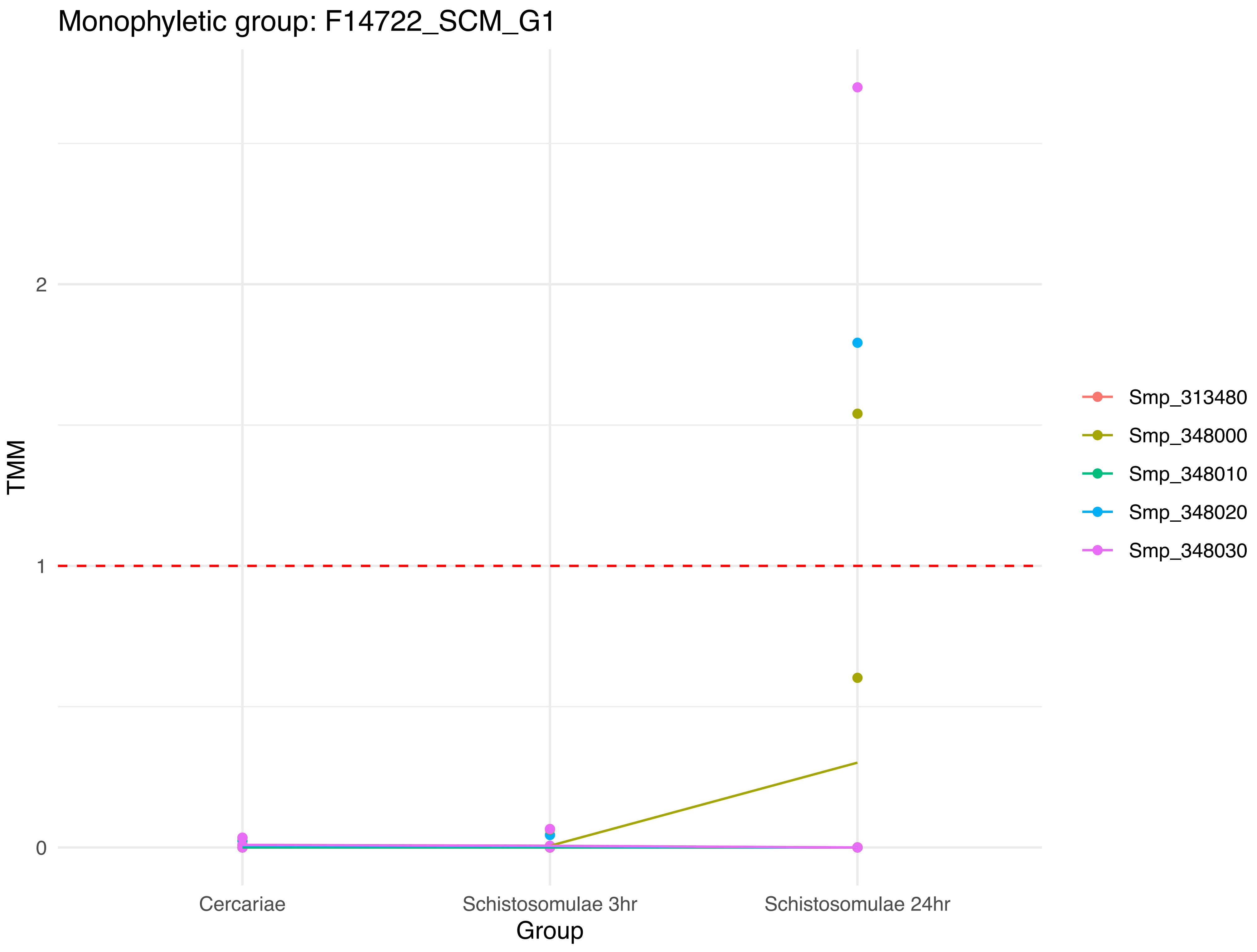
Monophyletic group: F14722_SCM_G1
TMM
0
1
2
Cercariae
Schistosomulae 3hr
Schistosomulae 24hr
Group
Smp_313480
Smp_348000
Smp_348010
Smp_348020
Smp_348030

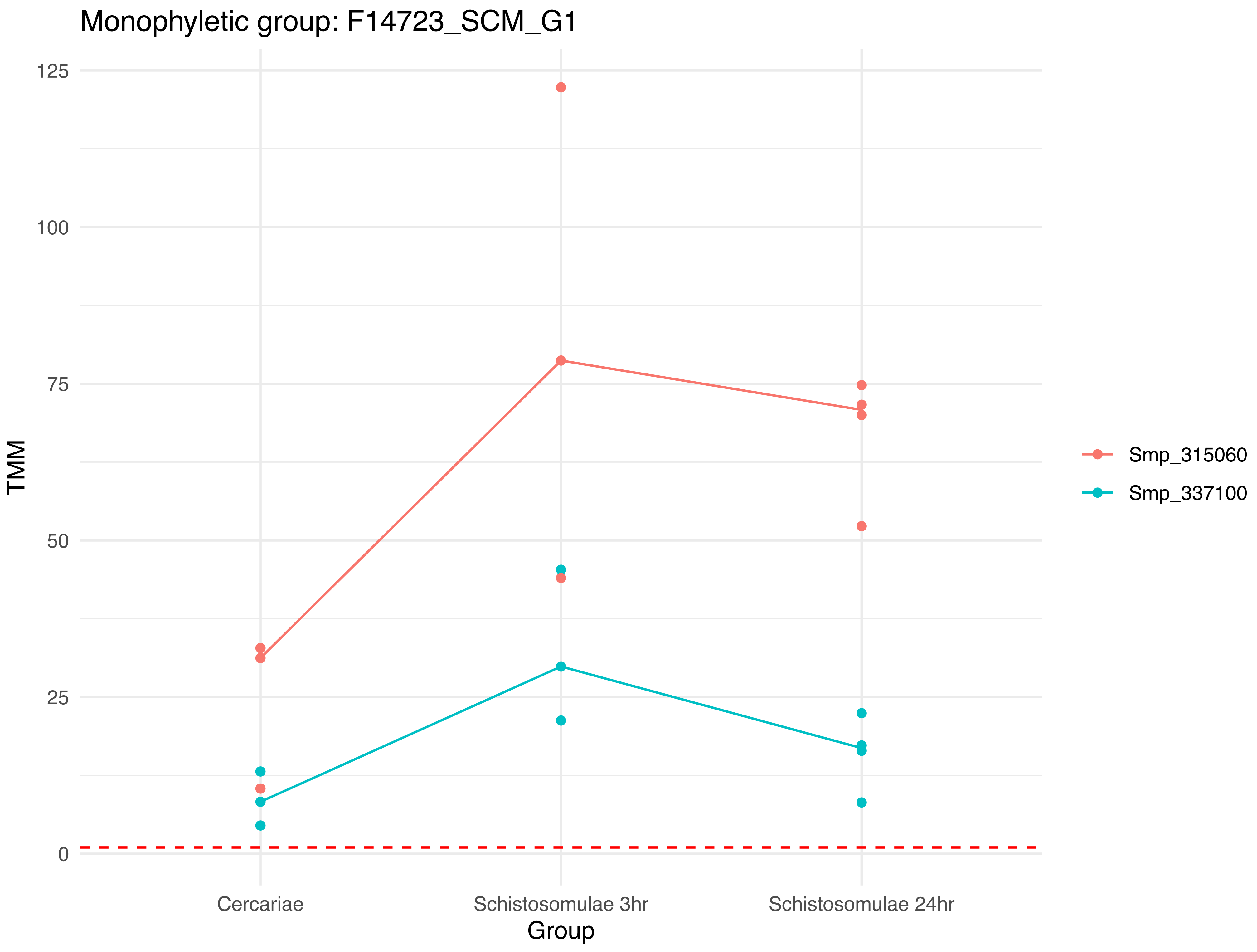

Monophyletic group: F14723_SCM_G1
125
100
75
50
25
0
TMM
Cercariae
Schistosomulae 3hr
Schistosomulae 24hr
Group
Smp_315060
Smp_337100

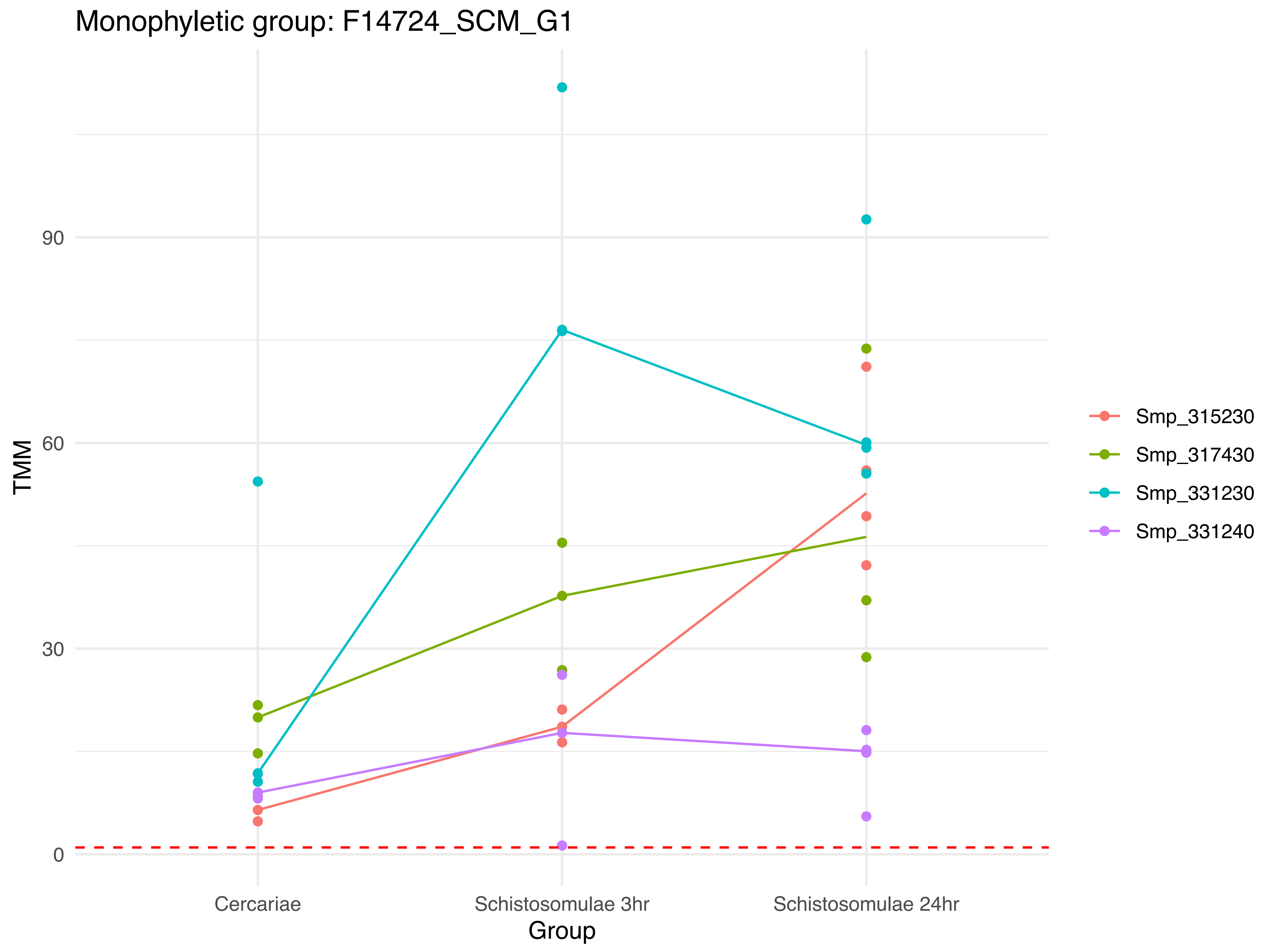

Monophyletic group: F14724_SCM_G1
TMM
90
60
30
0
Cercariae
Schistosomulae 3hr
Schistosomulae 24hr
Group
Smp_315230
Smp_317430
Smp_331230
Smp_331240

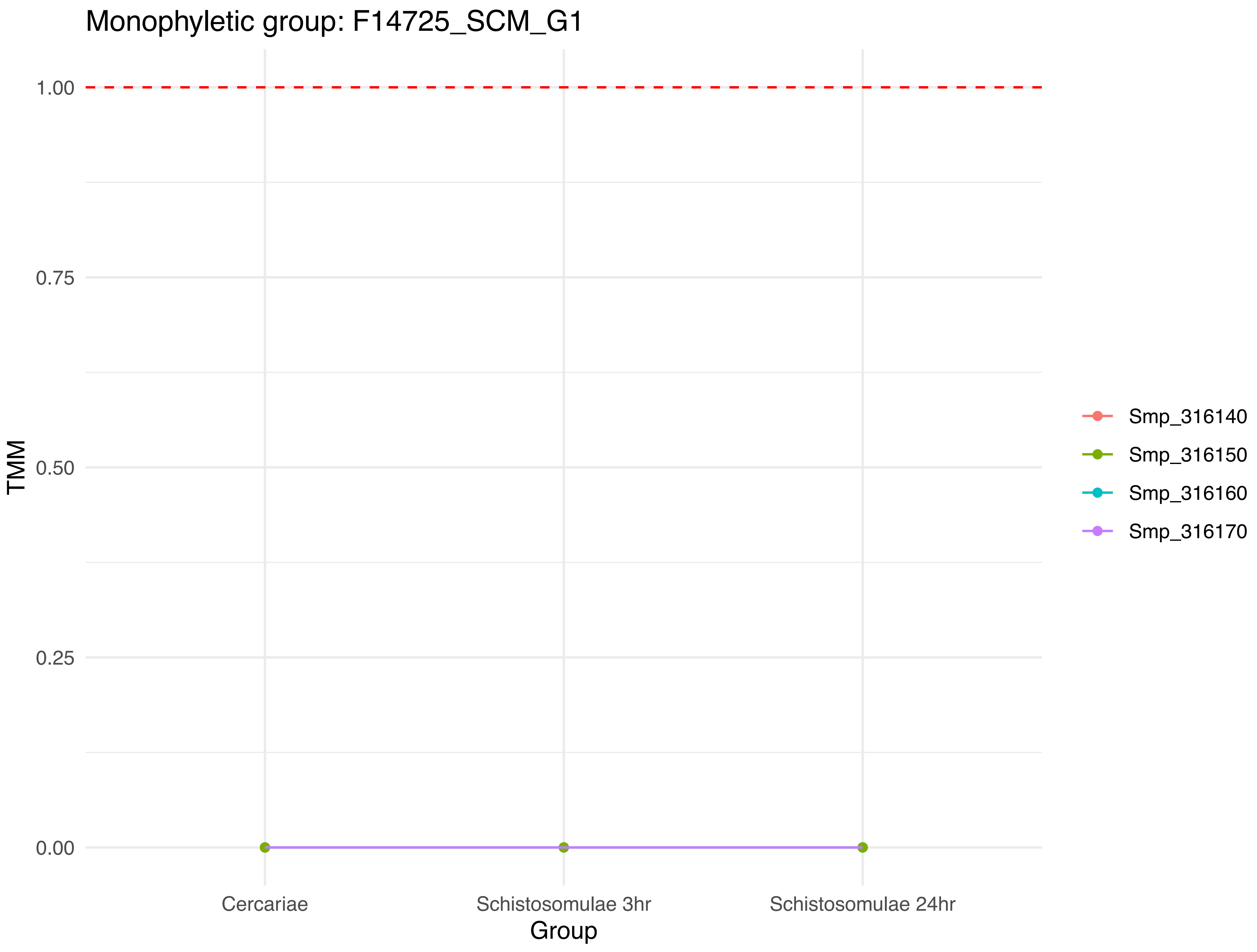

Monophyletic group: F14725_SCM_G1
1.00
0.75
0.50
0.25
0.00
TMM
Cercariae
Schistosomulae 3hr
Schistosomulae 24hr
Group
Smp_316140
Smp_316150
Smp_316160
Smp_316170

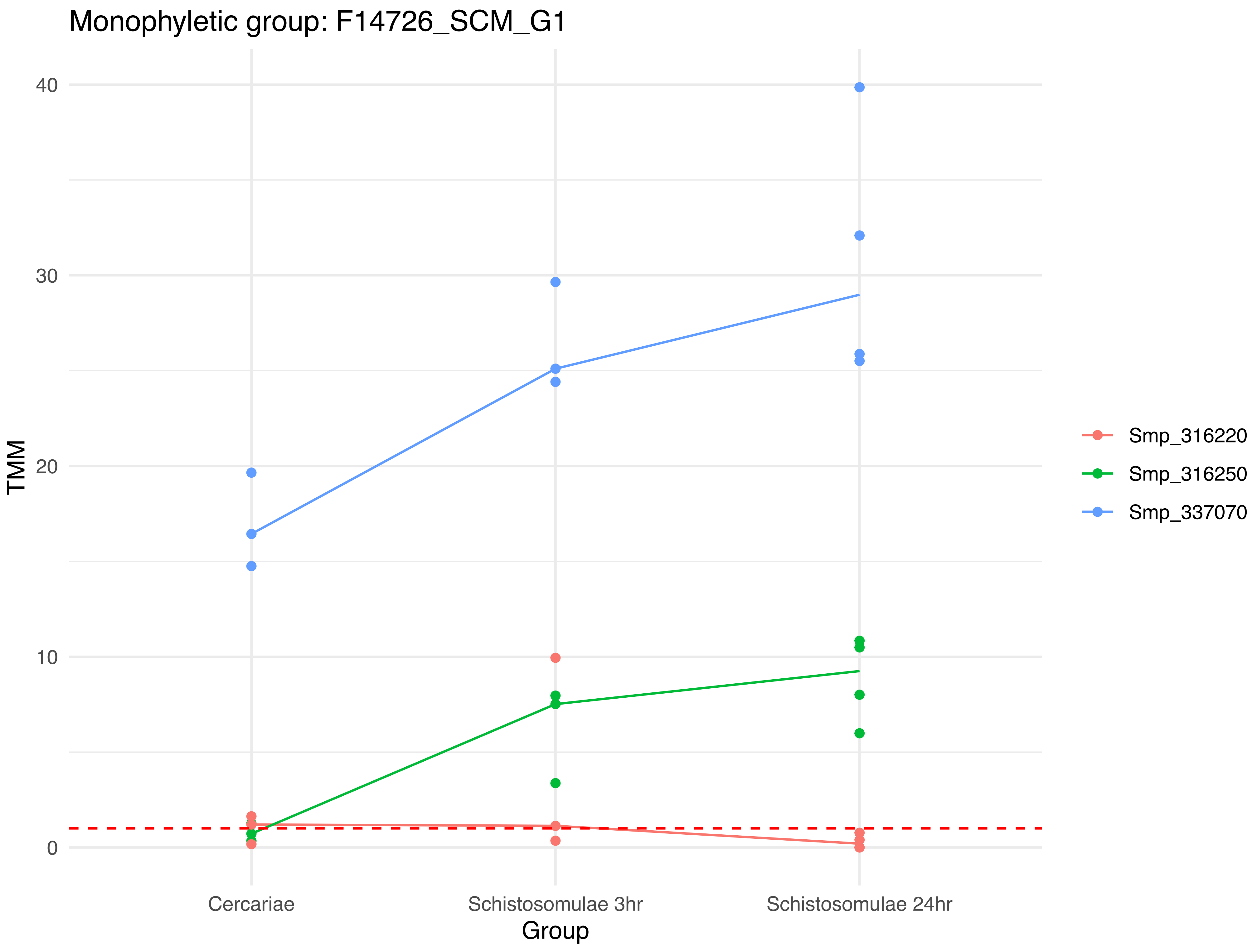
Monophyletic group: F14726_SCM_G1
40
30
20
10
0
TMM
Cercariae
Schistosomulae 3hr
Schistosomulae 24hr
Group
Smp_316220
Smp_316250
Smp_337070

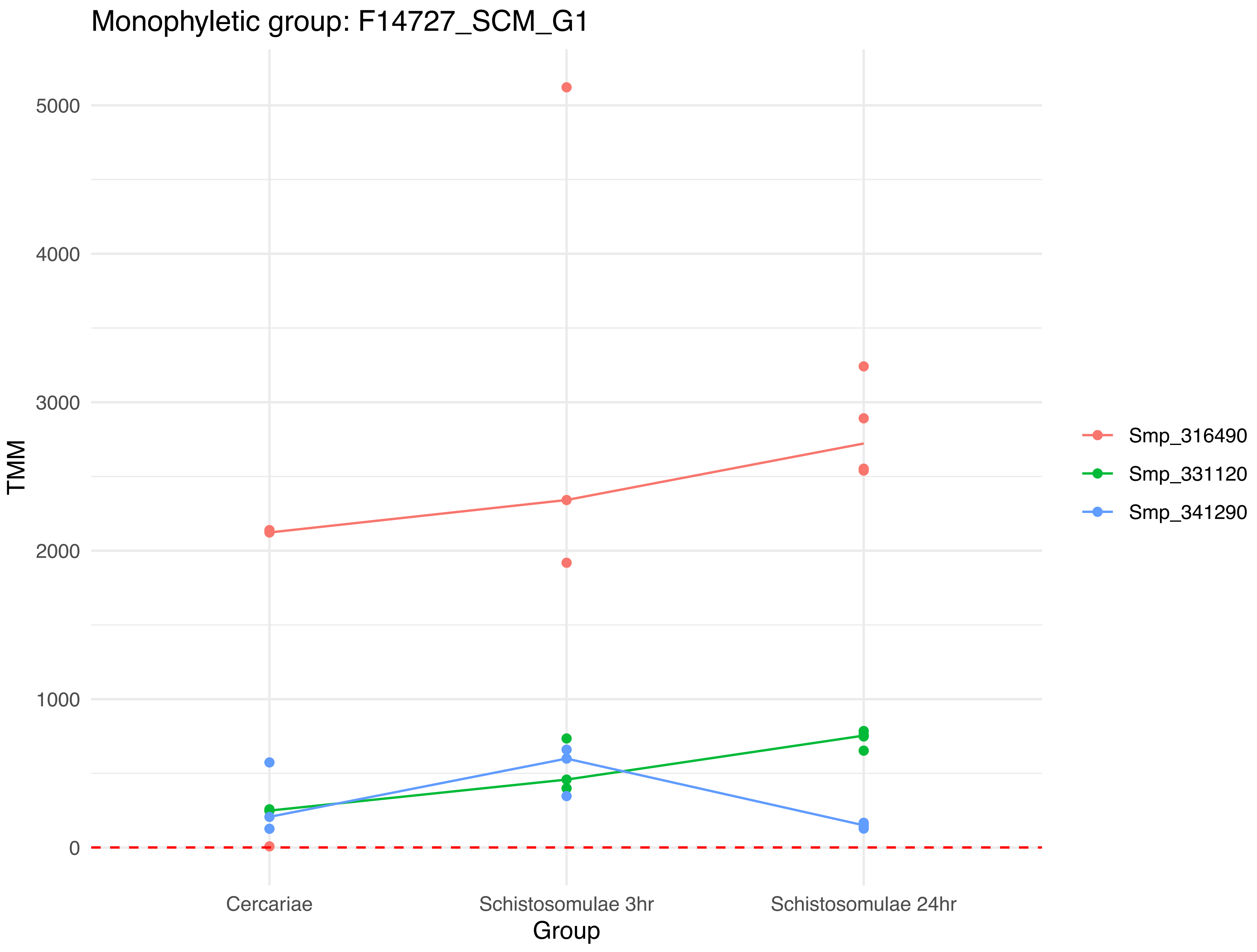
Monophyletic group: F14727_SCM_G1
5000
4000
3000
2000
1000
0
TMM
Cercariae
Schistosomulae 3hr
Schistosomulae 24hr
Group
Smp_316490
Smp_331120
Smp_341290

Monophyletic group: F14728_SCM_G1

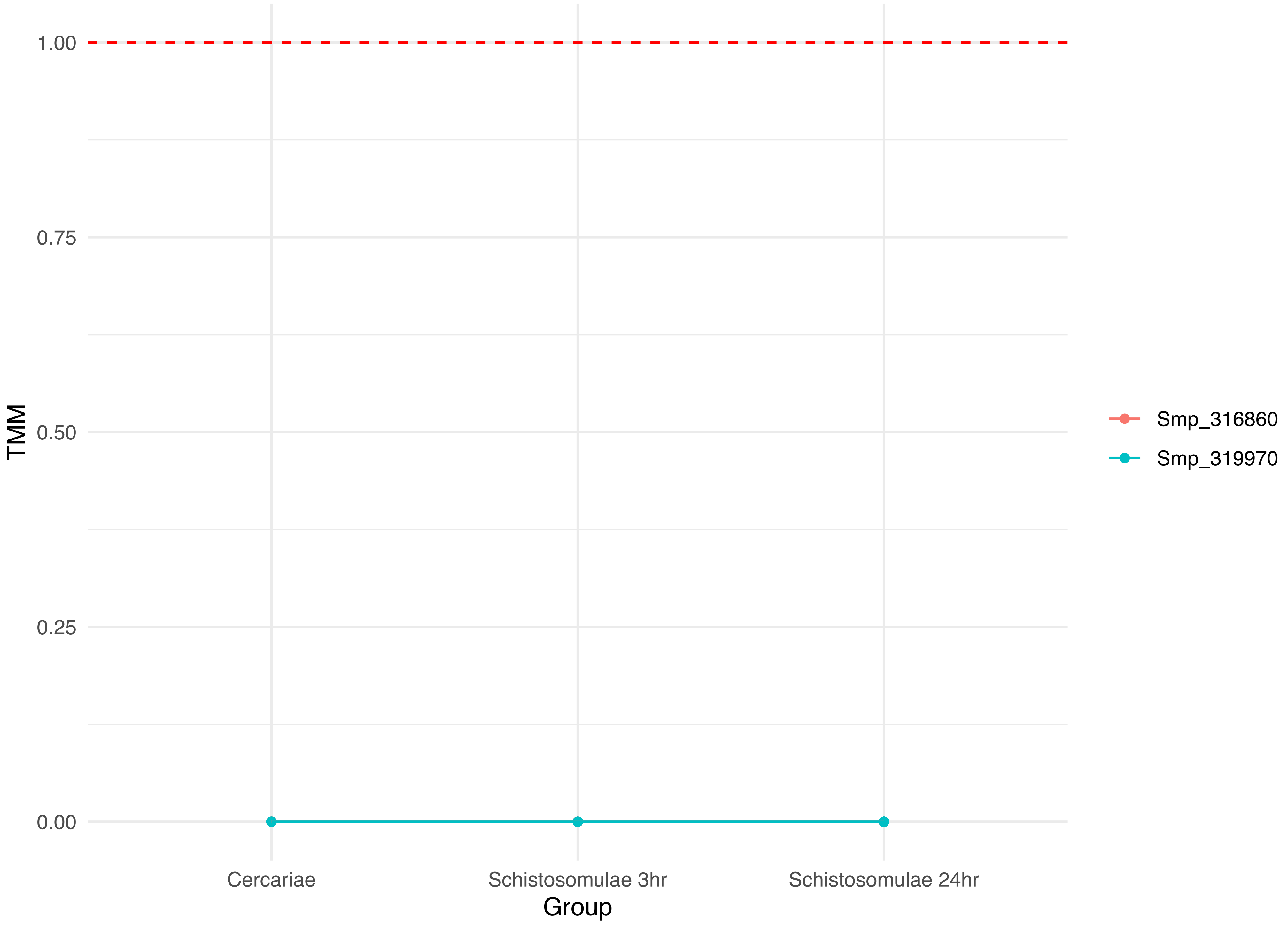

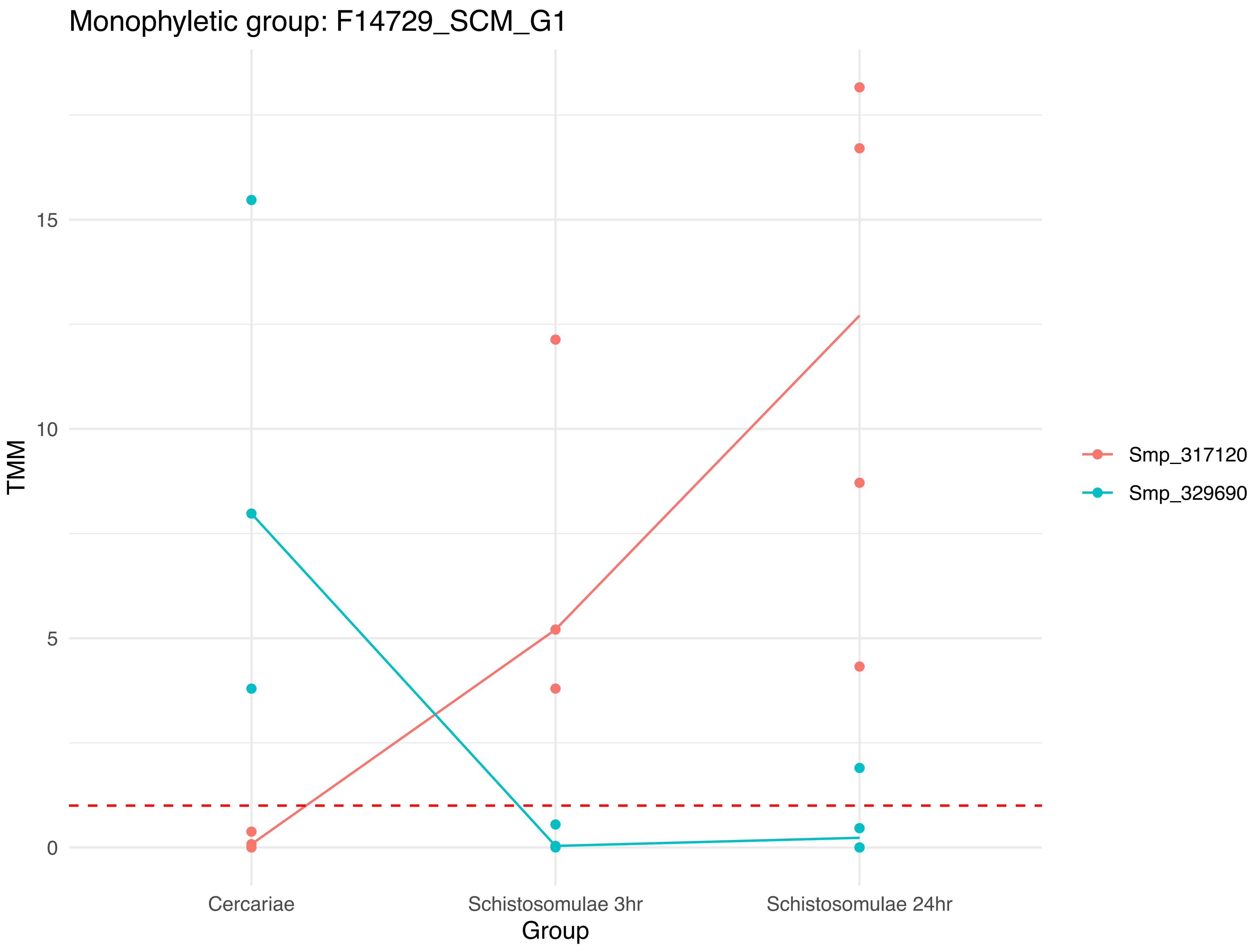
Monophyletic group: F14729_SCM_G1
TMM
15
10
5
0
Cercariae
Schistosomulae 3hr
Schistosomulae 24hr
Group
Smp_317120
Smp_329690

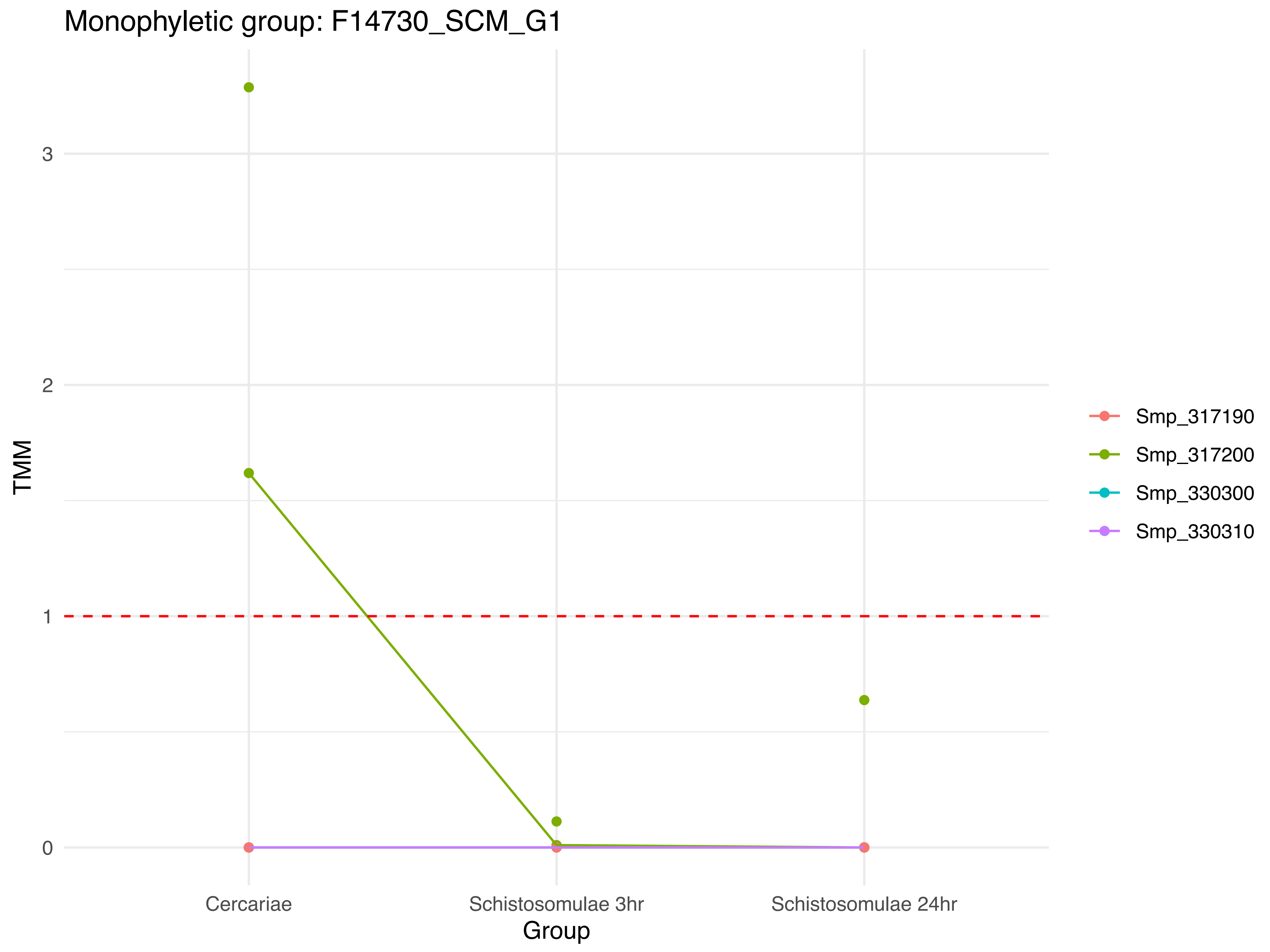
Monophyletic group: F14730_SCM_G1
TMM
3
2
1
0
Cercariae
Schistosomulae 3hr
Schistosomulae 24hr
Group
Smp_317190
Smp_317200
Smp_330300
Smp_330310

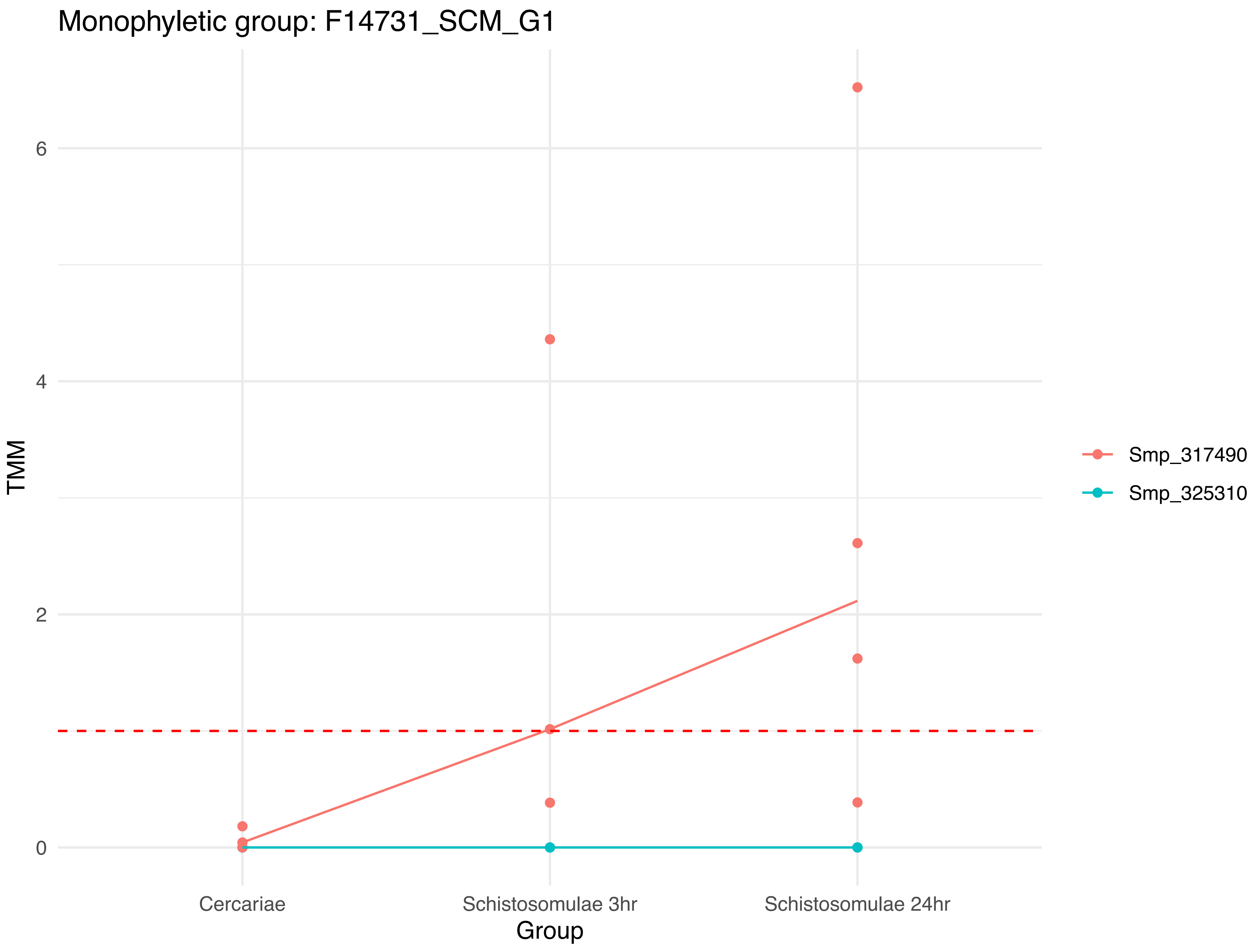
Monophyletic group: F14731_SCM_G1
TMM
0
2
4
6
Cercariae
Schistosomulae 3hr
Schistosomulae 24hr
Group
Smp_317490
Smp_325310

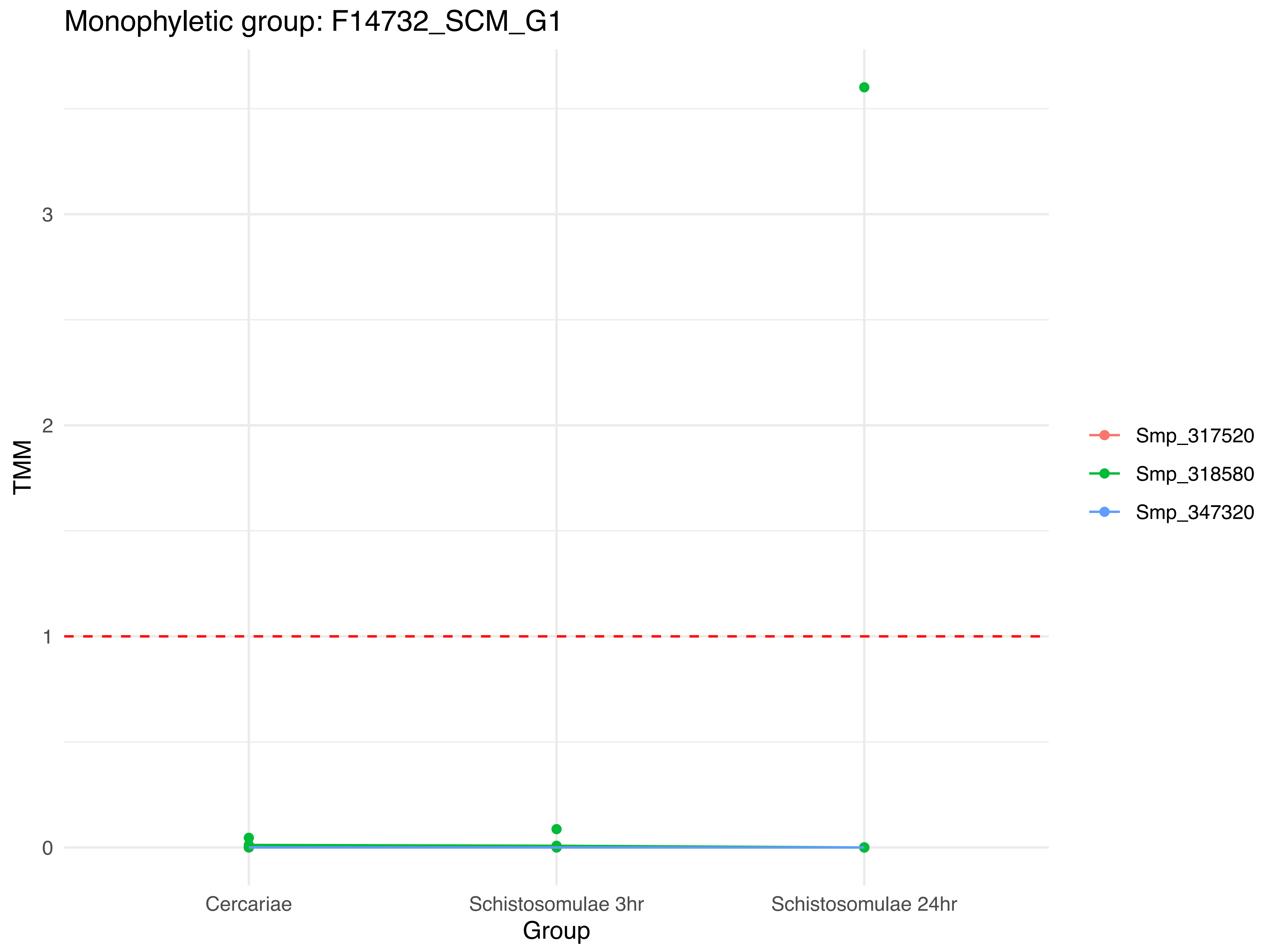
Monophyletic group: F14732_SCM_G1
TMM
3
2
1
0
Cercariae
Schistosomulae 3hr
Schistosomulae 24hr
Group
Smp_317520
Smp_318580
Smp_347320

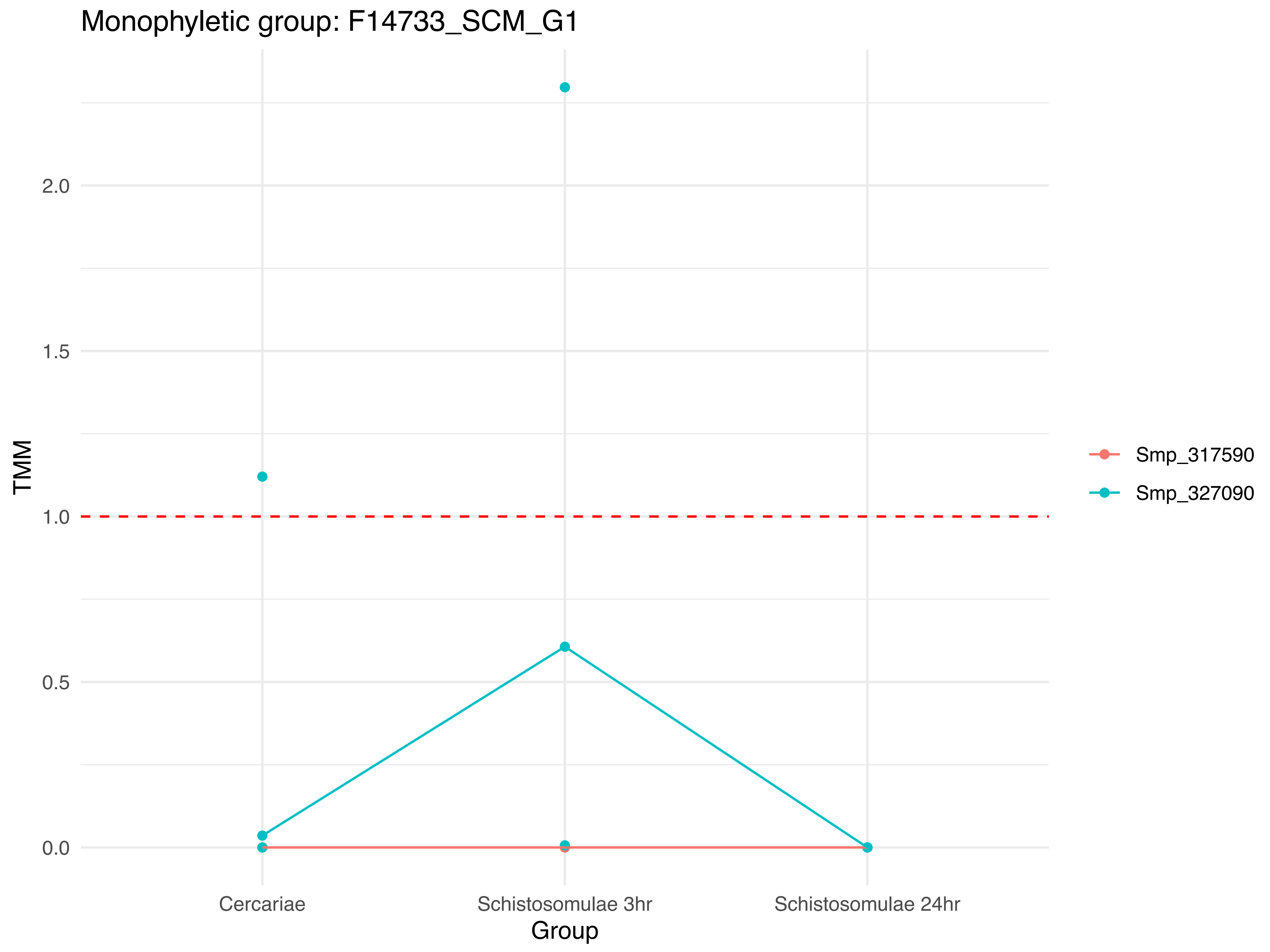

Monophyletic group: F14733_SCM_G1
TMM
2.0
1.5
1.0
0.5
0.0
Cercariae
Schistosomulae 3hr
Schistosomulae 24hr
Group
Smp_317590
Smp_327090

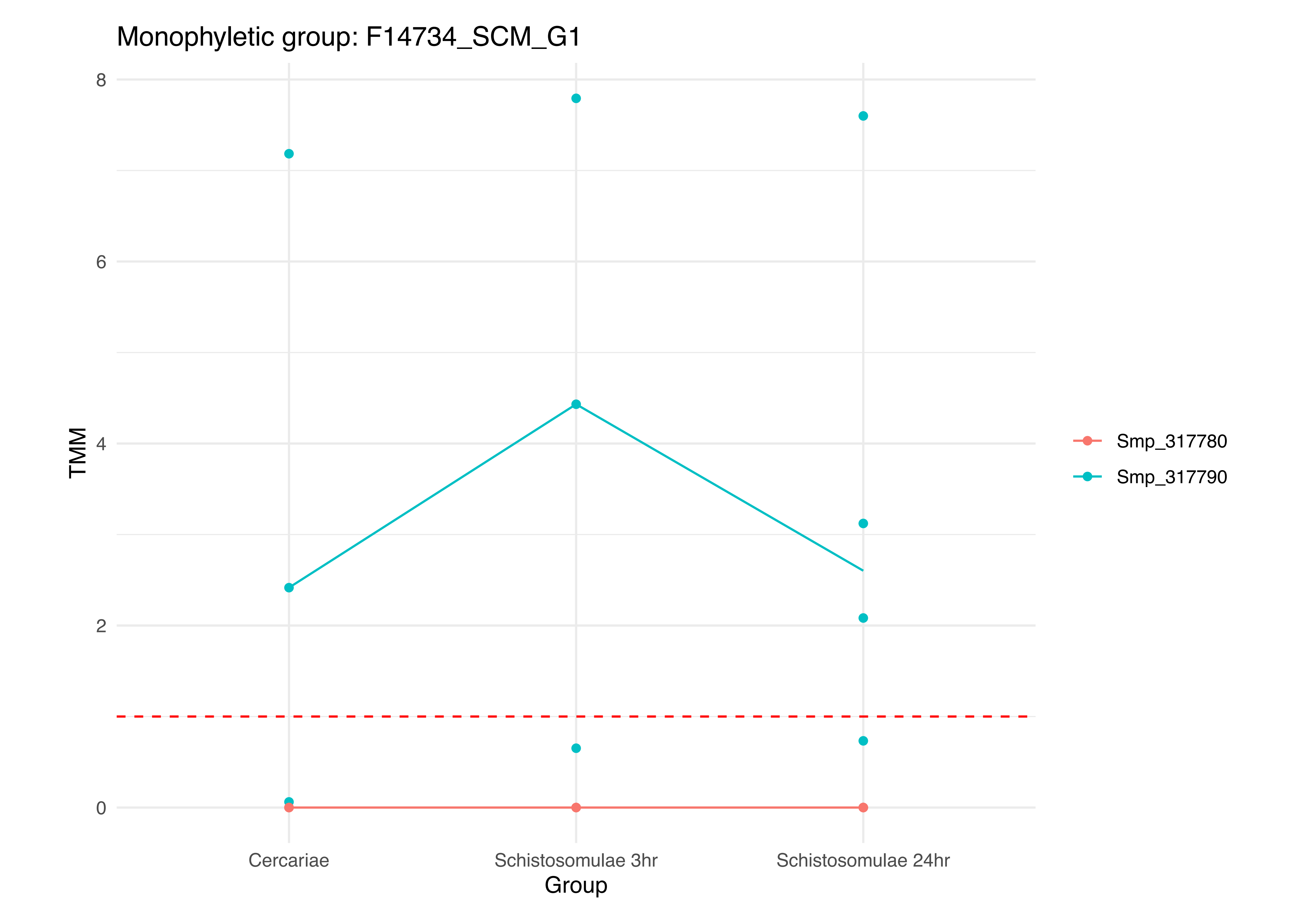

Monophyletic group: F14734_SCM_G1
TMM
8
6
4
2
0
Cercariae
Schistosomulae 3hr
Schistosomulae 24hr
Group
Smp_317780
Smp_317790

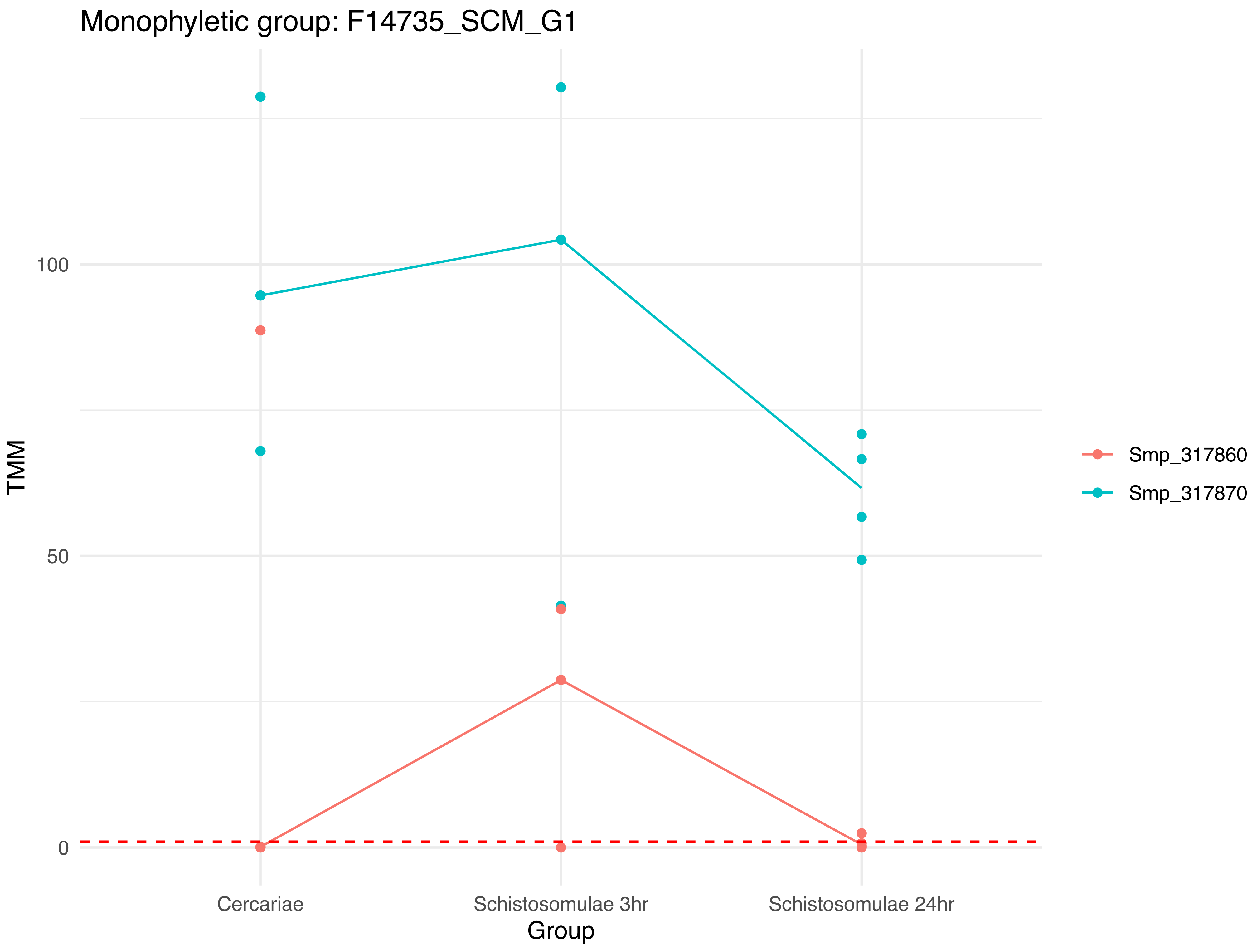

Monophyletic group: F14735_SCM_G1
TMM
100
50
0
Cercariae
Schistosomulae 3hr
Schistosomulae 24hr
Group
Smp_317860
Smp_317870

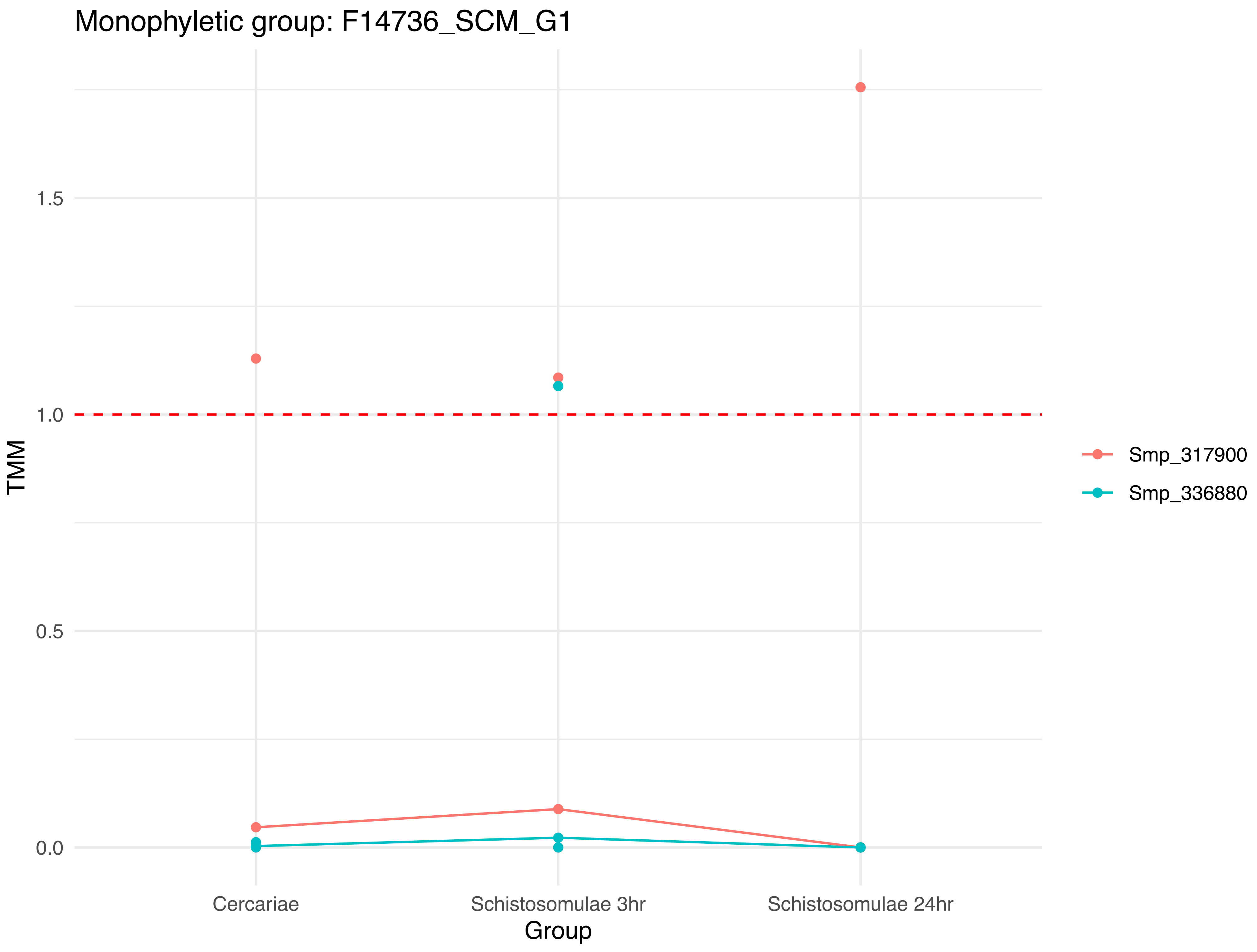
Monophyletic group: F14736_SCM_G1
1.5
1.0
0.5
0.0
TMM
Cercariae
Schistosomulae 3hr
Schistosomulae 24hr
Group
Smp_317900
Smp_336880

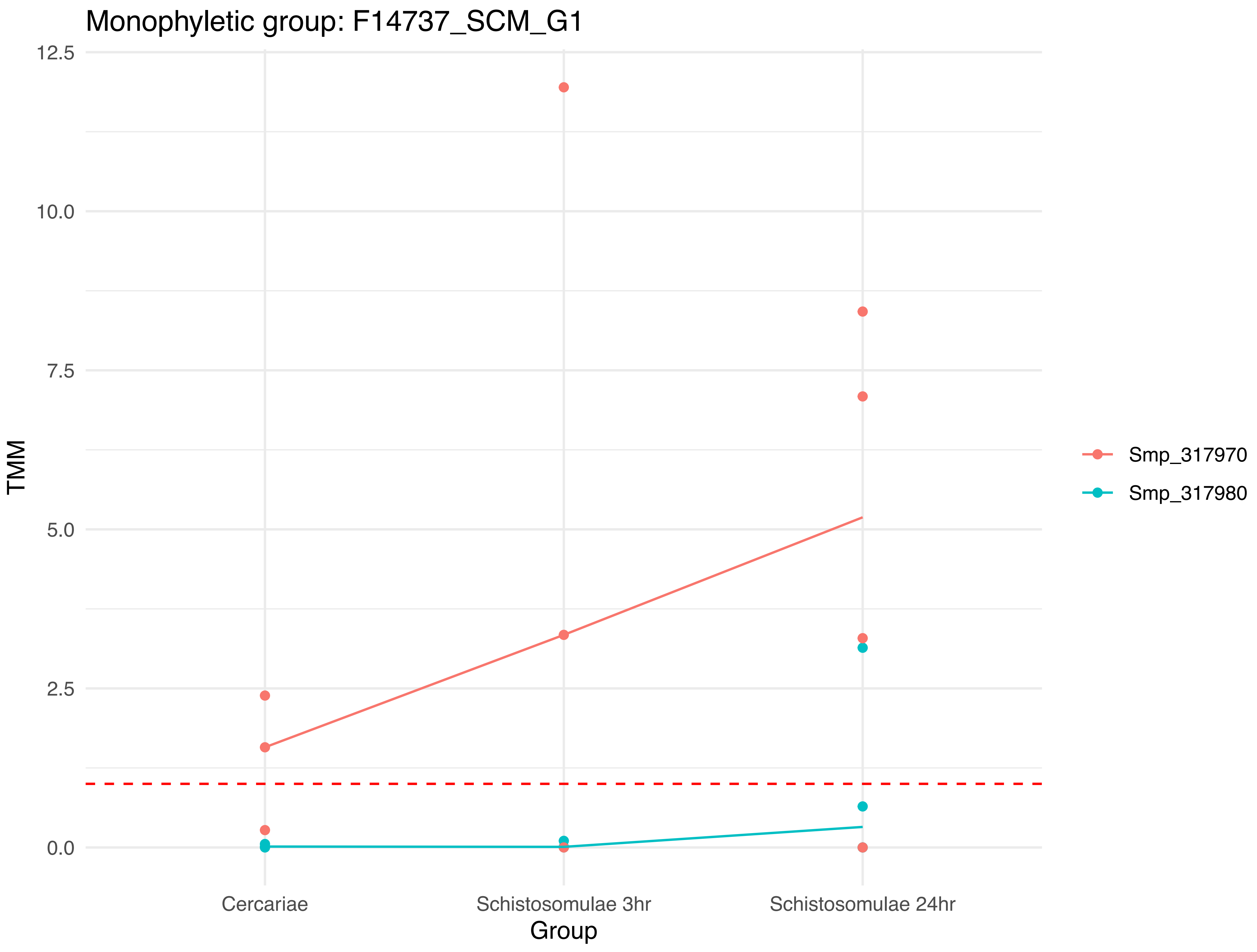

Monophyletic group: F14737_SCM_G1
12.5
10.0
7.5
5.0
2.5
0.0
TMM
Cercariae
Schistosomulae 3hr
Schistosomulae 24hr
Group
Smp_317970
Smp_317980

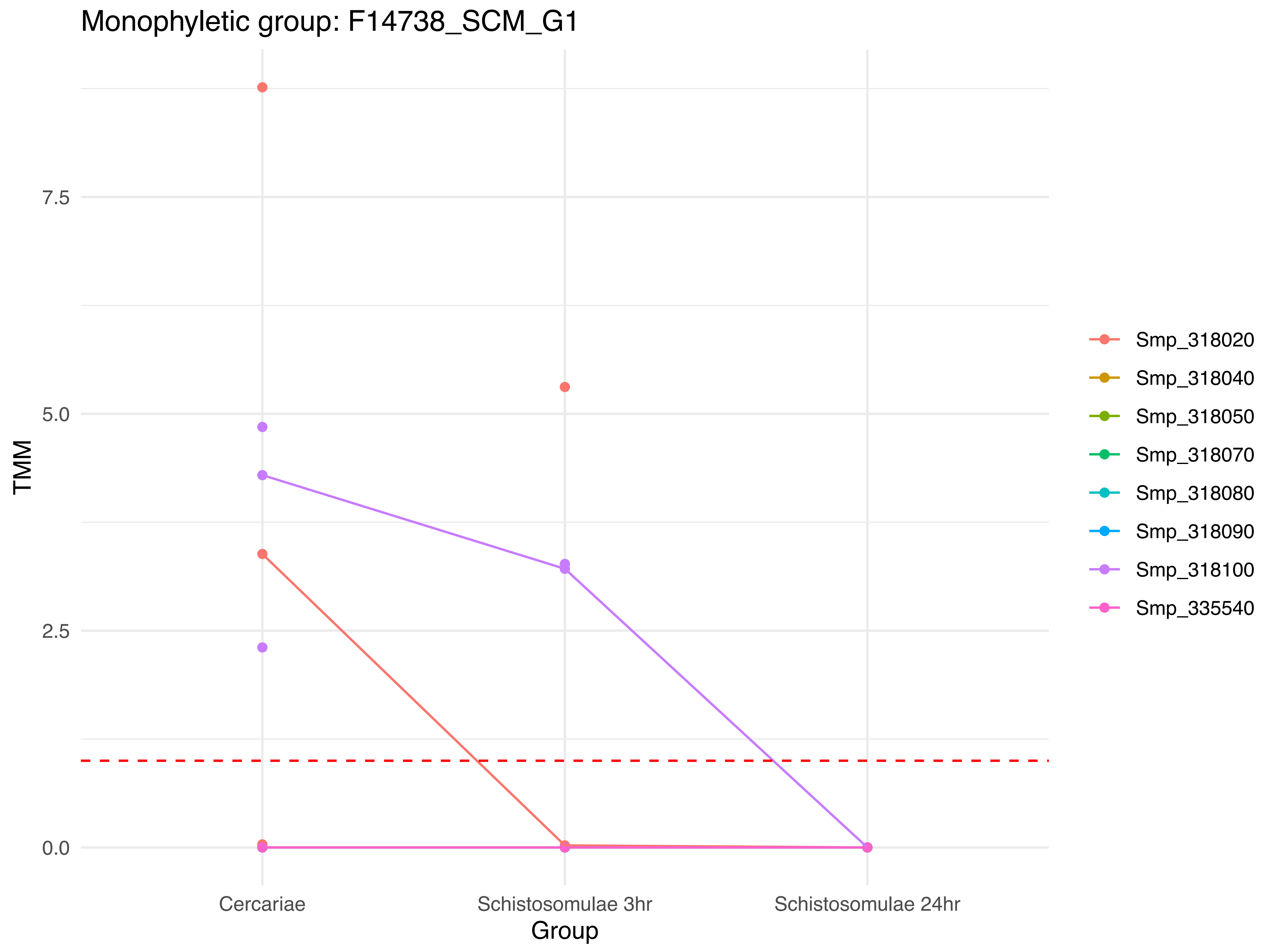
Monophyletic group: F14738_SCM_G1
TMM
7.5
5.0
2.5
0.0
Cercariae
Schistosomulae 3hr
Schistosomulae 24hr
Group
Smp_318020
Smp_318040
Smp_318050
Smp_318070
Smp_318080
Smp_318090
Smp_318100
Smp_335540

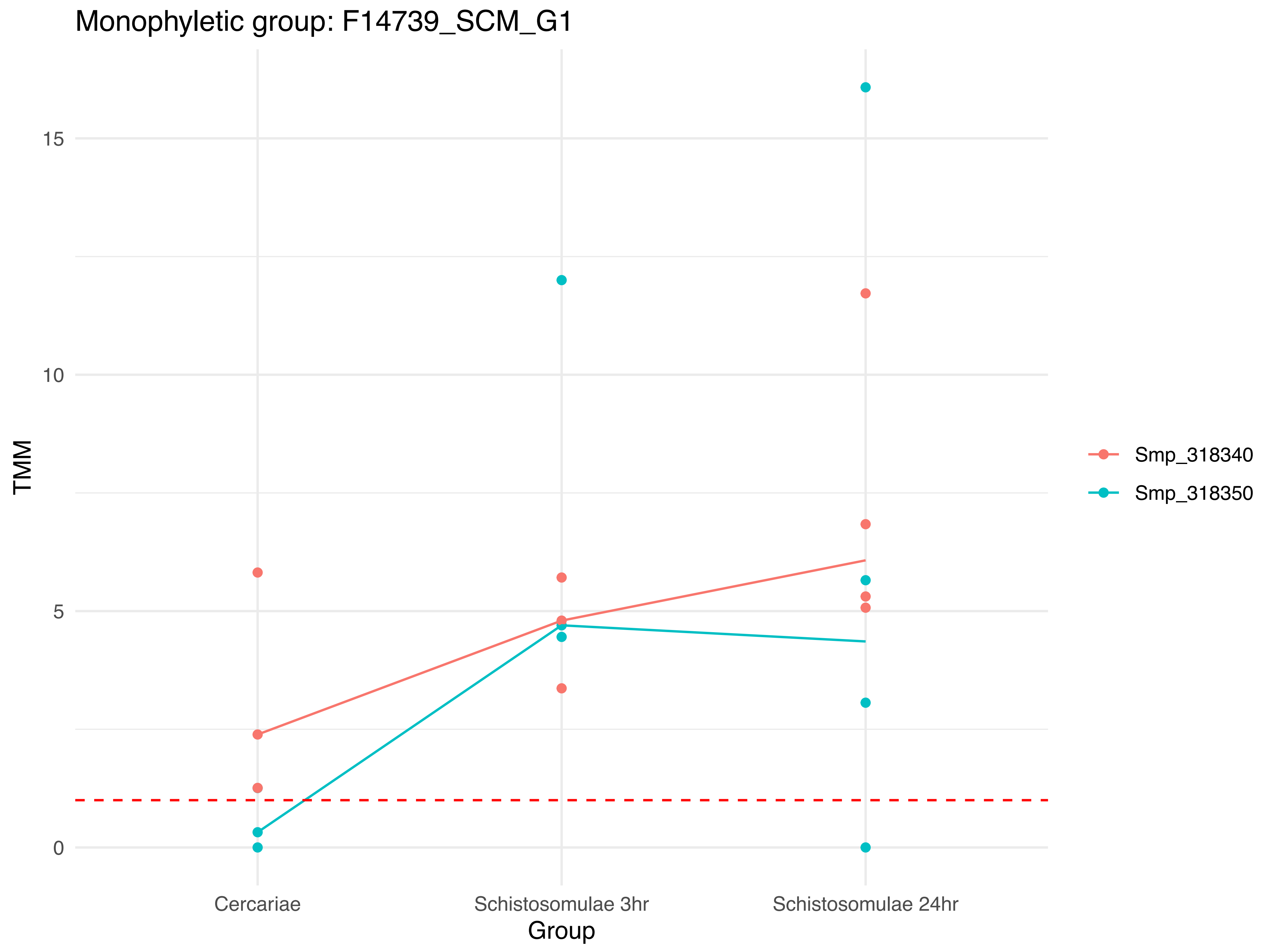
Monophyletic group: F14739_SCM_G1
15
10
5
0
TMM
Cercariae
Schistosomulae 3hr
Schistosomulae 24hr
Group
Smp_318340
Smp_318350

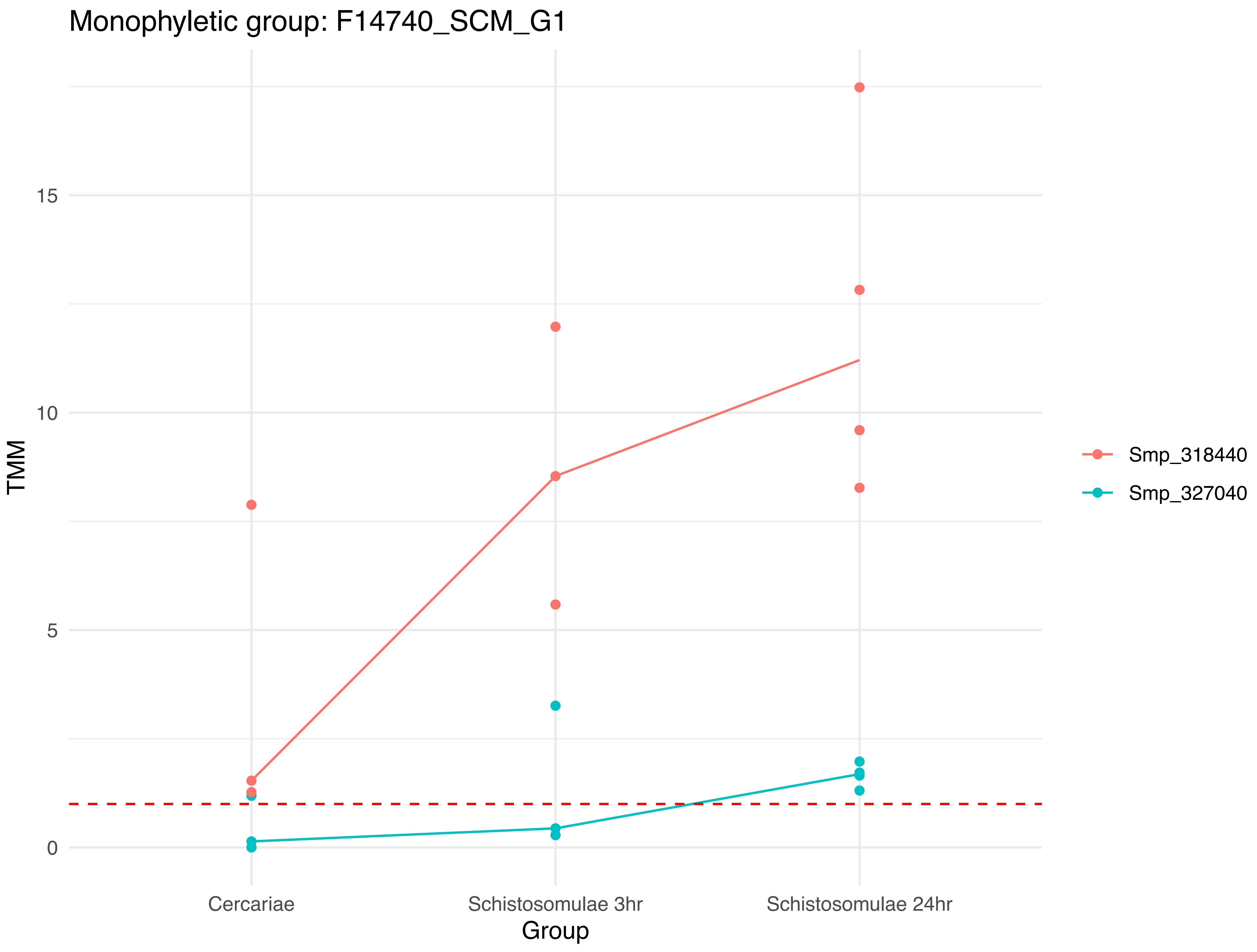

Monophyletic group: F14740_SCM_G1
TMM
15
10
5
0
Cercariae
Schistosomulae 3hr
Schistosomulae 24hr
Group
Smp_318440
Smp_327040

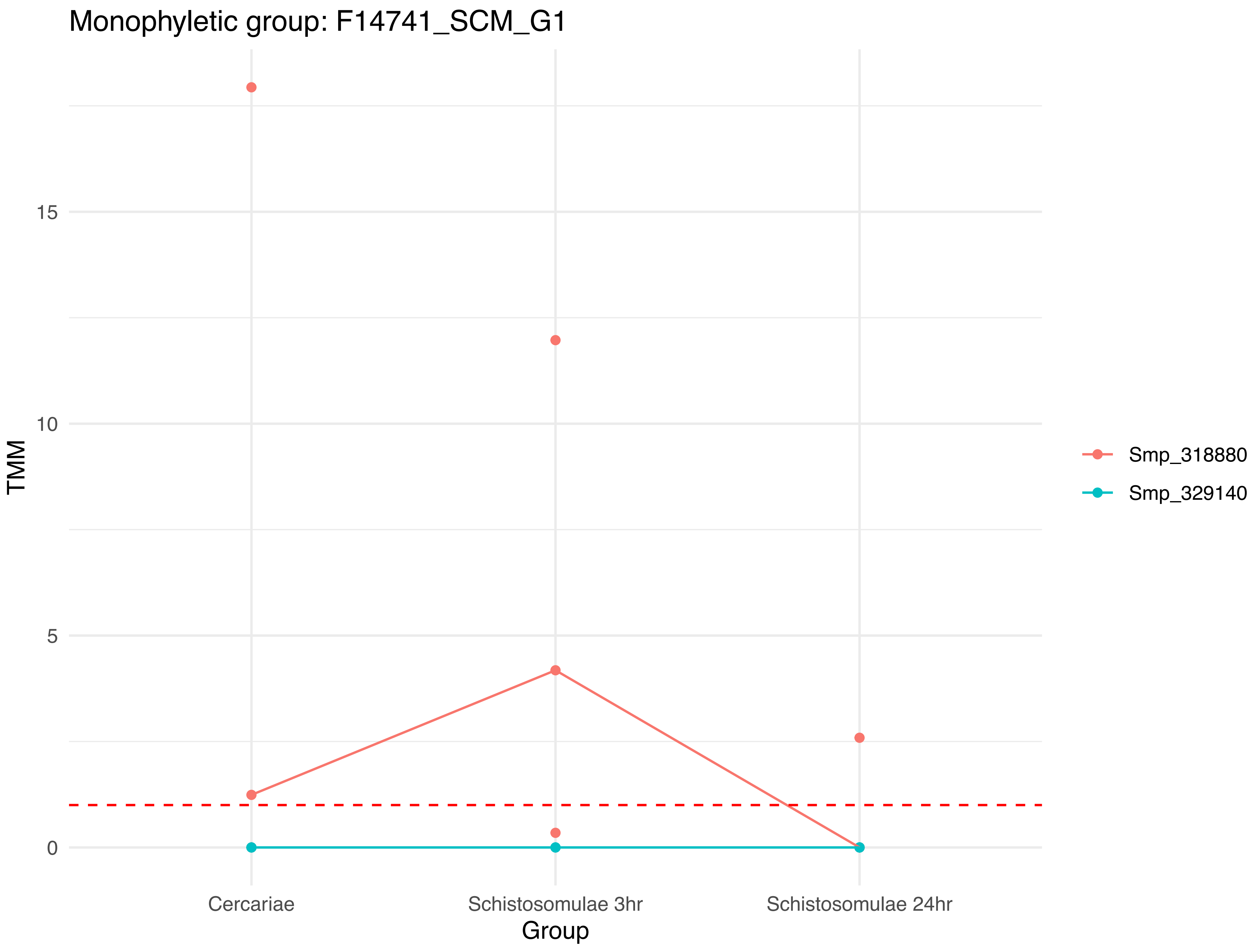
Monophyletic group: F14741_SCM_G1
TMM
15
10
5
0
Cercariae
Schistosomulae 3hr
Schistosomulae 24hr
Group
Smp_318880
Smp_329140

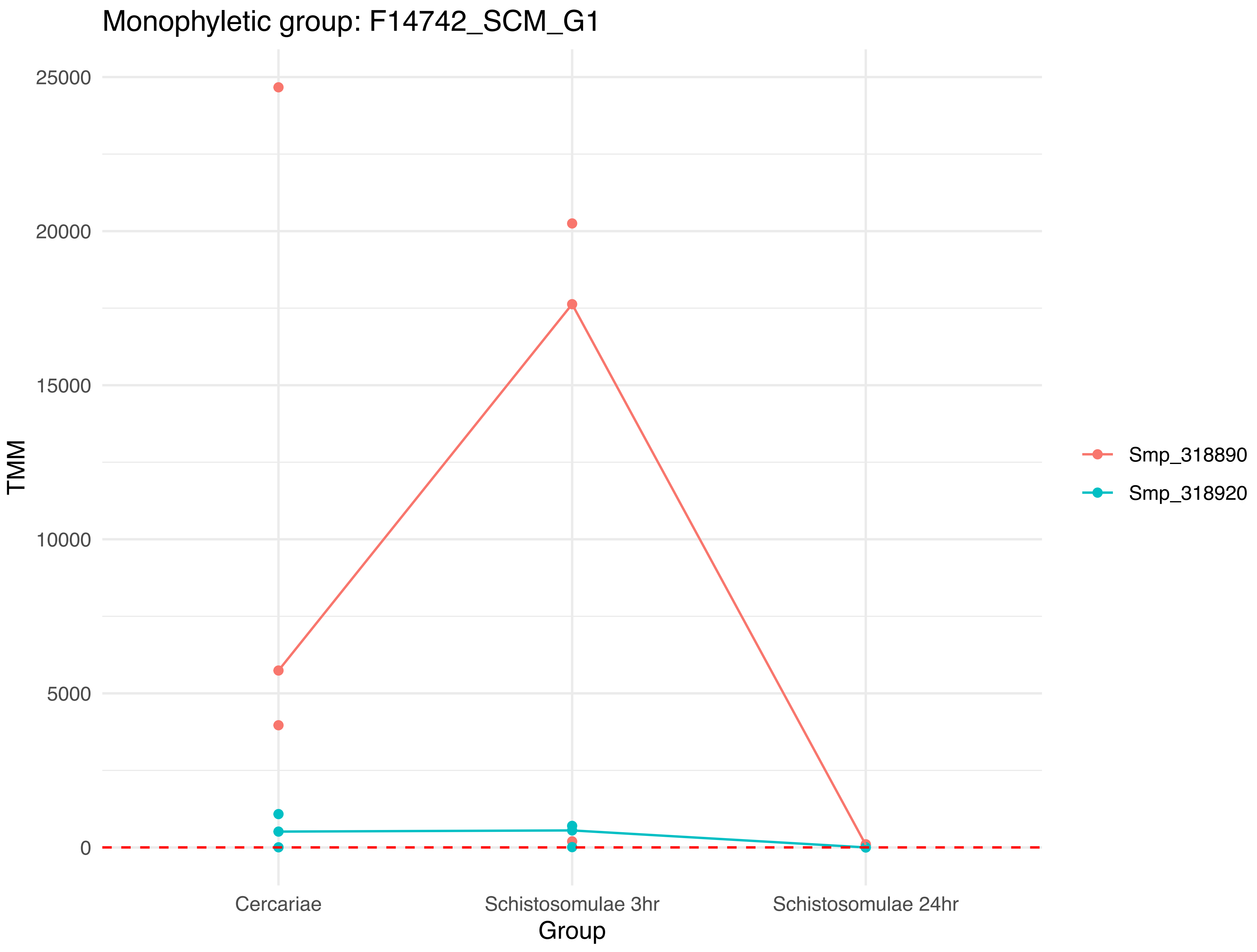

Monophyletic group: F14742_SCM_G1
25000
20000
15000
10000
5000
0
TMM
Cercariae
Schistosomulae 3hr
Schistosomulae 24hr
Group
Smp_318890
Smp_318920

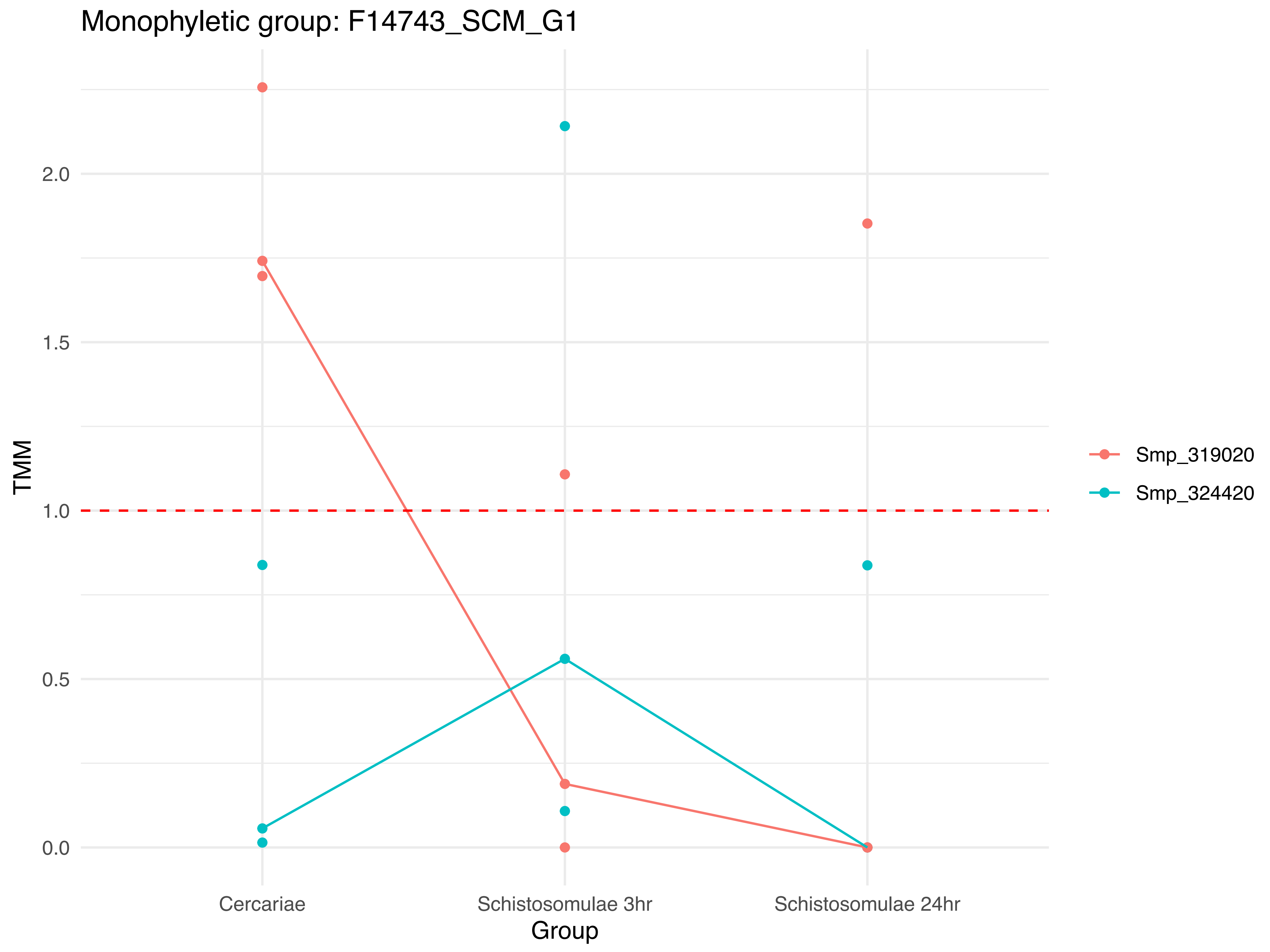
Monophyletic group: F14743_SCM_G1
TMM
2.0
1.5
1.0
0.5
0.0
Cercariae
Schistosomulae 3hr
Schistosomulae 24hr
Group
Smp_319020
Smp_324420

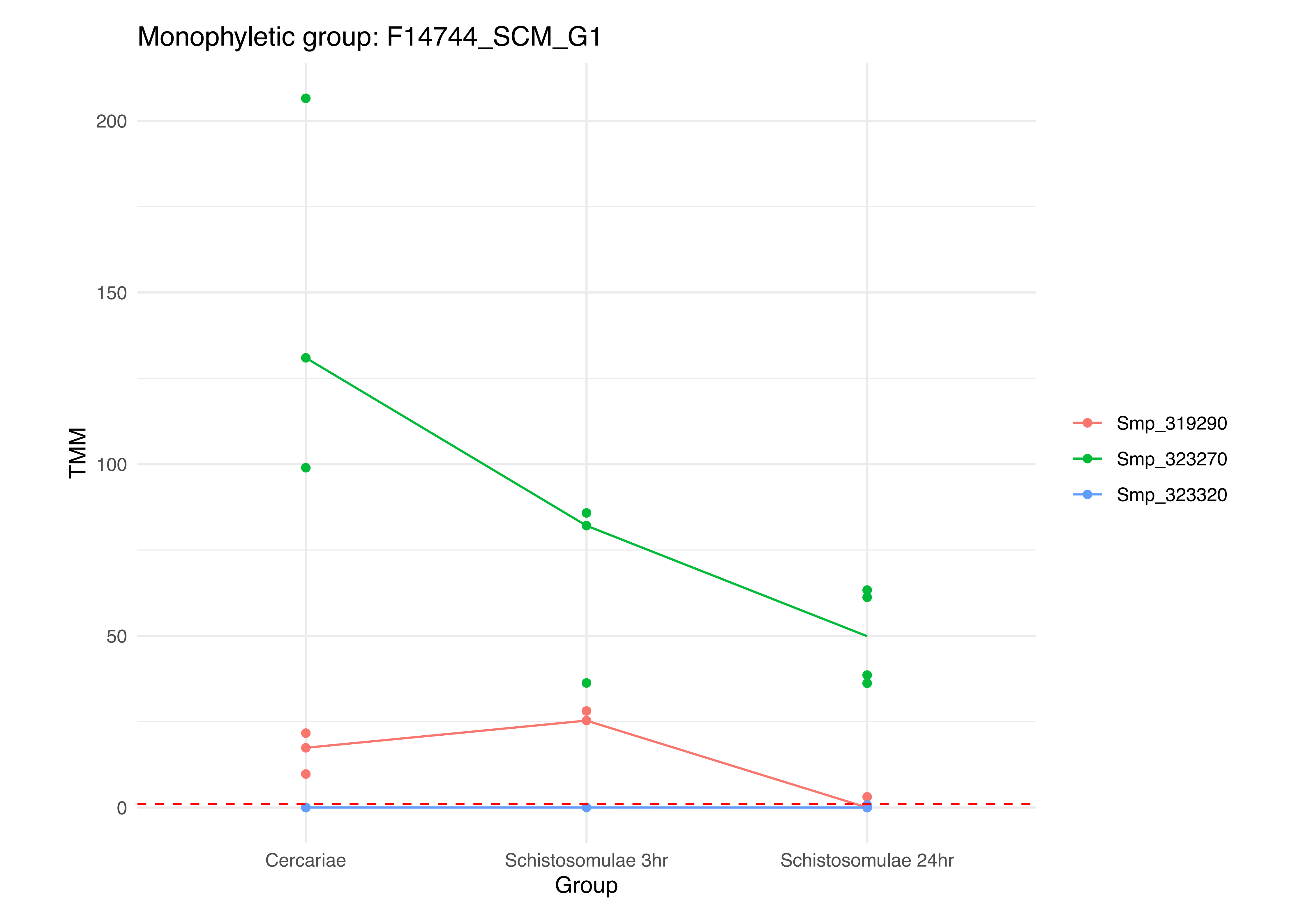

Monophyletic group: F14744_SCM_G1
TMM
200
150
100
50
0
Cercariae
Schistosomulae 3hr
Schistosomulae 24hr
Group
Smp_319290
Smp_323270
Smp_323320

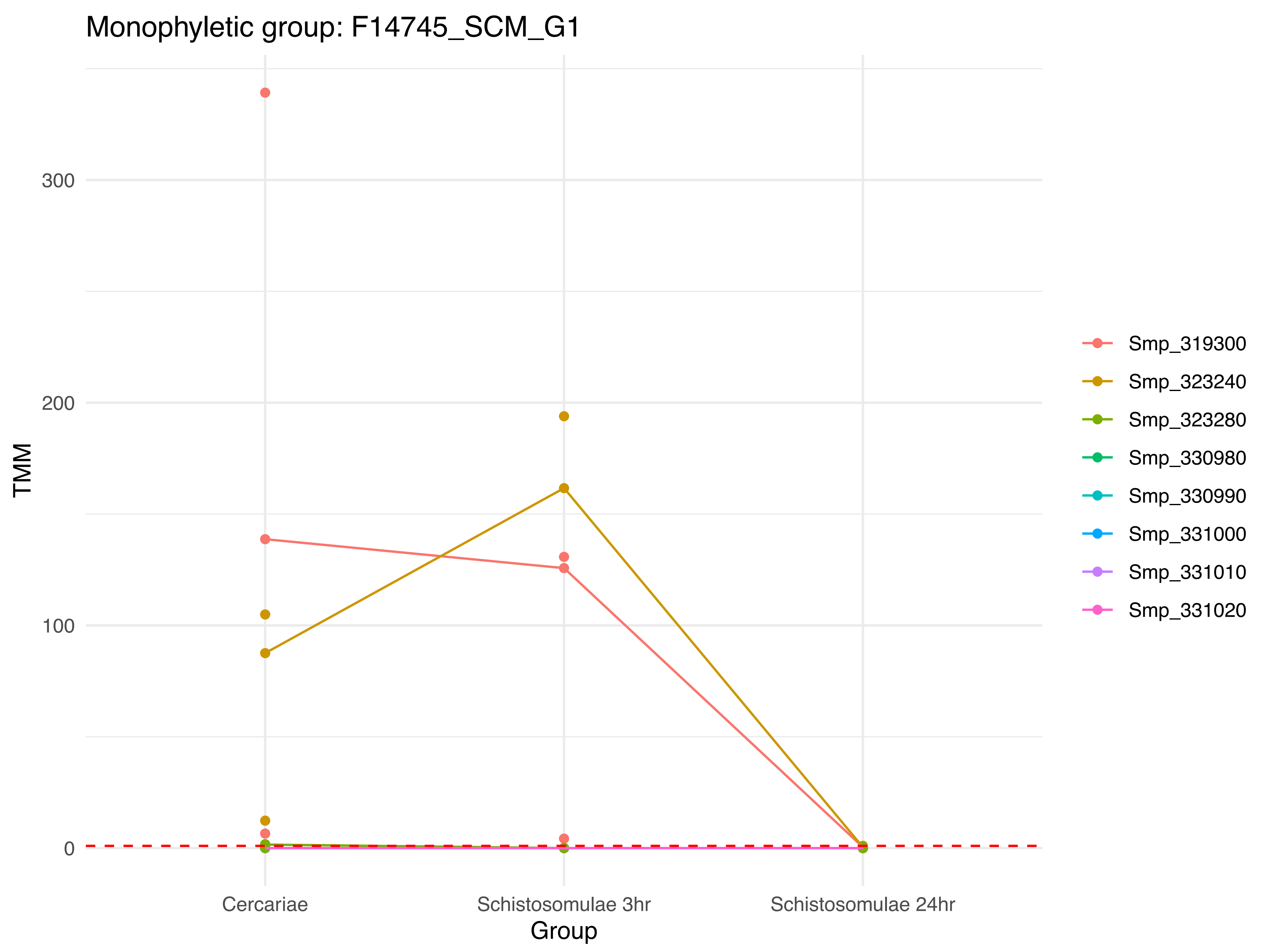

Monophyletic group: F14745_SCM_G1
TMM
300
200
100
0
Cercariae
Schistosomulae 3hr
Schistosomulae 24hr
Group
Smp_319300
Smp_323240
Smp_323280
Smp_330980
Smp_330990
Smp_331000
Smp_331010
Smp_331020

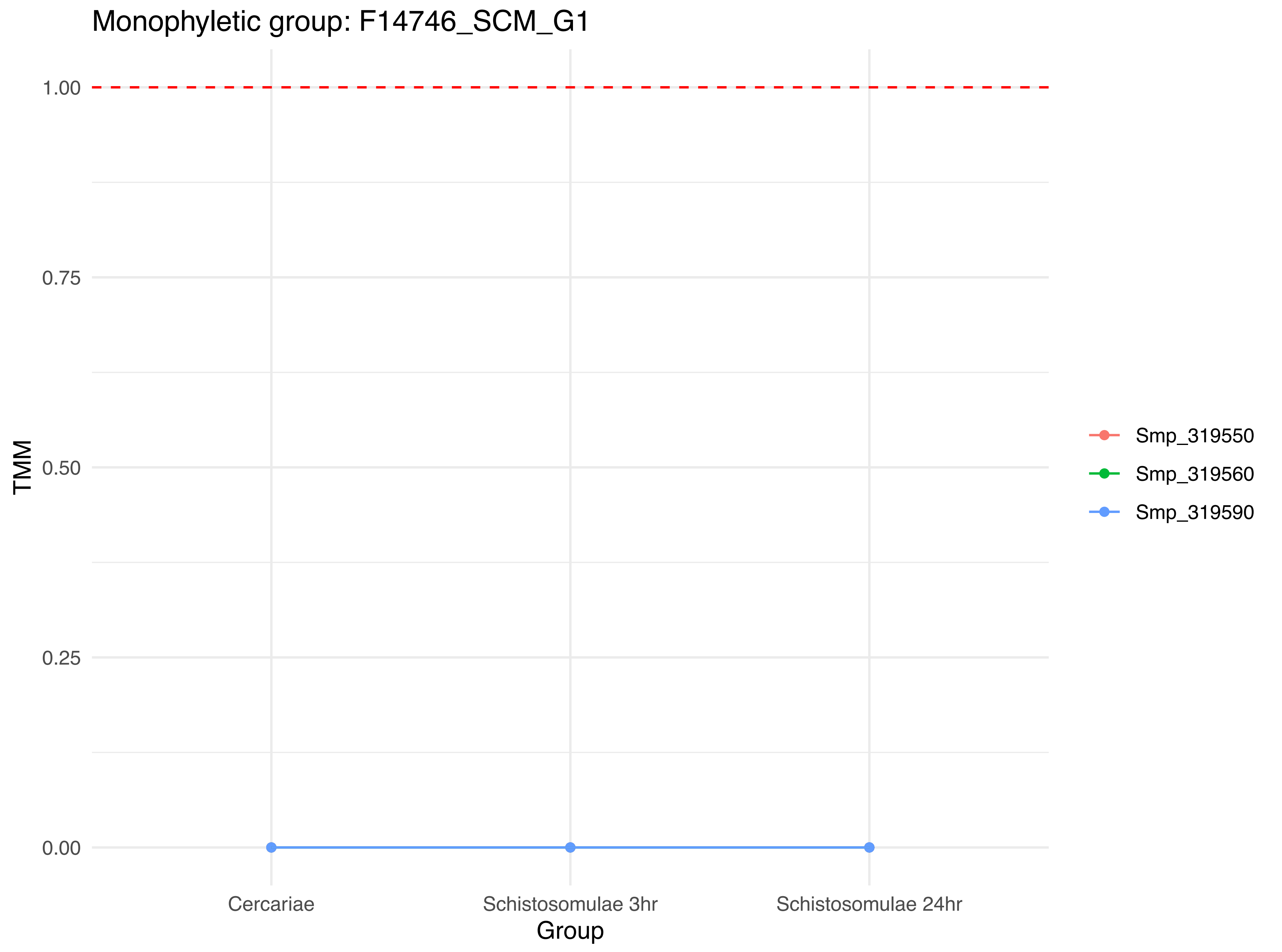

Monophyletic group: F14746_SCM_G1
TMM
1.00
0.75
0.50
0.25
0.00
Cercariae
Schistosomulae 3hr
Schistosomulae 24hr
Group
Smp_319550
Smp_319560
Smp_319590

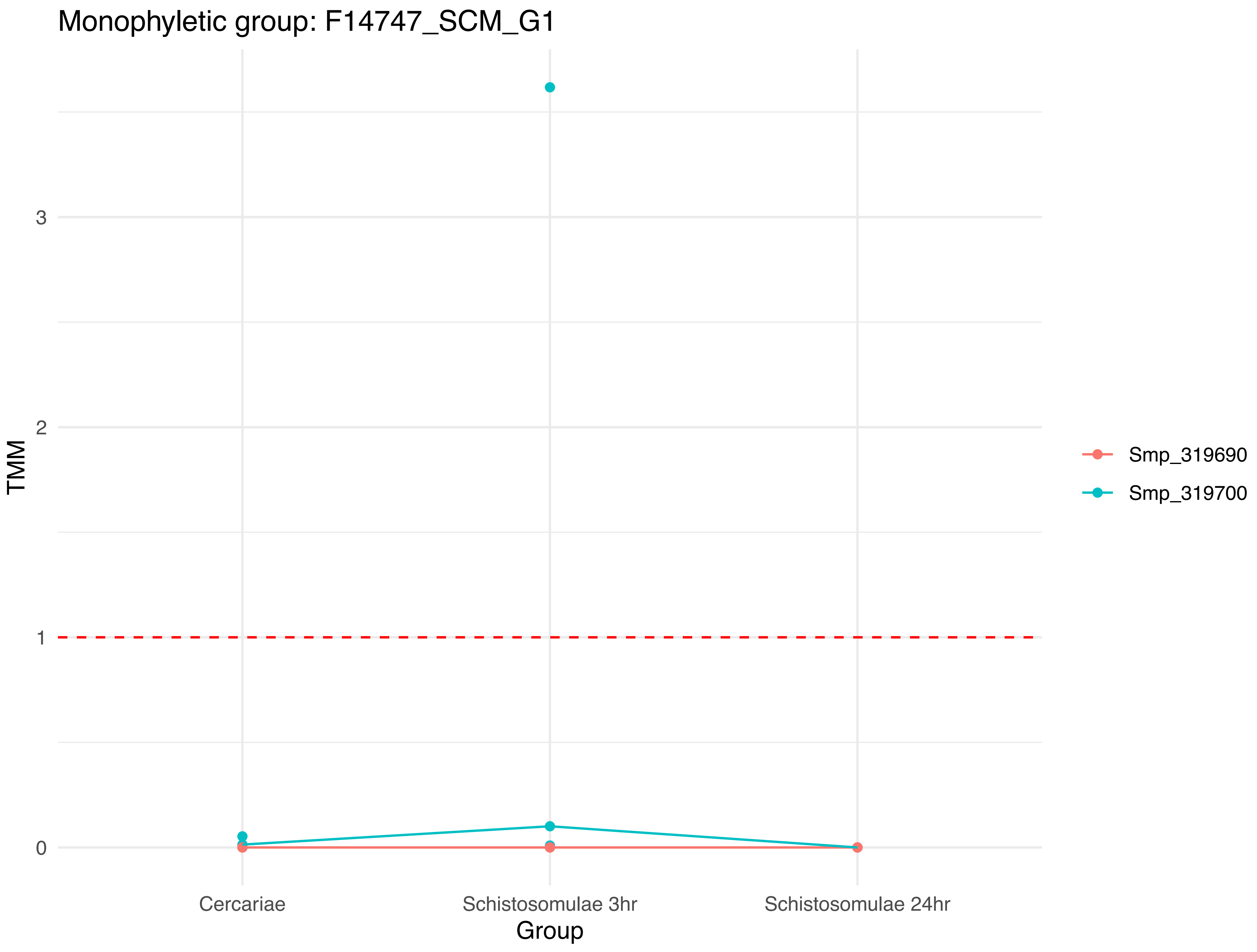

Monophyletic group: F14747_SCM_G1
TMM
3
2
1
0
Cercariae
Schistosomulae 3hr
Schistosomulae 24hr
Group
Smp_319690
Smp_319700

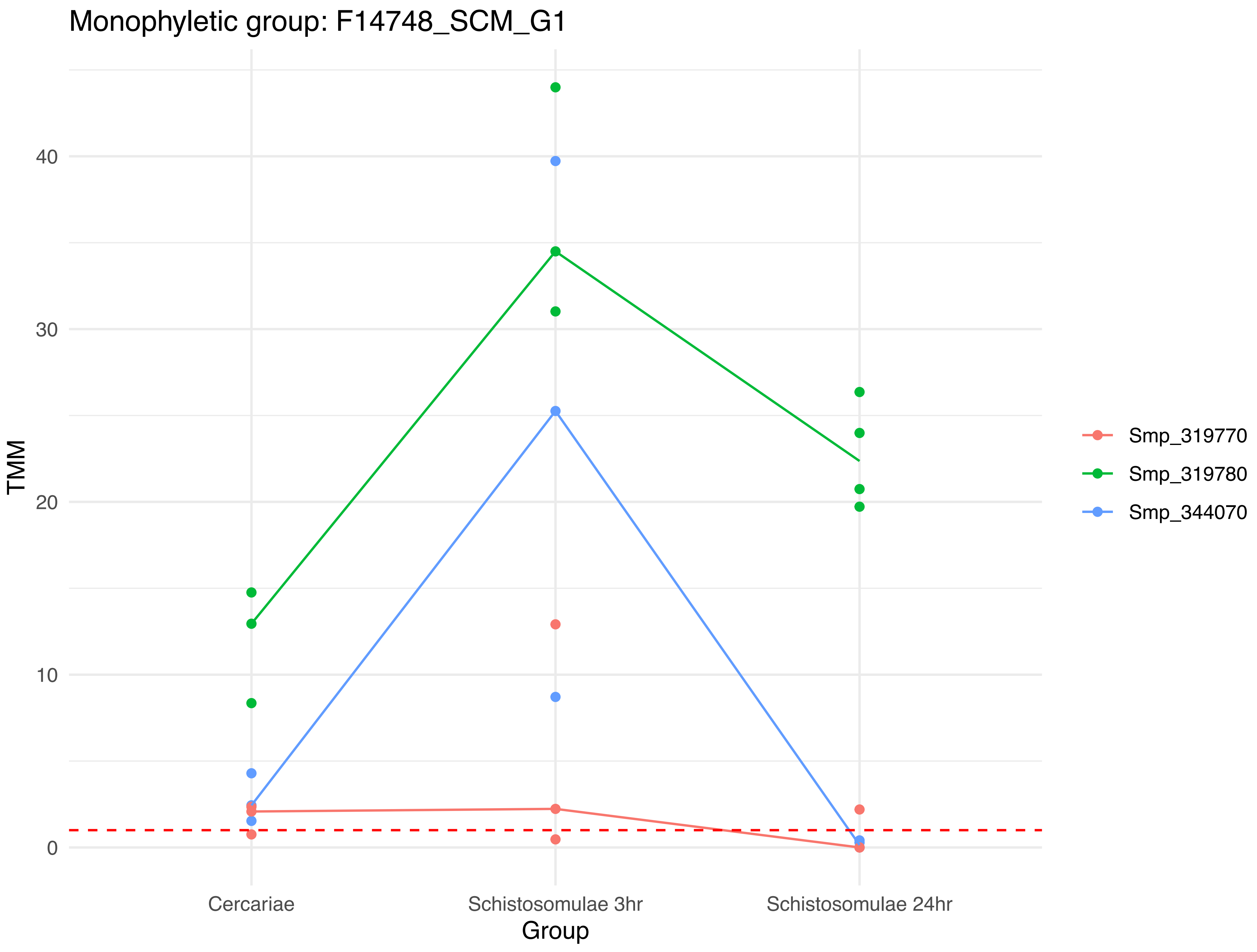

Monophyletic group: F14748_SCM_G1
TMM
40
30
20
10
0
Cercariae
Schistosomulae 3hr
Schistosomulae 24hr
Group
Smp_319770
Smp_319780
Smp_344070

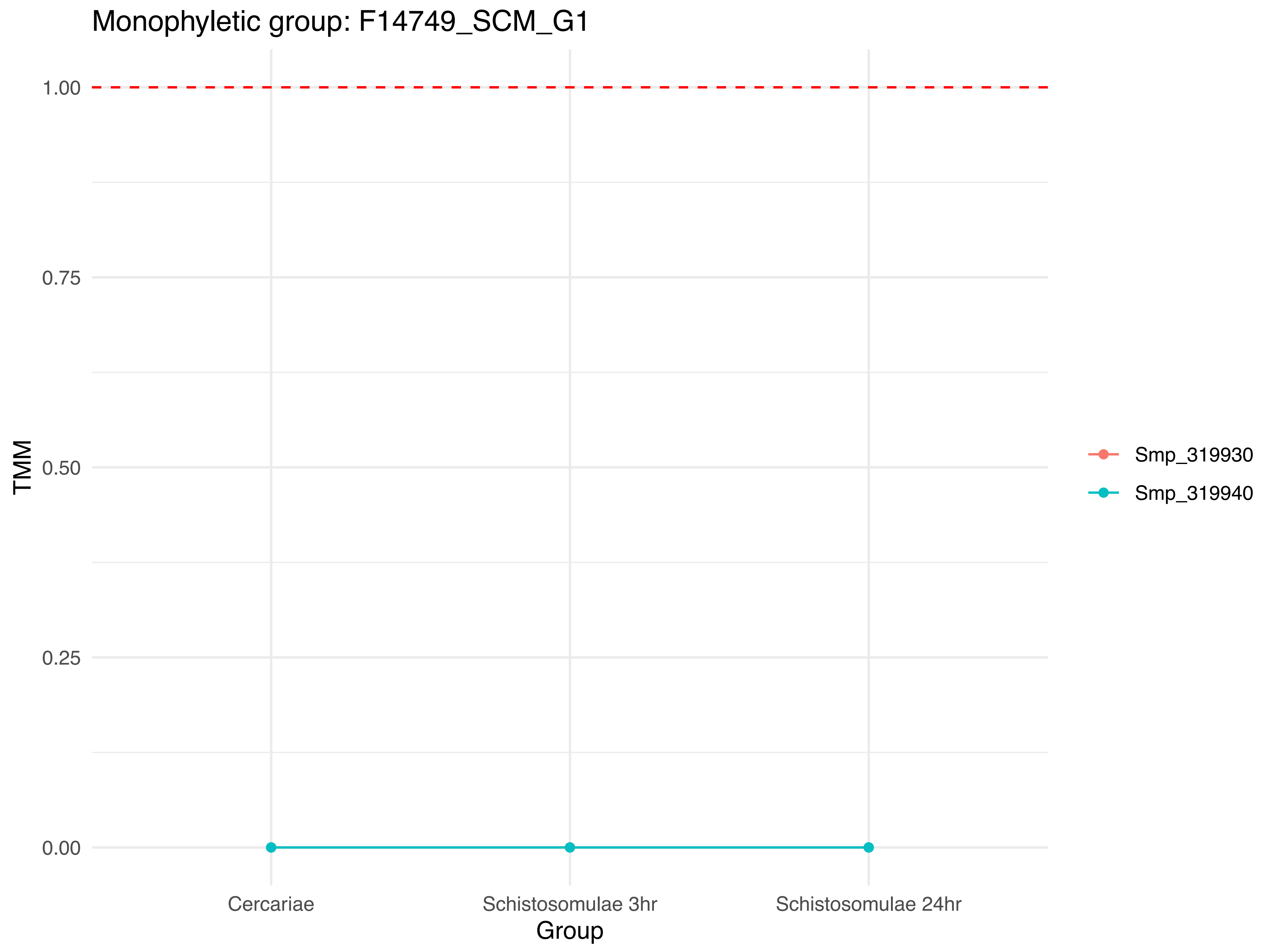
Monophyletic group: F14749_SCM_G1
TMM
1.00
0.75
0.50
0.25
0.00
Cercariae
Schistosomulae 3hr
Schistosomulae 24hr
Group
Smp_319930
Smp_319940

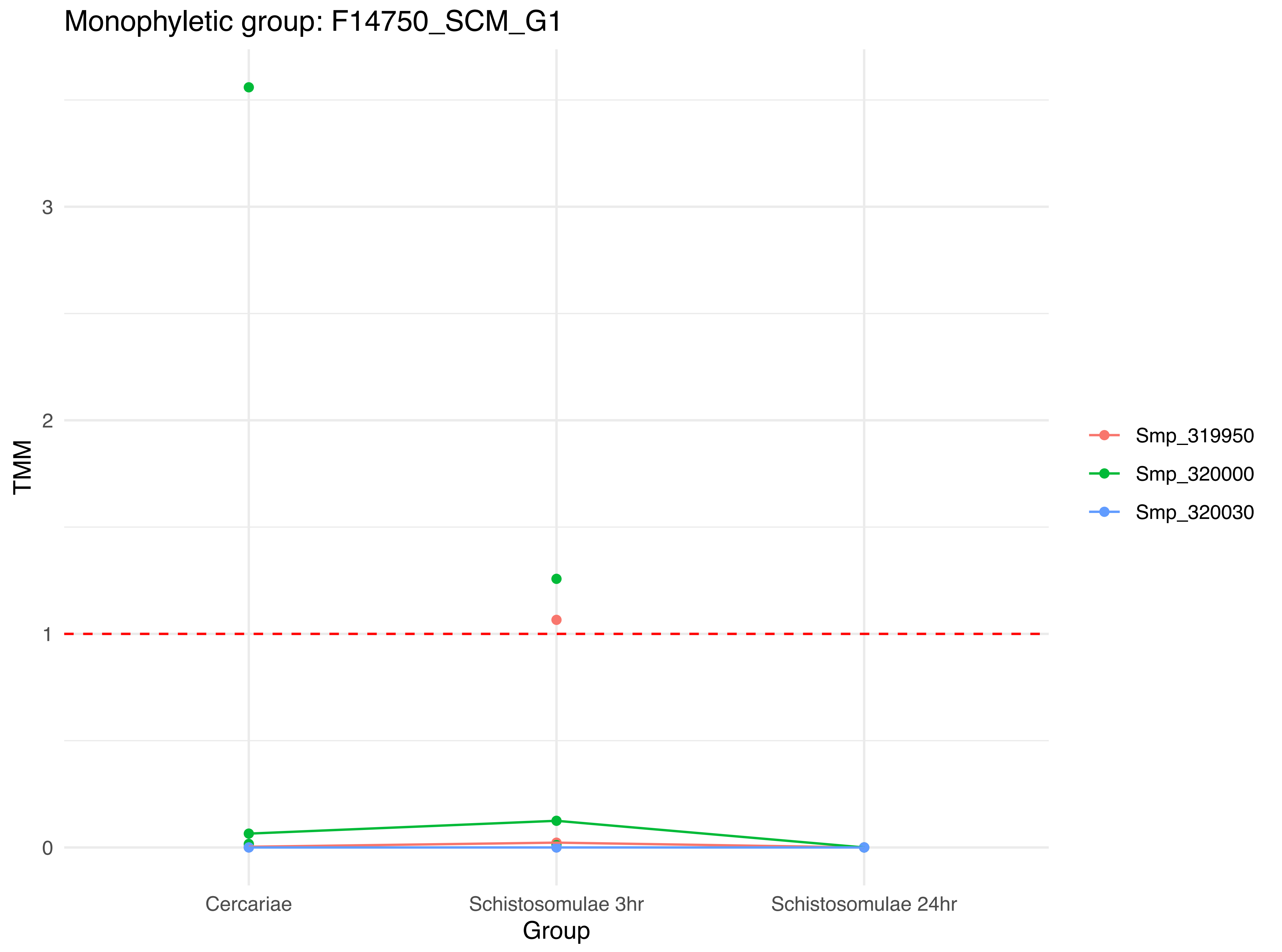

Monophyletic group: F14750_SCM_G1
TMM
3
2
1
0
Cercariae
Schistosomulae 3hr
Schistosomulae 24hr
Group
Smp_319950
Smp_320000
Smp_320030

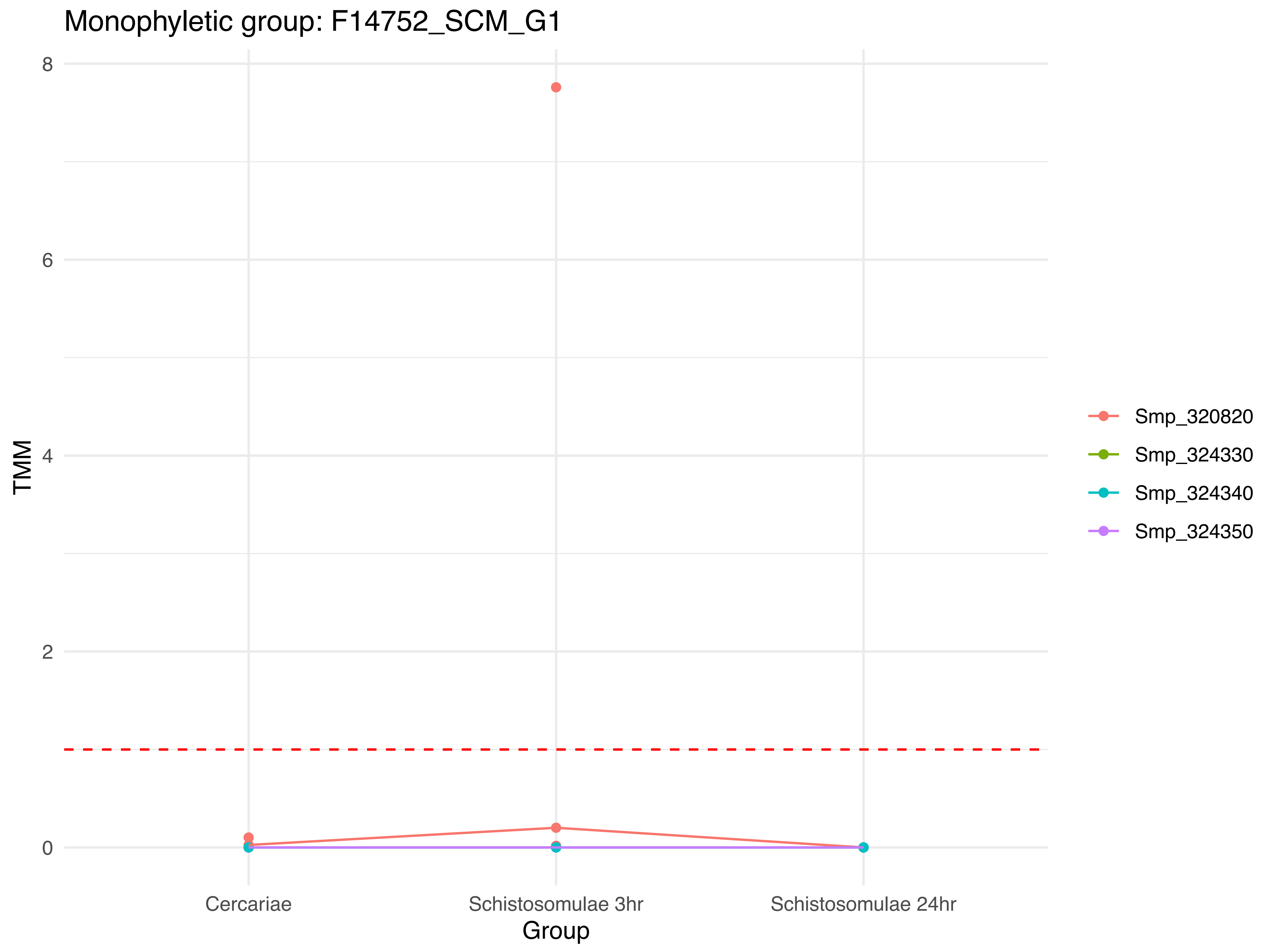
Monophyletic group: F14752_SCM_G1
TMM
8
6
4
2
0
Cercariae
Schistosomulae 3hr
Schistosomulae 24hr
Group
Smp_320820
Smp_324330
Smp_324340
Smp_324350

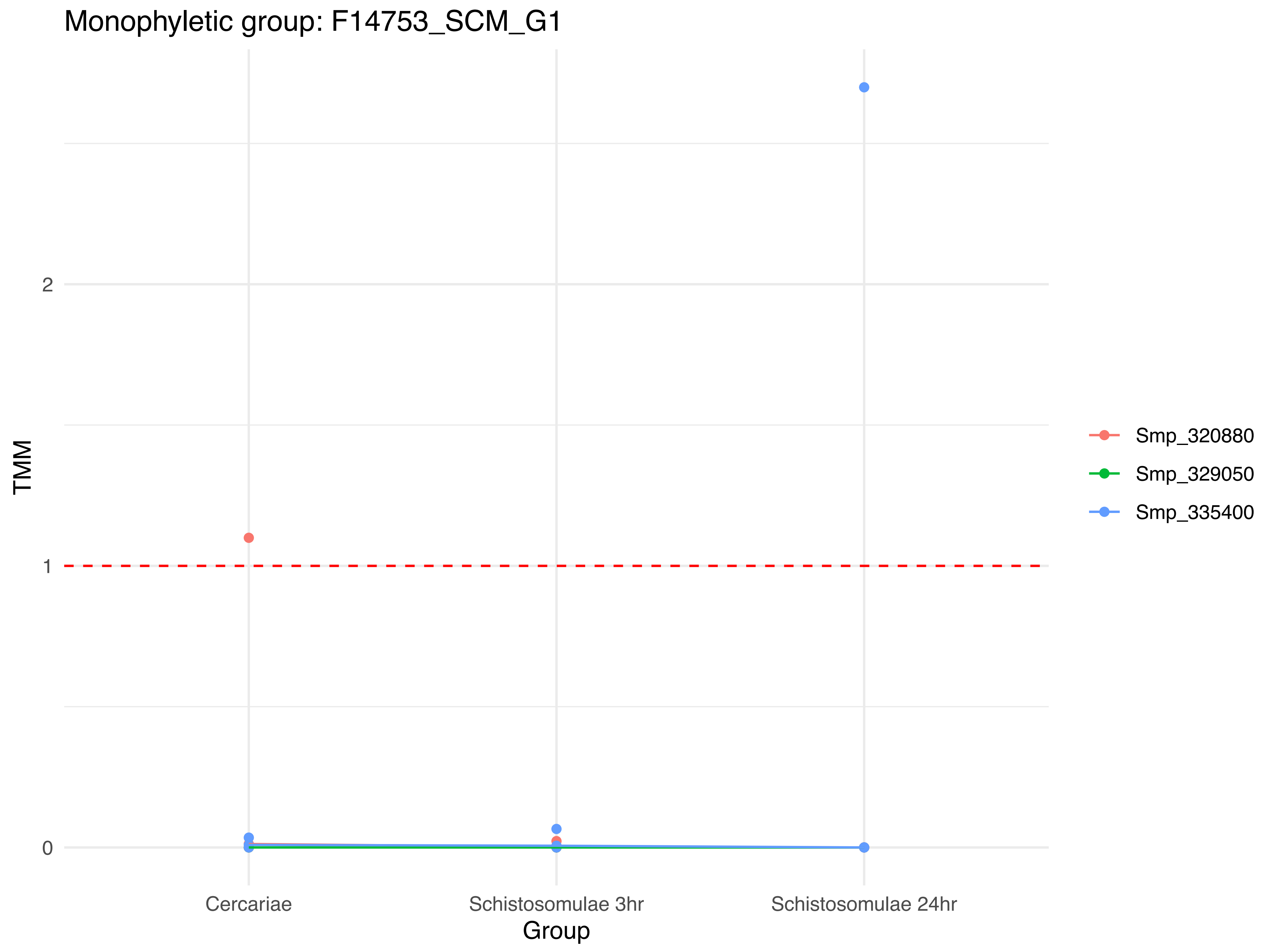
Monophyletic group: F14753_SCM_G1
TMM
2
1
0
Cercariae
Schistosomulae 3hr
Schistosomulae 24hr
Group
Smp_320880
Smp_329050
Smp_335400

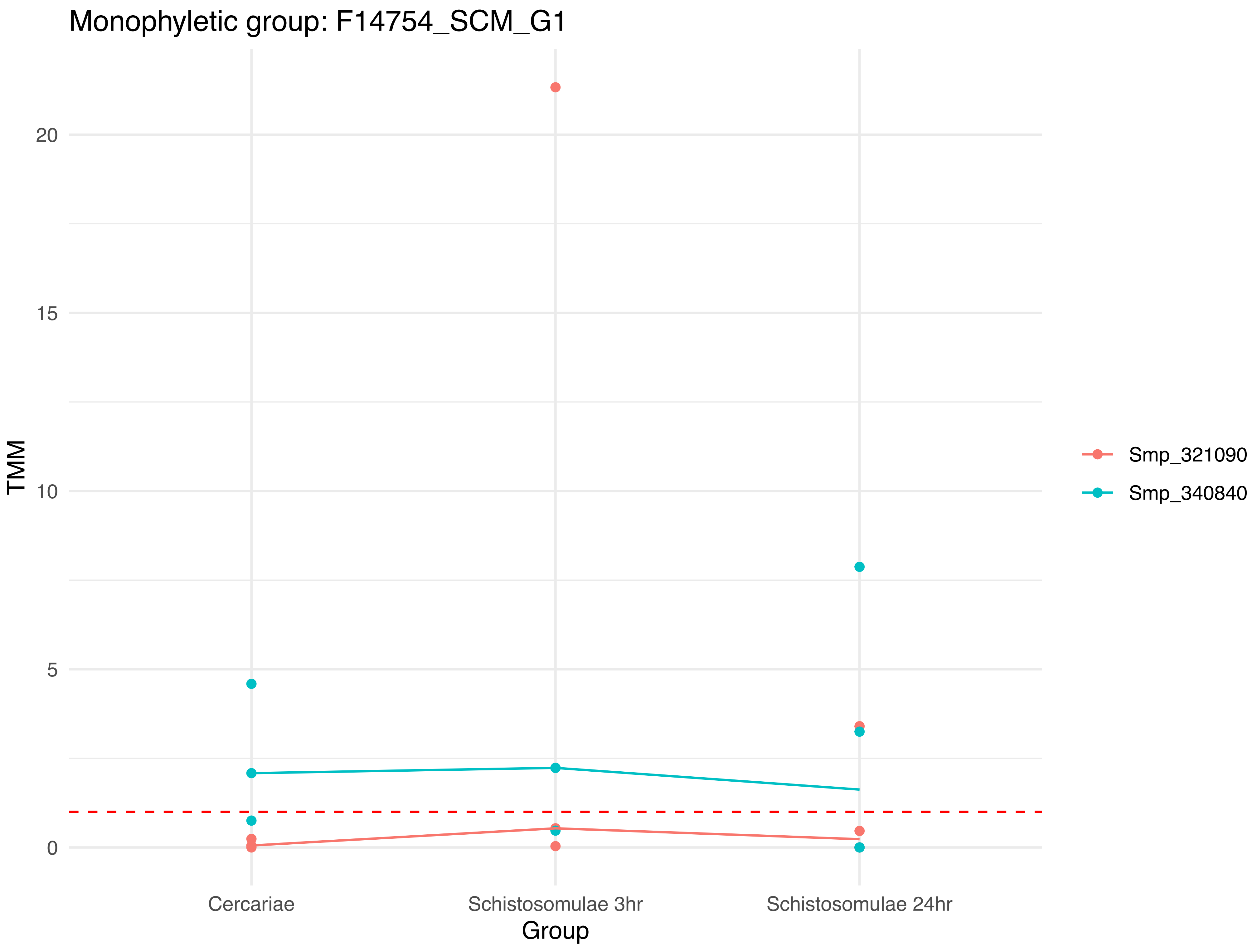
Monophyletic group: F14754_SCM_G1
20
15
10
5
0
TMM
Cercariae
Schistosomulae 3hr
Schistosomulae 24hr
Group
Smp_321090
Smp_340840

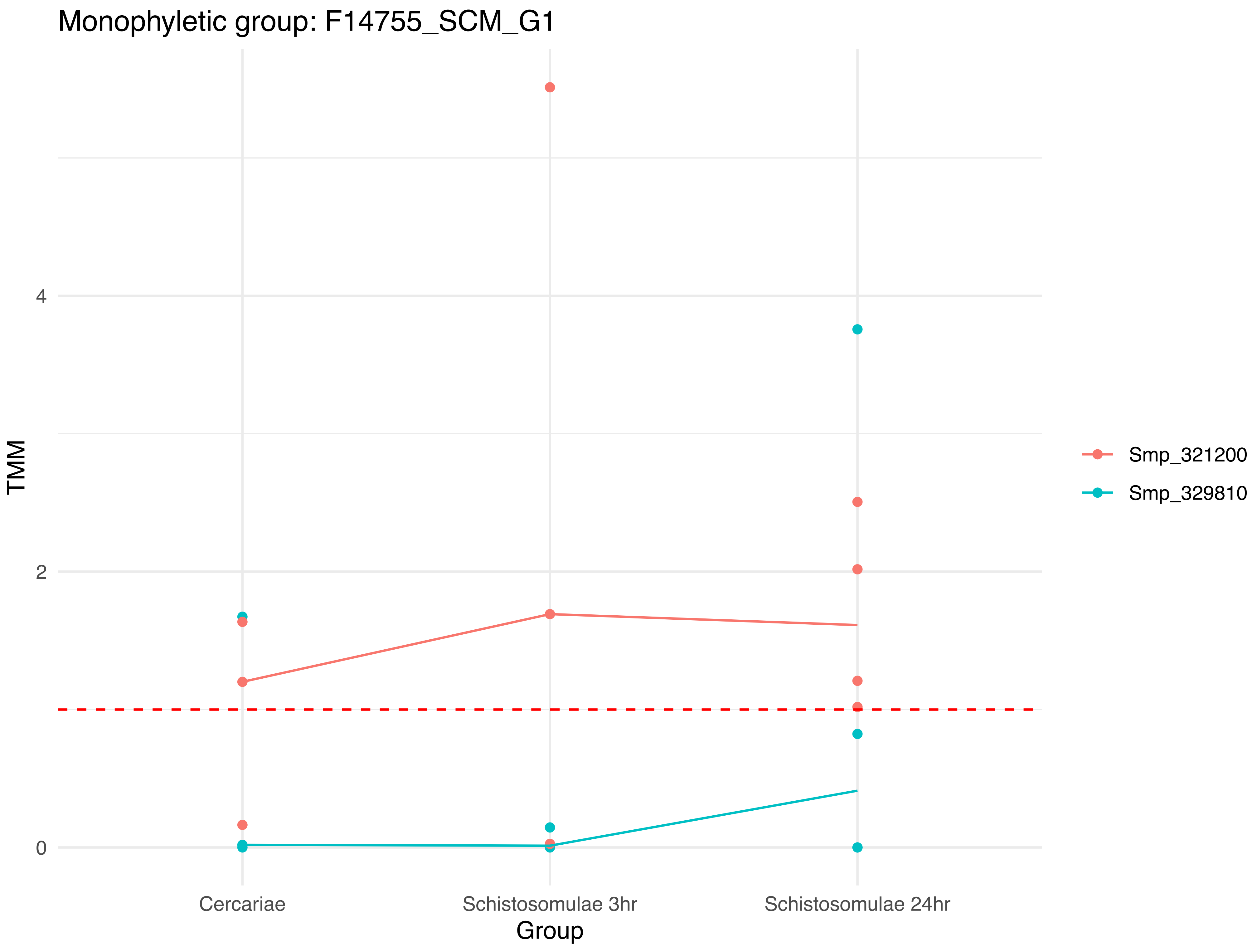

Monophyletic group: F14755_SCM_G1
TMM
4
2
0
Cercariae
Schistosomulae 3hr
Schistosomulae 24hr
Group
Smp_321200
Smp_329810

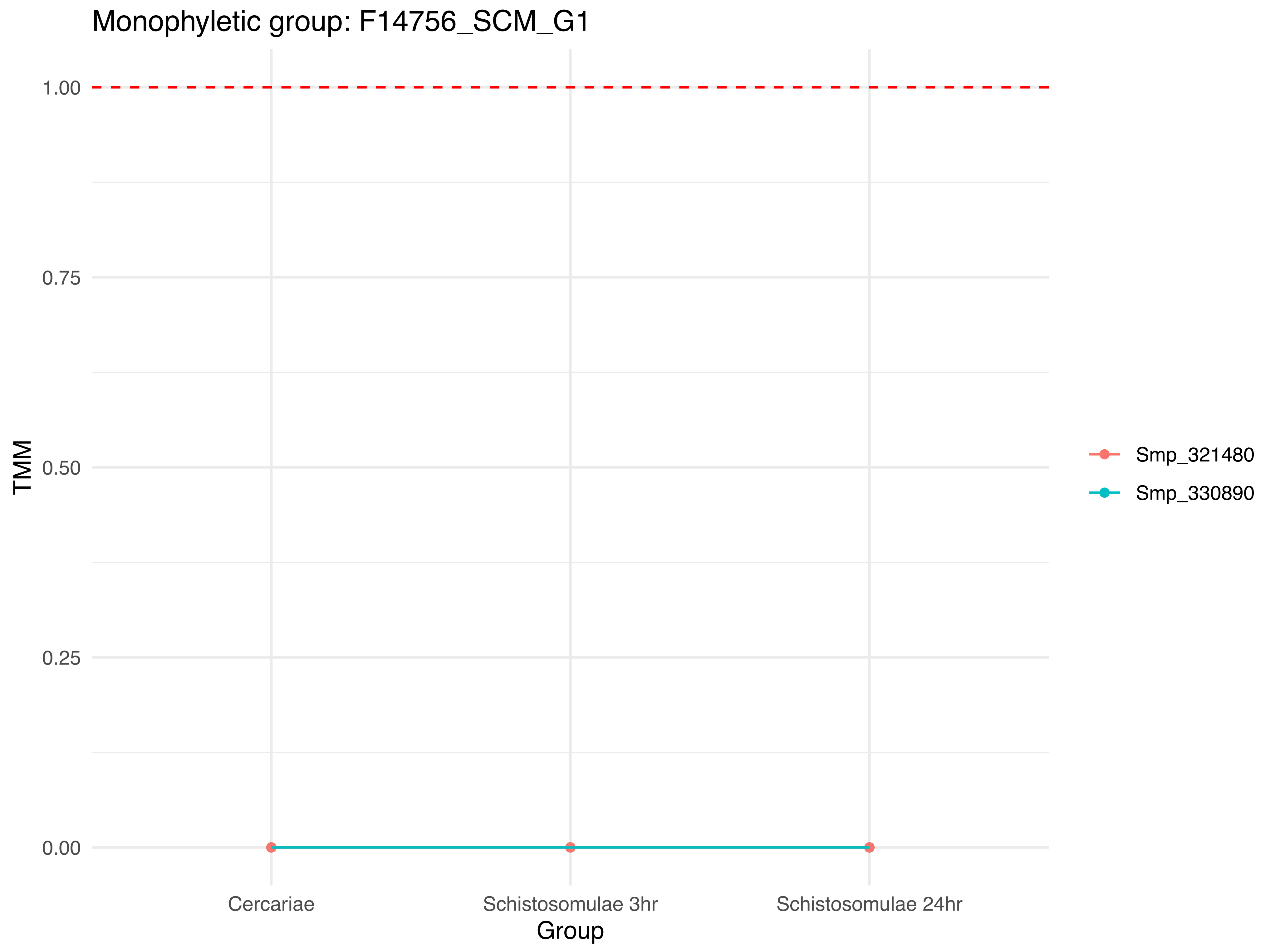
Monophyletic group: F14756_SCM_G1
TMM
1.00
0.75
0.50
0.25
0.00
Cercariae
Schistosomulae 3hr
Schistosomulae 24hr
Group
Smp_321480
Smp_330890

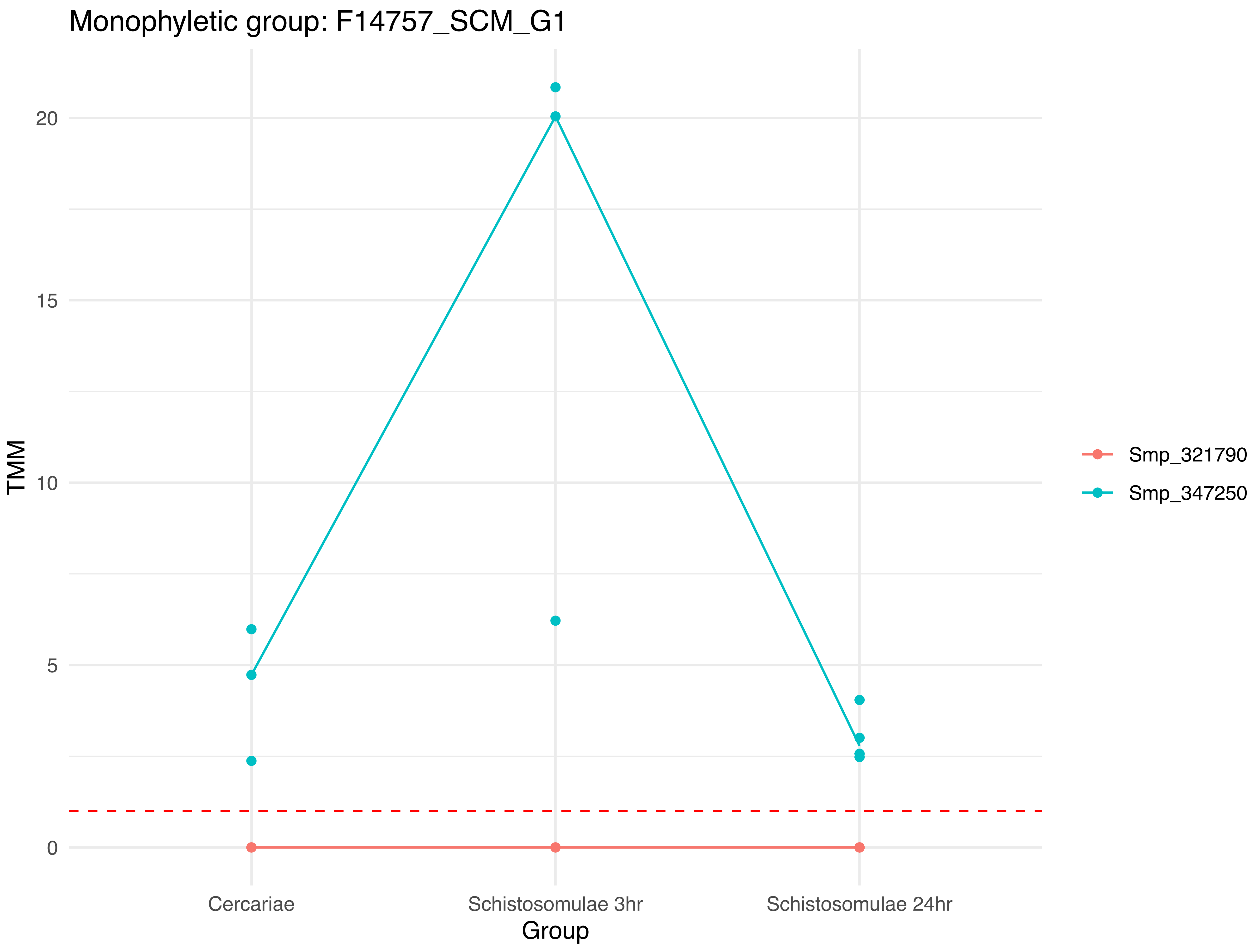

Monophyletic group: F14757_SCM_G1
TMM
20
15
10
5
0
Cercariae
Schistosomulae 3hr
Schistosomulae 24hr
Group
Smp_321790
Smp_347250

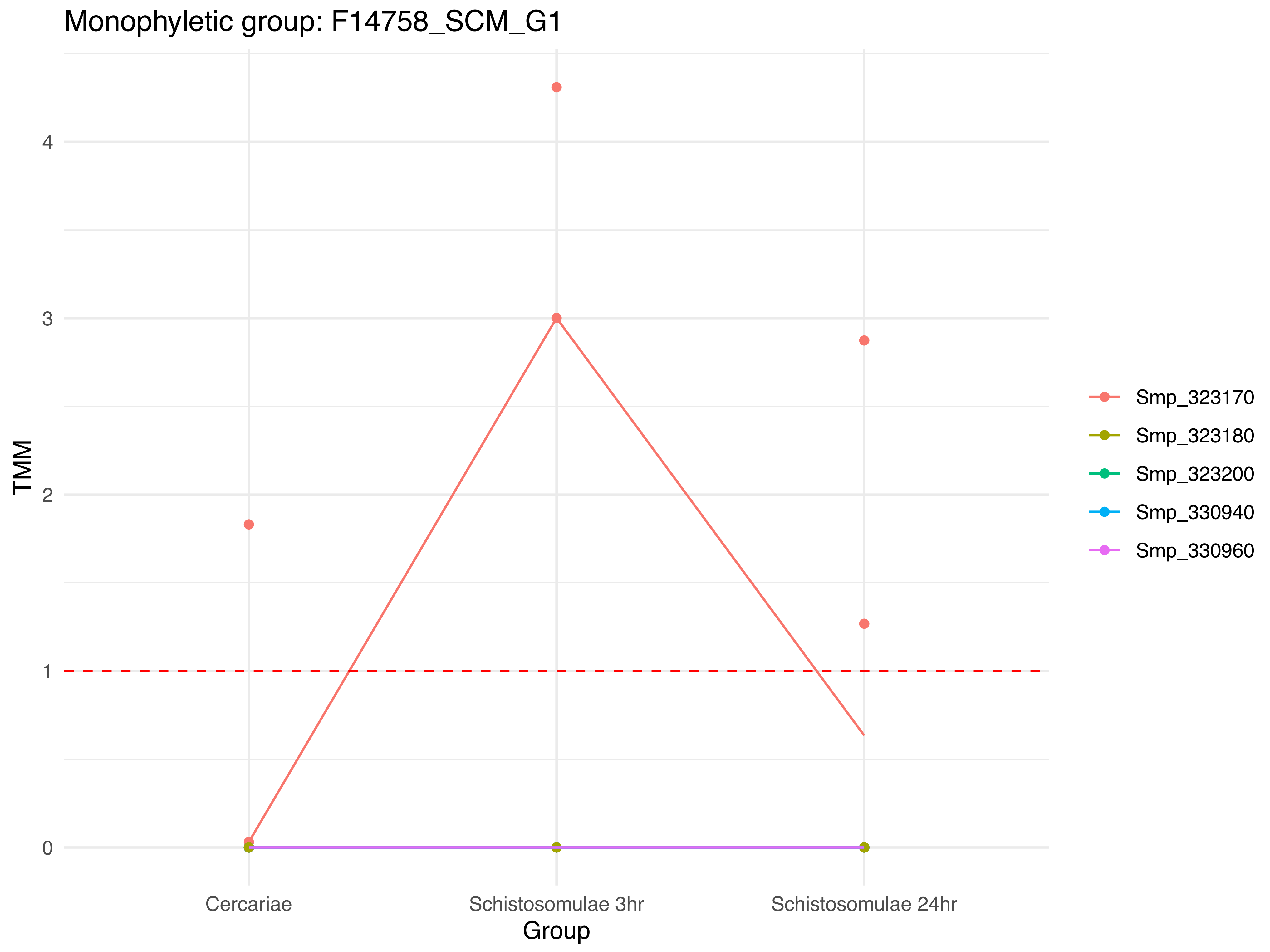

Monophyletic group: F14758_SCM_G1
TMM
4
3
2
1
0
Cercariae
Schistosomulae 3hr
Schistosomulae 24hr
Group
Smp_323170
Smp_323180
Smp_323200
Smp_330940
Smp_330960

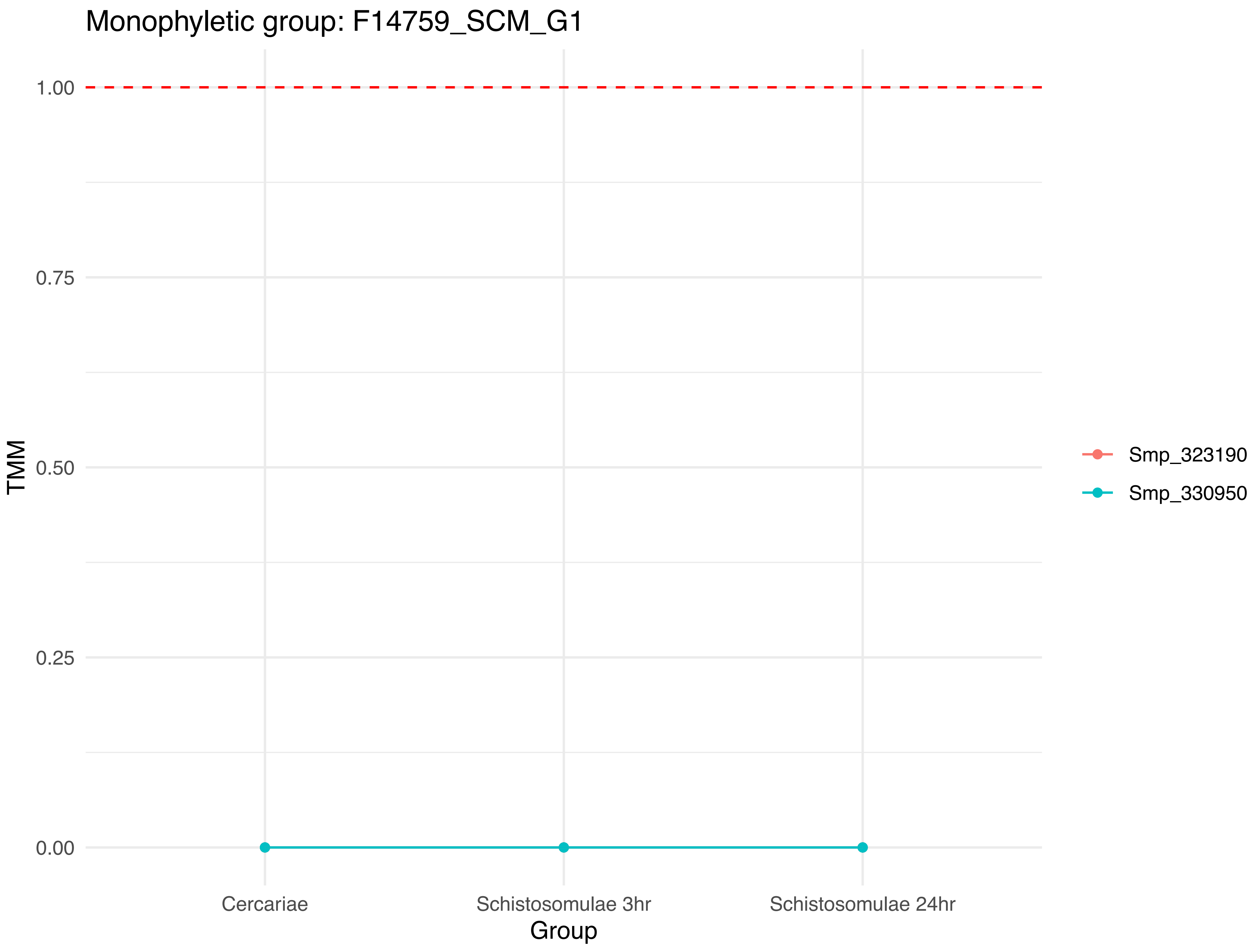
Monophyletic group: F14759_SCM_G1
1.00
0.75
0.50
0.25
0.00
TMM
Cercariae
Schistosomulae 3hr
Schistosomulae 24hr
Group
Smp_323190
Smp_330950

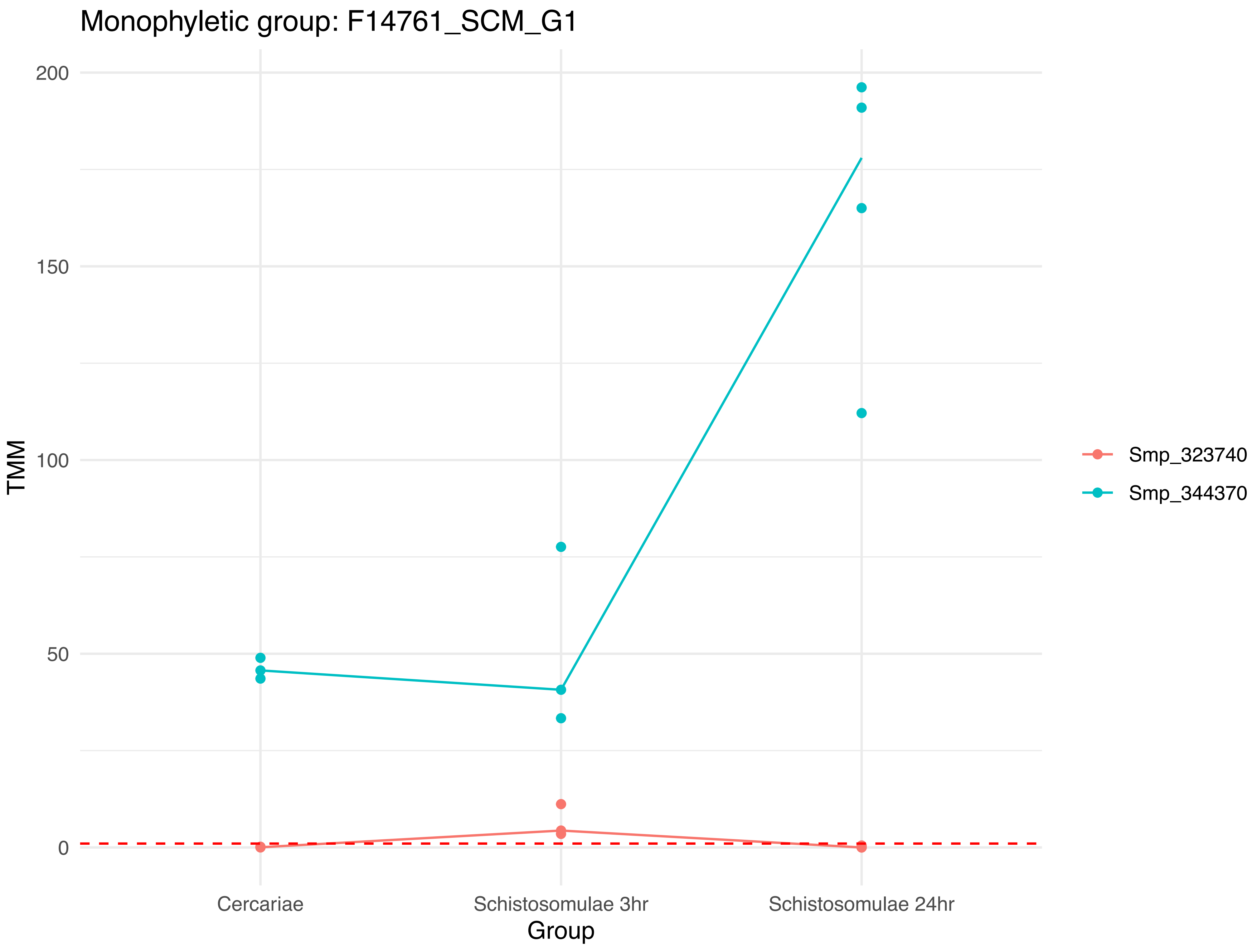

Monophyletic group: F14761_SCM_G1
200
150
100
50
0
TMM
Cercariae
Schistosomulae 3hr
Schistosomulae 24hr
Group
Smp_323740
Smp_344370

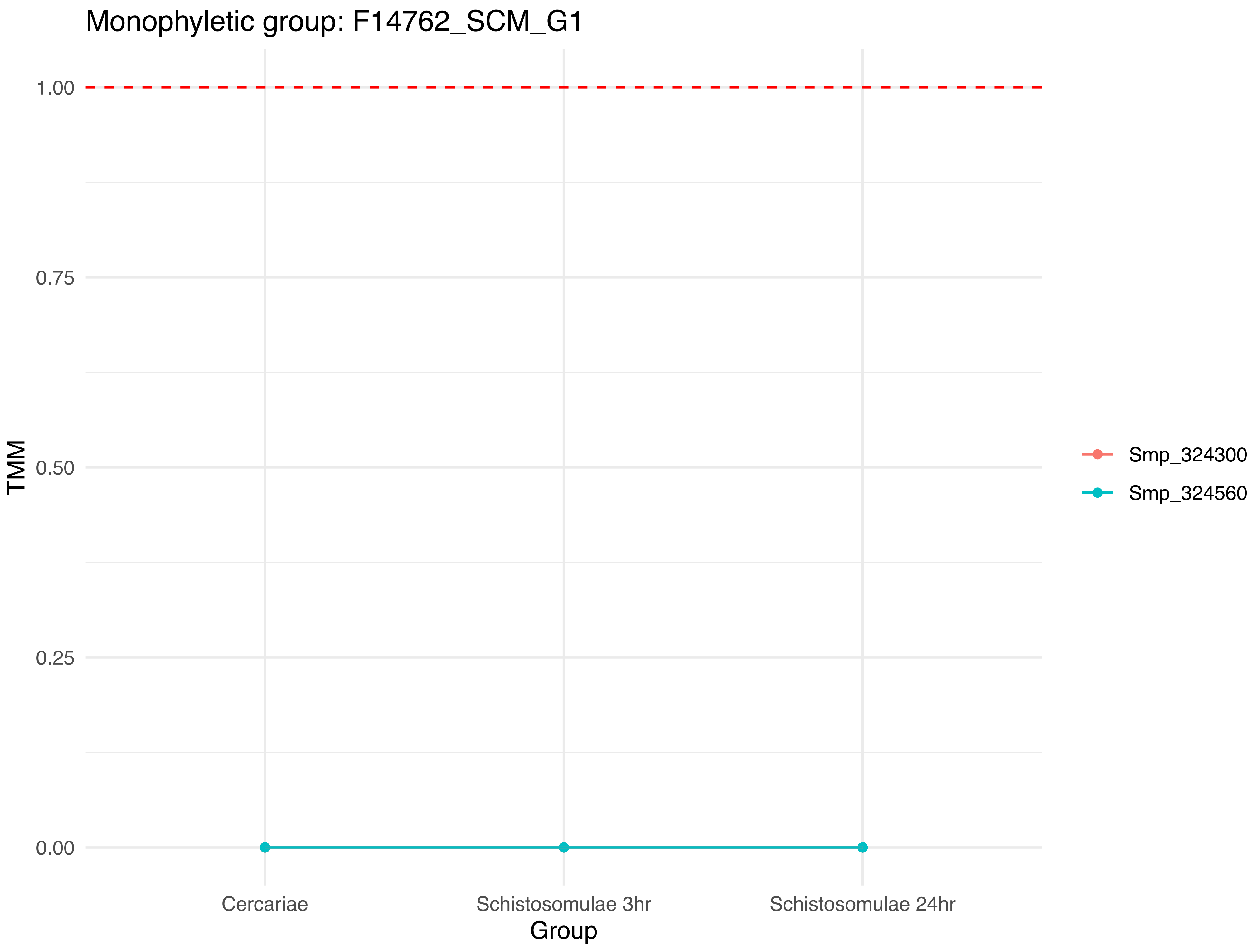
Monophyletic group: F14762_SCM_G1
1.00
0.75
0.50
0.25
0.00
TMM
Cercariae
Schistosomulae 3hr
Schistosomulae 24hr
Group
Smp_324300
Smp_324560

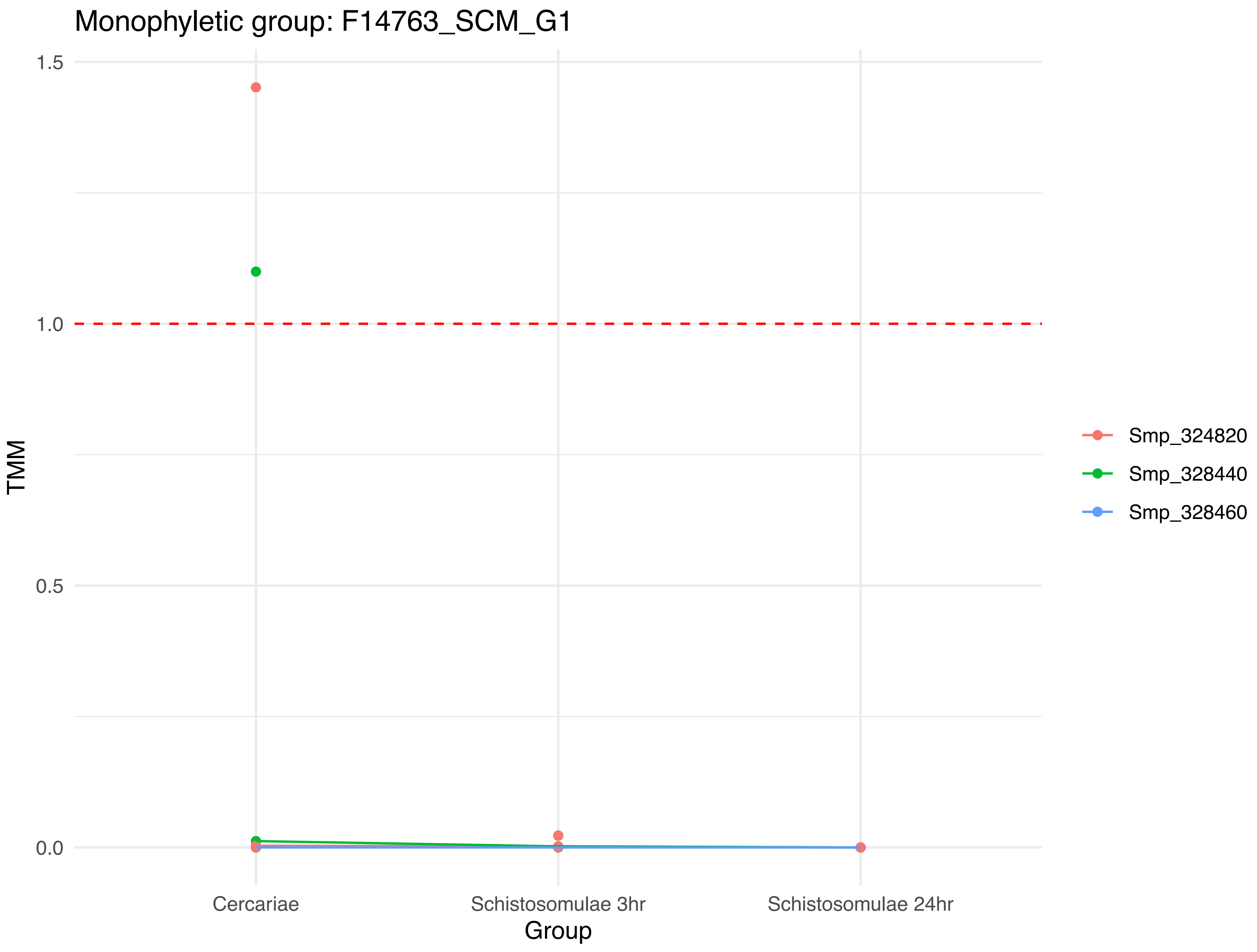

Monophyletic group: F14763_SCM_G1
1.5
1.0
0.5
0.0
TMM
Cercariae
Schistosomulae 3hr
Schistosomulae 24hr
Group
Smp_324820
Smp_328440
Smp_328460

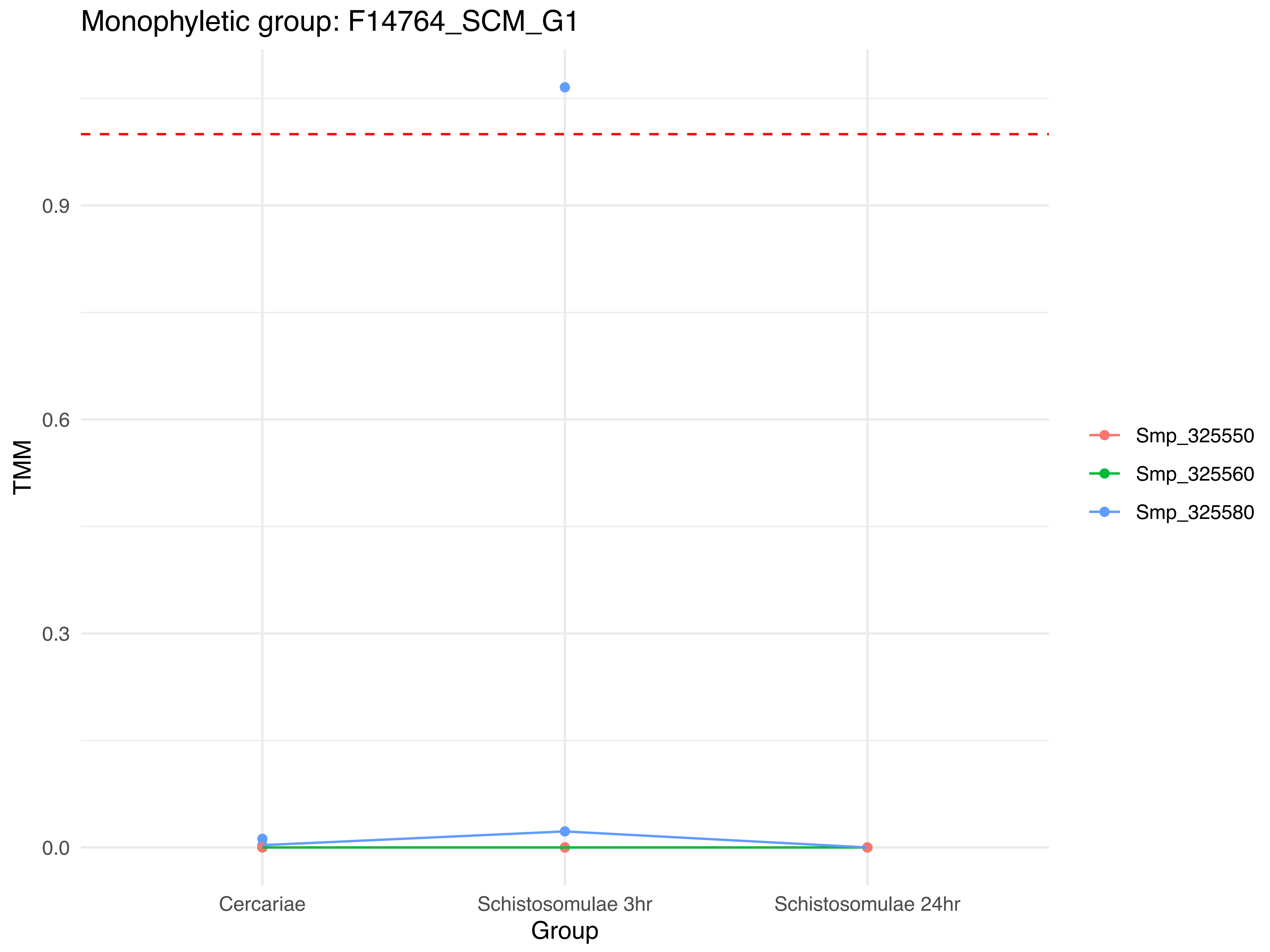

Monophyletic group: F14764_SCM_G1
TMM
0.9
0.6
0.3
0.0
Cercariae
Schistosomulae 3hr
Schistosomulae 24hr
Group
Smp_325550
Smp_325560
Smp_325580

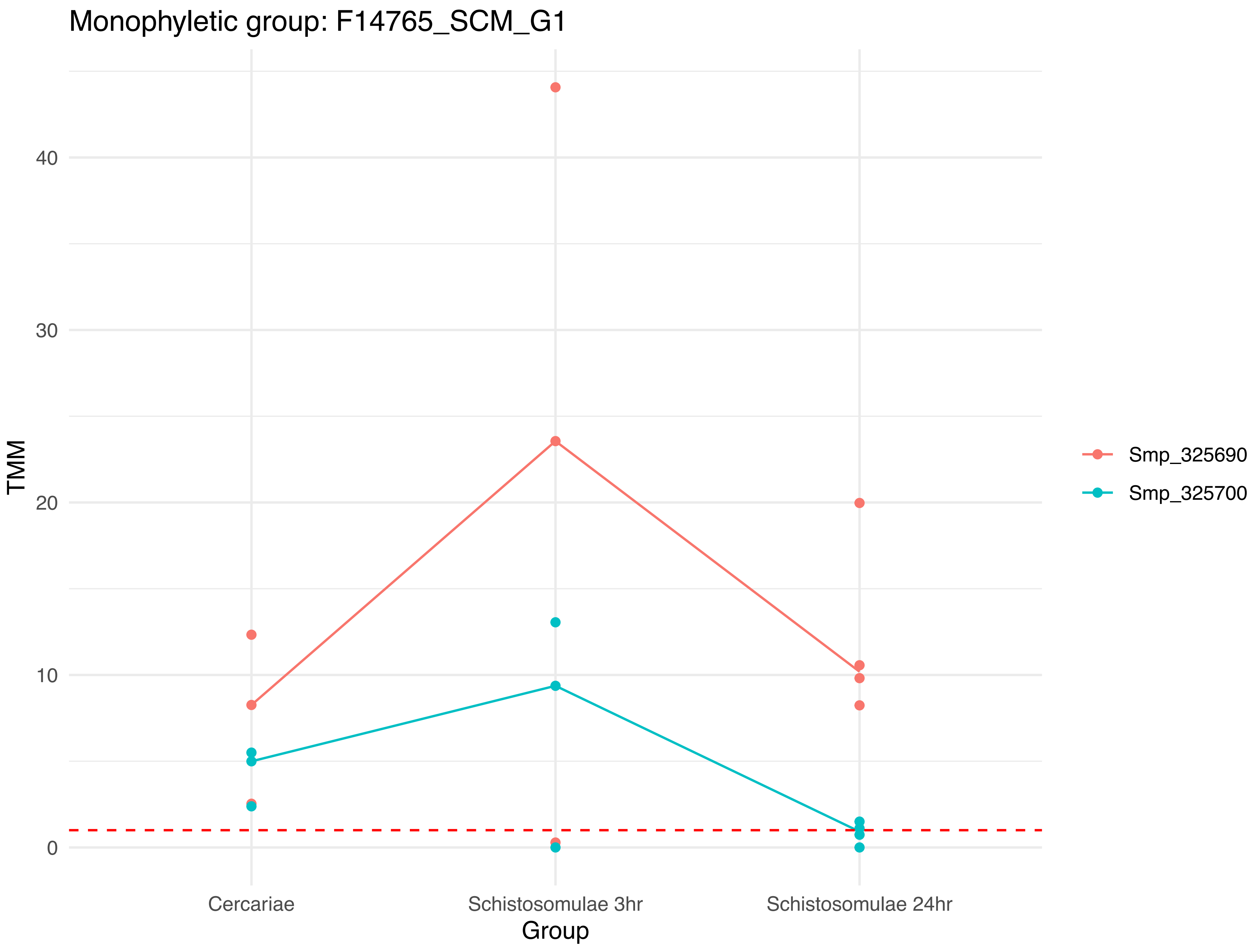
Monophyletic group: F14765_SCM_G1
TMM
40
30
20
10
0
Cercariae
Schistosomulae 3hr
Schistosomulae 24hr
Group
Smp_325690
Smp_325700

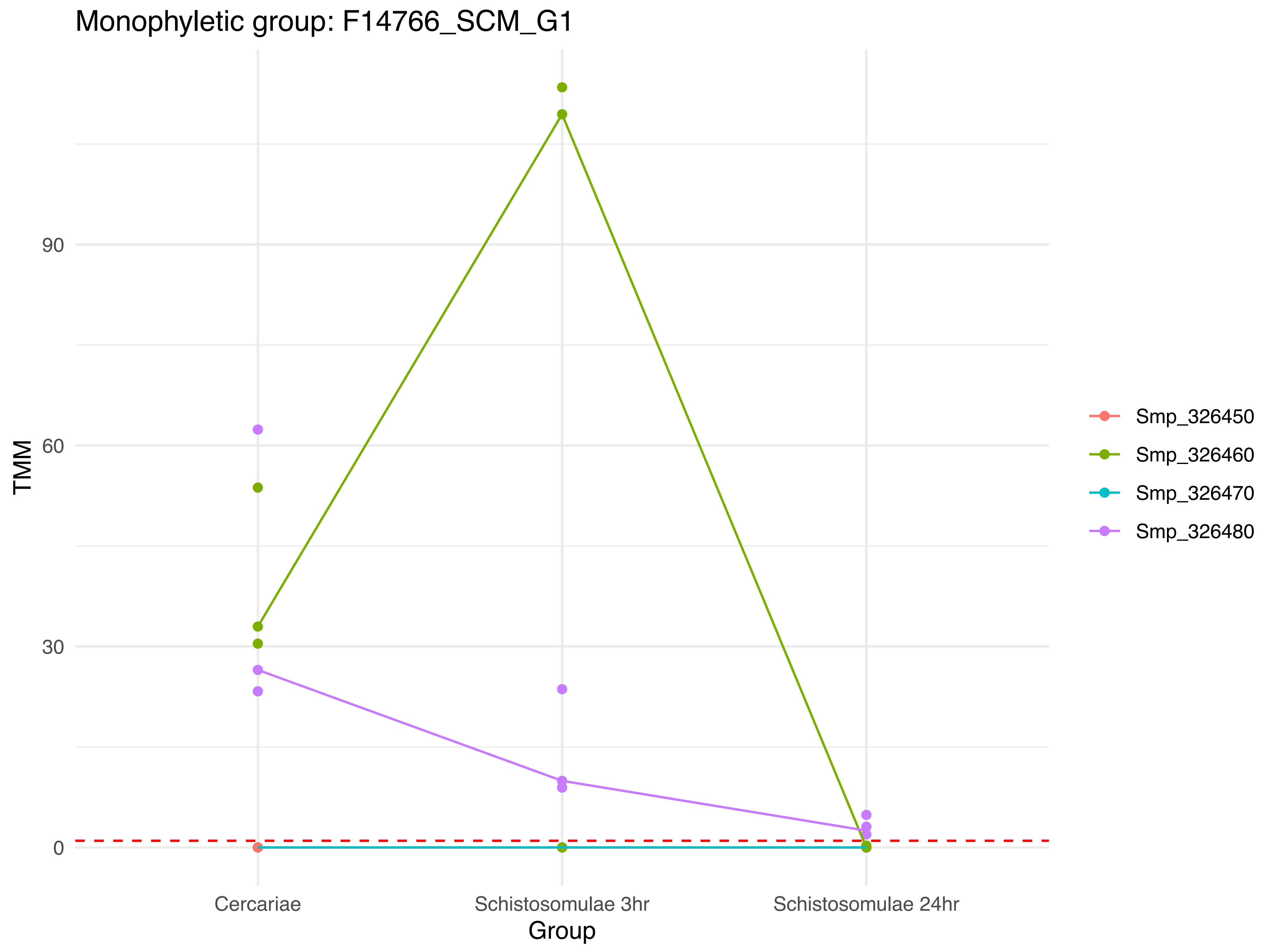

Monophyletic group: F14766_SCM_G1
TMM
90
60
30
0
Cercariae
Schistosomulae 3hr
Schistosomulae 24hr
Group
Smp_326450
Smp_326460
Smp_326470
Smp_326480

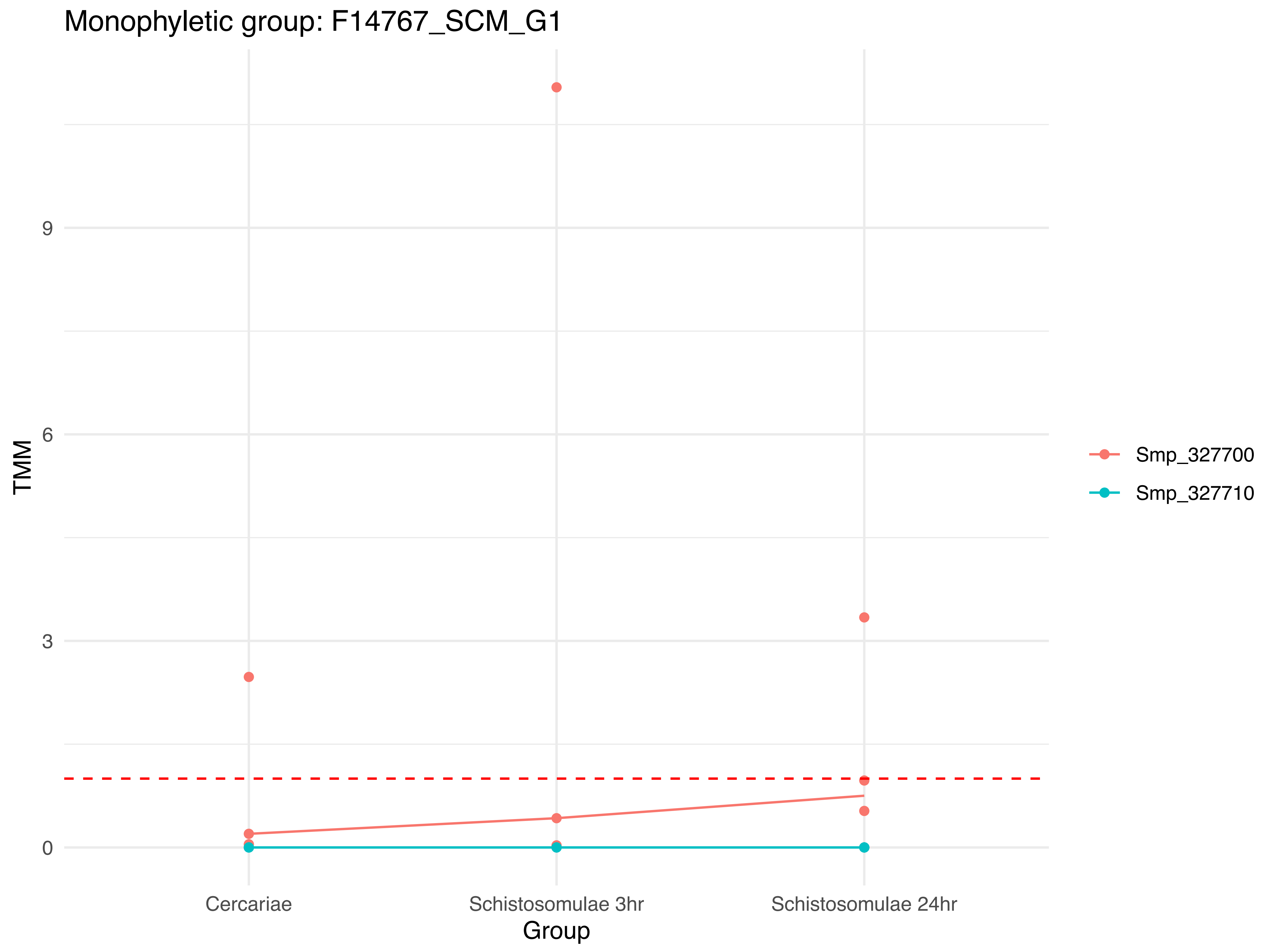
Monophyletic group: F14767_SCM_G1
TMM
0
3
6
9
Cercariae
Schistosomulae 3hr
Schistosomulae 24hr
Group
Smp_327700
Smp_327710

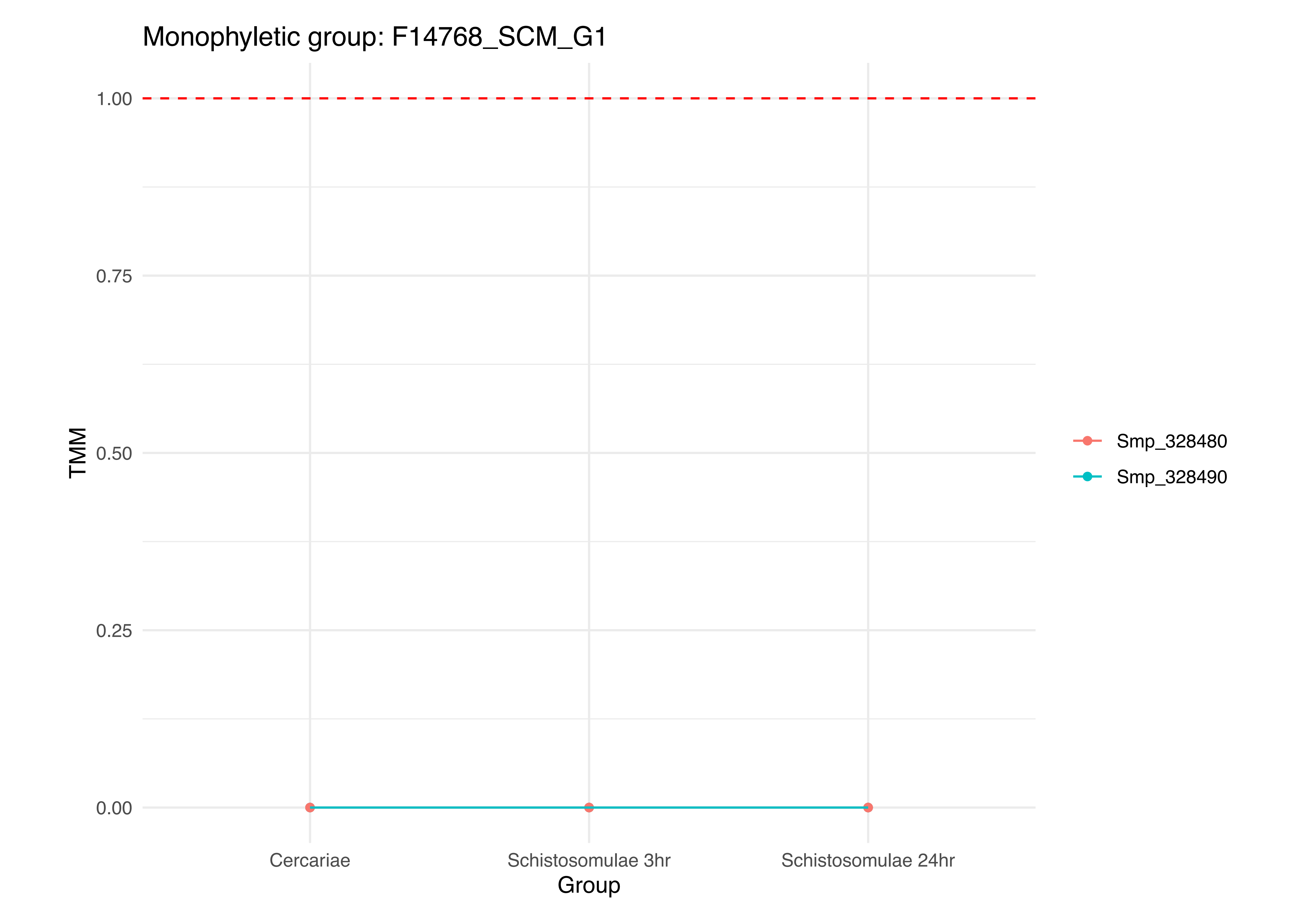

Monophyletic group: F14768_SCM_G1
1.00
0.75
0.50
0.25
0.00
TMM
Cercariae
Schistosomulae 3hr
Schistosomulae 24hr
Group
Smp_328480
Smp_328490

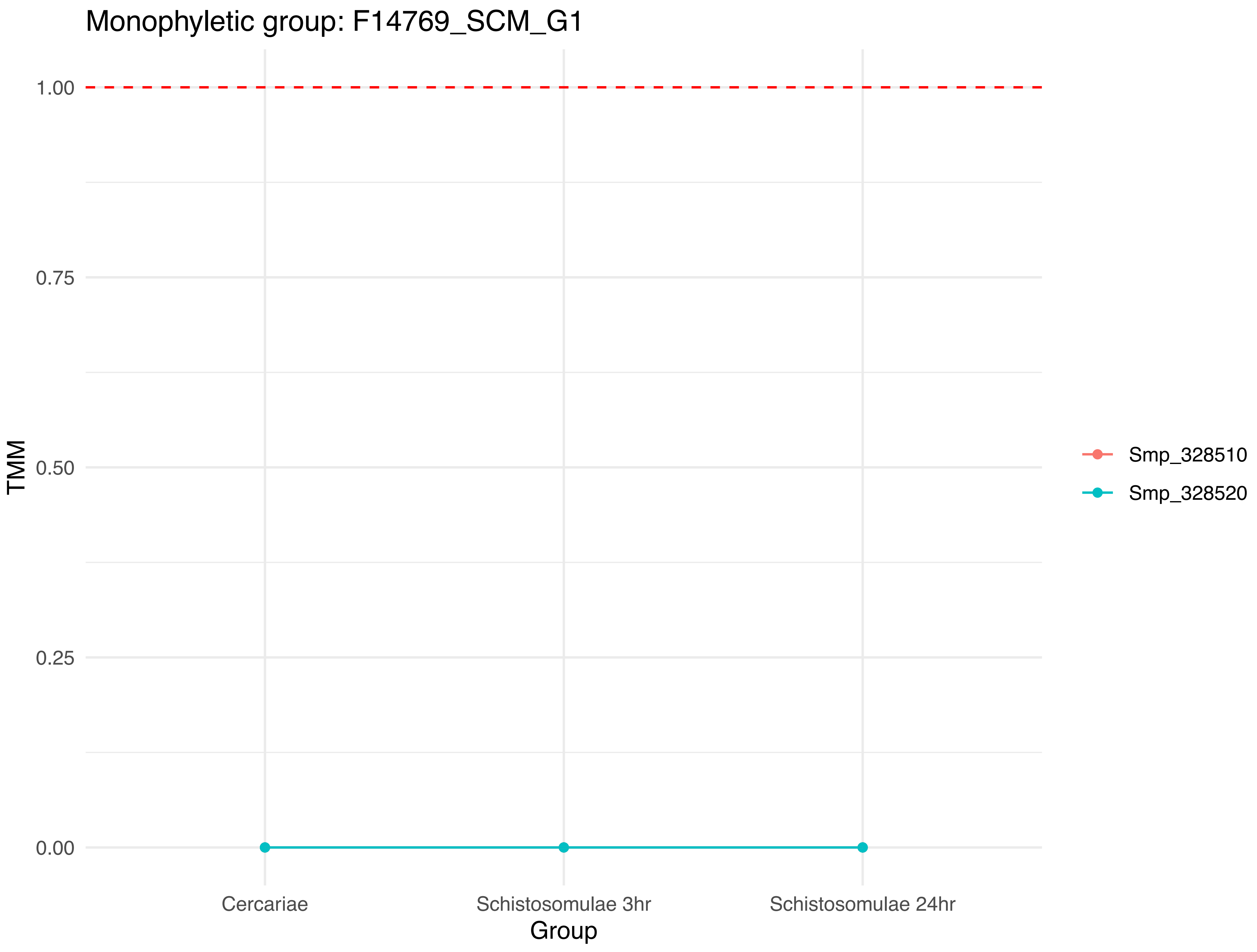

Monophyletic group: F14769_SCM_G1
1.00
0.75
0.50
0.25
0.00
TMM
Cercariae
Schistosomulae 3hr
Schistosomulae 24hr
Group
Smp_328510
Smp_328520

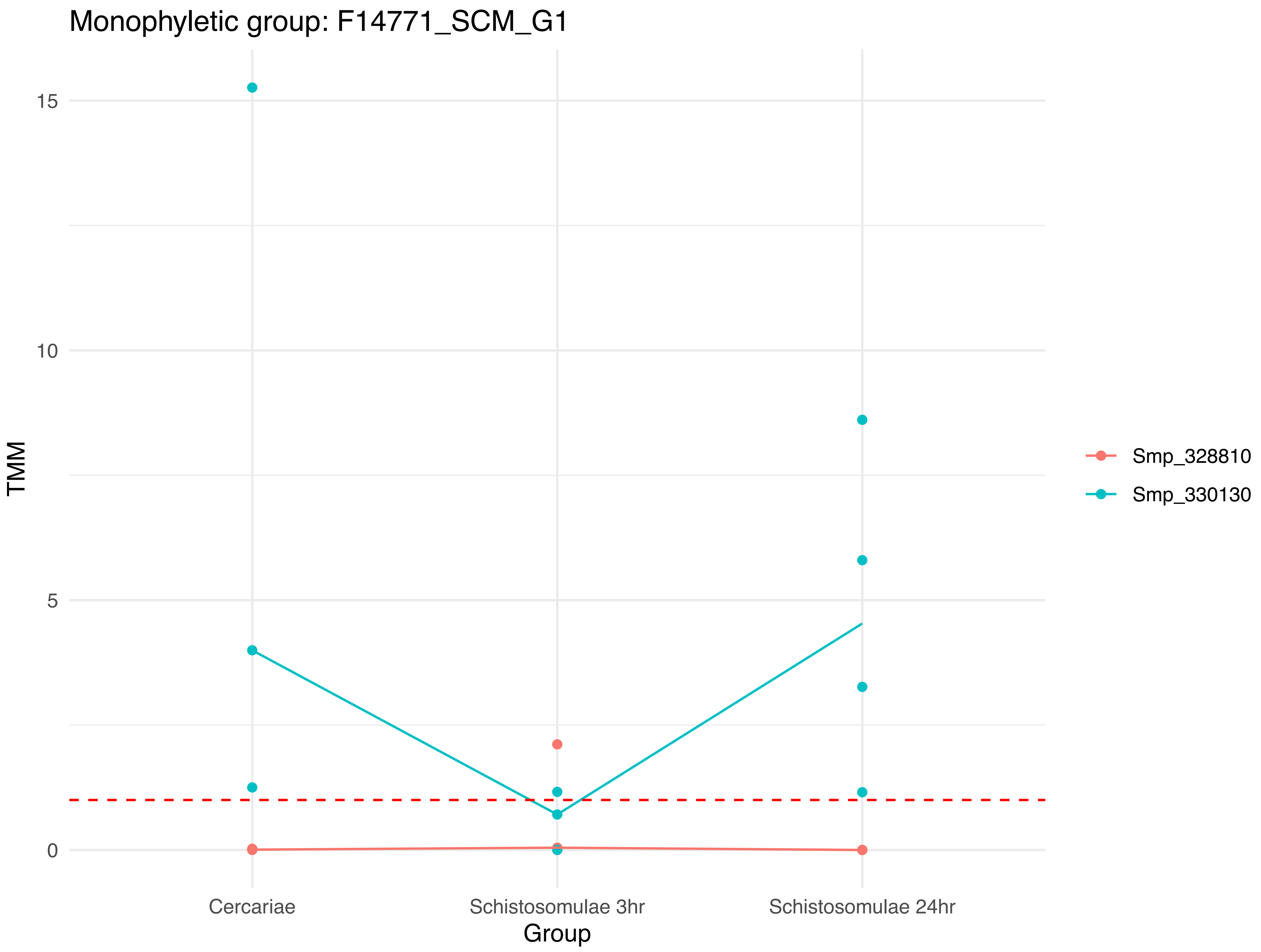

Monophyletic group: F14771_SCM_G1
TMM
15
10
5
0
Cercariae
Schistosomulae 3hr
Schistosomulae 24hr
Group
Smp_328810
Smp_330130

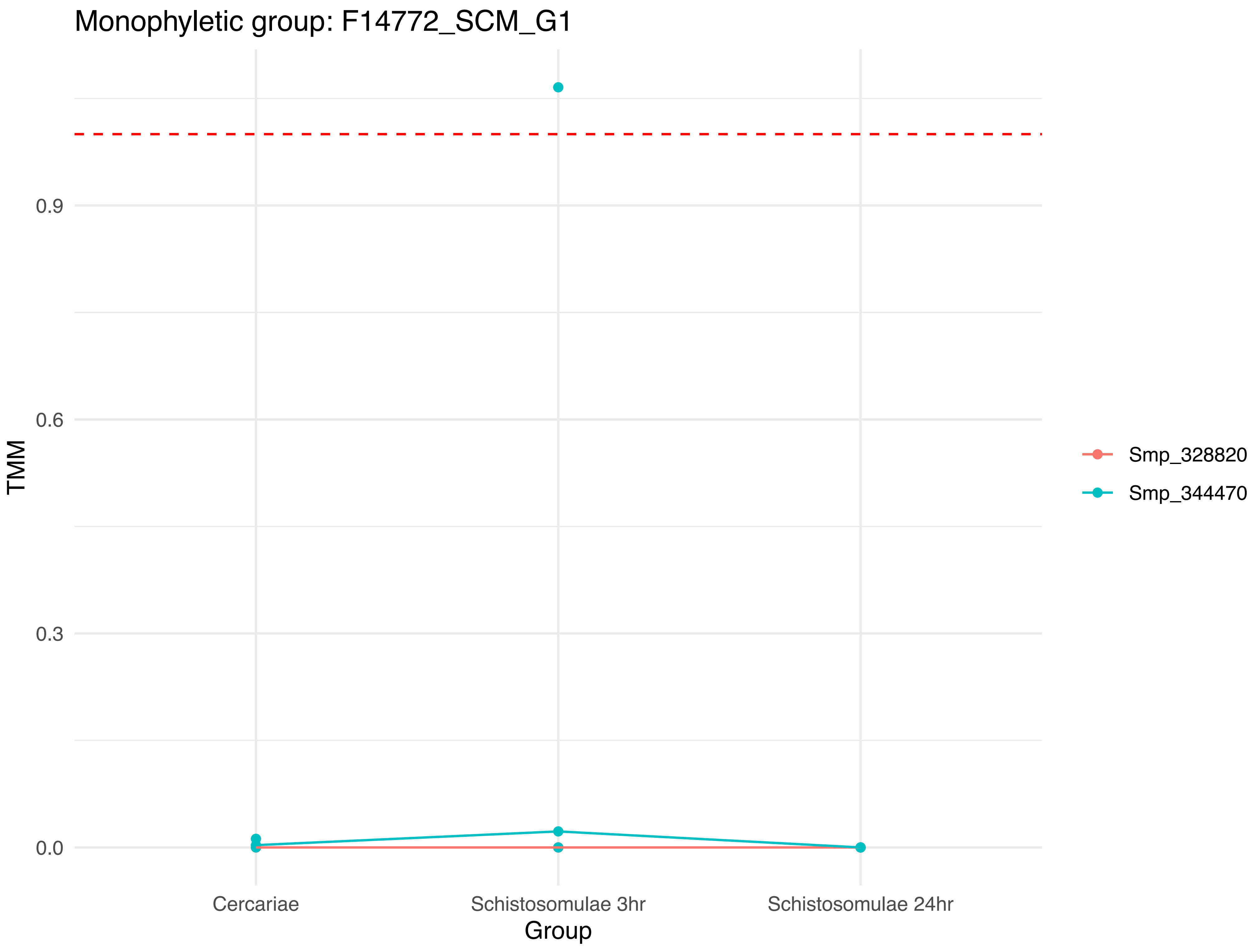

Monophyletic group: F14772_SCM_G1
TMM
0.9
0.6
0.3
0.0
Cercariae
Schistosomulae 3hr
Schistosomulae 24hr
Group
Smp_328820
Smp_344470

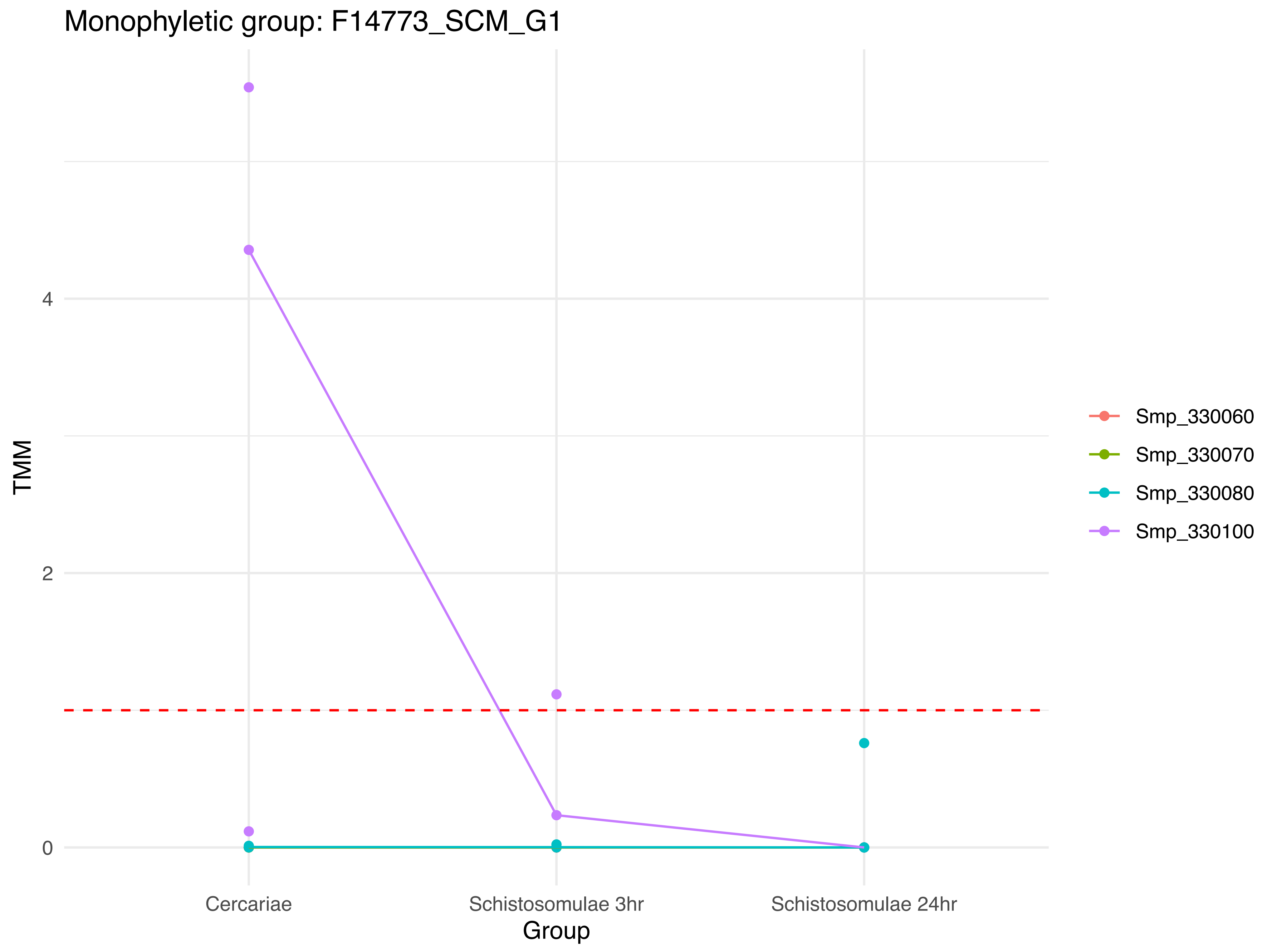
Monophyletic group: F14773_SCM_G1
TMM
0
2
4
Cercariae
Schistosomulae 3hr
Schistosomulae 24hr
Group
Smp_330060
Smp_330070
Smp_330080
Smp_330100

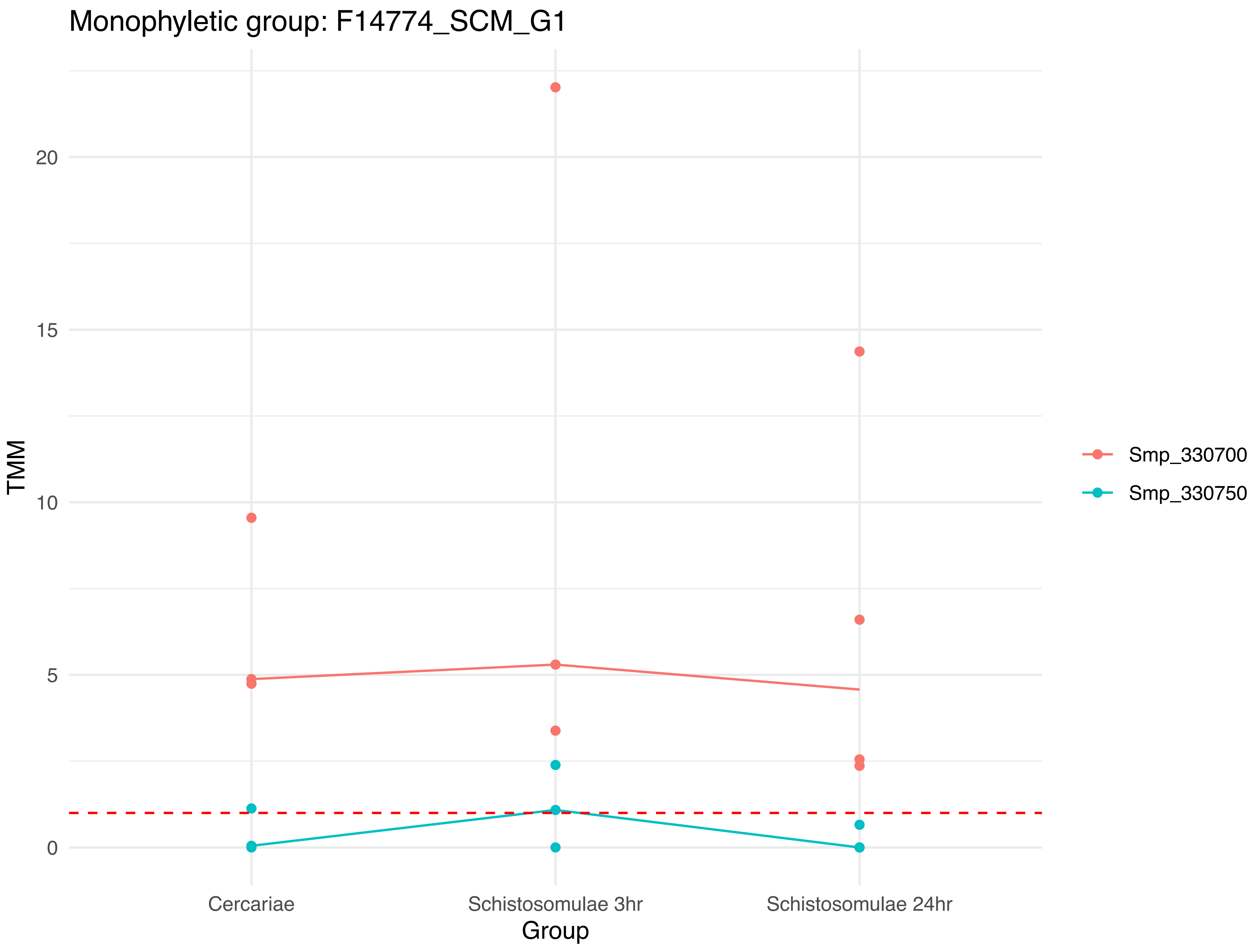
Monophyletic group: F14774_SCM_G1
TMM
20
15
10
5
0
Cercariae
Schistosomulae 3hr
Schistosomulae 24hr
Group
Smp_330700
Smp_330750

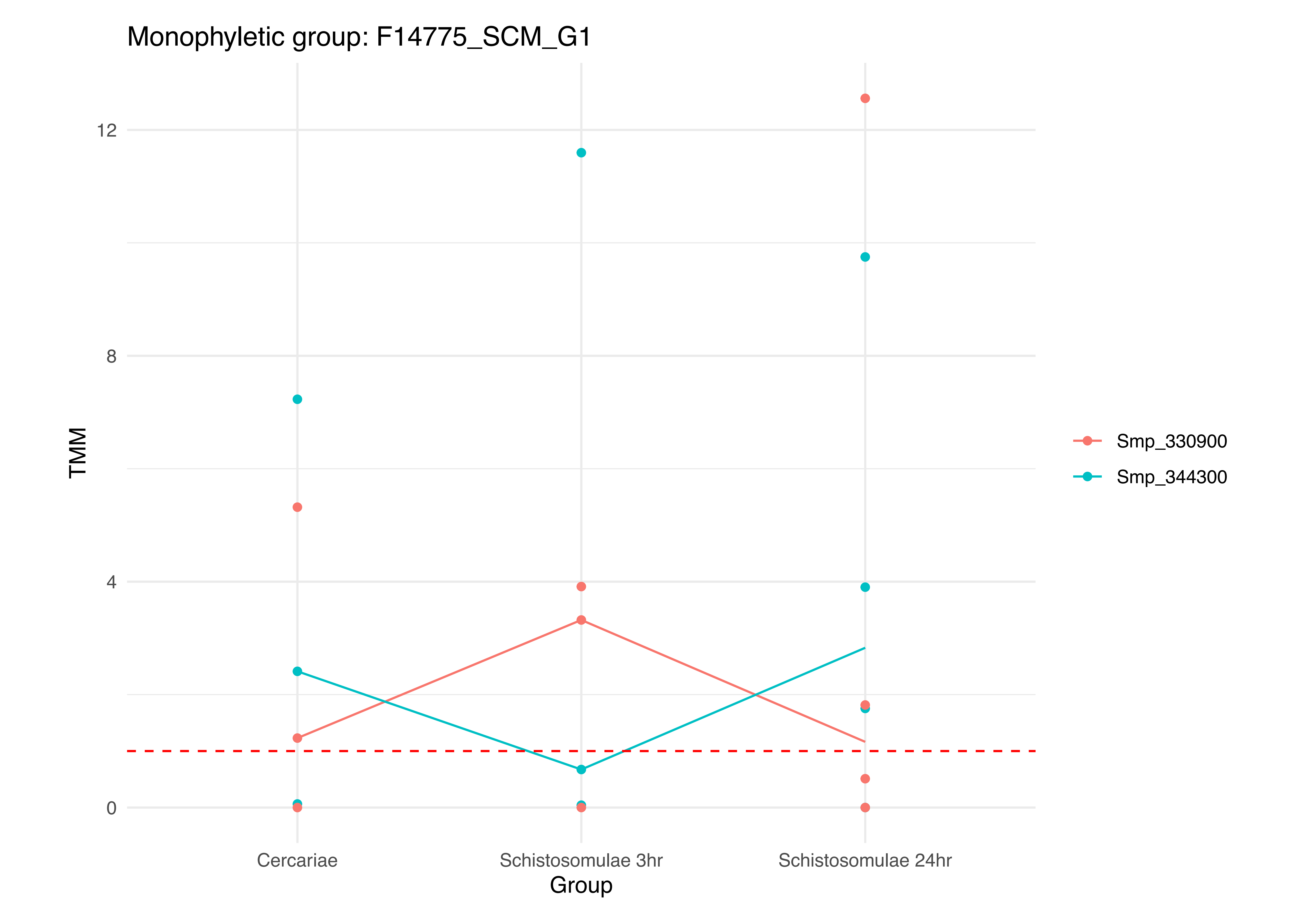
Monophyletic group: F14775_SCM_G1
12
8
4
0
TMM
Cercariae
Schistosomulae 3hr
Schistosomulae 24hr
Group
Smp_330900
Smp_344300

Monophyletic group: F14776_SCM_G1

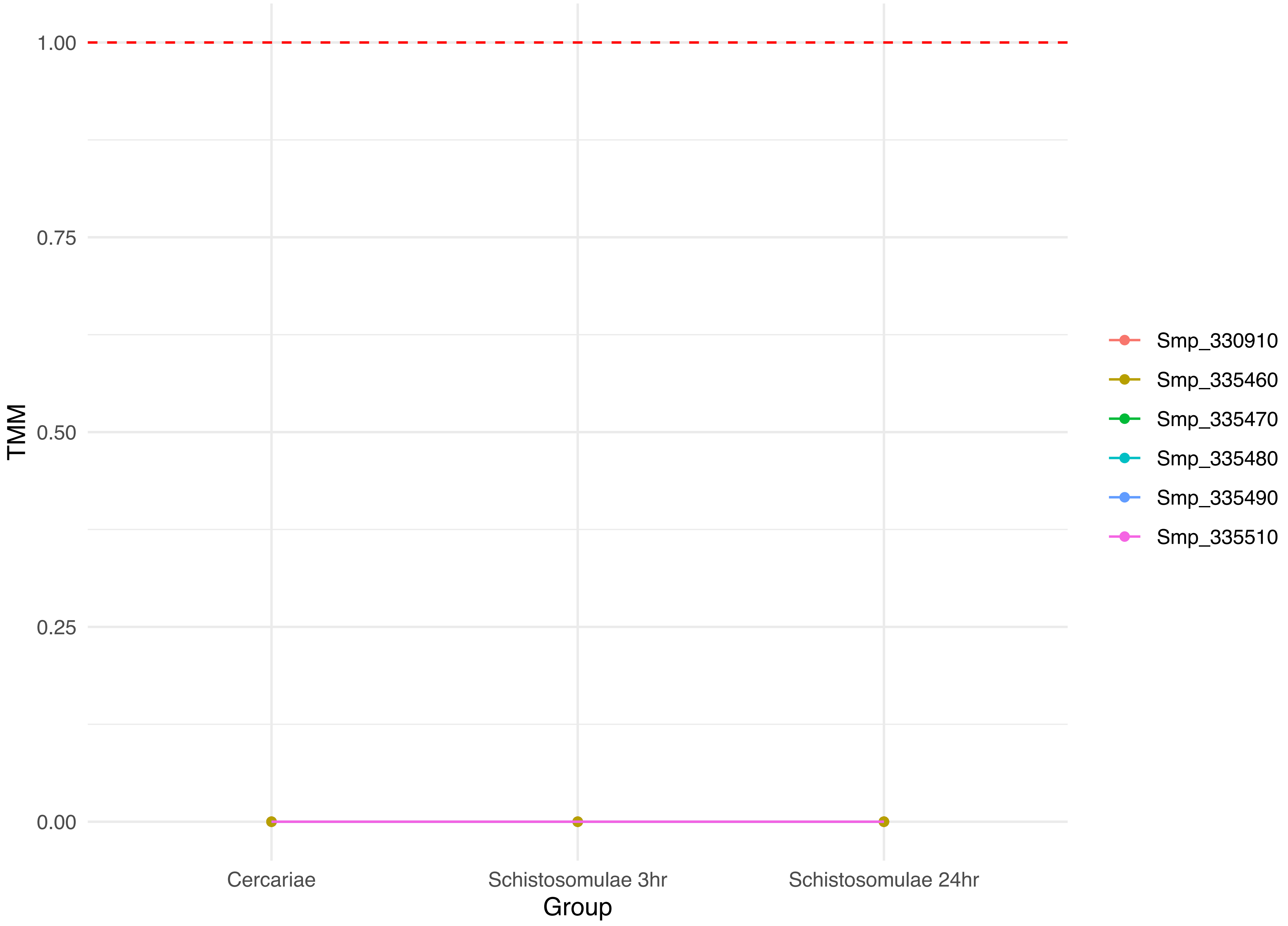

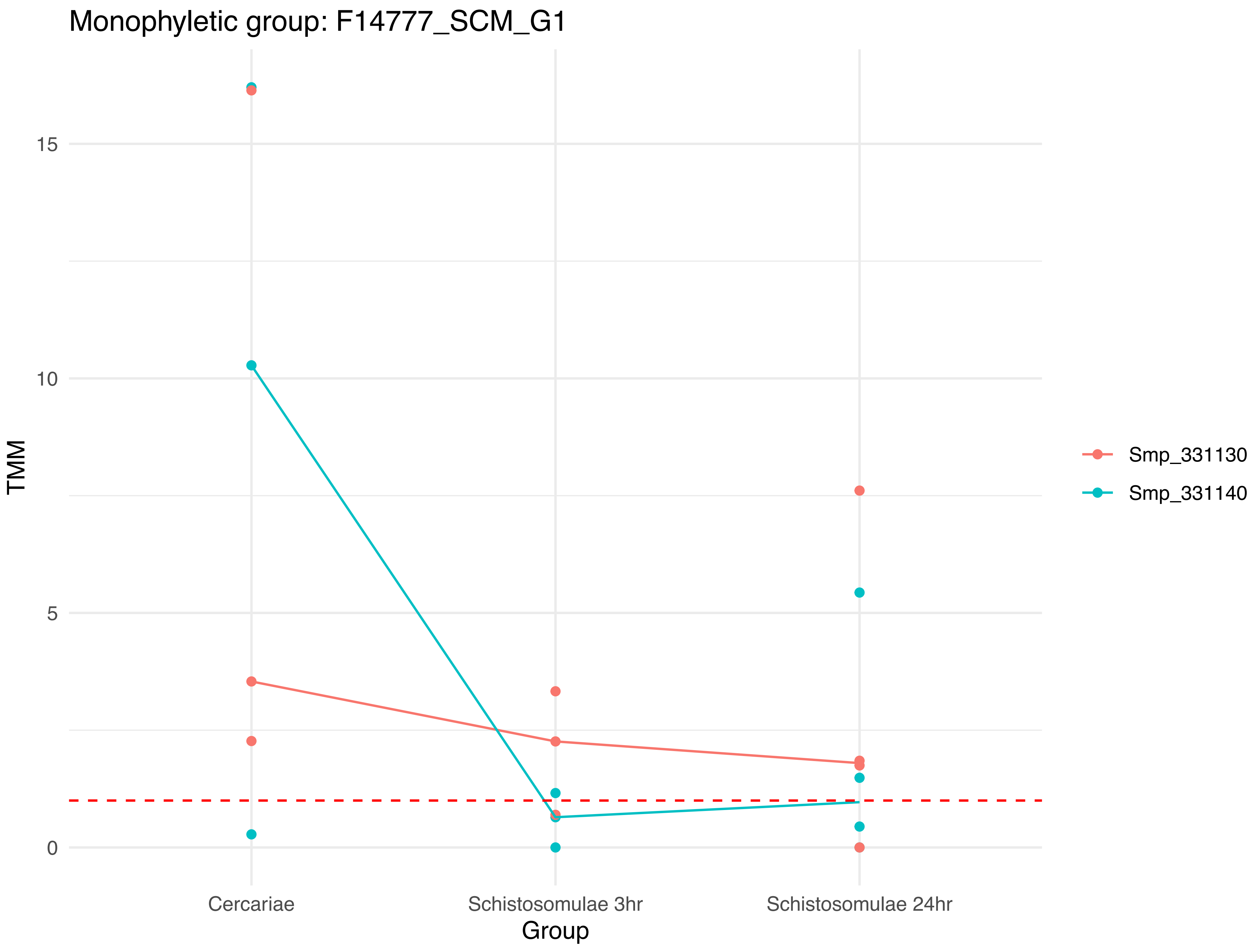
Monophyletic group: F14777_SCM_G1
TMM
15
10
5
0
Cercariae
Schistosomulae 3hr
Schistosomulae 24hr
Group
Smp_331130
Smp_331140

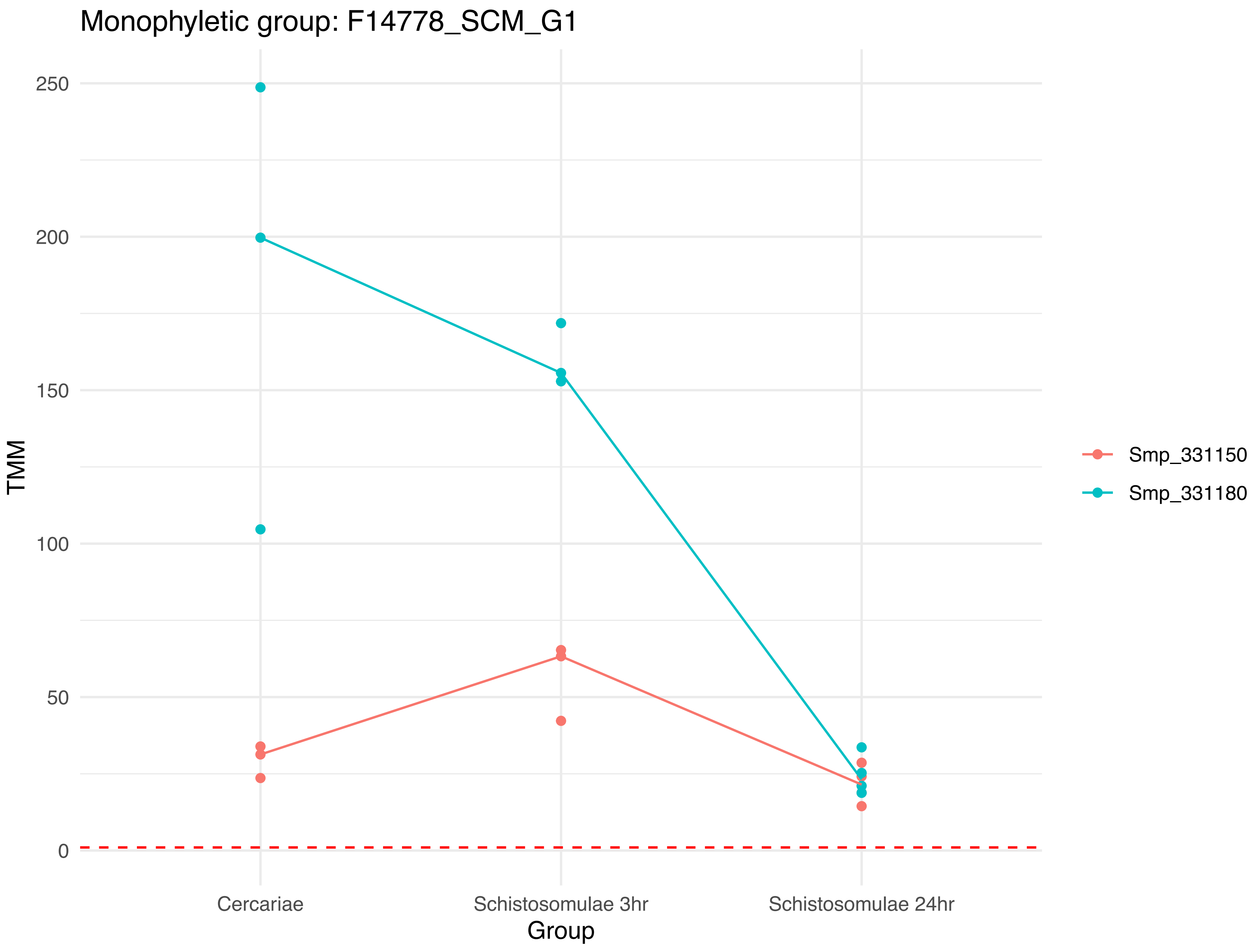
Monophyletic group: F14778_SCM_G1
250
200
150
100
50
0
TMM
Cercariae
Schistosomulae 3hr
Schistosomulae 24hr
Group
Smp_331150
Smp_331180

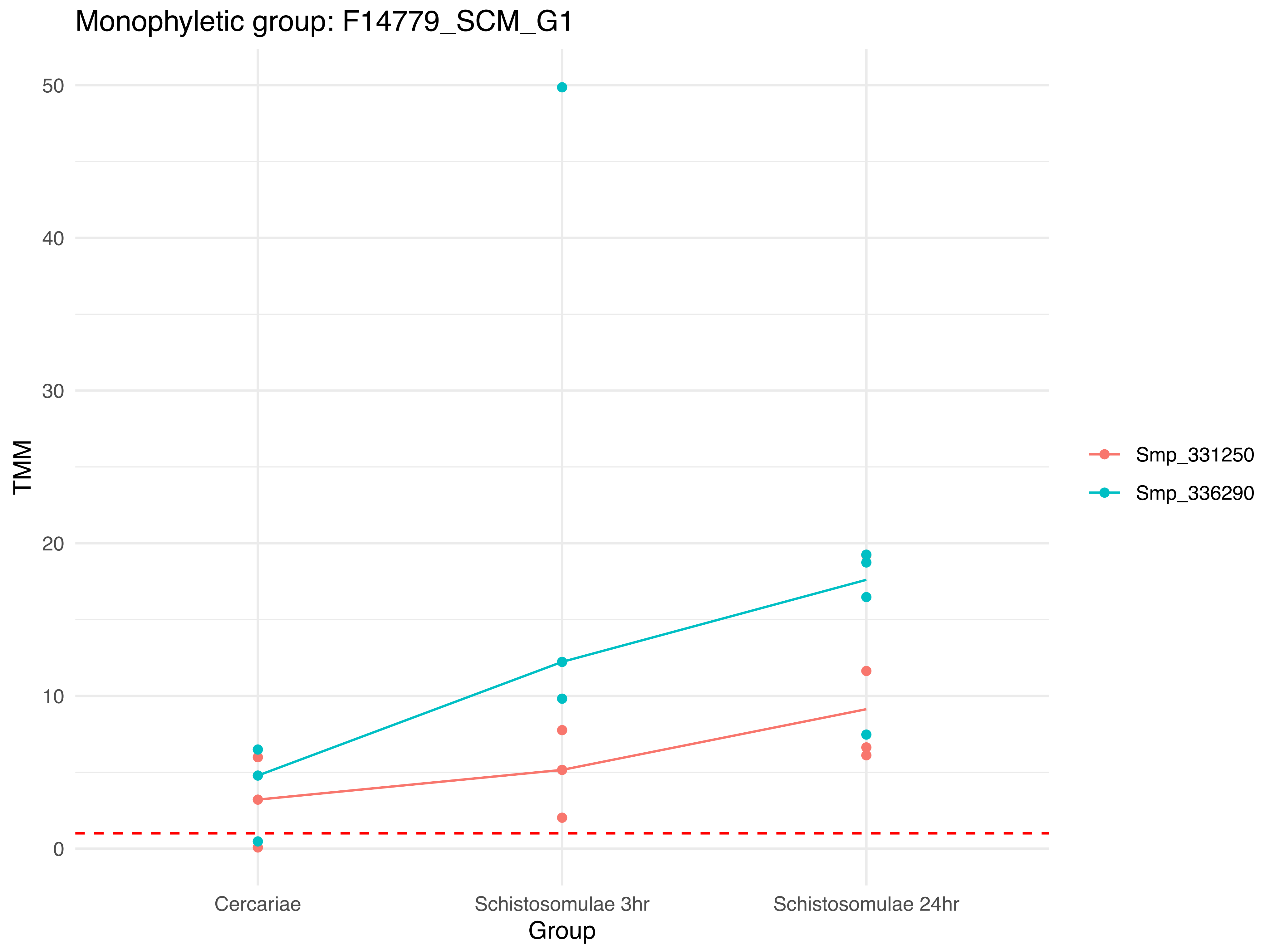
Monophyletic group: F14779_SCM_G1
TMM
50
40
30
20
10
0
Cercariae
Schistosomulae 3hr
Schistosomulae 24hr
Group
Smp_331250
Smp_336290

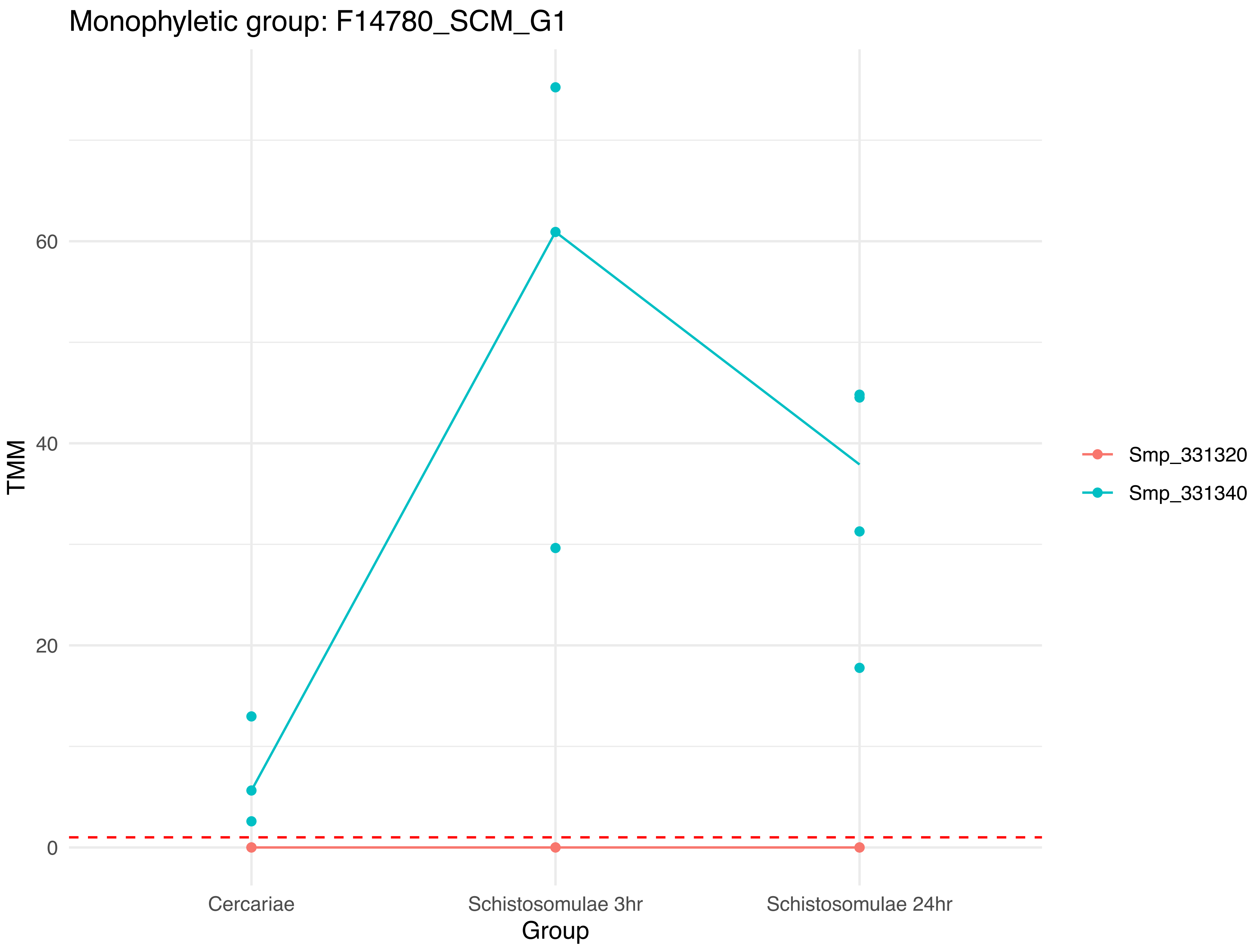

Monophyletic group: F14780_SCM_G1
TMM
60
40
20
0
Cercariae
Schistosomulae 3hr
Schistosomulae 24hr
Group
Smp_331320
Smp_331340

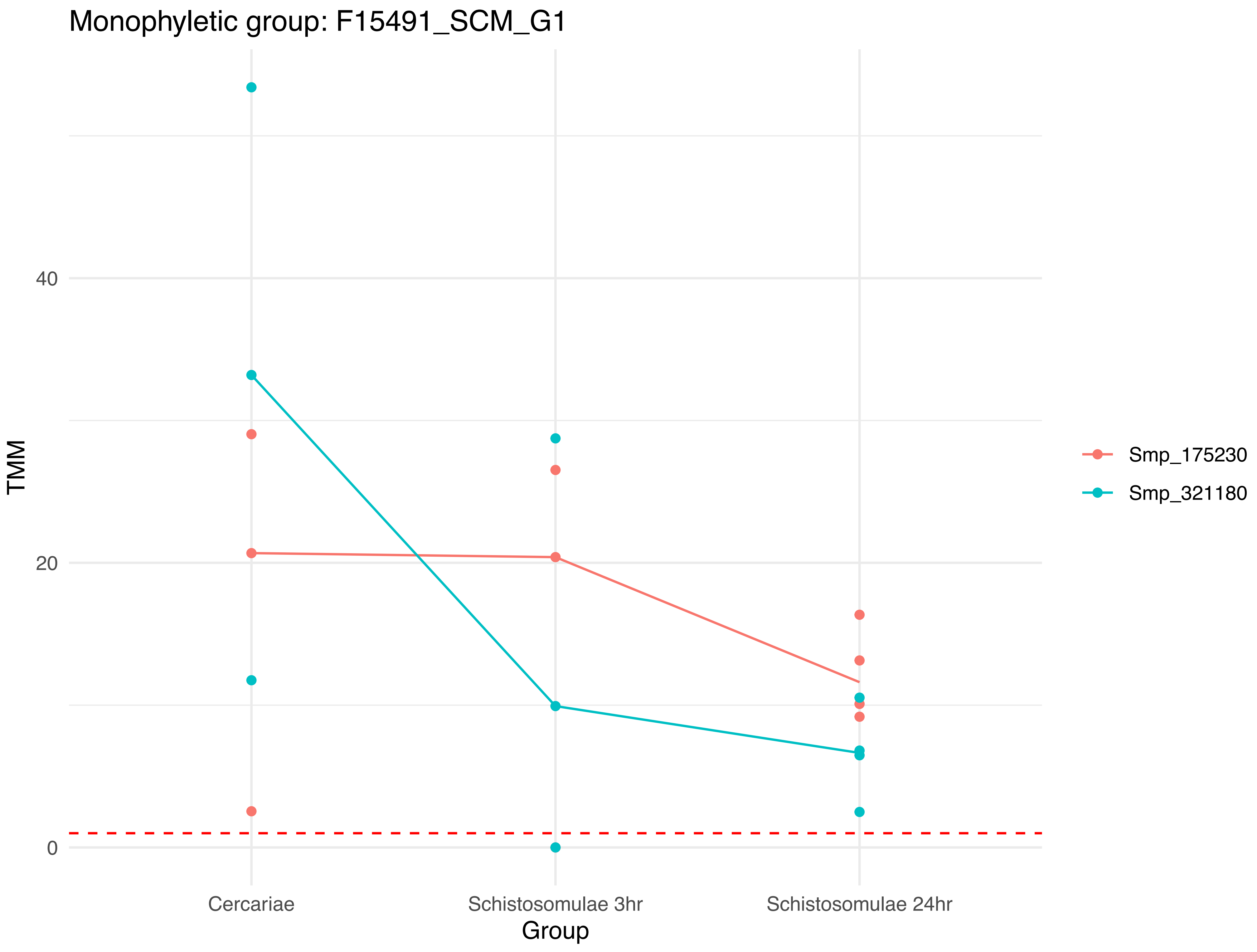

Monophyletic group: F15491_SCM_G1
TMM
40
20
0
Cercariae
Schistosomulae 3hr
Schistosomulae 24hr
Group
Smp_175230
Smp_321180

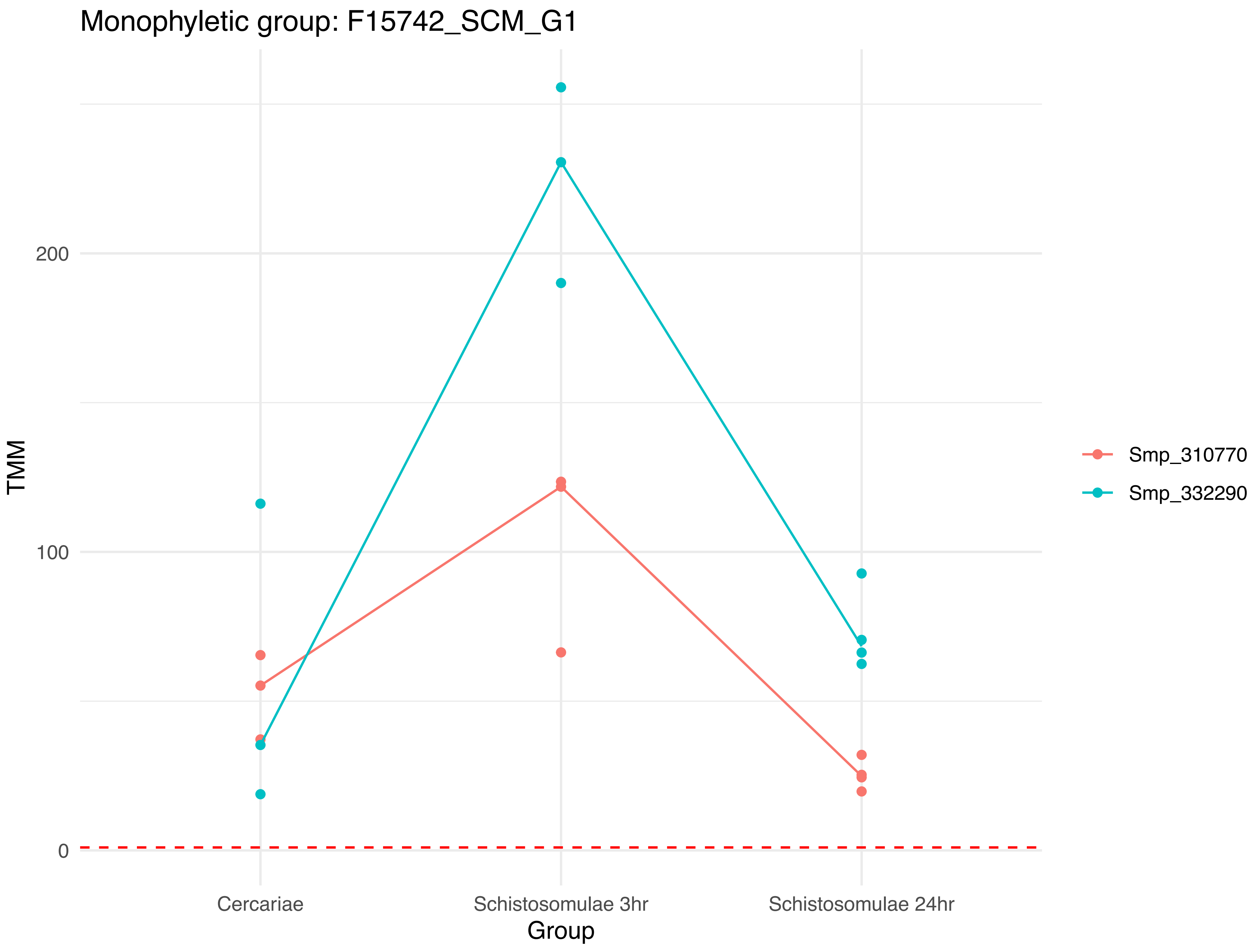

Monophyletic group: F15742_SCM_G1
TMM
200
100
0
Cercariae
Schistosomulae 3hr
Schistosomulae 24hr
Group
Smp_310770
Smp_332290

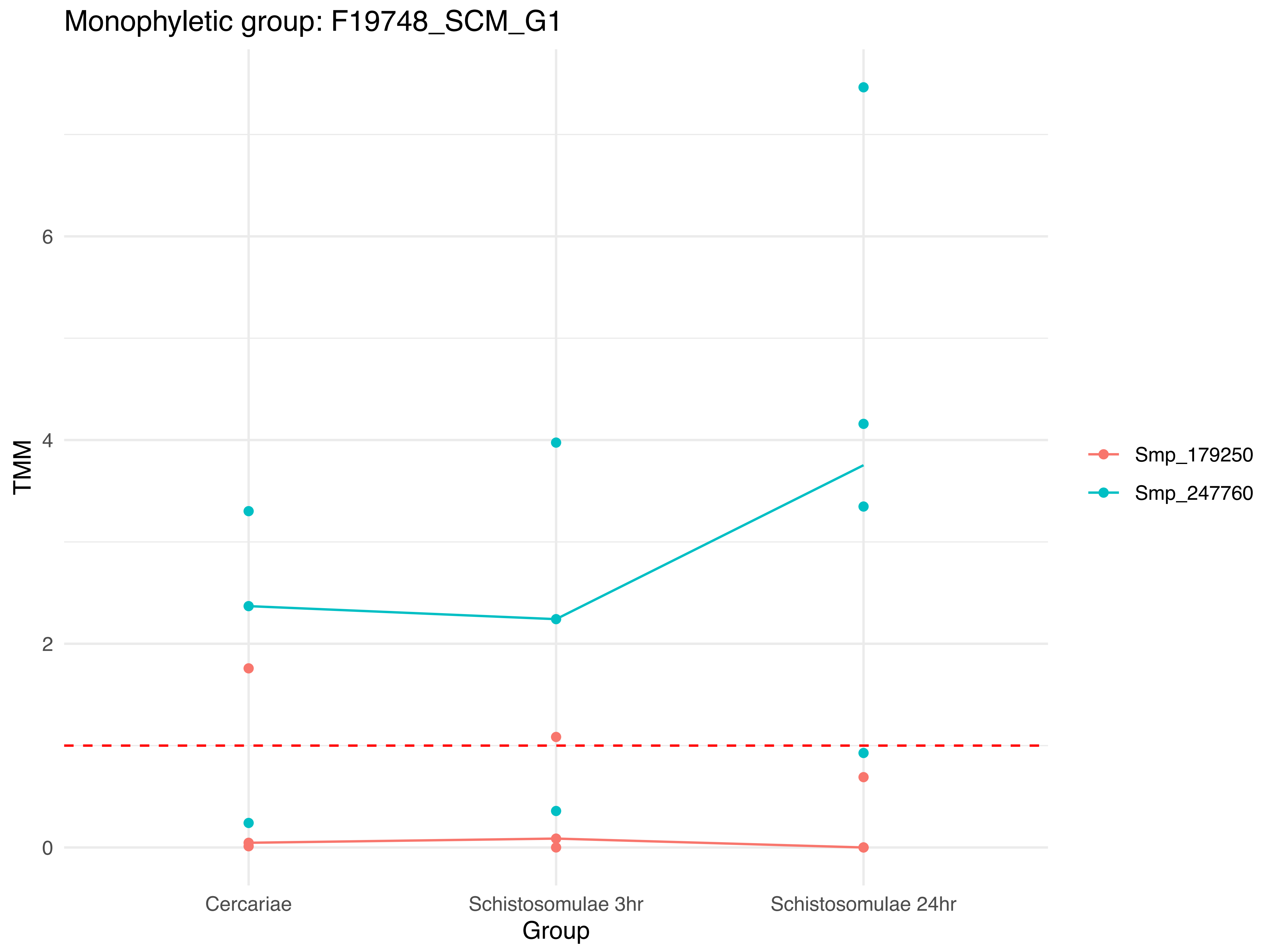

Monophyletic group: F19748_SCM_G1
TMM
6
4
2
0
Cercariae
Schistosomulae 3hr
Schistosomulae 24hr
Group
Smp_179250
Smp_247760

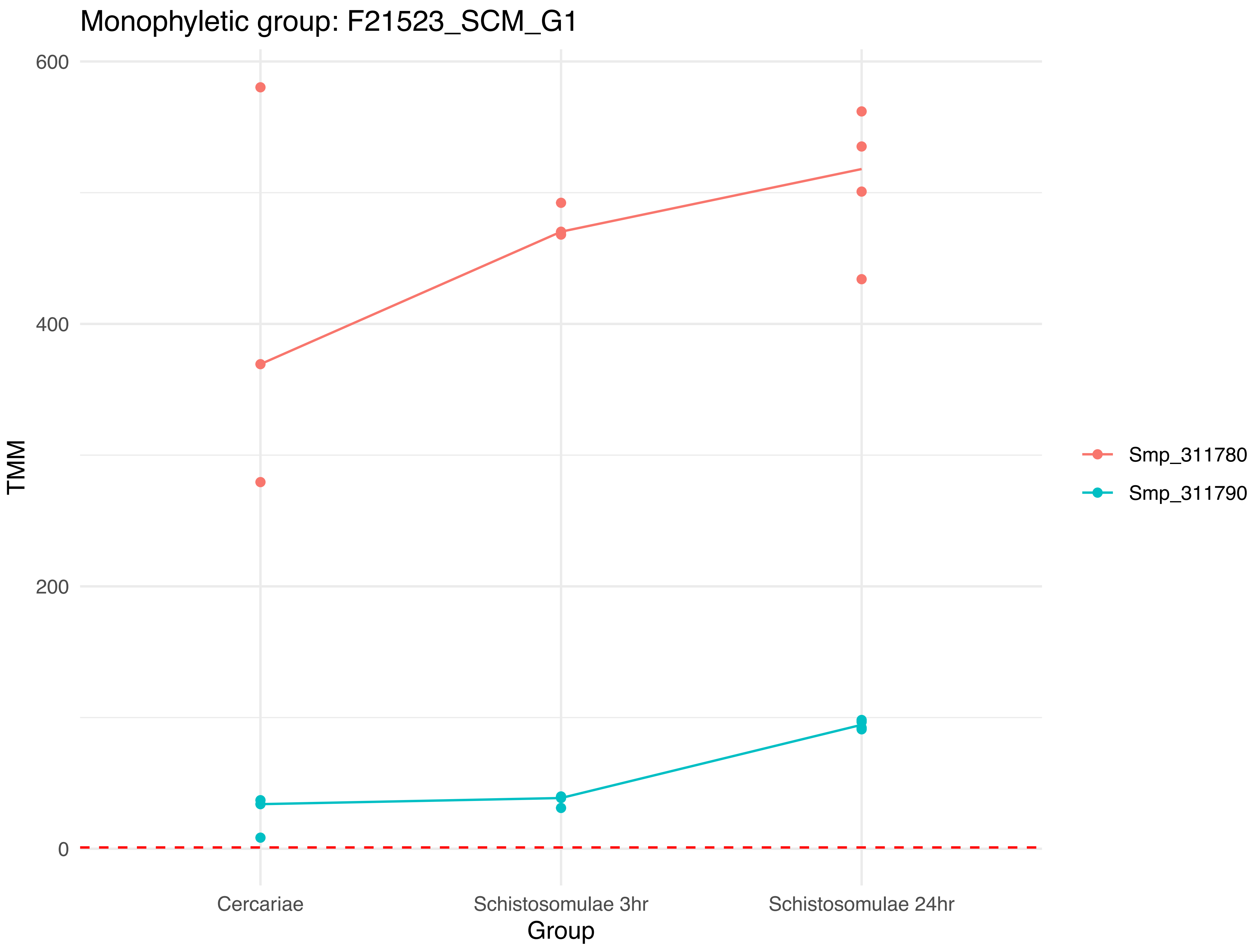
Monophyletic group: F21523_SCM_G1
600
400
200
0
TMM
Cercariae
Schistosomulae 3hr
Schistosomulae 24hr
Group
Smp_311780
Smp_311790

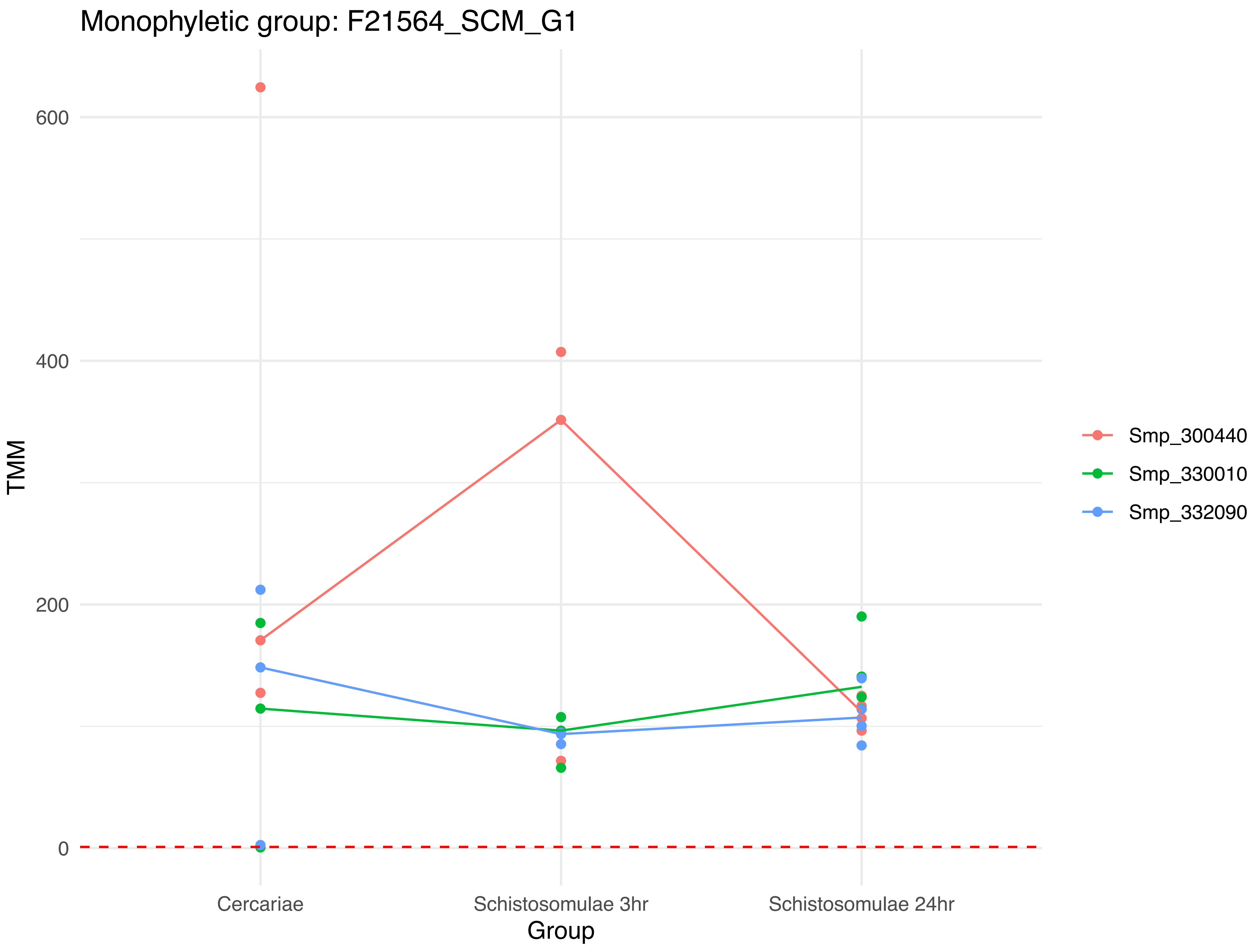

Monophyletic group: F21564_SCM_G1
600
400
200
0
TMM
Cercariae
Schistosomulae 3hr
Schistosomulae 24hr
Group
Smp_300440
Smp_330010
Smp_332090

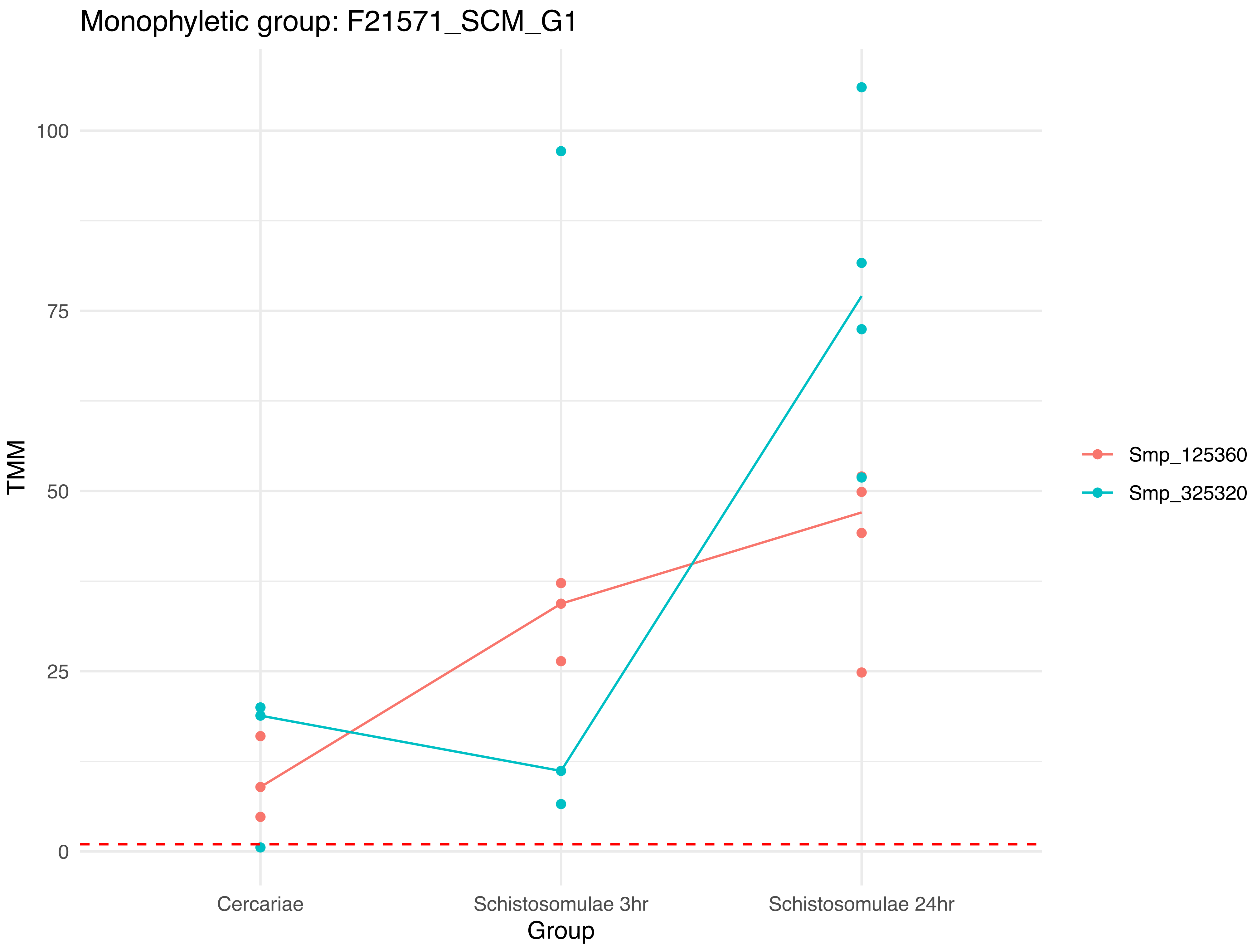

Monophyletic group: F21571_SCM_G1
TMM
100
75
50
25
0
Cercariae
Schistosomulae 3hr
Schistosomulae 24hr
Group
Smp_125360
Smp_325320

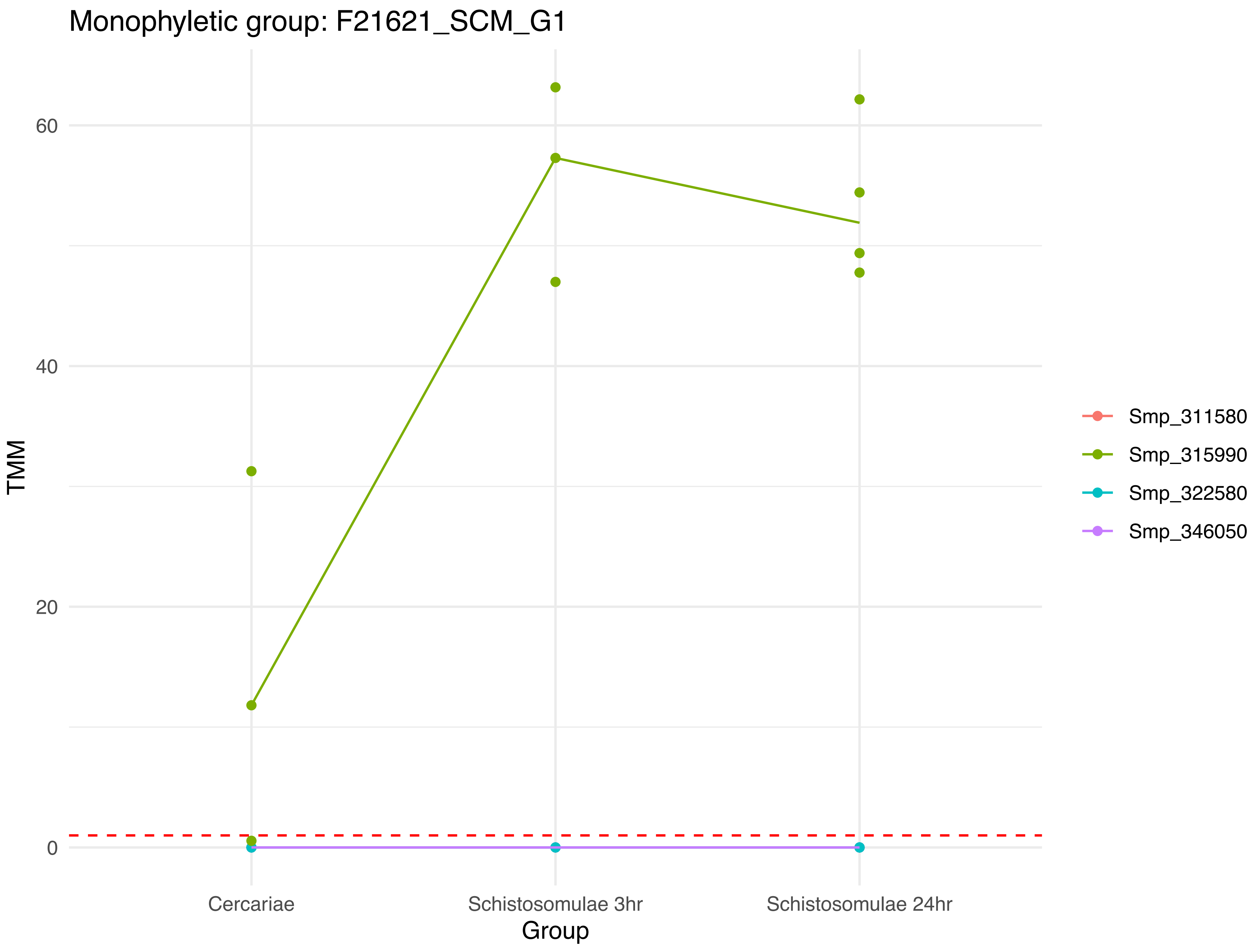

Monophyletic group: F21621_SCM_G1
TMM
60
40
20
0
Cercariae
Schistosomulae 3hr
Schistosomulae 24hr
Group
Smp_311580
Smp_315990
Smp_322580
Smp_346050

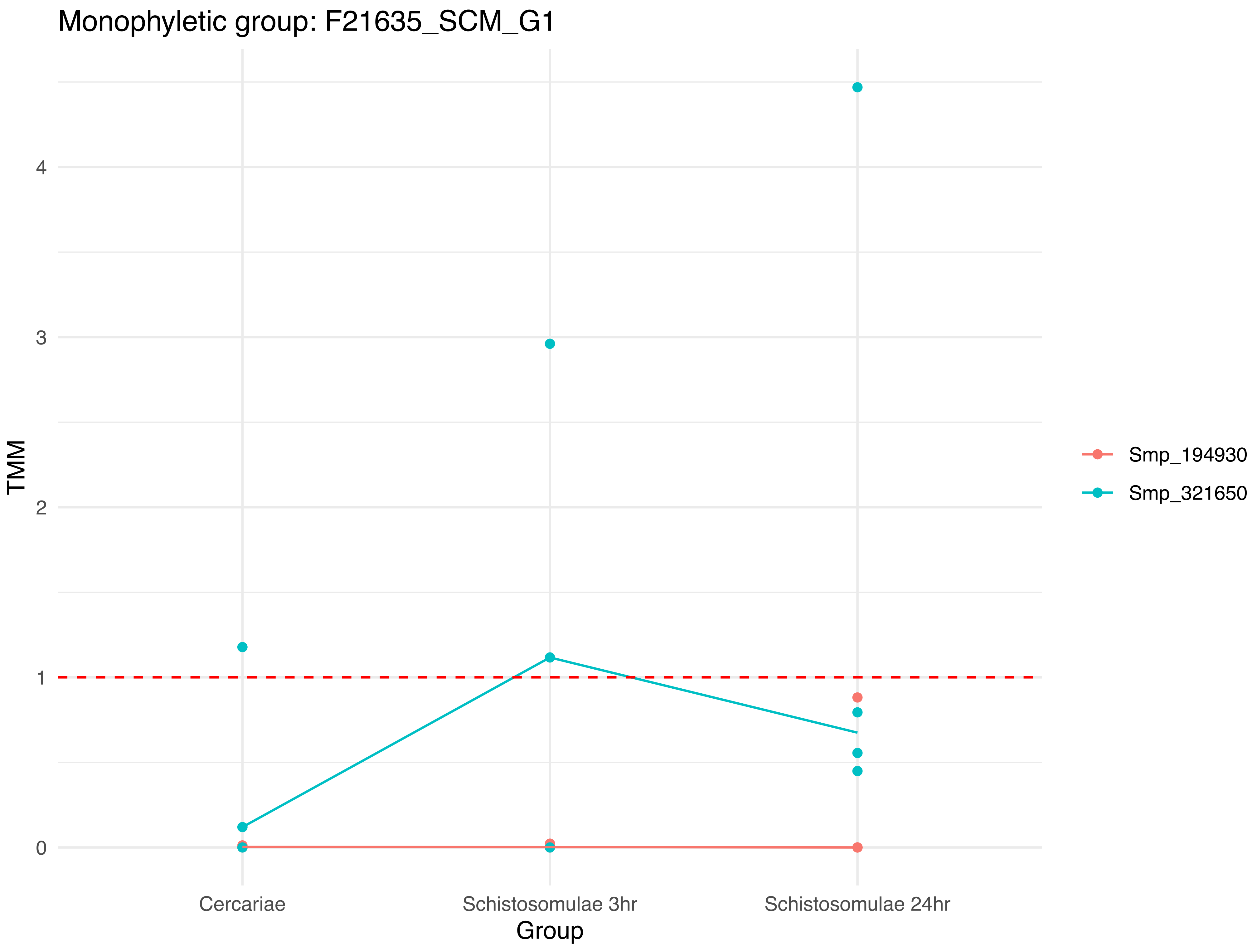
Monophyletic group: F21635_SCM_G1
TMM
0
1
2
3
4
Cercariae
Schistosomulae 3hr
Schistosomulae 24hr
Group
Smp_194930
Smp_321650

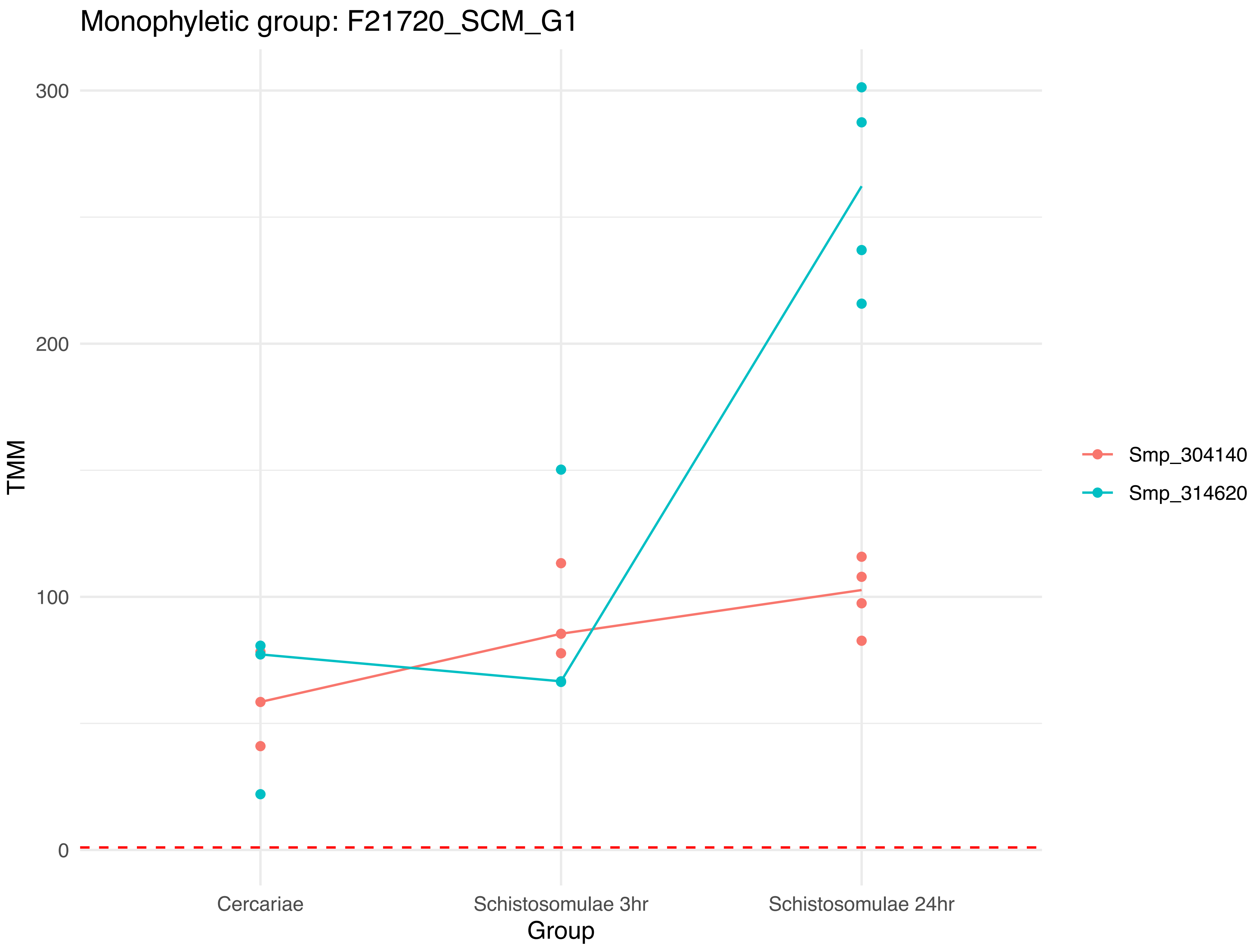
Monophyletic group: F21720_SCM_G1
300
200
100
0
TMM
Cercariae
Schistosomulae 3hr
Schistosomulae 24hr
Group
Smp_304140
Smp_314620

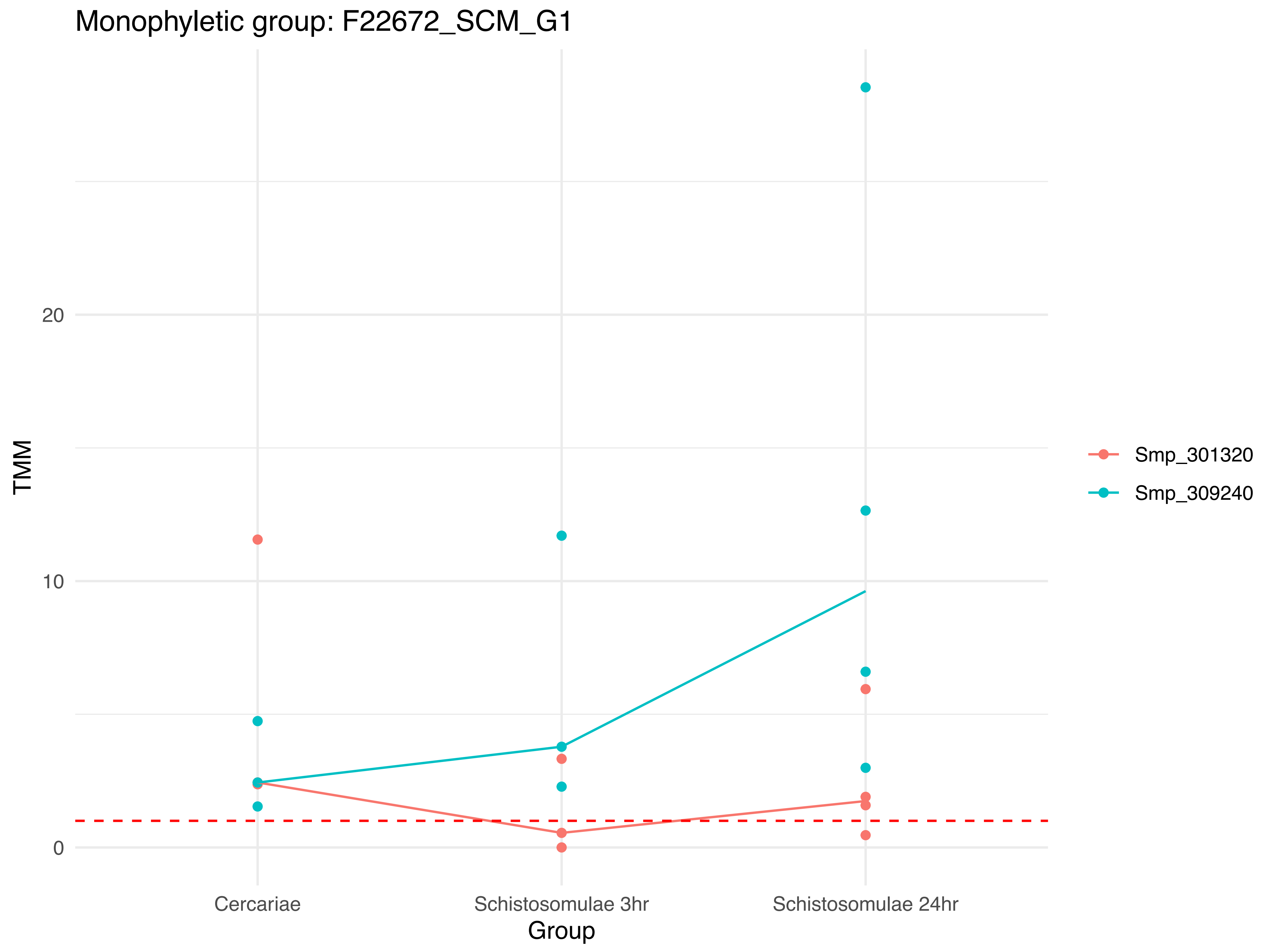

Monophyletic group: F22672_SCM_G1
TMM
20
10
0
Cercariae
Schistosomulae 3hr
Schistosomulae 24hr
Group
Smp_301320
Smp_309240

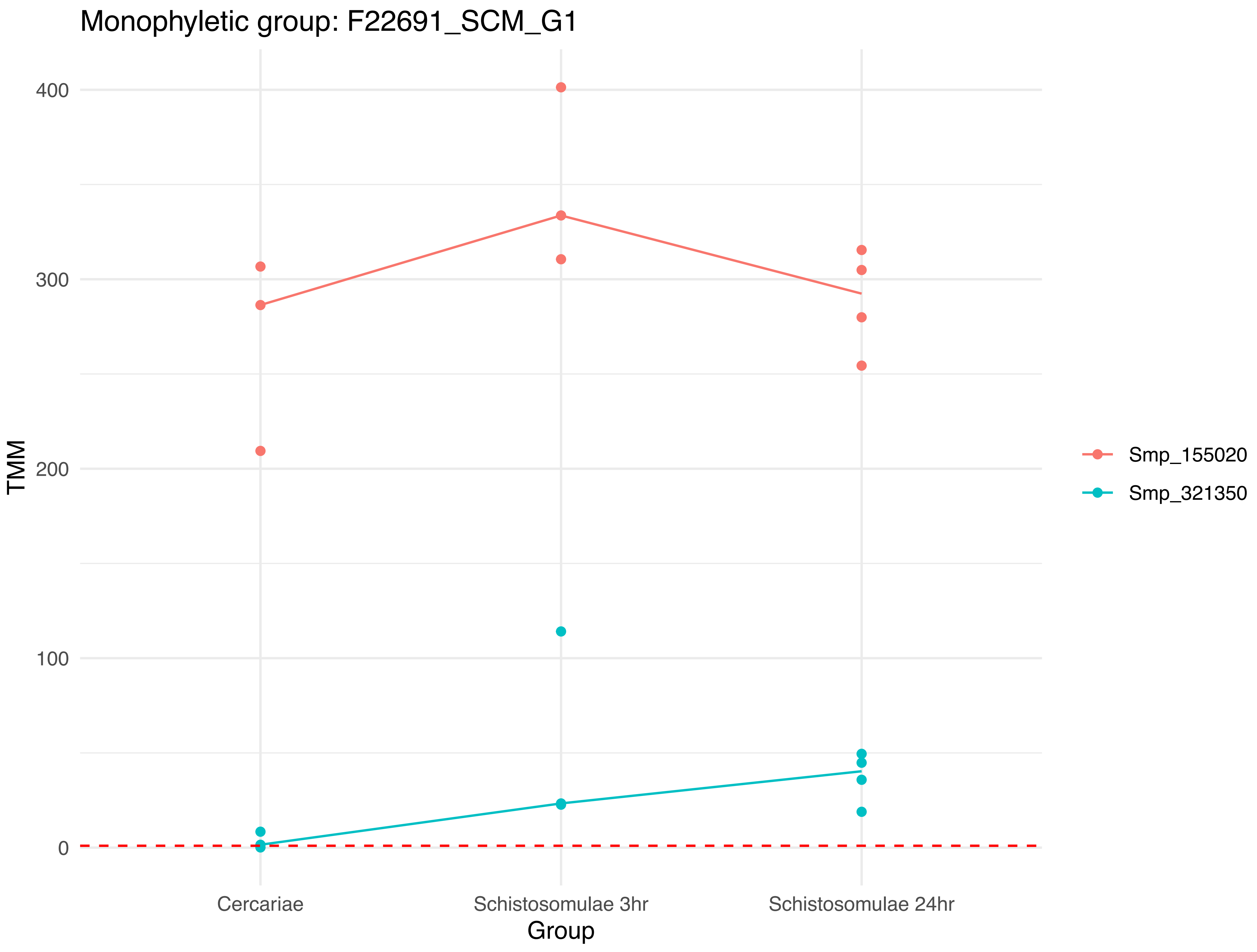

Monophyletic group: F22691_SCM_G1
400
300
200
100
0
TMM
Cercariae
Schistosomulae 3hr
Schistosomulae 24hr
Group
Smp_155020
Smp_321350

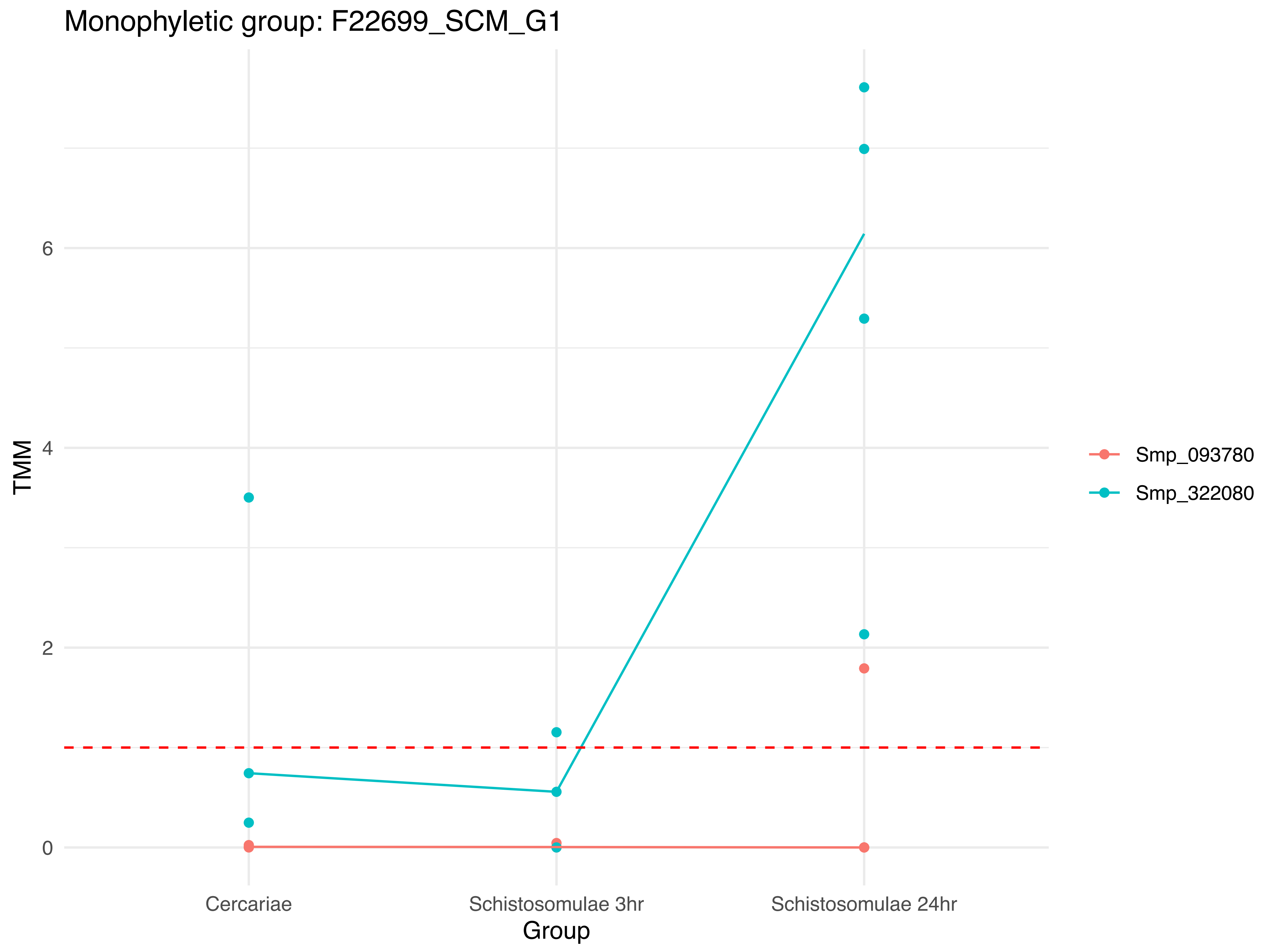
Monophyletic group: F22699_SCM_G1
TMM
6
4
2
0
Cercariae
Schistosomulae 3hr
Schistosomulae 24hr
Group
Smp_093780
Smp_322080

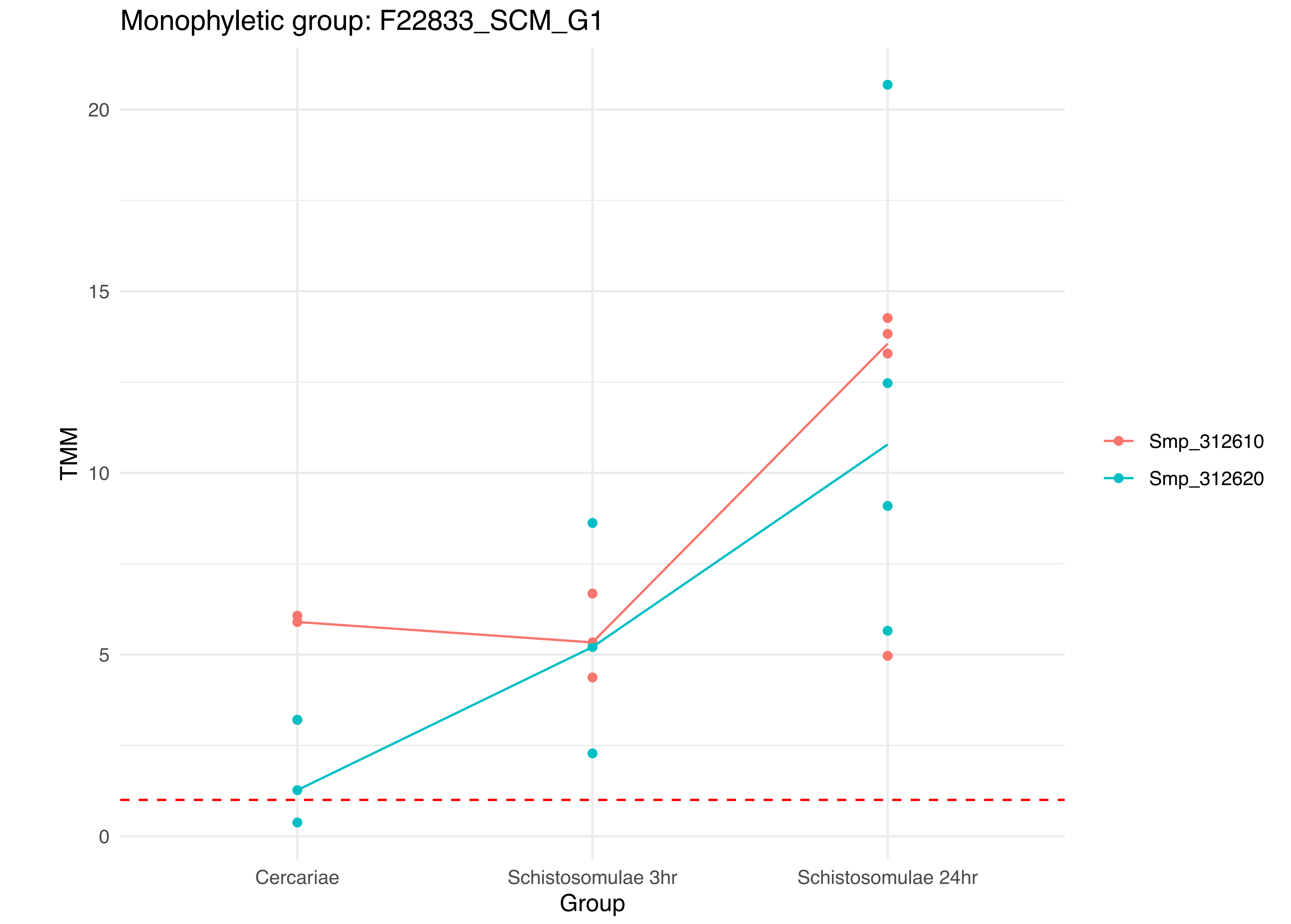

Monophyletic group: F22833_SCM_G1
20
15
10
5
0
TMM
Cercariae
Schistosomulae 3hr
Schistosomulae 24hr
Group
Smp_312610
Smp_312620

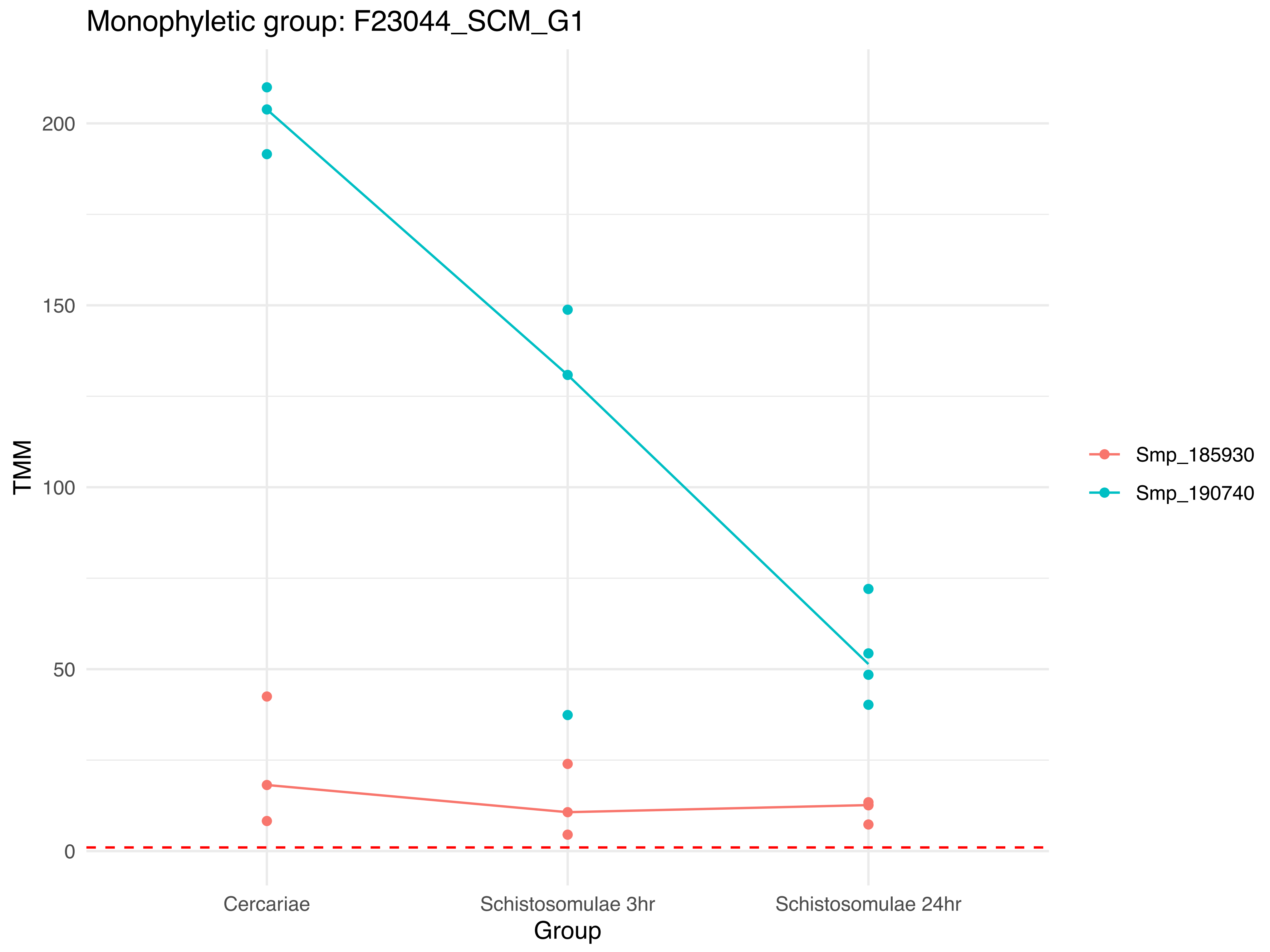
Monophyletic group: F23044_SCM_G1
TMM
200
150
100
50
0
Cercariae
Schistosomulae 3hr
Schistosomulae 24hr
Group
Smp_185930
Smp_190740

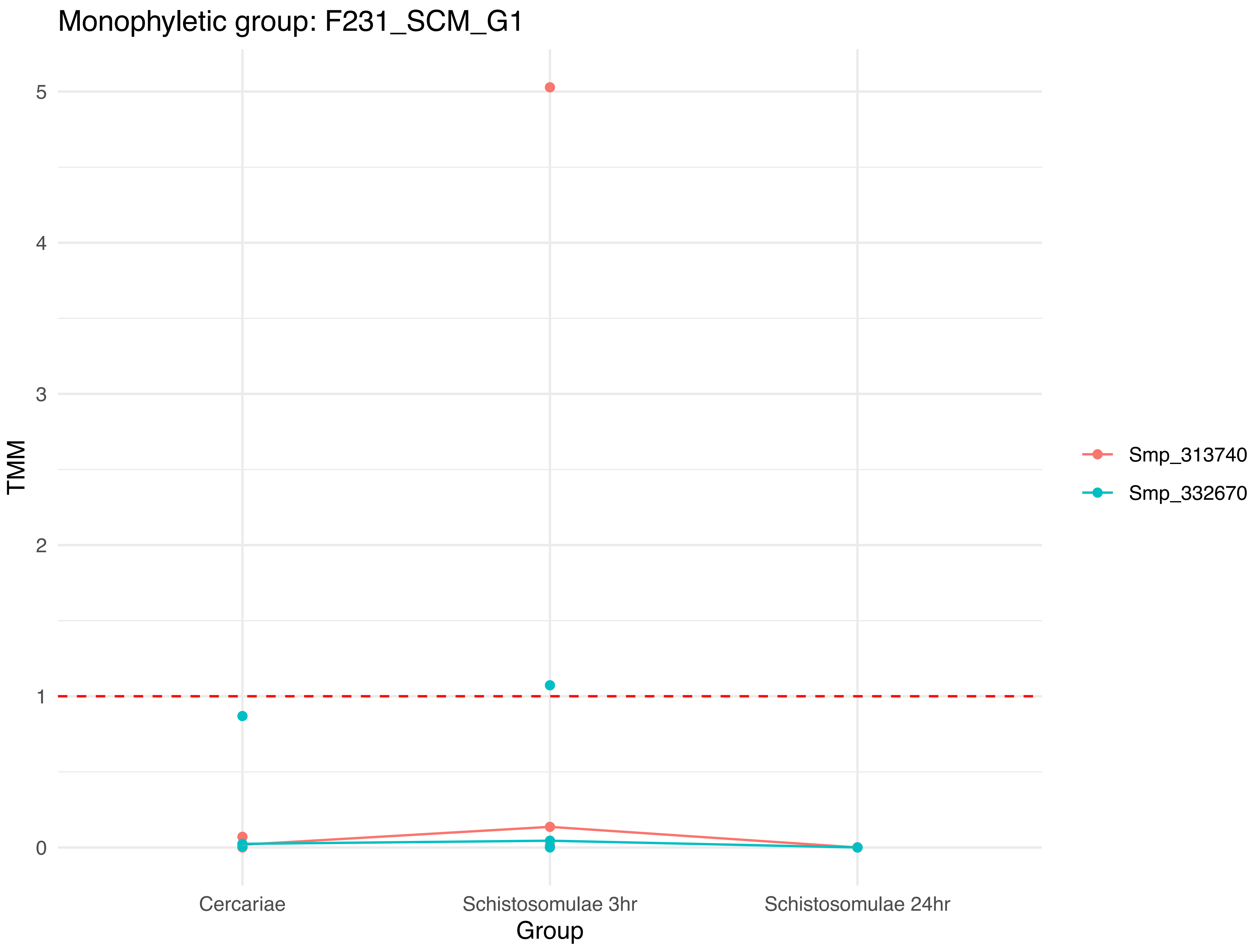

Monophyletic group: F231_SCM_G1
TMM
5
4
3
2
1
0
Cercariae
Schistosomulae 3hr
Schistosomulae 24hr
Group
Smp_313740
Smp_332670

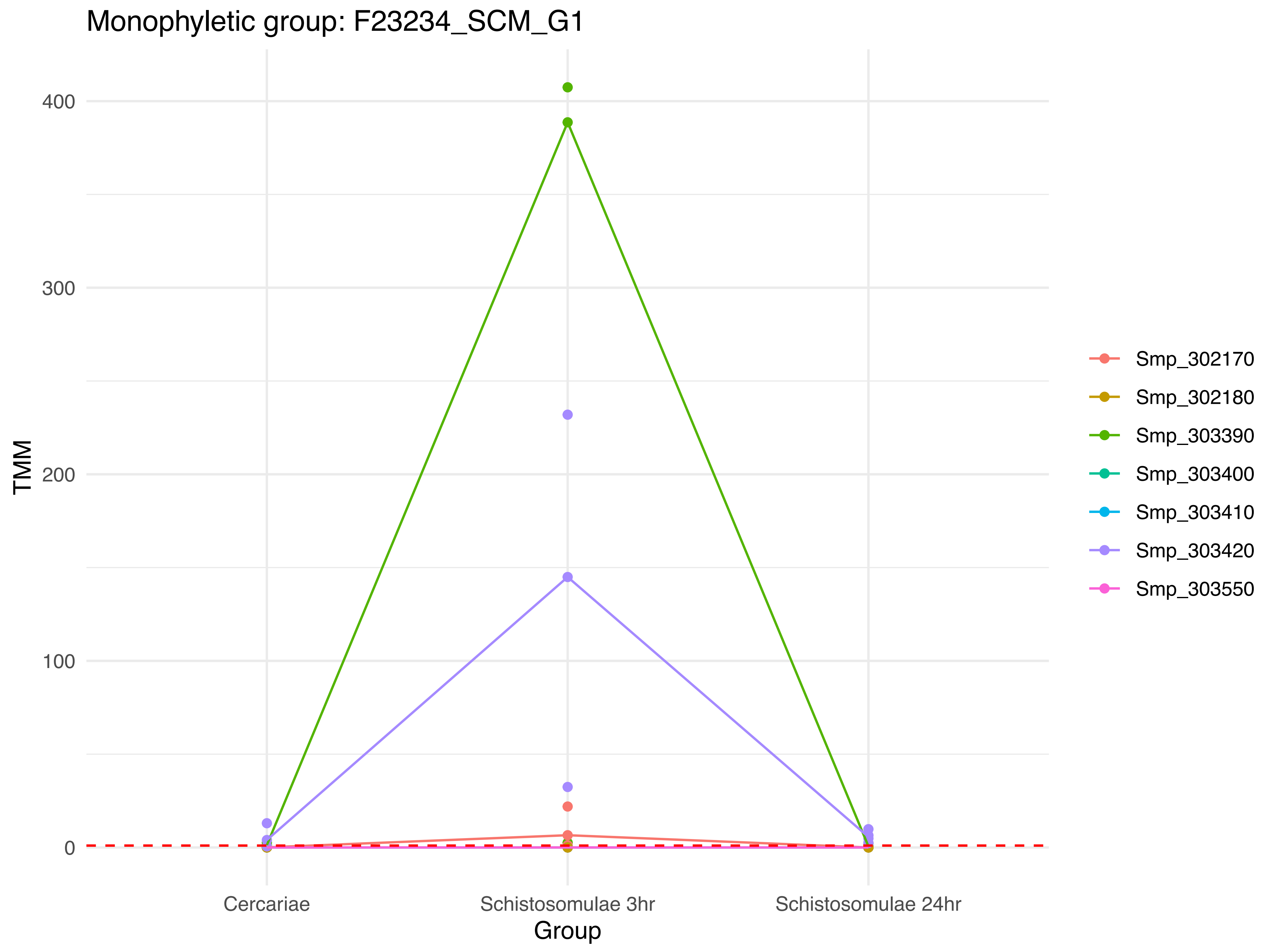

Monophyletic group: F23234_SCM_G1
400
300
200
100
0
TMM
Cercariae
Schistosomulae 3hr
Schistosomulae 24hr
Group
Smp_302170
Smp_302180
Smp_303390
Smp_303400
Smp_303410
Smp_303420
Smp_303550

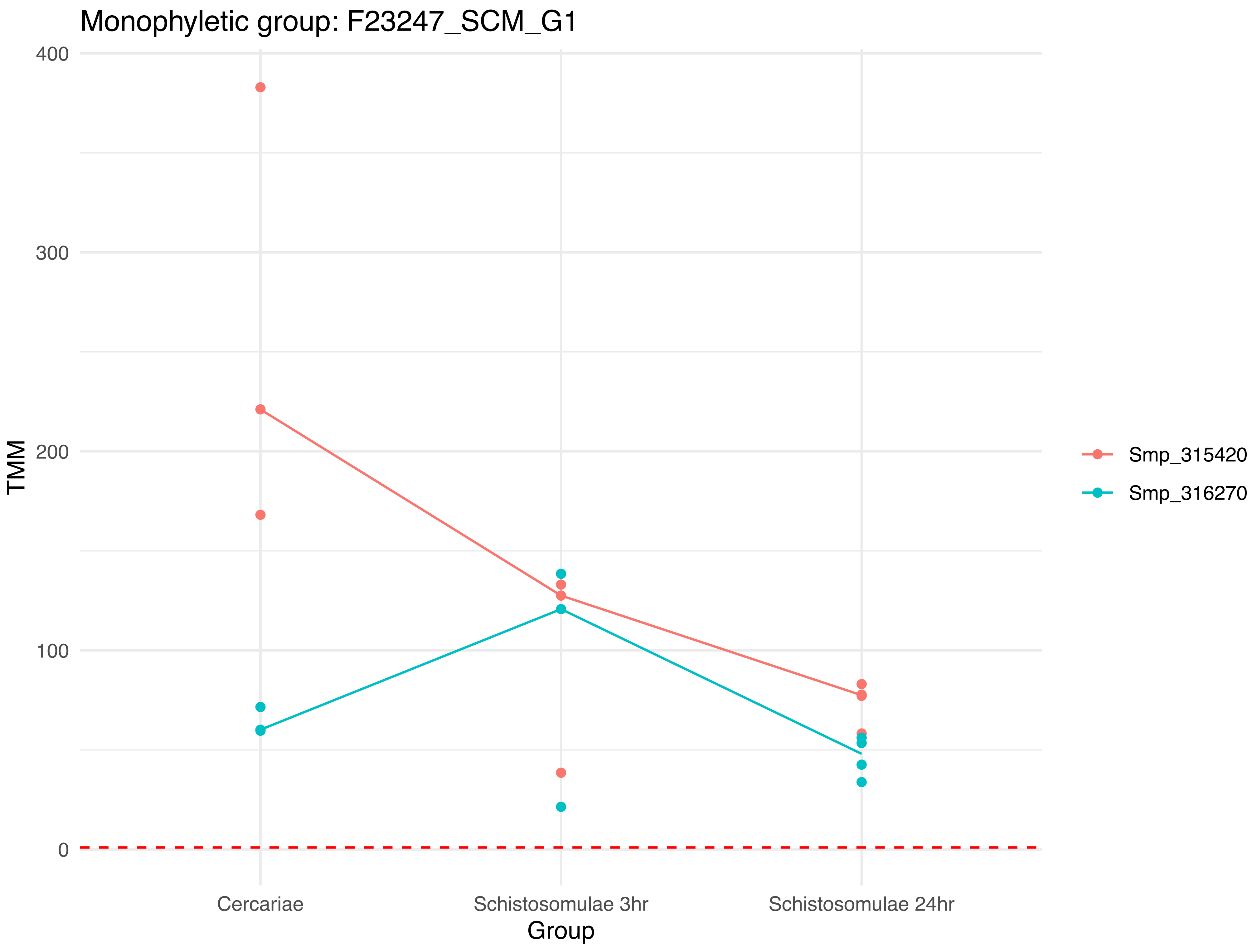
Monophyletic group: F23247_SCM_G1
TMM
400
300
200
100
0
Cercariae
Schistosomulae 3hr
Schistosomulae 24hr
Group
Smp_315420
Smp_316270

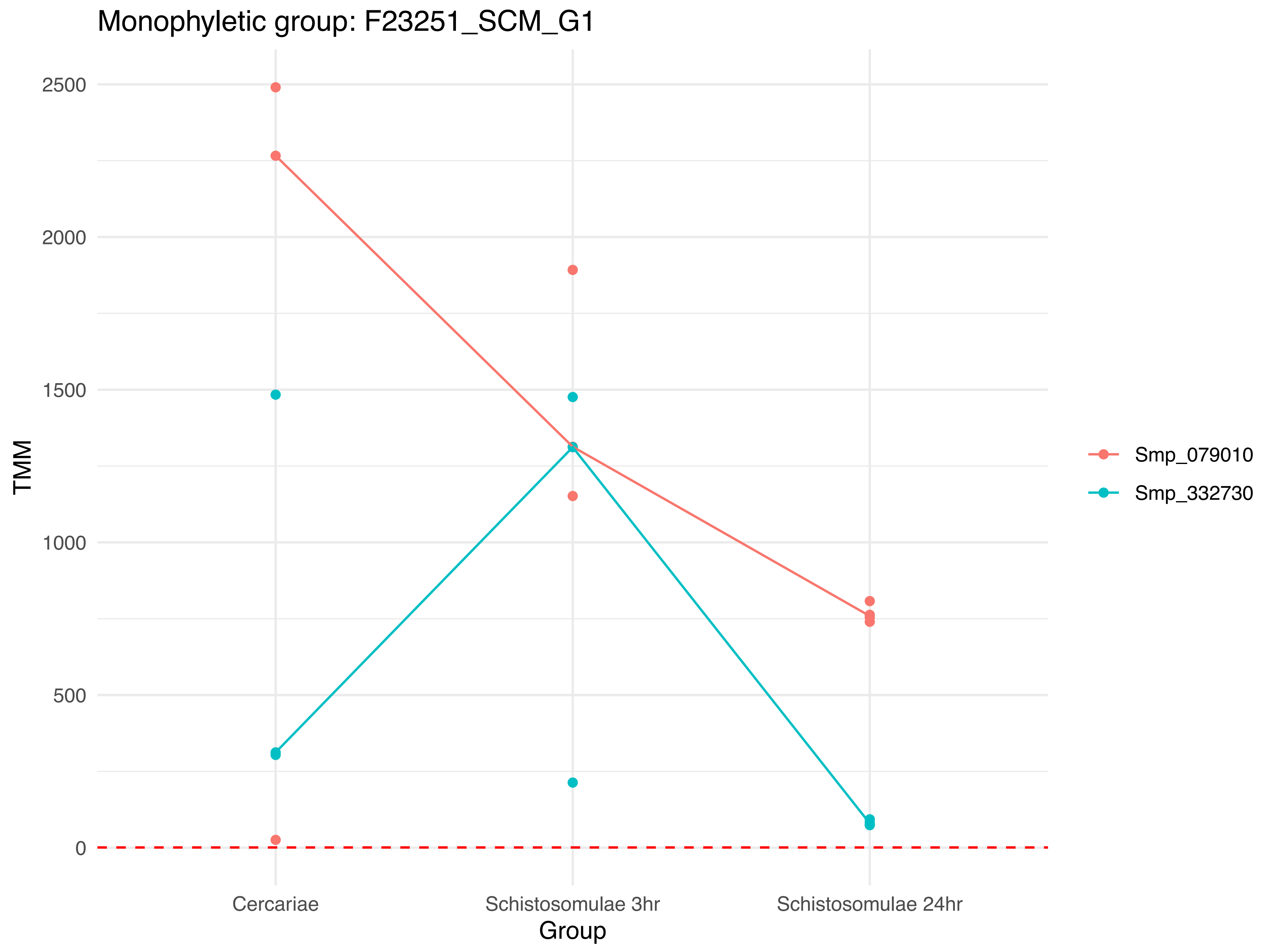

Monophyletic group: F23251_SCM_G1
2500
2000
1500
1000
500
0
TMM
Cercariae
Schistosomulae 3hr
Schistosomulae 24hr
Group
Smp_079010
Smp_332730

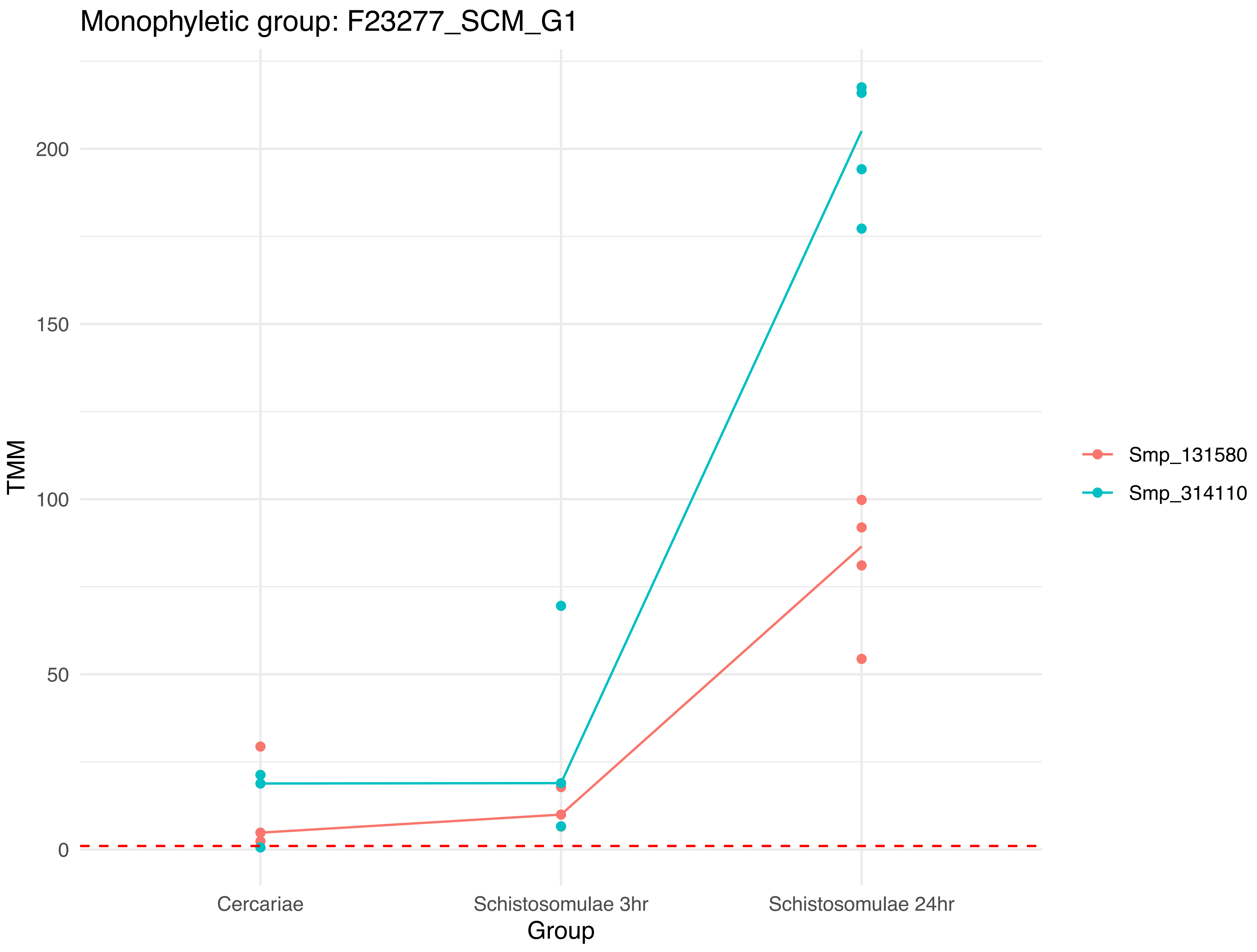

Monophyletic group: F23277_SCM_G1
TMM
200
150
100
50
0
Cercariae
Schistosomulae 3hr
Schistosomulae 24hr
Group
Smp_131580
Smp_314110

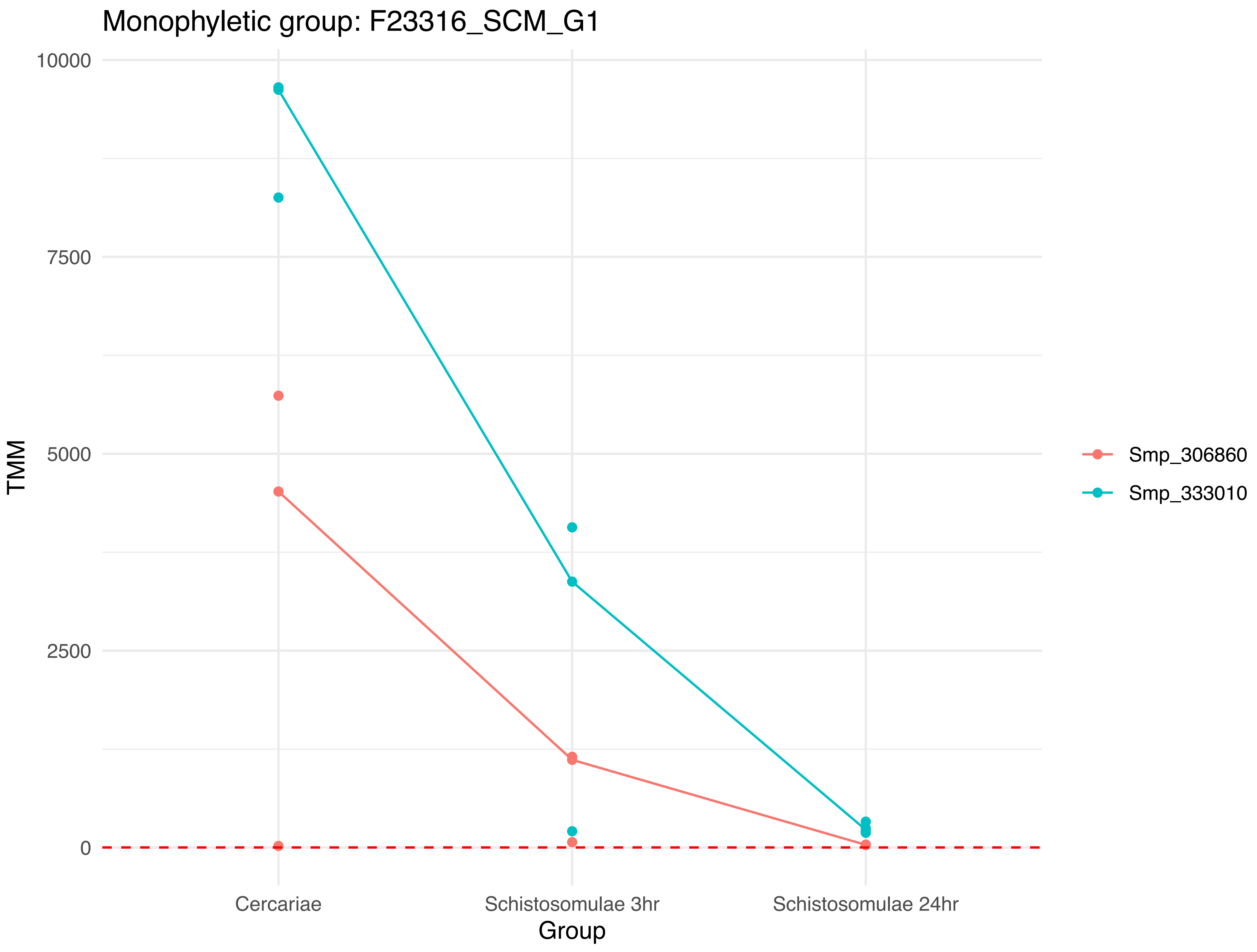

Monophyletic group: F23316_SCM_G1
10000
7500
5000
2500
0
TMM
Cercariae
Schistosomulae 3hr
Schistosomulae 24hr
Group
Smp_306860
Smp_333010

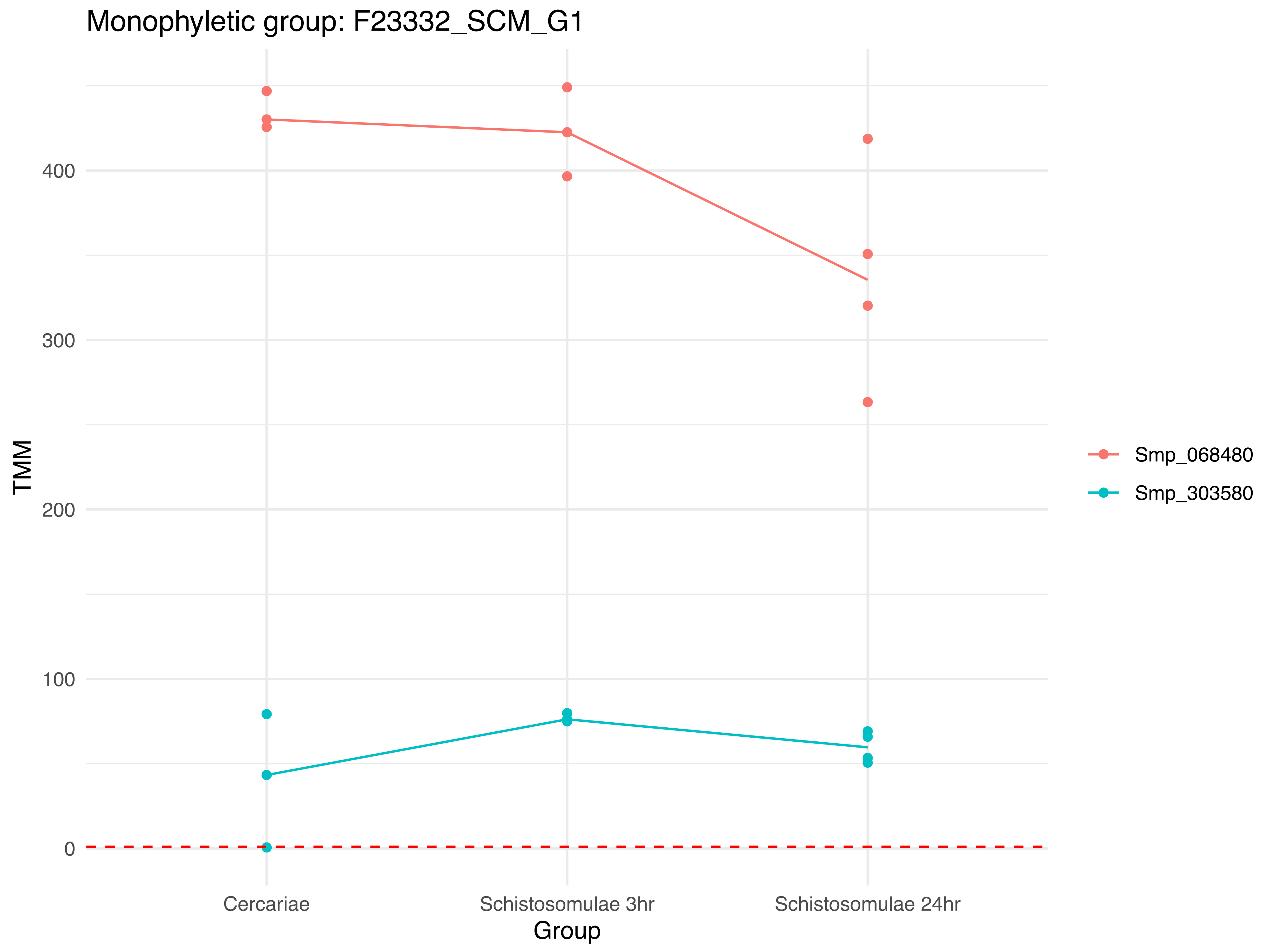

Monophyletic group: F23332_SCM_G1
TMM
400
300
200
100
0
Cercariae
Schistosomulae 3hr
Schistosomulae 24hr
Group
Smp_068480
Smp_303580

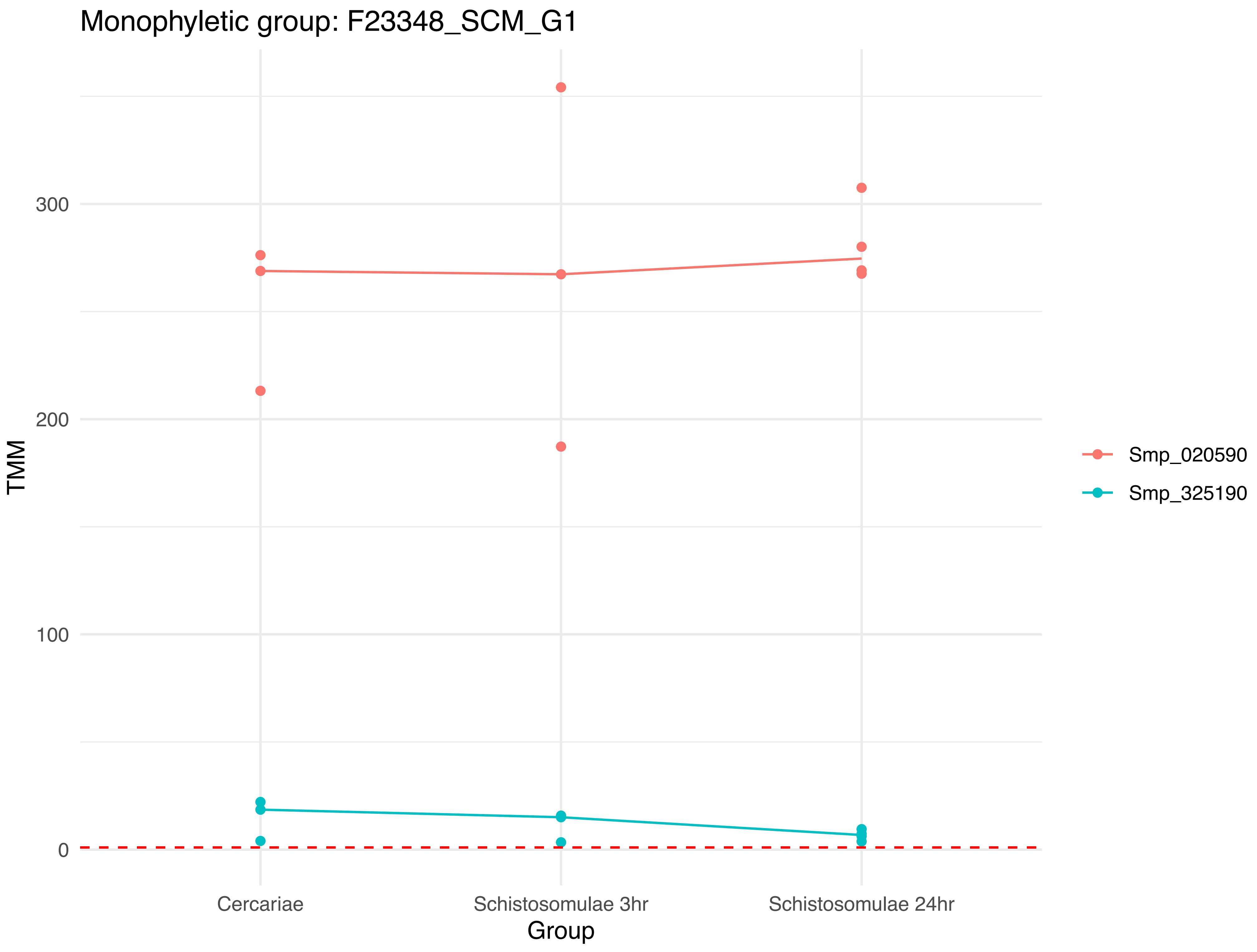

Monophyletic group: F23348_SCM_G1
TMM
300
200
100
0
Cercariae
Schistosomulae 3hr
Schistosomulae 24hr
Group
Smp_020590
Smp_325190

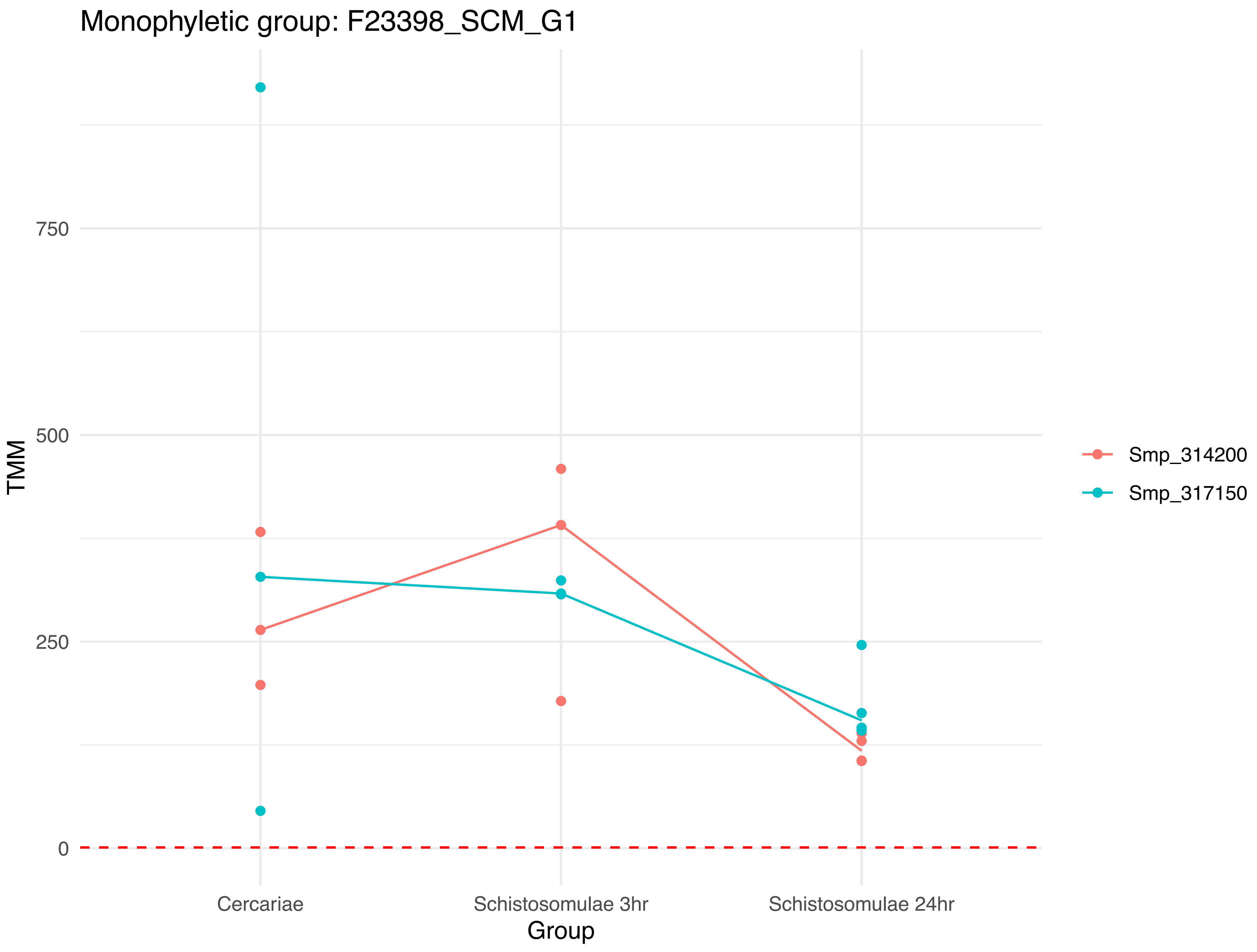

Monophyletic group: F23398_SCM_G1
TMM
750
500
250
0
Cercariae
Schistosomulae 3hr
Schistosomulae 24hr
Group
Smp_314200
Smp_317150

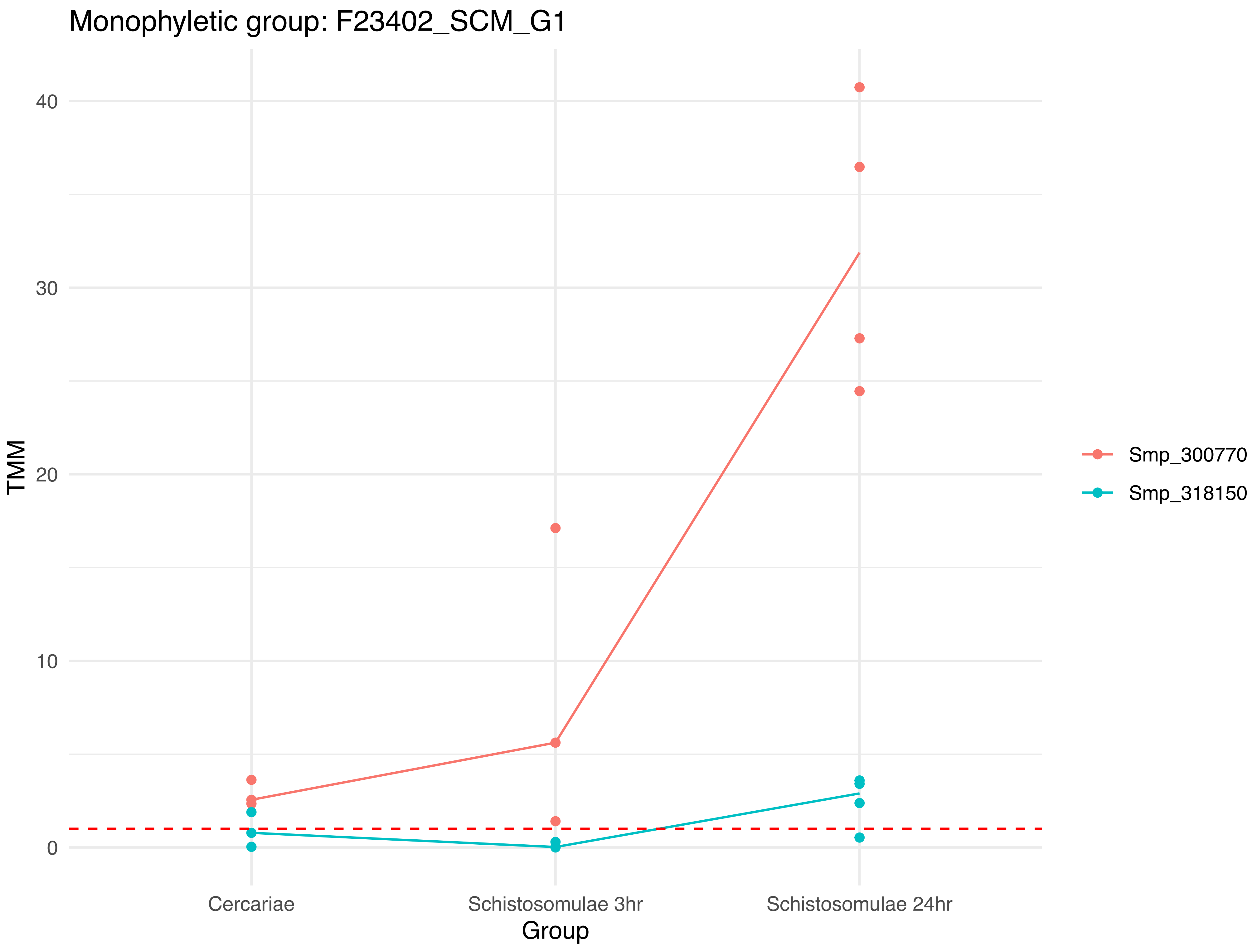

Monophyletic group: F23402_SCM_G1
TMM
40
30
20
10
0
Cercariae
Schistosomulae 3hr
Schistosomulae 24hr
Group
Smp_300770
Smp_318150

Monophyletic group: F23436_SCM_G1

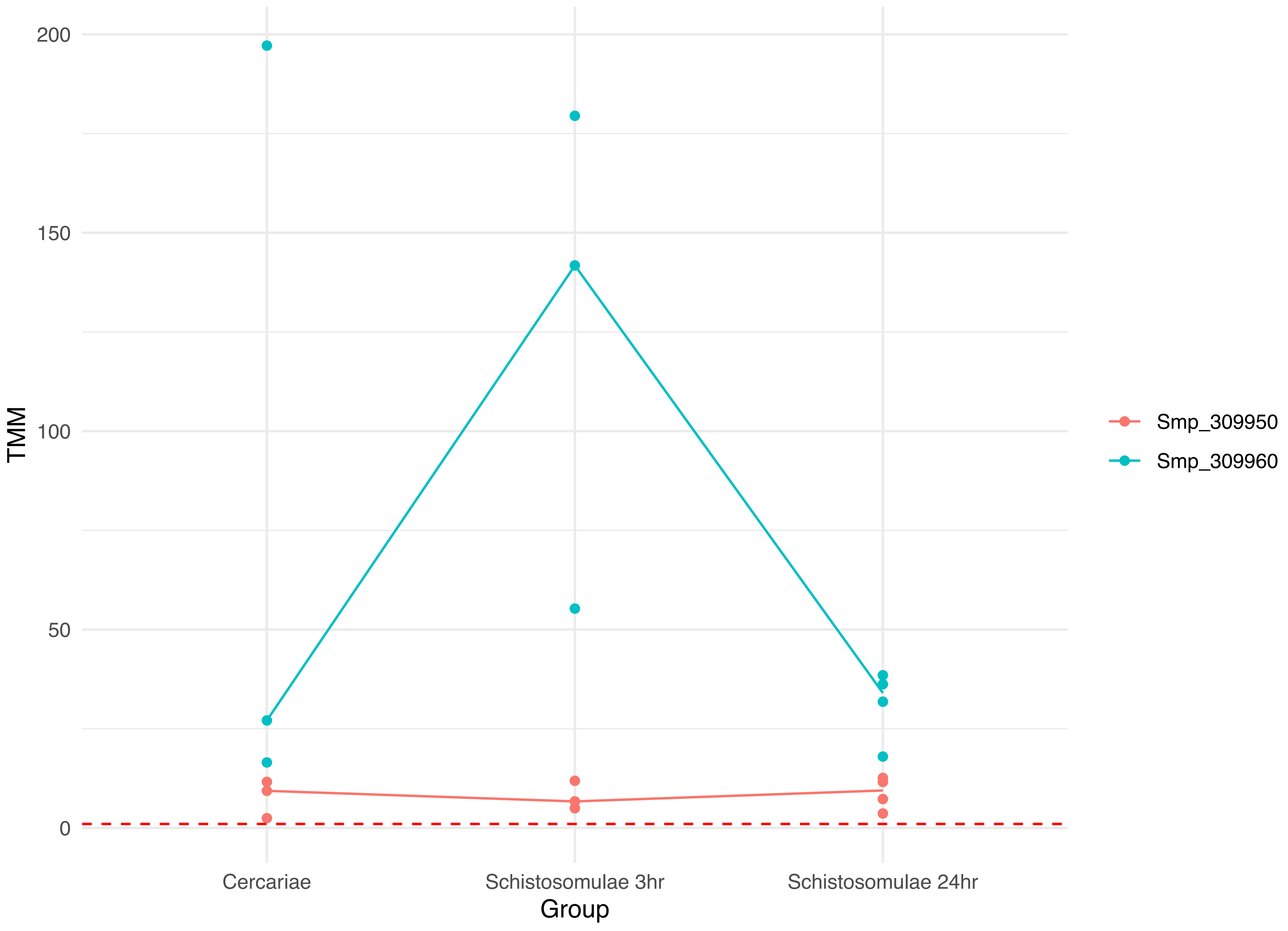

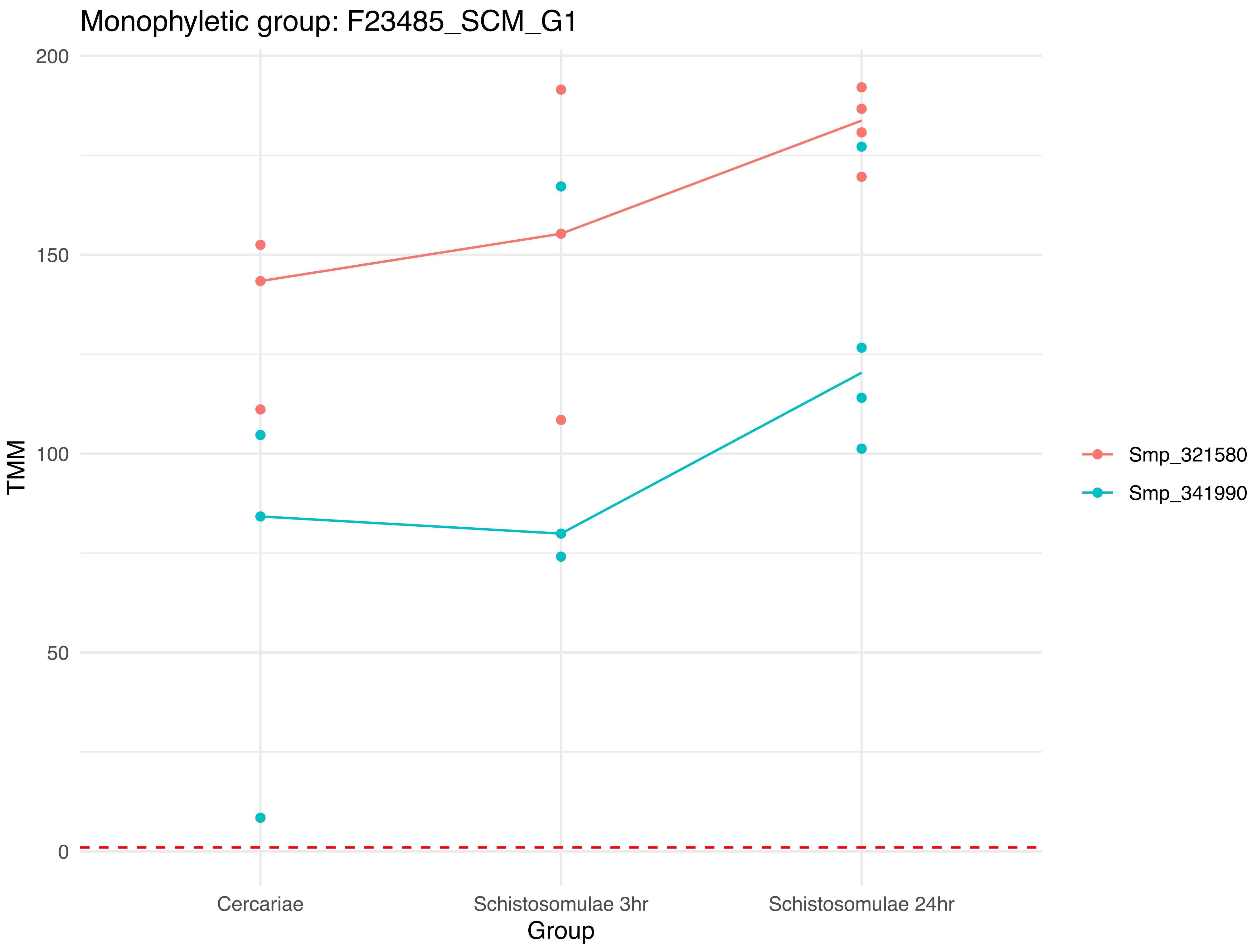

Monophyletic group: F23485_SCM_G1
200
150
100
50
0
TMM
Cercariae
Schistosomulae 3hr
Schistosomulae 24hr
Group
Smp_321580
Smp_341990

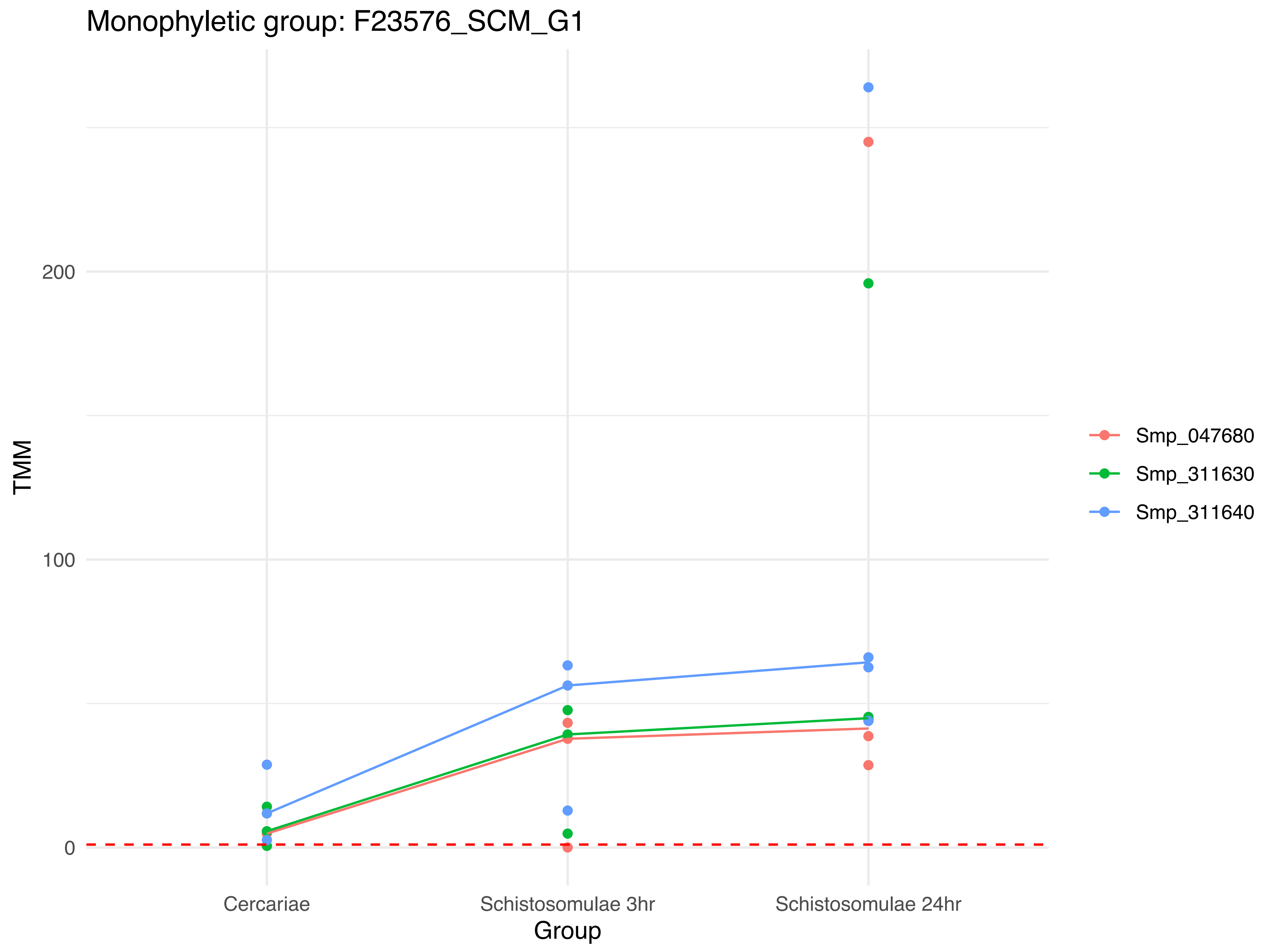

Monophyletic group: F23576_SCM_G1
TMM
200
100
0
Cercariae
Schistosomulae 3hr
Schistosomulae 24hr
Group
Smp_047680
Smp_311630
Smp_311640

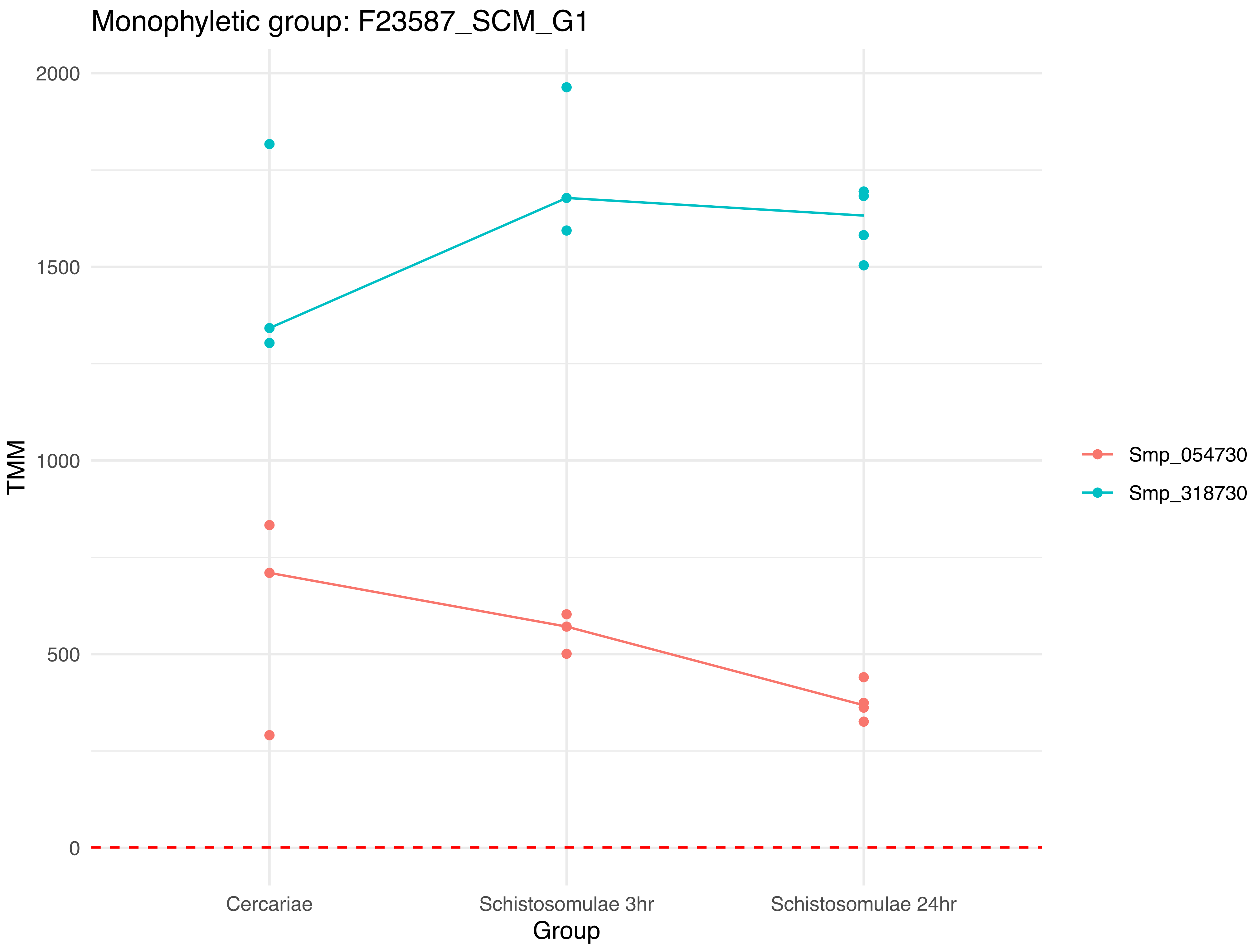
Monophyletic group: F23587_SCM_G1
2000
1500
1000
500
0
TMM
Cercariae
Schistosomulae 3hr
Schistosomulae 24hr
Group
Smp_054730
Smp_318730

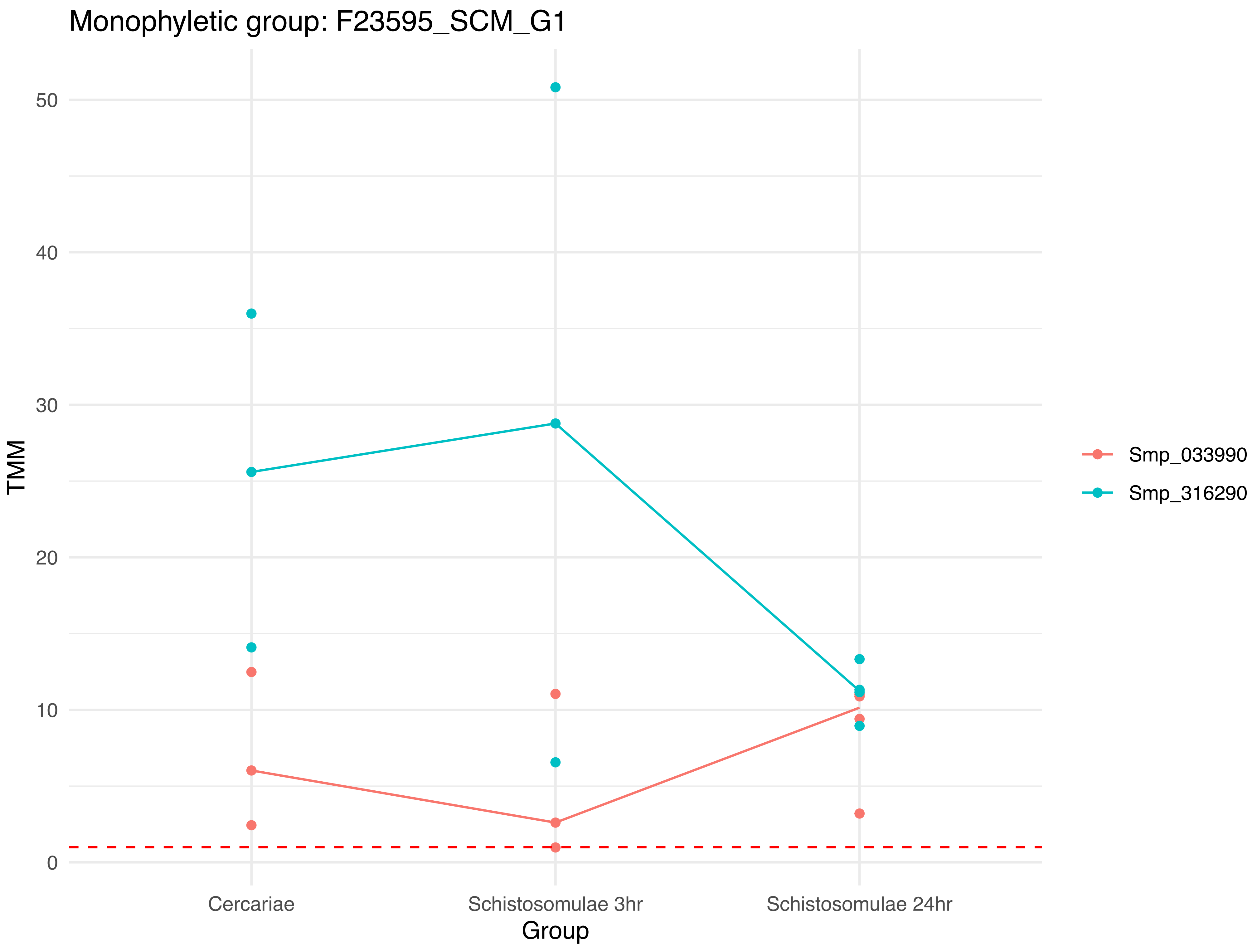
Monophyletic group: F23595_SCM_G1
TMM
50
40
30
20
10
0
Cercariae
Schistosomulae 3hr
Schistosomulae 24hr
Group
Smp_033990
Smp_316290

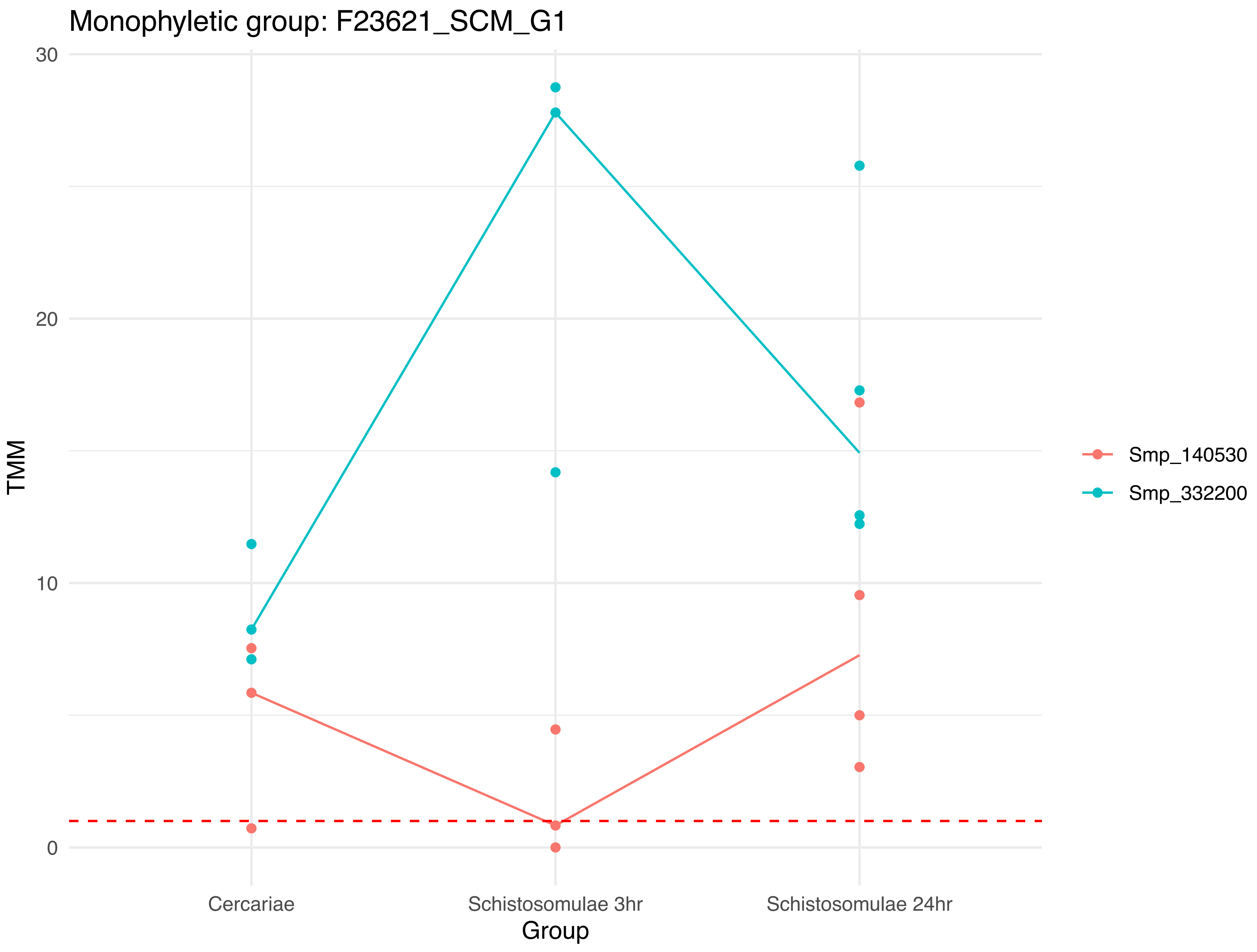

Monophyletic group: F23621_SCM_G1
30
20
10
0
TMM
Cercariae
Schistosomulae 3hr
Schistosomulae 24hr
Group
Smp_140530
Smp_332200

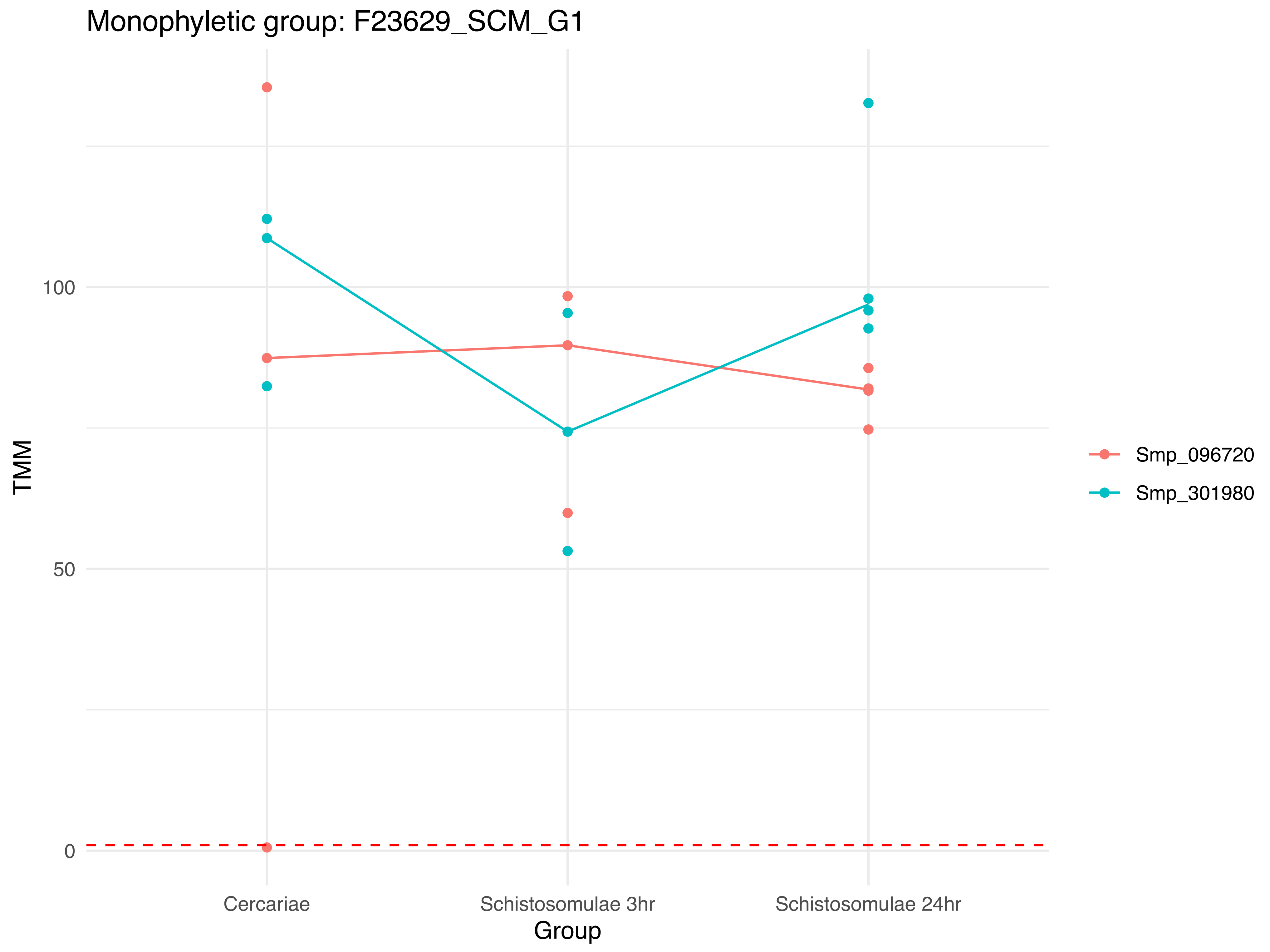
Monophyletic group: F23629_SCM_G1
TMM
100
50
0
Cercariae
Schistosomulae 3hr
Schistosomulae 24hr
Group
Smp_096720
Smp_301980

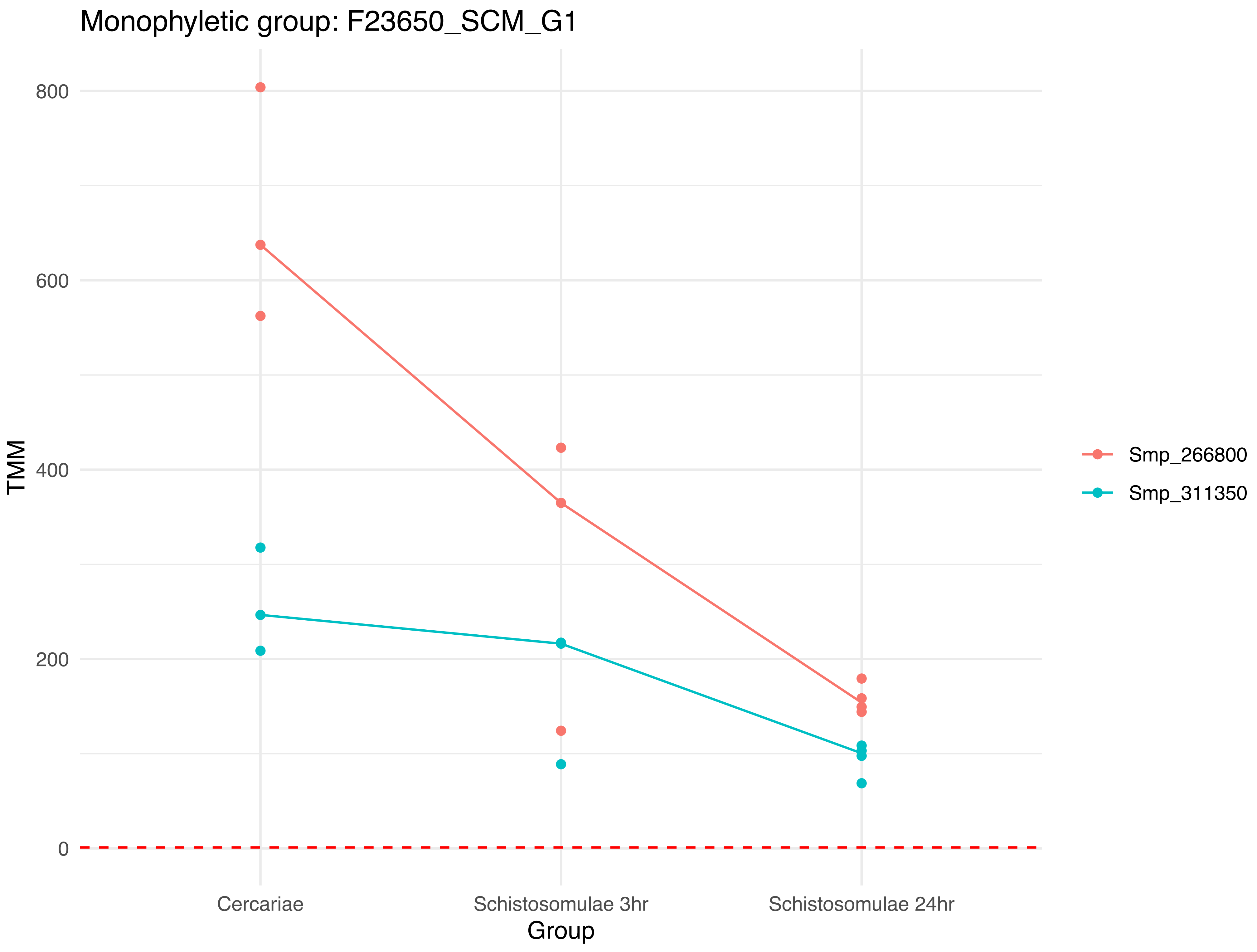
Monophyletic group: F23650_SCM_G1
TMM
800
600
400
200
0
Cercariae
Schistosomulae 3hr
Schistosomulae 24hr
Group
Smp_266800
Smp_311350

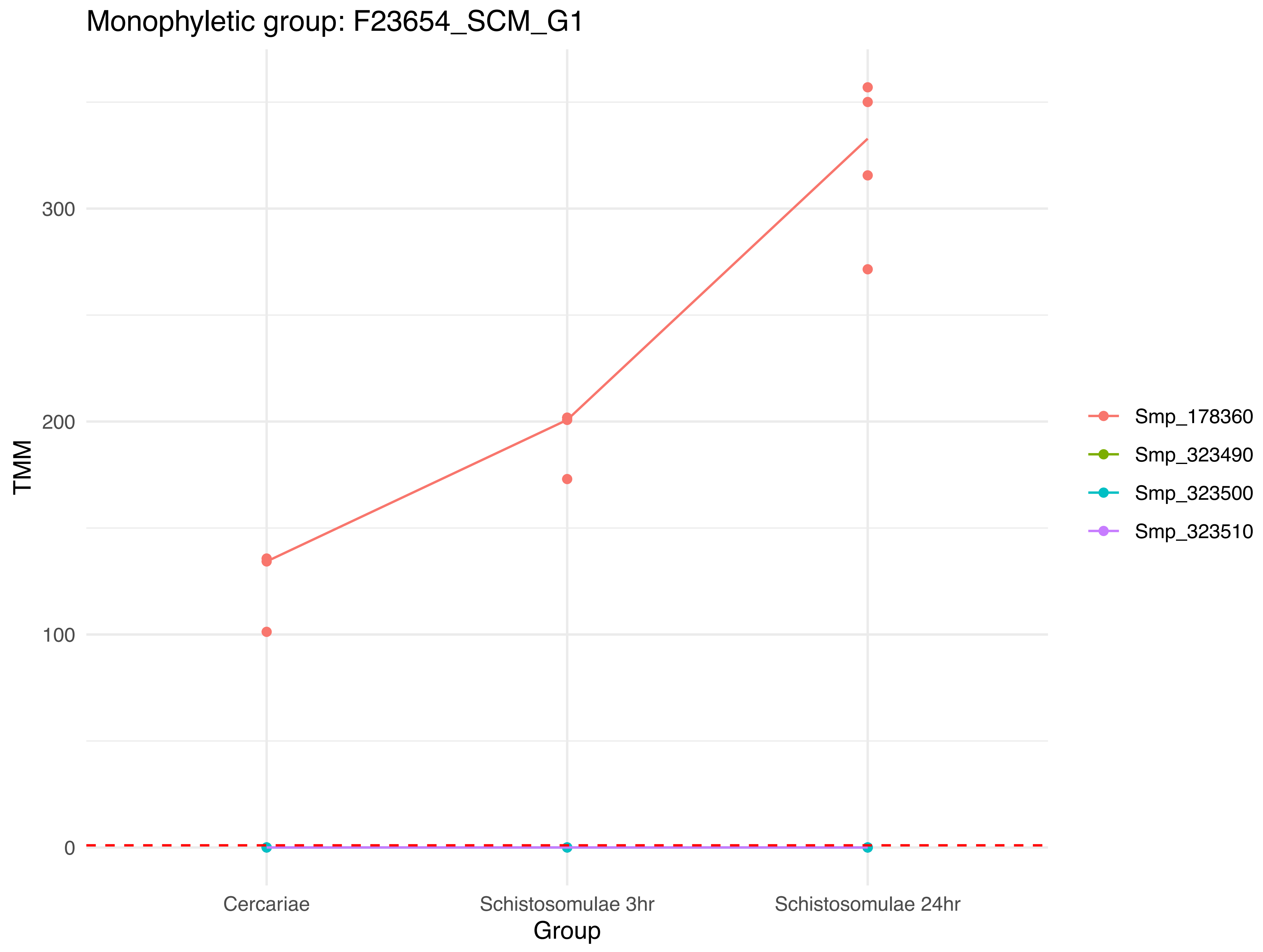

Monophyletic group: F23654_SCM_G1
TMM
300
200
100
0
Cercariae
Schistosomulae 3hr
Schistosomulae 24hr
Group
Smp_178360
Smp_323490
Smp_323500
Smp_323510

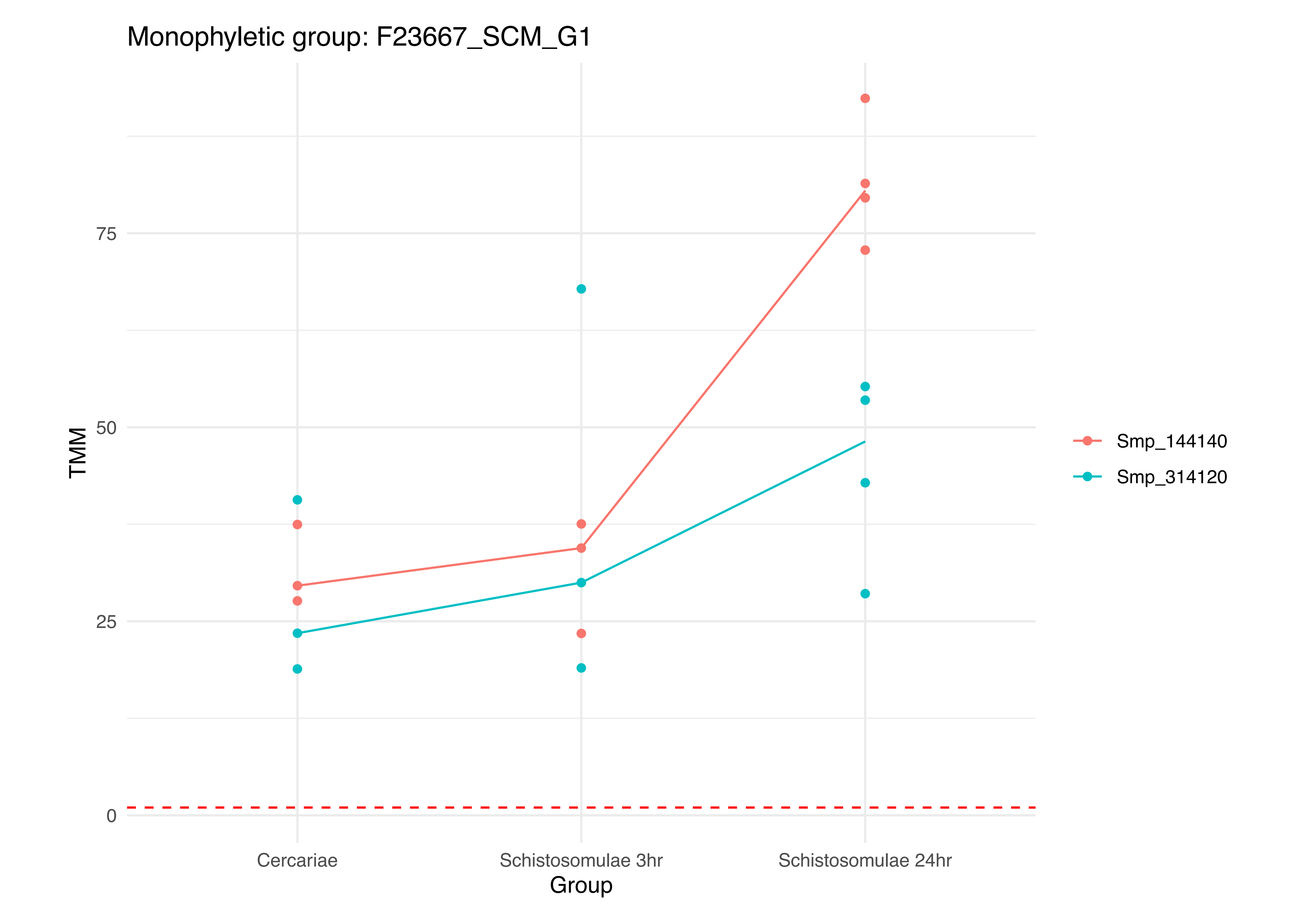
Monophyletic group: F23667_SCM_G1
TMM
75
50
25
0
Cercariae
Schistosomulae 3hr
Schistosomulae 24hr
Group
Smp_144140
Smp_314120

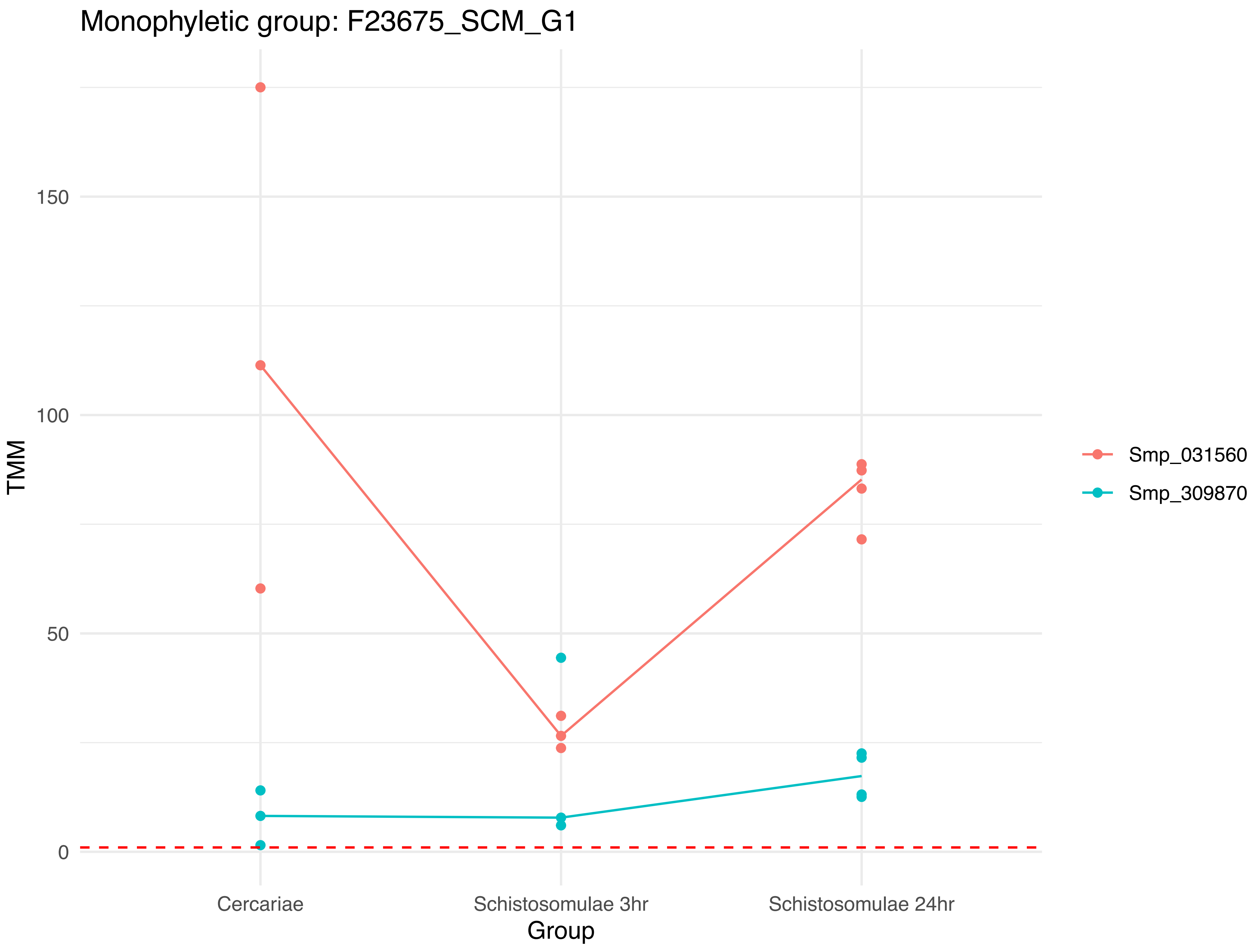

Monophyletic group: F23675_SCM_G1
TMM
150
100
50
0
Cercariae
Schistosomulae 3hr
Schistosomulae 24hr
Group
Smp_031560
Smp_309870

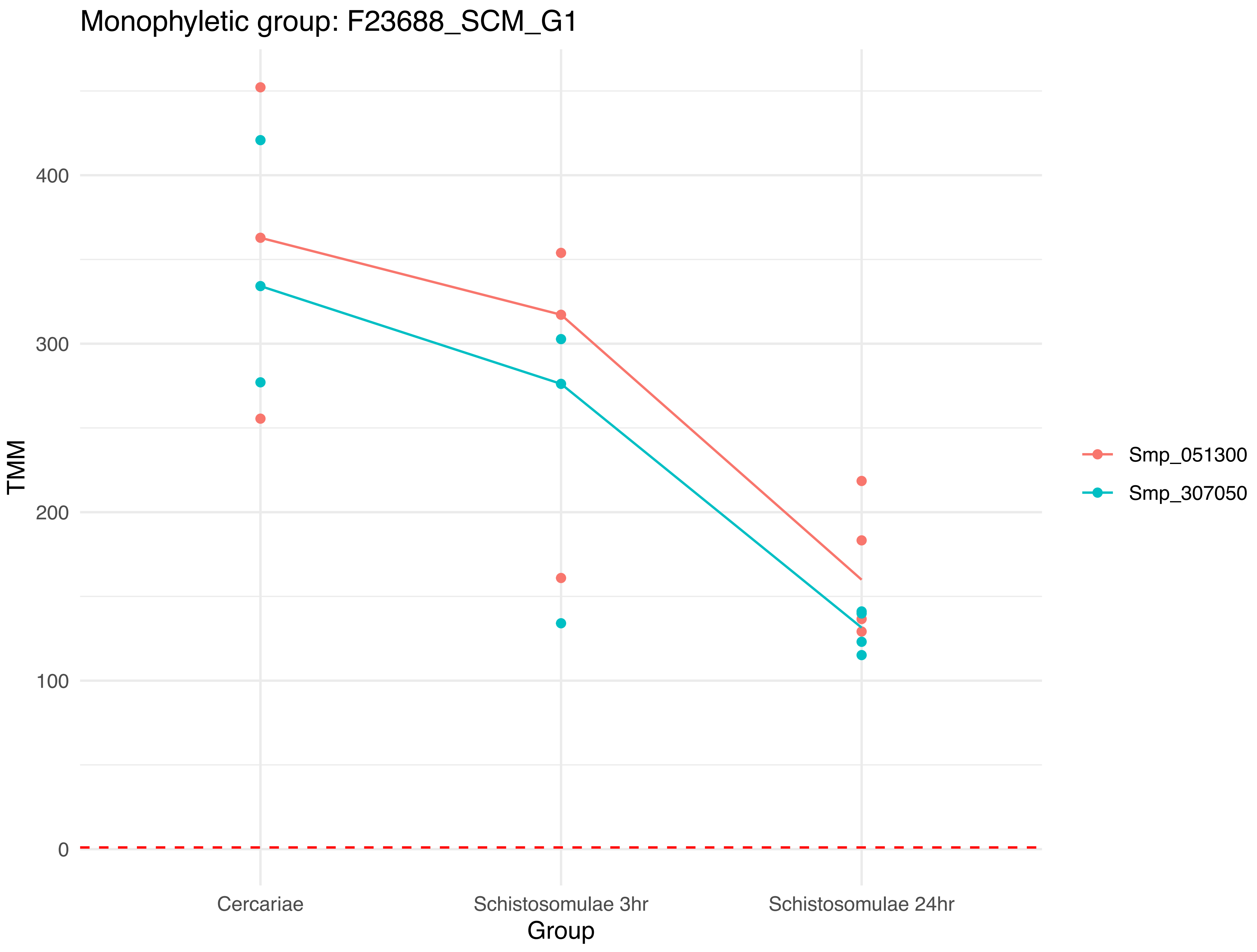
Monophyletic group: F23688_SCM_G1
TMM
400
300
200
100
0
Cercariae
Schistosomulae 3hr
Schistosomulae 24hr
Group
Smp_051300
Smp_307050

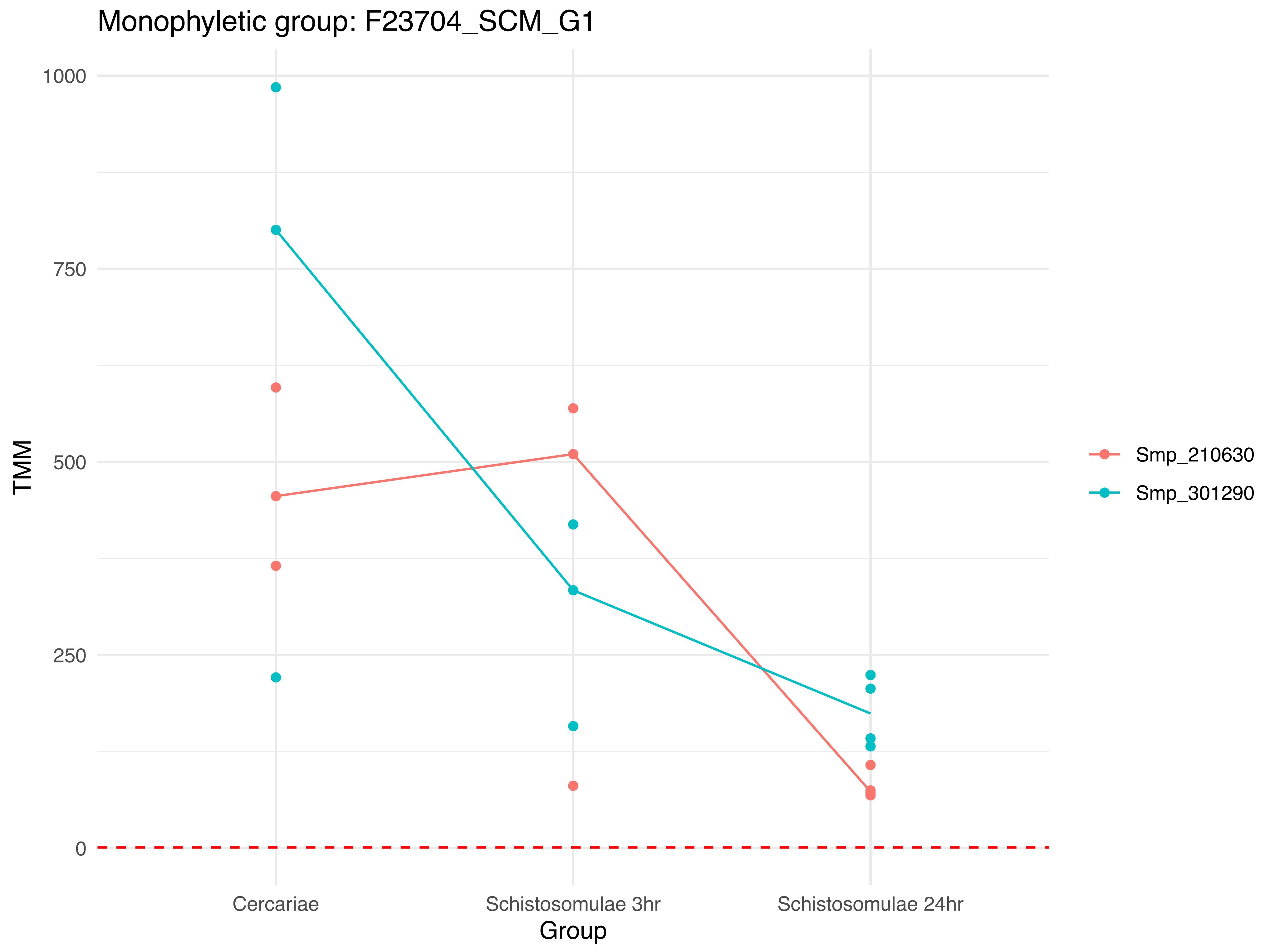

Monophyletic group: F23704_SCM_G1
1000
750
500
250
0
TMM
Cercariae
Schistosomulae 3hr
Schistosomulae 24hr
Group
Smp_210630
Smp_301290

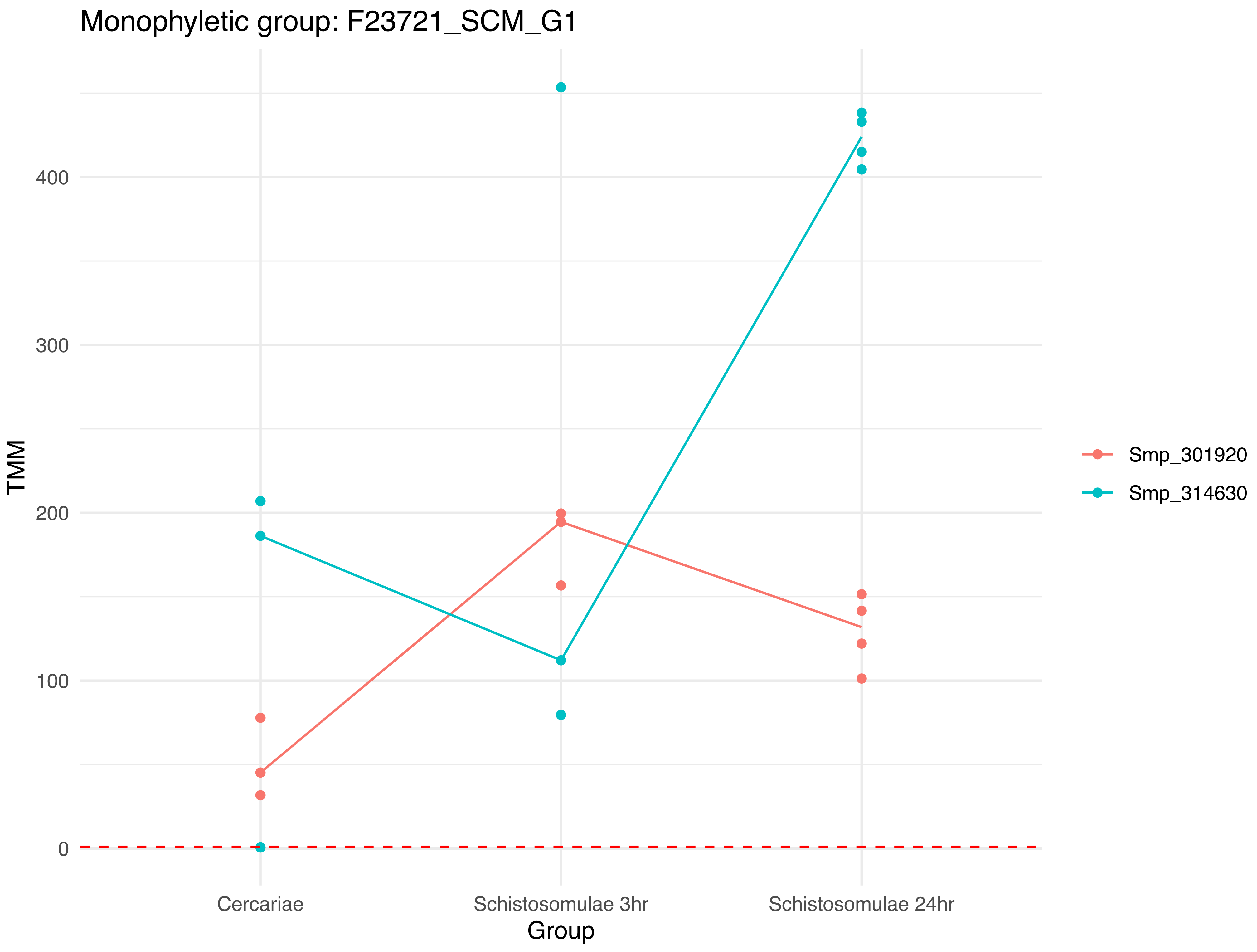

Monophyletic group: F23721_SCM_G1
TMM
400
300
200
100
0
Cercariae
Schistosomulae 3hr
Schistosomulae 24hr
Group
Smp_301920
Smp_314630

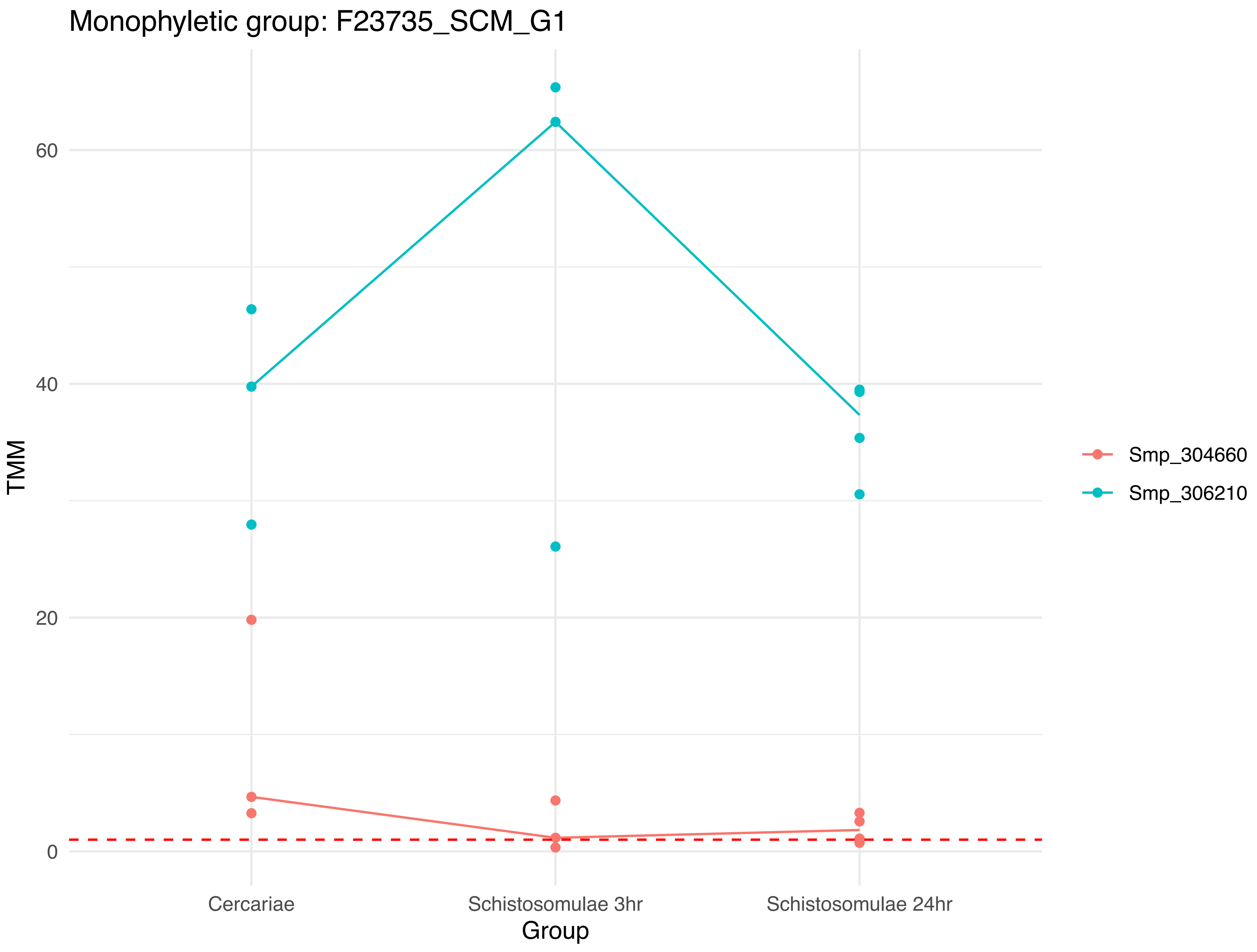

Monophyletic group: F23735_SCM_G1
TMM
60
40
20
0
Cercariae
Schistosomulae 3hr
Schistosomulae 24hr
Group
Smp_304660
Smp_306210

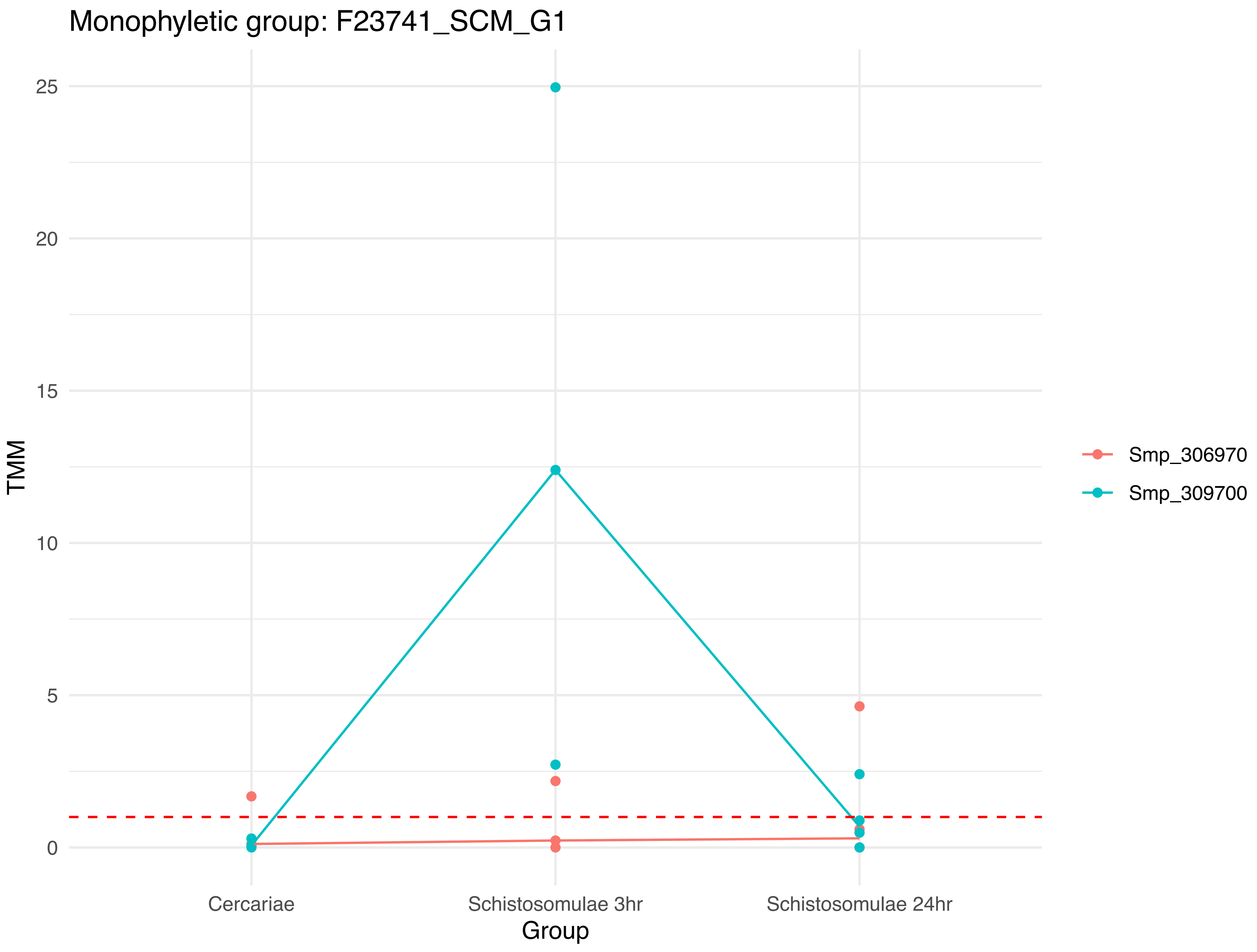
Monophyletic group: F23741_SCM_G1
TMM
25
20
15
10
5
0
Cercariae
Schistosomulae 3hr
Schistosomulae 24hr
Group
Smp_306970
Smp_309700

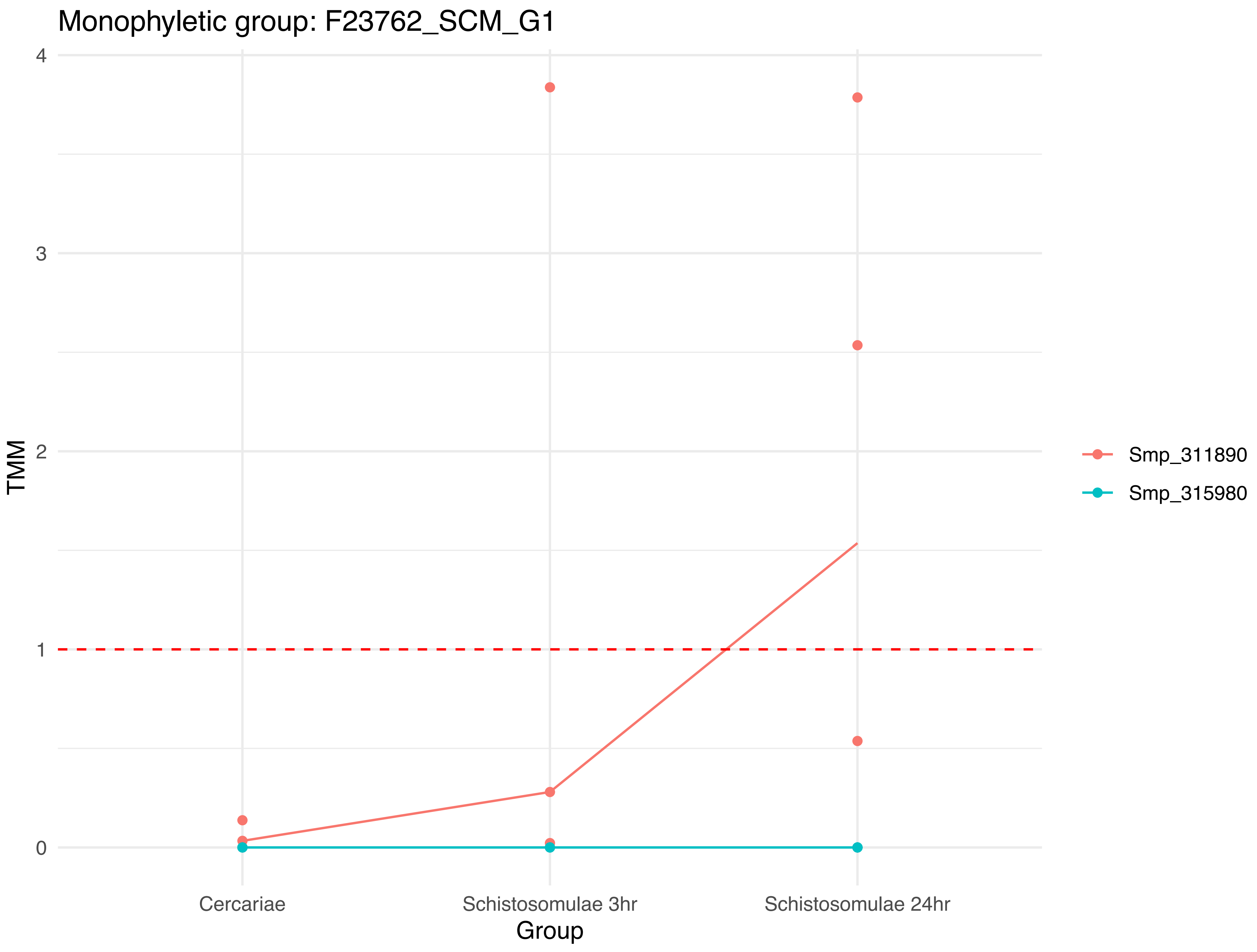

Monophyletic group: F23762_SCM_G1
TMM
4
3
2
1
0
Cercariae
Schistosomulae 3hr
Schistosomulae 24hr
Group
Smp_311890
Smp_315980

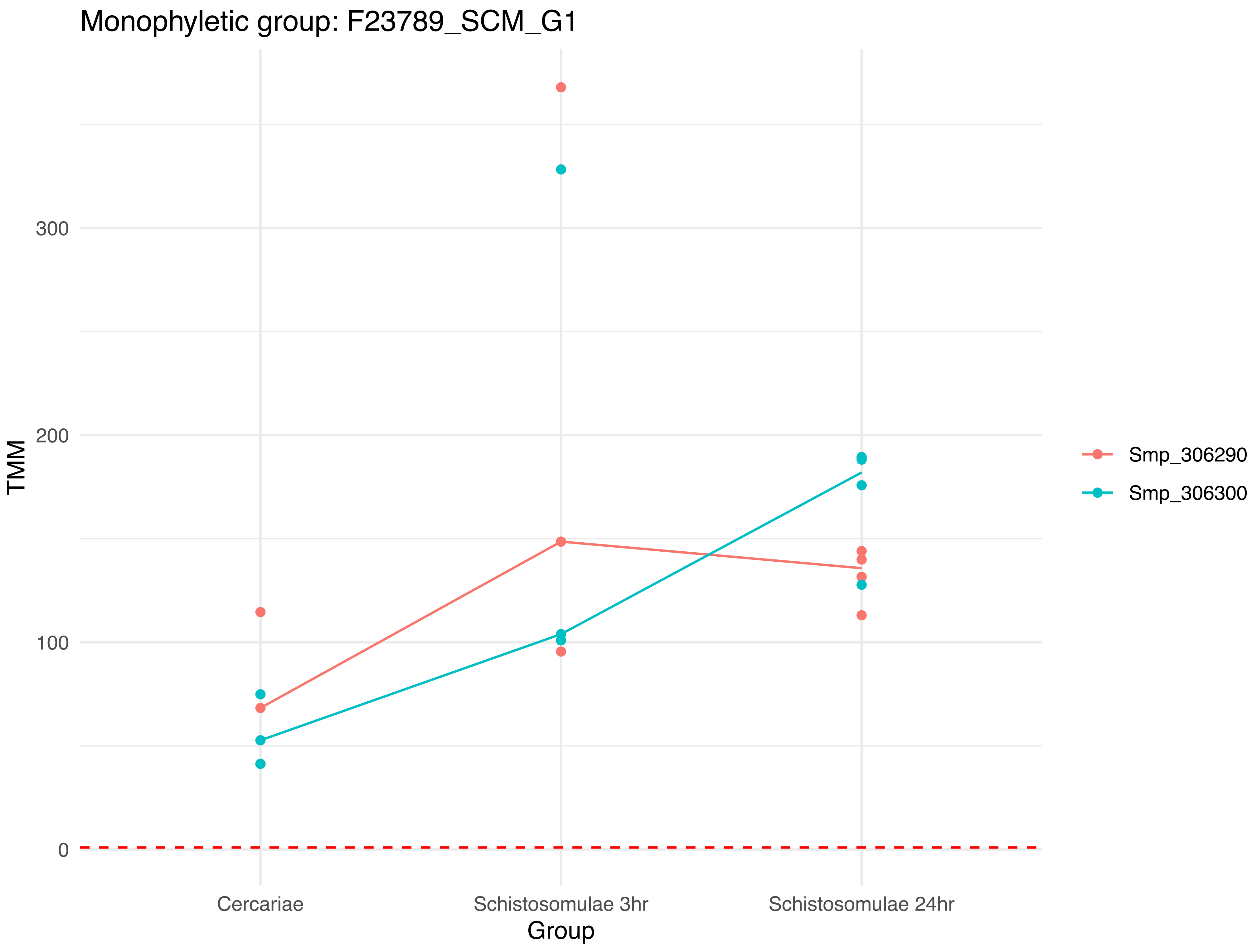
Monophyletic group: F23789_SCM_G1
TMM
300
200
100
0
Cercariae
Schistosomulae 3hr
Schistosomulae 24hr
Group
Smp_306290
Smp_306300

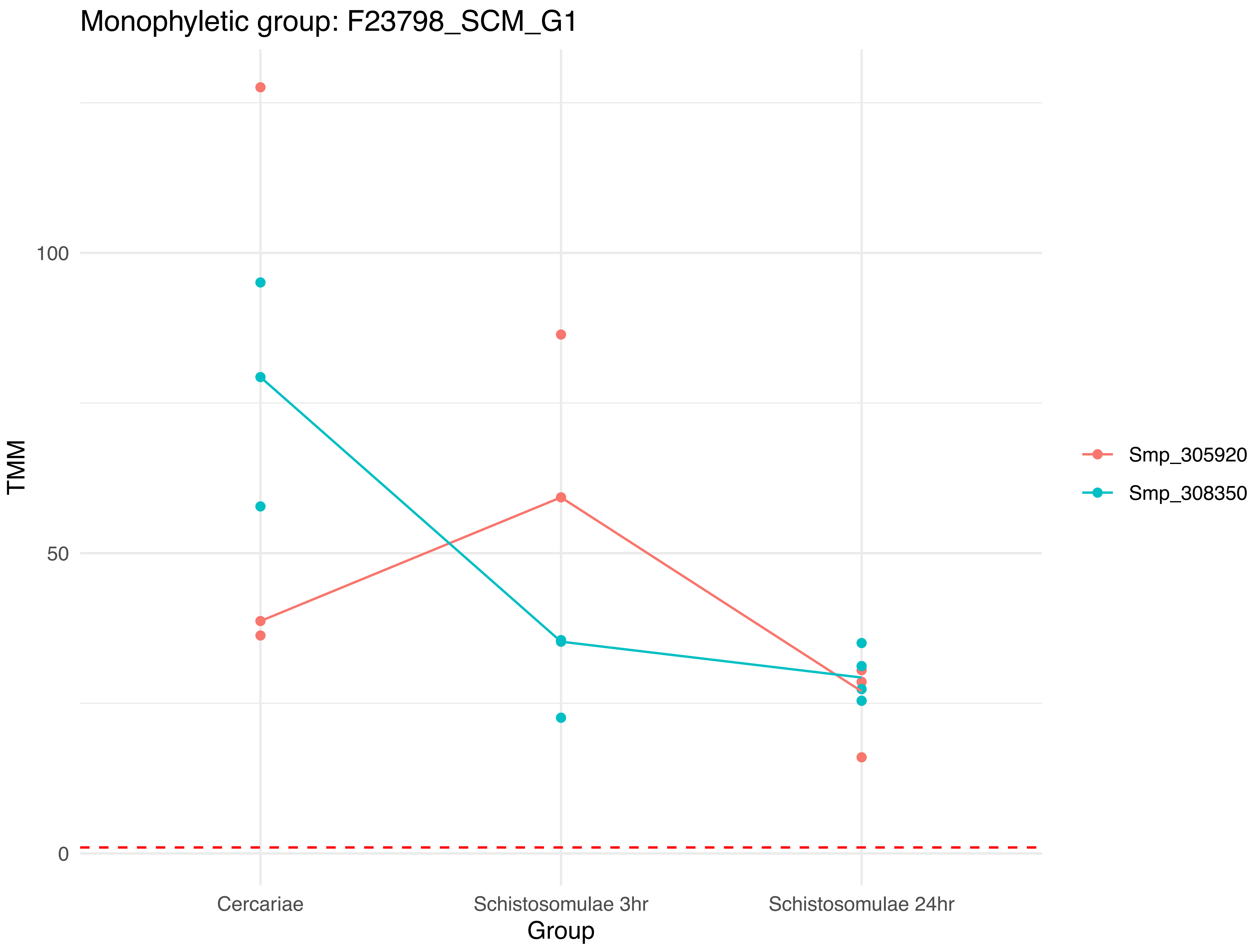

Monophyletic group: F23798_SCM_G1
TMM
100
50
0
Cercariae
Schistosomulae 3hr
Schistosomulae 24hr
Group
Smp_305920
Smp_308350

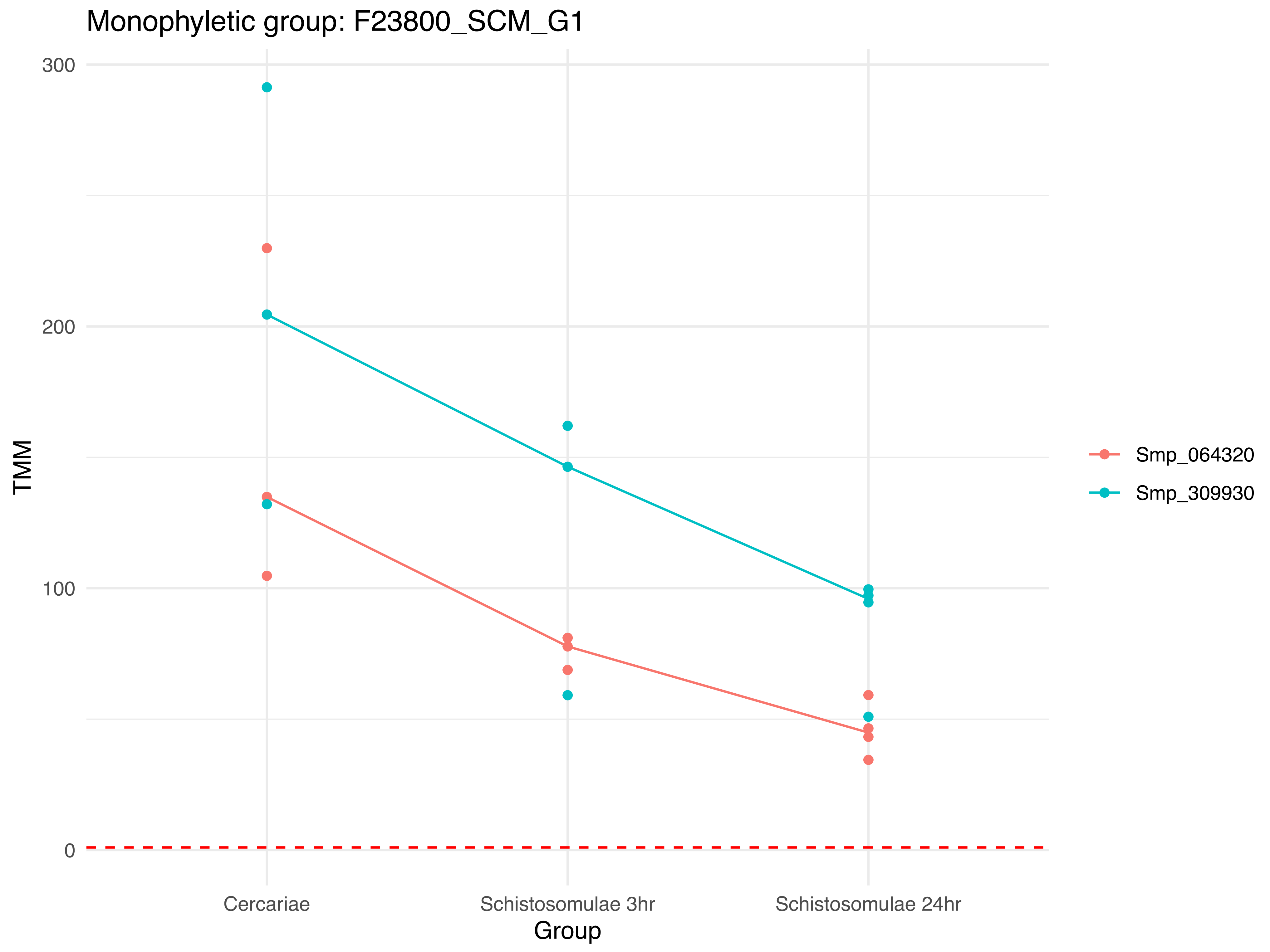
Monophyletic group: F23800_SCM_G1
TMM
300
200
100
0
Cercariae
Schistosomulae 3hr
Schistosomulae 24hr
Group
Smp_064320
Smp_309930

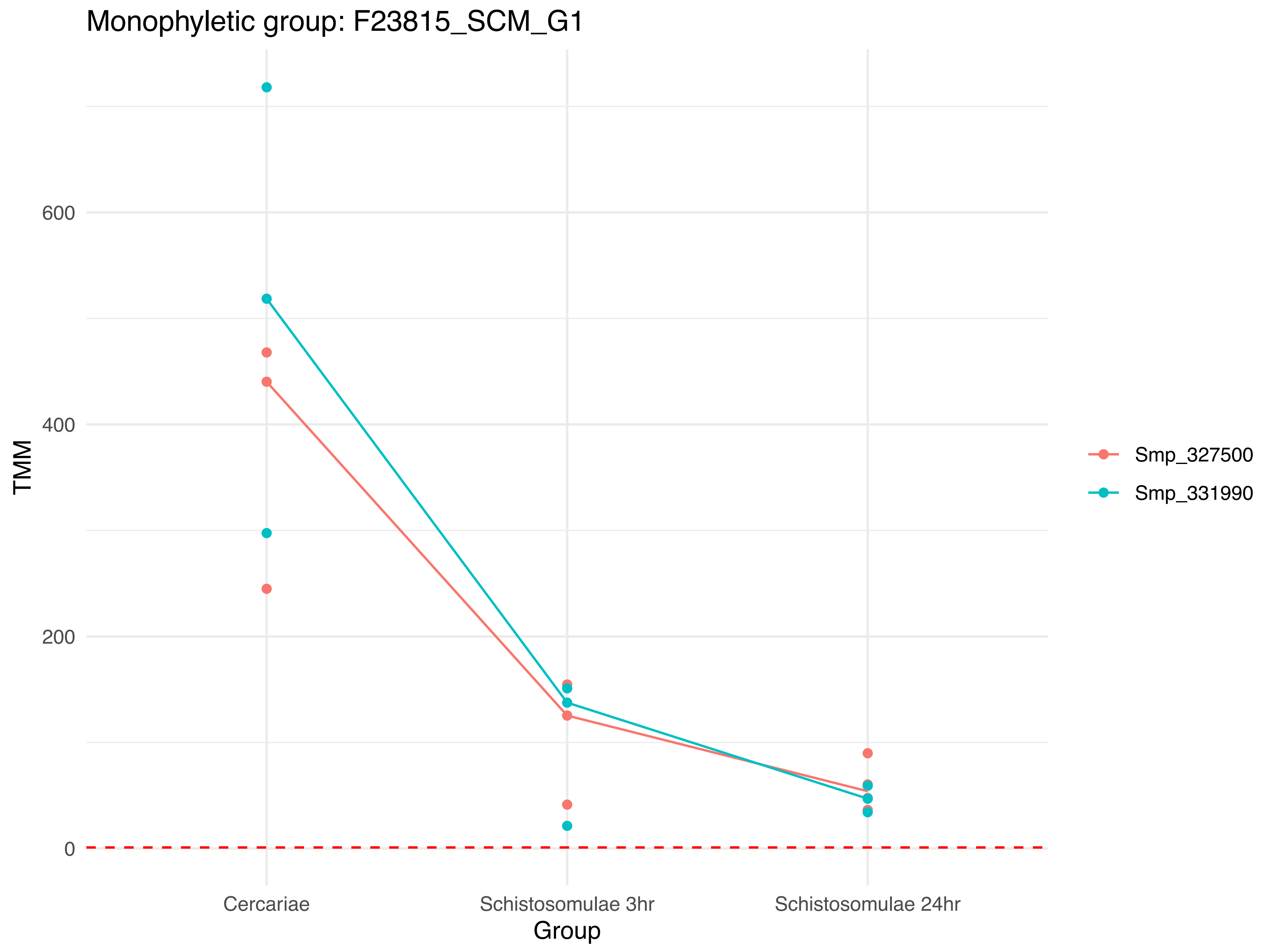

Monophyletic group: F23815_SCM_G1
600
400
200
0
TMM
Cercariae
Schistosomulae 3hr
Schistosomulae 24hr
Group
Smp_327500
Smp_331990

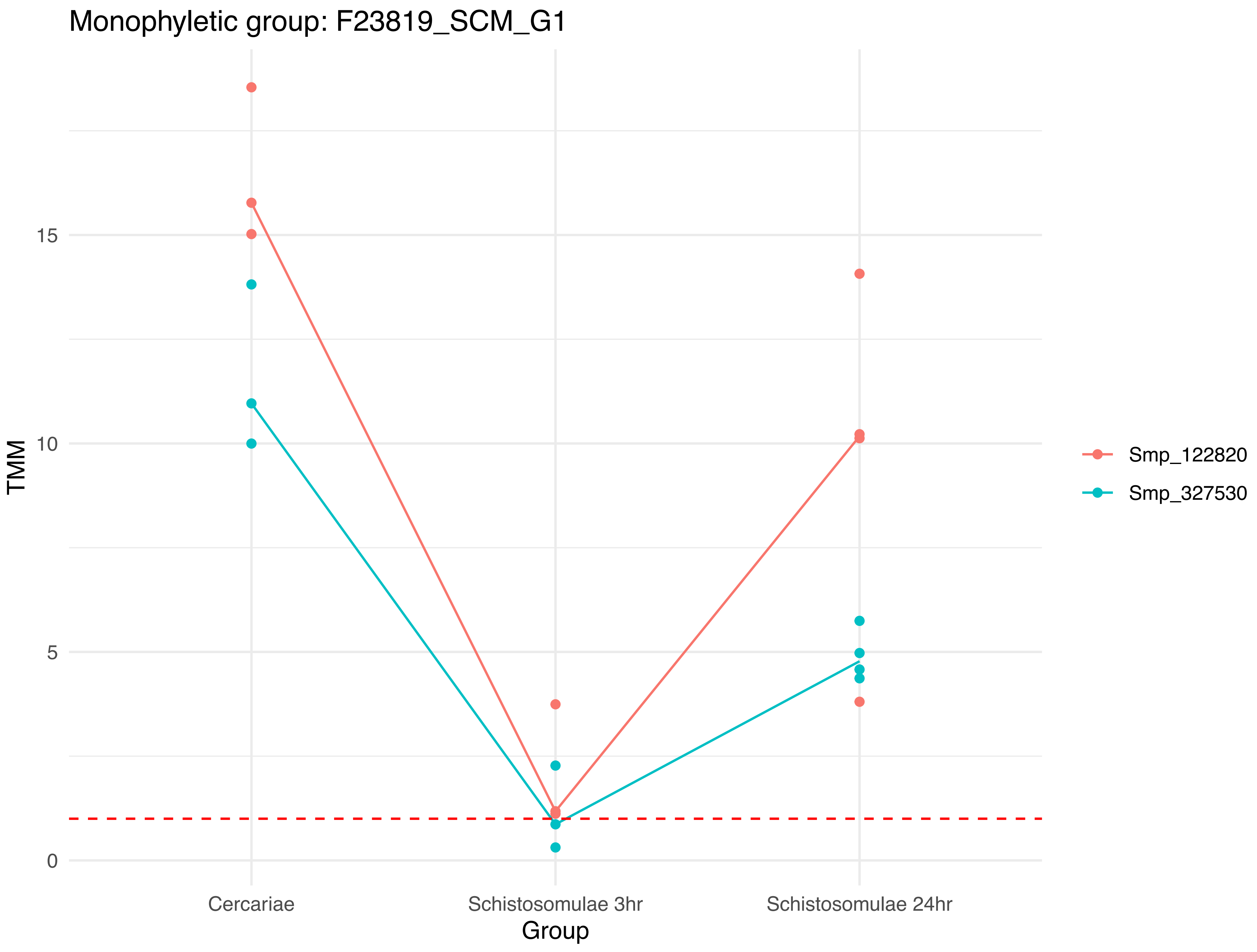

Monophyletic group: F23819_SCM_G1
TMM
15
10
5
0
Cercariae
Schistosomulae 3hr
Schistosomulae 24hr
Group
Smp_122820
Smp_327530

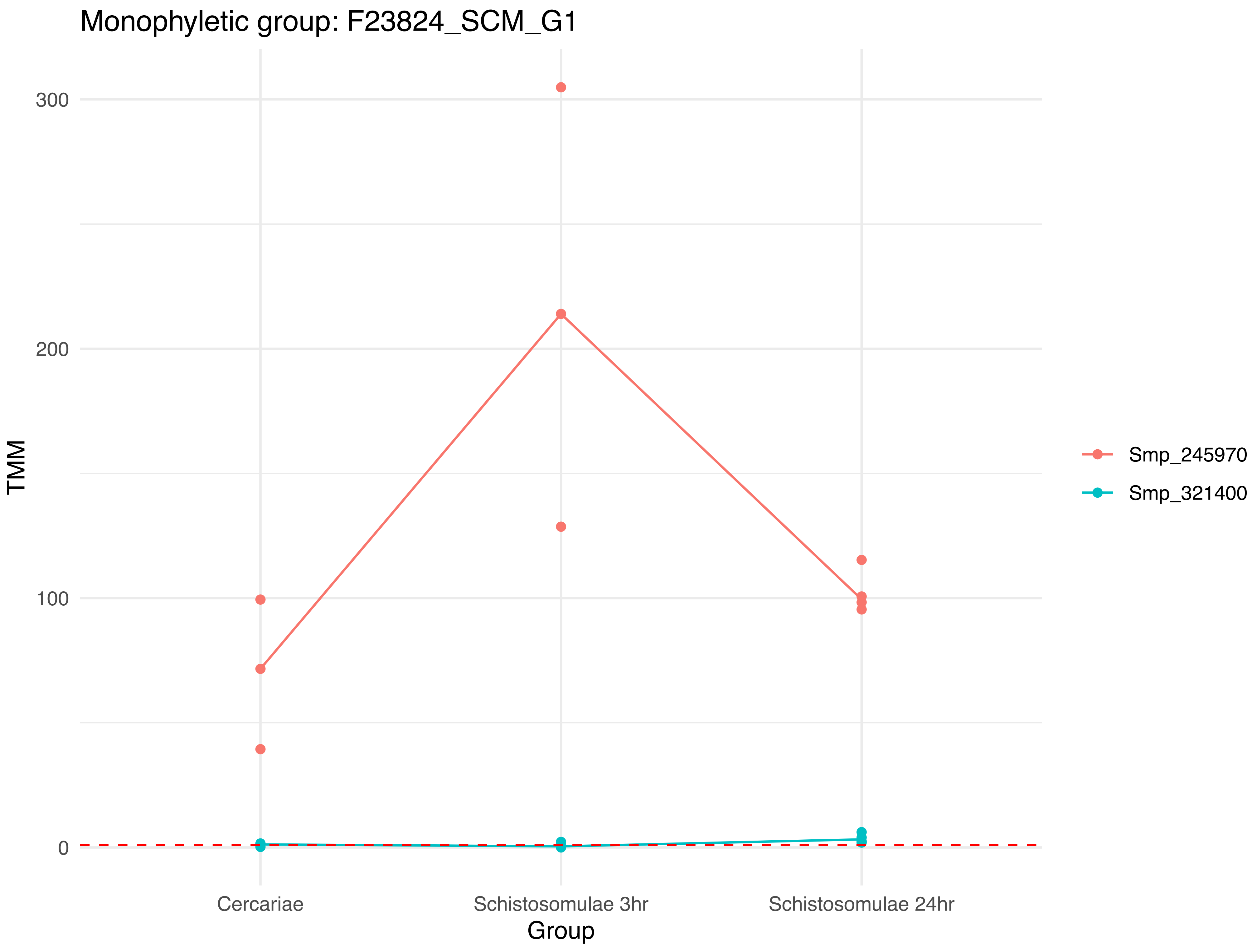

Monophyletic group: F23824_SCM_G1
300
200
100
0
TMM
Cercariae
Schistosomulae 3hr
Schistosomulae 24hr
Group
Smp_245970
Smp_321400

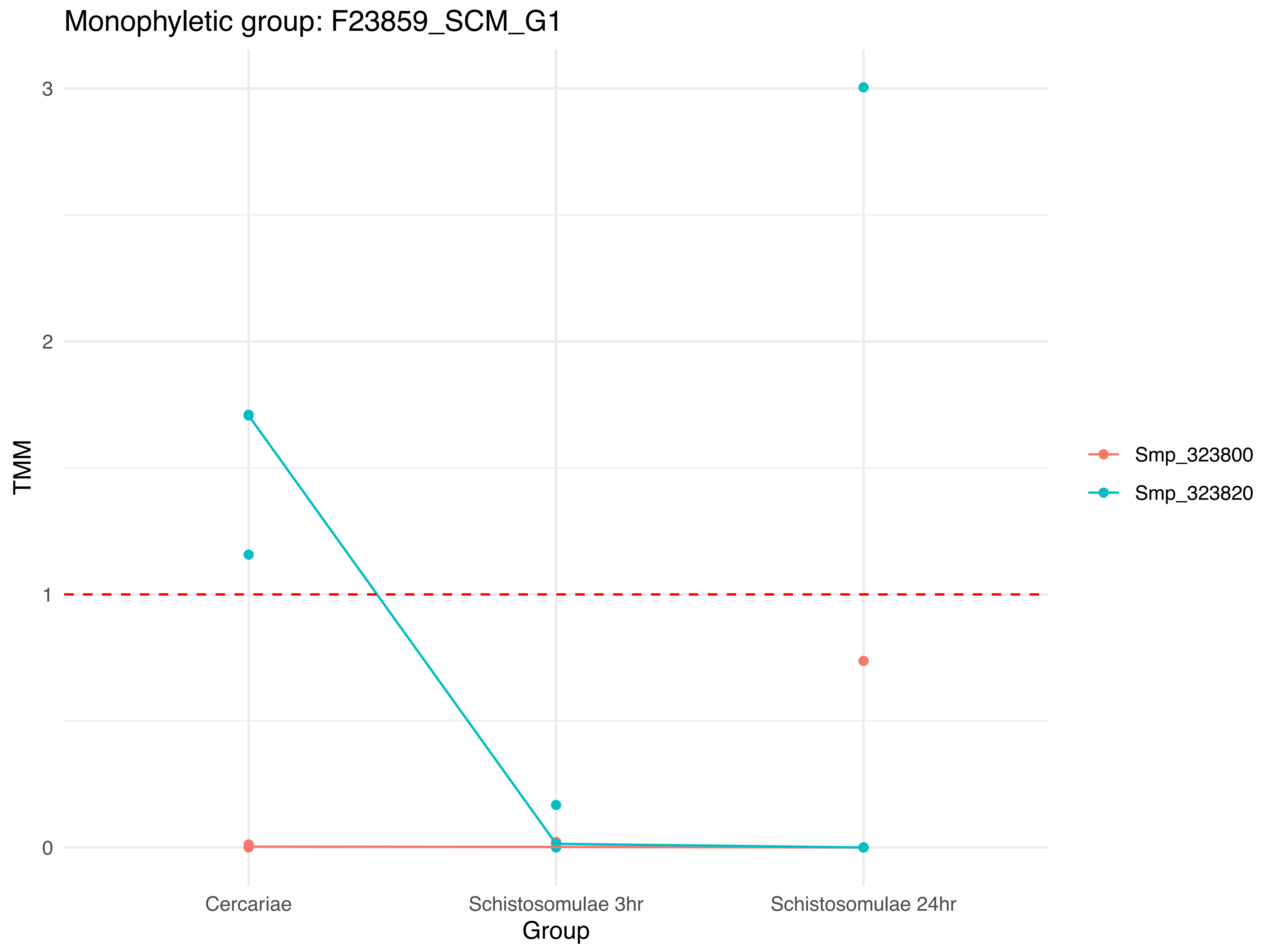

Monophyletic group: F23859_SCM_G1
TMM
3
2
1
0
Cercariae
Schistosomulae 3hr
Schistosomulae 24hr
Group
Smp_323800
Smp_323820

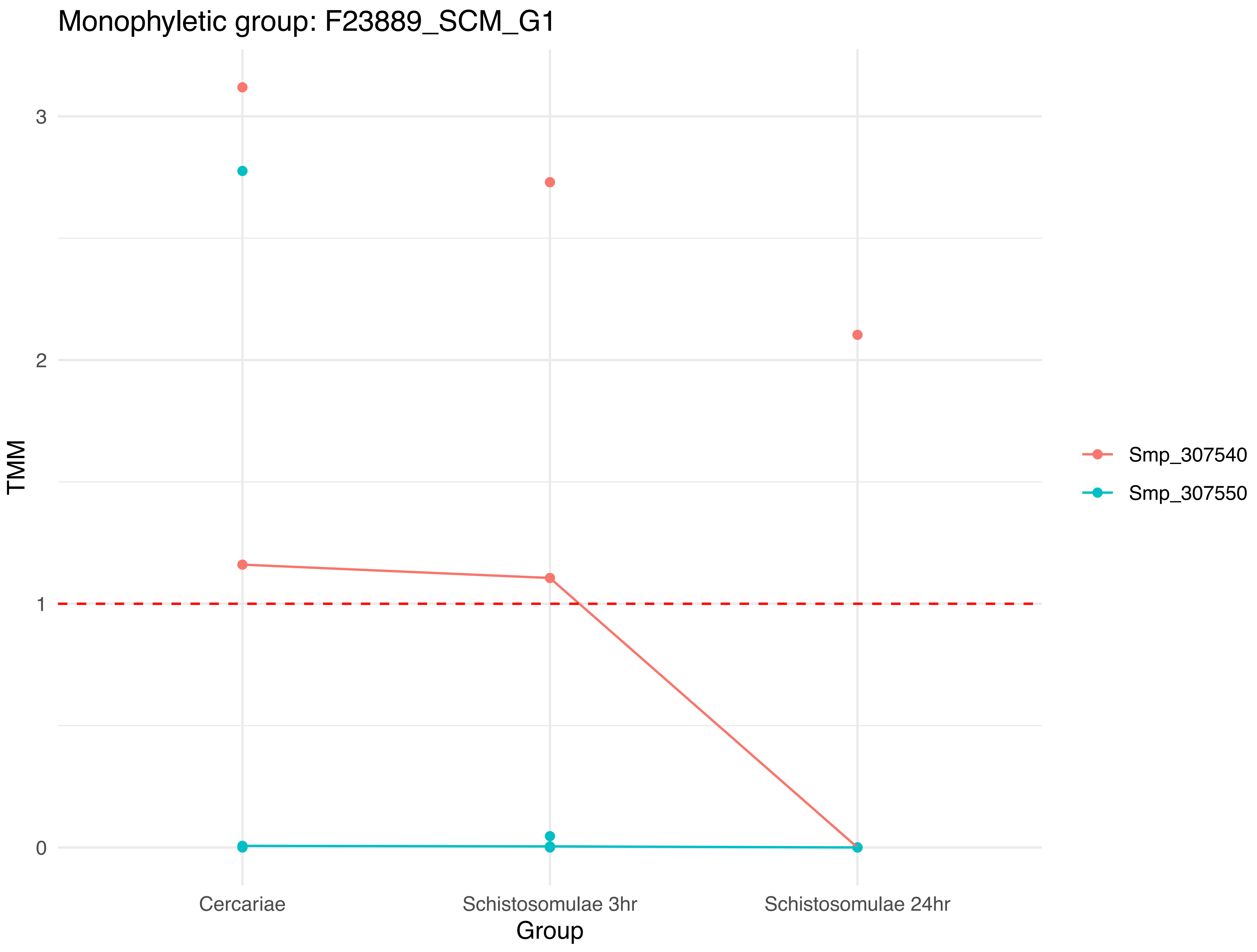

Monophyletic group: F23889_SCM_G1
TMM
3
2
1
0
Cercariae
Schistosomulae 3hr
Schistosomulae 24hr
Group
Smp_307540
Smp_307550

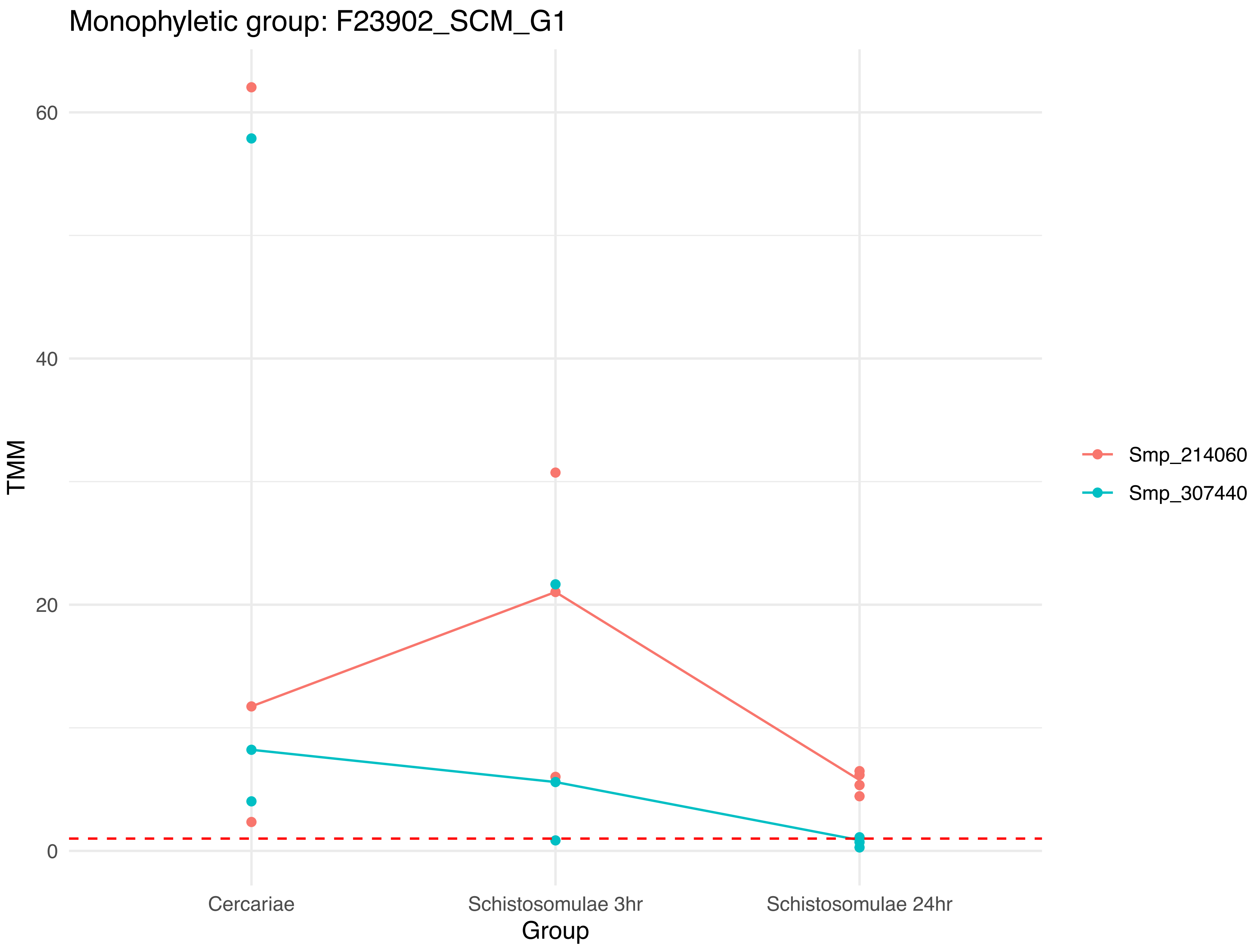

Monophyletic group: F23902_SCM_G1
TMM
60
40
20
0
Cercariae
Schistosomulae 3hr
Schistosomulae 24hr
Group
Smp_214060
Smp_307440

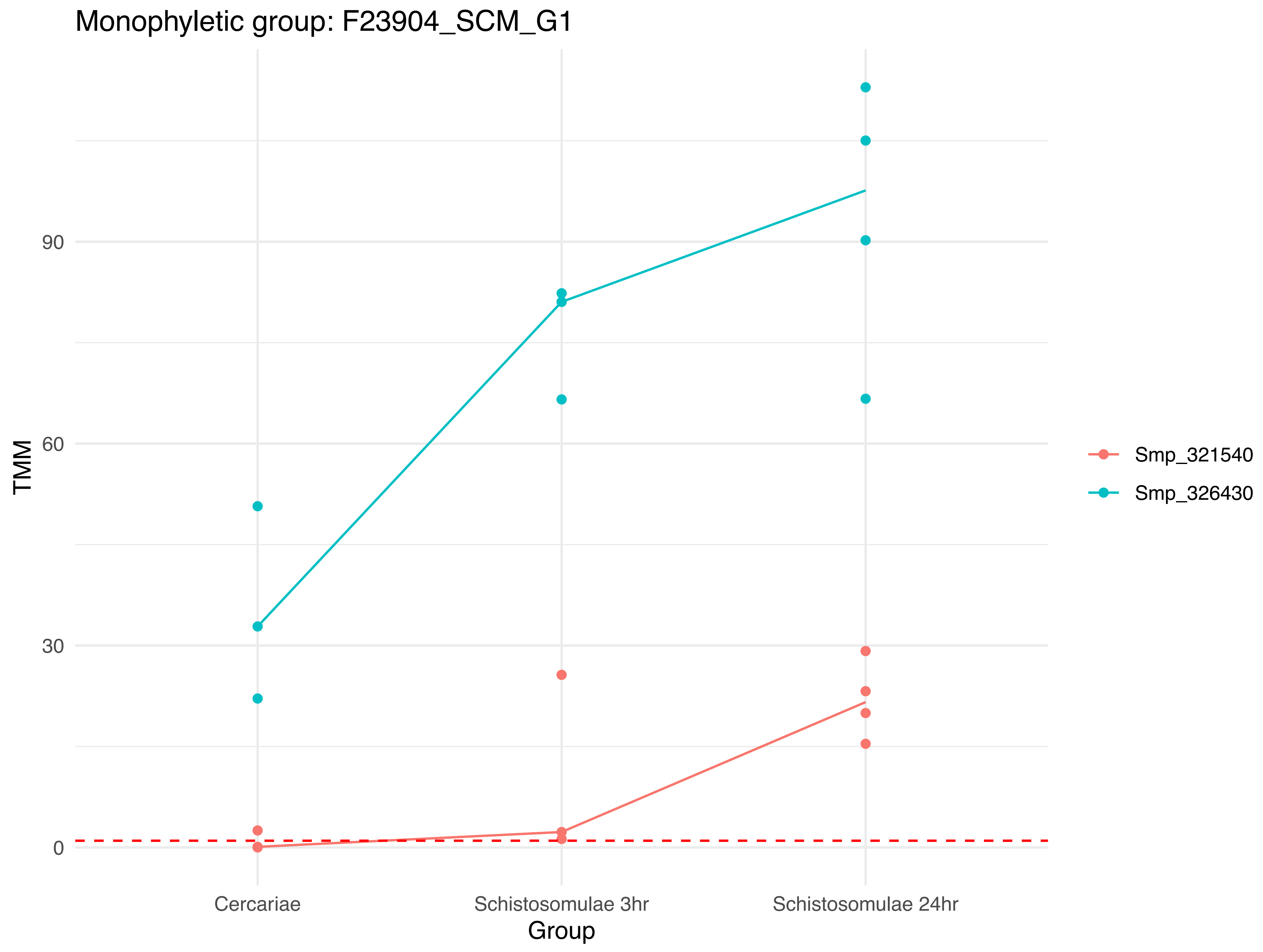

Monophyletic group: F23904_SCM_G1
TMM
90
60
30
0
Cercariae
Schistosomulae 3hr
Schistosomulae 24hr
Group
Smp_321540
Smp_326430

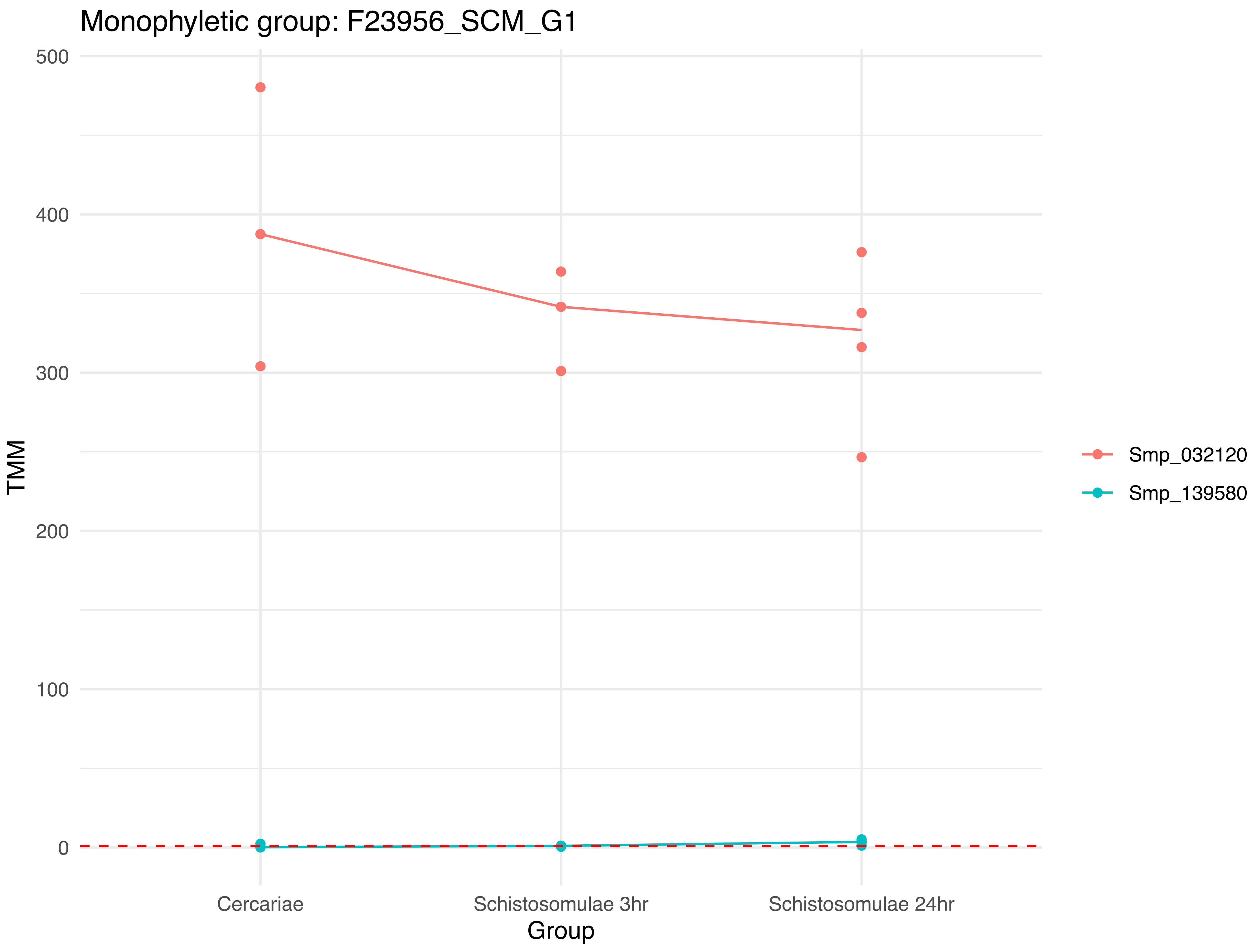

Monophyletic group: F23956_SCM_G1
500
400
300
200
100
0
TMM
Cercariae
Schistosomulae 3hr
Schistosomulae 24hr
Group
Smp_032120
Smp_139580

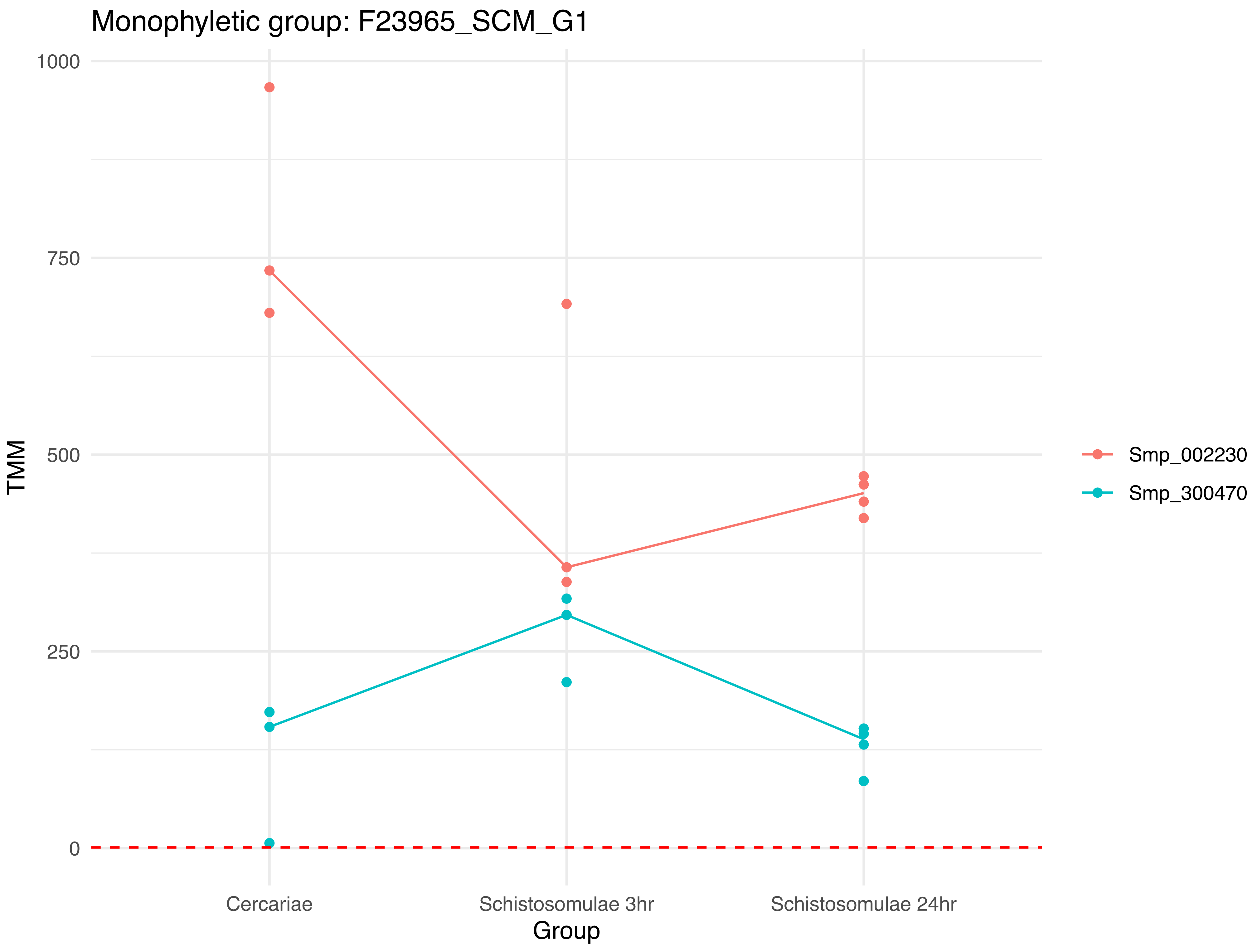

Monophyletic group: F23965_SCM_G1
1000
750
500
250
0
TMM
Cercariae
Schistosomulae 3hr
Schistosomulae 24hr
Group
Smp_002230
Smp_300470

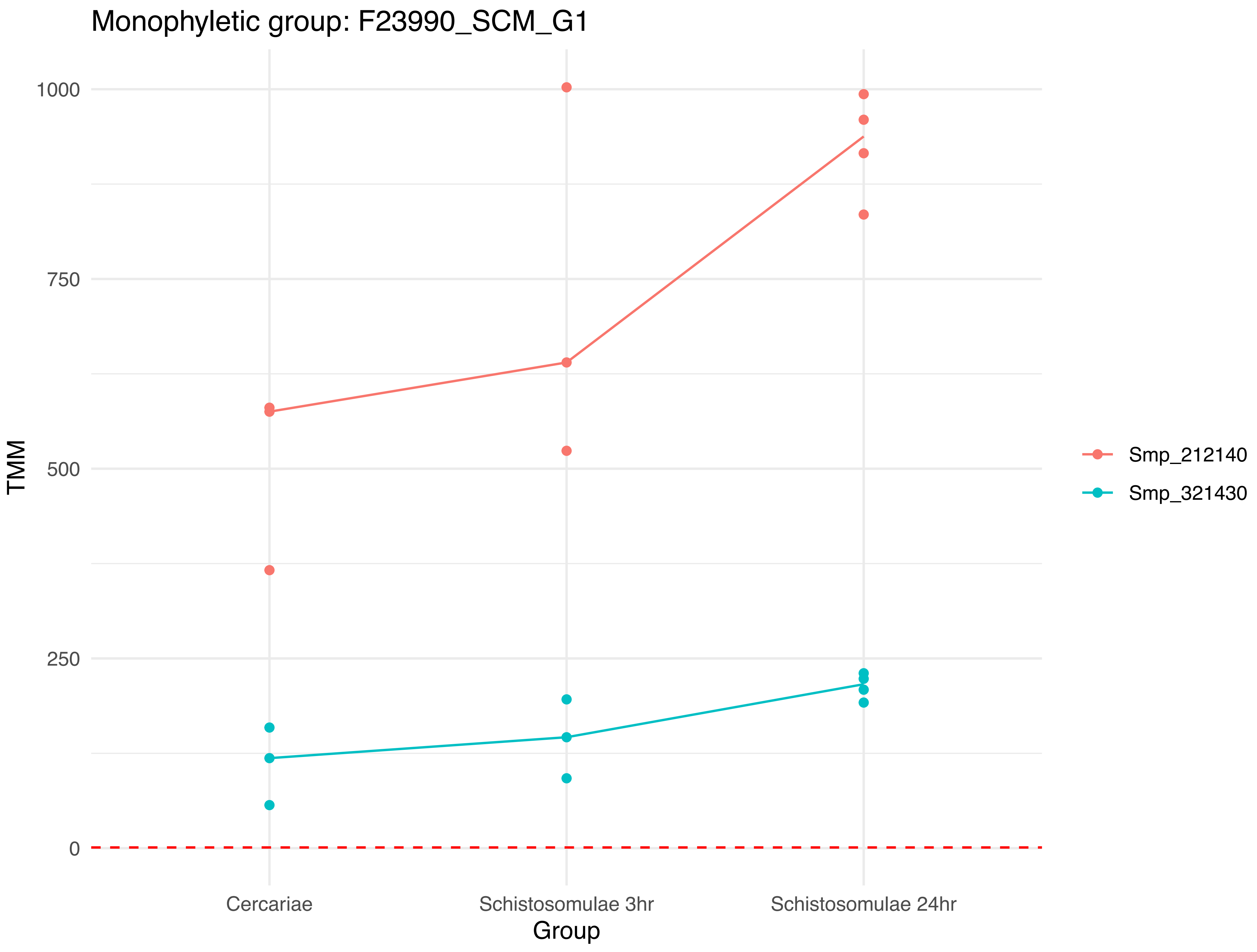
Monophyletic group: F23990_SCM_G1
1000
750
500
250
0
TMM
Cercariae
Schistosomulae 3hr
Schistosomulae 24hr
Group
Smp_212140
Smp_321430

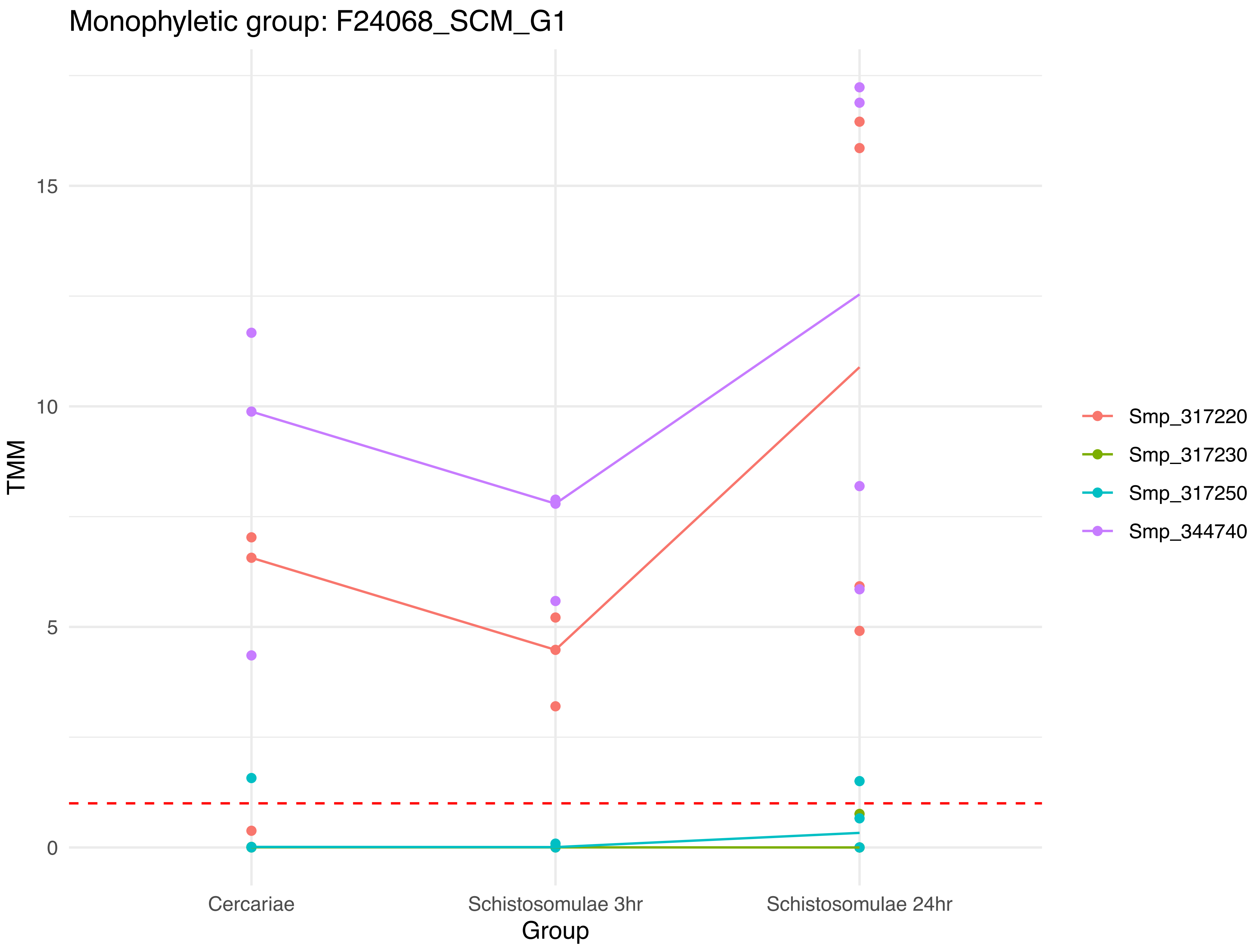
Monophyletic group: F24068_SCM_G1
TMM
15
10
5
0
Cercariae
Schistosomulae 3hr
Schistosomulae 24hr
Group
Smp_317220
Smp_317230
Smp_317250
Smp_344740

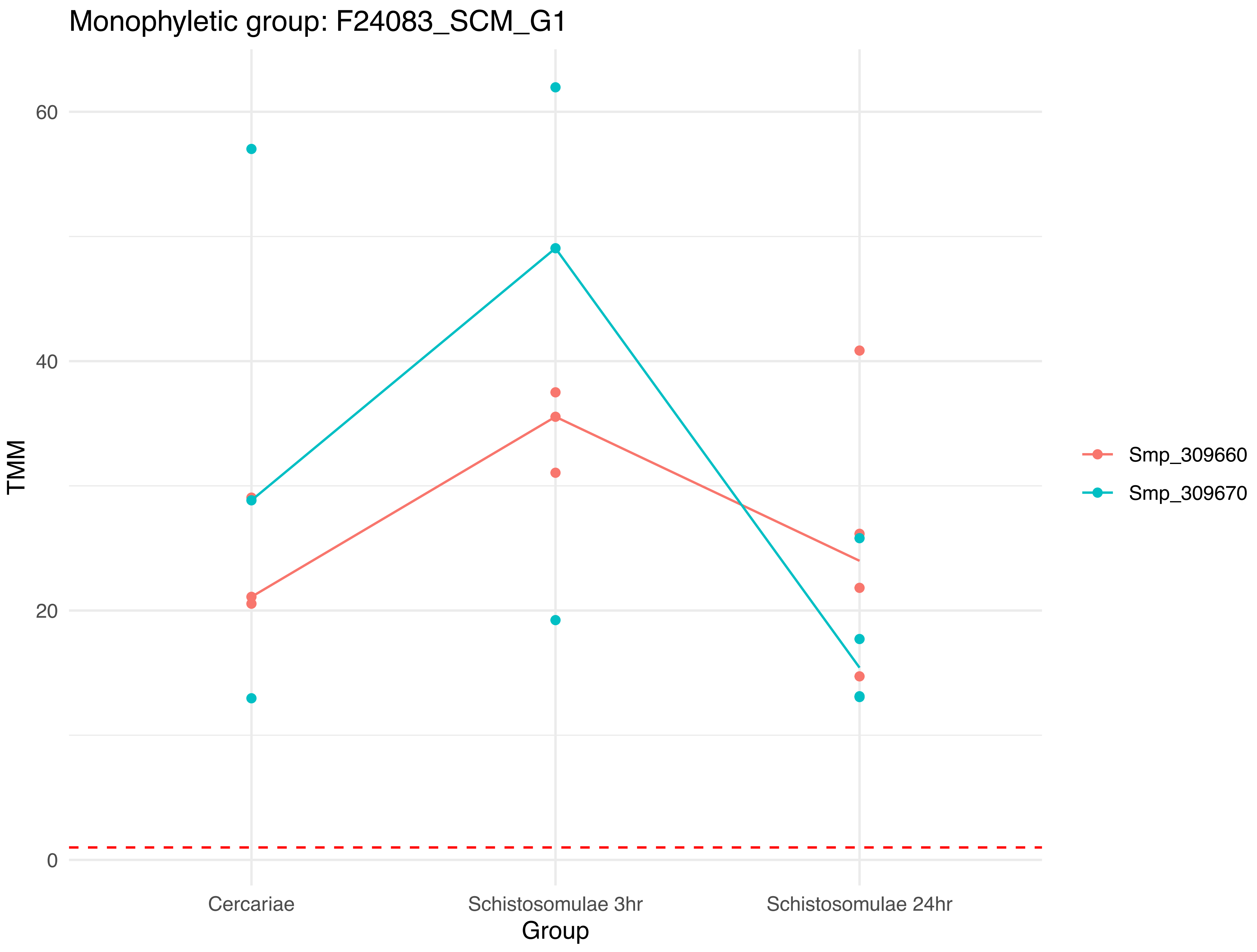

Monophyletic group: F24083_SCM_G1
TMM
60
40
20
0
Cercariae
Schistosomulae 3hr
Schistosomulae 24hr
Group
Smp_309660
Smp_309670

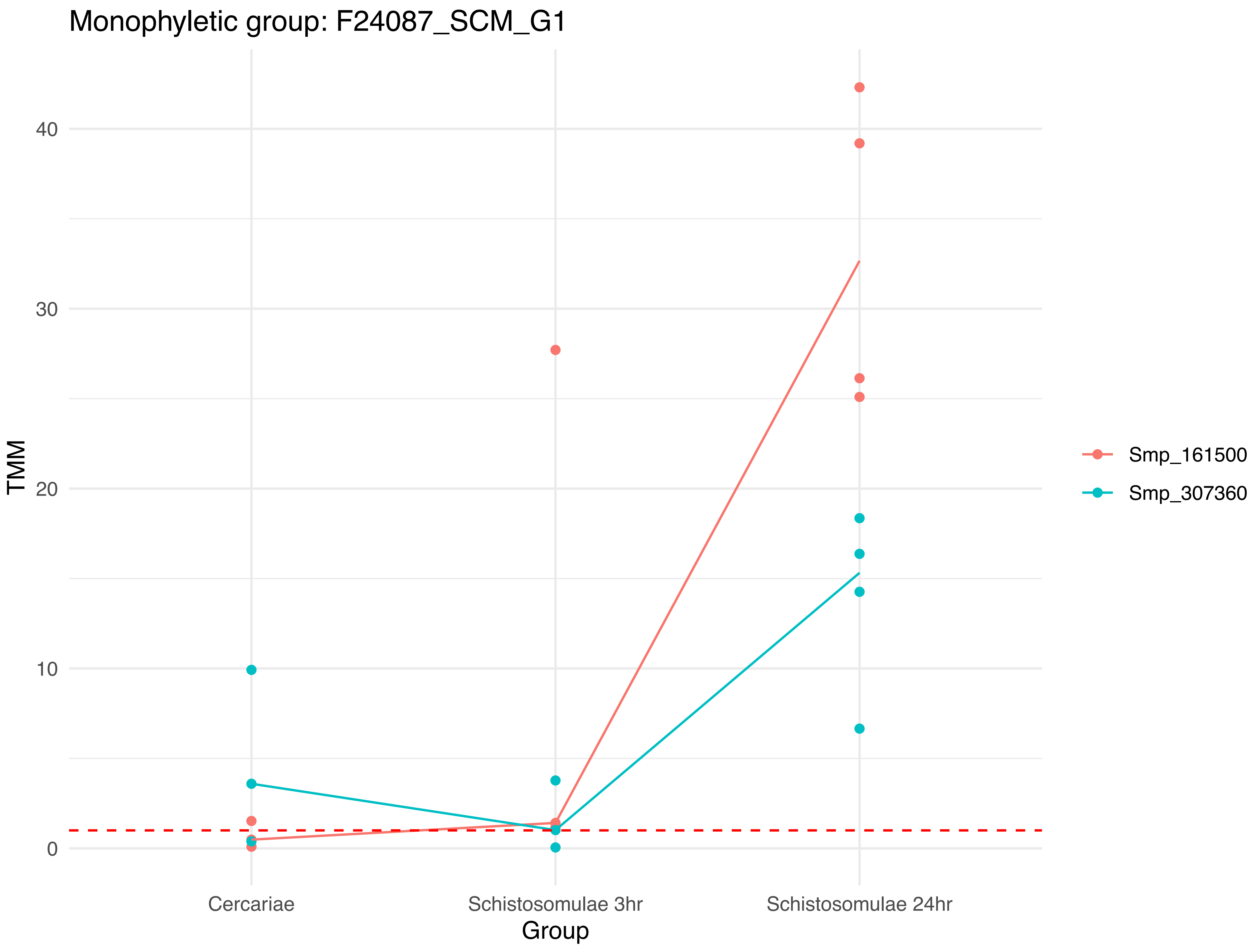

Monophyletic group: F24087_SCM_G1
TMM
40
30
20
10
0
Cercariae
Schistosomulae 3hr
Schistosomulae 24hr
Group
Smp_161500
Smp_307360

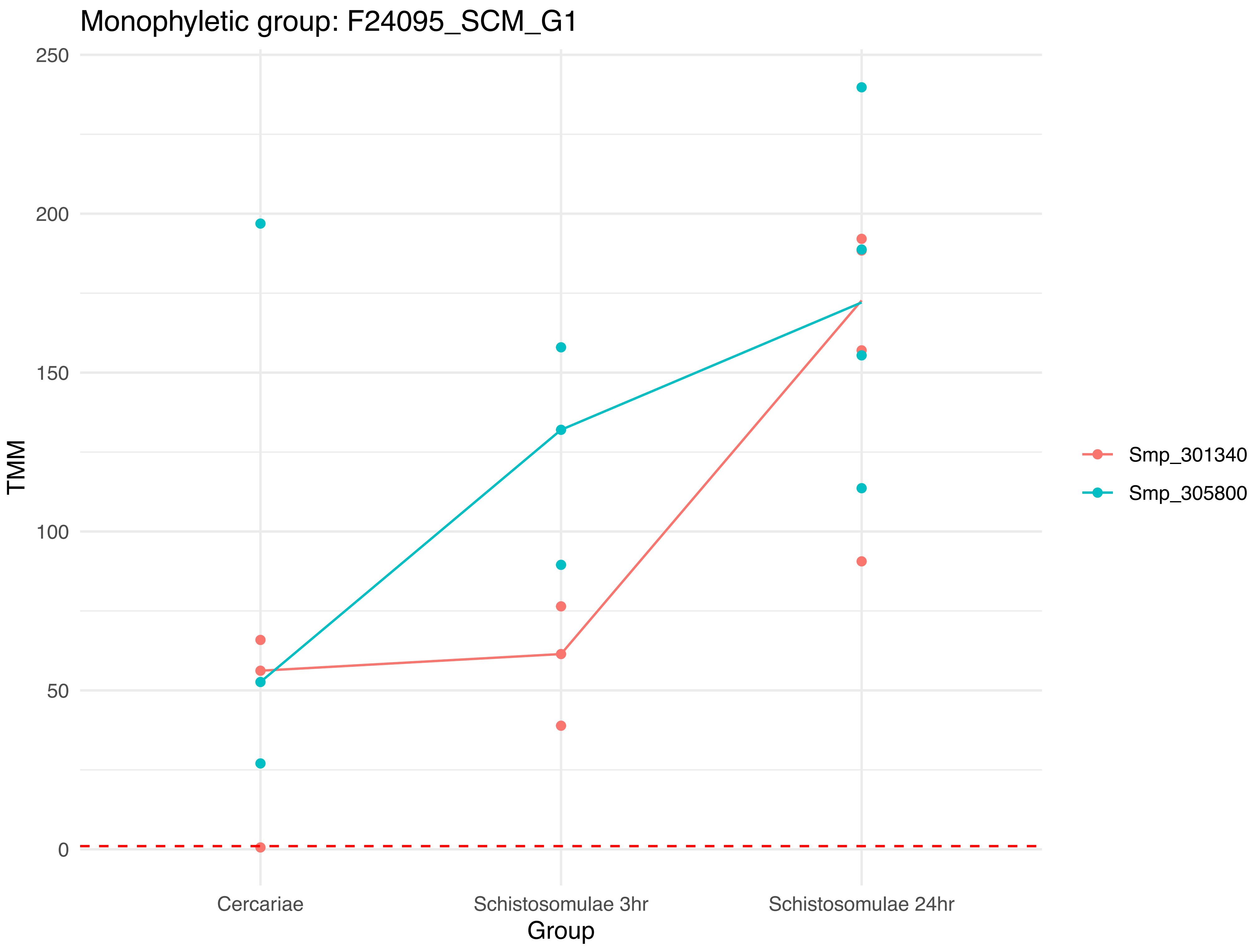

Monophyletic group: F24095_SCM_G1
TMM
250
200
150
100
50
0
Cercariae
Schistosomulae 3hr
Schistosomulae 24hr
Group
Smp_301340
Smp_305800

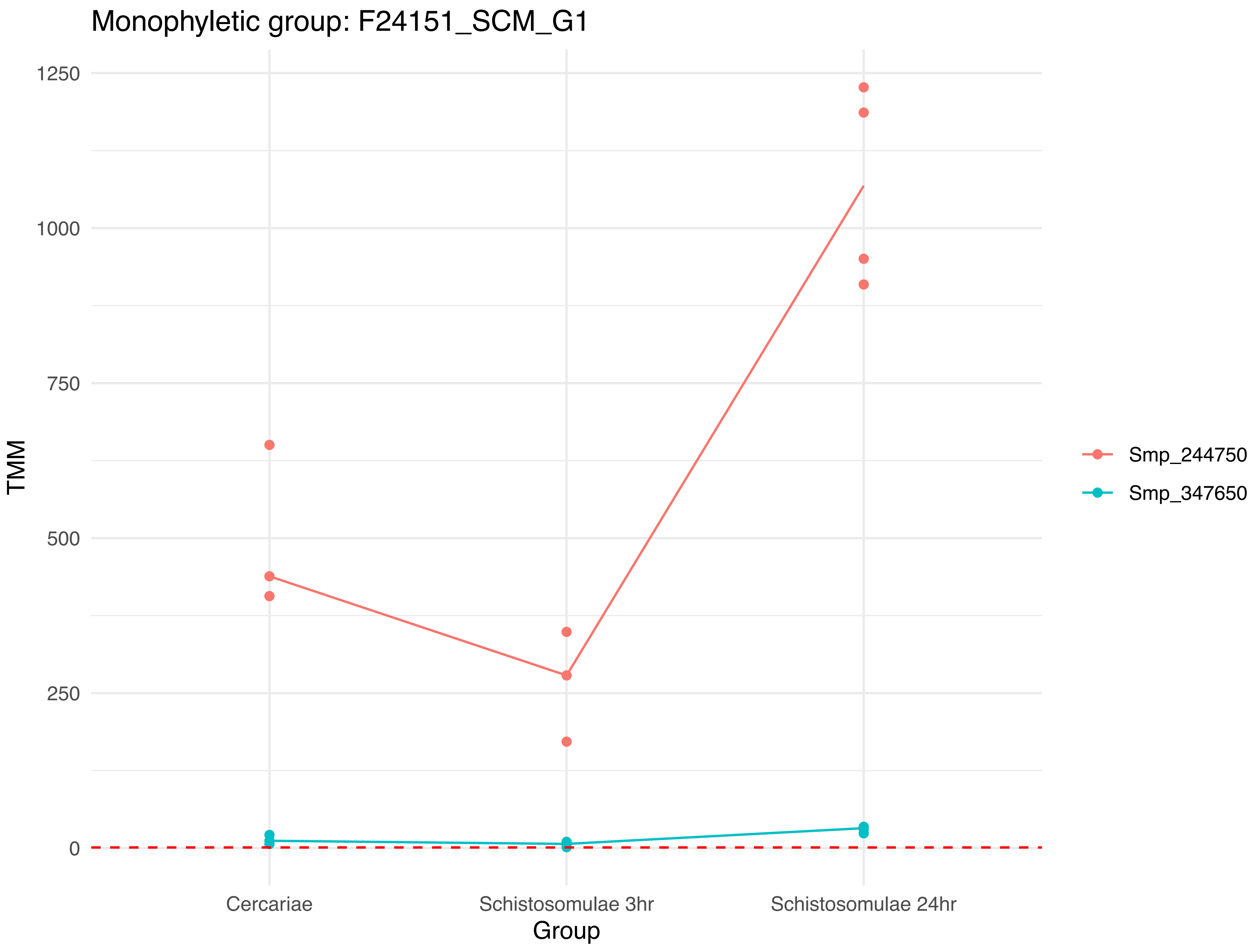

Monophyletic group: F24151_SCM_G1
1250
1000
750
500
250
0
TMM
Cercariae
Schistosomulae 3hr
Schistosomulae 24hr
Group
Smp_244750
Smp_347650

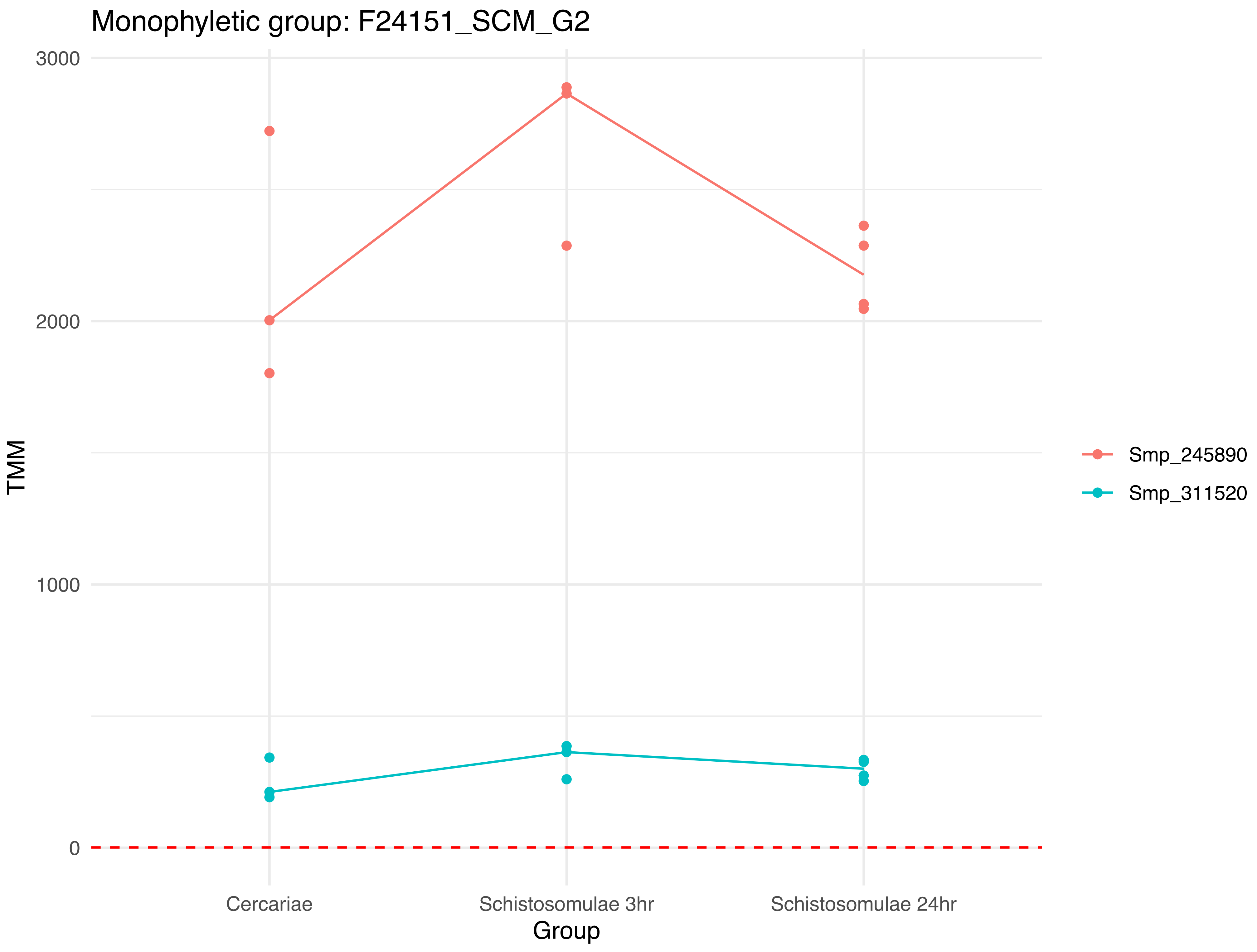
Monophyletic group: F24151_SCM_G2
3000
2000
1000
0
TMM
Cercariae
Schistosomulae 3hr
Schistosomulae 24hr
Group
Smp_245890
Smp_311520

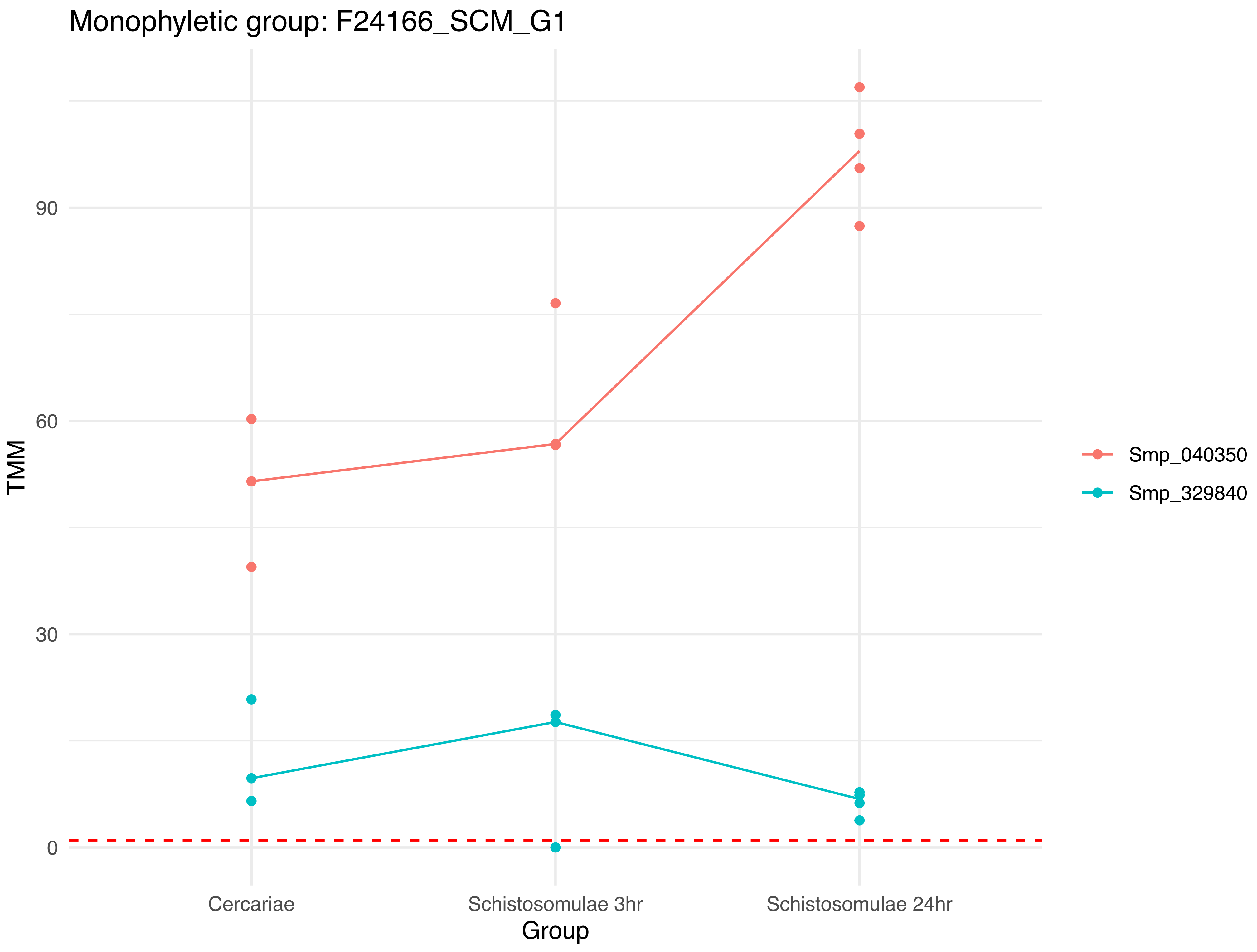

Monophyletic group: F24166_SCM_G1
TMM
90
60
30
0
Cercariae
Schistosomulae 3hr
Schistosomulae 24hr
Group
Smp_040350
Smp_329840

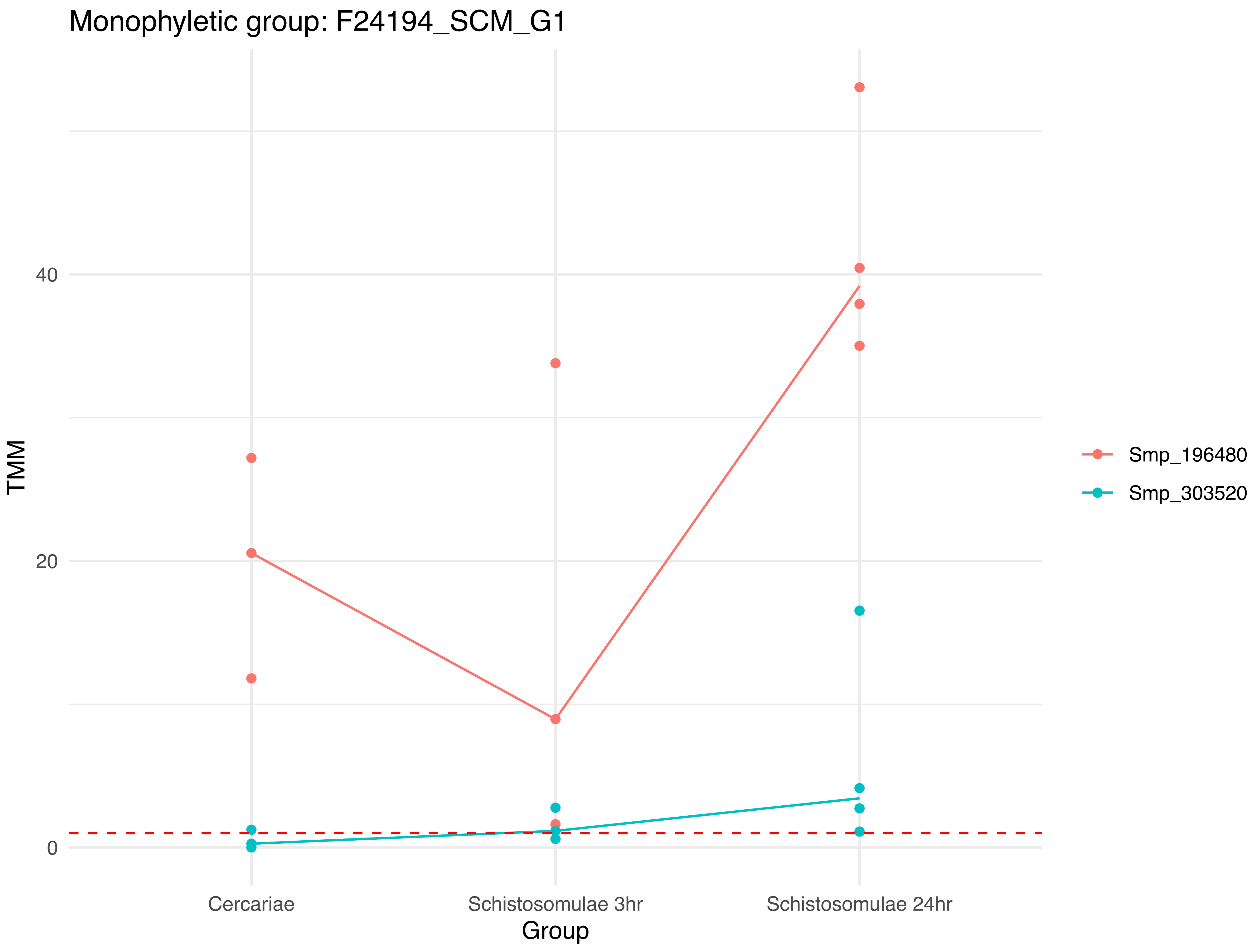

Monophyletic group: F24194_SCM_G1
TMM
40
20
0
Cercariae
Schistosomulae 3hr
Schistosomulae 24hr
Group
Smp_196480
Smp_303520

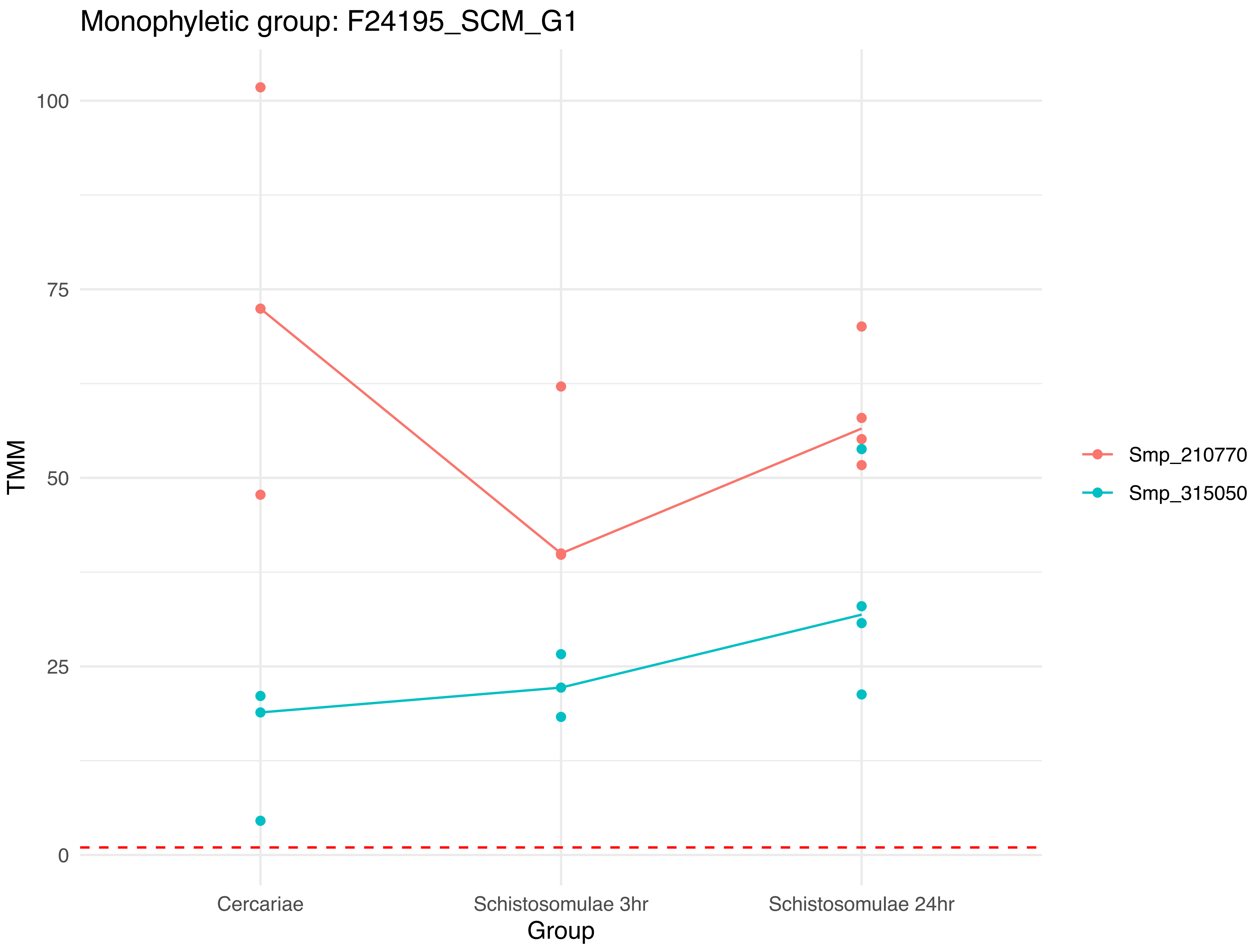

Monophyletic group: F24195_SCM_G1
100
75
50
25
0
TMM
Cercariae
Schistosomulae 3hr
Schistosomulae 24hr
Group
Smp_210770
Smp_315050

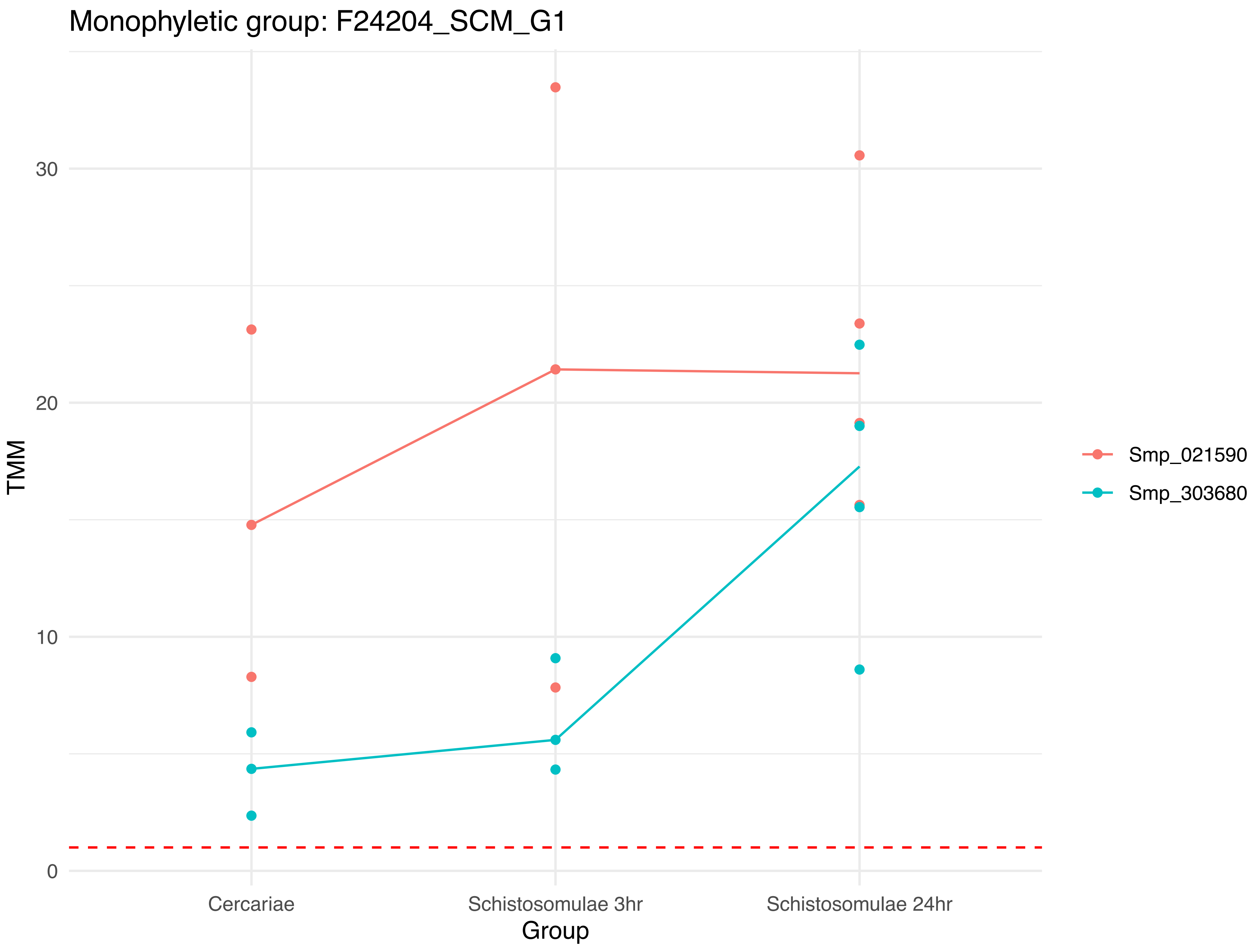

Monophyletic group: F24204_SCM_G1
TMM
30
20
10
0
Cercariae
Schistosomulae 3hr
Schistosomulae 24hr
Group
Smp_021590
Smp_303680

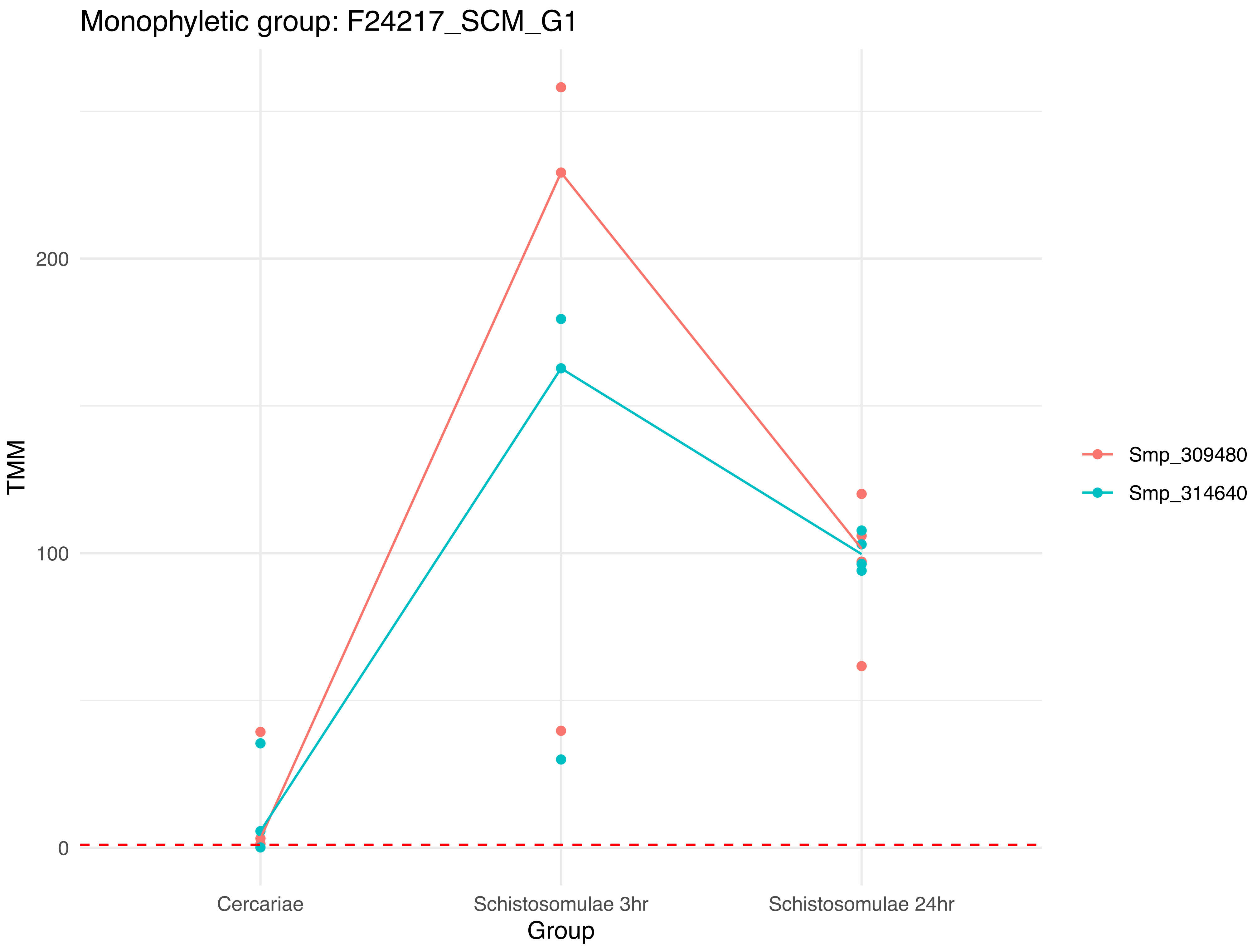

Monophyletic group: F24217_SCM_G1
TMM
200
100
0
Cercariae
Schistosomulae 3hr
Schistosomulae 24hr
Group
Smp_309480
Smp_314640

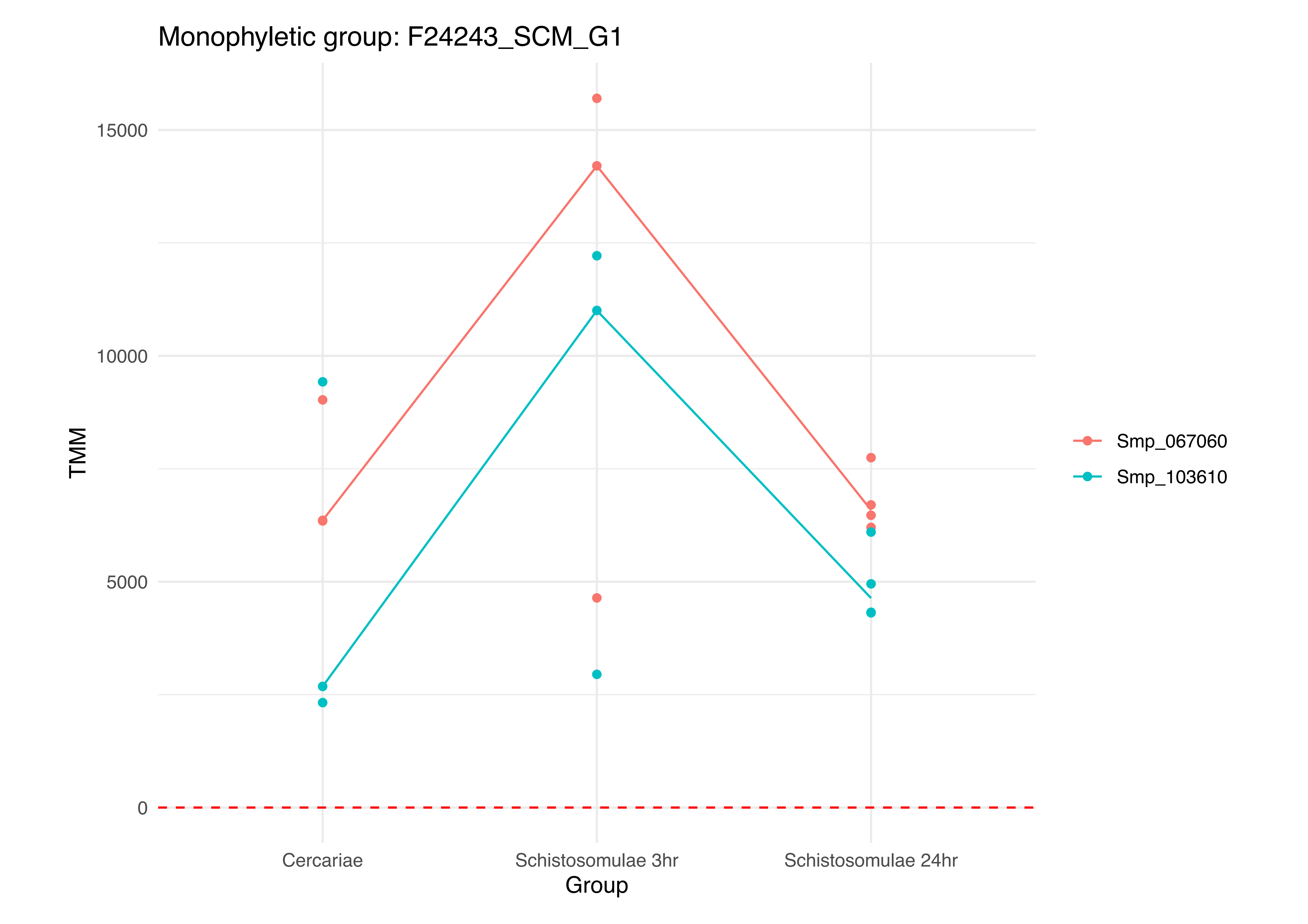
Monophyletic group: F24243_SCM_G1
15000
10000
5000
0
TMM
Cercariae
Schistosomulae 3hr
Schistosomulae 24hr
Group
Smp_067060
Smp_103610

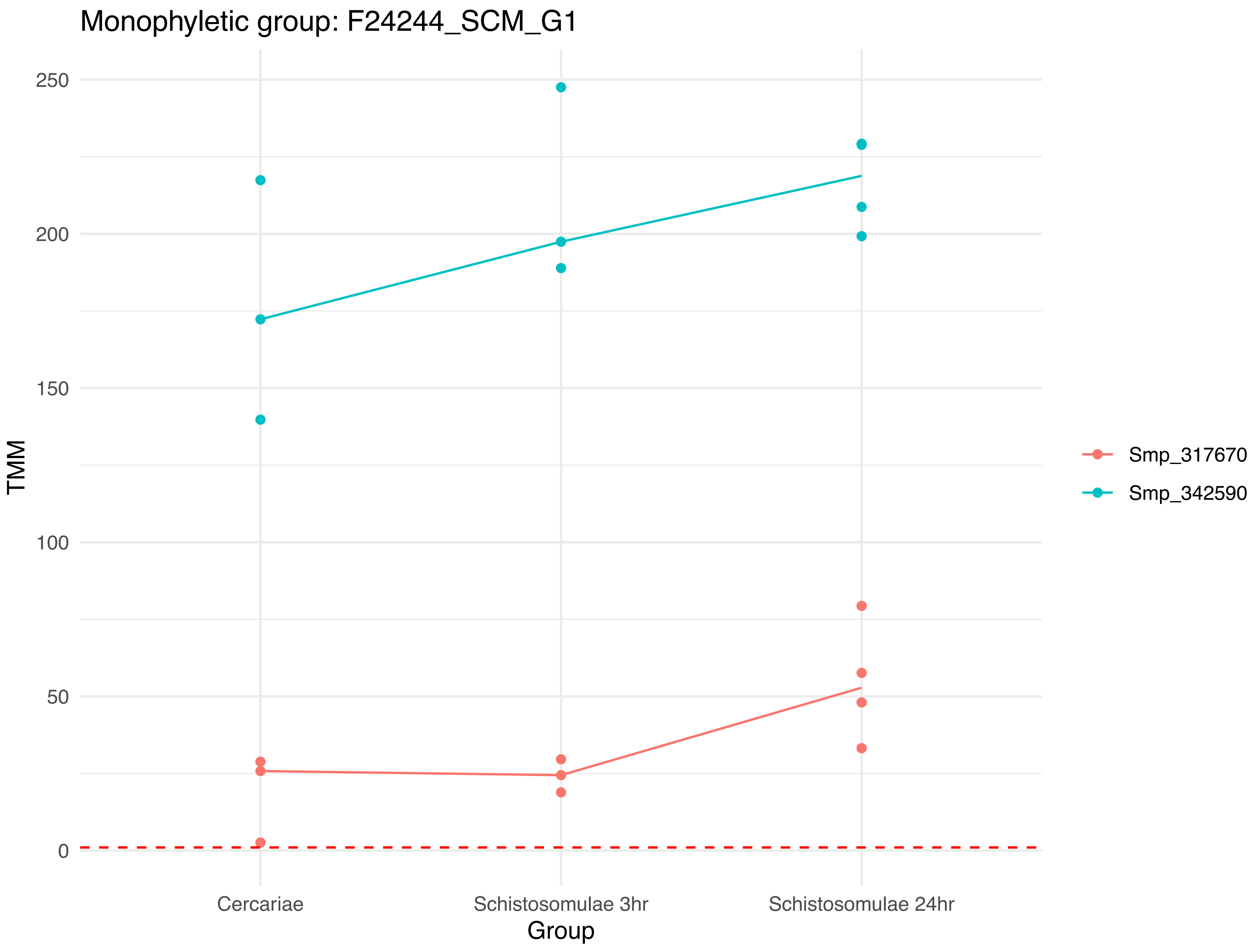

Monophyletic group: F24244_SCM_G1
TMM
250
200
150
100
50
0
Cercariae
Schistosomulae 3hr
Schistosomulae 24hr
Group
Smp_317670
Smp_342590

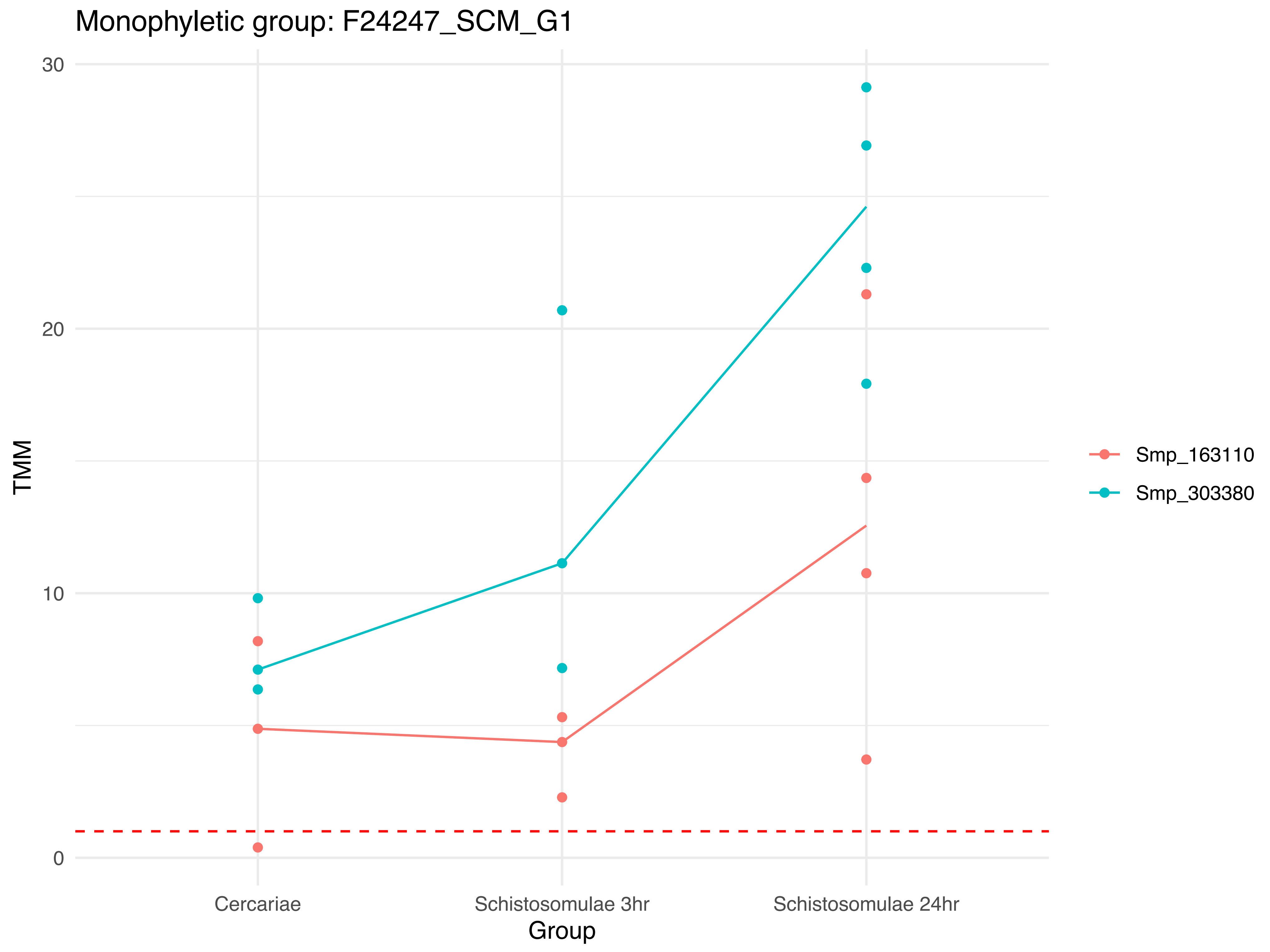
Monophyletic group: F24247_SCM_G1
TMM
30
20
10
0
Cercariae
Schistosomulae 3hr
Schistosomulae 24hr
Group
Smp_163110
Smp_303380

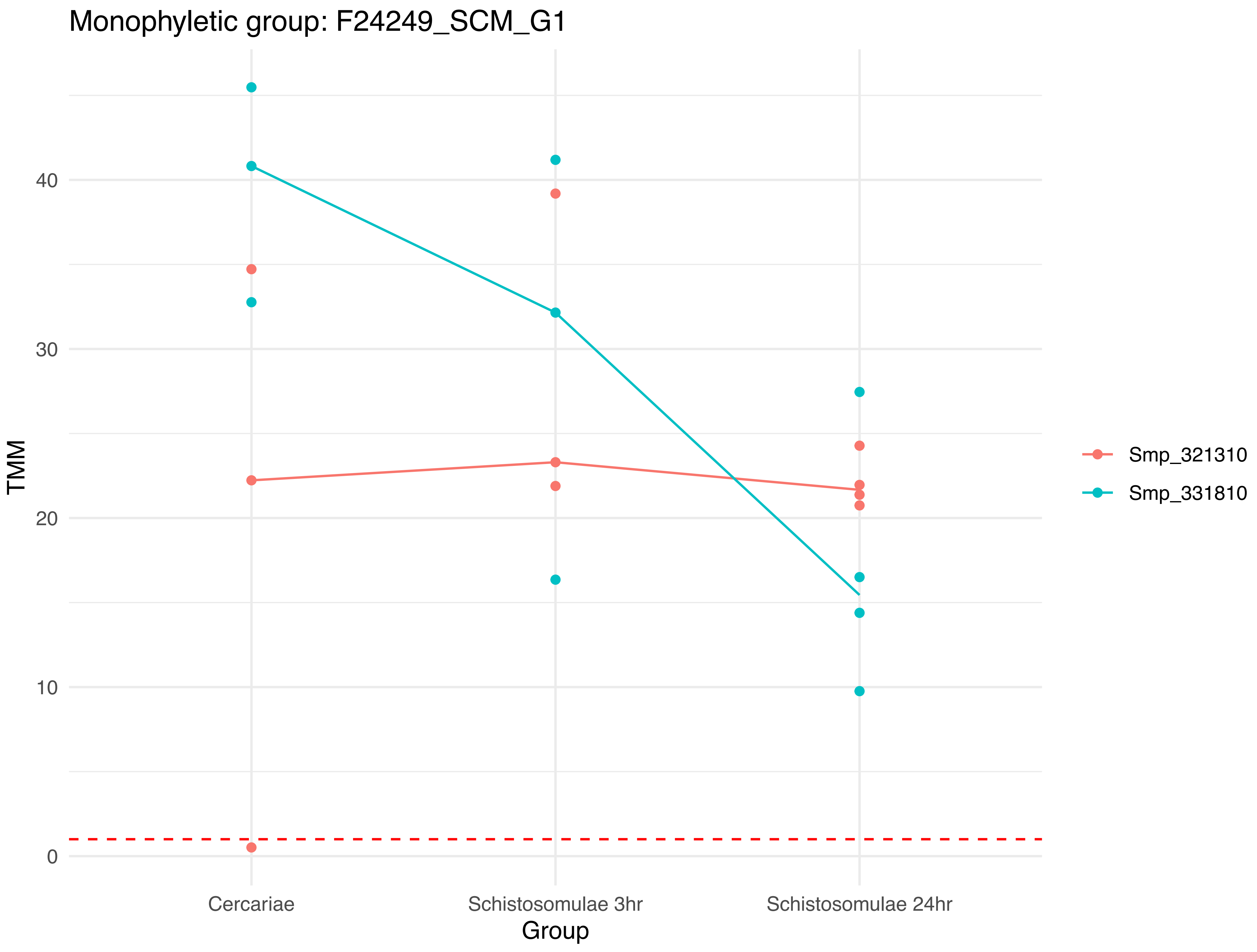
Monophyletic group: F24249_SCM_G1
TMM
40
30
20
10
0
Cercariae
Schistosomulae 3hr
Schistosomulae 24hr
Group
Smp_321310
Smp_331810

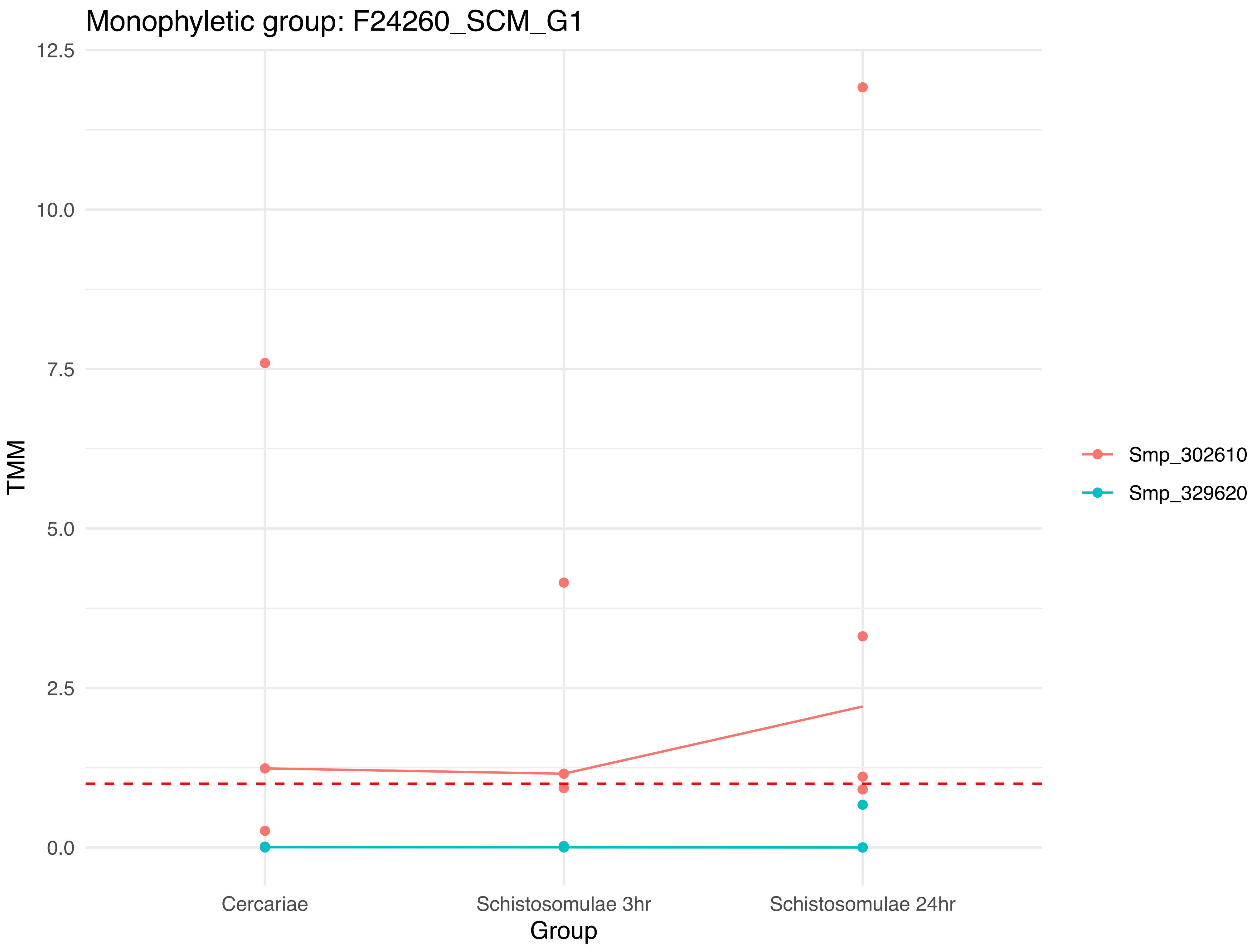

Monophyletic group: F24260_SCM_G1
12.5
10.0
7.5
5.0
2.5
0.0
TMM
Cercariae
Schistosomulae 3hr
Schistosomulae 24hr
Group
Smp_302610
Smp_329620

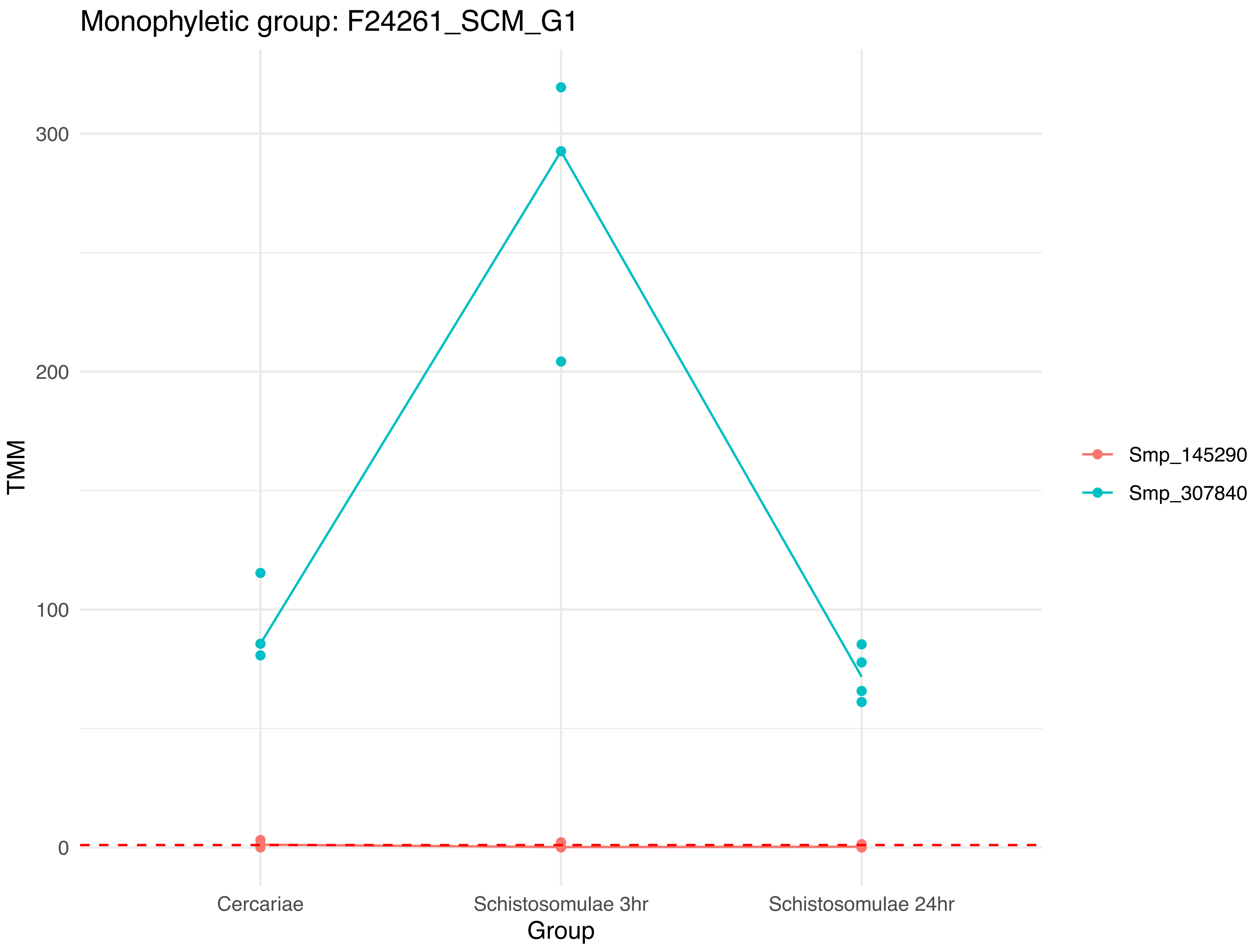

Monophyletic group: F24261_SCM_G1
300
200
100
0
TMM
Cercariae
Schistosomulae 3hr
Schistosomulae 24hr
Group
Smp_145290
Smp_307840

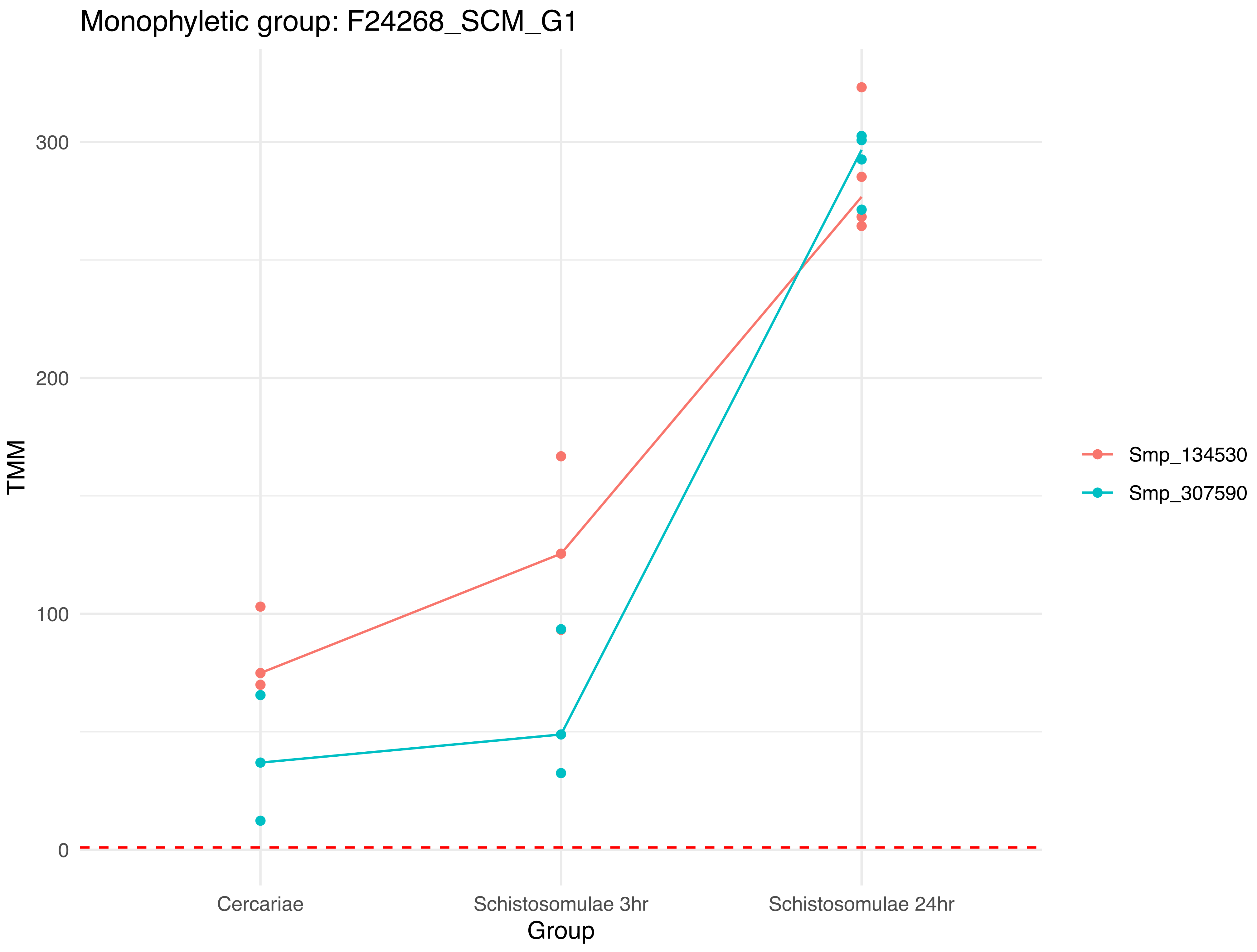

Monophyletic group: F24268_SCM_G1
300
200
100
0
TMM
Cercariae
Schistosomulae 3hr
Schistosomulae 24hr
Group
Smp_134530
Smp_307590

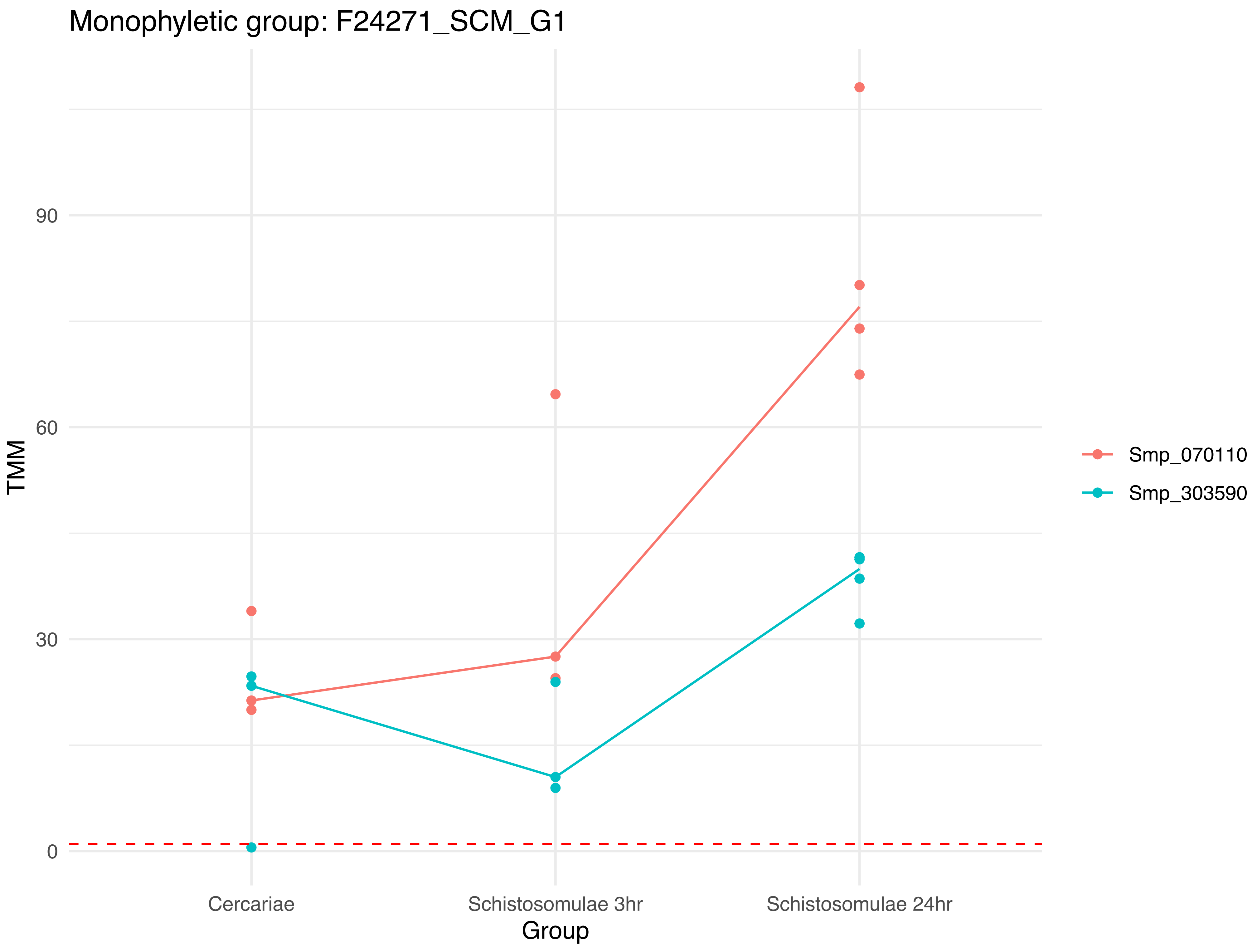

Monophyletic group: F24271_SCM_G1
TMM
90
60
30
0
Cercariae
Schistosomulae 3hr
Schistosomulae 24hr
Group
Smp_070110
Smp_303590

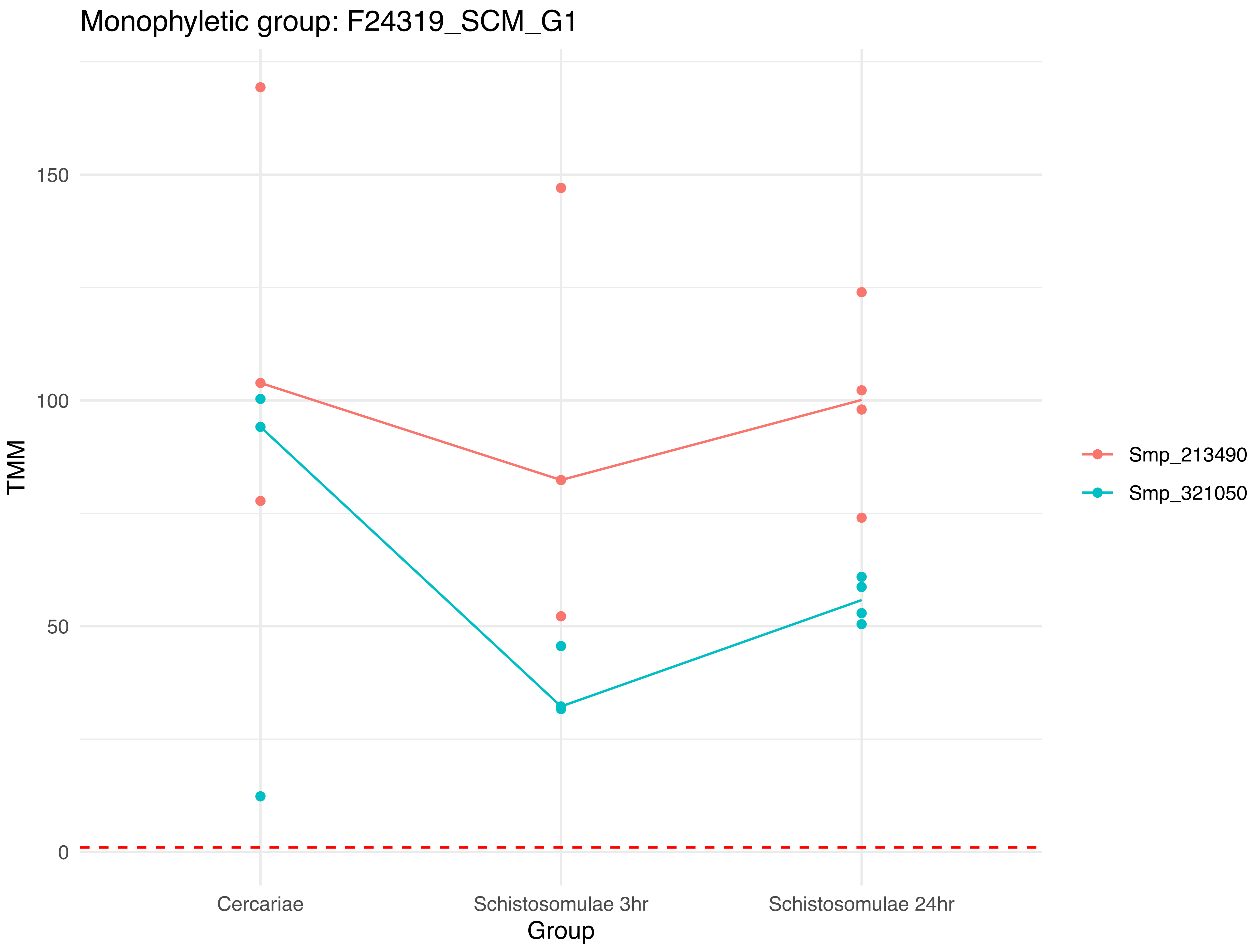

Monophyletic group: F24319_SCM_G1
150
100
50
0
TMM
Cercariae
Schistosomulae 3hr
Schistosomulae 24hr
Group
Smp_213490
Smp_321050

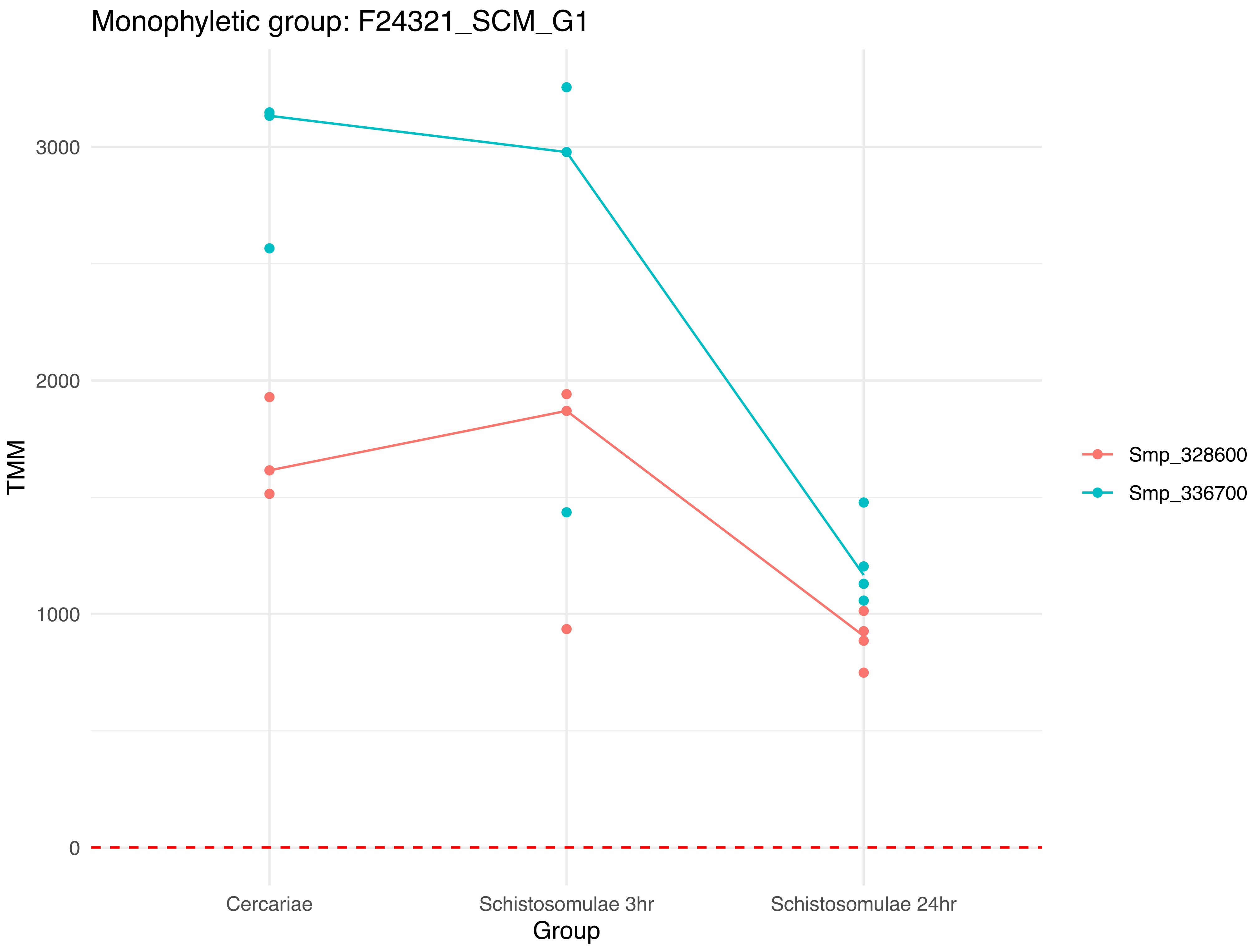

Monophyletic group: F24321_SCM_G1
3000
2000
1000
0
TMM
Cercariae
Schistosomulae 3hr
Schistosomulae 24hr
Group
Smp_328600
Smp_336700

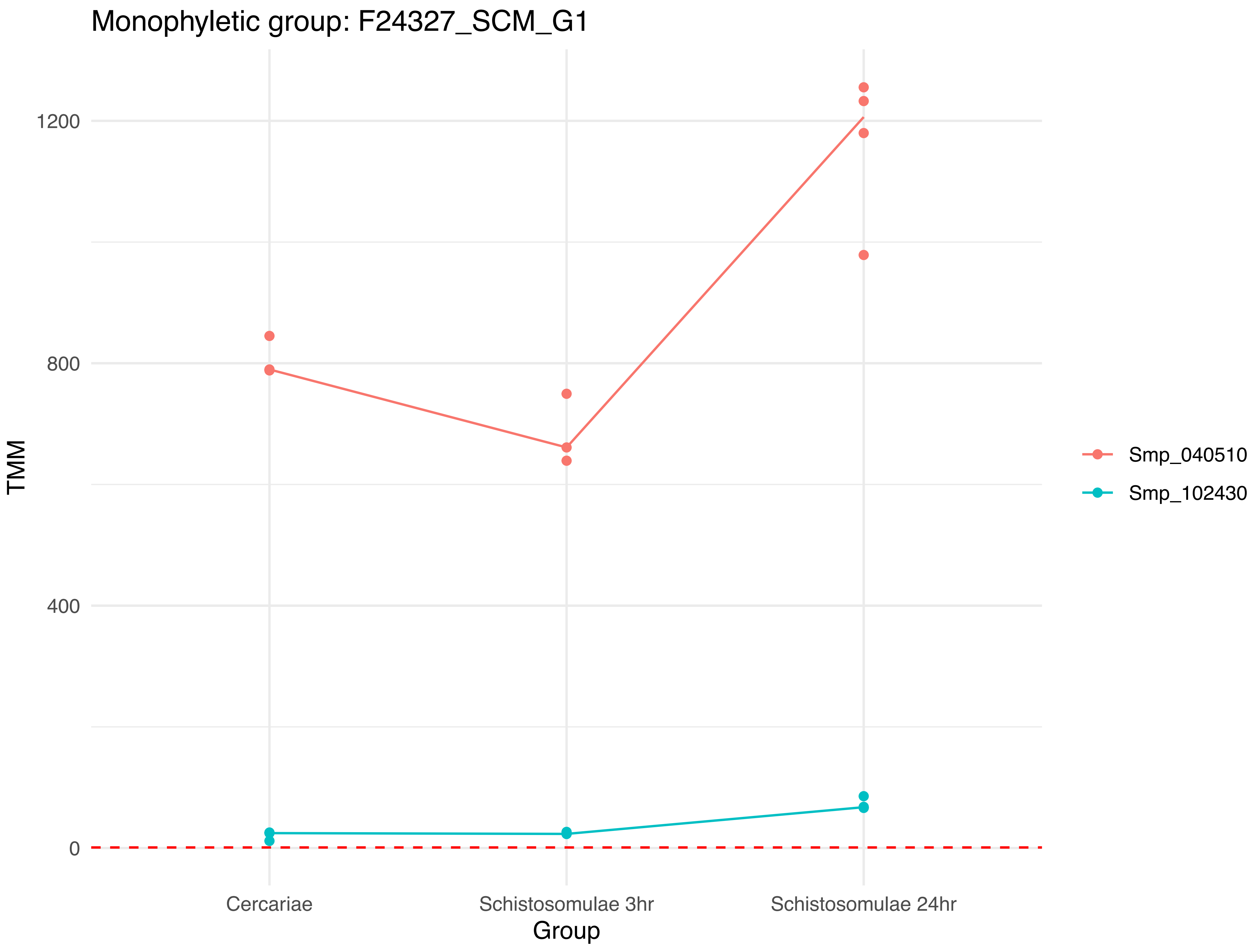
Monophyletic group: F24327_SCM_G1
1200
800
400
0
TMM
Cercariae
Schistosomulae 3hr
Schistosomulae 24hr
Group
Smp_040510
Smp_102430

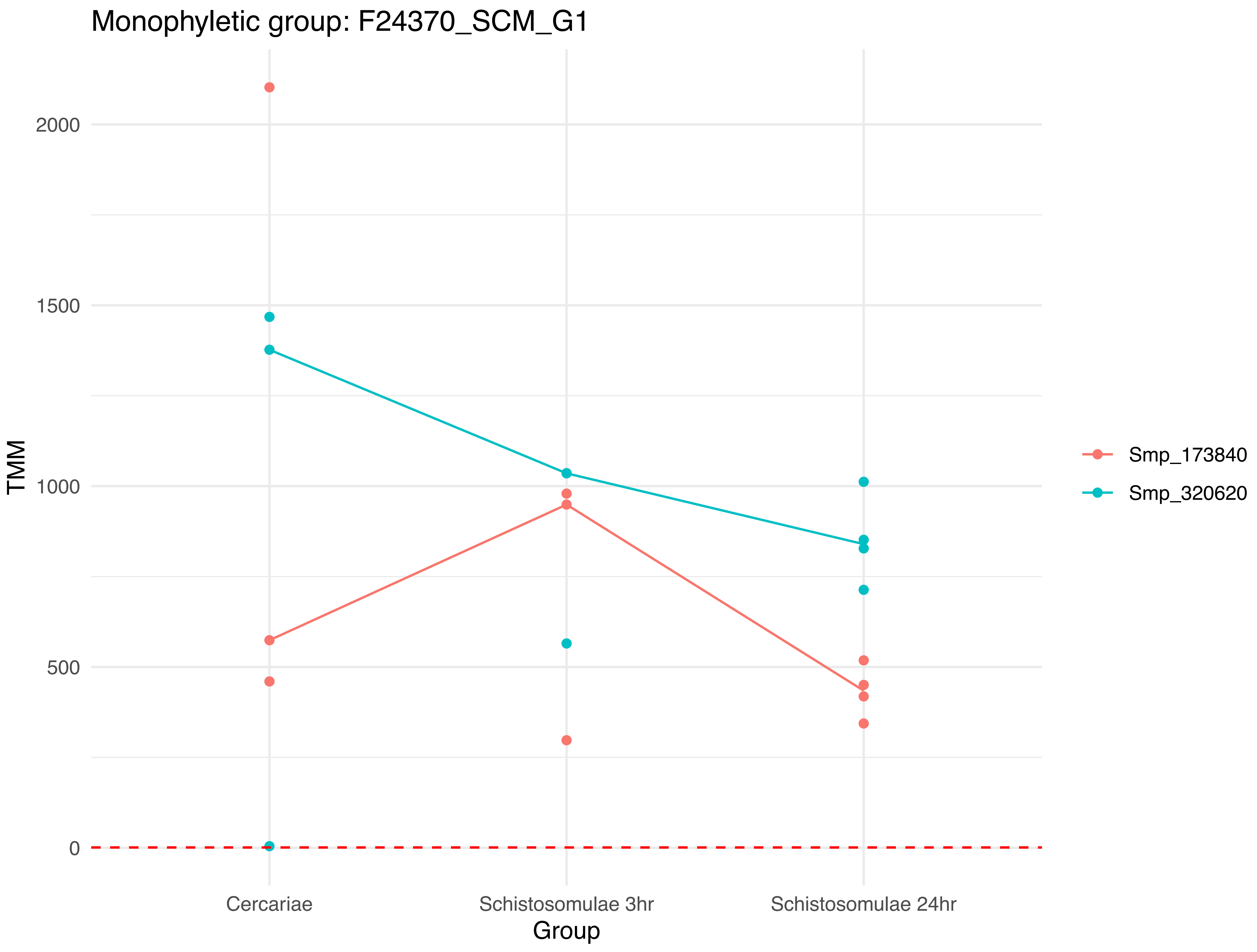
Monophyletic group: F24370_SCM_G1
TMM
2000
1500
1000
500
0
Cercariae
Schistosomulae 3hr
Schistosomulae 24hr
Group
Smp_173840
Smp_320620

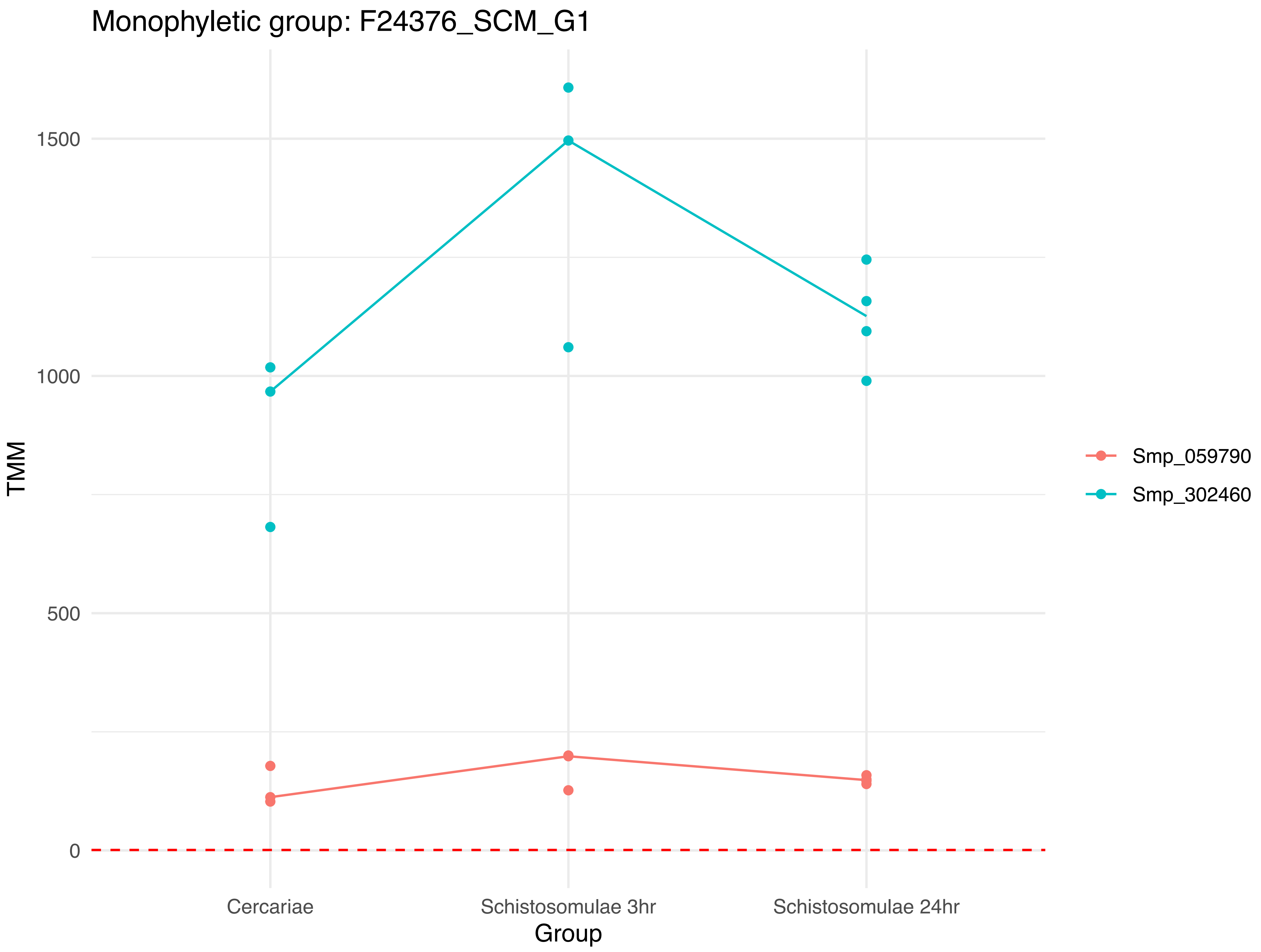

Monophyletic group: F24376_SCM_G1
1500
1000
500
0
TMM
Cercariae
Schistosomulae 3hr
Schistosomulae 24hr
Group
Smp_059790
Smp_302460

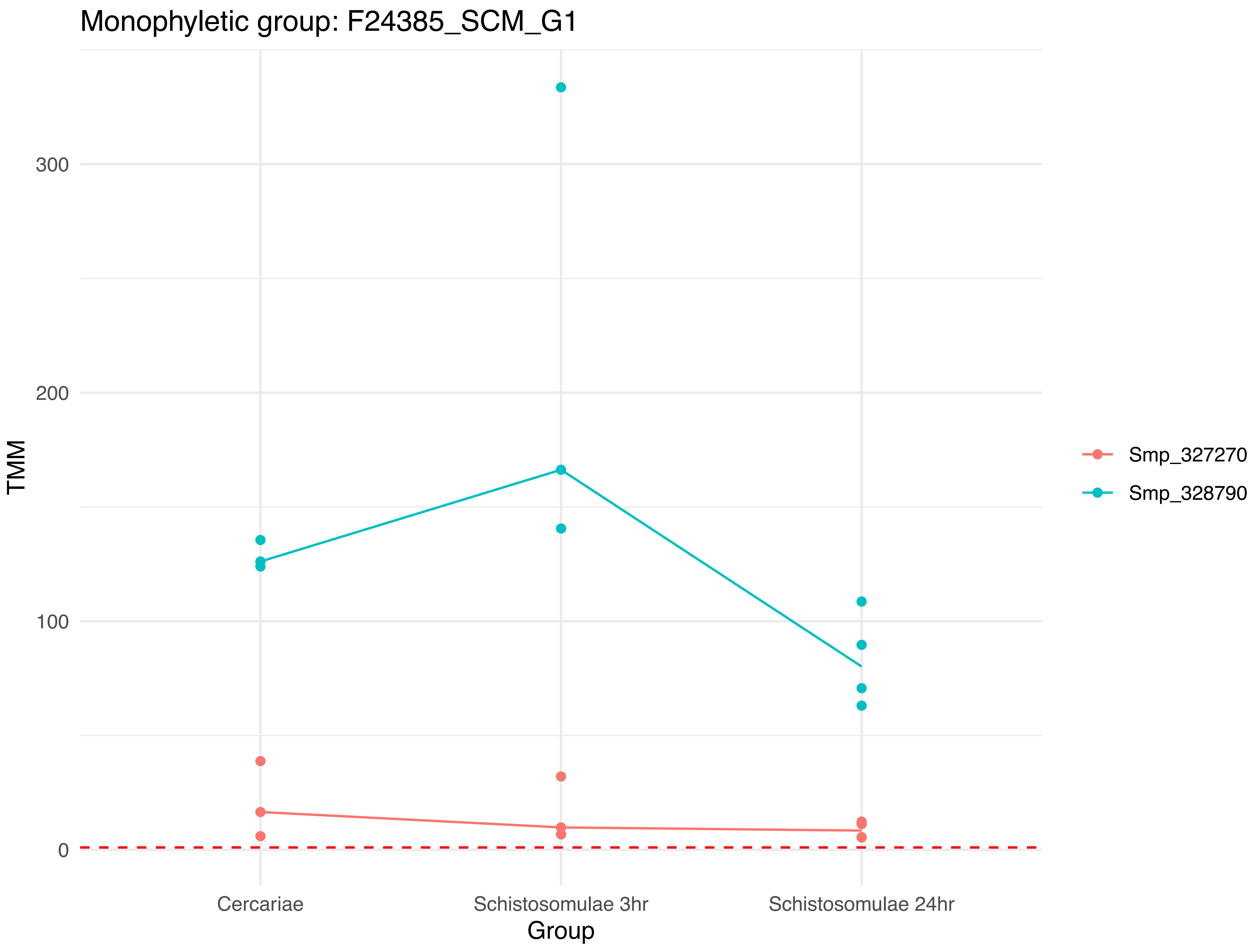

Monophyletic group: F24385_SCM_G1
TMM
300
200
100
0
Cercariae
Schistosomulae 3hr
Schistosomulae 24hr
Group
Smp_327270
Smp_328790

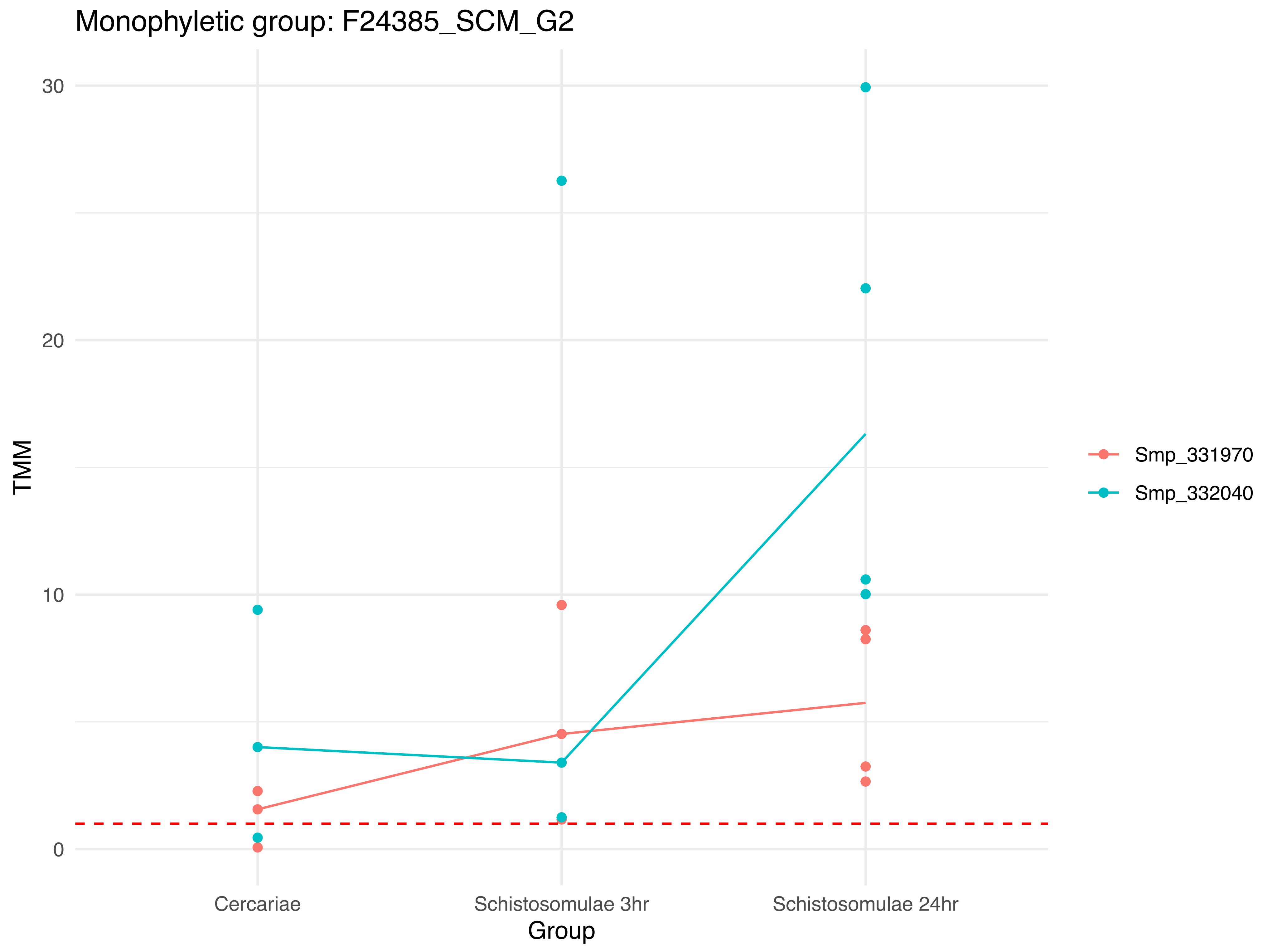

Monophyletic group: F24385_SCM_G2
30
20
10
0
TMM
Cercariae
Schistosomulae 3hr
Schistosomulae 24hr
Group
Smp_331970
Smp_332040

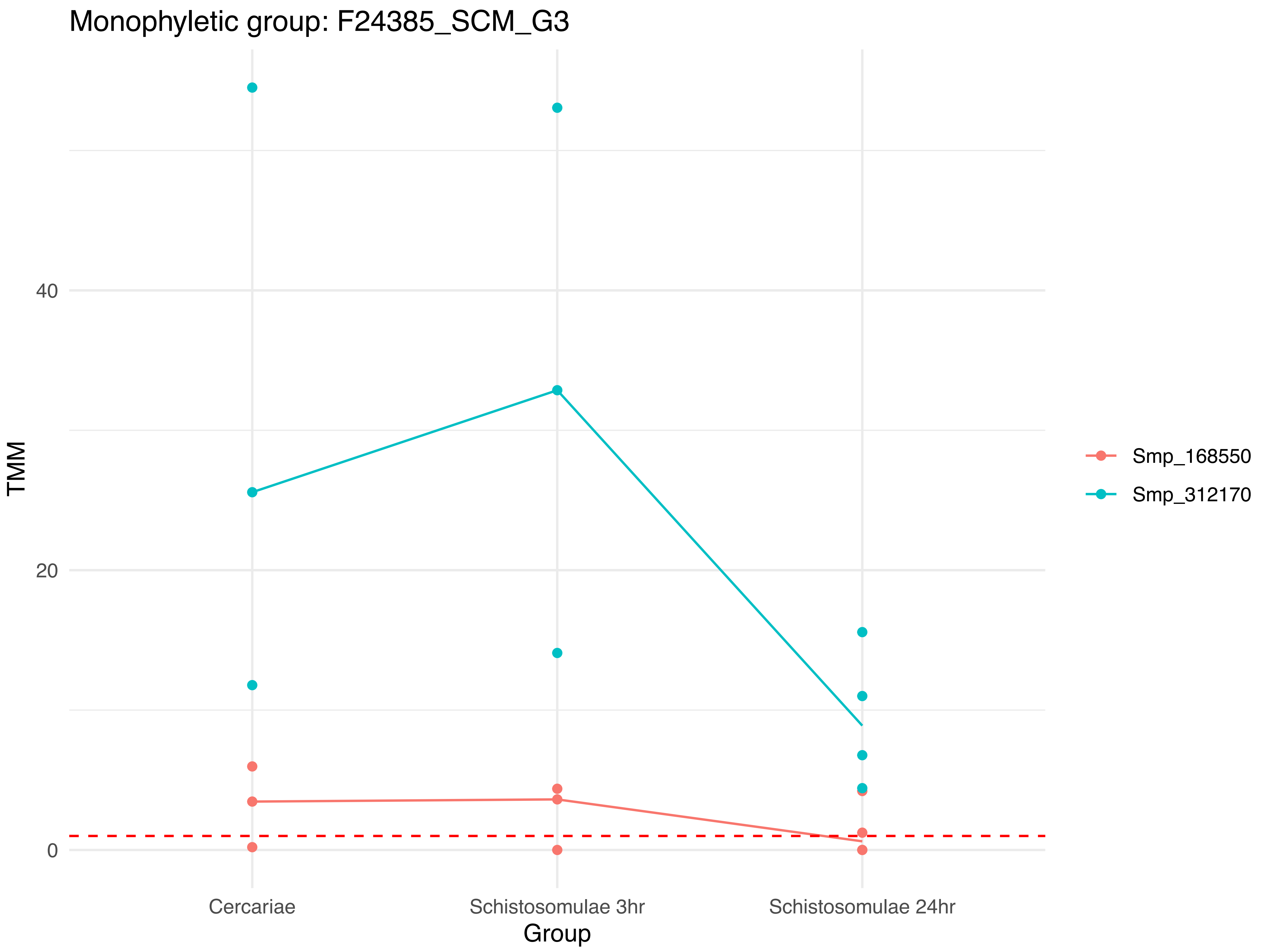

Monophyletic group: F24385_SCM_G3
TMM
40
20
0
Cercariae
Schistosomulae 3hr
Schistosomulae 24hr
Group
Smp_168550
Smp_312170

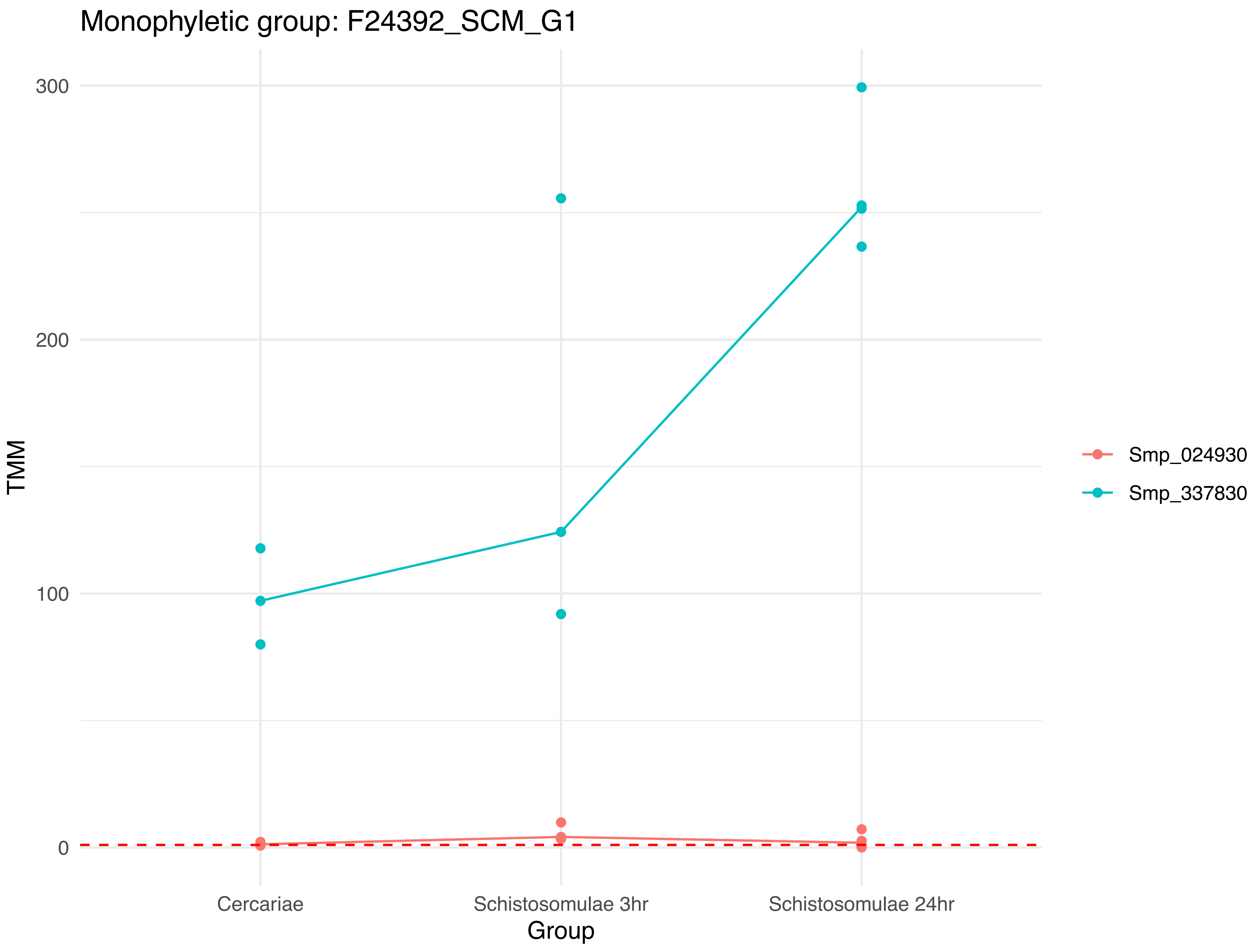

Monophyletic group: F24392_SCM_G1
300
200
100
0
TMM
Cercariae
Schistosomulae 3hr
Schistosomulae 24hr
Group
Smp_024930
Smp_337830

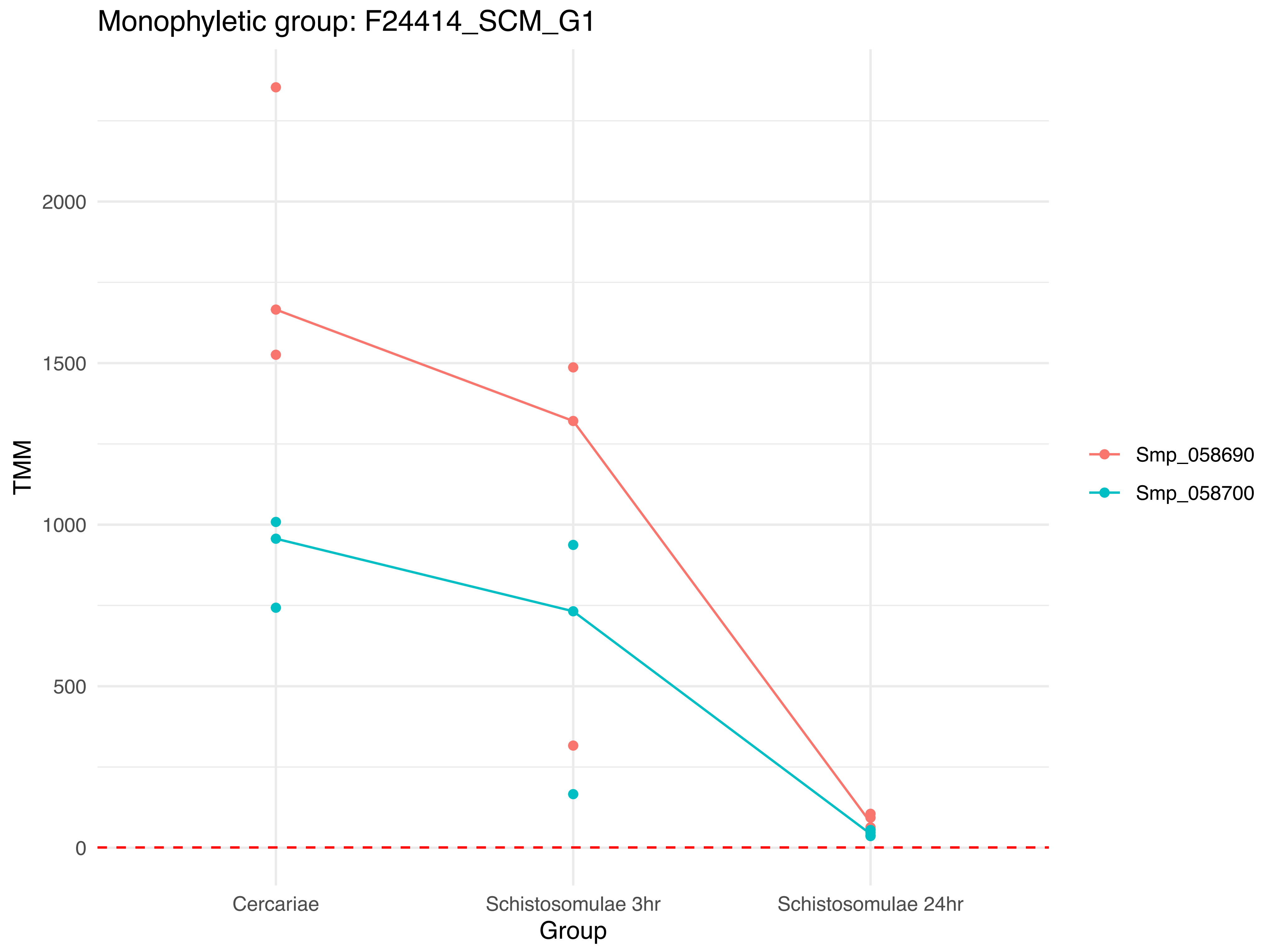
Monophyletic group: F24414_SCM_G1
TMM
2000
1500
1000
500
0
Cercariae
Schistosomulae 3hr
Schistosomulae 24hr
Group
Smp_058690
Smp_058700

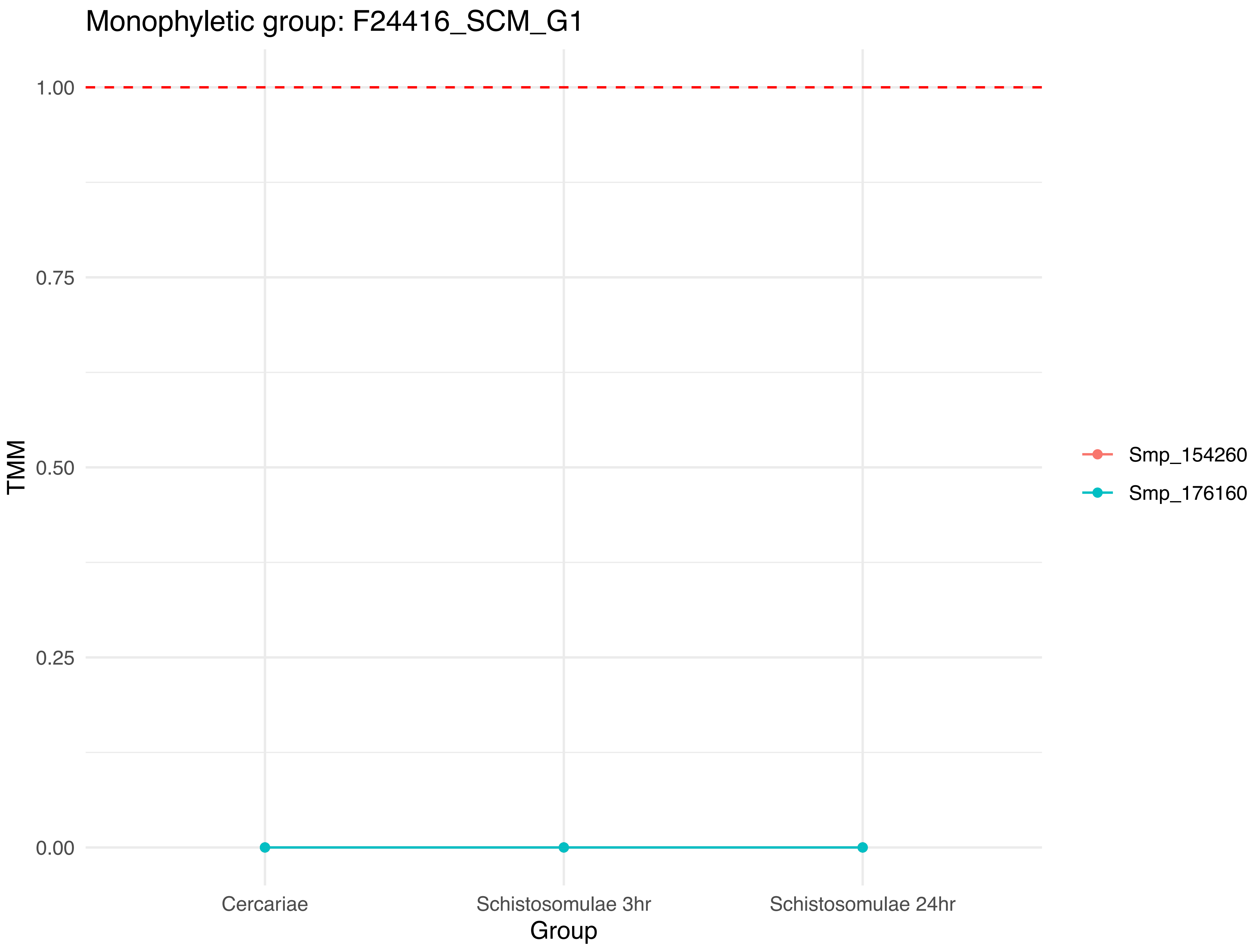

Monophyletic group: F24416_SCM_G1
1.00
0.75
0.50
0.25
0.00
TMM
Cercariae
Schistosomulae 3hr
Schistosomulae 24hr
Group
Smp_154260
Smp_176160

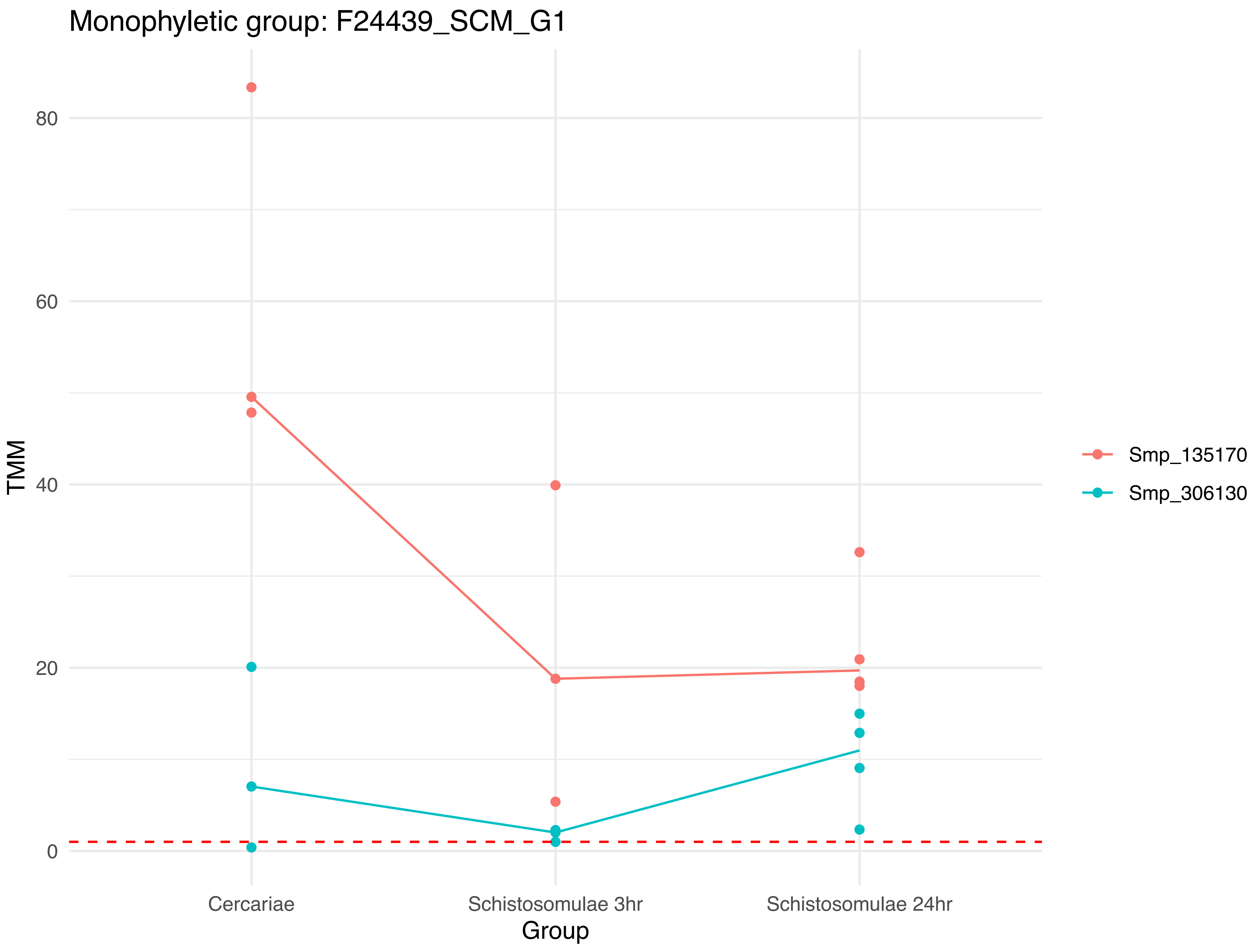
Monophyletic group: F24439_SCM_G1
TMM
80
60
40
20
0
Cercariae
Schistosomulae 3hr
Schistosomulae 24hr
Group
Smp_135170
Smp_306130

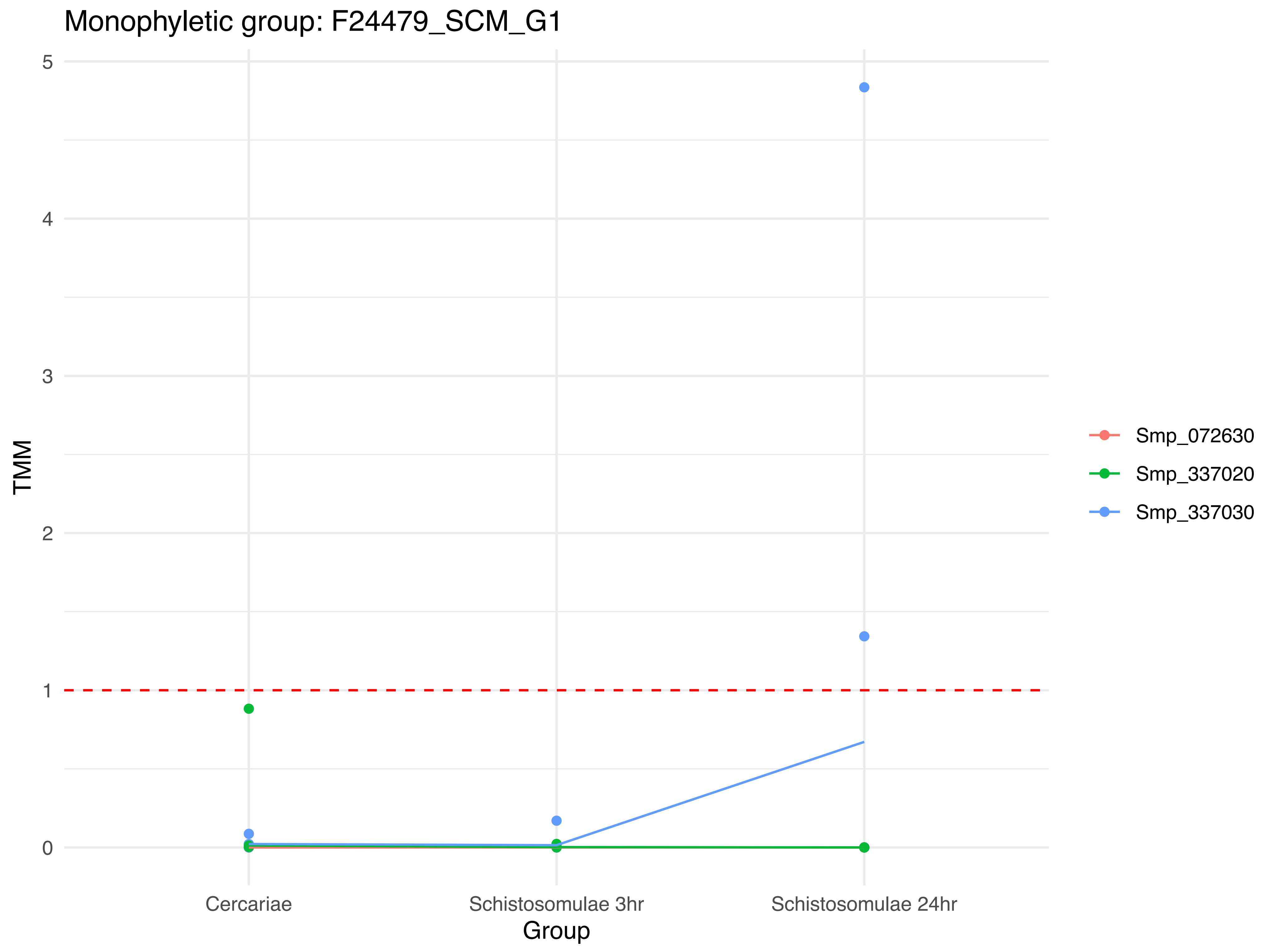

Monophyletic group: F24479_SCM_G1
TMM
5
4
3
2
1
0
Cercariae
Schistosomulae 3hr
Schistosomulae 24hr
Group
Smp_072630
Smp_337020
Smp_337030

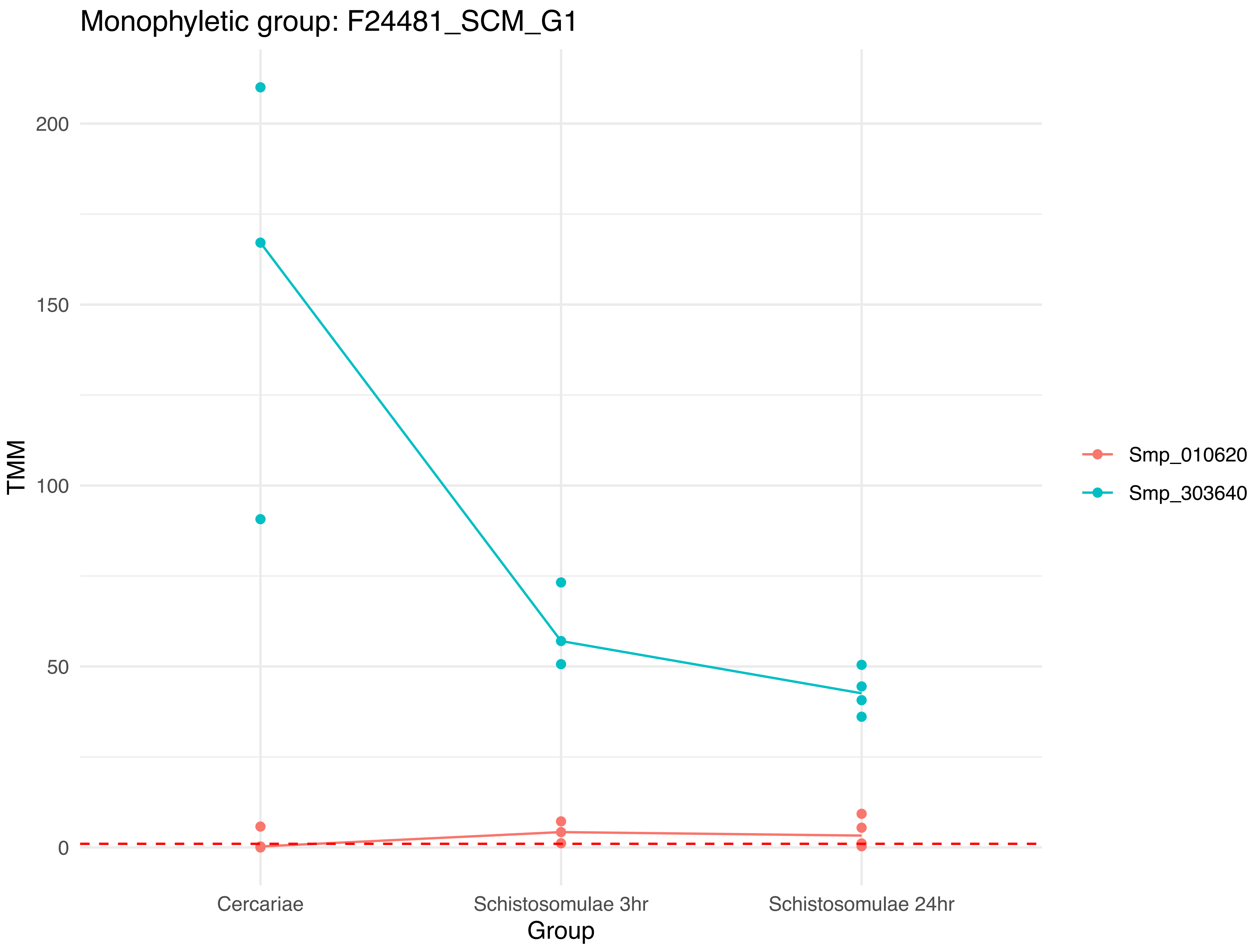

Monophyletic group: F24481_SCM_G1
200
150
100
50
0
TMM
Cercariae
Schistosomulae 3hr
Schistosomulae 24hr
Group
Smp_010620
Smp_303640

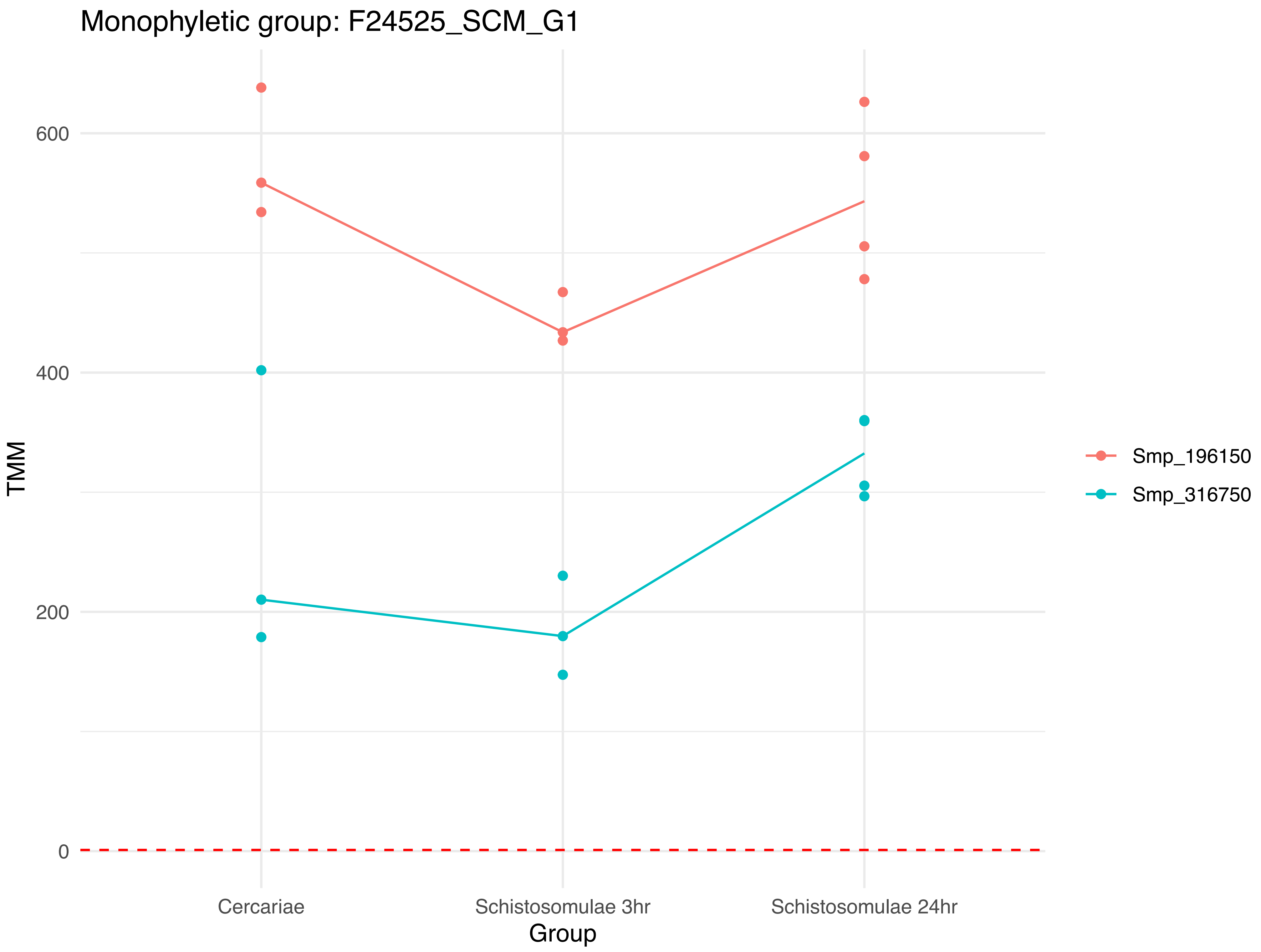
Monophyletic group: F24525_SCM_G1
600
400
200
0
TMM
Cercariae
Schistosomulae 3hr
Schistosomulae 24hr
Group
Smp_196150
Smp_316750

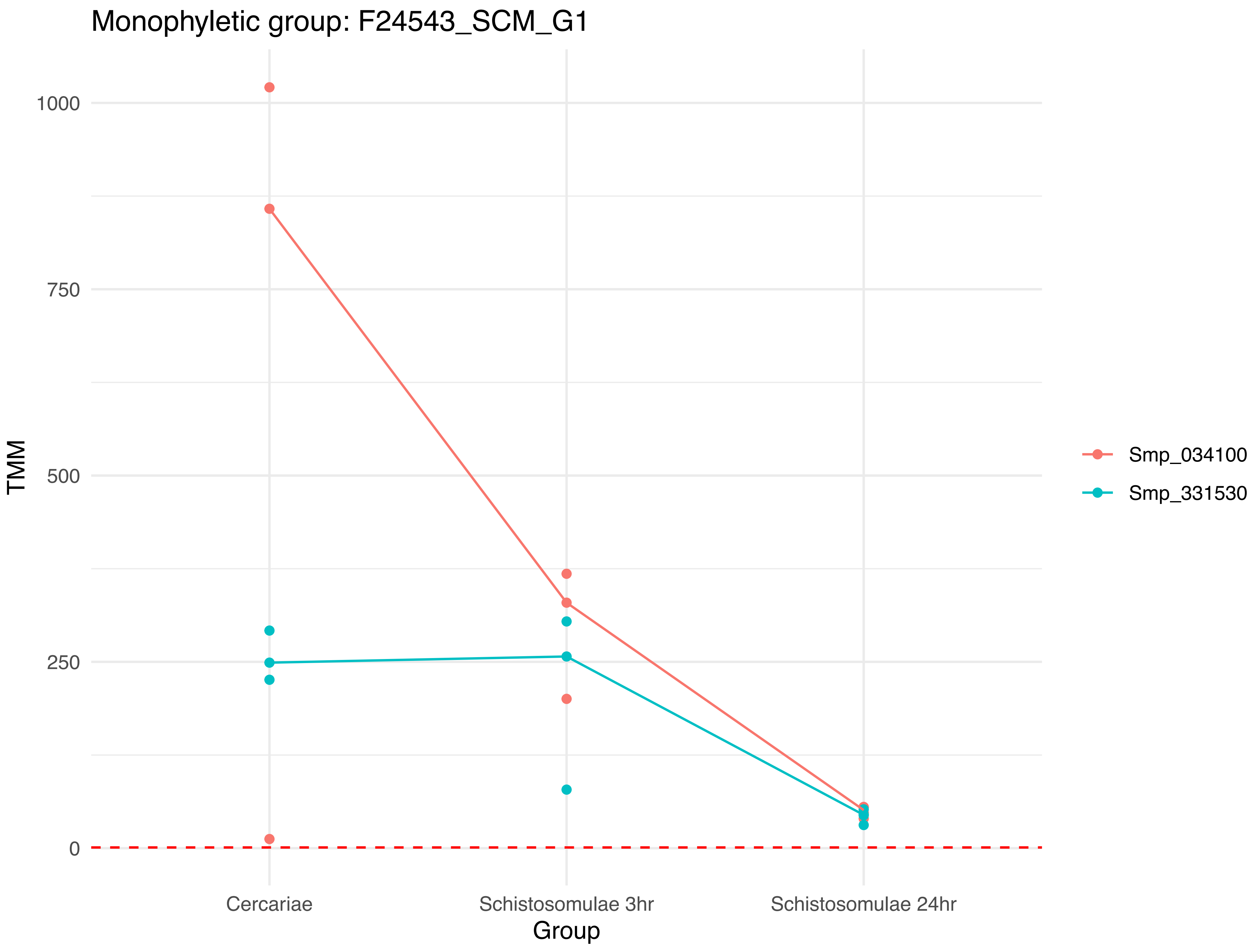

Monophyletic group: F24543_SCM_G1
1000
750
500
250
0
TMM
Cercariae
Schistosomulae 3hr
Schistosomulae 24hr
Group
Smp_034100
Smp_331530

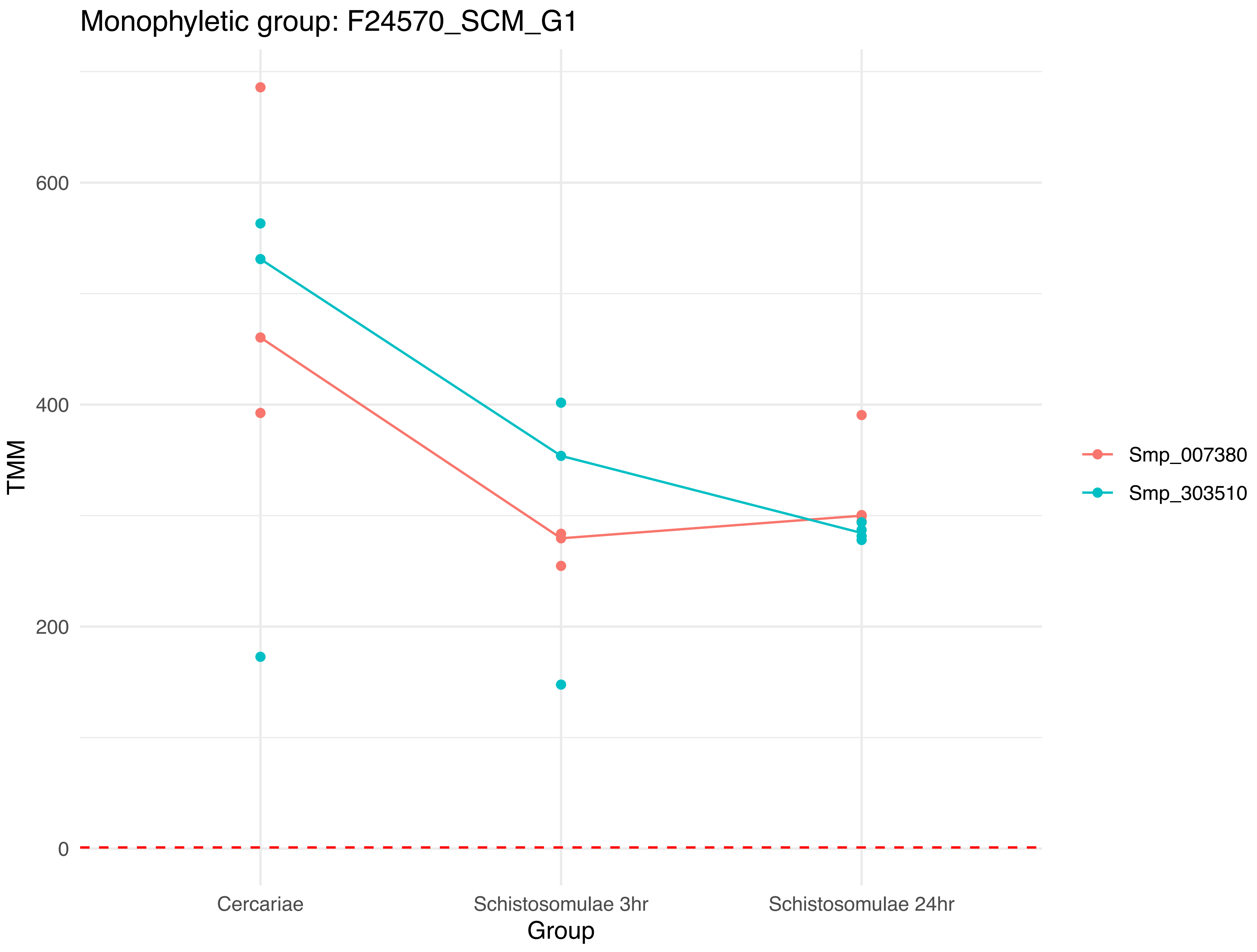
Monophyletic group: F24570_SCM_G1
TMM
600
400
200
0
Cercariae
Schistosomulae 3hr
Schistosomulae 24hr
Group
Smp_007380
Smp_303510

Monophyletic group: F24574_SCM_G1

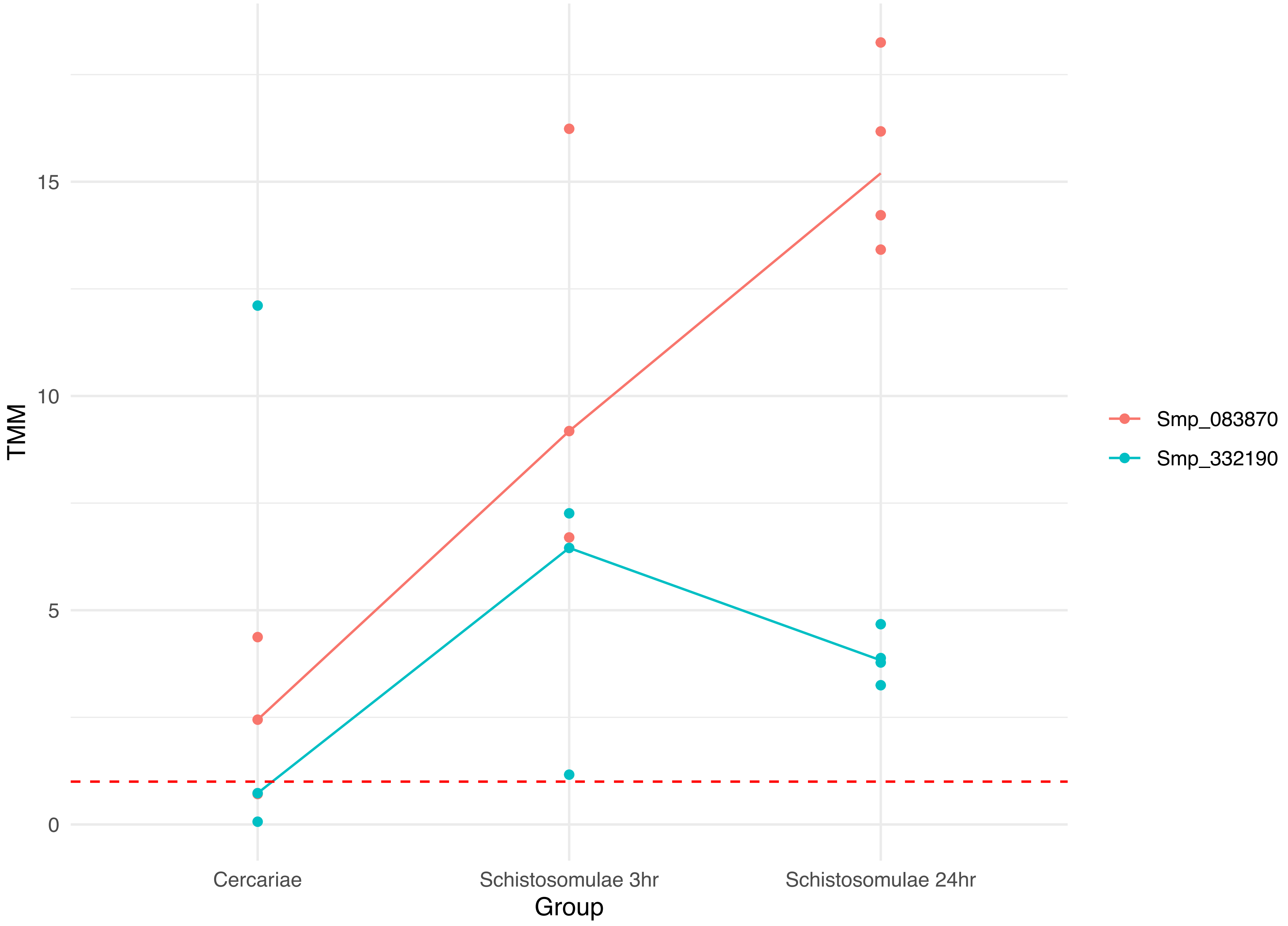

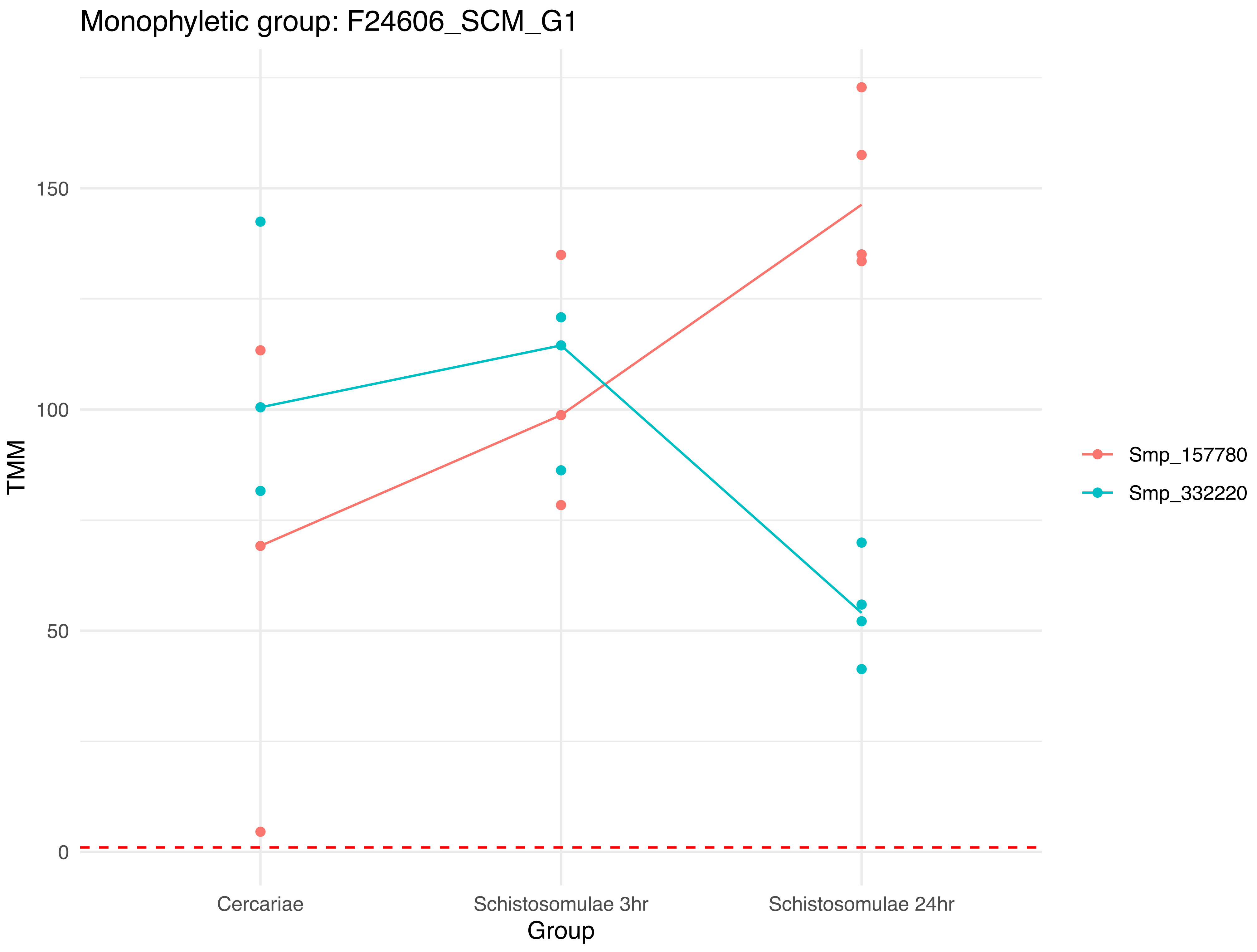

Monophyletic group: F24606_SCM_G1
TMM
150
100
50
0
Cercariae
Schistosomulae 3hr
Schistosomulae 24hr
Group
Smp_157780
Smp_332220

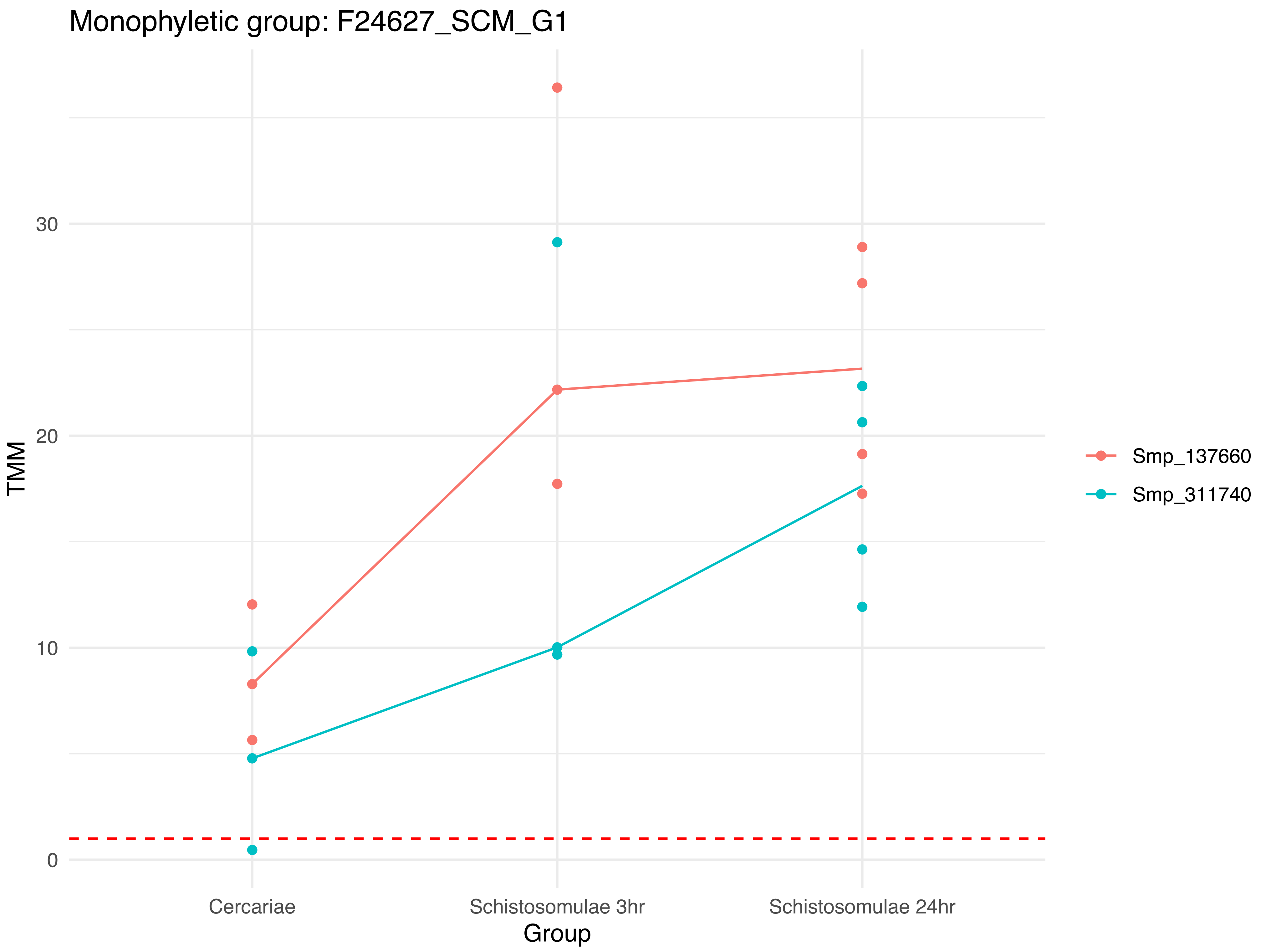

Monophyletic group: F24627_SCM_G1
TMM
30
20
10
0
Cercariae
Schistosomulae 3hr
Schistosomulae 24hr
Group
Smp_137660
Smp_311740

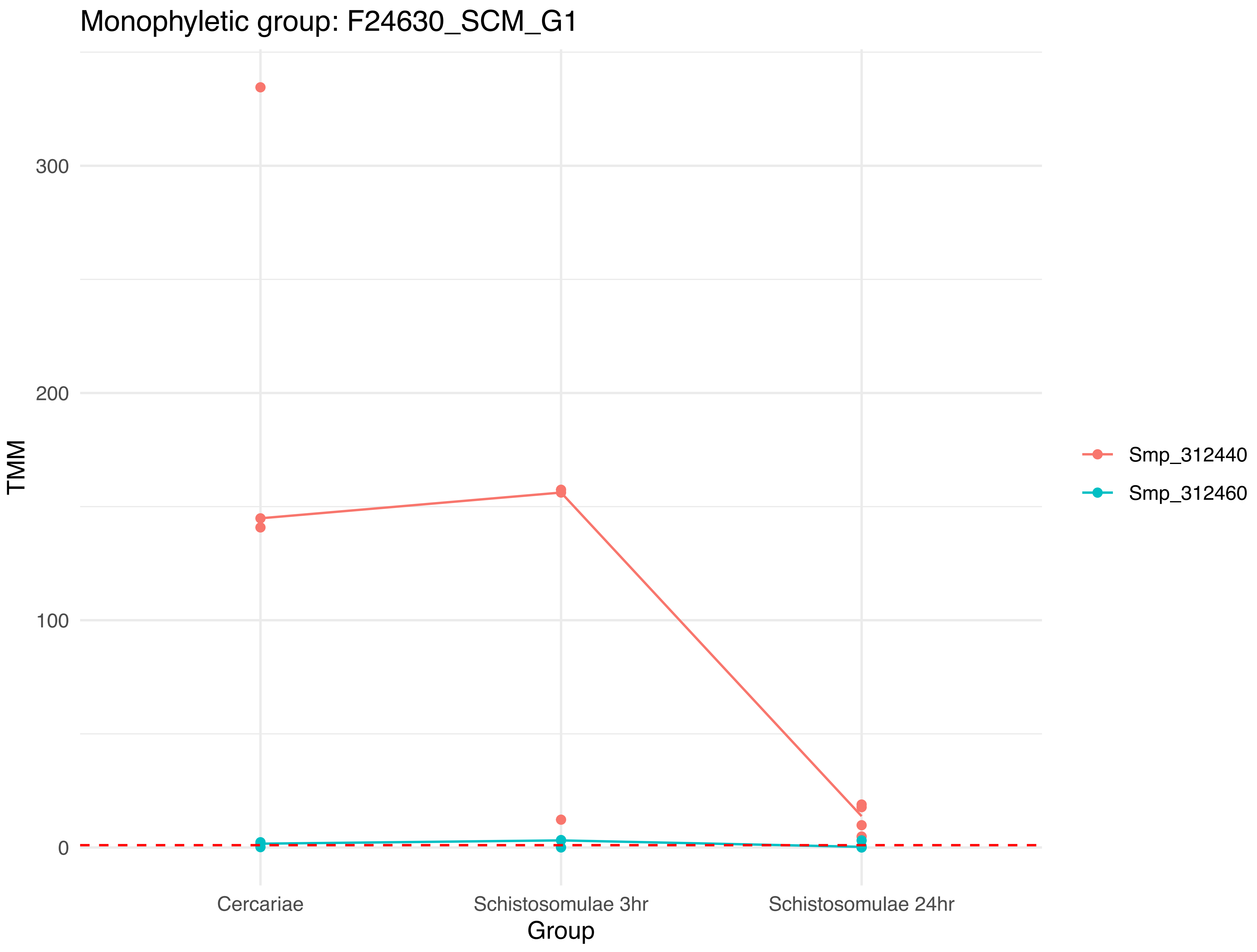

Monophyletic group: F24630_SCM_G1
TMM
300
200
100
0
Cercariae
Schistosomulae 3hr
Schistosomulae 24hr
Group
Smp_312440
Smp_312460

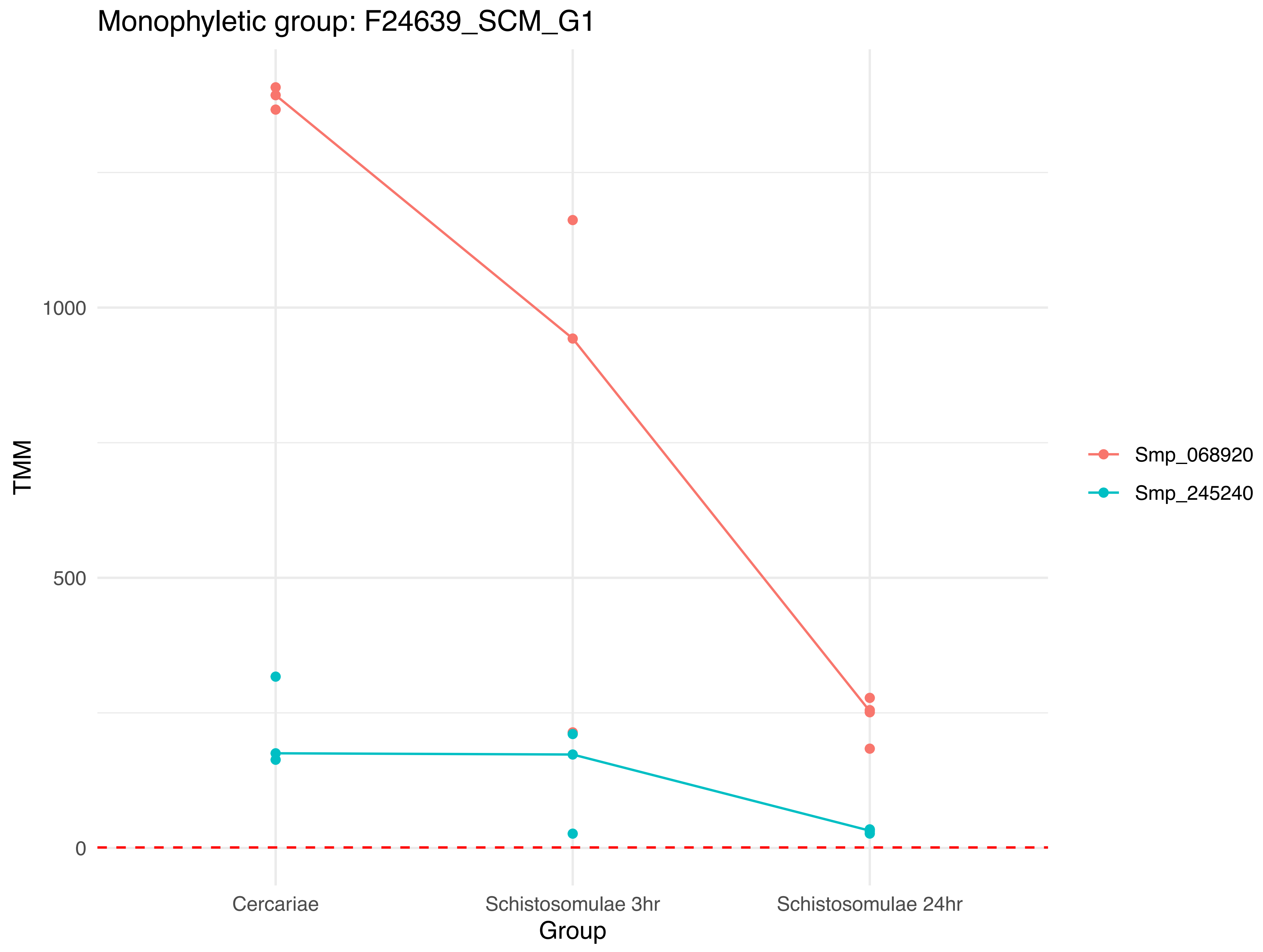
Monophyletic group: F24639_SCM_G1
TMM
1000
500
0
Cercariae
Schistosomulae 3hr
Schistosomulae 24hr
Group
Smp_068920
Smp_245240

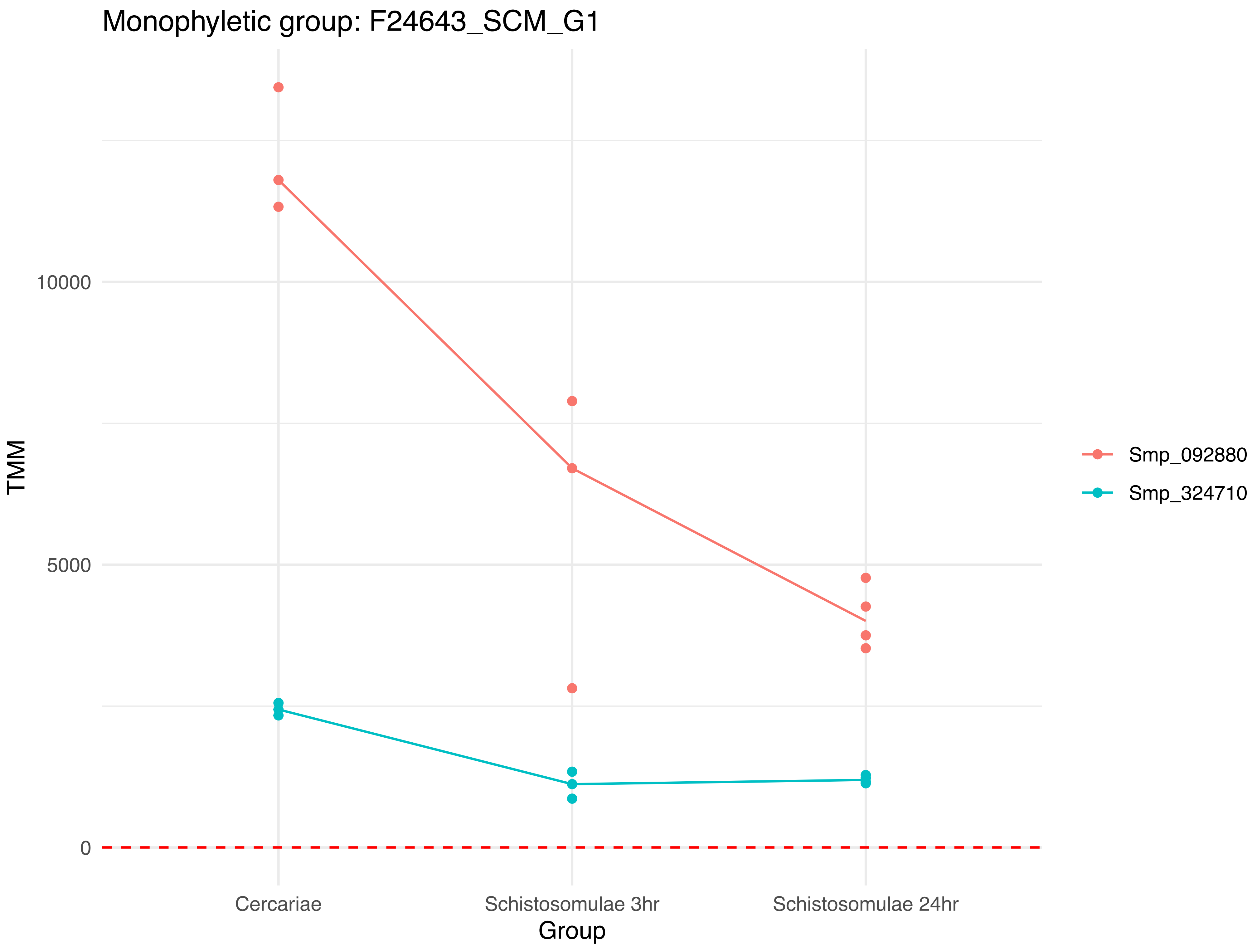
Monophyletic group: F24643_SCM_G1
10000
5000
0
TMM
Cercariae
Schistosomulae 3hr
Schistosomulae 24hr
Group
Smp_092880
Smp_324710

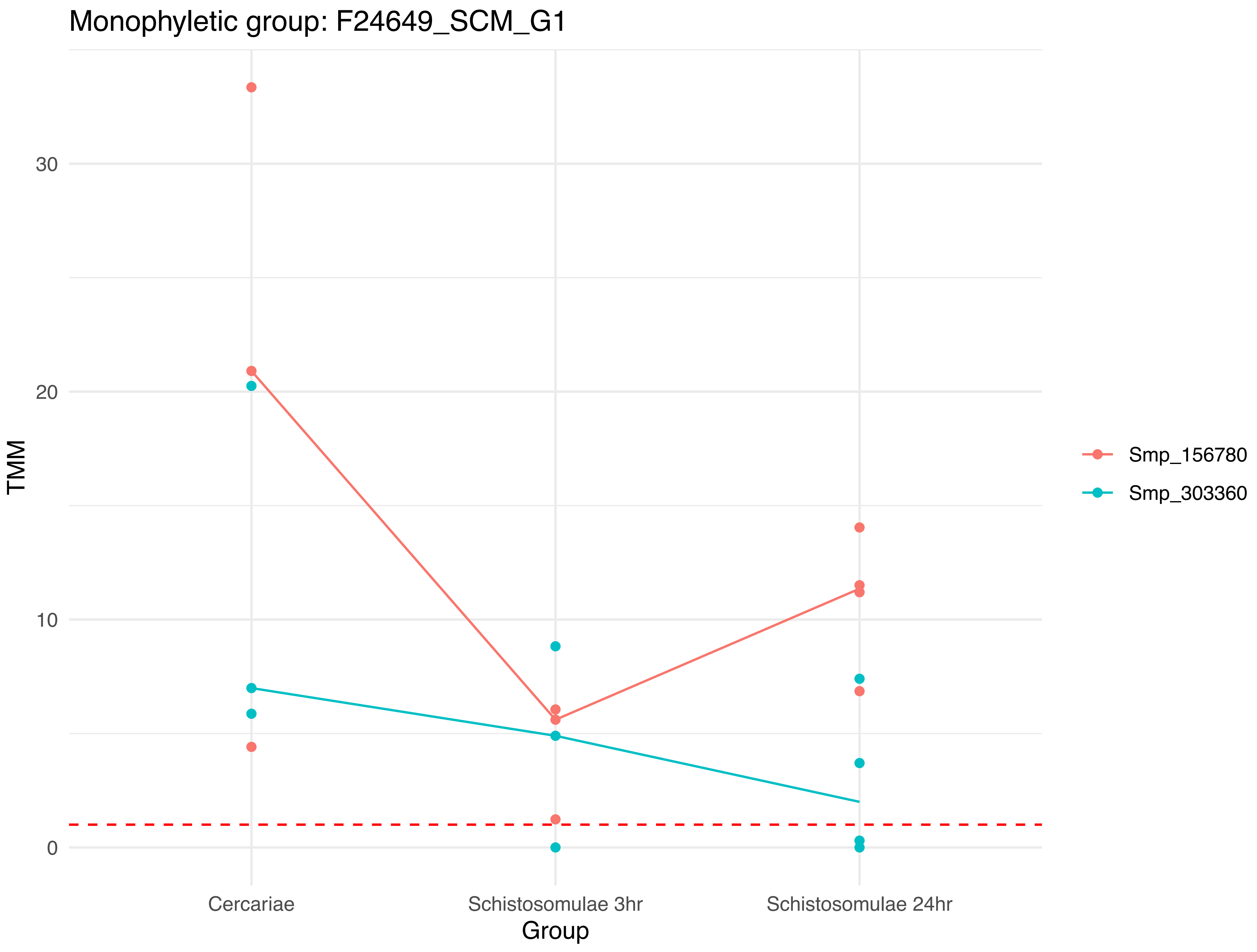

Monophyletic group: F24649_SCM_G1
30
20
10
0
TMM
Cercariae
Schistosomulae 3hr
Schistosomulae 24hr
Group
Smp_156780
Smp_303360

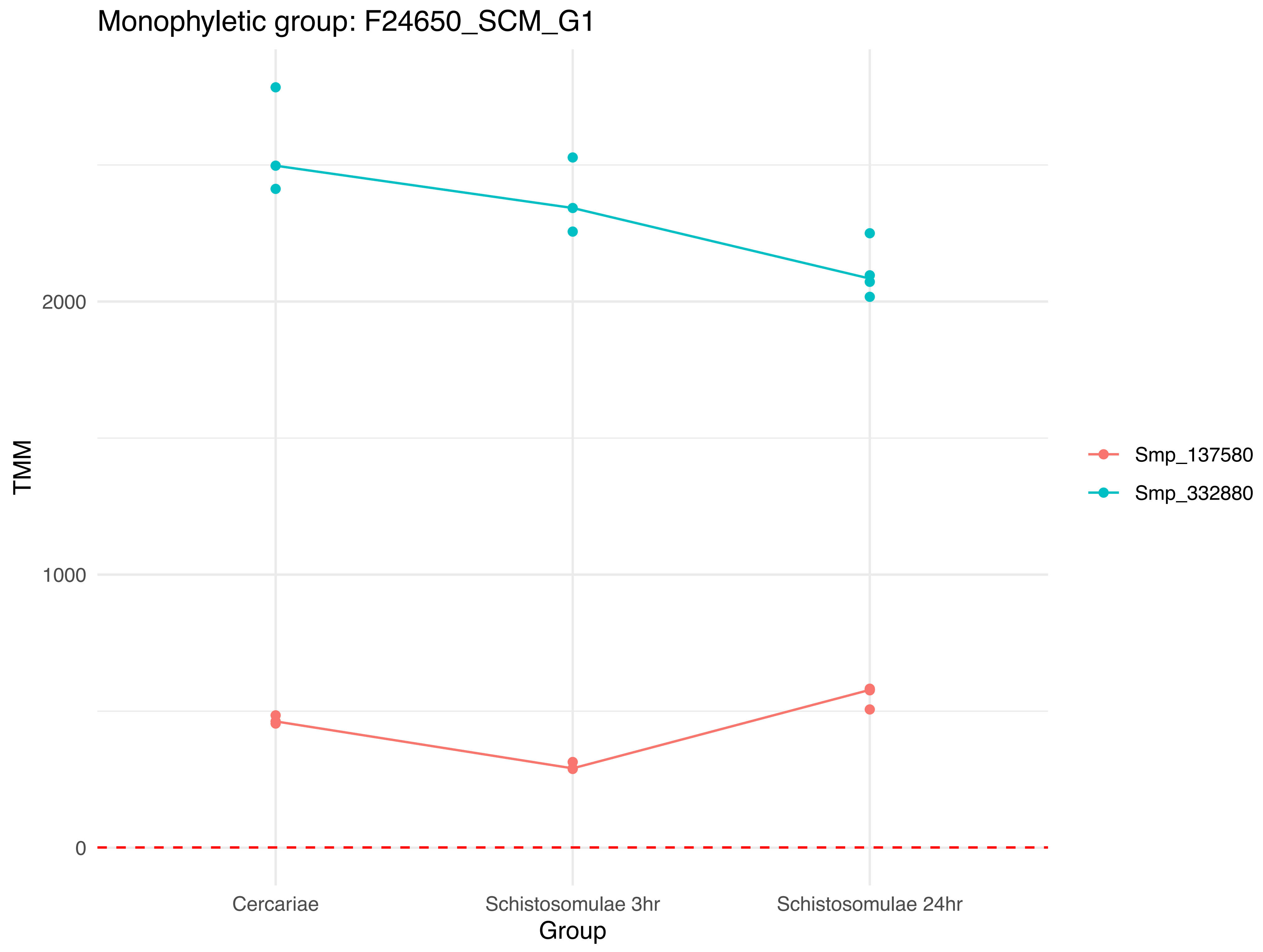

Monophyletic group: F24650_SCM_G1
TMM
2000
1000
0
Cercariae
Schistosomulae 3hr
Schistosomulae 24hr
Group
Smp_137580
Smp_332880

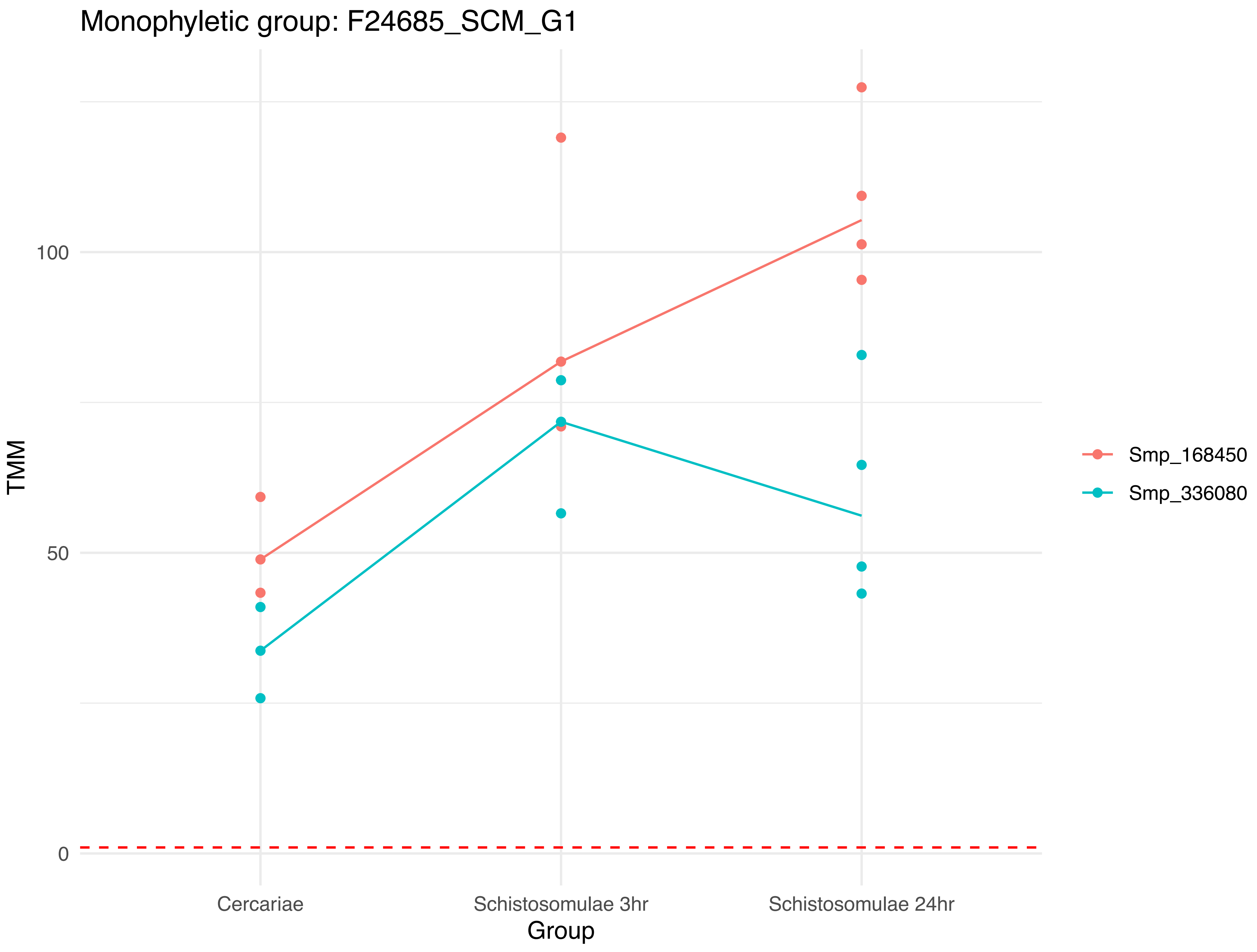

Monophyletic group: F24685_SCM_G1
TMM
100
50
0
Cercariae
Schistosomulae 3hr
Schistosomulae 24hr
Group
Smp_168450
Smp_336080

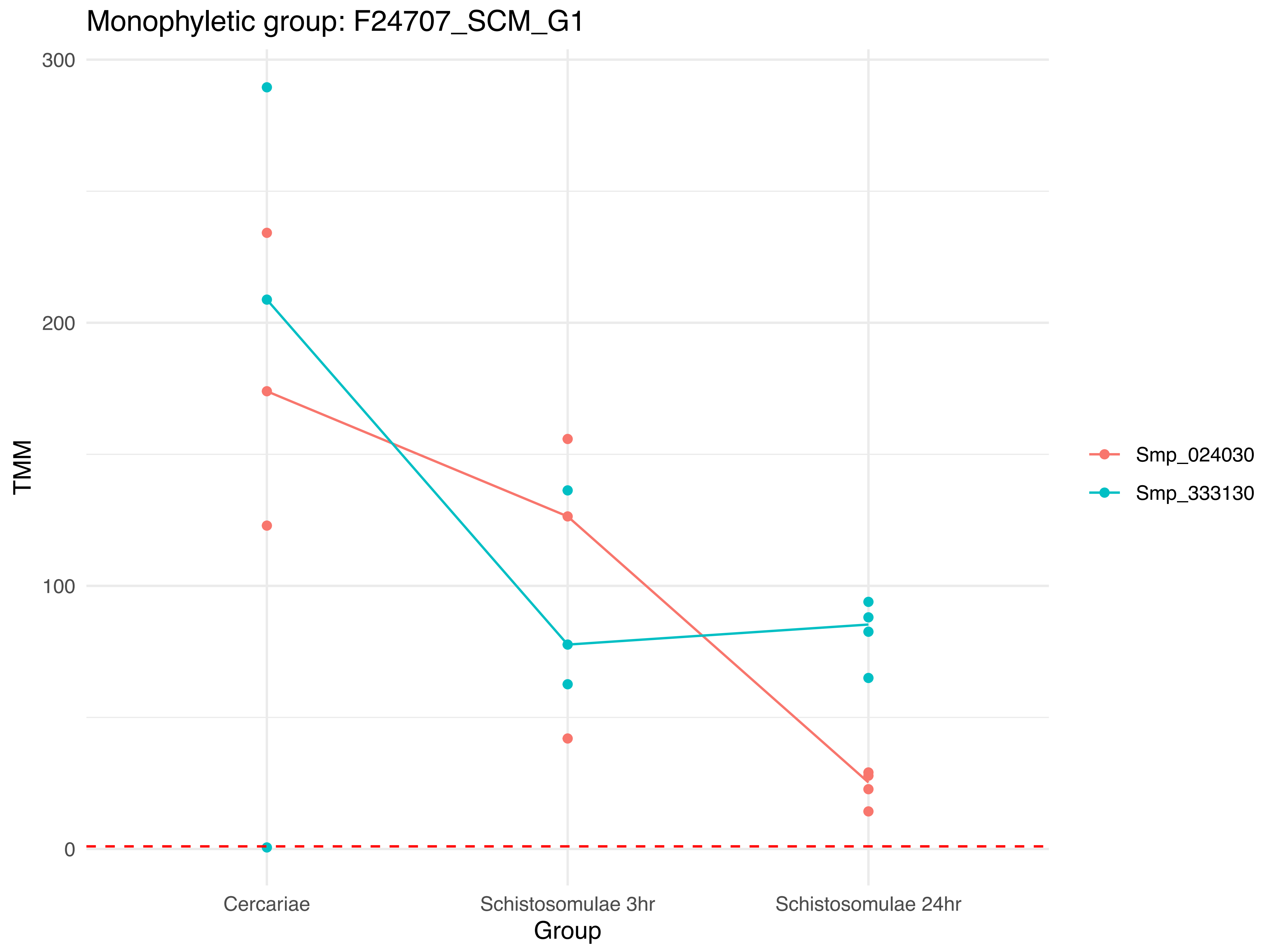

Monophyletic group: F24707_SCM_G1
300
200
100
0
TMM
Cercariae
Schistosomulae 3hr
Schistosomulae 24hr
Group
Smp_024030
Smp_333130

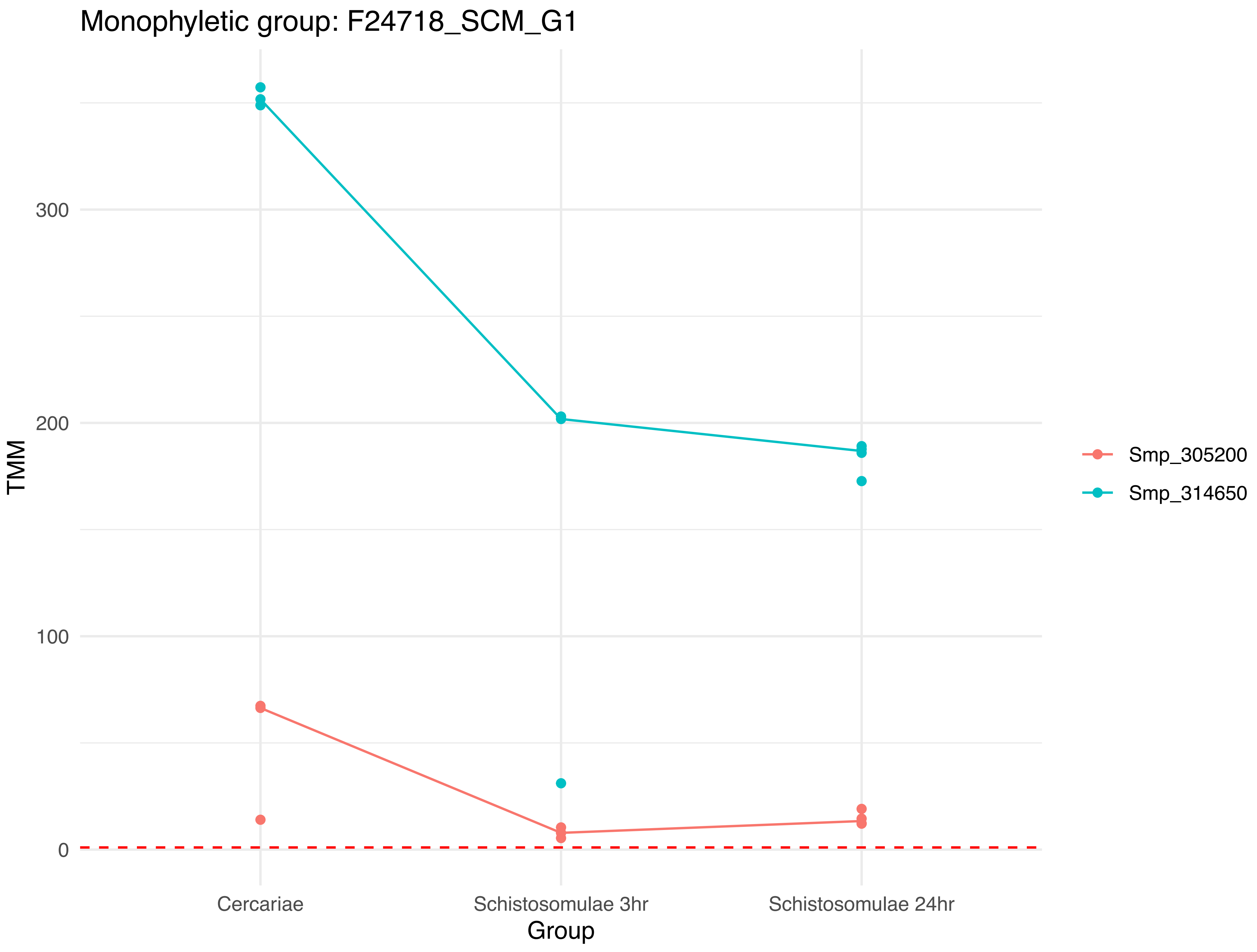

Monophyletic group: F24718_SCM_G1
TMM
300
200
100
0
Cercariae
Schistosomulae 3hr
Schistosomulae 24hr
Group
Smp_305200
Smp_314650

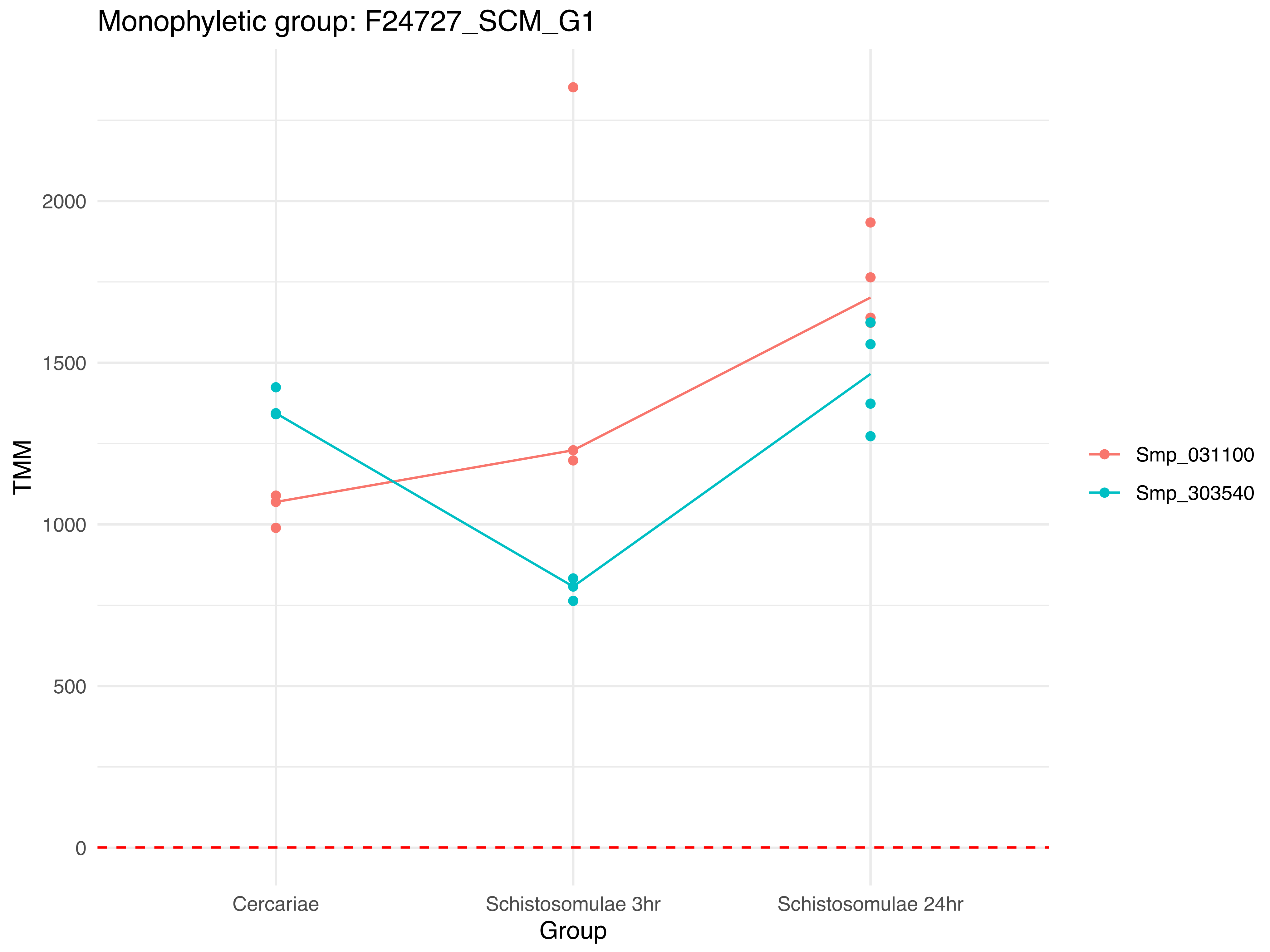

Monophyletic group: F24727_SCM_G1
TMM
2000
1500
1000
500
0
Cercariae
Schistosomulae 3hr
Schistosomulae 24hr
Group
Smp_031100
Smp_303540

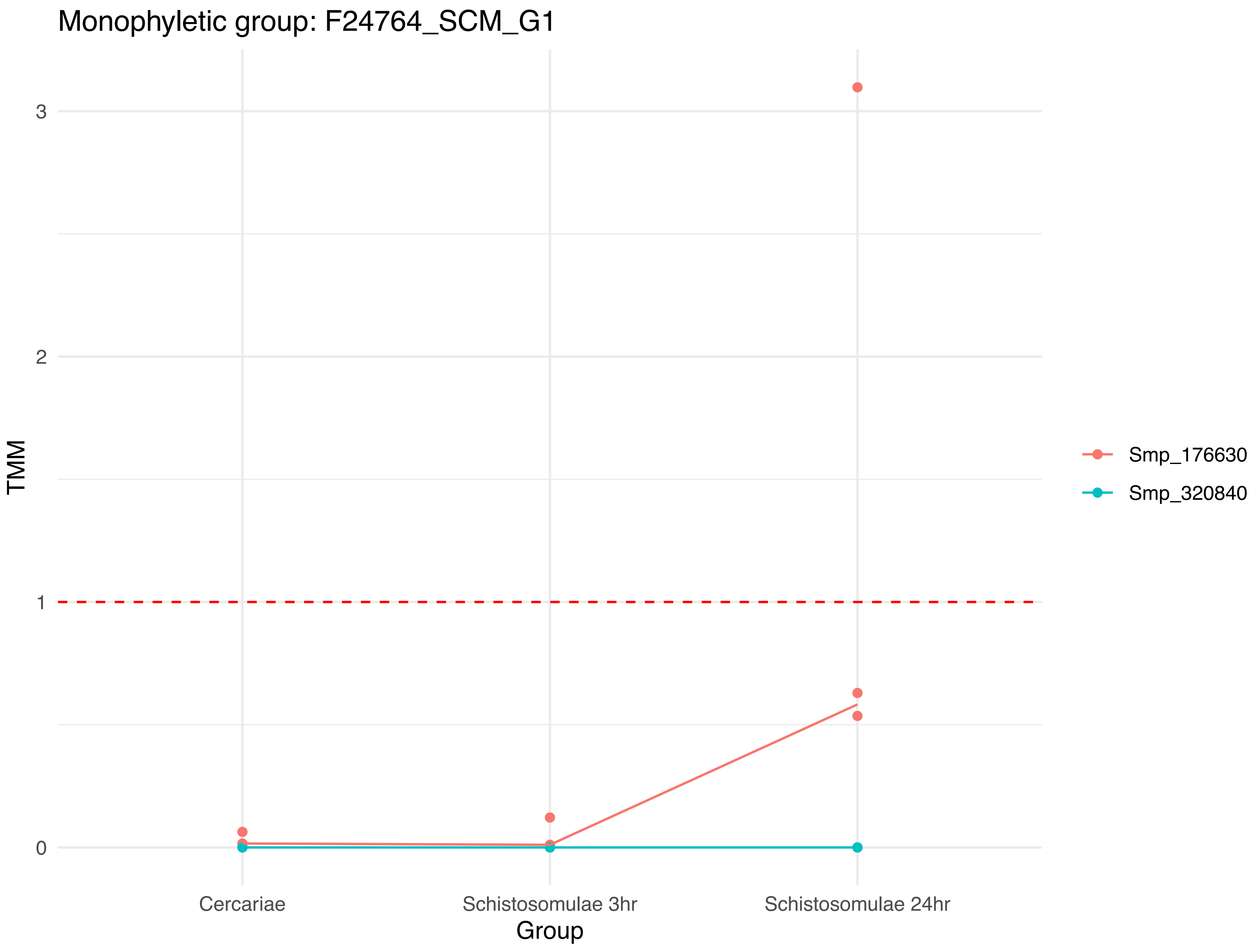

Monophyletic group: F24764_SCM_G1
TMM
3
2
1
0
Cercariae
Schistosomulae 3hr
Schistosomulae 24hr
Group
Smp_176630
Smp_320840

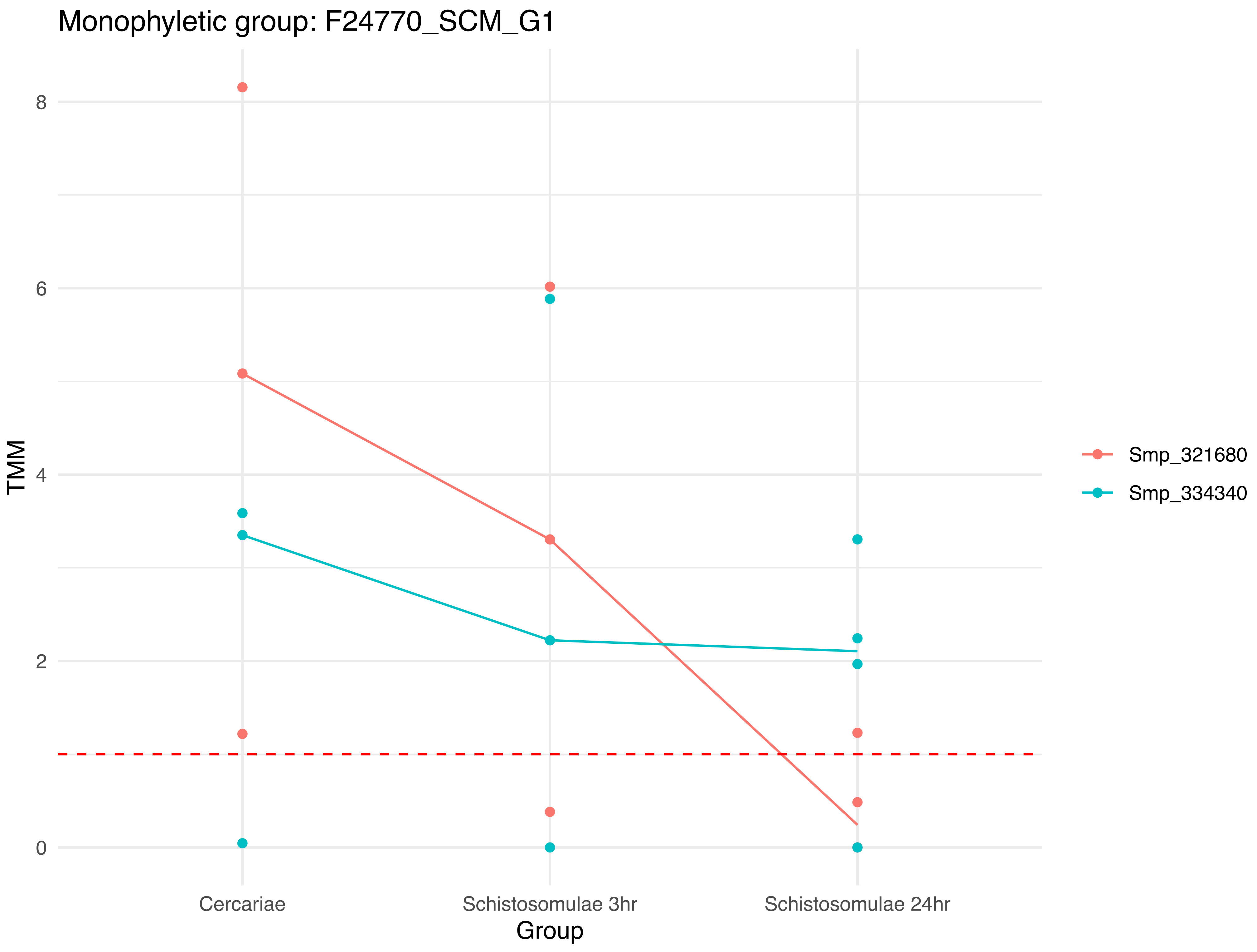

Monophyletic group: F24770_SCM_G1
8
6
4
2
0
TMM
Cercariae
Schistosomulae 3hr
Schistosomulae 24hr
Group
Smp_321680
Smp_334340

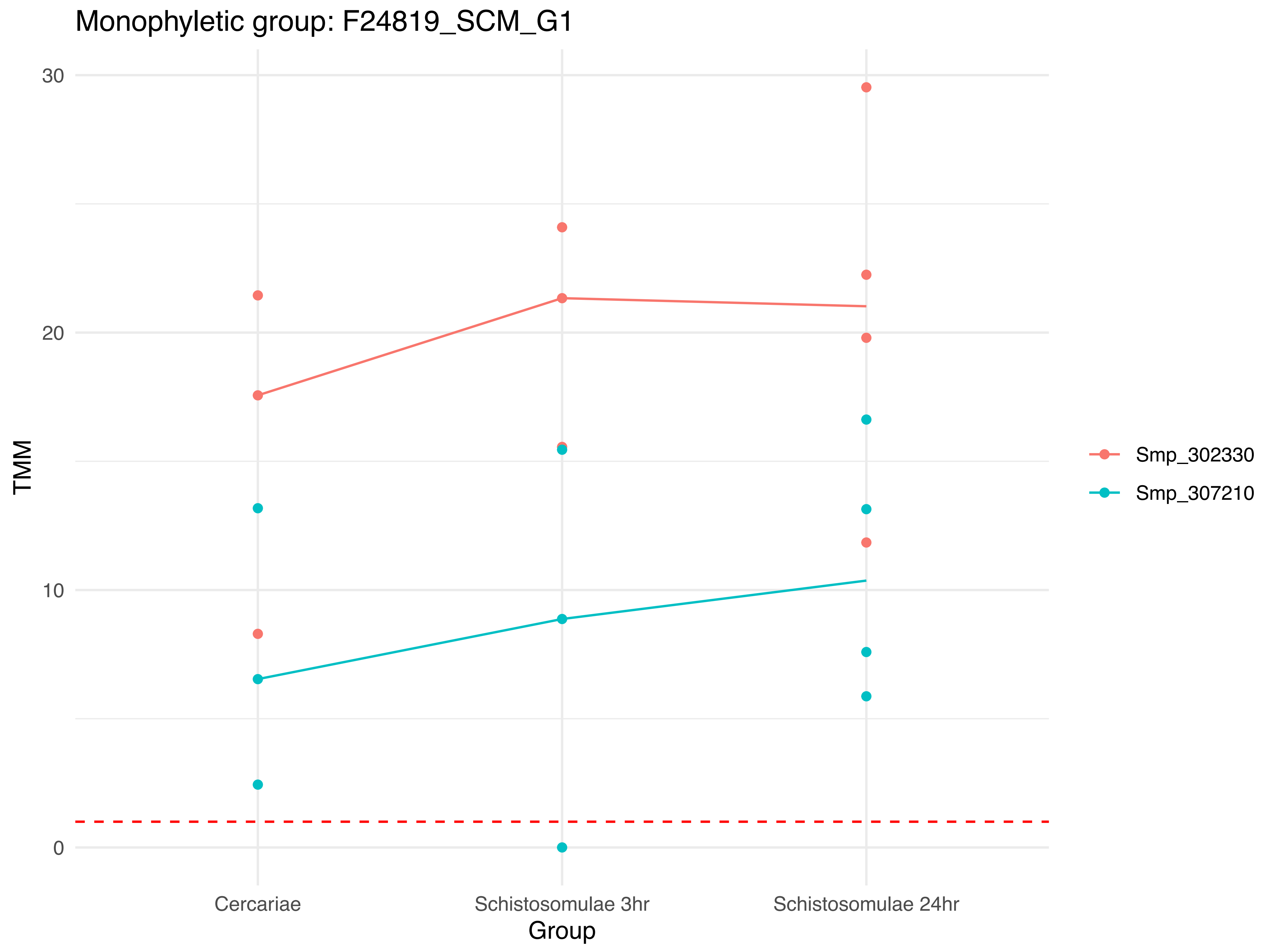

Monophyletic group: F24819_SCM_G1
30
20
10
0
TMM
Cercariae
Schistosomulae 3hr
Schistosomulae 24hr
Group
Smp_302330
Smp_307210

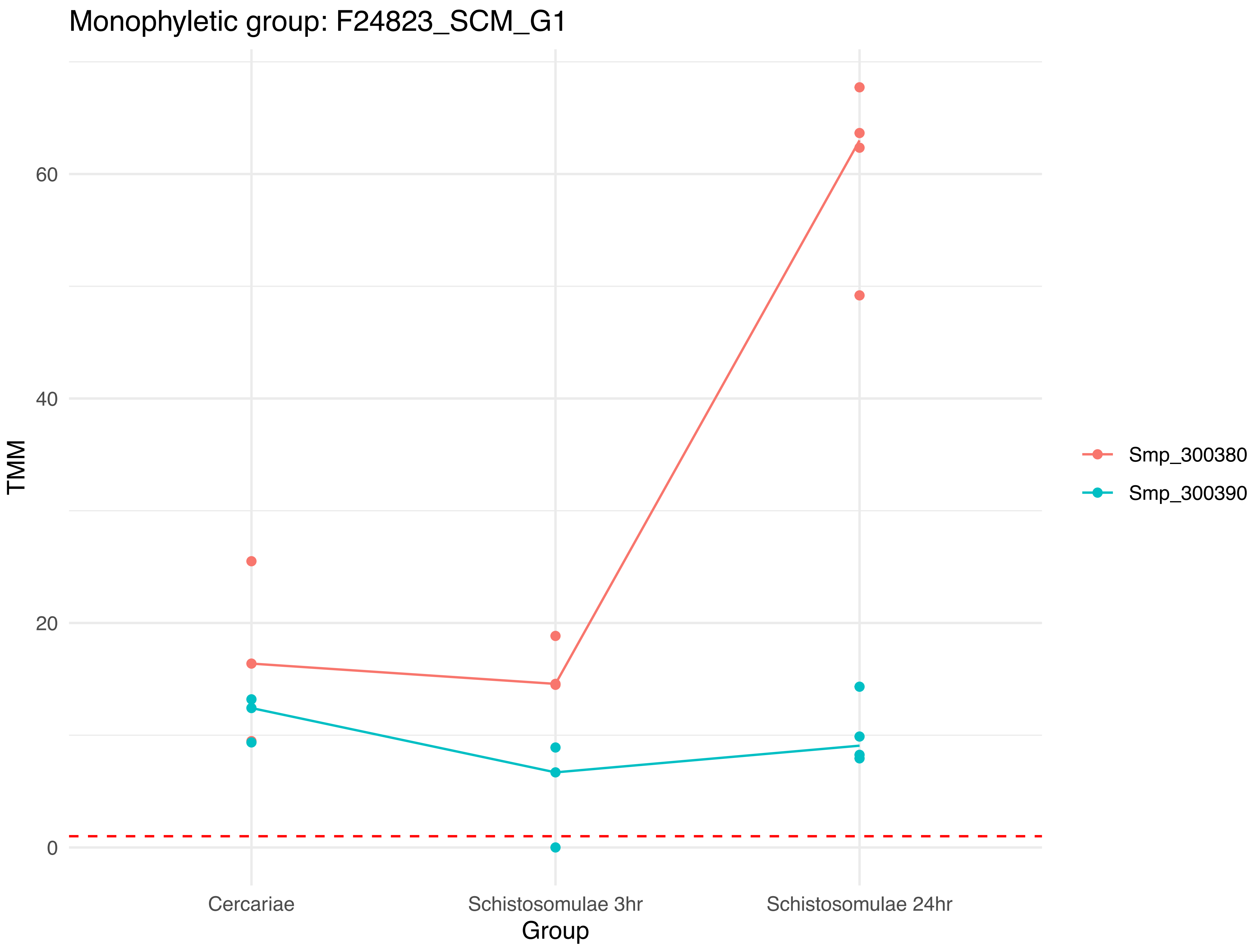
Monophyletic group: F24823_SCM_G1
TMM
60
40
20
0
Cercariae
Schistosomulae 3hr
Schistosomulae 24hr
Group
Smp_300380
Smp_300390

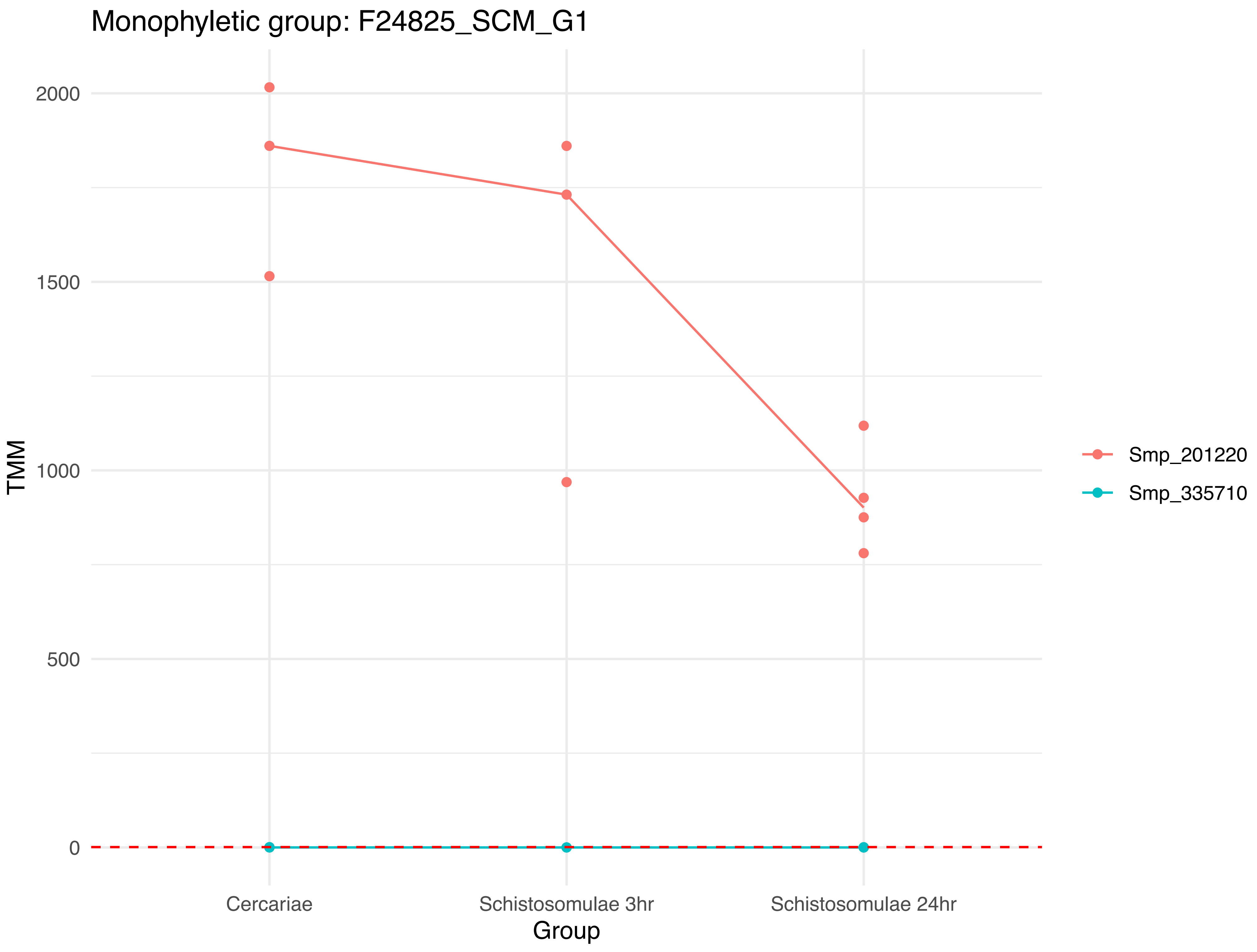
Monophyletic group: F24825_SCM_G1
2000
1500
1000
500
0
TMM
Cercariae
Schistosomulae 3hr
Schistosomulae 24hr
Group
Smp_201220
Smp_335710

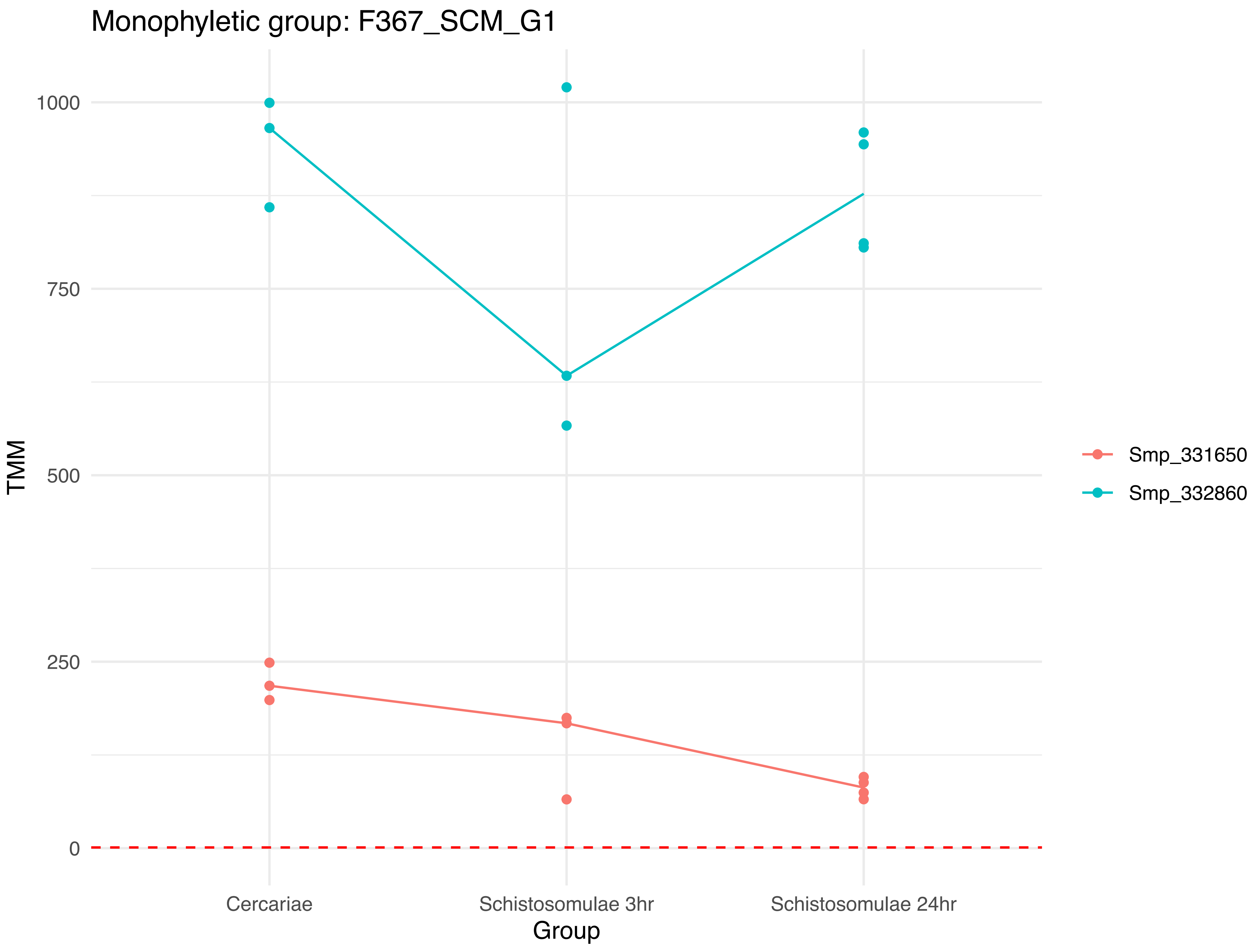
Monophyletic group: F367_SCM_G1
TMM
1000
750
500
250
0
Cercariae
Schistosomulae 3hr
Schistosomulae 24hr
Group
Smp_331650
Smp_332860

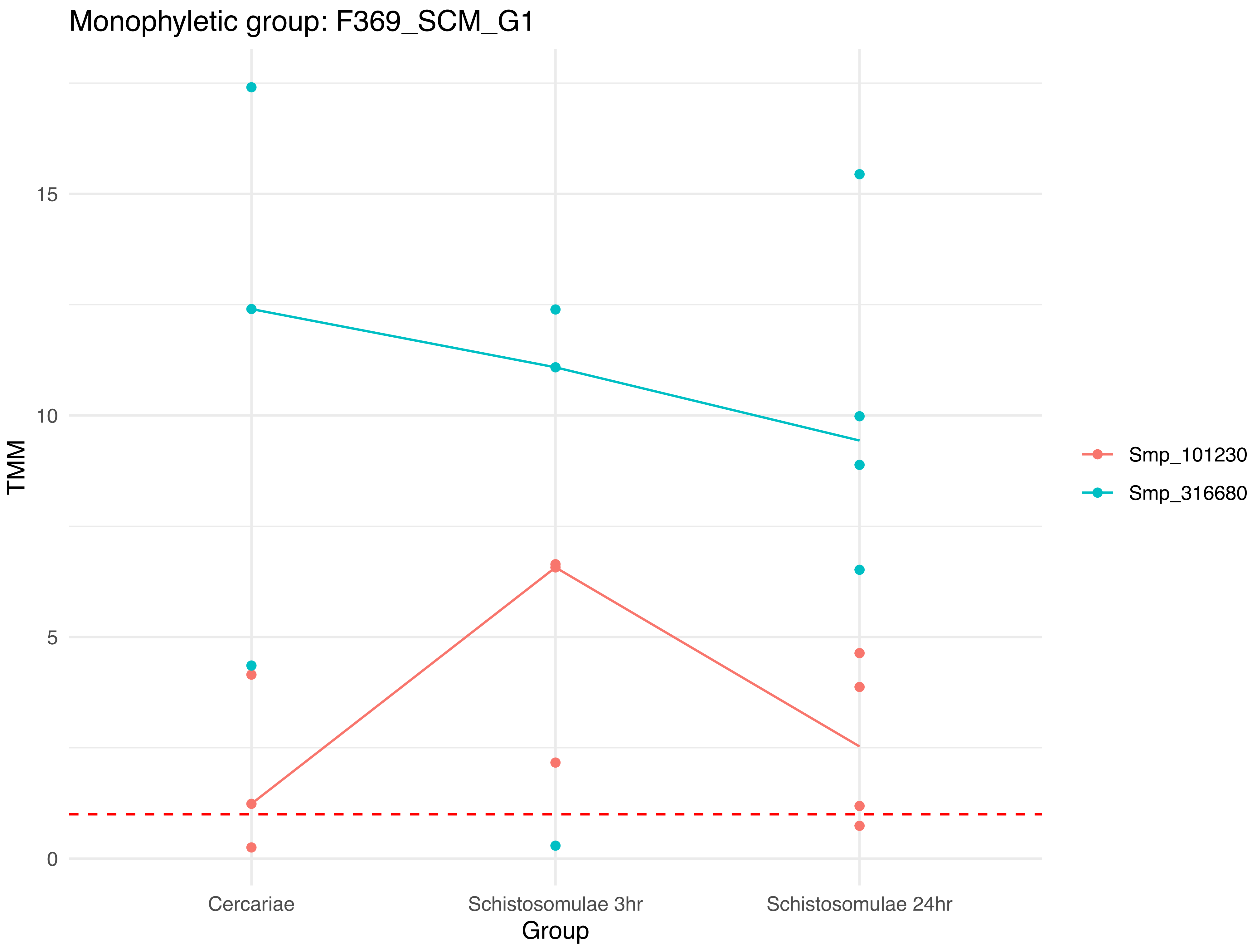

Monophyletic group: F369_SCM_G1
15
10
5
0
TMM
Cercariae
Schistosomulae 3hr
Schistosomulae 24hr
Group
Smp_101230
Smp_316680

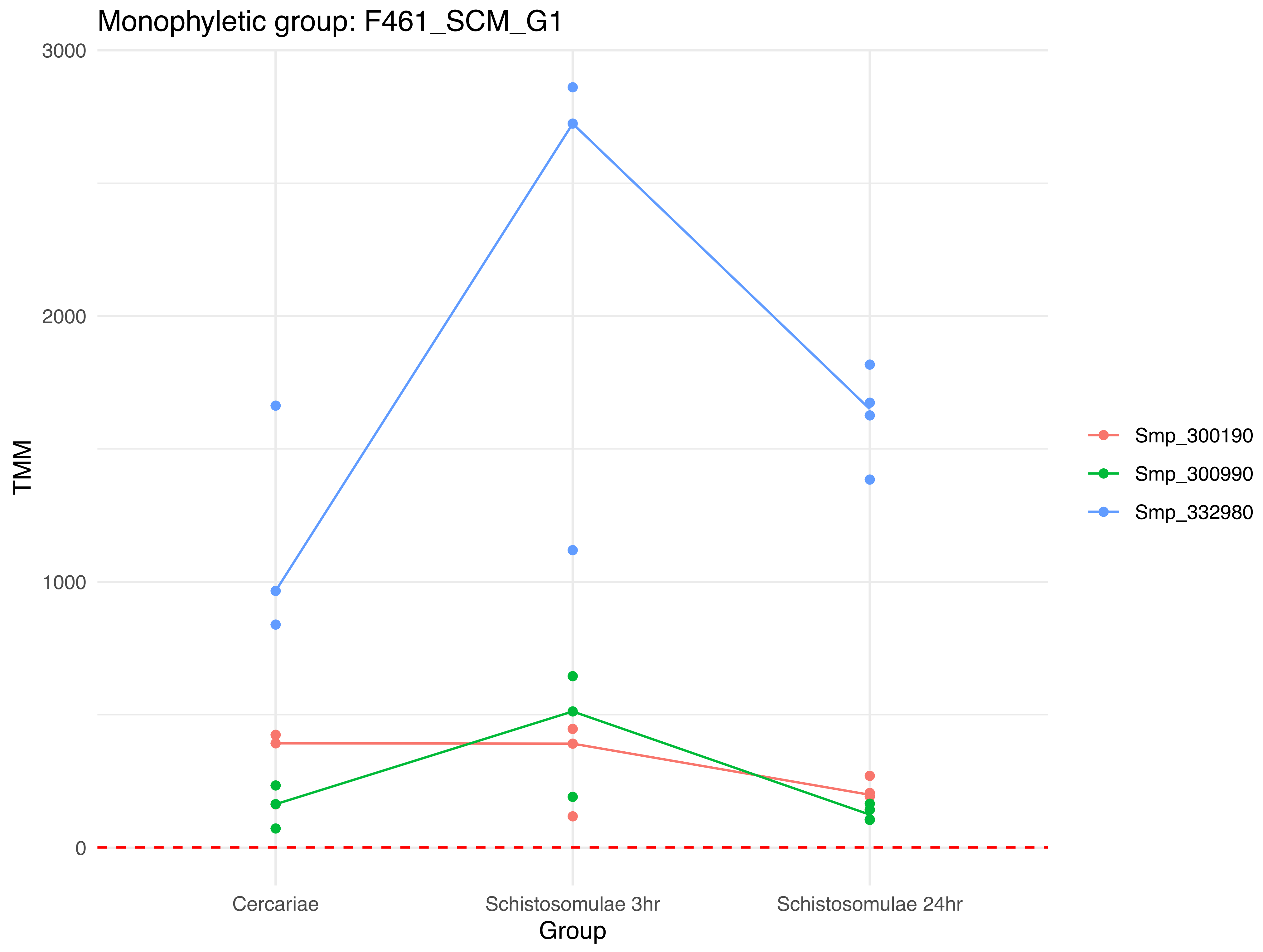
Monophyletic group: F461_SCM_G1
3000
2000
1000
0
TMM
Cercariae
Schistosomulae 3hr
Schistosomulae 24hr
Group
Smp_300190
Smp_300990
Smp_332980

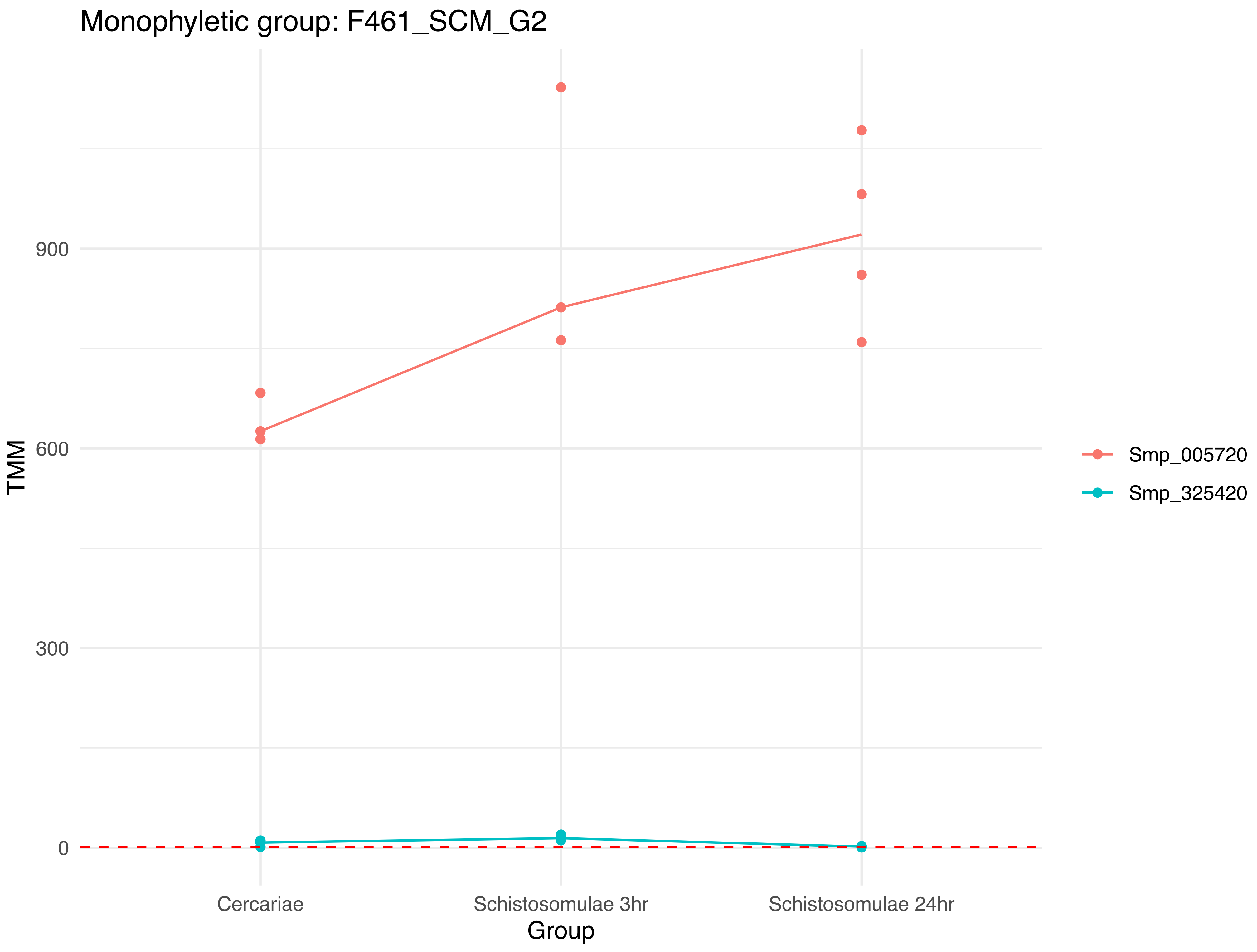

Monophyletic group: F461_SCM_G2
TMM
900
600
300
0
Cercariae
Schistosomulae 3hr
Schistosomulae 24hr
Group
Smp_005720
Smp_325420

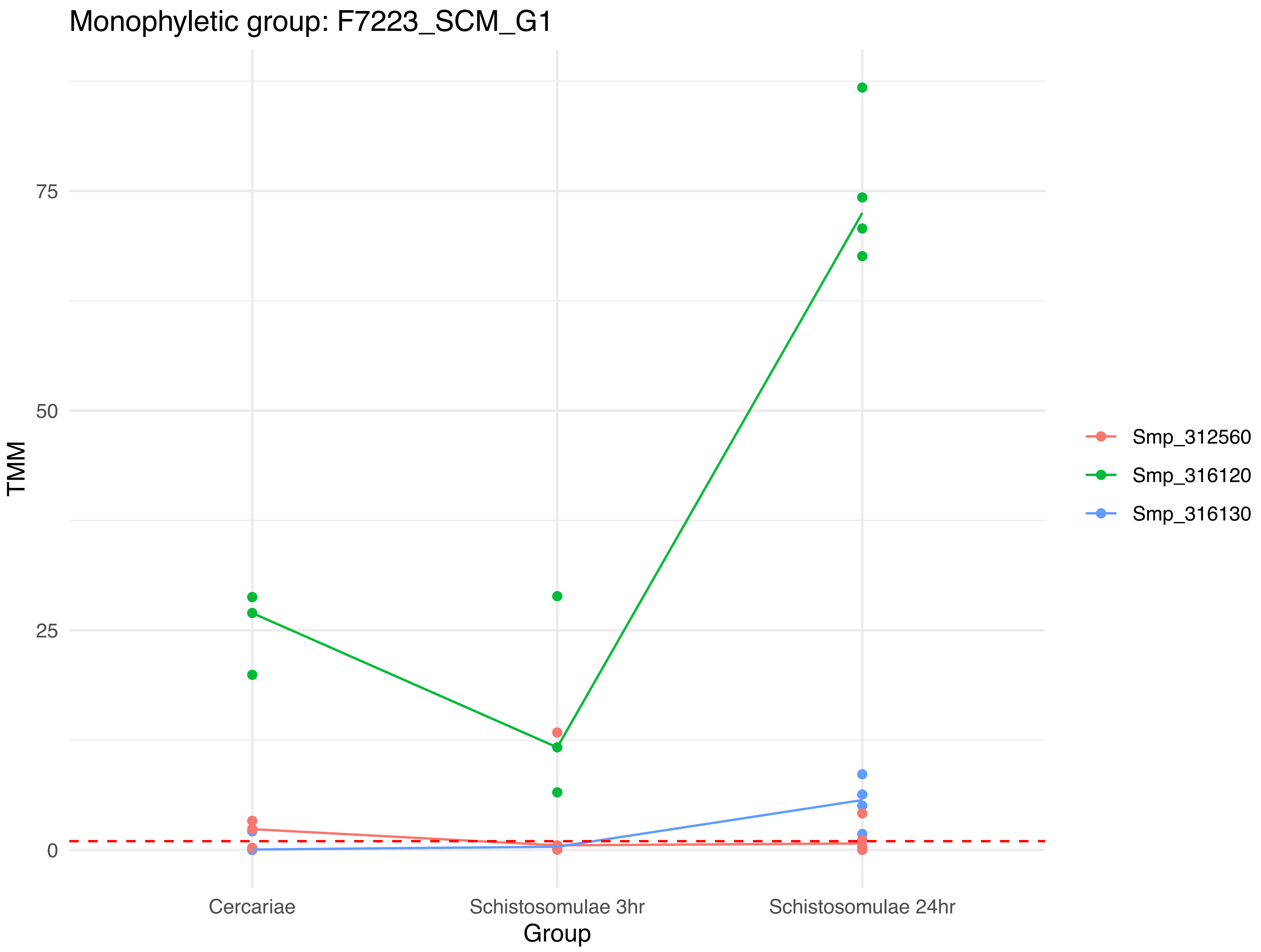

Monophyletic group: F7223_SCM_G1
TMM
75
50
25
0
Cercariae
Schistosomulae 3hr
Schistosomulae 24hr
Group
Smp_312560
Smp_316120
Smp_316130

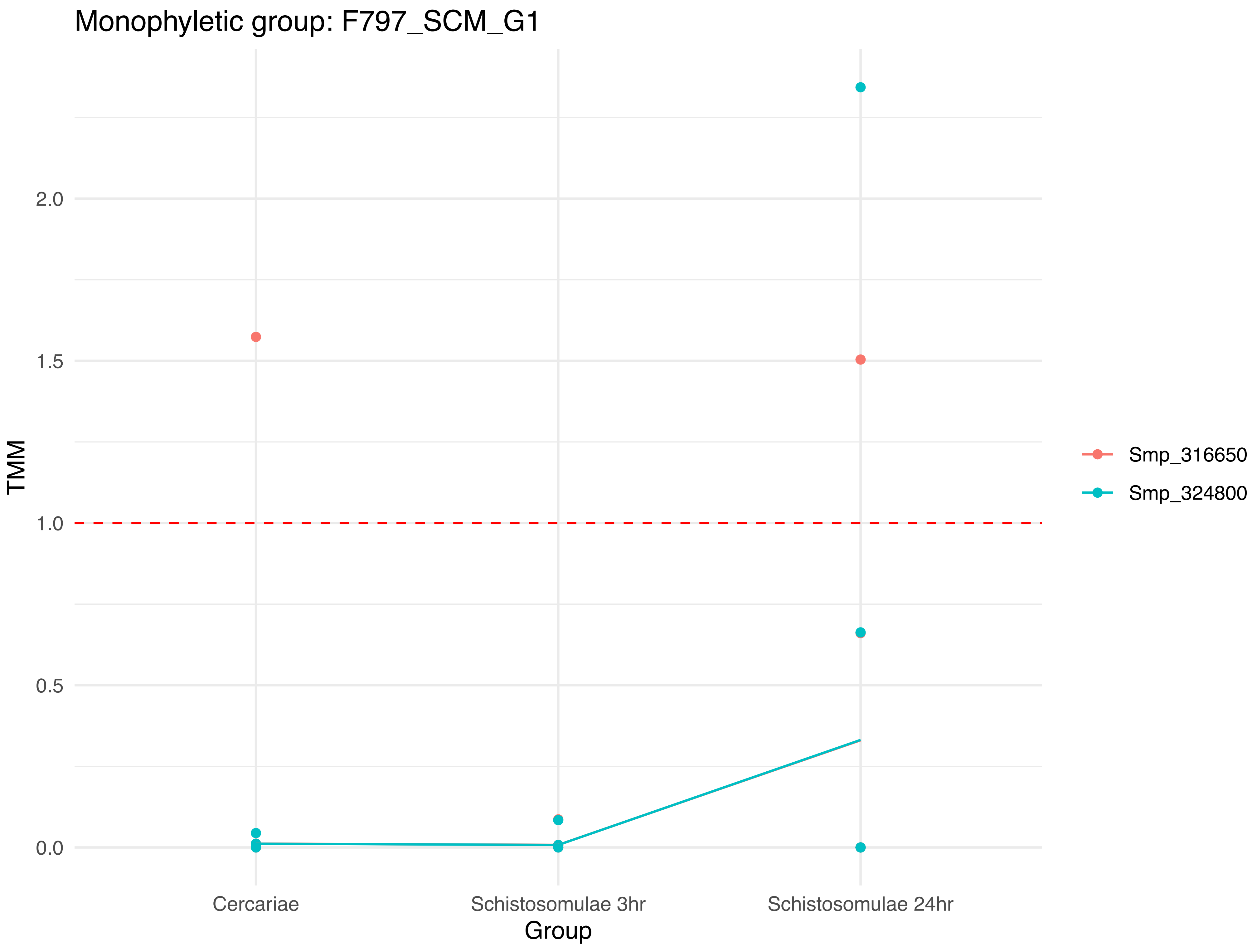

Monophyletic group: F797_SCM_G1
2.0
1.5
1.0
0.5
0.0
TMM
Cercariae
Schistosomulae 3hr
Schistosomulae 24hr
Group
Smp_316650
Smp_324800

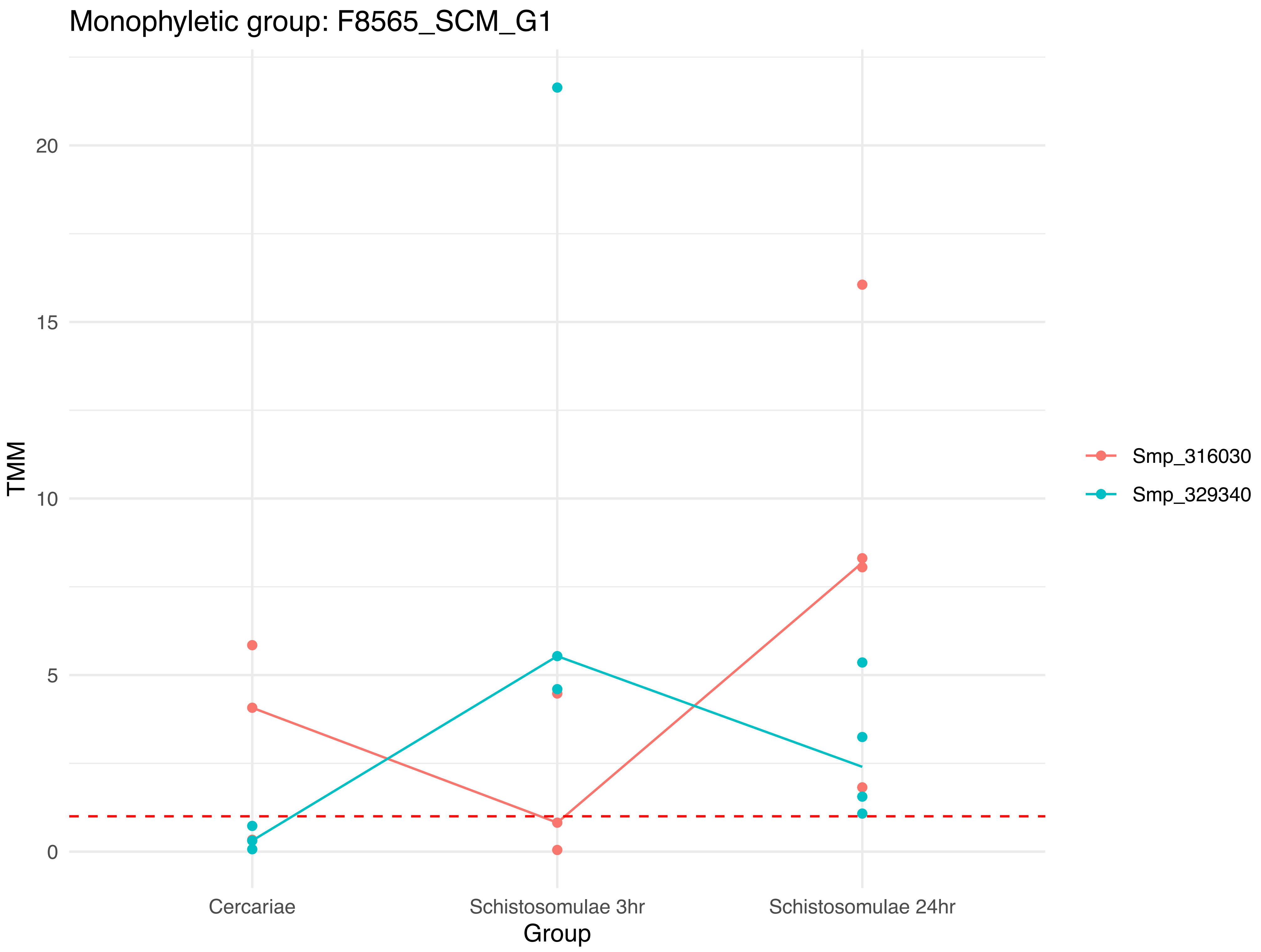

Monophyletic group: F8565_SCM_G1
TMM
20
15
10
5
0
Cercariae
Schistosomulae 3hr
Schistosomulae 24hr
Group
Smp_316030
Smp_329340
