## Supplementary Figure 8. Box-plots showing the distribution of Pearson correlation coefficients (r) of expression level of inparalogs by distance. for "Evolutionary analysis of genome-specific duplications in flatworm genomes"

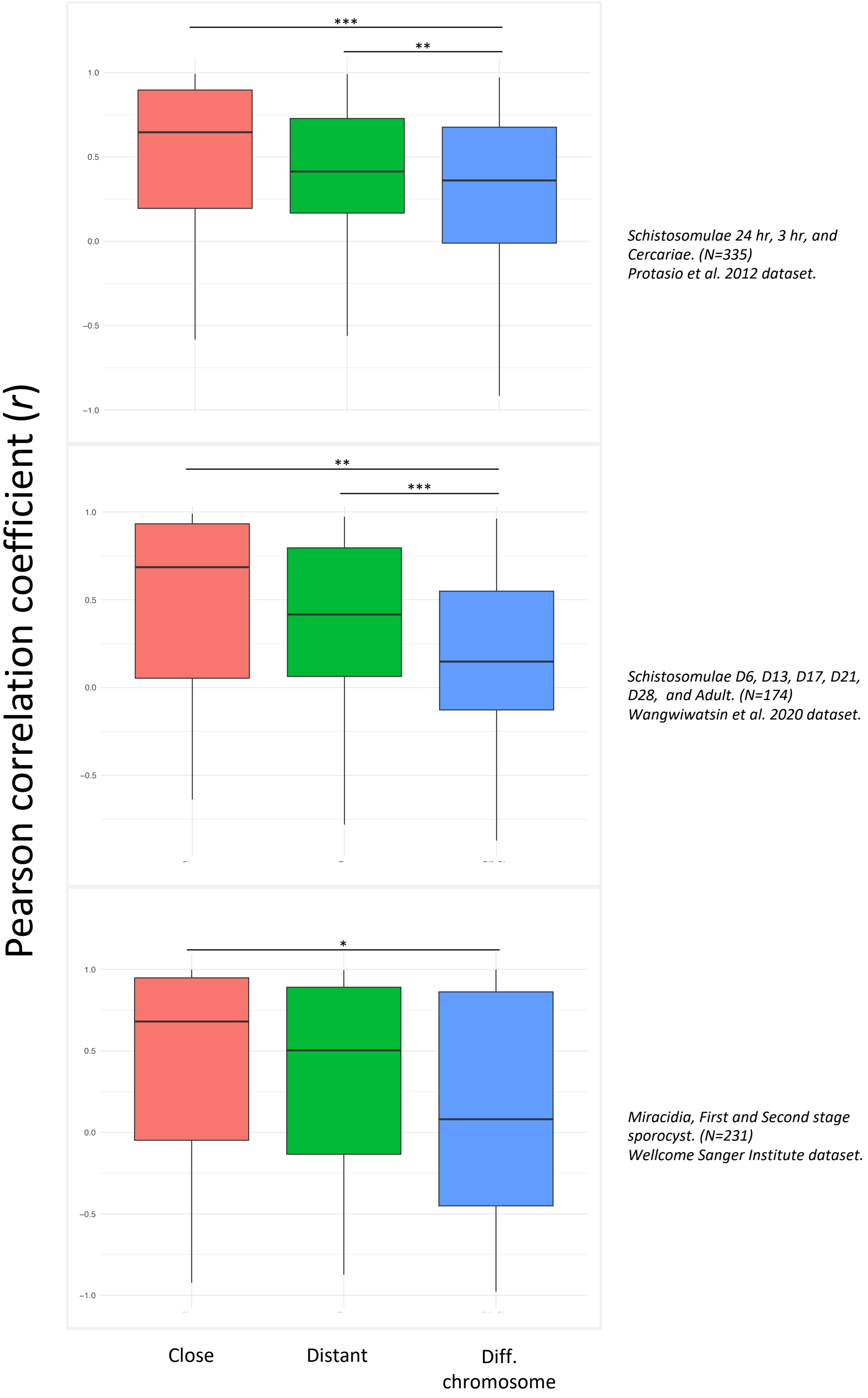


**Supplementary Figure 8**. Box-plots showing the distribution of Pearson correlation coefficients (r) of expression level (TMM) of inparalogs classified according to physical distance in the genome. All pairs of inparalogs were classified in "Close", "Distant", and "Diff. chromosome" using a threshold of 10 kbp. Only genes showing a median TMM above one in all time points, within each study, were considered in the analysis. The number of pairwise comparisons included is shown between brackets. Statistical significance was assessed by Mann-Whitney U-test and indicated by asterisks: $p \leq 0.1$ (*), $p \leq 0.05$ (**) , and $p \leq 0.01$ (***).
