## Supplementary Figure 9. Summary of results of the molecular evolution tests. for "Evolutionary analysis of genome-specific duplications in flatworm genomes"

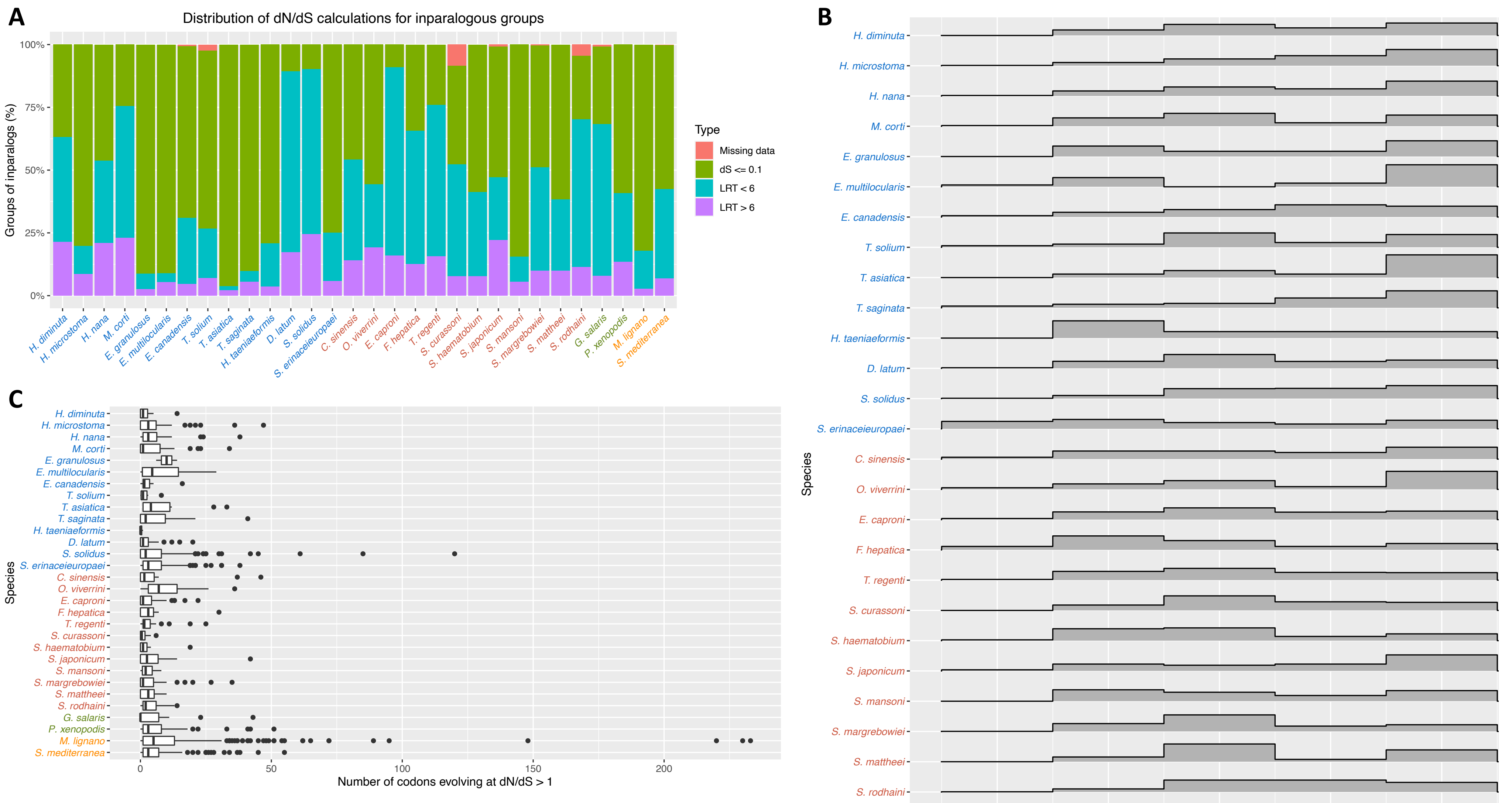


Supplementary Figure 9. Summary of results of the molecular evolution tests. A) Distribution of inparalogous groups according to the dN/dS test results. All the inparalogous groups with a total dS estimation below 0.1 were not considered for the dN/dS test and are indicated as a green bar. The neutral model M1a, with no sites evolving under positive selection, was compared with the positive selection model M2a using the Likelihood Ratio Test in each imparalogous group. The sky blue bar indicates those groups in which the neutral model fits significantly better than the positive selection model. The magenta bar indicates those groups in which the positive selection model fits significantly better than the neutral model. Missing data, groups in which no test could be applied, were indicated in pink. B) Histogram showing the dN/dS rate distribution estimated in inparalogous groups in each species. Note that the dN/dS was estimated in each group according to the site model that fits better and that groups with a dS < 0.1 were not analyzed. C) Box plot of the number of sites evolving under positive selection in each group is indicated. The number of sites with dN/dS > 1 in a cluster was only considered if the positive selection model was significantly more probable than the neutral model.
